# Single-Cell Transcriptomic Signatures Enable Stratified Combination Therapy for Platinum-Resistant Ovarian Cancer

**DOI:** 10.64898/2026.03.04.709546

**Authors:** Laura Gall-Mas, Katrin Kleinmanns, Anna Pirttikoski, Mara Santarelli, Gina Stangeland, Jun Dai, David Fontaneda-Arenas, Matias M. Falco, Carola Doerr, Sampsa Hautaniemi, Johanna Hynninen, Emmet McCormack, Krister Wennerberg, Line Bjørge, Anna Vähärautio, Benno Schwikowski

**Affiliations:** Biotech Research & Innovation Centre (BRIC), University of Copenhagen, Copenhagen, Denmark; Centre for Cancer Biomarkers CCBIO, Department of Clinical Science, University of Bergen, Norway; Department of Obstetrics and Gynecology, Haukeland University Hospital, Bergen, Norway; Research Program in Systems Oncology, Research Programs Unit, Faculty of Medicine, University of Helsinki, Helsinki, Finland; Computational Systems Biomedicine Lab, Institut Pasteur, Université Paris Cité, 25–28 Rue du Dr Roux, 75015 Paris, France; Sorbonne Université, CNRS, LIP6, Paris, France; Department of Obstetrics and Gynecology, University of Turku and Turku University Hospital, Turku, Finland; Foundation for the Finnish Cancer Institute, Helsinki, Finland

**Author notes:** Corresponding authors **Correspondence and requests for materials** should be addressed to Benno Schwikowski. These authors contributed equally: Laura Gall-Mas, Katrin Kleinmanns, Anna Pirttikoski, Mara Santarelli.

## Abstract

In high-grade serous carcinoma (HGSC), extensive intra-tumoral heterogeneity hinders complete eradication and remains a major obstacle to developing combination therapies capable of eliminating subpopulations resistant to standard-of-care treatment. Using single-cell RNA sequencing of 72 samples from 54 HGSC patients spanning treatment-naïve, post-neoadjuvant chemotherapy and relapse stages, we established a carboplatin-anchored framework that identifies transcriptional signatures of intrinsic (pre-existing) and adaptive (therapy-induced) resistance in individual tumors and prioritizes mechanistically matched drugs to potentiate carboplatin efficacy. Candidate compounds were ranked by integrating orthogonal resources—viability (GDSC, PRISM) and perturbational transcriptomics (L1000, Perturb-seq)—to reduce context bias. Among 64 candidates, three carboplatin adjuvants enhanced long-term efficacy in patient-derived organoids (PDOs), and pevonedistat further significantly reduced tumor burden in orthotopic xenografts. This tiered validation pipeline—from short-term and long-term PDOs and *in vivo* orthoptic xenografts—establishes a translational framework linking single cell resistance programs to actionable, tumor-specific, carboplatin-anchored combinations for HGSC.

## Main

Emerging resistance to initially effective therapies is one of the key challenges of cancer care. High-grade serous carcinoma (HGSC), the most common and aggressive subtype of ovarian cancer, is typically diagnosed at advanced stages and marked by extensive intra- and inter-tumor heterogeneity. Standard first-line treatment—cytoreductive surgery followed by carboplatin- and paclitaxel-based chemotherapy—induces responses in most patients, but recurrent disease driven by carboplatin resistance remains the principal cause of mortality (Papageorgiou et al. 2025). Current biomarkers primarily assess the tumor’s DNA damage repair capacity (Soslow and Tornos 2011; Perez-Villatoro et al. 2022), or folate receptor expression (Van Gorp et al. 2025). However, recurrence management remains determined by prior carboplatin sensitivity and treatment-related toxicities, and no biomarkers exist to guide carboplatin-anchored therapeutic decisions—an unmet clinical need.

The broad initial response followed by frequent relapses reflects tumor subpopulations with differential chemosensitivity, including cells that persist and drive resistant recurrences. Despite extensive multi-omics efforts, targetable genetic drivers of platinum resistance remain elusive. This absence implicates non-genetic mechanisms—most notably dynamic cell state plasticity, intratumoral heterogeneity and microenvironmental influence—as critical contributors to therapeutic failure. Addressing resistance requires integrated high-resolution profiling, computational modeling of targetable heterogeneity, and patient-derived models for drug screening and validation. Single-cell RNA sequencing (scRNA-seq) enables high-resolution dissection of tumor heterogeneity and is increasingly incorporated into precision oncology to target resistant cell states. Computational tools such as BeyondCell (Fustero-Torre et al. 2021), scDrug (Hsieh et al. 2023), DREEP (Pellecchia et al. 2023) and scTherapy (Ianevski et al. 2024) link single-cell transcriptomes to drug sensitivity. BeyondCell introduced the concept of “therapeutic clusters”, groups of cells with shared drug-response profiles that enable subpopulation-specific therapy design. Similarly, DREEP and scDrug integrate large pharmacogenomic screens (e.g., GDSC, CTRP, PRISM) to predict single-cell drug sensitivity and associated expression signatures, highlighting the potential of scRNA-seq to reveal actionable vulnerabilities beyond bulk profiling. Although powerful, these frameworks (i) are mostly not anchored to standard-of-care regimens such as carboplatin, (ii) prioritize discrete cell clusters rather than continuous resistance programs, (iii) lack temporal precision to distinguish intrinsic from adaptive states, (iv) often rely on drug-response resources with variable lineage and assay contexts, and (v) rarely extend beyond short-term *in vitro* assays to long-term patient-derived organoids (PDOs) and *in vivo* testing in patient-derived xenograft (PDX) models.

To address these gaps, we present a carboplatin-anchored discovery framework integrating scRNA-seq, multimodal in-silico drug prioritization, and tiered patient-derived model validation. By analyzing scRNA-seq profiles from 54 HGSC patients—including treatment-naïve, post-NACT, and relapse—we resolved intrinsic and adaptive resistance programs as continuous cell-state spectra. These programs enabled patient-specific mapping of vulnerabilities and guided selection of drug candidates predicted to enhance carboplatin activity. Systematic testing in short- and long-term PDOs and orthotopic PDX identified three synergistic adjuvants, including pevonedistat, which reduced tumor burden *in vivo*. Because the resistance programs can be quantified in individual tumors, our pipeline establishes a foundation for personalized carboplatin-anchored combination therapy in HGSC.

## Results

### A multi-stage pipeline to identify carboplatin-anchored drug combinations

We established a multi-stage pipeline to discover drug combinations that overcome platinum resistance in HGSC (Fig. 1). Single-cell RNA-seq of treatment-naïve, post-NACT, and relapse tumors, plus normal fallopian tube epithelium, mapped epithelial cell-state heterogeneity. Two analytical branches generated resistance-associated transcriptional signatures.

**Figure 1.**
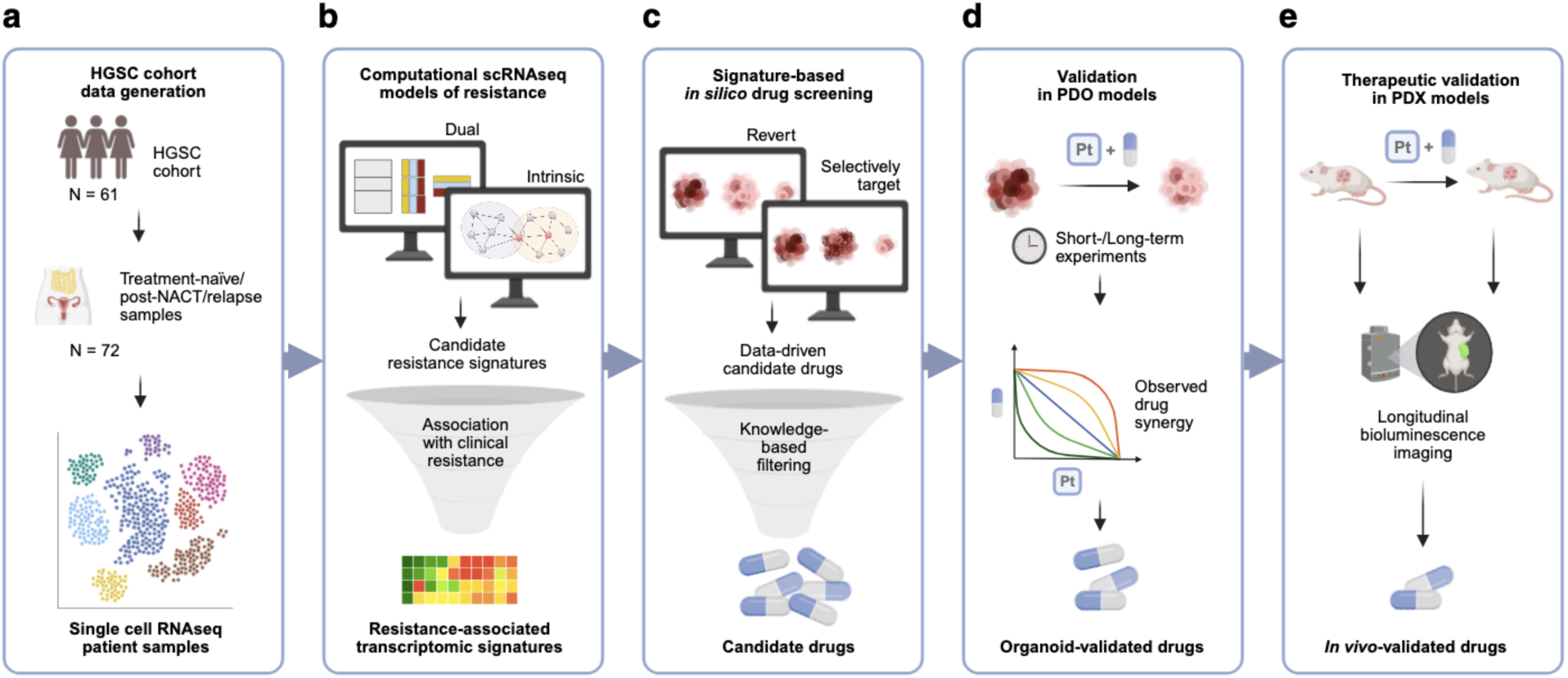
Pipeline for identifying and validating drug combinations to overcome platinum resistance in HGSC. **a,** Single-cell RNA-seq profiling of patient tumors (treatment-naïve, post-NACT, relapse) and healthy fallopian tube controls captures intratumoral heterogeneity. **b,** Complementary “Dual”/“Intrinsic” analyses generate transcriptomic signatures linked to pre-treatment/treatment-induced platinum resistance. **c,** *In silico* drug predictions using drug-response databases and perturbation profiles nominates candidate drugs targeting these signatures. **d,** Synergy testing in PDOs identifies carboplatin-sensitizing agents **e,** which are subsequently validated in PDX models, confirming *in vivo* efficacy and enabling clinical translation.

These signatures were applied to four orthogonal drug-response datasets (GDSC, PRISM, L1000, Perturb-seq) to prioritize drugs predicted either to reverse resistance-associated programs or selectively target resistant subpopulations. Candidate adjuvants were screened in short- and long-term PDO assays, and top combinations were evaluated in orthotopic PDX models.

### Discovery of Signatures Representing Intrinsic and Adaptive Resistance in HGSC

To investigate intratumoral heterogeneity underlying resistance, we analysed single-cell RNA-seq profiles from well-annotated clinical samples (n=72): treatment-naive (n=52), post-NACT (n=17), or relapsed tumors (n=3), along with control samples from healthy postmenopausal women (n=7) (Supplementary Table 1 and Supplementary Table 2). Using this dataset, we applied two complementary approaches to identify key transcriptional signatures of intratumoral heterogeneity (Fig. 2a).

**Figure 2.**
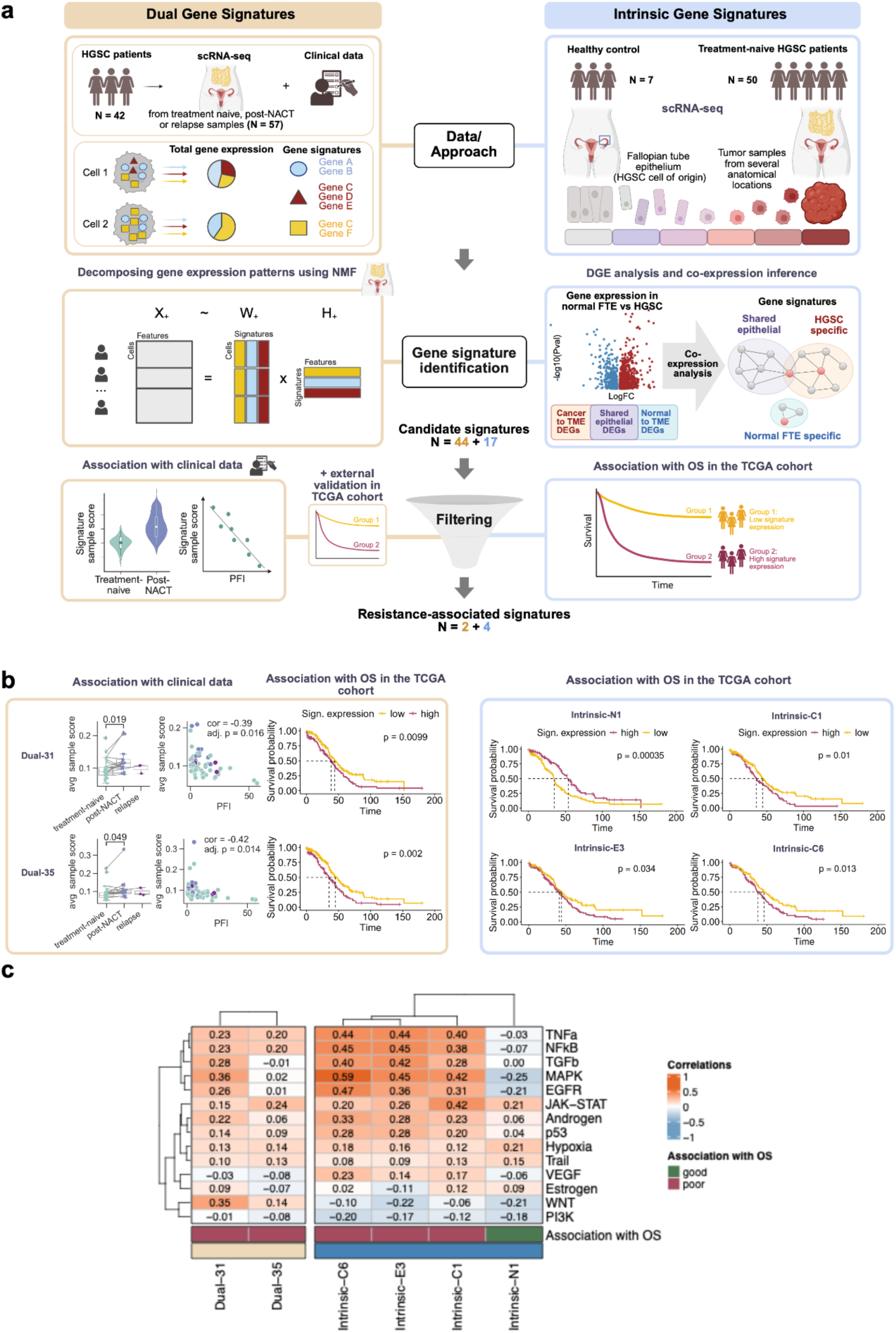
Identification of adaptive and intrinsic transcriptional signatures associated with platinum resistance. **a,** Overview of the dataset and analytical framework to identify cancer cell subpopulations driving platinum resistance. Dual signatures are derived from a global decomposition of all gene expression data into 44 signatures, while 17 intrinsic signatures are constructed from co-expressed genes differentially expressed between cancer and normal epithelia. Both sets were filtered for association with clinical outcomes, yielding six resistance-associated signatures. **b,** Kaplan-Meier survival curves of the six signatures significantly associated with OS in the TCGA cohort (n=258). For dual signatures, boxplots show expression in post-NACT versus treatment-naive and relapsed samples (n=54), and scatterplots display correlation with PFI (n=57). **c,** Single-cell level Spearman correlations between resistance-associated signatures and PROGENy pathway activities. The correlations were calculated at single-cell level (n=36,229 cells for dual; n=30,236 cells for intrinsic).

The *dual* approach identifies transcriptional signatures associated with early adaptive and intrinsic resistance by integrating data from treatment-naive and post-treatment samples. We applied non-negative matrix factorization (NMF; Lee and Seung 1999) to decompose tumor gene expression profiles into 44 gene signatures (Methods, Extended Data Fig. 1, Supplementary Table 3). Signatures were retained if they met three criteria: differential expression after chemotherapy (Extended Data Fig. 2), association with short treatment response within the study cohort (Extended Data Fig. 2), and association with overall survival (OS) in The Cancer Genome Atlas (TCGA) treatment-naive tumors, suggestive of intrinsic resistance. Two signatures, Dual-31 and Dual-35, fulfilled all criteria. Both were enriched in post-NACT samples (Fig. 2b), consistent with either expansion of signature-positive cells or induction of the signatures after chemotherapy. Across all treatment phases, both signatures negatively correlated with platinum free interval (PFI; Fig. 2b). In the TCGA cohort, tumors with high Dual-31 and Dual-35 expression had significantly shorter OS than low-expressing tumors (Fig. 2b).

The *intrinsic* approach compares treatment-naïve tumors with normal fallopian tube epithelium to identify co-expression signatures that capture intrinsic resistance and reveal programs lost, maintained, or gained during malignant transformation. This strategy is built on our recent framework for mapping HGSC cell-state dynamics and resistance (Pirttikoski et al. 2025). We first identified genes enriched in (i) normal fallopian tube epithelial (FTE) cells, (ii) cancer epithelial cells, or (iii) core epithelial patterns conserved across normal and tumor subpopulations, all overexpressed relatively to stromal and immune cells within the tumor microenvironment (TME). Co-expression patterns within individual tumors were clustered to mitigate inter-tumoral heterogeneity, yielding 17 signatures: four normal FTE-specific, nine cancer-specific, and four shared epithelial programs (see Methods; Extended Data Fig. 3 ; Supplementary Table 4). To isolate signatures linked to intrinsic platinum resistance, we focused on those associated with overall survival in the TCGA cohort of treatment-naive tumors. High expression of Intrinsic-C1, Intrinsic-C6 and Intrinsic-E3 correlated with shorter OS, whereas the normal FTE-specific Intrinsic-N1 signature correlated with longer OS (Fig. 2b).

To explore the biological functions represented by the six resistance-associated signatures, we correlated signature expression with inferred PROGENy pathway activities in clinical specimens (Fig. 2c; signature expression in Extended Data Fig. 4). Strong correlations (P < 1.0 × 10⁻¹⁶) were observed between Dual-31, Intrinsic-C6, Intrinsic-E3 and Intrinsic-N1 with the MAPK pathway (Spearman ⍴= 0.36, 0.59, 0.45, and –0,25 respectively), and between Intrinsic-C1 and Dual-35 with the JAK-STAT pathway (Spearman ⍴= 0.42 and 0.24, respectively). Comparison with Cancer Hallmark pathways revealed two major clusters, one enriched for inflammatory and growth factor response and the other for metabolic and DNA repair activity (Extended Data Fig. 5). Notably, all resistance-associated signatures aligned with the inflammatory/growth factor cluster, whereas the only signature associated with improved survival, Intrinsic-N1, clustered with the metabolic and DNA repair cluster, consistent with our previous analysis (Zhang et al. 2022). The heterogeneous expression of these signatures across patients suggests that different tumors may require distinct adjuvant strategies, enabling signature-based patient stratification.

### Computational Prediction of Drugs Synergizing with Carboplatin to Target Molecular Vulnerabilities in Ovarian Cancer

To identify drugs targeting platinum-resistant phenotypes, we used two complementary strategies (Fig. 3a). The *signature reversion* strategy screened perturbational profiles to identify compounds or druggable genes whose signatures anti-correlate with resistant profiles, suggesting potential to convert resistant phenotypes into sensitive states. Perturbation datasets included LINCS L1000 and Perturb-seq (Subramanian et al. 2017; Szalai et al. 2019; Replogle et al. 2022). The *signature-based killing* strategy used unperturbed expression and cell line sensitivity data to identify compounds predicted to target platinum-resistant phenotypes. We leveraged two large drug screening datasets: the Genomics of Drug Sensitivity in Cancer (GDSC) dataset (Garnett et al. 2012) and the Profiling Relative Inhibition Simultaneously in Mixtures (PRISM) Repurposing dataset (Corsello et al. 2020). Candidate drugs and targets were filtered for HGSC relevance, including tumor expression, druggability, translational potential, and non-redundancy to generate a refined list for experimental testing (see Methods). The final list comprised 64 drugs (Fig. 3b). Of these 55% (n=35) targeted individual signatures, whereas the 45% (n=29) targeted multiple signatures (Extended Data Fig. 6 a). Most drugs (80%, n=51) were predicted by a single data source, while 20% (n=13) were supported by two or three sources (Extended Data Fig. 6 b). GDSC contributed the largest fraction of predictions (52%, n=33), and included all 13 drugs predicted by multiple data sources. The candidates span diverse drug classes, with signal transduction inhibitors being most prevalent.

**Figure 3.**
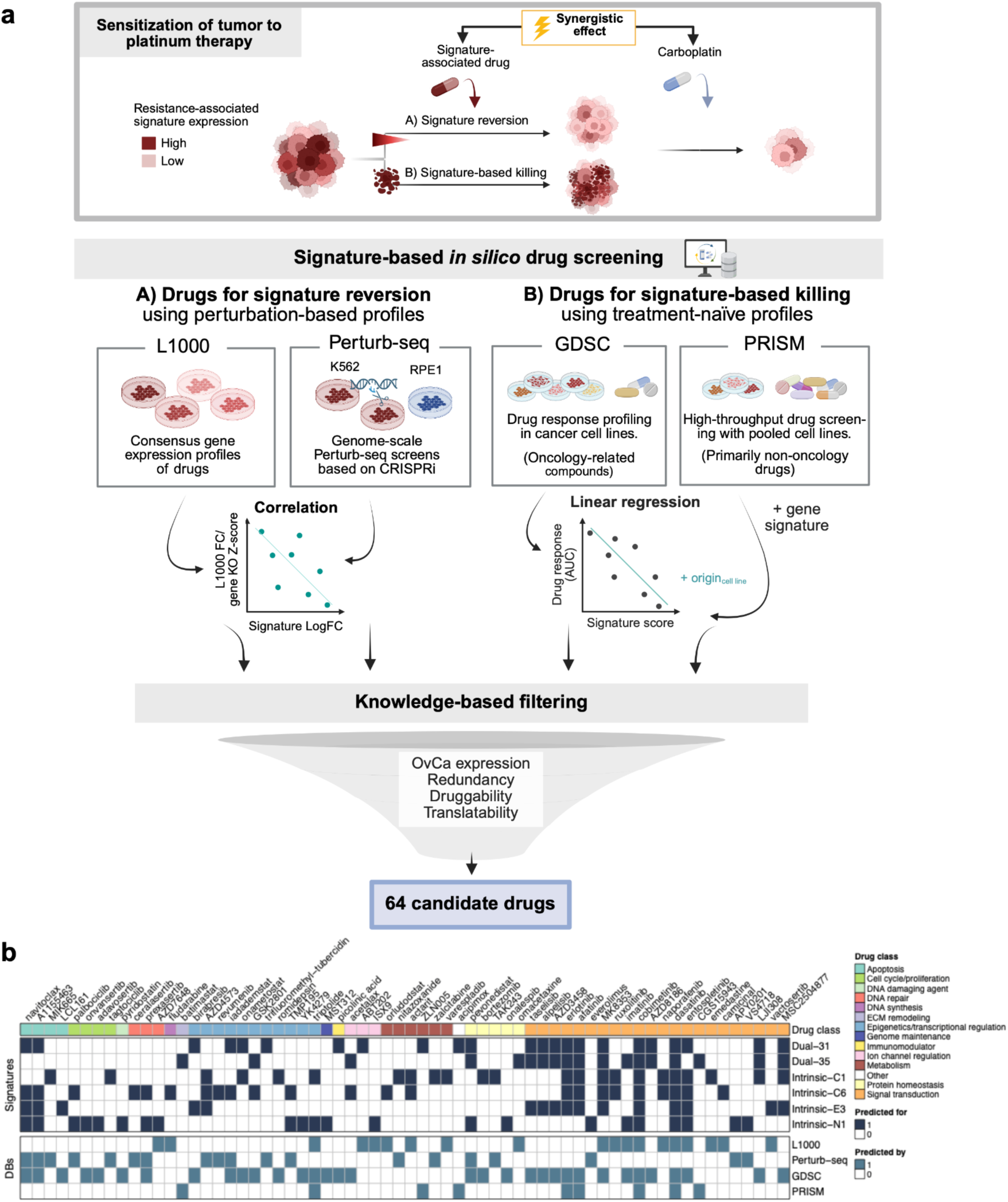
Algorithmic identification of synergistic drugs from resistance-associated signatures. **a,** Overview of two computational strategies to identify drugs targeting carboplatin-resistant phenotypes. The first leverages perturbation-based datasets to nominate compounds predicted to revert resistant signatures to a sensitive state, whereas the second uses drug screening datasets to identify compounds selectively cytotoxic to resistant subpopulations in treatment-naive profiles. **b,** Heatmap of the final 64 candidate drugs after knowledge-based filtering, showing their associated signatures and the database of origin. Drugs are categorized by their primary biological functions.

### Validation of Drug Combinations in HGSC PDOs Identifies Carboplatin Sensitizers

After nominating 64 candidate drugs, we assessed carboplatin synergy in validated HGSC PDOs established from primary tumor material (treatment-naïve) (Senkowski et al. 2023) (Supplementary Fig. 1). To identify carboplatin sensitizers, we screened two chemo-naïve PDOs derived from ascites with distinct resistant signature profiles (Fig. 4a–b + Source data_Fig. 4). Each candidate drug was tested at two concentrations with a four-fold difference, selected to yield a weak-to-moderate single-agent effect. The concentrations were chosen based on previous knowledge and/or bibliographic reviewing. As the standard carboplatin partner in HGSC treatment, Paclitaxel was included as a reference.

**Figure 4.**
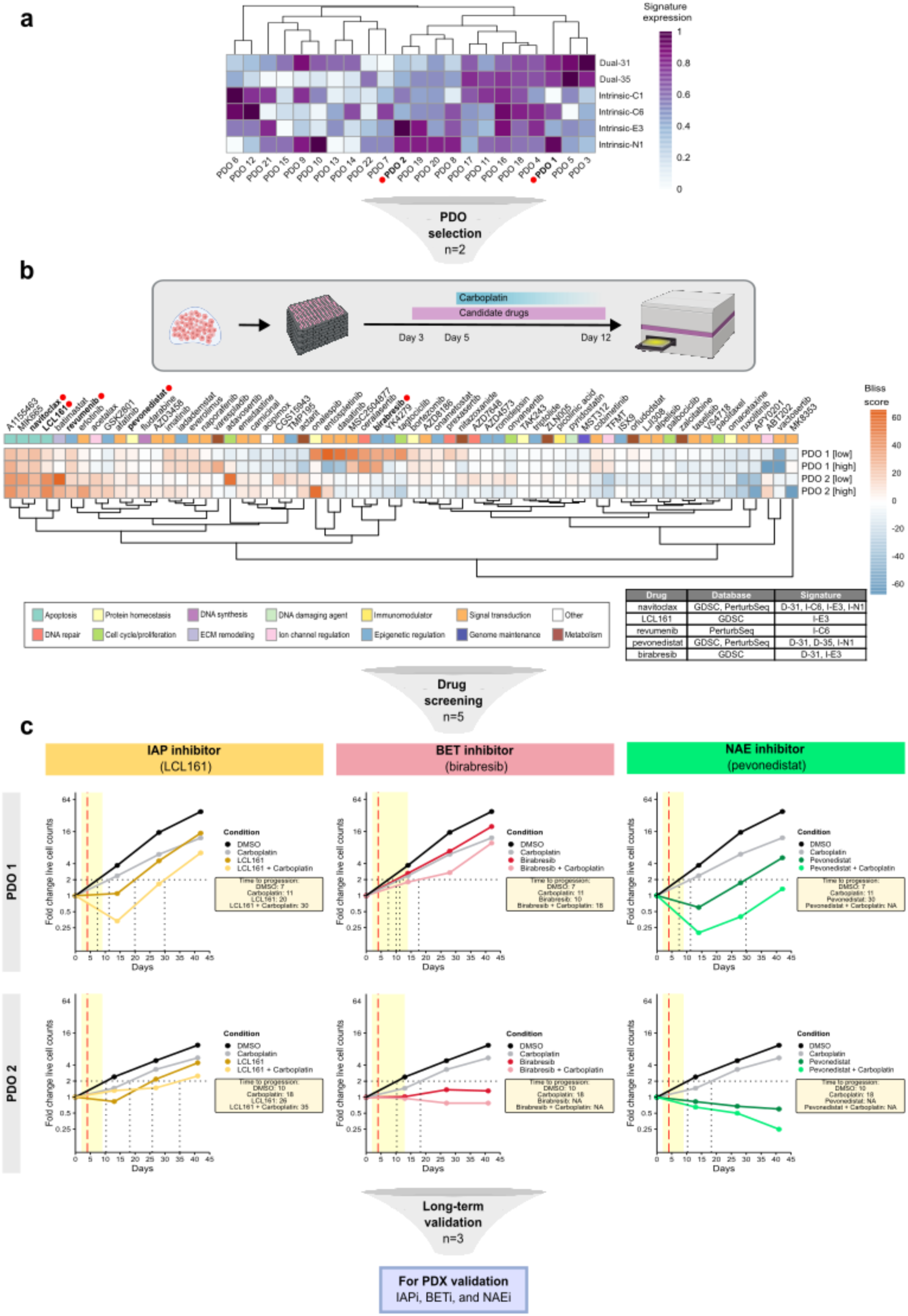
Drug screening and long-term validation of candidate drugs in HGSC PDOs. **a,** Normalized expression of resistance-associated signatures in 22 HGSC PDO models from RNA-Seq. **b,** Drug screening pipeline and heatmap of the Bliss scores in PDO1 and 2. Columns are hierarchically clustered, based on Euclidean distances. **c,** Long-term growth plots for PDO1 and 2 treated with drugs that synergize with carboplatin. The red dashed line indicates carboplatin treatment, and the yellow shaded area indicates treatment with the drug combination. Time-to-progression corresponds to the number of days required for the PDO cultures to double the initial number of viable cells seeded. Abbreviations: D; dual. I; intrinsic.

Following the screen in PDO1, the doses of 12 out of 65 drugs were adjusted due to excessive single-agent toxicities (Supplementary Table 5). PDOs were pre-treated for two days before carboplatin, and viability was quantified 7 days later. The effect of the combinatorial treatments was evaluated using the Bliss independence model (Bliss 1939) to determine the interaction type between each candidate drug and carboplatin. Most (80.3%) combinations were additive (–10 < Bliss < 10), but 16.5% were synergistic (Bliss ≥ 10) when aggregating the synergy calls from both concentrations (Extended Data Fig. 7 + Source data Extended Data Fig. 7). The identified drug synergies were either model-specific or shared between both PDOs (Fig. 4b + Source data_Fig. 4). Shared synergistic carboplatin combinations were found through inhibition of anti-apoptotic proteins via navitoclax (Bcl-2/Bcl-xL/Bcl-W inhibitor) and inhibition of IAPs via LCL161 (pan-IAP inhibitor; XIAP/cIAP1/cIAP2). Revumenib, a menin-MLL interaction inhibitor, also showed synergy in the short-term screening. Model-specific effects included synergy with the BET inhibitor birabresib (BRD2/3/4) in PDO1. Although carboplatin combined with the neddylation activating enzyme inhibitor (NAEi) pevonedistat only elicited an additive effect in PDO2, the drug was selected for further validation based on prior evidence of carboplatin sensitization in the HGSC cell line Kuramochi (Dai et al. 2024). The number of predicted synergistic interactions varied across the databases, with PertubSeq yielding the highest discovery rate of synergistic drugs (24.3%), followed by GDSC (21.2%) and PRISM (16.7%). On the contrary, drugs predicted from the L1000 database were rarely synergistic (7.9%) (Extended Data Fig. 7 + Source data Extended Data Fig. 7).

In summary, the five drugs that validated in short-term PDOs (birabresib, LCL161, navitoclax, pevonedistat, revumenib) originated from the per-cell line GDSC dataset and/or from gene knockdown-based Perturb-Seq data, reflecting their closer tissue/context alignment and mechanistic proximity to the pathways inferred from scRNA-seq. In contrast, predictions derived from the broader PRISM or L1000 resources—dominated by pooled viability assays or bulk compound signatures with greater polypharmacology, less specific drug collections (PRISM), and limited ovarian representation (L1000)—did not reproduce in the PDO models tested.

To assess the long-term effect of the top drug-screening hits (n=5) on tumor re-growth, we conducted survival assays with PDOs. PDO were treated with candidate drugs, as monotherapy and in combination with carboplatin, and replated at the initial cell density at each passage for 41-42 days. Inhibition of neddylation, induction of apoptosis through IAP inhibition, and suppression of oncogene transcription through BRD inhibition impaired cancer cell re-growth when combined with carboplatin (Fig. 4c + Source data Fig. 4 + Supplementary Table 6). This effect was observed in both PDO models, as indicated by extended periods with cancer cell regression or suppressed growth. By contrast, Bcl-2 family inhibition via navitoclax and menin-MLL blockade via revumenib failed to produce durable growth suppression. In fact, the latter even promoted long-term growth in one of the PDOs (Extended Data Fig. 8 + Source data Fig. 4). Based on these findings, BET, NAE, and IAP inhibitors were advanced to orthotopic PDX studies in combination with carboplatin. To enhance the translational relevance for *in vivo* studies, we replaced birabresib with the pan-BET inhibitor ZEN-3694 (BRD2/3/4/T), which was under evaluation in several phase 1 and 2 clinical trials for HGSC patients with recurrent disease or carboplatin resistance (NCT05950464, NCT04840589, NCT05071937). Similarly, LCL161 was substituted with xevinapant, another monovalent pan-IAP inhibitor in late-stage clinical testing for head and neck cancer (NCT06145412). Both substituted compounds demonstrated synergy comparable to the original compounds in five additional PDO models (Extended Data Fig. 9 + Source data Extended Data Fig. 9 + Supplementary Table 7), arguing for on-target effects.

### Validation of Drug Combinations in HGSC PDXs suggests Pevonedistat as Carboplatin Sensitizer

Next, we examined BET (ZEN-3694), NAE (pevonedistat), and IAP (xevinapant) inhibitors (Fig. 4c) in orthotopic PDX models following the workflow described earlier (Popa et al. 2022). Tumor-specific nucleotide polymorphisms were conserved across passages from donor tumors, confirming preservation of the patient-specific molecular landscape in the corresponding PDX models (Fig. 5a). PDX models were selected for their distinct expression of dual and intrinsic transcriptional signatures associated with platinum resistance (Fig. 5b). PDX material was implanted orthotopically at week –4 and tumor growth was monitored by bioluminescence imaging (BLI). By week –1, all mice exhibited detectable signals in the right ovary ovarian tumor (Fig. 5d). Following confirmation of stable engraftment, 32 mice were randomized and allocated into five groups with comparable baseline BLI (Fig. 5c, d + Extended Data Fig. 10 a). PDX mice receiving combinatorial treatment were pretreated with the candidate drugs for 3 days before initiation of carboplatin therapy.

**Figure 5.**
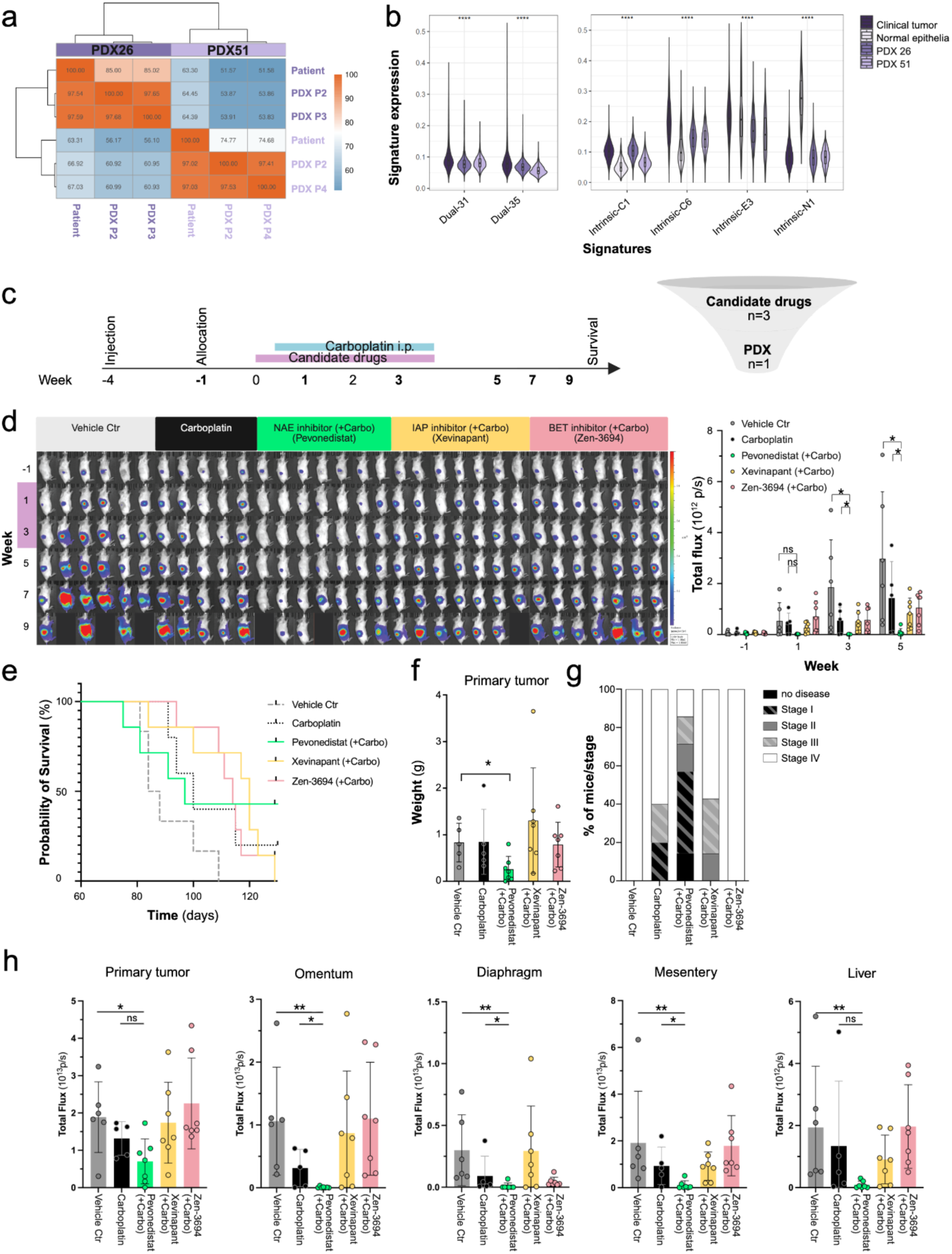
Therapeutic validation in PDX models. **a.** Percentage of SNPs shared between patient-derived HGSC xenografts and the original patient tumors, calculated row-wise. **b.** Signature expression at single-cell level contrasting (left) dual signature expression in patient-derived HGSC xenograft models compared to clinical data set (n= 36229 from 42 clinical ovarian tumors; n = 10252 from PDX26; n = 1866 from PDX51) with (right) intrinsic signature expression (n= 30236 from 50 clinical ovarian tumors; n=1251 normal epithelial cells from seven healthy controls, PDX26: n = 10252; PDX51: n = 1866). Statistical significance of signature expression between groups was assessed using a Kruskal–Wallis test; all comparisons shown were significant (p ≤ 0.0001). **c.** Experimental timeline for orthotopic PDX generation, allocation to five treatment groups, biweekly bioluminescence imaging (BLI)) and treatment (n=32). **d.** Biweekly BLI of PDX26-implanted mice in the lateral position. Each column represents an individual mouse (left). The total lateral photon flux before, during and after treatment of each mouse presented for each treatment group (right). Unpaired t-test, two-tailed; p<0.05. **e.** Kaplan-Meier curves for tested drugs. **f.** Endpoint tumor burden across treatment groups, shown as primary tumor weight. Unpaired t-test, two-tailed; p<0.05. **g.** Disease stage (I-IV) at survival endpoint, based on ex vivo BLI and necropsy-confirmed tumor dissemination presented as percentage of mice across disease stages. **h**. Bioluminescence signal intensity (total photon flux) of each organ *ex vivo*. Unpaired Mann-Whitney test.

Longitudinal biweekly BLI of PDX26 showed consistent tumor growth and progressive disease in the control mice, whereas PDX26 mice treated with carboplatin plus the NAE inhibitor pevonedistat exhibited reduced BLI at week 3 and 5 throughout treatment (Fig. 5d + Extended Data Fig. 10 b,c). Although metastatic spread was first detected in the vehicle control arm (Extended Data Fig. 10 c), the other two candidate drugs, ZEN-3694 (BETi) and xevinapant (IAPi) in combination with carboplatin showed no additional benefit in reducing the primary tumor size (Fig. 5d), inhibiting metastatic dissemination (Extended Data Fig. 10 c), or improving survival (Fig. 5e). The tumor growth suppression observed in the pevonedistat plus carboplatin group resumed after treatment discontinuation at week 5 and did not confer a significant survival benefit (Fig. 5e + Extended Data Fig. 10 b,c). However, at the study endpoint, *ex vivo* measurements showed that the combination of pevonedistat and carboplatin significantly reduced weight of the primary tumor (Fig. 5f), while the other treatment groups did not show significant differences. To quantify the extent of the metastatic spread across the groups in PDX26 in a clinically meaningful manner, we applied a modified FIGO staging classification based on BLI and necropsy findings (Fig. 5g, Extended Data Fig. 10 d). Importantly, this revealed a lower overall disease stage in the carboplatin-pevonedistat group compared to other groups (Fig 5g). The *ex vivo* BLI and necropsy reports exposed extensive dissemination to multiple abdominal organs in vehicle control and ZEN-3694-treated mice (Fig 5g,h, Extended Data Fig. 10 d). Furthermore, the total intensity of the bioluminescence signal across all organs was significantly lower in the pevonedistat plus carboplatin cohort, indicating a reduced extent of disease dissemination involving only the diaphragm, omentum and contralateral side, respectively in one animal each (Fig. 5h, Extended Data Fig. 10 d), consistent with *in vivo* observations. Although this combination did not improve overall survival, pevonedistat effectively reduced metastatic dissemination. This indicates that selective inhibition of chemoresistance-associated transcriptional programs can potentiate carboplatin efficacy in preclinical HGSC models. The alignment between signature-defined vulnerabilities and *in vivo* responses illustrates how our discovery framework can nominate different combinations for different patients depending on the resistance programs active in their tumors.

## Discussion

Resistance to platinum-based chemotherapy remains the central obstacle to cures or durable disease control in high-grade serous carcinoma (HGSC), even though many patients initially experience a complete or partial response. Because platinum resistance arises from both pre-existing and therapy-induced cellular programs, systematic approaches are needed to disentangle this heterogeneity and identify targetable vulnerabilities alongside standard-of-care treatment. Here, we developed a carboplatin-anchored framework that leverages single-cell transcriptomic data to resolve intrinsic (pre-existing) and adaptive (therapy-induced) resistance programs in clinical tumors, yielding six signatures associated with overall survival in an independent cohort. These signatures guided two complementary *in silico* screens across four resources (L1000, Perturb-seq, GDSC, PRISM) to nominate agents predicted either to revert resistance programs or selectively target resistant subpopulations. From the 64 candidates functionally screened in PDOs, three compounds enhanced carboplatin activity also in long-term PDO assays, of which one drug, pevonedistat, significantly reduced tumor burden *in vivo*.

Several recent tools, such as BeyondCell, scDrug, DREEP, and scTherapy (Fustero-Torre et al. 2021; Hsieh et al. 2023; Pellecchia et al. 2023; Ianevski et al. 2024), use single-cell transcriptomes to inform treatment selection. Our study advances this concept in five key ways: (i) clinical anchoring—predictions explicitly centered on carboplatin, the backbone of HGSC therapy; (ii) program-level targeting of continuous, non-binary resistance states rather than discrete clusters; (iii) temporal precision by incorporating intrinsic and dual signatures, including post-NACT patterns, to enable stage-specific intervention; (iv) orthogonal data integration by combining viability datasets (GDSC, PRISM) with perturbational transcriptomics (L1000, Perturb-seq) to identify mechanism-matched, context-specific, and less widely explored agents; and (v) a systematic validation funnel—from short- and long-term PDO assays to orthotopic PDXs—that brings predictions to preclinical proof-of-concept. This combination distinguishes our approach as a directly translatable framework for platinum resistance in HGSC.

Unlike approaches that map drugs to discrete cell clusters (Ianevski et al. 2024; Hsieh et al. 2023), we focused on continuous transcriptional programs that capture subtle state gradients underlying platinum resistance. Program-level signatures generalized across patients and retained prognostic value, enabling mechanism-informed drug nomination beyond what cluster labels typically provide. By explicitly distinguishing signatures present at baseline from those induced after neoadjuvant chemotherapy, our framework supports stage-specific intervention strategies: adjuvants can be selected to pre-empt baseline-resistant states or to counteract adaptive reprogramming emerging after platinum exposure.

Like previous approaches used for drug prediction (Fustero-Torre et al. 2021; Hsieh et al. 2023; Pellecchia et al. 2023), we used multiple databases, integrating viability data (GDSC, PRISM) with perturbational transcriptomics (L1000, Perturb-seq) to mitigate biases inherent to any single dataset. We did not observe consistent differences between databases presenting viability and perturbational transcriptomics readouts. However, drugs predicted from GDSC and Perturb-Seq validated in short-term PDOs better reflect relevant tissue context and functional drivers of resistance, respectively. In contrast, predictions from PRISM and L1000—based on pooled or bulk assays with broader, less specific compound profiles and limited HGSC representation—did not reproduce in PDO models. This finding is consistent with DREEP validation in head and neck squamous cell carcinoma, where GDSC-based predictions outperformed those from PRISM (Pellecchia et al. 2023). Overall, these observations indicate that lineage representation, assay design, and perturbation type are all key determinants of which *in silico* predictions translate experimentally, underscoring that database selection and calibration to tumor context are critical design decisions in signature-based drug discovery (see Supplementary Table 10 for a comparative overview).

PDO validation revealed several recurring mechanisms. Compounds targeting the Bcl-2 family (navitoclax, A1155463, MIK665) or IAP proteins (LCL161, xevinapant) enhanced carboplatin response, highlighting apoptotic resistance as a key driver of short-term drug resistance. However, Bcl-2 inhibition was not synergistic in the long-term cultures. Synergy with pro-apoptotic agents is also frequently observed in large-scale screens (Jaaks et al. 2022), raising the concern that such combinatorial effects may be generic rather than cancer cell-specific. Consistent with this, clinical development of most agents in this class, except for the hematopoietic-selective inhibitor venetoclax, has been constrained by narrow therapeutic windows. BET inhibitors (birabresib, ZEN-3694) also potentiated carboplatin activity in long-term survival assays with PDOs, underscoring epigenetic regulation as an actionable feature. BET inhibitors are currently being evaluated in clinical trials across multiple haematological and solid malignancies, including ovarian cancer, often in combination with the PARP inhibitors olaparib and rucaparib, due to reported drug synergy independent of the HR status (Andrikopoulou et al. 2021). Agents that disrupt proteostasis (pevonedistat, onalespib) similarly enhanced carboplatin response and induced apoptosis among multiple reported mechanisms (Fu and Wang 2023). Pevonedistat has been extensively tested in clinical trials, including in combination with carboplatin-paclitaxel chemotherapy, with promising phase Ib/II results in solid tumors, though its development has been limited by toxicities (Fu and Wang 2023)(Lockhart et al. 2019; Millett et al. 2022). In contrast, drugs that inhibit proliferation, such as paclitaxel, were not synergistic with carboplatin, highlighting that *in vitro* synergy can be highly context-dependent and does not always mirror clinical benefit (Palmer and Sorger 2017). Finally, the carboplatin synergies observed with birabresib and LCL161 were reproduced with ZEN-3694 and xevinapant, respectively, supporting on-target effects. Overall, PDO assays revealed actionable resistance mechanisms (apoptosis, epigenetics, proteostasis), and filtered out transient drug effects. Orthotopic PDX models then enabled assessment of tumor control and tolerability *in vivo*. Together, this tiered strategy prioritized pevonedistat as the most consistent carboplatin adjuvant and illustrates a pragmatic path from single-cell programs to translational candidates.

In PDOs, pevonedistat enhanced the long-term growth-suppressive effect of carboplatin, and, in PDX26, the combination significantly reduced tumor growth and metastatic spread, even at a submaximal dose (Kong et al. 2022). By contrast, carboplatin combined with ZEN-3694 or xevinapant did not yield comparable benefit. All regimens were tolerated, but pevonedistat’s formulation and dosing constraints (poor solubility, administration route, conservative dosing) likely underestimated achievable efficacy, highlighting both the promise and the practical challenges of translating signature-predicted vulnerabilities into clinically viable regimens.

Despite the strengths of our integrative pipeline, several experimental and analytical constraints temper the interpretation of our findings. The main PDO cohort was small, ascites-derived, and consisting solely of cancer cells, with an HR-proficient profile. Our PDX studies prioritized tumor dissemination and control, rather than survival outcomes. Finally, signatures derived from epithelial cancer cells may not capture immune- or mesenchymal stromal cell-derived resistance mechanisms that remain inaccessible in these preclinical systems.

This study establishes a framework for overcoming platinum resistance in HGSC through a carboplatin-anchored strategy that integrates molecular profiling and functional modeling. By directly linking patient-derived resistance signature profiles to effective drug combinations, this approach enables rational design of synergistic regimens with preclinical efficiency in PDOs and PDX models. Advancing stratified, carboplatin-anchored combinations will require expanding PDO and PDX diversity—including immune-competent and humanized systems (Kleinmanns et al. 2022)—together with proteomic and spatial profiling to uncover non-transcriptional resistance mechanisms. Parallel optimization of clinically feasible dosing (e.g., for pevonedistat), and translation of molecular signatures into actionable biomarkers will enable patient-tailored treatment adaptation. More broadly, this work illustrates how systematic integration of molecular profiling and functional modeling can transform platinum-chemotherapy from a fixed standard into a more dynamic, precision-guided platform with potential applications across resistant malignancies.

## Methods

### Patient-Derived Single-Cell Transcriptomics

Tumor and ascites (n=72) along with normal fallopian tube or fimbriae specimens (n=7), were collected from 61 patients enrolled in the prospective clinical trial DECIDER NCT04846933 at Turku University Hospital, Finland. Samples were dissociated and processed for scRNA-seq as previously described (Zhang et al. 2022). Detailed information regarding samples, associated clinical data, anatomical locations, and previous publications presenting subsets of the scRNA-seq data is summarized in Supplementary Table 1. Written informed consent was obtained from all patients contributing HGSC or normal FTE samples for the study. The DECIDER study was approved by the Wellbeing Services County of Southwest Finland (VARHA/28314/13.02.02/2023).

### Signature Expression Scores

We calculated signature expression scores similar as in AUCell (Aibar et al. 2017). For the deconvolution-based signatures from the *dual approach* (see below), which are defined by continuous weights, we employed a weighted version of AUCell utilizing the normalized signature gene scores. In contrast, the signatures from the *intrinsic approach* (see below) were scored using the unweighted version of AUCell method implemented through the UCell R package v2.8.0 with default settings (Andreatta and Carmona 2021). We used this rank-based approach to calculate single-cell expression scores from gene sets, as it effectively quantified gene signature expression across diverse samples and experimental datasets.

To obtain signature expression in scRNA-seq data, we considered the top 1,500 genes with highest rank (as defined in UCell) to capture only expressed genes and avoid introducing noise from low or unexpressed genes. For bulk RNA-seq data, we increased the number of genes to 20,000, as bulk data captures a larger number of expressed genes.

### Computational Identification of Transcriptomic Signatures

We used two approaches - *dual* and *intrinsic* - to derive resistance-associated gene expression signatures from largely identical subsets of the scRNA-seq data.

#### Dual approach

The dual approach focused exclusively on HGSC samples from various treatment stages, comprising 57 samples from 42 patients. This included treatment-naive samples (n=37) collected at laparoscopy or primary debulking surgery (PDS), post-NACT samples (n=17) collected at interval debulking surgery, and relapse samples (n=3) collected at progression. For 27 patients, only a single sample was available, while 15 patients had paired samples collected at different time points during their disease trajectory. For these patients, clinical data included the platinum-free interval (PFI), defined as time from completion of primary treatment to relapse.

The scRNA-seq data for these samples were preprocessed following the same workflow described in Launonen et al. (Launonen et al. 2024). We further filtered the preprocessed scRNA-seq data to retain only cells annotated as cancer cells and exclude genes detected in fewer than 30 cells.

We then applied non-negative matrix factorization (NMF) (Lee and Seung 1999) to decompose the scRNA-seq transcriptome of each cell approximately as a weighted sum of canonic gene signatures. Specifically, NMF represents the gene expression matrix as the product of two non-negative matrices: the *signature matrix H* that represents gene signatures by assigning a weight to each gene reflecting its contribution to a signature, and the *usage matrix W* that quantifies the level of expression of each signature in each cell.

We used the NMF implementation from scikit-learn (v.1.0.2) (Pedregosa et al. 2011) in Python, with default parameters except for init=”random”, max_iter=1000 and n_components=44, and applied it to the full count matrix after combining the samples by concatenating all cells. The number of components (signatures) was determined by running NMF across a range of n_components values (range 5 to 100, step size=3) and selecting an elbow point in the curve of the reconstruction error versus n_components (see Extended Data Fig. 1).

Prior to applying NMF, we used sctransform (v.0.3.4) (Hafemeister and Satija 2019) for variance stabilization and correction of technical variation. sctransform produces normalized expression values as Pearson residuals, which can include negative values. Because NMF requires non-negative inputs, we shifted the residuals by adding the model-derived fitted means for each gene and set any remaining negative values to zero. After applying NMF, we performed post-processing on signature and usage matrices. The result of NMF is invariant to scaling. Therefore, to ensure that the signature usages *W* are comparable within cells, we multiplied the columns of *W* by the sum of rows in the signature matrix *H*. Additionally, we ℓ_1_-normalized the signature usages for each cell (row) in *W*.

To make signature gene weights comparable within a signature, we computed normalized gene scores, which we used for all subsequent analyses. Since we can not mean-center the data prior to NMF, the gene weights in H are dependent on the scale of the gene expression. We thus computed an *R*^2^-like score that reflects the contribution of gene j in signature k to the overall variability in the expression of the gene. This normalized gene score for gene *j* and signature *k* was calculated as:

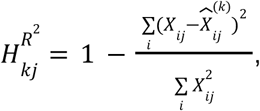

where *i* is the index for cells, and 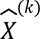 is the reconstructed matrix using only signature *k*. To avoid variation driven by noise, we then replaced values lower than 0.03 by 0.

From the resulting canonical NMF signatures, we selected those associated with resistance within the patient cohort using a two-step filtering approach. Sample-level signature expression was computed by averaging the cellular signature expression scores (see above) across all cells within the sample. *Step 1*: Assuming that chemotherapy enriches for resistant cells, resistance-related signatures are expected to show differential expression between treatment-naive and post-NACT samples. To test this, we compared the distribution of sample-level signature expression between treatment-naïve and post-NACT samples using a Mann-Whitney U test (scipy.stats.mannwhitneyu, v.1.8.0) (Virtanen et al. 2020). However, this test cannot distinguish between resistance effects or unrelated treatment-induced effects and we also applied a second filtering step. *Step 2*: We further assessed correlations between sample-level signature expression and PFI, using Spearman correlations (scipy.stats.spearmanr, v.1.8.0) for all signatures that passed the first filtering step. P-values from both tests were adjusted for multiple testing using the Benjamini-Hochberg false discovery rate (FDR) procedure, as implemented in statsmodels.stats.multitest.multipletests (v.0.13.2) with method=’fdr_bh’.

#### Intrinsic approach

The intrinsic approach builds on our previous study of HGSC cell state dynamics and resistance, in which we investigated the epithelial origin of drug resistance using conserved gene modules (Pirttikoski et al. 2025). In this approach, we incorporated a single-cell RNA-seq dataset preprocessed in that study, comprising 59 samples from 57 patients—including 52 treatment-naïve HGSC tumor samples and 7 samples from postmenopausal healthy women serving as controls. HGSC samples were obtained from diverse anatomical sites such as the peritoneum, omentum, mesentery, ovary, ascites, adnexa, and fallopian tube, while control samples were collected from fallopian tubes or fimbriae.

Signature identification in the intrinsic approach consisted of three main steps: differential gene expression analysis, calculation of gene–gene associations, and clustering of genes based on these associations. These steps are described in detail in the following paragraphs.

Differential gene expression (DGE) analysis was performed using the MAST framework via the FindMarkers function in Seurat v4, with sample identity included as a latent variable. Differentially expressed genes (DEGs) were identified in three comparisons: between cancer cells and normal FTE cells, as well as between cancer cells and tumor microenvironment (TME) consisting of immune and stromal cells, and between normal secretory cells and TME. To reduce computational load, a maximum of 200 immune and stromal cells per sample were included in DGE testing. Genes differentially expressed in both cancer and normal FTE compared to TME were defined as shared epithelial genes. DEGs were filtered using a Bonferroni-adjusted p-value > 0.05 and averaged log2FC > 0.1. Only protein-coding genes were retained,excluding genes encoding ribosomal proteins and mitochondrial genes. To ensure generalizability, DGEs were further required to have more than one count in ≥ 30% of cells across ≥ 30% samples. After filtering, the final gene sets included 220 normal FTE-specific, 726 cancer-specific, and 301 shared epithelial genes.

Gene-gene associations were calculated using the R package proper, applying the proportionality metric p (Quinn et al. 2017). Because log-ratio transformation cannot handle zero values, one was added to the original count matrices prior to analysis. Associations were calculated separately for each sample. A cancer-specific gene-gene association matrix was constructed by averaging associations between cancer-related genes across cancer samples. Similarly, a normal FTE-specific gene-gene association matrix was created by averaging associations between normal-specific genes in the control samples. Due to the imbalance in the sample counts, associations among shared epithelial genes were calculated separately in cancer and control samples. The two resulting matrices were then combined by averaging their means. Samples with low cell counts were excluded from the analysis: four cancer samples (sample2, sample8, sample36, and sample48) and one control sample (control7).

Gene signatures were constructed by clustering genes with similar expression profiles, based on the associations calculated in the previous step. Clustering was performed using the Leiden algorithm through the FindClusters function in Seurat, with preprocessing according to the standard Seurat workflow. Clustering was performed with several resolutions, and the results were visualized with the R package clustree (v0.4.4) (Zappia and Oshlack 2018). This visualization guided the selection of stable resolutions for the different gene groups (cancer-specific, normal-specific, and shared epithelial), although it was challenging to explicitly define the optimal threshold. A clustering resolution of 0.4 was set for normal-specific and shared epithelial genes, and 0.9 for cancer-specific genes. The silhouette approach was used to validate consistency within the clusters and estimate the average distance between the clusters. The R package factoextra (v1.0.7) was used for the cluster validation (Kassambara 2020). Silhouette information was computed with the function silhouette, and all genes having a negative silhouette width were removed from the clusters. In total, 12 normal-specific, 126 cancer-specific, and 36 shared epithelial genes have a negative silhouette width.

#### Pathway inference

We used the PROGENy v1.26.0 R package to infer the expression of 14 signaling pathways on our single-cell data set (Schubert et al. 2018). Expression of Hallmark gene sets (Liberzon et al. 2015) was calculated using UCell. We compared pathways and gene set expression with our signature expression using Spearman correlations, calculated on the single-cell level, to investigate the biological properties of the identified gene signatures. Correlation P-values were adjusted for multiple testing using the Benjamini-Hochberg procedure.

#### Prognostic significance based on bulk TCGA RNA-seq

The prognostic relevance of the gene signatures was evaluated using deconvoluted bulk TCGA RNA-seq data from primary tumors of 258 HGSC patients (grade: G3 and G4) with available OS information (Zhang et al. 2022; Häkkinen et al. 2021). OS was defined as time from diagnosis of HGSC to death or last follow-up.

Patients were stratified into two groups based on optimized gene signature expression thresholds, determined using the max.stat (v.0.7.25) R package (Hothorn and Lausen 2003). Survival analysis was performed using the survival (v3.7.0) and survminer (v0.4.9) R packages.

### Computational Prediction of Drugs

To prioritize carboplatin adjuvant drugs on the basis of the identified putative resistance signatures, we applied two complementary strategies: (1) identifying drugs that revert resistance signatures and (2) identifying drugs that selectively target and eliminate cells expressing resistance signatures.

#### Signature reversion strategy

To identify compounds capable of reversing gene signature expression, we used perturbation-driven gene expression dataset from the LINCS L1000 project, compromising 2866 drug consensus signatures across multiple cell lines, doses, and time points based on 978 landmark genes (Subramanian et al. 2017; Szalai et al. 2019). Based on the Perturb-seq screens, our objective was to identify genes acting as drivers of the signatures and connect these genes to drugs that specifically target them. The Perturb-seq datasets consisted of transcriptomic profiles resulting from genetic perturbations, which were measured in chronic myeloid leukemia (CML) K562 cell lines and retinal pigment epithelial (RPE1) cell lines (Replogle et al. 2022). We used both essential- and genome-wide CML K562 datasets including 2057 and 9867 unique genetic perturbations respectively, as well as essential-wide RPE1 dataset including 2393 genetic perturbations.

We compared our transcriptional signatures to drug consensus signatures from LINCS L1000 and perturbed genes from Perturb-seq screens by transforming narrow gene signature groups into genome-wide ordered gene lists. To achieve this, we identified the top 25% of cells with the highest signature expression as signature-active cells and calculated differential expression between these cells and all other cells using the Seurat function FindMarkers(test.use=’LR’, latent.vars=c(“sample”, “S.Score”, “G2M.Score”). We calculated Spearman correlation using logFCs between the signature gene list and L1000 drugs as well as perturbed genes, saving those with an adjusted p-value of less than 0.05. For L1000 drugs, we applied a correlation coefficient threshold of −0.2, while a threshold of −0.1 was used for genes derived from Perturb-seq. These thresholds were chosen to balance the number of candidate targets identified across both databases, addressing the observation that there were fewer significant hits in the comparison with Perturb-seq than with L1000 data. As one of the gene signatures demonstrated a positive association with survival and as we aimed to promote the associated cellular state, we used positive correlation coefficient thresholds for this signature. The candidate driver genes from Perturb-seq comparison were further required to be differentially expressed in the cells showing high signature expression compared to the other cells.

#### Signature-based killing strategy

To identify drugs that showed specificity for cells with high (or low) signature expression, we used two publicly available datasets from large-scale drug screening studies: Profiling Relative Inhibition Simultaneously in Mixtures (PRISM) (Corsello et al. 2020) and Genomics of Drug Sensitivity in Cancer (GDSC) (Garnett et al. 2012). Both datasets contain treatment-naïve bulk gene expression profiles from human cancer cell lines and drug response for these cell lines measured as area under the dose-response curve (AUC). We used the secondary PRISM Repurposing dataset, where a broad drug library of approximately 1,500 drugs, primarily composed of non-oncology drugs, along with chemotherapeutics and targeted oncology agents, was tested across approximately 500 cell lines using a multiplexed barcoding approach, with cell viability inferred from changes in barcode abundance following drug treatment. The GDSC dataset 2 (GDSC) focused on about 300 oncology-related compounds, including targeted therapies and chemotherapies, which were tested on approximately 1,000 cell lines using a standardized ATP-based luminescence viability assay. We used only cell lines established from solid tumors (thus, excluding cell lines from haematological malignancies: leukemia, lymphoma, myeloma) and excluded all drugs screened in less than 100 cell lines or where the R2-value of the AUC was below 0.3, indicating insufficient goodness-of-fit of the dose-response model and unreliable AUC estimates.

To identify drugs for which the signature expression was a significant predictor of drug response, we employed a linear regression model, where drug response was modeled as a function of the signature expression, adjusted for the tissue of origin of the cell lines for all non-ovarian tissue origins. Signature expression scores were calculated as explained above on the treatment-naïve expression profiles of the cell lines. We created indicator variables for all categories of tissue origin with at least 10 cell lines, except for ovary-related tissue (urogenital_system for GDSC, ovary for PRISM). The model was implemented using the statsmodels library in Python and P-values were adjusted for multiple testing using the Benjamini-Hochberg FDR approach (statsmodels.stats.multitest.multipletests, v. 0.13.2).

For each signature, we retained only the predicted drugs with an adjusted p-value ≤ 0.05. Furthermore, we applied a sign filter to the predicted drugs, based on the association of the signature with survival outcomes. Specifically, for signatures associated with worse survival (i.e., resistance), we only considered drugs with negative coefficients, indicating that cells with higher signature expression are more likely to be killed. Conversely, for signatures associated with better survival, we retained the predicted drugs with positive coefficients to preferably target the low-signature expressing cells.

#### Tissue enrichment analysis

To evaluate tissue-specific drug sensitivity, we performed a tissue enrichment analysis. For each drug, a one-sided Mann-Whitney U test was used to determine whether the drug response (AUC) values of cell lines from a given tissue type were significantly lower than those of all other cell lines, indicating higher sensitivity. Resulting P-values were adjusted using the Benjamini–Hochberg false discovery rate (FDR) method.

#### Knowledge-based filtering of predicted drugs

Perturb-seq-derived targets were required to be expressed in HGSC and have a druggable product, ideally with clinically traceable compounds. For GDSC and PRISM candidates, tissue enrichment analysis was performed to assess whether drug-responsive cell lines were significantly enriched for HGSC (Supplementary Table 8 and Supplementary Table 9), whereas drugs showing preferential activity unrelated to tissue type were excluded. Finally, drugs with overlapping target profiles were removed to generate a refined list for experimental testing.

### Experimental Validations With Patient-Derived HGSC Organoid And Xenograft Models

#### Bulk RNA-Seq in HGSC PDOs

Confluent organoids growing in 6-well plates were harvested in technical duplicates by mechanical and enzymatic dissociation like in a regular passage. After centrifugation and supernatant aspiration, pellets were homogenized in 400 μL of 1X DNA/RNA Shield reagent (#R1200, Zymo Research), transferred to a microcentrifuge tube, and stored at −20°C. Total RNA was isolated with the Quick-RNA Miniprep Plus Kit (#R1058, Zymo Research) following the “Total RNA isolation” protocol from the kit’s manual. RNA concentration and purity was determined with a spectrophotometer (NanoDrop2000, Thermo Scientific). Long-non coding RNA (lncRNA) was sequenced with the NovaSeq X Plus Series (PE150) at Novogene Europe (Planegg, Germany) after additional quality control of the RNA integrity (RIN) and concentration re-assessment (Qubit). Library preparation included rRNA depletion. The sequencing depth was 40 M read pairs, leading to 12 Gbp of raw data per sample.

In the initial data exploration, paired-end RNA reads were quality controlled with FastQC (v0.12.1) (Andrews, S. 2010). The raw data was processed with SePIA (Icay, K. et al. 2016), a comprehensive data processing workflow for RNA sequencing. The first 12 and the last 5 bases of all reads were trimmed due to uneven base sequence content with Trimmomatic (v0.33) (Bolger, A.M. et al. 2014). Additionally, all leading bases with a quality score lower than 20 and all trailing bases with a quality score lower than 30 were removed. Then, all reads were scanned with a 3-base sliding window, cutting when the average quality dropped below 20 and discarding all sequences shorter than 20bp after trimming. All trimmed reads were aligned to the human reference genome (GRCh38.d1.vd1) using STAR (v2.7.6a) (Dobin, A. et al. 2013), allowing up to 10 mismatches and otherwise default parameters. Lastly, gene expression was quantified using eXpress (v1.5.1-linux_x86_64) (Roberts, A. 2013) and expressed as log2(TPM+1).

#### High-content drug screening in HGSC PDOs

Prior to drug screening, HGSC PDO cultures were re-established from cryopreserved cells using our standard culture method (Senkowski et al. 2023), and, at least, one passage was conducted to recover and expand the cultures. Briefly, PDO cells were thawed at 37°C, washed twice at 300×g for 5 min with 10 mL of M1 culture medium supplemented with 5 µM Y-27632 Rho kinase inhibitor (ROCKi) (MedChemExpress, #HY-10071) only in the second wash. Cell pellets were resuspended in basement membrane extract type 2 (BME-2) hydrogel (R&D Systems, #3533-005-02) diluted to a protein concentration of 8 mg/mL with ice-cold PBS and seeded in pre-warmed 6-well plates (Thermo Fisher Scientific, #140685), as 10 domes of 20 µL per well. Plates were incubated for 45 min to allow complete solidification followed by addition of a culture medium supplemented with 5 µM ROCKi. Medium was changed every 2-3 days without ROCKi.

For drug screening, confluent PDOs were harvested by mechanical and enzymatic dissociation like in a regular passage. Approximately 1 confluent well of a 6-well plate was used to seed each 384-well plate, to achieve an optimal cell density of the cultures. After medium aspiration, hydrogels were rinsed with 1 mL of pre-warmed PBS prior to adding 2 mL of TrypLE Express Enzyme (Gibco, #12604013). Domes were detached from the bottom of the plate with a cell lifter (Corning, #3008), pipetted 10-14 times using a P1000 pipette, and incubated at 37°C for 15 min. The cell digest was collected, wells were rinsed with 1 mL of pre-warmed PBS, and pelleted by centrifugation (300×g for 5 min). Cell pellets were resuspended in 8 mg/mL BME-2 and 10 μL were seeded in pre-cooled ultra-low attachment 384-well plates (30 min at −20°C + 15 min on ice) (Corning, #4588). Plates were incubated on ice for 15 min and at 37°C for 30 min to solidify the hydrogels. Pre-warmed culture medium supplemented with 5 µM ROCKi was added (40 μL per well), and plates were briefly centrifuged (300×g for 30 s) (Senkowski et al. 2023). Medium was changed on days 3, 5, and 9 with a microplate washer (Agilent, BioTek 405 LS Washer) using 30 µL/well without ROCKi. Drugs were dispensed at the time of media change using an acoustic liquid handler (Labcyte, Echo 550). Each treatment condition was tested in technical triplicates at two anchoring concentrations with a four-fold difference. Controls (0.1% DMSO; negative, 10 μM bortezomib; positive) and carboplatin (20 μM PDO1; 10 μM PDO2) were tested in 12 technical replicates. Viability was assessed on day 12 using the CellTiter-Glo 3D Cell Viability Assay (Promega, #G9683). After aspirating 25 μL medium, 25 μL of reagent was added per well. Plates were shaken for 5 min, incubated for 30 min in the dark, and luminescence measured with a plate reader (SpectraMax Paradigm, Molecular Devices) according to the manufacturer’s instructions.

#### Drug synergy quantification

For data analysis, raw luminescence intensity data was first inspected for outliers. For single and combination drug treatments, triplicates with high intrinsic variability were flagged using a normalized standard deviation (SD/mean CTG signal of the triplicate) threshold of 0.35 (manually determined). Within each flagged triplicate, outliers were identified by calculating the absolute difference of the CTG signal in pairs. For the controls (0.1% DMSO, 10 μM bortezomib) and carboplatin, which were performed in 12 replicates, outliers were identified by comparing each replicate to the group mean and SD. Intensity values deviating by 1.25-2 SD from the mean (manually determined and experiment-dependent) were excluded. Assay performance and plate quality were subsequently assessed by calculating Z’-factors of each screening plate.

Drug interactions between carboplatin and the candidate drugs were evaluated using the Bliss independence model (Bliss 1939). After data curation, viability was calculated by normalizing the luminescence of each triplicate to the negative and positive controls. In carboplatin, the mean viability across replicates was calculated. Then, all viability values were capped at 0 and 100. Expected drug effect under the Bliss assumption for combinatorial treatments was calculated as the product of the single-agent viabilities. Observed viability for carboplatin combinatorial treatments was determined as described above.

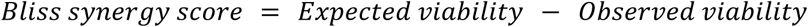

Bliss scores were calculated for each replicate and averaged across triplicates, and were used to assess the type of drug interaction: synergistic (Bliss ≥ 10), additive (–10 < Bliss < 10), and antagonistic (Bliss ≤ –10).

#### Long-term validation of drug screening hits in HGSC PDOs

Selected drug hits of clinical relevance were validated in long-term viability assays with PDOs. Confluent PDOs were harvested as described above. After, the digested PDO suspension was homogenized with a P1000 pipette and two 10 μL samples were collected to determine the number of live cells with a trypan blue exclusion test (Invitrogen, #T10282) and a Bürker chamber (Brand, #718905). Based on the standard split ratios, 3 × 10^5^ and 4 × 10^5^ live cells per well were seeded for PDO1 and PDO2, respectively, as 10 × 20-μL domes per well of a 6-well plate (1 well per condition). After the first medium change, PDOs were treated with the candidate drugs for 48 h followed by carboplatin as in the drug screening. Drug concentrations and schedules were determined based on clinical usage, drug screening results, and prior experimental experience (Supplementary Table 6). Drug treatments were applied only during the first passage. Afterwards, PDOs were maintained in a drug-free medium. PDOs were passaged when negative control reached confluency (day 13 for PDO2 and day 14 for PDO1). At each passage, live cells were counted in technical duplicates as explained above, and viable cells were re-seeded at the initial density, when possible. Otherwise, all available live cells were re-plated. PDOs were passed twice, yielding an experiment duration of 41-42 days.

#### *In vivo* evaluation of candidate drug combinations in orthotopic HGSC PDX models

Patient-derived HGSC xenograft models were established from treatment-naïve primary HGSC samples collected during primary debulking surgery at the Department of Obstetrics and Gynecology Haukeland University Hospital Bergen. The tumor samples were obtained through the Gynaecologic Cancer Biobank at the Women’s Clinic, Haukeland University Hospital, Bergen, Norway and were immediately processed into a single-cell suspension, cryopreserved and stored in liquid nitrogen. The biobank and associated research project were approved by the Regional Committee for Medical and Health Research Ethics (IDs: 2014/1907 and 2018/72), and written informed consents were obtained before data and tissue collection.

All animal experiments were performed in accordance with procedures approved by the Norwegian Commission for Laboratory Animals and the Norwegian Food Safety Authority (ID: 30736). Mice were bred at the University of Bergen animal facility and enrolled at 6-12 weeks of age. Immunodeficient female NOD-*scid* IL2*rγ^null^*(NSG) mice were orthotopically engrafted with 1 × 10^5^ HGSC cells derived from two established luciferase-positive PDX models (Kleinmanns, Bischof, et al. 2020; Popa et al. 2022). Primary patient cells had been transduced with red-shifted firefly luciferase reporter gene (Perkin Elmer, #CLS960003). Briefly, cells from previously engrafted tumors were seeded in 48-well plates and allowed to attach for 24 h, followed by transduction with RED-FLuc-GFP lentiviral particles at a multiplicity of infection (MOI) of 5 in the presence of 0.5% v/v Vectofusion-1 (MiltenyiBiotec, #130-111-163) to enhance transduction. Attached cells were spinoculated at 700×g for 90 min at 32°C and incubated for 16-24 h at 37°C. Cells were detached using accutase (Sigma-Aldrich, #SCR005), centrifuged, resuspended in saline, and mixed with 1:1 Matrigel/RPMI medium prior to propagation in mice. Following tumor engraftment, primary tumors were harvested, dissociated into a single-cell suspension, sorted for GFP-high cells (Sony Biotechnology, SH800S Cell Sorter), and cryopreserved in liquid nitrogen for subsequent use. For orthotopic injection of the patient material into the ovary, mice were anaesthetised and placed in left lateral recumbency. Analgesia was administered via subcutaneous injection of 0.1 mg/kg buprenorphine hydrochloride (Temgesic) (Indivior UK Limited, #561634) and 5 mg/kg meloxicam (Metacam) (Boehringer Ingelheim Animal Health Nordics, #386860). Single-cell suspensions (1 × 10^5^ cells) from two different primary patient tumours were resuspended in 5 µL saline and separately mixed with 5 µL of 1:1 Matrigel/RPMI medium (Corning, #08-774-391). Injection of the 10-µL mixture into the ovarian bursa at the oviduct was performed using a 30 G needle and a 10x microscope (PDX26 n=32). Abdominal muscles and skin were closed separately, using 6.0 absorbable polyglactin sutures (Vicryl) (Ethicon Surgical Technologies, #V492H Vicryl). Animals received subcutaneous saline post-operatively and recovered in a warmed environment before returning to the home cage. Mice were housed in groups of 4-6 in individually ventilated cages on a 12-h dark/night cycle and weighted three times per week.

To monitor disease progression, non-invasive bioluminescence imaging (BLI) was performed biweekly (weeks −1, 1, 3, 5, 7, 9) on all luciferase-transduced PDX models. BLI was performed 10 min after intraperitoneal administration of 150 mg/kg of D-luciferin (Biosynth, #L-8220) using the IVIS Spectrum In Vivo Imaging System (PerkinElmer). Mice were allocated to five treatment groups once a stable bioluminescence signal was detected in the right (injected) ovary in the lateral view.

The control groups received vehicles (10% DMSO, 40% PEG300, 5% Tween-80, 45% saline) orally. A standard-of-care control group received carboplatin monotherapy (12 mg/kg intraperitoneally dissolved in PBS [Fresenius Kabi, Hospital Pharmacy Norway, 10 mg/mL]) in addition to the vehicle. The candidate drugs pevonedistat, xevinapant, and ZEN-3694 were administered in combination with carboplatin following a three-day monotherapy treatment (Fig. 5d). Pevonedistat (MedChemTronica, #HY-70062) was dissolved in 10% DMSO, 40% PEG300, 5% Tween-80, 45% saline and administered intraperitoneally at a dose of 30 mg/kg (Shapiro et al. 2023). Xevinapant (MedChemTronica, #HY-15454,) was dissolved in 10% DMSO, 40% PEG300, 5% Tween-80, 45% saline while ZEN-3694 (Selleckchem, #E1517) was dissolved in 5% DMSO, 40% PEG300, 5% Tween-80, and 50% saline. They were both administered at a dose of 100 mg/kg orally (Cai et al. 2011) A dose-range finding study of all candidate drugs was performed to determine a safe study dose (Supplementary Fig. 2). Treatment with 60 mg/kg pevonedistat induced toxicity in mice, while 40 mg/kg was well tolerated. Carboplatin was given intraperitoneally twice weekly for three weeks (Q2W×3) either as monotherapy or in combination with candidate drugs. Pevonedistat was injected intraperitoneally twice daily, 5 days per week. Xevinapant, ZEN-3694, and the oral vehicle control were administered once daily, five days per week for a total of three weeks and three days. Disease progression and drug response were evaluated biweekly by BLI, palpation, and body weight measurements (Extended Data Fig. 10 e).

Mice were scored based on reduced activity, appearance, tumor burden by optical imaging and palpation, pain assessment, body weight loss (>10%) or body weight gain (tumor burden) and euthanised when reaching a score above 10 or at the end of the study period (15 weeks after treatment initiation.) Post-mortem examination included macroscopic description of primary and metastatic tumors, *ex vivo* BLI of all organs, measurements of primary tumor (size,weight), and collection of tumor samples.

We previously developed a modified staging system based on the FIGO classification for ovarian, fallopian tube, and peritoneal cancer (Berek et al. 2018; Kleinmanns, Fosse, et al. 2020) assigning disease stages I–IV to each mouse based on pre-surgical whole-body BLI and intraoperative assessment. Since animals in this study did not undergo surgery, we further adapted the system: stage I was defined to disease to the right ovary only, stage II involved spread to the lower abdomen (e.g., mesentery, contralateral side), stage III indicated metastatic spread to the omentum and/or diaphragm (peritoneum), and stage IV signified disease involving the spleen and/or liver. The *ex vivo* bioluminescence signal of all organs was quantified using regions of interest (ROI) using the IVIS Spectrum In Vivo Imaging System (PerkinElmer). Data were analysed using the Mann Whitney U test and are shown as mean +/– standard deviation (SD). Values of p<0.05 were considered statistically significant. All statistical analyses were performed in GraphPad Prism v10.

#### Variant overlap analysis between patient tumors and PDX models

SNP overlap between patient-derived HGSC xenografts and the corresponding primary tumors was assessed using bcftools (v1.13). VCF files were restricted to SNPs, and both total SNP counts and unique variant positions were quantified to account for multi-allelic sites. Reference and alternate allele distributions were summarized across samples. Shared and sample-specific SNPs between matched tumor-PDX pairs were identified using bcftools isec. The percentage of shared SNPs was calculated row-wise as the proportion of overlapping SNP positions relative to the total number of SNPs detected per sample and used for visualization.

## Data and code availability

Raw human scRNA-seq data are available through the EGA under accession EGAS00001005010. Processed scRNA-seq datasets for the intrinsic and dual approaches have been deposited in GEO (GSE307231* and GSE318568*). Processed PDX scRNA-seq data will be released upon publication (GSEXXXXX*). *In vivo* bioluminescence datasets are accessible at synapse.org (syn71720400). All software and custom code used for bioinformatic analyses are available on Zenodo (https://doi.org/10.5281/zenodo.17911214).

\**Data will be fully accessible upon publication*.

## Acknowledgments

We are deeply grateful to the patients who participated in this study and to patient representative Jeanette Hoel (Gynekreftforeningen, Norway) for her essential contributions. We also thank Merja Heinäniemi and Alexandra Leary for insightful discussions. We thank CSC–IT Center for Science, Biocenter Finland, and the BRIC High Content CRISPR Screens core facility (Novo Nordisk Foundation NNF20OC0061734) for computational and technical support.

## Author Contributions

EMC, KW, LB, AV, and BS conceived and designed the study, and, together with SH and JH, acquired funding. LGM, KK, AP, MS, together with KW, LB, AV, and BS wrote the inital draft, together with EMC, managed the project, and, with additional input from CD, developed methodologies. LGM, KK, AP, MS, and DFA created the visualizations, and, with GS and JH, curated data. LGM, KK, AP, MS, GS, MMF, DFA, CD and BS performed formal analysis. LGM, KK, AP, MS, GS, JD, CD, SH, and JH conducted research and investigation. EMC, KW, LB, AV, and BS provided resources, together with JH and SH. They supervised research together with CD. AP and MS developed software. GM, KK, MS, GS, KW and LB performed validation. All authors reviewed, edited and approved the final manuscript.

## Additional information

**Extended data** is available for this paper below.

**Supplementary information** The online version contains supplementary material.

---

**Extended Data Fig. 1.**
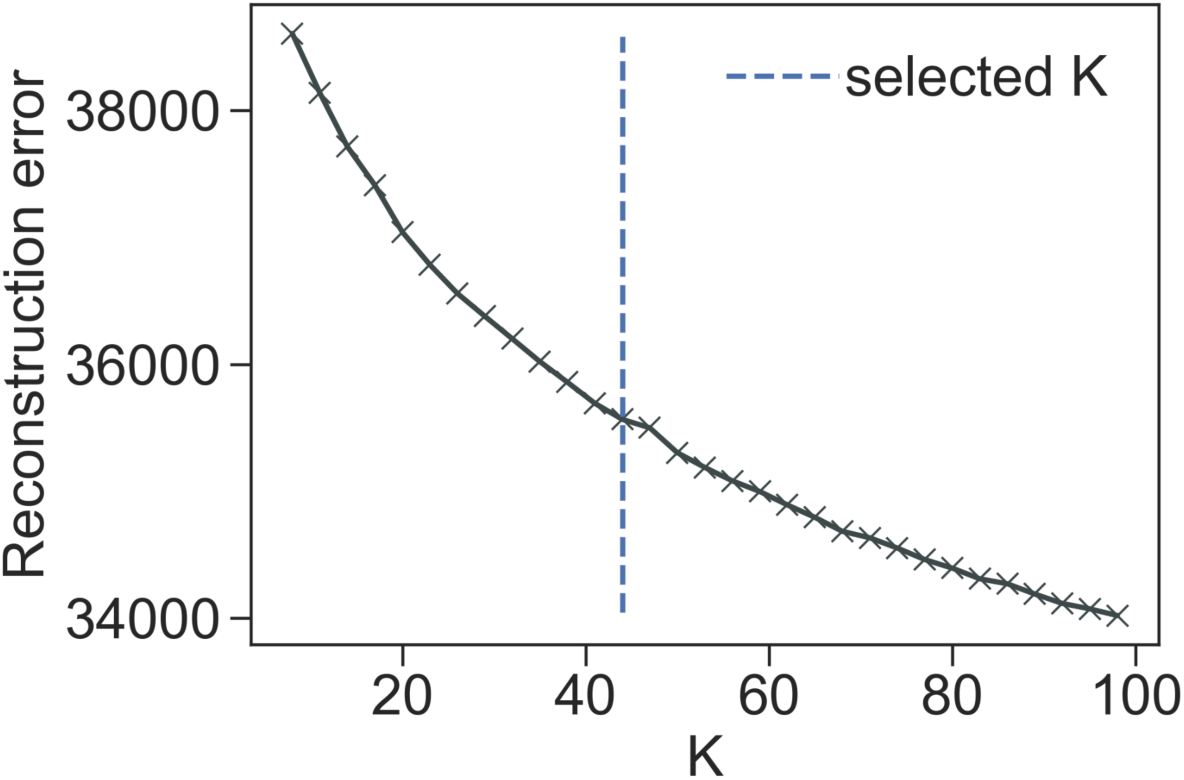
Selecting the number of signatures (*K*) for the dual approach. NMF reconstruction error across increasing values of *K*. The selected *K* corresponds to the elbow of the curve.

**Extended Data Fig. 2.**
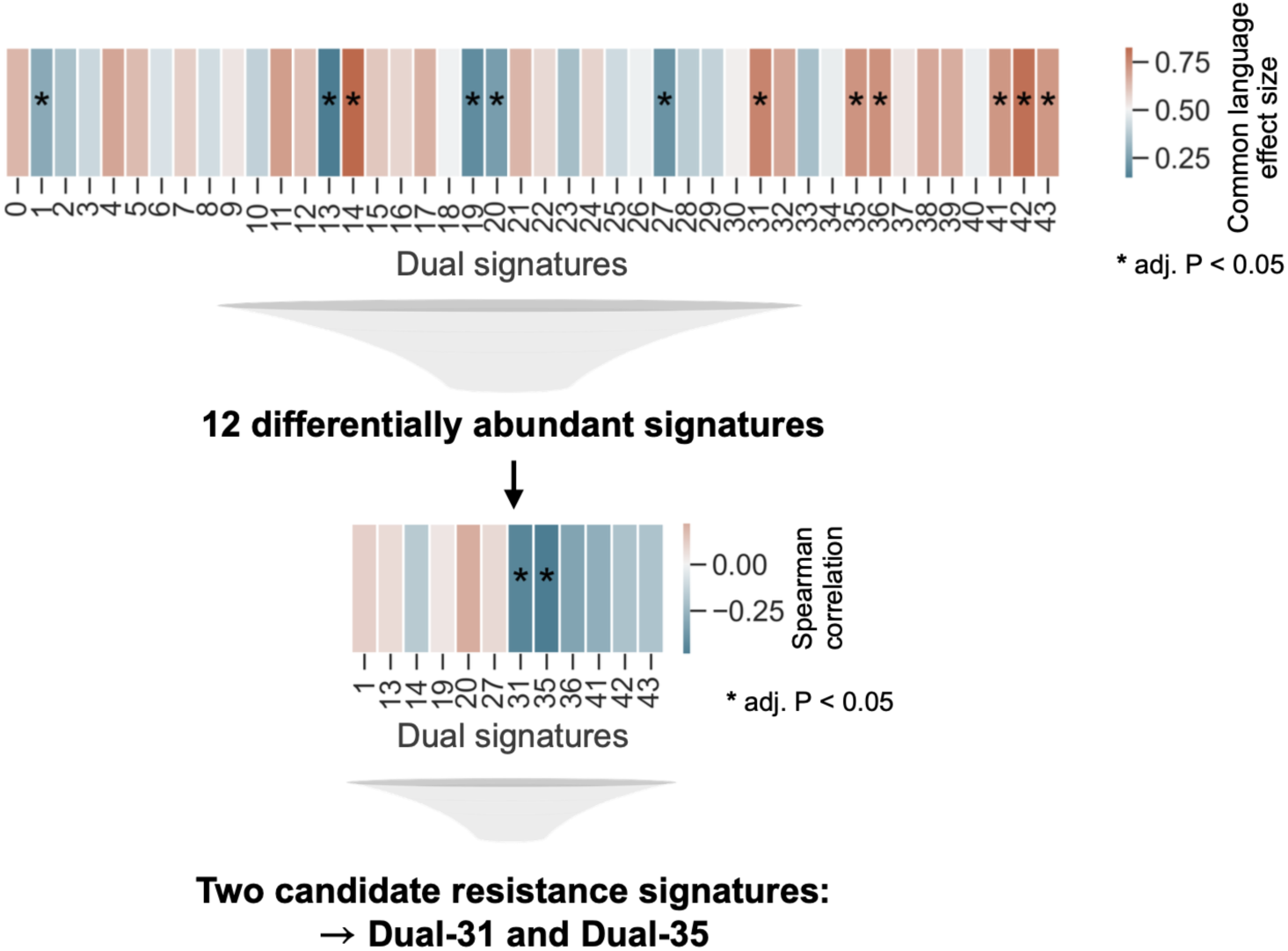
Statistical framework for the identification of candidate resistance-associated signatures for the dual approach. Common language effect sizes from Mann-Whitney U tests comparing signature expression scores between treatment-naïve (N = 37) and post-NACT tumors (N = 17) are shown. Signatures significant in this first step (12 total) were further tested for association with PFI using Spearman correlation. Two candidate resistance signatures remained significant after both filtering steps (adjusted P < 0.05).

**Extended Data Fig. 3.**
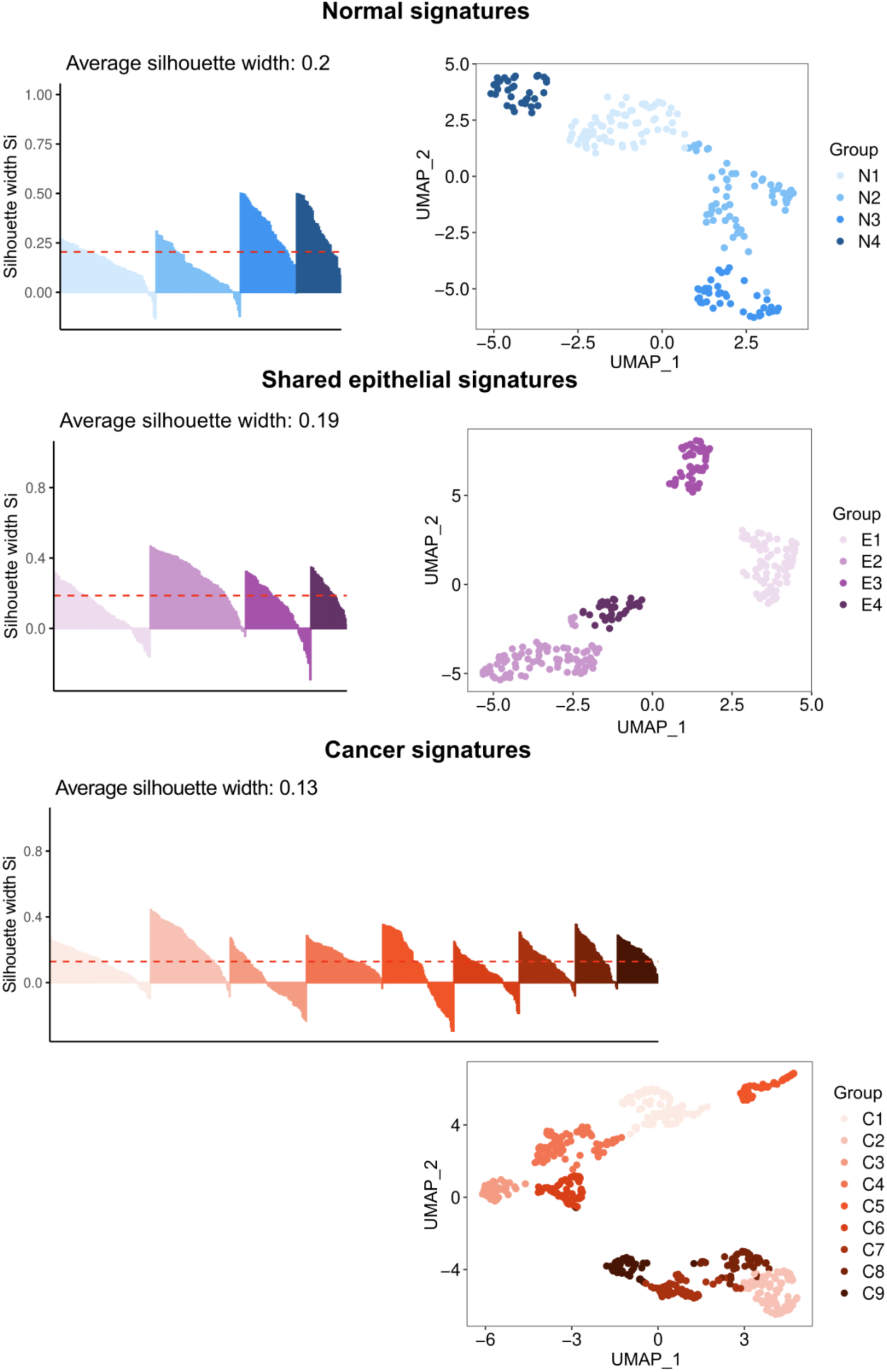
Silhouette-based refinement and UMAP visualization of intrinsic signatures. Genes differentially expressed between cancer and normal epithelial cells were classified as normal-enriched, cancer-enriched, or shared. Clusters were refined using silhouette width analysis, excluding genes with negative values. The silhouette plot is shown on the left, and the final gene clusters are visualized by UMAP on the right.

**Extended Data Fig. 4.**
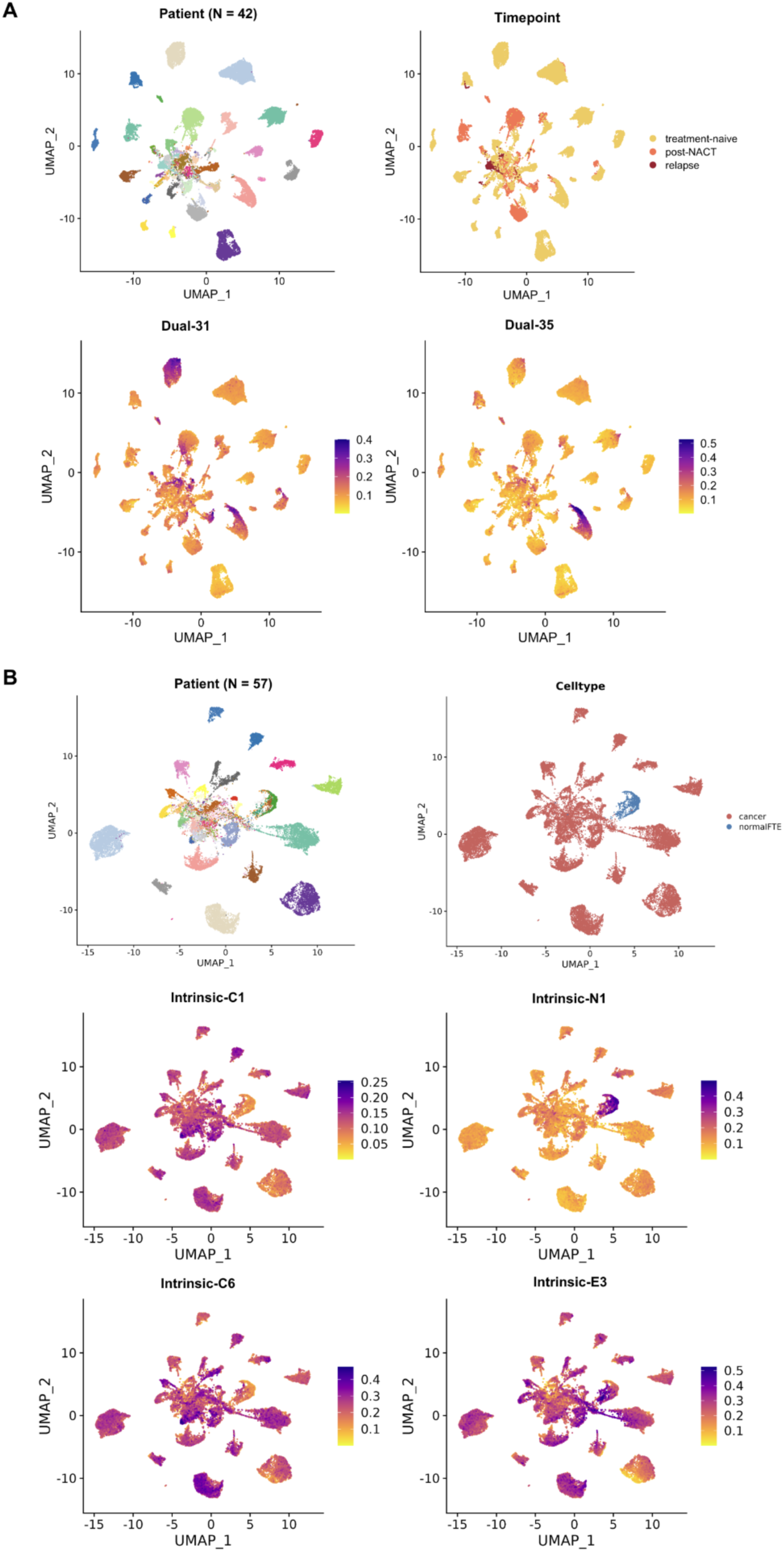
UMAP visualization of datasets analyzed using dual and intrinsic approaches. **a,** UMAPs of cells included in the dual approach, colored by patient, treatment phase, and signature activity calculated using a weighted AUCell method. **b,** UMAPs of cells included in the intrinsic approach, colored by patient, cell type, and signature activity calculated using UCell.

**Extended Data Fig. 5.**
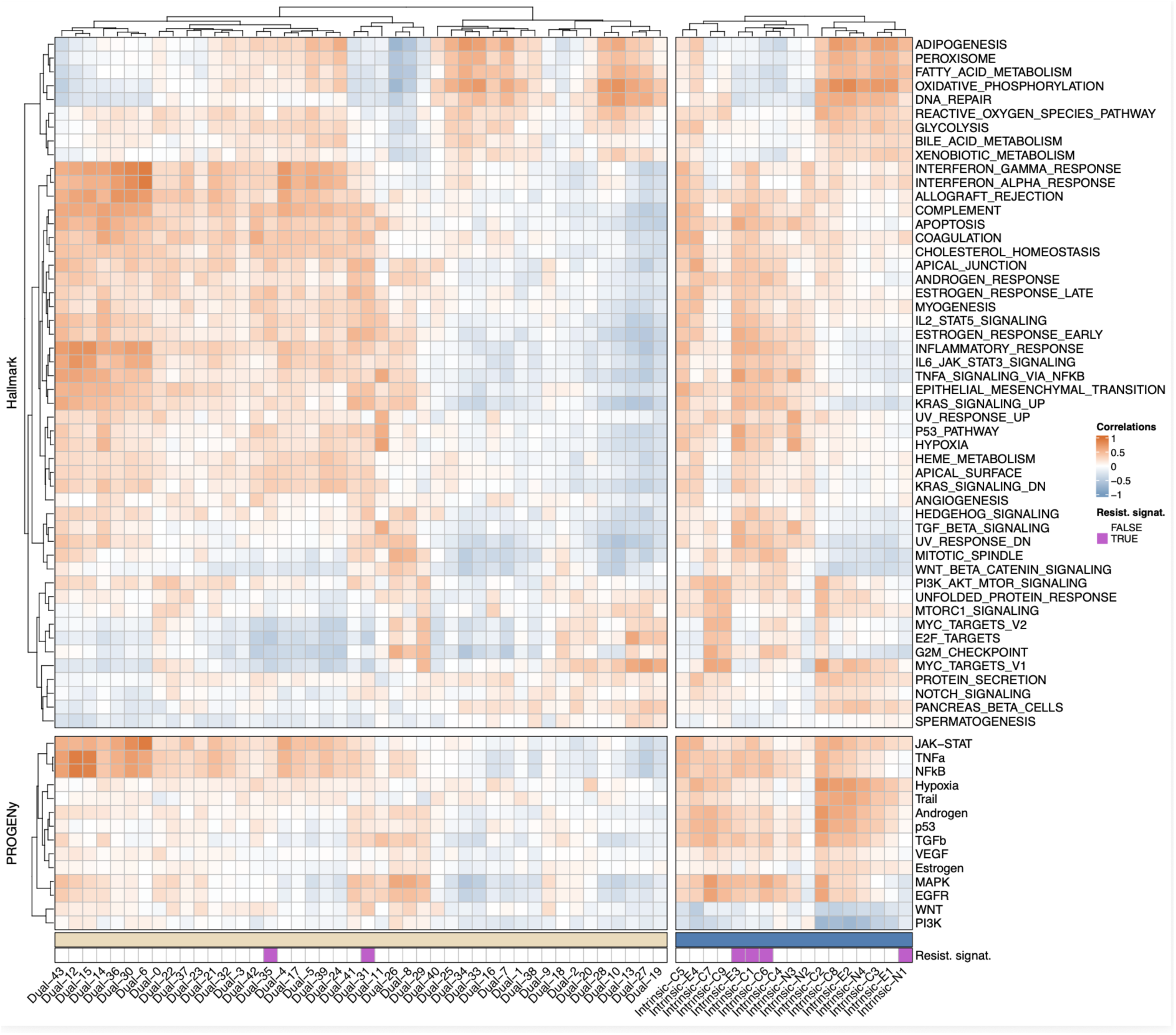
Correlation of signature expression with Hallmark and PROGENy pathway activity in clinical specimens. Correlation between the expressions of all dual and intrinsic signatures (horizontal) with Cancer Hallmark pathway activities (top) and inferred PROGENy pathway activities (bottom), calculated at the single-cell level.

**Extended Data Fig. 6.**
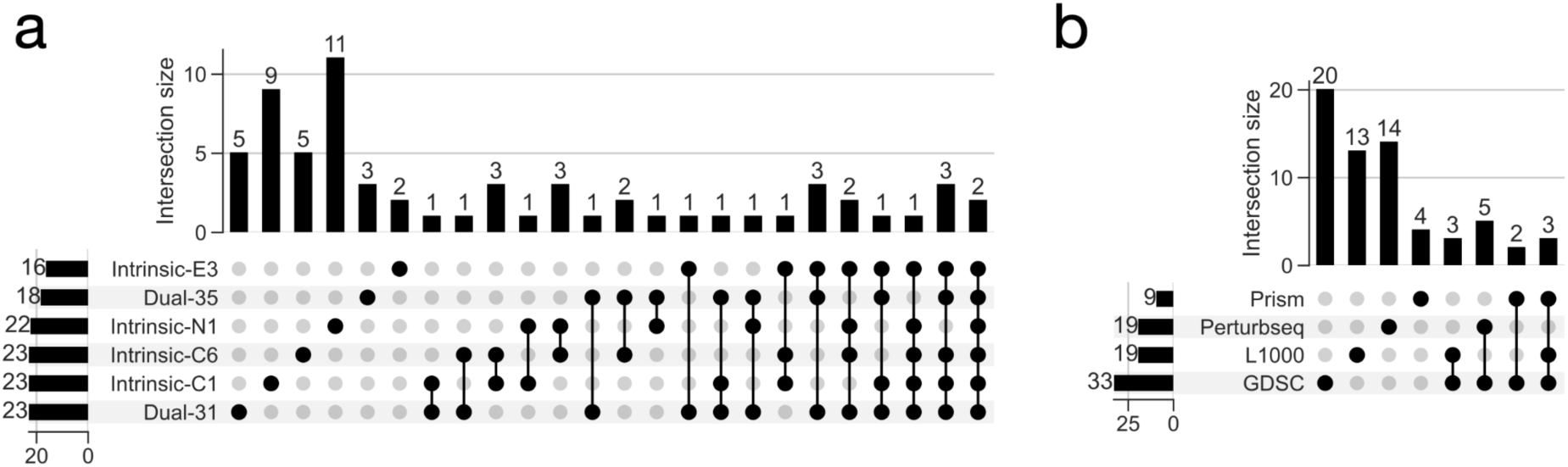
Overlap among drug predictions. UpSet plots showing the number of individually and jointly predicted drugs, among **a,** resistance-associated signatures and **b,** databases.

**Extended Data Fig. 7.**
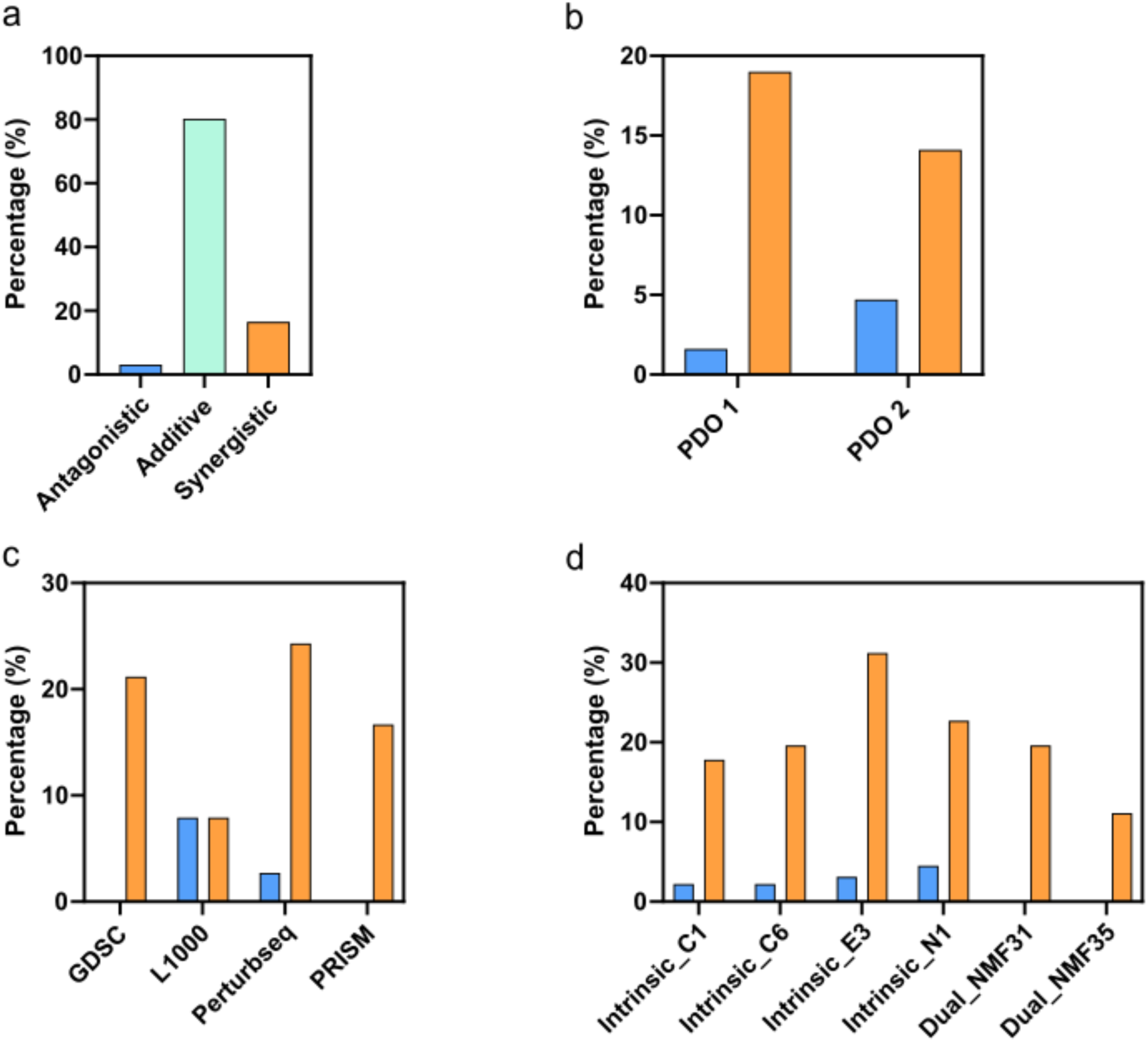
Drug screening results per PDO, database, and signature. **a,** Percentage of antagonistic, additive, and synergistic drug combinations with carboplatin in the drug screens with the candidate drugs (n=64) in PDO 1 and 2. The plot contains the results from PDO 1 and 2, after summarizing the drug interaction type (n=128). Synergy in any of the concentrations was summarized as synergy. Similarly, additivity was prioritized over antagonism. Paclitaxel results were excluded in the analysis, as it was included as a reference combination. **b,** Percentage of synergistic and antagonistic drug combinations in each PDO model. **c,** Percentage of synergistic and antagonistic drug combinations per database. **d,** Percentage of synergistic and antagonistic drugs across signatures.

**Extended Data Fig. 8.**
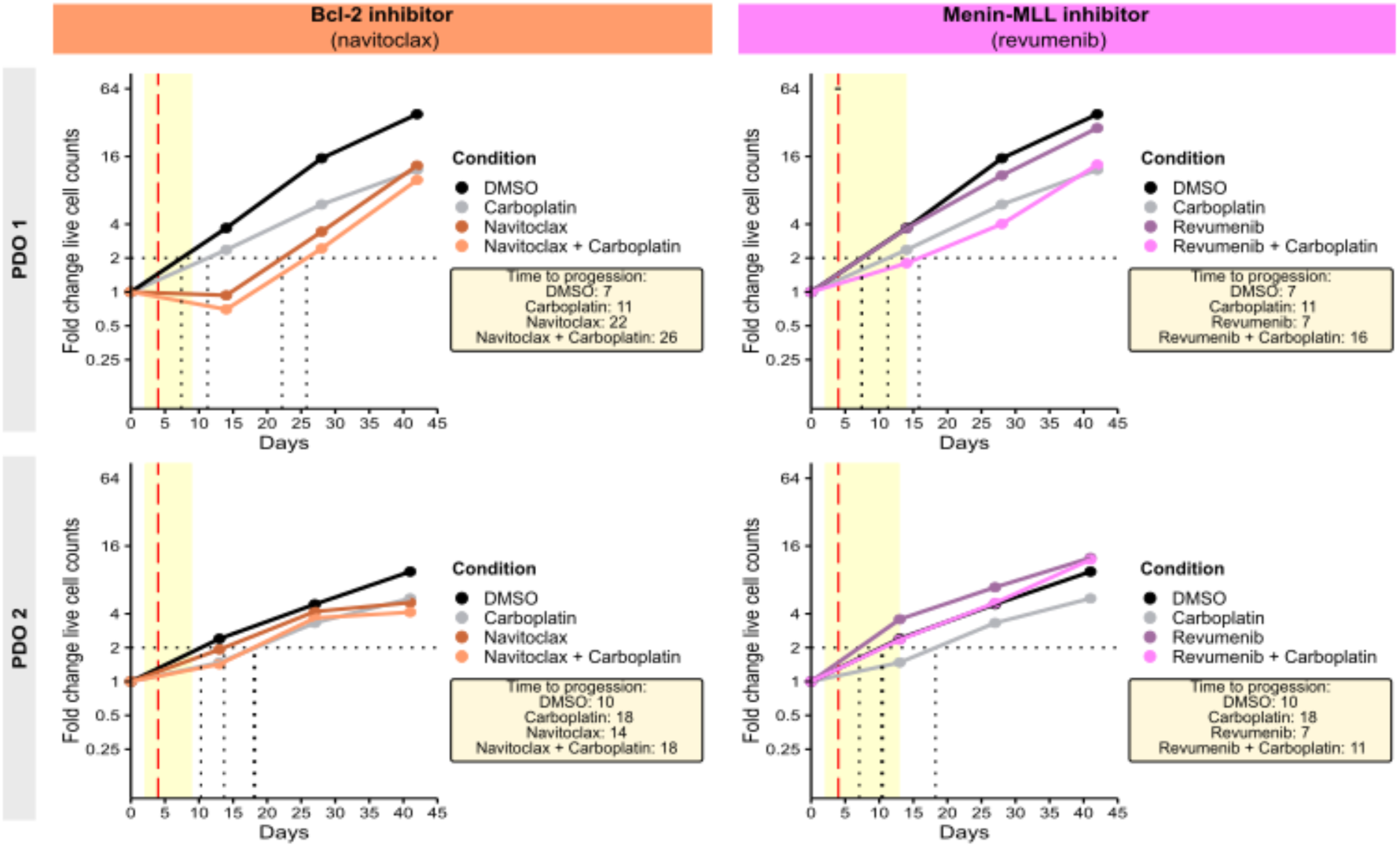
Additional long-term survival in PDO 1 and PDO 2. Long-term survival in PDO 1 and 2. The red dashed line indicates carboplatin treatment and the yellow shaded area indicates treatment duration. Time-to-progression corresponds to the number of days required for the PDOs to double the initial number of viable cells seeded.

**Extended Data Fig. 9.**
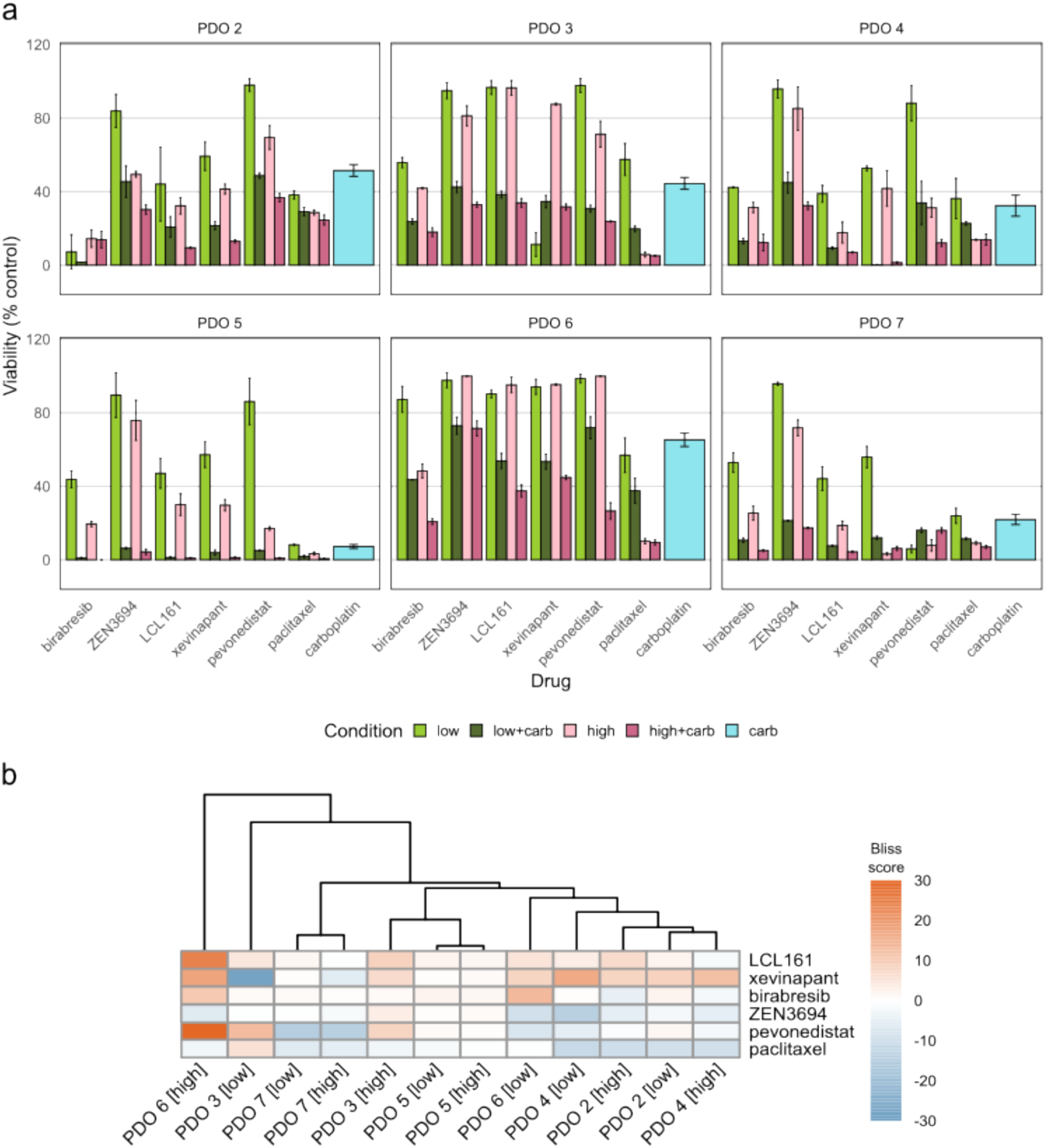
Validation of replacement drugs in additional PDOs. **a,** Viability of the candidate drugs used in the PDO validations (birabresib, LCL161, and pevonedistat) and their replacements (ZEN-3694 for birabresib, xevinapant for LCL161) for PDX studies. Paclitaxel was included as a control. Carboplatin concentrations for PDOs 2, 3, 4, 5, and 7 was 10 µM whereas for PDO 6 was 20 µM. Bortezomib (positive control) was used at a final concentration of 10 µM and DMSO (negative control) 0.1% v/v. Each drug treatment was conducted in technical triplicates. Viability was indirectly assessed by CellTiter-Glo 3D. Experimental conditions and analysis were the same as in the drug screening with the candidate drugs (Fig. 4b). **b,** Bliss scores. Columns are ordered by hierarchical clustering using Euclidean distance.

**Extended Data Fig. 10.**
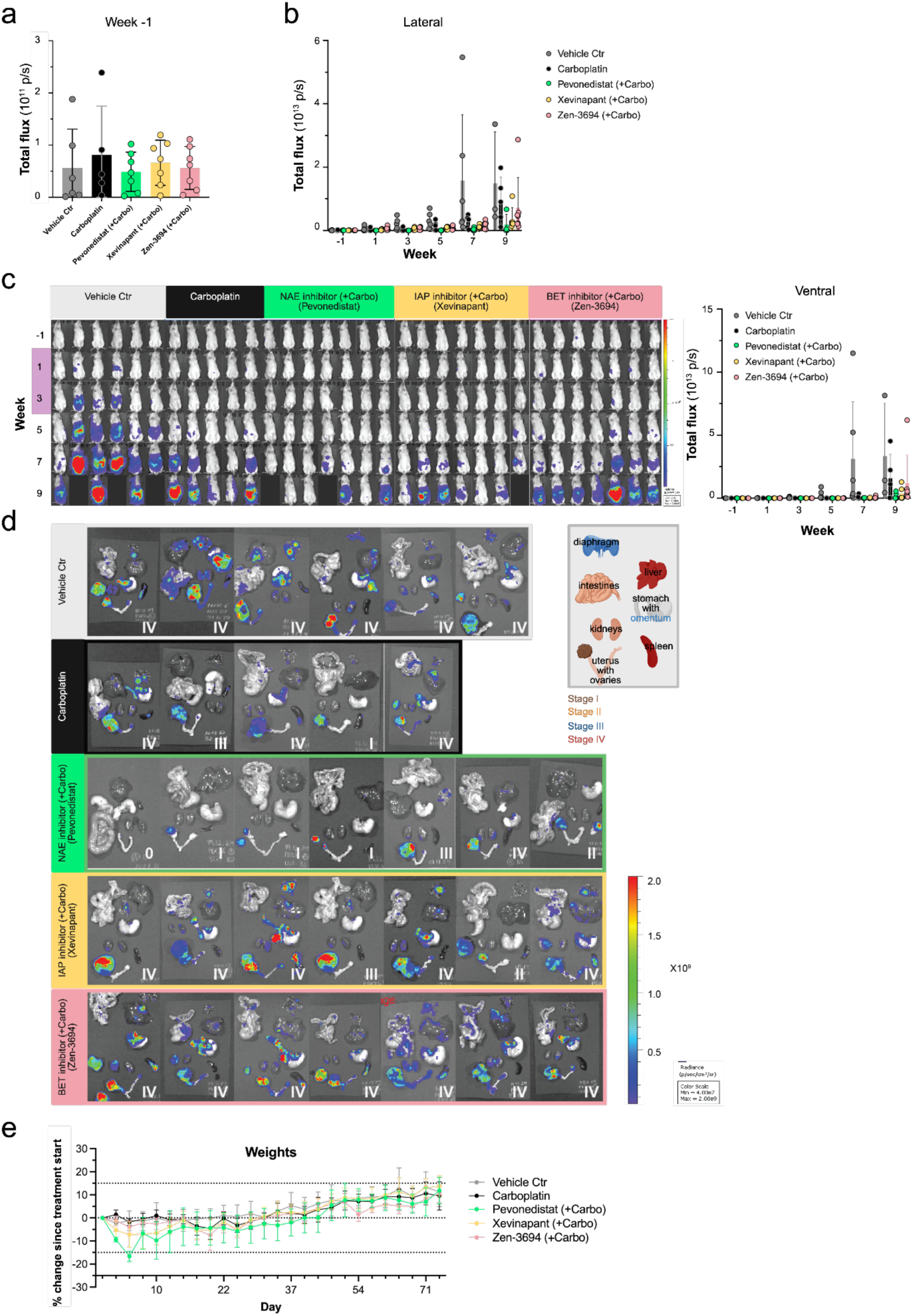
Validation of drug combinations in HGSC PDX models. **a,** Total photon flux (lateral view) at group assignment at week −1 and **b,** Total photon flux (lateral view) during and after treatment. **c,** Ventral bioluminescence images from all study mice (each column represents an individual mouse) and longitudinally plotted ventral photon flux for each treatment group from week –1 to week 9 visualizing primary and metastatic tumor growth. **d,** *Ex vivo* bioluminescence intensity of organs harvested at the study endpoint from mice bearing PDX26 tumors, stratified by treatment group. A modified staging system (Stage I-IV) was applied based on *ex vivo* imaging intensity across all organs. **e,** Relative change in body weight related to pretreatment weight (day 0) shown for all treatment days (day 3—day 22) and during the survival period (day 24—74).

