## Supplementary Figures and Tables for "Single-Cell Transcriptomic Signatures Enable Stratified Combination Therapy for Platinum-Resistant Ovarian Cancer"

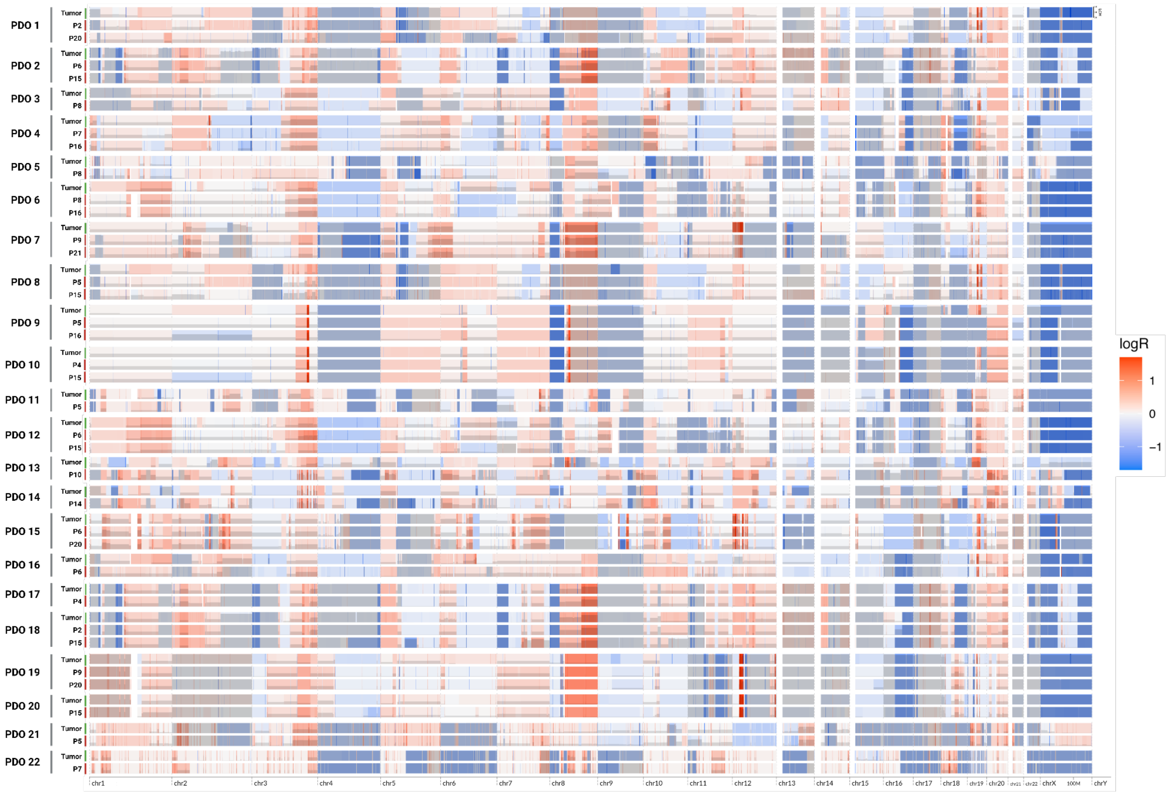


**Supplementary Fig. 1. Validation of the long-term HGSC PDO models.** Copy number variation (CNV) analysis from whole genome sequencing (WGS) of all PDOs used in the experimental procedures (n=22), with their matching tumors of origin. PDOs were profiled at early and late passages, showing retention of the genetic features of their tumor of origin, with copy number profiles matching the corresponding patient samples even after extended culture. Passage numbers (P) are indicated for PDO cultures. Copy number changes are expressed as logR, with amplifications shown in red and deletions in blue. The loss of heterozygosity (LOH) score of each segment is displayed with gray bars. Data was visualized using the GenomeSpy toolkit (https://genomespy.app/).

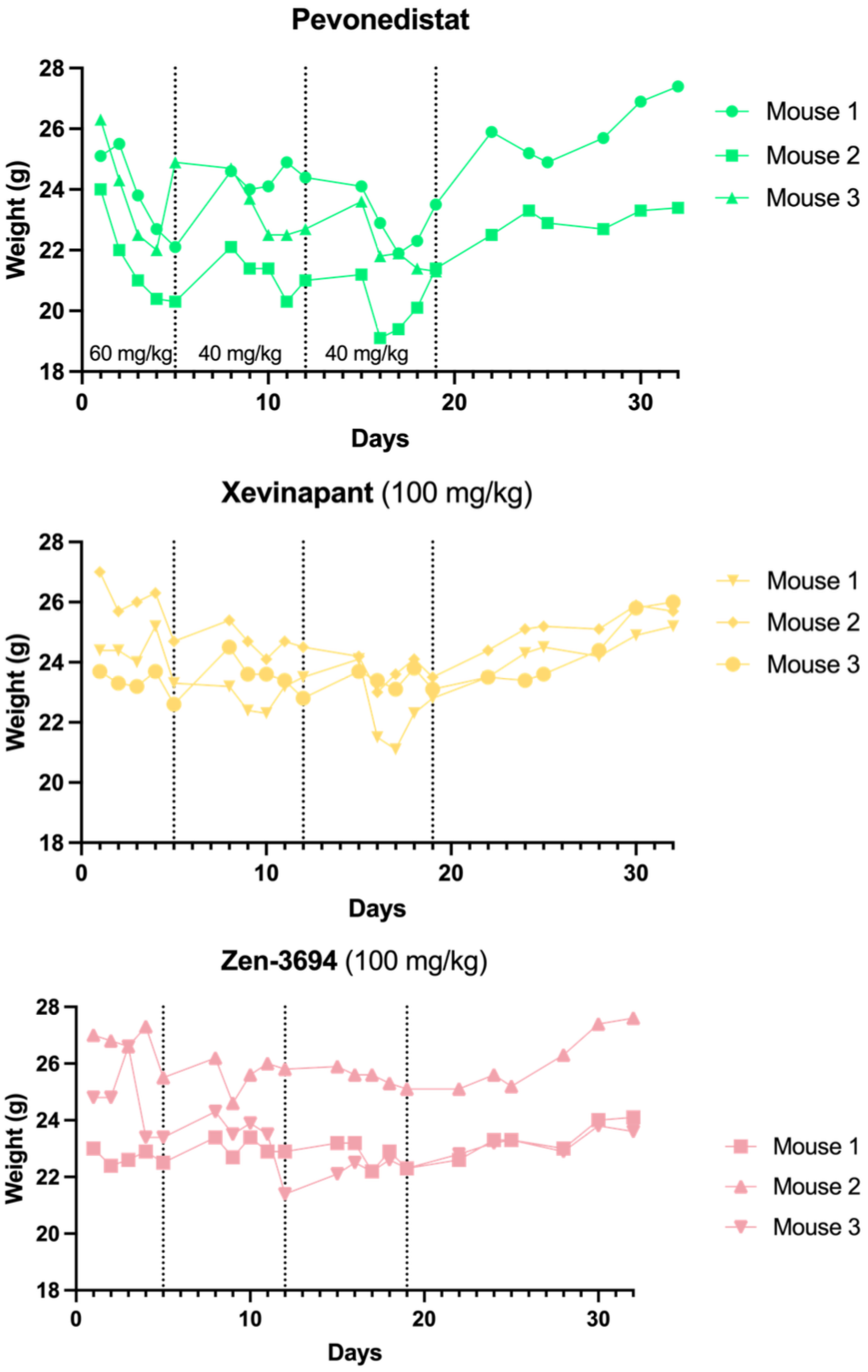


**Supplementary Fig. 2. Toxicity assessment of three candidate drugs (Pevonedistat, Xevinapant and Zen-3694) in non-tumor bearing NSG mice.** Body weights of all mice during three treatment cycles, each consisting of five consecutive days of drug administration followed by two days off. Cycles are separated by vertical lines. The dose of the candidate drug pevonedistat was reduced from 60 mg/kg to 40 mg/kg after the first treatment cycle due to weight loss. Pevonedistat was administered intraperitoneally, whereas xevinapant and Zen-3694 were administered orally.

Suppl. Table 1

| PatientID | Sample | Treatment_phase | Tissue_site | Stage | Age_at_Diagnosis | PFI (thresholded due to privacy concerns) | Published |
|---|---|---|---|---|---|---|---|
| Patient1 | sample1 | primary | Tubules | IVA | 57 | 6-12months | H√§kkinen_et_al_Bioinformatics |
| Patient2 | sample2 | primary | Ascites | IIIC | 72 | | H√§kkinen_et_al_Bioinformatics |
| Patient3 | sample3 | primary | Omentum | IVB | 81 | >12months | H√§kkinen_et_al_Bioinformatics |
| Patient4 | sample4 | primary | Omentum | IVB | 77 | >12months | Launonen_et_al_CancerCell |
| Patient5 | sample5; sample53 | primary; interval | Peritoneum; Ovary | IIIC | 60 | <6months | Launonen_et_al_CancerCell |
| Patient6 | sample10 | primary | Omentum | IIIC | 78 | 6-12months | Launonen_et_al_CancerCell |
| Patient7 | sample6 | primary | Omentum | IIIC | 74 | >12months | Launonen_et_al_CancerCell |
| Patient8 | sample7 | primary | Peritoneum | IVB | 68 | <6months | Launonen_et_al_CancerCell |
| Patient9 | sample8; sample55 | primary; interval | Omentum; Tumor | IIIC | 75 | 6-12months | Hippen_et_al_PlosCompBiol; no |
| Patient10 | sample9 | primary | Ovary | IIIC | 75 | >12months | Hippen_et_al_PlosCompBiol |
| Patient11 | sample11 | primary | Ascites | IVA | 67 | >12months | H√§kkinen_et_al_Bioinformatics |
| Patient12 | sample12 | primary | Mesenterium | IIIC | 77 | >12months | Launonen_et_al_CancerCell |
| Patient13 | sample13 | primary | Peritoneum | IIIC | 74 | 6-12months | H√§kkinen_et_al_Bioinformatics |
| Patient14 | sample14 | primary | Mesenterium | IIIC | 70 | <6months | Pirttikoski_et_al |
| Patient15 | sample15 | primary | Ovary | IIIC | 78 | >12months | Launonen_et_al_CancerCell |
| Patient16 | sample16; sample56 | primary; interval | Peritoneum; Peritoneum | IIIC | 68 | >12months | Zhang_et_al_SciAdv |
| Patient17 | sample17 | primary | Ovary | IIIC | 69 | >12months | Pirttikoski_et_al |

Suppl. Table 1

| PatientID | Sample | Treatment_phase | Tissue_site | Stage | Age_at_Diagnosis | PFI (thresholded due to privacy concerns) | Published |
|---|---|---|---|---|---|---|---|
| Patient18 | sample18; sample57 | primary; interval | Omentum; Ovary | IIIC | 81 | >12months | Launonen_et_al_CancerCell |
| Patient19 | sample19; sample58 | primary; interval | Omentum; Omentum | IVA | 54 | <6months | Zhang_et_al_SciAdv |
| Patient20 | sample20 | primary | Tumor | IIIC | 63 | <6months | Hippen_et_al_PlosCompBiol |
| Patient21 | sample21 | primary | Tumor | IIIC | 62 | <6months | Hippen_et_al_PlosCompBiol |
| Patient22 | sample22; sample59 | primary; interval | Omentum; Omentum | IIIC | 62 | <6months | Zhang_et_al_SciAdv |
| Patient23 | sample23; sample24 | primary; interval | Adnexa, Ascites | IIIC | 74 | <6months | Senkowski_et_al_DevCell; Senkowski_et_al_DevCell |
| Patient24 | sample25; sample60 | primary; interval | Peritoneum; Omentum | IVA | 67 | <6months | Zhang_et_al_SciAdv |
| Patient25 | sample26; sample61 | primary; interval | Peritoneum; Omentum | IIIC | 62 | <6months | Zhang_et_al_SciAdv |
| Patient26 | sample27; sample62 | primary; interval | Mesentery; Omentum | IVA | 64 | >12months | Zhang_et_al_SciAdv |
| Patient27 | sample28 | primary | Peritoneum | IIIC | 68 | <6months | Pirttikoski_et_al (part of Micoli_et_al_CancerDiscov) |
| Patient28 | sample29; sample64 | primary; relapse | Peritoneum; Ascites | IVB | 72 | <6months | Launonen_et_al_CancerCell; no |
| Patient30 | sample30; sample65 | primary; interval | Peritoneum; Omentum | IVA | 72 | <6months | Zhang_et_al_SciAdv |
| Patient31 | sample31; sample66 | primary; interval | Peritoneum; Tumor | IVA | 73 | <6months | Zhang_et_al_SciAdv |

Suppl. Table 1

| PatientID | Sample | Treatment_phase | Tissue_site | Stage | Age_at_Diagnosis | PFI (thresholded due to privacy concerns) | Published |
|---|---|---|---|---|---|---|---|
| Patient32 | sample32; sample67 | primary; interval | Omentum; Omentum | IVA | 78 | >12months | Zhang_et_al_SciAdv |
| Patient33 | sample33 | primary | Omentum | IIIC | 79 | | Senkowski_et_al_DevCell |
| Patient34 | sample34 | primary | Peritoneum | IIIC | 72 | <6months | Launonen_et_al_CancerCell |
| Patient35 | sample35; sample68 | primary; interval | Peritoneum; Omentum | IVB | 67 | <6months | Zhang_et_al_SciAdv |
| Patient36 | sample36; sample69 | primary; interval | Omentum; Omentum | IIIC | 71 | <6months | Launonen_et_al_CancerCell |
| Patient37 | sample37 | primary | Omentum | IIIC | 75 | 6-12months | Launonen_et_al_CancerCell |
| Patient38 | sample38 | primary | Peritoneum | IIIC | 85 | <6months | Pirttikoski_et_al |
| Patient39 | sample39; sample70 | primary; interval | Omentum; Omentum | IVA | 74 | 6-12months | Zhang_et_al_SciAdv |
| Patient40 | sample40; sample41 | primary; primary | Omentum | IIIC | 68 | 6-12months | Pirttikoski_et_al (part of Micoli_et_al_CancerDiscov) |
| Patient41 | sample42 | primary | Omentum | IIIC | 77 | | Launonen_et_al_CancerCell |
| Patient42 | sample43 | primary | Ovary | IIIC | 73 | <6months | Pirttikoski_et_al (part of Micoli_et_al_CancerDiscov) |
| Patient43 | sample44 | primary | Omentum | IVB | 55 | | Pirttikoski_et_al |
| Patient44 | sample45 | primary | Peritoneum | IIIC | 78 | | Launonen_et_al_CancerCell |
| Patient45 | sample46 | primary | Omentum | IIIC | 73 | | Pirttikoski_et_al (part of Micoli_et_al_CancerDiscov) |
| Patient46 | sample47 | primary | Ascites | IIIC | 73 | | Pirttikoski_et_al |
| Patient47 | sample48 | primary | Ascites | IIIB | 67 | | Pirttikoski_et_al |
| Patient48 | sample49 | primary | Ascites | IIIC | 68 | | Launonen_et_al_CancerCell |

Suppl. Table 1

| PatientID | Sample | Treatment_phase | Tissue_site | Stage | Age_at_Diagnosis | PFI (thresholded due to privacy concerns) | Published |
|---|---|---|---|---|---|---|---|
| Patient49 | sample50 | primary | Omentum | IIIC | 70 | | Pirttikoski_et_al (part of Micoli_et_al_CancerDiscov) |
| Patient50 | sample51 | primary | Omentum | IIIC | 75 | | Pirttikoski_et_al |
| Patient51 | sample52 | primary | Omentum | IVA | 68 | | Pirttikoski_et_al |
| Patient52 | control1 | control | Fimbria | NA | NA | | Pirttikoski_et_al |
| Patient53 | control2 | control | Fimbria | NA | NA | | Perki√∂_et_al_SciRep |
| Patient54 | control3 | control | Tube | NA | NA | | Perki√∂_et_al_SciRep |
| Patient55 | control4 | control | Fimbria | NA | NA | | Perki√∂_et_al_SciRep |
| Patient56 | control5 | control | Fimbria | NA | NA | | Perki√∂_et_al_SciRep |
| Patient57 | control6 | control | Tube | NA | NA | | Perki√∂_et_al_SciRep |
| Patient58 | control7 | control | Fimbria | NA | NA | | Pirttikoski_et_al |
| Patient59 | sample54 | interval | Omentum | IIIC | 67 | 6-12months | H√§kkinen_et_al_Bioinformatics |
| Patient60 | sample63 | interval | Ovary | IIIC | 67 | 6-12months | no |
| Patient61 | sample71 | relapse | Mesentery | IVB | 68 | >12months | no |
| Patient62 | sample72 | relapse | Tumor | IIIB | 58 | >12months | Micoli_et_al_CancerDiscov |

Suppl. Table 2

| SampleID | Cancer_intrinsic | NormalFTE_intrinsic | Immune_intrinsic |
|---|---|---|---|
| sample1 | 211 | NA | 189 |
| sample2 | 9 | NA | 691 |
| sample3 | 198 | NA | 127 |
| sample4 | 50 | NA | 433 |
| sample5 | 52 | NA | 195 |
| sample6 | 224 | NA | 721 |
| sample7 | 51 | NA | 185 |
| sample8 | 7 | NA | 85 |
| sample9 | 245 | NA | 79 |
| sample10 | 65 | NA | 282 |
| sample11 | 673 | NA | 2 |
| sample12 | 2631 | NA | 63 |
| sample13 | 157 | NA | 1231 |
| sample14 | 2686 | NA | 175 |
| sample15 | 877 | NA | 1422 |
| sample16 | 777 | NA | 479 |
| sample17 | 831 | NA | 943 |
| sample18 | 3533 | NA | 610 |
| sample19 | 1701 | NA | 710 |
| sample20 | 46 | NA | 835 |
| sample21 | 131 | NA | 562 |
| sample22 | 186 | NA | 1717 |
| sample23 | 775 | NA | 113 |
| sample24 | 511 | NA | 3085 |
| sample25 | 178 | NA | 3238 |
| sample26 | 200 | NA | 1194 |
| sample27 | 200 | NA | 1650 |
| sample28 | 499 | NA | 1023 |
| sample29 | 816 | NA | 1601 |
| sample30 | 3545 | NA | 856 |
| sample31 | 591 | NA | 3152 |
| sample32 | 400 | NA | 856 |
| sample33 | 509 | NA | 4949 |
| sample34 | 66 | NA | 1800 |
| sample35 | 49 | NA | 1712 |
| sample36 | 7 | NA | 5144 |
| sample37 | 548 | NA | 669 |
| sample38 | 425 | NA | 1727 |
| sample39 | 49 | NA | 2277 |
| sample40 | 540 | NA | 536 |
| sample41 | 228 | NA | 2319 |
| sample42 | 223 | NA | 2286 |
| sample43 | 947 | NA | 2437 |
| sample44 | 156 | NA | 74 |

Suppl. Table 2

| SampleID | Cancer_intrinsic | NormalFTE_intrinsic | Immune_intrinsic |
|---|---|---|---|
| sample45 | 78 | NA | 2278 |
| sample46 | 116 | NA | 3246 |
| sample47 | 1303 | NA | 281 |
| sample48 | 3 | NA | 2374 |
| sample49 | 22 | NA | 2885 |
| sample50 | 408 | NA | 1195 |
| sample51 | 186 | NA | 393 |
| sample52 | 1317 | NA | 792 |
| sample53 | NA | NA | NA |
| sample54 | NA | NA | NA |
| sample55 | NA | NA | NA |
| sample56 | NA | NA | NA |
| sample57 | NA | NA | NA |
| sample58 | NA | NA | NA |
| sample59 | NA | NA | NA |
| sample60 | NA | NA | NA |
| sample61 | NA | NA | NA |
| sample62 | NA | NA | NA |
| sample63 | NA | NA | NA |
| sample64 | NA | NA | NA |
| sample65 | NA | NA | NA |
| sample66 | NA | NA | NA |
| sample67 | NA | NA | NA |
| sample68 | NA | NA | NA |
| sample69 | NA | NA | NA |
| sample70 | NA | NA | NA |
| sample71 | NA | NA | NA |
| sample72 | NA | NA | NA |
| control1 | NA | 252 | 2270 |
| control2 | NA | 143 | 1879 |
| control3 | NA | 75 | 1790 |
| control4 | NA | 575 | 2433 |
| control5 | NA | 41 | 889 |
| control6 | NA | 158 | 2624 |
| control7 | NA | 7 | 1001 |

Suppl. Table 2

| Stroma_intrinsic | Cancer_dual |
|---|---|
| 240 | 382 |
| 0 | NA |
| 110 | 215 |
| 51 | 65 |
| 9 | 54 |
| 32 | 237 |
| 42 | 51 |
| 348 | NA |
| 149 | 239 |
| 128 | 302 |
| 0 | 649 |
| 2 | 2483 |
| 1003 | 247 |
| 48 | 2525 |
| 127 | 936 |
| 224 | 840 |
| 865 | 1036 |
| 324 | 2290 |
| 521 | 1929 |
| 413 | 44 |
| 2079 | 193 |
| 322 | 295 |
| 2288 | 623 |
| 4 | NA |
| 406 | 257 |
| 361 | 224 |
| 1207 | 355 |
| 233 | 368 |
| 94 | 994 |
| 67 | 3323 |
| 416 | 662 |
| 583 | 509 |
| 447 | NA |
| 331 | 126 |
| 640 | 92 |
| 206 | NA |
| 265 | 614 |
| 70 | 422 |
| 41 | 79 |
| 41 | 411 |
| 162 | 370 |
| 140 | NA |
| 26 | 933 |
| 41 | NA |

Suppl. Table 2

| Stroma_intrinsic | Cancer_dual |
|---|---|
| 8 | NA |
| 51 | NA |
| 1 | NA |
| 162 | NA |
| 6 | NA |
| 32 | NA |
| 30 | NA |
| 79 | NA |
| NA | 259 |
| NA | 639 |
| NA | 306 |
| NA | 278 |
| NA | 1220 |
| NA | 274 |
| NA | 227 |
| NA | 508 |
| NA | 529 |
| NA | 2867 |
| NA | 1693 |
| NA | 31 |
| NA | 466 |
| NA | 556 |
| NA | 132 |
| NA | 89 |
| NA | 35 |
| NA | 207 |
| NA | 475 |
| NA | 64 |
| 1128 | NA |
| 186 | NA |
| 634 | NA |
| 54 | NA |
| 1 | NA |
| 7 | NA |
| 1489 | NA |

Suppl. Table 3

| Dual-0 | Dual-1 | Dual-2 | Dual-3 | Dual-4 | Dual-5 | Dual-6 | Dual-7 | Dual-8 | Dual-9 | Dual-10 | Dual-11 |
|---|---|---|---|---|---|---|---|---|---|---|---|
| S100A2 | BCYRN1 | LTB | MT1G | SAA1 | HP | CXCL10 | S100A4 | CALCA | H19 | PPP1R14A | FOS |
| CISH | MT-TV | TESC | MT1H | SAA2 | CAPN13 | CXCL11 | VPS28 | JAG1 | BIRC2 | PSMD8 | JUN |
| KRT5 | MT-TI | LHFPL3-AS2 | MT1F | AKR1C1 | PPP1R1B | IFIT3 | HCFC1R1 | TNNT3 | TMEM123 | PSMC4 | JUNB |
| CHCHD7 | MT-TL1 | KCNS3 | MT1E | SAA4 | IFITM2 | IFIT2 | NDUFB9 | ARHGAP29 | YAP1 | POLR2I | IER2 |
| COX7A1 | MT-TP | LIMD2 | MT2A | CFB | HPR | IFIT1 | EEF1D | ELP5 | PAGE2 | EIF3K | EGR1 |
| PRKAG2-AS1 | MT-TE | LAPTM4A | MT1M | SLPI | NNMT | OASL | S100A13 | HES4 | CPXM2 | BET1 | DUSP1 |
| SPTSSA | MT-TK | LL22NC03-N | MT1X | AKR1C2 | DEFB1 | ISG15 | TMEM141 | HLA-DMB | SYT8 | ECH1 | ATF3 |
| KCNN4 | MT-ATP8 | DDX52 | MT1L | IKBKE | CHI3L1 | NT5C3A | UPK1B | EMID1 | FGFR2 | GTSF1 | ZFP36 |
| MAPKAPK3 | ZSCAN16-AS | KIZ | MIEN1 | TNFAIP6 | GPX3 | TAP1 | GPX1 | CDH2 | RP11-732A21 | MRPL52 | FOSB |
| MALSU1 | MT-TS2 | RRAD | MT1JP | PDZK1IP1 | DUSP23 | ETV7 | VAMP8 | NOTCH3 | RP3-417G15. | TIMM50 | NR4A1 |
| CDKN2A | MT-TH | SLCO4A1 | ERBB2 | FAM133A | FGL1 | SAMD9L | ANXA2 | IQGAP1 | MCRIP1 | IL32 | CCNL1 |
| CSNK1A1 | PCP4 | COL6A2 | CEND1 | CEND1 | SRD5A3 | OAS1 | OCIAD2 | ABCD3 | ARHGDIA | SPINT2 | GADD45B |
| CLDN4 | ATP5I | TRABD2B | MT1P3 | AKR1C3 | QPCT | TNFSF10 | FIS1 | FOXC1 | RAC3 | EIF2S2 | BTG2 |
| PDK4 | RPL36A | CDH16 | NDUFA1 | LCN2 | IFITM1 | IFIH1 | TATDN1 | COL4A2 | P4HB | RBM42 | KLF10 |
| SLC14A1 | ROMO1 | IL23A | ASS1 | CSTB | CLDN10 | GBP1 | S100A6 | AC116614.1 | AGO2 | NFKBIB | RHOB |
| AGT | NDUFA1 | ICAM2 | MT1A | SERPINB7 | TMEM37 | CMPK2 | CYHR1 | AEN | BMP2 | MED29 | KLF6 |
| CCND1 | MT-ND4L | SLC15A1 | SYCE1L | CHI3L2 | SERPINA1 | GBP4 | EDF1 | CLEC4M | SIRT7 | CDK2AP2 | UBC |
| FAM107A | NUPR2 | LAD1 | MT2P1 | IDO1 | LINC01320 | PARP14 | PPP1R16A | FBN2 | RAB40B | GDF15 | CSRNP1 |
| PID1 | JUND | COMT | AC004057.1 | IER3IP1 | PDZK1IP1 | RSAD2 | QPRT | ATXN1 | PLEC | GNG11 | HES1 |
| RPL22L1 | CBX3 | CYS1 | LRRC55 | CYP24A1 | SLC40A1 | WARS | COX6C | ENC1 | PCGF5 | PSME2 | TRA2B |

Suppl. Table 3

| Dual-12 | Dual-13 | Dual-14 | Dual-15 | Dual-16 | Dual-17 | Dual-18 | Dual-19 | Dual-20 | Dual-21 | Dual-22 | Dual-23 |
|---|---|---|---|---|---|---|---|---|---|---|---|
| CXCL1 | TPX2 | ANKRD1 | CCL20 | HSPA6 | S100A9 | STMN2 | GAGE2A | NDUFA4L2 | PI3 | MMP7 | SST |
| CXCL3 | NUSAP1 | KRTAP2-3 | CXCL8 | DNAJB1 | S100A8 | DCX | GAGE12H | NXPH4 | SLPI | TMEM123 | XIST |
| CXCL8 | BIRC5 | NNMT | SAT1 | HSPA1B | LCN2 | ELAVL3 | GAGE1 | PGK1 | KRT6B | PAGE2 | GBA3 |
| CXCL2 | PTTG1 | HLA-DRB1 | CFLAR | BAG3 | S100A12 | RTN1 | NDUFS4 | SLC2A1 | AKR1C1 | P4HB | SAT1 |
| BIRC3 | UBE2C | MGP | HOTAIRM1 | HSPA8 | S100A7 | INSM1 | GAGE12J | BNIP3 | BASP1 | BIRC2 | BRI3 |
| SOD2 | TOP2A | HLA-DRB5 | FAM134B | HSP90AA1 | S100A7A | TAGLN3 | TUBB2B | P4HA1 | MANF | YAP1 | MEIS2 |
| IFNGR2 | HMGB2 | HLA-DPB1 | TNFAIP6 | HSPA1A | S100P | NEUROD1 | LHX2 | FTLP3 | LAMB3 | ARHGDIA | IQGAP1 |
| ADAMTS9 | MKI67 | LGALS3BP | POU5F1 | HSPA7 | APOBEC3A | MIAT | FAM49B | CA9 | CDSN | CPXM2 | PON2 |
| CAV2 | CDC20 | THBS1 | TANK | CACYBP | PDZK1IP1 | NKX2-2 | NREP | LINC01503 | SBSN | BIRC3 | CLEC2B |
| TNIP1 | CENPF | HLA-DPA1 | PDLIM5 | HSPH1 | DHRS9 | CELF3 | RP11-475C1( | SAT2 | TIMP1 | MCRIP1 | CFLAR |
| SEMA3C | CENPA | CCDC71L | TFPI2 | ZFAND2A | SLC16A3 | NHLH2 | GAGE10 | CTDNEP1 | AKR1C3 | RPS19BP1 | RNASET2 |
| IL1RN | NUF2 | HLA-DRA | RIPK2 | IER5 | S100A11 | GNG8 | RPS23P8 | RNASET2 | MRPS12 | POMP | ABAT |
| GJB2 | CCNB1 | RP11-54H7.4 | LITAF | HSPB1 | NCCRP1 | ELAVL4 | SLC8A1 | TPI1 | CD40 | CCND1 | SEC61B |
| ICAM1 | CDCA3 | HLA-DMA | SQRDL | SERPINH1 | LY6E | GNG3 | LRRC75A-AS | TMEM45A | C19orf33 | PRLR | GPNMB |
| SGPP2 | CDKN3 | ADPRHL1 | IL1B | HSPE1 | PGLYRP4 | DPYSL5 | RPL18AP3 | ENO1 | TGFB1 | NETO2 | ADRM1 |
| LAMB3 | ASPM | STK17A | CD200 | DEDD2 | PRSS27 | NHLH1 | UBE2E3 | ALDOA | KRT6A | TSPAN14 | ABCD3 |
| MGLL | CCNA2 | CD74 | DAPP1 | DNAJB4 | PTGES | TUBB2B | CCT8 | KRT10 | CSTB | IGFL1 | ABCA4 |
| WTAP | AURKA | MAFB | SGPP2 | CHORDC1 | MAPK13 | LBX1 | ERBB4 | NME2P1 | MGLL | CRELD2 | POU5F1 |
| CXCL6 | SMC4 | RUNX1 | TNIP1 | DYNLL1 | PRSS22 | STMN4 | RPS21P4 | RP11-371A2: | TGM1 | ABLIM3 | HOTAIRM1 |
| PTX3 | DLGAP5 | ZFP36L1 | ATP1B1 | DNAJA4 | ADGRF1 | MLLT11 | SNRPE | FTH1P8 | SERP1 | LGMN | TMEM243 |

Suppl. Table 3

| Dual-24 | Dual-25 | Dual-26 | Dual-27 | Dual-28 | Dual-29 | Dual-30 | Dual-31 | Dual-32 | Dual-33 | Dual-34 | Dual-35 |
|---|---|---|---|---|---|---|---|---|---|---|---|
| LTF | KRT81 | RBM39 | PLD3 | RP11-338I21 | NAA38 | GRB7 | FN1 | G0S2 | RPL13 | REG1A | LYPD2 |
| RARRES3 | AARD | SFPQ | LDHB | GTSF1 | POLR2A | MIEN1 | AC006262.5 | NRP2 | RPL18A | GPNMB | LYNX1 |
| GLIPR2 | ALDH2 | EIF4G2 | ISLR | CTAG2 | SAT2 | PPP1R1B | MUC5B | MTSS1 | RPS6 | LINC01320 | MUC4 |
| MESP1 | BEST1 | EIF5 | MGST1 | BET1 | GSTP1 | MX2 | SPOCK2 | SLC25A38 | RPL29 | CFI | FAM83A |
| CHCHD10 | KLK8 | EIF3A | COMT | RP11-88I21.1 | RIOK1 | TERB2 | CST6 | MKNK2 | RPSA | ASRGL1 | TMEM176A |
| C3 | SPDEF | GTPBP4 | SRP9 | CSTA | SP8 | MIA | FOXC1 | CCDC109B | RPS8 | HGD | PTGES |
| TMC5 | MT-TM | DDX5 | SNHG5 | GNG11 | MTHFD1L | IFITM1 | MUC4 | RP11-700H1 | RPS11 | C4BPA | SCEL |
| TTC39A | PPP1R1A | LRRFIP1 | FCGRT | SIPA1L3 | POP7 | LAMP3 | PTGES | C4orf3 | RPS12 | SCGB2A1 | TMEM176B |
| STXBP6 | DNPH1 | HSP90AB1 | NAP1L1 | GART | CTDNEP1 | FAM153B | ABCA4 | FAM177A1 | RPL13A | PPIC | KRT4 |
| KLK7 | CCR1 | CEBPZ | PRDX3 | POLR2J | PHF23 | CMPK2 | LMTK3 | AC105339.1 | RPS16 | C10orf11 | AOC1 |
| NPC2 | RP11-435D7. | PRRC2C | HOXB-AS3 | TFF2 | NUP93 | DAPL1 | CTSD | SNCG | RPL28 | LAMTOR5 | ASPG |
| GM2A | SH2D1B | HNRNPA0 | UBE2L3 | CLEC2L | WDR74 | FOXA2 | MELTF | RHOF | RPL14 | TMEM37 | MAL2 |
| C1orf21 | MT1X | SYNCRIP | RANBP1 | PFDN4 | HMGA1 | CCL2 | ITGAV | PTPRS | RPL24 | PRUNE2 | ATP6V1B1 |
| NFIB | KLK5 | BZW1 | C1QBP | BLVRB | RP11-889L3. | DEFB4A | ADGRG1 | RP11-362F1 | RPL15 | SLC3A1 | SDC1 |
| GLUL | KRT5 | WAC | ACP1 | PSME2 | LYAR | FAM153A | PXDN | C19orf43 | RPL32 | C2orf88 | MUC1 |
| DDT | SLC34A2 | ASH1L | FBXO17 | POLR2F | YBX3 | SLFN5 | NBL1 | RP11-632C1 | RPS25 | QPCT | CXCL17 |
| CCDC107 | MT-TR | HNRNPK | STRA6 | MRPL52 | KMT5B | PIGR | SPARC | INSIG1 | RPS5 | FGL1 | PHACTR3 |
| NTN4 | MIF | MALAT1 | IGHM | ROMO1 | ODC1 | IGKC | RAB31 | GMPR | RPL7 | ERP27 | FXYD3 |
| BBC3 | FLNA | KIF5B | NPW | CSTB | LRPAP1 | RTP4 | C9orf16 | AKT1 | RPL11 | CCDC146 | LY6D |
| MUC1 | HEBP2 | NORAD | HOXB9 | SNRPA | PSMD11 | NT5C3A | TRIM29 | SLC25A3P1 | RPL18 | MAOA | GDPD3 |

Suppl. Table 3

| Dual-36 | Dual-37 | Dual-38 | Dual-39 | Dual-40 | Dual-41 | Dual-42 | Dual-43 |
|---|---|---|---|---|---|---|---|
| CCL3 | PTGDS | C20orf85 | RARRES1 | LAMA5 | WFDC21P | CRYAB | KRT6B |
| CCL3L3 | IFI6 | FAM183A | GLIPR2 | PLEC | SPRR2A | HSPB1 | CTC-575D19.1 |
| TYROBP | SCG3 | PIFO | RARRES3 | FAM83H | S100P | NUPR1 | SERPINB1 |
| C1QA | EEF2 | C5orf49 | NPC2 | CD81 | AQP3 | ATP6V0E1 | SERPINB2 |
| APOC1 | CHGB | C1orf194 | PDZK1IP1 | COL18A1 | KRT6A | CD59 | TICAM1 |
| C1QC | RPL36A | EFCAB1 | TMC5 | SERF2 | SERPINB5 | HLA-C | KRT17 |
| LAPTM5 | WFDC2 | CAPSL | KLK5 | GLIS2 | SPRR2E | LGALS3 | RAB3IP |
| C1QB | APOA1 | C11orf88 | EVA1C | SCRIB | KRT13 | KRT7 | IRAK3 |
| FCER1G | SRRM3 | ROPN1L | HEBP2 | C17orf89 | SPRR2D | CAPG | OAS3 |
| FCGR2A | MSLN | TEKT1 | VTCN1 | NCOR2 | PRSS27 | S100A1 | MGLL |
| ZEB2 | BEX1 | C9orf24 | KLK7 | EIF3B | GPRC5A | HSP90AA1 | HSFY1P1 |
| LILRB4 | RTBDN | ZMYND10 | CHCHD10 | PLXND1 | LYPD3 | B2M | PDLIM1 |
| SPP1 | PARP14 | FAM92B | NFIB | MDM2 | KRT4 | VAMP8 | TRAF3IP2 |
| LCP2 | HNRNPA1 | CFAP126 | DUSP23 | RBM38 | KRT23 | HLA-B | TPRA1 |
| RASSF8-AS1 | BSG | RSPH1 | GAS6 | CDV3 | ZFP36L2 | CST6 | TMBIM4 |
| LYZ | PCOLCE | SNTN | PNRC1 | MT-ND4 | GNA15 | UCA1 | CDSN |
| CCL4 | CD63 | TUBA4B | TC2N | COX5A | UPK2 | MAL | TP53BP2 |
| GPR183 | REC8 | CCDC78 | NTN4 | ATP5I | S100A11 | DNAJB1 | PLSCR1 |
| KRAS | ATP6V1B1 | CAPS | GLUL | INF2 | S100A14 | NBL1 | IFIT1 |
| SRGN | FBXO2 | RSPH9 | C3 | PGAM2 | KRT10 | SFTA2 | RAC1 |

Suppl. Table 4

| N1 | N2 | N3 | N4 | C1 | C2 | C3 | C4 | C5 |
|---|---|---|---|---|---|---|---|---|
| ABHD14B | AC013461.1 | ANXA1 | ADIRF | AKAP12 | ADRM1 | ALG3 | ANAPC5 | AGRN |
| APP | AGR2 | ATF3 | APRT | ARHGAP5 | ANXA2 | ANXA7 | ASAP1 | APOA1 |
| AR | AKAP9 | BAG3 | ATP5G2 | ARL16 | AP2M1 | BAX | B4GALT5 | APOE |
| ARL6IP5 | ALDH1A1 | BTG1 | BSG | ARL4C | ARF1 | C4orf3 | BCCIP | CCL20 |
| ASRGL1 | APOBEC3C | BTG2 | CIRBP | ASPH | ARPC2 | CACYBP | BCL7B | CLDN6 |
| BORCS7 | C1orf168 | CCNL1 | COX4I1 | BACE2 | ATP5B | CAPG | BID | CREB5 |
| CAPS | C2orf88 | CLK1 | CYB5A | BLVRB | ATP5E | CD81 | BLCAP | CRYAB |
| CCNG1 | CCL2 | CSRNP1 | EDF1 | BST2 | ATP5L | CDKN2A | BLOC1S2 | CTGF |
| CD74 | CFI | CYR61 | EEF2 | BZW2 | ATP6V0B | CISD1 | CCDC6 | CXCL8 |
| CD99 | CHD1 | DDX3X | EID1 | C1orf186 | BRK1 | CKB | CHRAC1 | DDIT4 |
| CLU | CHD9 | DNAJB1 | EIF3E | C1orf52 | C14orf2 | COA4 | CITED4 | DEFB1 |
| CNDP2 | CLDN10 | DUSP1 | EIF3L | CAB39 | C19orf53 | DNAJC19 | CNIH1 | FBXO32 |
| CRISP3 | CXCL2 | EGR1 | EIF4A2 | CAV2 | C20orf24 | ECH1 | COMMD2 | FOLR1 |
| CSRP1 | CXCR4 | EHF | EIF4B | CDC42EP3 | CAMTA1 | ECI1 | COPS2 | GAL |
| CTSH | DDX17 | FOS | GABARAPL2 | CITED2 | CD63 | FAM162A | COTL1 | GBP1 |
| DPP7 | ECHDC2 | FOSB | GSTK1 | CKS2 | CFL1 | HLA-B | CPSF6 | HDAC9 |
| DUT | ELF1 | FUS | HINT1 | CLIC4 | CHCHD2 | HMGN3 | DCUN1D5 | HIST3H2A |
| EIF2S3 | EMX2 | GADD45B | HNRNPA1 | CLIP1 | CHMP4B | IDH3B | DDX39A | HMGA2 |
| ERP27 | FNBP1 | HSP90AA1 | HSP90AB1 | COX19 | CLIC1 | KRT7 | DIAPH1 | HMGB2 |
| ESD | FRMD4B | HSPA1B | IFITM2 | CRK | CLTA | LAMP1 | DKC1 | IFI6 |
| ETFB | GADD45G | HSPA8 | ITM2B | DBN1 | CNBP | LAMTOR2 | DNPEP | IGF2 |
| FAM107A | H2AFX | HSPB1 | NAP1L1 | DCP1A | COPE | LDHB | EHD1 | IGFBP3 |
| FAM200B | HLA-DPA1 | HSPH1 | PCBD1 | DDX49 | COPS9 | LRPAP1 | EMP1 | IRF7 |
| FBXO21 | HLA-DQB1 | ID4 | PFDN5 | DFFA | COX5A | LSM2 | EXOSC4 | ISG20 |
| FCGRT | HLA-DRA | IER2 | PHPT1 | DNAJB9 | COX6A1 | MDK | FNBP4 | KDM6B |
| FGGY | HLA-DRB1 | IFRD1 | RBMX | DTD1 | COX6C | NDUFB5 | FNTA | KLK10 |
| GAS6 | HOOK2 | JUN | RBP1 | EPHA2 | CSNK2B | PAFAH1B3 | GARS | KLK7 |
| HLA-DMA | HSPA1A | JUNB | RNASET2 | ESF1 | CYC1 | PET100 | GNL2 | KRT17 |
| IGFBP7 | HSPB8 | KLF5 | SLC25A6 | ETS2 | EEF1D | PFDN6 | KCNK1 | LDLR |
| ISYNA1 | INTS6 | KMT2E | SMDT1 | FABP5 | EIF2S2 | PIGT | KPNA2 | MAL |
| KLK11 | LCN12 | MAFF | SNRPD2 | FAM208B | EIF6 | PLD3 | LAMB3 | MFAP2 |
| LITAF | LYPD6B | MAP1LC3B | ST13 | FAM50A | ENY2 | PLP2 | MAFG | MGST1 |
| LMAN1 | MPHOSPH8 | MYADM | TMEM219 | FAM84B | EPCAM | PNKD | MAP4K4 | MMP7 |
| LTC4S | NFAT5 | MYLIP | TSPO | FAM91A1 | GADD45GIP1 | PSMF1 | MAPK6 | MT1E |
| MGMT | OFD1 | NR4A1 | UXT | FOSL1 | GAPDH | RNF5 | MAX | MT1F |
| MGST2 | OVGP1 | PNRC1 | NA | GSK3B | GNB2 | SDHC | MPHOSPH10 | MT1G |
| MPST | PHIP | PPP1R15A | NA | H2AFJ | H3F3B | SLC39A4 | MYO10 | MT1X |
| NME3 | PIK3R1 | SOX17 | NA | HECTD1 | HNRNPC | SSNA1 | NELFCD | MUC16 |
| NSMCE1 | PNISR | TRA2A | NA | HES4 | MDH2 | STRA13 | NGDN | MX1 |
| NUCB2 | PPP1R10 | TRA2B | NA | HSPA4 | MLF2 | TPM1 | NOC2L | NUPR1 |
| OCIAD1 | PRKX | ZFP36 | NA | HSPB11 | NDUFB9 | TRAPPC1 | NOLC1 | PI3 |
| OSTC | RBM47 | ZFP36L1 | NA | HTATSF1 | PCBP2 | TSTA3 | NRBP1 | PIM1 |
| PABPC4 | RNASE1 | ZFP36L2 | NA | IER3 | PFDN2 | TUSC2 | PARP14 | PLA2G16 |
| PCM1 | RRAD | NA | NA | IFI27 | PKM | WFDC2 | PCGF5 | PLAU |
| PDLIM1 | RSRP1 | NA | NA | ISG15 | POMP | NA | PDLIM7 | PLTP |
| PKP2 | SLC25A25 | NA | NA | LARP6 | PPA1 | NA | PFKP | PRSS23 |
| PMVK | SLC25A29 | NA | NA | LRPPRC | PRELID1 | NA | PHAX | PTX3 |
| PRDX4 | SMC5 | NA | NA | LYN | PSMB5 | NA | PHF20 | RGCC |
| PSMA2 | SNCG | NA | NA | MARCH7 | PTP4A2 | NA | PLEC | TNF |
| QARS | SOCS3 | NA | NA | MEST | RAC1 | NA | PLSCR1 | TPM2 |
| QPRT | SORL1 | NA | NA | MSN | RALY | NA | PNO1 | TUBA1A |
| RASSF7 | SPARCL1 | NA | NA | MUC1 | RAN | NA | PPHLN1 | UBB |
| RBBP7 | STAT3 | NA | NA | NDFIP2 | RBM8A | NA | PRPF6 | UBL3 |
| RCN2 | STX18 | NA | NA | NPM3 | RHEB | NA | PSMC2 | UCHL1 |
| RNH1 | SYNE2 | NA | NA | OGFR | RHOA | NA | PSMD6 | NA |
| SCGB2A1 | TACC1 | NA | NA | PBDC1 | RNF7 | NA | PTPN11 | NA |
| SEPP1 | TANK | NA | NA | PDLIM2 | SEC61B | NA | PWP1 | NA |

Suppl. Table 4

| N1 | N2 | N3 | N4 | C1 | C2 | C3 | C4 | C5 |
|---|---|---|---|---|---|---|---|---|
| SERPINB6 | TERF2IP | NA | NA | PHLDA3 | SEC61G | NA | RAB3IP | NA |
| SLC40A1 | WEE1 | NA | NA | PITPNB | SERBP1 | NA | RIPK2 | NA |
| SYF2 | WT1 | NA | NA | PMEPA1 | SF3B6 | NA | RNF126 | NA |
| SYPL1 | ZNF83 | NA | NA | PML | SLC25A3 | NA | RRP15 | NA |
| TMC4 | NA | NA | NA | POR | SLC25A5 | NA | SLC25A33 | NA |
| TMEM141 | NA | NA | NA | PPFIA1 | SNRPB | NA | SMARCA4 | NA |
| TMEM256 | NA | NA | NA | PTK2 | SNRPE | NA | SMARCC1 | NA |
| TMEM98 | NA | NA | NA | PTPRA | SNRPF | NA | SPTSSA | NA |
| TPD52L1 | NA | NA | NA | RABGGTB | SNU13 | NA | SQLE | NA |
| TTC3 | NA | NA | NA | RAE1 | SOD1 | NA | SRPK2 | NA |
| TTYH1 | NA | NA | NA | RAI14 | SPINT2 | NA | STAT1 | NA |
| TXN2 | NA | NA | NA | RAP1B | SRSF9 | NA | SUPT16H | NA |
| NA | NA | NA | NA | RIF1 | SSU72 | NA | SVIL | NA |
| NA | NA | NA | NA | RNF19A | SUB1 | NA | TAF9 | NA |
| NA | NA | NA | NA | RNF213 | SUMO1 | NA | TMEM189 | NA |
| NA | NA | NA | NA | RRAS2 | TCEB1 | NA | TNFRSF21 | NA |
| NA | NA | NA | NA | RUNX1 | TCEB2 | NA | TP53RK | NA |
| NA | NA | NA | NA | S100A1 | TMA7 | NA | TRIAP1 | NA |
| NA | NA | NA | NA | SF3A3 | TMBIM6 | NA | TSN | NA |
| NA | NA | NA | NA | SFN | TMED2 | NA | TUBB6 | NA |
| NA | NA | NA | NA | SFSWAP | TMSB10 | NA | TWF1 | NA |
| NA | NA | NA | NA | SHOC2 | TOMM20 | NA | TWISTNB | NA |
| NA | NA | NA | NA | SIAH2 | TPI1 | NA | UBE2E2 | NA |
| NA | NA | NA | NA | SLC2A1 | TPM3 | NA | UCHL3 | NA |
| NA | NA | NA | NA | SLC9A3R2 | TRMT112 | NA | UGCG | NA |
| NA | NA | NA | NA | SLCO3A1 | UBA52 | NA | USP34 | NA |
| NA | NA | NA | NA | SMAD3 | UQCRFS1 | NA | UTP11 | NA |
| NA | NA | NA | NA | SMIM15 | VDAC2 | NA | VAMP3 | NA |
| NA | NA | NA | NA | SP100 | YBX1 | NA | WDR74 | NA |
| NA | NA | NA | NA | SPECC1 | YBX3 | NA | WWTR1 | NA |
| NA | NA | NA | NA | SPEN | YWHAB | NA | YKT6 | NA |
| NA | NA | NA | NA | ST3GAL1 | YWHAE | NA | ZNF22 | NA |
| NA | NA | NA | NA | TBCC | YWHAQ | NA | ZNF622 | NA |
| NA | NA | NA | NA | TIMM23 | YWHAZ | NA | NA | NA |
| NA | NA | NA | NA | TMEM70 | NA | NA | NA | NA |
| NA | NA | NA | NA | TNNT1 | NA | NA | NA | NA |
| NA | NA | NA | NA | TRIP12 | NA | NA | NA | NA |
| NA | NA | NA | NA | TSPAN14 | NA | NA | NA | NA |
| NA | NA | NA | NA | U2AF2 | NA | NA | NA | NA |
| NA | NA | NA | NA | UBAP1 | NA | NA | NA | NA |
| NA | NA | NA | NA | UBE2G2 | NA | NA | NA | NA |
| NA | NA | NA | NA | UBXN2A | NA | NA | NA | NA |
| NA | NA | NA | NA | UTP23 | NA | NA | NA | NA |
| NA | NA | NA | NA | YPEL3 | NA | NA | NA | NA |
| NA | NA | NA | NA | ZFAND1 | NA | NA | NA | NA |
| NA | NA | NA | NA | ZNF326 | NA | NA | NA | NA |
| NA | NA | NA | NA | ZNRD1 | NA | NA | NA | NA |

Suppl. Table 4

| **C6** | **C7** | **C8** | **C9** | **E1** | **E2** | **E3** | **E4** |
|---|---|---|---|---|---|---|---|
| ADAR | ANXA11 | ANAPC11 | ARL6IP1 | AC090498.1 | ACTG1 | ALCAM | ANXA11 |
| ANKLE2 | ATG101 | AP2S1 | BAG1 | ADI1 | ALKBH7 | AMOTL2 | ATP1A1 |
| ARHGAP29 | AUP1 | ARPC1B | CAPZA2 | ADIRF | ARL6IP4 | ANXA3 | BCAM |
| ARPP19 | BRD2 | ATP5G1 | CDC37 | AHCY | ATP5A1 | ARHGAP29 | CAPN2 |
| CCND1 | BRD4 | ATP5H | CHMP2B | APEX1 | ATP5C1 | ATF3 | CD151 |
| CEBPB | BUD31 | BCAP31 | DAZAP2 | ARL2 | ATP5D | ATP1B1 | CD24 |
| CEBPG | BZW1 | C11orf31 | DDX18 | BEX2 | ATP5F1 | BHLHE41 | CD47 |
| CEBPZ | CBX3 | CAPZB | EBNA1BP2 | BEX4 | ATP5G1 | BPTF | CD9 |
| CLDN4 | CCT2 | CD47 | EIF5A | C19orf48 | ATP5G2 | C6orf132 | CLDN3 |
| CNN3 | CCT3 | CD9 | FAM32A | CAPS | ATP5O | CCND1 | CLDN7 |
| DDX1 | CCT5 | COX7A2L | FUNDC2 | CCNG1 | ATPIF1 | CD2AP | CMTM6 |
| DDX27 | CCT6A | CST3 | GID8 | CETN2 | BANF1 | CLDN1 | CNPY2 |
| DENR | CDC42 | CUTA | HLA-E | CHMP3 | BEX3 | CP | DDAH2 |
| DHX9 | CLTB | DAD1 | HM13 | CLNS1A | BSG | CRYAB | DMKN |
| DNTTIP2 | CSDE1 | DYNLL1 | HNRNPF | CLU | C11orf31 | CTGF | EIF4A2 |
| EIF2AK2 | DDX21 | ERGIC3 | HNRNPR | CMTM7 | C19orf33 | CXADR | EMC10 |
| EIF3B | DDX5 | FKBP1A | HSF1 | COA4 | C6orf48 | CYR61 | FLOT1 |
| EIF4A3 | EIF3A | FKBP2 | IFT57 | COL18A1 | C8orf59 | DOK5 | HSPB1 |
| ELOF1 | EIF4G1 | FTL | IMP4 | CRABP2 | CCT3 | EDN1 | HSPD1 |
| ENAH | EIF4H | GHITM | LYPLA1 | CSRP1 | CLTA | EFNA1 | JUP |
| FAM103A1 | EIF5 | HEBP2 | MARCKSL1 | DNAJC4 | COX5B | EFNB2 | KRT19 |
| FAM204A | EIF5B | HSP90B1 | MEAF6 | DNPH1 | COX7A2L | EHF | KRT7 |
| FAM60A | EZR | HSPE1 | MED10 | DUT | COX7C | ELF3 | KRT8 |
| FOSL2 | FAM49B | KDELR2 | MMADHC | EBPL | CUTA | EMX2 | LSR |
| FYTTD1 | GTPBP4 | KRT18 | NAA20 | ECH1 | CYC1 | EPHA2 | MAL2 |
| GOLGA7 | HMGA1 | KRT8 | PAIP1 | FDFT1 | DDT | GADD45A | MDK |
| H1FX | HN1 | KRTCAP2 | POP7 | FKBP4 | DYNLL1 | HES1 | MSLN |
| HIF1A | HNRNPAB | MAL2 | PRELID3B | FOLR1 | ECHS1 | HOOK2 | NECTIN2 |
| ILF3 | HSPA9 | MGST3 | PSMA3 | FXYD3 | EIF3G | ID4 | PDLIM1 |
| ITGB8 | ITGB1 | NDUFA2 | PSMC1 | GAS6 | EIF3H | IER2 | PERP |
| JAK1 | KHDRBS1 | NDUFB10 | PSMD14 | GCSH | EIF3K | IGF2BP2 | PRMT1 |
| JUND | KIF5B | NDUFC1 | RALA | GGCT | EIF3M | IP6K2 | RALBP1 |
| KRR1 | KTN1 | NEDD8 | RALBP1 | GLO1 | EIF4B | LDLR | RCN2 |
| LLPH | LRRFIP1 | NPC2 | RNF114 | GRHPR | EPCAM | MMP7 | RNASET2 |
| LMO4 | METAP2 | PGK1 | RPN1 | GTF2H5 | ERGIC3 | NCOA7 | SYNGR2 |
| MAPK1IP1L | MTPN | PHB | SAR1A | HDDC2 | FBL | NET1 | TMEM205 |
| MLLT4 | NCL | POLR2J | SARS | IFT57 | FIS1 | PARD6B | WFDC2 |
| NAA50 | PA2G4 | POLR2L | SEPT7 | ISYNA1 | FKBP2 | PAWR | NA |
| NIFK | PFDN4 | PPDPF | SMS | KLK11 | GSTP1 | PBX1 | NA |
| NOP16 | POLR1D | PSAP | SNRPA1 | KRTCAP3 | HIGD2A | PNN | NA |
| PAX8 | PPM1G | RBX1 | SSB | LYPD1 | HINT1 | PRKCI | NA |
| PTBP1 | PPP1CC | SCAND1 | TCP1 | MAP3K13 | HMGN1 | PUM1 | NA |
| RAB18 | PRMT1 | SH3BGRL3 | TMEM123 | MGMT | HMGN2 | RBM47 | NA |
| RAB22A | PRPF40A | STOML2 | TMEM183A | MGST2 | HMGN3 | SCGB2A1 | NA |
| RAD21 | PSMD2 | TMEM50A | TOMM40 | MYL6B | LAMTOR4 | SDC4 | NA |
| RBBP6 | PUF60 | UQCR10 | TOMM5 | NDUFA10 | LAPTM4B | SERTAD1 | NA |
| RNF145 | RBM17 | UQCR11 | TUBB4B | NDUFB5 | LINC00493 | SLC25A36 | NA |
| STARD7 | RNPS1 | VDAC1 | ZNF593 | NDUFS4 | LSM4 | SNCG | NA |
| TACSTD2 | SBDS | NA | ZYX | OLA1 | MDH2 | SOX17 | NA |
| TARS | SNRPB2 | NA | NA | PGRMC1 | MGST3 | TACSTD2 | NA |
| TDG | SRSF2 | NA | NA | PMVK | MTCH1 | TJP1 | NA |
| TNIP1 | SRSF3 | NA | NA | POLR2G | MZT2A | TM4SF1 | NA |
| TRMT10C | STAU1 | NA | NA | PPP1R16A | MZT2B | TOB1 | NA |
| TSC22D1 | STRAP | NA | NA | PRDX3 | NAA38 | ZNF503 | NA |
| TXNRD1 | TAX1BP1 | NA | NA | PRKCSH | NACA | NA | NA |
| WDR43 | TIMM17A | NA | NA | PTRHD1 | NDUFA11 | NA | NA |
| YWHAG | TRAM1 | NA | NA | QPRT | NDUFA4 | NA | NA |

| **C6** | **C7** | **C8** | **C9** | **E1** | **E2** | **E3** | **E4** |
|---|---|---|---|---|---|---|---|
| ZFR | TUBA1C | NA | NA | RAB34 | NDUFB11 | NA | NA |
| NA | VAPA | NA | NA | RASSF7 | NDUFB7 | NA | NA |
| NA | WBSCR22 | NA | NA | RBBP7 | NDUFB9 | NA | NA |
| NA | ZC3H15 | NA | NA | RPAIN | NDUFC1 | NA | NA |
| NA | ZFAND5 | NA | NA | S100A1 | NENF | NA | NA |
| NA | NA | NA | NA | S100A14 | NHP2 | NA | NA |
| NA | NA | NA | NA | SH3YL1 | NPM1 | NA | NA |
| NA | NA | NA | NA | SMIM19 | NUCKS1 | NA | NA |
| NA | NA | NA | NA | SMIM22 | PCBD1 | NA | NA |
| NA | NA | NA | NA | SUCLG1 | PEBP1 | NA | NA |
| NA | NA | NA | NA | SYPL1 | PGLS | NA | NA |
| NA | NA | NA | NA | TCEAL4 | PHB | NA | NA |
| NA | NA | NA | NA | TKT | PHB2 | NA | NA |
| NA | NA | NA | NA | TMED3 | PHPT1 | NA | NA |
| NA | NA | NA | NA | TMED4 | PPA1 | NA | NA |
| NA | NA | NA | NA | TMEM141 | PRDX2 | NA | NA |
| NA | NA | NA | NA | TMEM261 | PRDX5 | NA | NA |
| NA | NA | NA | NA | TPD52L1 | PRDX6 | NA | NA |
| NA | NA | NA | NA | TPRKB | RACK1 | NA | NA |
| NA | NA | NA | NA | TSEN34 | S100A13 | NA | NA |
| NA | NA | NA | NA | UCP2 | SNRPD2 | NA | NA |
| NA | NA | NA | NA | UQCRC2 | SNRPE | NA | NA |
| NA | NA | NA | NA | ZCRB1 | SPINT2 | NA | NA |
| NA | NA | NA | NA | NA | STOML2 | NA | NA |
| NA | NA | NA | NA | NA | TCEB2 | NA | NA |
| NA | NA | NA | NA | NA | TIMM13 | NA | NA |
| NA | NA | NA | NA | NA | TMEM14B | NA | NA |
| NA | NA | NA | NA | NA | TMEM14C | NA | NA |
| NA | NA | NA | NA | NA | TMEM219 | NA | NA |
| NA | NA | NA | NA | NA | TSTD1 | NA | NA |
| NA | NA | NA | NA | NA | TUFM | NA | NA |
| NA | NA | NA | NA | NA | UFC1 | NA | NA |
| NA | NA | NA | NA | NA | UQCRH | NA | NA |
| NA | NA | NA | NA | NA | VDAC2 | NA | NA |
| NA | NA | NA | NA | NA | VPS28 | NA | NA |
| NA | NA | NA | NA | NA | WDR83OS | NA | NA |
| NA | NA | NA | NA | NA | YWHAE | NA | NA |
| NA | NA | NA | NA | NA | NA | NA | NA |
| NA | NA | NA | NA | NA | NA | NA | NA |
| NA | NA | NA | NA | NA | NA | NA | NA |
| NA | NA | NA | NA | NA | NA | NA | NA |
| NA | NA | NA | NA | NA | NA | NA | NA |
| NA | NA | NA | NA | NA | NA | NA | NA |
| NA | NA | NA | NA | NA | NA | NA | NA |
| NA | NA | NA | NA | NA | NA | NA | NA |
| NA | NA | NA | NA | NA | NA | NA | NA |
| NA | NA | NA | NA | NA | NA | NA | NA |

**Supplementary Table 5. Drug screening concentrations in PDO1 and PDO2**

| | PDO 1 | | PDO 2 | |
|---|---|---|---|---|
| Drug | Low concentration (µM) | High concentration (µM) | Low concentration (µM)2 | High concentration (µM)3 |
| taselisib | 0.1250 | 0.5000 | 0.12500 | 0.5000 |
| alpelisib | 0.2500 | 1.0000 | 0.25000 | 1.0000 |
| AZD3458 | 0.0625 | 0.2500 | 0.06250 | 0.2500 |
| ceralasertib | 0.2500 | 1.0000 | 0.25000 | 1.0000 |
| navitoclax | 0.2500 | 1.0000 | 0.25000 | 1.0000 |
| A1155463 | 0.1250 | 0.5000 | 0.12500 | 0.5000 |
| birabresib | 0.2500 | 1.0000 | 0.25000 | 1.0000 |
| palbociclib | 0.2500 | 1.0000 | 0.25000 | 1.0000 |
| AZD4573 | 0.0125 | 0.0500 | 0.00125 | 0.0050 |
| prexasertib | 0.0125 | 0.0500 | 0.00250 | 0.0100 |
| orludodstat | 0.0125 | 0.0500 | 0.01250 | 0.0500 |
| AZD7648 | 0.2500 | 1.0000 | 0.25000 | 1.0000 |
| erlotinib | 0.2500 | 1.0000 | 0.25000 | 1.0000 |
| afatinib | 0.1250 | 0.5000 | 0.12500 | 0.5000 |
| everolimus | 0.0125 | 0.0500 | 0.01250 | 0.0500 |
| MK8353 | 0.1250 | 0.5000 | 0.12500 | 0.5000 |
| fludarabine | 0.2500 | 1.0000 | 0.25000 | 1.0000 |
| ruxolitinib | 0.1250 | 0.5000 | 0.12500 | 0.5000 |
| imatinib | 0.2500 | 1.0000 | 0.25000 | 1.0000 |
| revumenib | 0.0500 | 0.2000 | 0.05000 | 0.2000 |
| iadademstat | 0.0250 | 0.1000 | 0.02500 | 0.1000 |
| MIK665 | 0.0625 | 0.2500 | 0.06250 | 0.2500 |
| cobimetinib | 0.1250 | 0.5000 | 0.12500 | 0.5000 |
| nitazoxanide | 2.5000 | 10.0000 | 2.50000 | 10.0000 |
| pevonedistat | 0.1250 | 0.5000 | 0.01250 | 0.0500 |
| AZD8186 | 0.1250 | 0.5000 | 0.12500 | 0.5000 |
| onvansertib | 0.1250 | 0.5000 | 0.01250 | 0.0500 |
| onametostat | 0.0250 | 0.1000 | 0.02500 | 0.1000 |
| bortezomib | 0.0025 | 0.0100 | 0.00250 | 0.0100 |
| naporafenib | 0.5000 | 2.0000 | 0.50000 | 2.0000 |
| dasatinib | 0.0625 | 0.2500 | 0.06250 | 0.2500 |
| entospletinib | 0.2500 | 1.0000 | 0.25000 | 1.0000 |
| TAK243 | 0.0625 | 0.2500 | 0.01250 | 0.0500 |
| adavosertib | 0.0375 | 0.1500 | 0.03750 | 0.1500 |
| LCL161 | 0.5000 | 2.0000 | 0.10000 | 0.4000 |
| acetalax | 1.2500 | 5.0000 | 1.25000 | 5.0000 |
| CGS15943 | 0.0625 | 0.2500 | 0.06250 | 0.2500 |
| GSK2801 | 1.2500 | 5.0000 | 1.25000 | 5.0000 |
| tagtociclib | 0.2500 | 1.0000 | 0.25000 | 1.0000 |
| trifluoromethyl-tubercidin | 1.2500 | 5.0000 | 0.12500 | 0.5000 |
| acipimox | 100.0000 | 400.0000 | 100.00000 | 400.0000 |
| romidepsin | 0.0063 | 0.0250 | 0.00063 | 0.0025 |
| TMP195 | 0.3750 | 1.5000 | 0.37500 | 1.5000 |
| emedastine | 2.5000 | 10.0000 | 2.50000 | 10.0000 |
| onalespib | 0.0625 | 0.2500 | 0.01250 | 0.0500 |
| actarit | 2.5000 | 10.0000 | 2.50000 | 10.0000 |
| batimastat | 0.2500 | 1.0000 | 0.25000 | 1.0000 |
| camicinal | 0.0750 | 0.3000 | 0.07500 | 0.3000 |
| ABT202 | 2.5000 | 10.0000 | 2.50000 | 10.0000 |
| ZLN005 | 5.0000 | 20.0000 | 1.25000 | 5.0000 |
| picolinic acid | 1.2500 | 5.0000 | 1.25000 | 5.0000 |
| APY0201 | 0.0250 | 0.1000 | 0.02500 | 0.1000 |
| zalcitabine | 1.2500 | 5.0000 | 1.25000 | 5.0000 |
| omacetaxine | 0.0125 | 0.0500 | 0.01250 | 0.0500 |
| VS4718 | 0.2500 | 1.0000 | 0.25000 | 1.0000 |
| pyridostatin | 1.0000 | 4.0000 | 1.00000 | 4.0000 |
| YK4279 | 1.2500 | 5.0000 | 1.25000 | 5.0000 |
| triptolide | 0.2500 | 1.0000 | 0.02500 | 0.1000 |
| LJI308 | 1.2500 | 5.0000 | 1.25000 | 5.0000 |
| MST312 | 1.2500 | 5.0000 | 1.25000 | 5.0000 |
| vactosertib | 0.1250 | 0.5000 | 0.12500 | 0.5000 |
| MSC2504877 | 0.0250 | 0.1000 | 0.02500 | 0.1000 |
| varespladib | 1.2500 | 5.0000 | 1.25000 | 5.0000 |
| ISX9 | 6.2500 | 25.0000 | 6.25000 | 25.0000 |
| paclitaxel | 0.0250 | 0.1000 | 0.00250 | 0.0100 |
| bortezomib (positive control) | 10.0 | | 10.0 | |
| DMSO (negative control) | 0.1% v/v | | 0.1% v/v | |
| carboplatin | 20.0 | | 10.0 | |

**Legend**

Adjusted concentrations between experiments

**Supplementary Table 6. Drug concentrations for the long-term survival assays in PDO1 and PDO2**

| | PDO 1 | | PDO 2 | |
|---|---|---|---|---|
| Drug | Concentration (µM) | Treatment duration | Concentration (µM)2 | Treatment duration3 |
| navitoclax | 1.00 | 7 days | 1.00 | 7 days |
| birabresib | 0.25 | 11 days | 0.25 | 12 days |
| revumenib | 0.20 | 11 days | 0.20 | 12 days |
| pevonedistat | 0.10 | 7 days | 0.10 | 7 days |
| LCL161 | 0.20 | 7 days | 0.20 | 7 days |
| carboplatin | 20.00 | 8 hours | 10.00 | 8 hours |
| DMSO | 0.1% | 11 days | 0.1% | 12 days |

### Supplementary Table 7. Drug screening concentrations in PDO 2-7

| Drug | Low concentration (µM) | High concentration (µM) |
|---|---|---|
| Pevonedistat | 0.0125 | 0.050 |
| Xevinapant | 0.1000 | 0.400 |
| ZEN-3694 | 0.2500 | 1.000 |
| Paclitaxel | 0.0025 | 0.010 |
| Birabresib | 0.2500 | 1.000 |
| LCL161 | 0.1000 | 0.400 |

| Carboplatin | Concentration (µM) |
|---|---|
| organoid 3 | 10.00 |
| organoid 5 | 10.00 |
| organoid 4 | 10.00 |
| organoid 2 | 10.00 |
| organoid 7 | 10.00 |
| organoid 6 | 20.00 |
| Bortezomib (positive control) | 10.00 |
| DMSO (negative control) | 0.1% v/v |

| DRUG_ID | DRUG_NAME | SYNONYMS | nervous_system_MWU_STAT | nervous_system_PVAL | nervous_system_PVAL_ADJ | soft_tissue_MWU_STAT | soft_tissue_PVAL | soft_tissue_PVAL_ADJ | bone_MWU_STAT | bone_PVAL | bone_PVAL_ADJ | skin_MWU_STAT | skin_PVAL | skin_PVAL_ADJ | urogenital_system_MWU_STAT | urogenital_system_PVAL | urogenital_system_PVAL_ADJ | lung_NSCLC_MWU_STAT | lung_NSCLC_PVAL | lung_NSCLC_PVAL_ADJ | lung_SCLC_MWU_STAT | lung_SCLC_PVAL | lung_SCLC_PVAL_ADJ | lung_MWU_STAT | lung_PVAL | lung_PVAL_ADJ | neuroblastoma_MWU_STAT | neuroblastoma_PVAL | neuroblastoma_PVAL_ADJ | pancreas_MWU_STAT | pancreas_PVAL | pancreas_PVAL_ADJ | aero_dig_tract_MWU_STAT | aero_dig_tract_PVAL | aero_dig_tract_PVAL_ADJ | breast_MWU_STAT | breast_PVAL | breast_PVAL_ADJ | kidney_MWU_STAT | kidney_PVAL | kidney_PVAL_ADJ |
|---|---|---|---|---|---|---|---|---|---|---|---|---|---|---|---|---|---|---|---|---|---|---|---|---|---|---|---|---|---|---|---|---|---|---|---|---|---|---|---|---|---|
| 1003 | Camptothecin | Camptothecine, (+)-Camptothecin | 17189 | 0.069 | 0.159 | 5528 | 0.009 | 0.046 | 8492 | 0.000 | 0.000 | 21011 | 0.759 | 1.000 | 33003 | 0.287 | 0.574 | 40912.5 | 0.993 | 1.000 | 18093 | 0.066 | 0.159 | 9146 | 0.788 | 1.000 | 5178 | 0.000 | 0.000 | 13639 | 0.954 | 1.000 | 27498 | 0.671 | 1.000 | 25973 | 1.000 | 1.000 | 9677 | 0.019 | 0.070 |
| 1004 | Vinblastine | Velban | 8988 | 0.062 | 0.199 | 3891 | 0.026 | 0.131 | 7448 | 0.018 | 0.131 | 8960 | 0.501 | 0.890 | 23114 | 0.660 | 0.961 | 24504 | 0.964 | 1.000 | 8670 | 0.033 | 0.131 | 6751 | 0.998 | 1.000 | 4394 | 0.002 | 0.037 | 11668 | 1.000 | 1.000 | 14122 | 0.102 | 0.271 | 14785 | 0.982 | 1.000 | 5822 | 0.753 | 1.000 |
| 1005 | Cisplatin | cis-Diammineplatinum(II) dichloride, Platinol, CIS-DDP | 9272 | 0.041 | 0.130 | 3923 | 0.021 | 0.083 | 5132 | 0.000 | 0.000 | 8853 | 0.359 | 0.639 | 21368 | 0.096 | 0.257 | 25175 | 0.944 | 1.000 | 9577 | 0.016 | 0.083 | 6219 | 0.971 | 1.000 | 3249 | 0.000 | 0.000 | 11404 | 0.995 | 1.000 | 15240 | 0.179 | 0.410 | 21736 | 1.000 | 1.000 | 4953.5 | 0.250 | 0.500 |
| 1006 | Cytarabine | Ara-Cytidine, Arabinosyl Cytosine, U-19920 | 11257 | 0.730 | 0.899 | 2646 | 0.000 | 0.002 | 7212 | 0.009 | 0.049 | 7072 | 0.023 | 0.090 | 25329 | 0.952 | 1.000 | 22988 | 0.715 | 0.899 | 10689 | 0.527 | 0.899 | 5547 | 0.871 | 0.995 | 4477 | 0.003 | 0.024 | 8272 | 0.429 | 0.899 | 16281 | 0.651 | 0.899 | 17581 | 1.000 | 1.000 | 4575 | 0.151 | 0.403 |
| 1007 | Docetaxel | RP-56976, Taxotere | 14456 | 0.001 | 0.006 | 5986 | 0.055 | 0.176 | 10711.5 | 0.013 | 0.068 | 22190 | 0.927 | 1.000 | 35465.5 | 0.736 | 1.000 | 36966.5 | 0.743 | 1.000 | 20915 | 0.596 | 1.000 | 11936.5 | 1.000 | 1.000 | 8404 | 0.020 | 0.079 | 15491 | 0.999 | 1.000 | 18231 | 0.000 | 0.000 | 20317.5 | 0.919 | 1.000 | 14878 | 0.980 | 1.000 |
| 1008 | Methotrexate | Abitrexate, Amethopterin, Rheumatrex, Trexall, Folex | 21562.5 | 0.999 | 0.999 | 5732 | 0.076 | 0.202 | 7479 | 0.002 | 0.019 | 21077 | 0.986 | 0.999 | 28635 | 0.201 | 0.379 | 28487 | 0.058 | 0.199 | 20175 | 0.989 | 0.999 | 6649 | 0.213 | 0.379 | 11361.5 | 0.998 | 0.999 | 8367 | 0.018 | 0.094 | 24251 | 0.873 | 0.999 | 15637.5 | 0.681 | 0.999 | 13751.5 | 0.976 | 0.999 |
| 1009 | Tretinoin | ATRA, Vesanoid, Renova, Atralin, Tretin-X, Avita | 716 | 0.003 | 0.051 | 170 | 0.520 | 0.933 | 466 | 0.700 | 0.933 | 1360 | 0.015 | 0.119 | 1136.5 | 0.617 | 0.933 | 1758 | 0.642 | 0.933 | 2030 | 0.996 | 0.996 | 599 | 0.826 | 0.944 | 540 | 0.669 | 0.933 | 107 | 0.194 | 0.619 | 1436 | 0.654 | 0.933 | 202 | 0.027 | 0.145 | 1169.5 | 0.533 | 0.933 |
| 1010 | Gefitinib | ZD-1839, Iressa | 21559 | 0.947 | 1.000 | 9880.5 | 0.992 | 1.000 | 17693 | 0.999 | 1.000 | 23673.5 | 0.994 | 1.000 | 32771.5 | 0.305 | 0.781 | 28761 | 0.001 | 0.009 | 29879.5 | 1.000 | 1.000 | 9058.5 | 0.787 | 1.000 | 12670 | 0.948 | 1.000 | 9581 | 0.060 | 0.240 | 10768 | 0.000 | 0.000 | 20768.5 | 0.979 | 1.000 | 8757.5 | 0.003 | 0.017 |
| 1011 | Navitoclax | ABT-263, ABT263, ABT 263 | 25620 | 1.000 | 1.000 | 8133 | 0.715 | 1.000 | 11302 | 0.037 | 0.098 | 28207 | 1.000 | 1.000 | 37211.5 | 0.928 | 1.000 | 39773 | 0.975 | 1.000 | 8470 | 0.000 | 0.000 | 11030 | 0.996 | 1.000 | 5316 | 0.000 | 0.000 | 9546 | 0.084 | 0.193 | 34778.5 | 1.000 | 1.000 | 13144 | 0.001 | 0.002 | 14109 | 0.927 | 1.000 |
| 1012 | Vorinostat | Zolinza, SAHA, suberanilohydroxamic acid, suberoylanilide hydroxamic acid, MK-0683 | 22771 | 0.991 | 1.000 | 6897 | 0.264 | 0.691 | 10291 | 0.006 | 0.030 | 18891.5 | 0.302 | 0.691 | 39566.5 | 0.997 | 1.000 | 41694.5 | 0.999 | 1.000 | 8985 | 0.000 | 0.000 | 10445 | 0.983 | 1.000 | 3411 | 0.000 | 0.000 | 17018 | 1.000 | 1.000 | 28810 | 0.927 | 1.000 | 19785 | 0.872 | 1.000 | 13583 | 0.866 | 1.000 |
| 1013 | Nilotinib | Tasigna, AMN 107 | 18410.5 | 0.353 | 1.000 | 8122 | 0.737 | 1.000 | 14376 | 0.732 | 1.000 | 12591 | 0.000 | 0.000 | 41292.5 | 1.000 | 1.000 | 39220 | 0.971 | 1.000 | 19520.5 | 0.465 | 1.000 | 8656 | 0.662 | 1.000 | 11370.5 | 0.803 | 1.000 | 12810 | 0.868 | 1.000 | 29819 | 0.980 | 1.000 | 6785 | 0.000 | 0.000 | 8509 | 0.002 | 0.009 |
| 1014 | Refametinib | RDEA119, BAY-86-9766, BAY 869766 | 21045 | 0.952 | 1.000 | 8371 | 0.833 | 1.000 | 20356 | 1.000 | 1.000 | 5557 | 0.000 | 0.000 | 36756 | 0.960 | 1.000 | 30662 | 0.055 | 0.146 | 34644 | 1.000 | 1.000 | 8955 | 0.785 | 1.000 | 10282 | 0.497 | 0.995 | 7751 | 0.006 | 0.035 | 22096 | 0.030 | 0.095 | 23507 | 1.000 | 1.000 | 12469 | 0.634 | 1.000 |
| 1016 | Temsirolimus | CCI-779, Torisel | 11840 | 0.000 | 0.000 | 6313.5 | 0.143 | 0.457 | 10257 | 0.018 | 0.097 | 19850 | 0.731 | 0.998 | 33830 | 0.711 | 0.998 | 28944 | 0.013 | 0.097 | 24536 | 0.998 | 0.998 | 8601 | 0.707 | 0.998 | 13312 | 0.997 | 0.998 | 12241 | 0.967 | 0.998 | 26464.5 | 0.818 | 0.998 | 15043.5 | 0.261 | 0.695 | 9883.5 | 0.050 | 0.200 |
| 1017 | Olaparib | AZD2281, KU0059436, Lynparza | 17048 | 0.060 | 0.160 | 4670 | 0.002 | 0.007 | 6338 | 0.000 | 0.000 | 24548.5 | 0.998 | 1.000 | 34702 | 0.606 | 0.969 | 35595.5 | 0.506 | 0.969 | 13752 | 0.000 | 0.000 | 8211 | 0.464 | 0.969 | 4234 | 0.000 | 0.000 | 13693.5 | 0.959 | 1.000 | 34227.5 | 1.000 | 1.000 | 23713 | 1.000 | 1.000 | 14722 | 0.974 | 1.000 |
| 1018 | Veliparib | ABT-888, ABT888, ABT 888 | 17824.5 | 0.327 | 0.616 | 7054.5 | 0.346 | 0.616 | 13482 | 0.510 | 0.742 | 22676 | 0.988 | 0.988 | 36024.5 | 0.915 | 0.988 | 31068 | 0.060 | 0.358 | 23528.5 | 0.983 | 0.988 | 7011 | 0.130 | 0.415 | 9729 | 0.308 | 0.616 | 7741 | 0.006 | 0.099 | 28911.5 | 0.970 | 0.988 | 17892.5 | 0.599 | 0.799 | 10618 | 0.122 | 0.415 |
| 1019 | Bosutinib | SKI-606, Bosulif | 23153.5 | 0.999 | 1.000 | 9625 | 0.988 | 1.000 | 13727 | 0.584 | 1.000 | 21496 | 0.932 | 1.000 | 32415 | 0.351 | 0.803 | 29222 | 0.007 | 0.040 | 30851 | 1.000 | 1.000 | 8698 | 0.701 | 1.000 | 13481 | 0.997 | 1.000 | 8900 | 0.066 | 0.263 | 14859.5 | 0.000 | 0.000 | 20752 | 0.985 | 1.000 | 6249 | 0.000 | 0.000 |
| 1020 | Lenalidomide | CDC-501, CC-5013, Revlimid | 17809.5 | 0.324 | 0.740 | 7751 | 0.625 | 0.919 | 16095 | 0.978 | 0.989 | 22770 | 0.989 | 0.989 | 33776.5 | 0.610 | 0.919 | 30938.5 | 0.053 | 0.279 | 23115.5 | 0.968 | 0.989 | 6540.5 | 0.056 | 0.279 | 8863.5 | 0.104 | 0.279 | 8345.5 | 0.024 | 0.279 | 26245 | 0.654 | 0.919 | 18509 | 0.747 | 0.919 | 12752.5 | 0.711 | 0.919 |
| 1021 | Axitinib | AG-13736, Inlyta | 16971 | 0.092 | 0.245 | 6593 | 0.173 | 0.396 | 8038 | 0.000 | 0.000 | 28713 | 1.000 | 1.000 | 37870 | 0.980 | 1.000 | 37053 | 0.795 | 1.000 | 11860 | 0.000 | 0.000 | 6672 | 0.063 | 0.245 | 3123 | 0.000 | 0.000 | 15755 | 1.000 | 1.000 | 29183 | 0.952 | 1.000 | 20176 | 0.918 | 1.000 | 13525 | 0.856 | 1.000 |
| 1022 | AZD7762 | AZD-7762, AZD 7762 | 18203 | 0.283 | 0.646 | 4950 | 0.004 | 0.023 | 7773 | 0.000 | 0.000 | 25241 | 1.000 | 1.000 | 33834 | 0.471 | 0.837 | 34431 | 0.325 | 0.649 | 22057 | 0.924 | 1.000 | 9758 | 0.925 | 1.000 | 6313 | 0.000 | 0.001 | 16020 | 1.000 | 1.000 | 22196 | 0.015 | 0.058 | 23049 | 0.999 | 1.000 | 13664 | 0.874 | 1.000 |
| 1023 | GW441756 | GW 441756 | 21313.5 | 0.915 | 1.000 | 6868 | 0.279 | 0.563 | 13715.5 | 0.581 | 0.830 | 23898 | 0.999 | 1.000 | 30210.5 | 0.057 | 0.228 | 29376.5 | 0.009 | 0.137 | 26656.5 | 1.000 | 1.000 | 7621 | 0.298 | 0.563 | 11591 | 0.965 | 1.000 | 8814.5 | 0.017 | 0.137 | 26091.5 | 0.622 | 0.830 | 15962 | 0.352 | 0.563 | 11097 | 0.218 | 0.563 |
| 1024 | Lestaurtinib | CEP-701, SP-924, SPM-924, A-154475, KT-555 | 19059 | 0.646 | 0.999 | 7318 | 0.454 | 0.909 | 9399 | 0.001 | 0.008 | 17772 | 0.190 | 0.506 | 39379 | 0.999 | 0.999 | 35360 | 0.745 | 0.999 | 18256 | 0.122 | 0.488 | 9428 | 0.895 | 0.999 | 3976 | 0.000 | 0.000 | 13779 | 0.996 | 0.999 | 27956 | 0.914 | 0.999 | 19859 | 0.944 | 0.999 | 11094 | 0.221 | 0.506 |
| 1025 | SB216763 | SB-216763, SB 216763 | 17668 | 0.157 | 0.279 | 6332.5 | 0.122 | 0.247 | 17023.5 | 0.996 | 1.000 | 21249 | 0.899 | 1.000 | 31229.5 | 0.123 | 0.247 | 30190.5 | 0.021 | 0.083 | 28752.5 | 1.000 | 1.000 | 5834.5 | 0.011 | 0.057 | 11679.5 | 0.967 | 1.000 | 8047.5 | 0.003 | 0.022 | 26583 | 0.699 | 1.000 | 24622 | 1.000 | 1.000 | 6997 | 0.000 | 0.000 |
| 1026 | Tanespimycin | 17-AAG, BMS-722782 | 13381 | 0.000 | 0.003 | 6820 | 0.262 | 0.628 | 19215 | 1.000 | 1.000 | 17765 | 0.184 | 0.590 | 35375 | 0.855 | 1.000 | 31814 | 0.116 | 0.462 | 28958 | 1.000 | 1.000 | 8547 | 0.647 | 1.000 | 14460 | 1.000 | 1.000 | 9960 | 0.275 | 0.628 | 16807 | 0.000 | 0.000 | 16807 | 0.317 | 0.635 | 12908 | 0.753 | 1.000 |
| 1029 | Motesanib | AMG-706, AMG 706, AMG706 | 17957 | 0.359 | 0.821 | 8003 | 0.719 | 0.980 | 13358 | 0.472 | 0.945 | 22295.5 | 0.977 | 0.980 | 33622.5 | 0.581 | 0.980 | 28542.5 | 0.003 | 0.034 | 23434 | 0.980 | 0.980 | 7110 | 0.151 | 0.541 | 11134 | 0.766 | 0.980 | 7595 | 0.004 | 0.034 | 27784 | 0.894 | 0.980 | 20537.5 | 0.978 | 0.980 | 11590 | 0.352 | 0.821 |
| 1030 | KU-55933 | KU55933 | 20330.5 | 0.760 | 1.000 | 6894 | 0.281 | 0.643 | 15912 | 0.968 | 1.000 | 23371.5 | 0.996 | 1.000 | 30424.5 | 0.064 | 0.209 | 31250 | 0.065 | 0.209 | 25701.5 | 1.000 | 1.000 | 7934.5 | 0.404 | 0.689 | 10887.5 | 0.874 | 1.000 | 11181.5 | 0.431 | 0.689 | 29129 | 0.974 | 1.000 | 13970.5 | 0.014 | 0.112 | 8792 | 0.004 | 0.063 |
| 1031 | Elesclomol | STA-4783 | 1177 | 0.401 | 0.777 | 149 | 0.402 | 0.777 | 424 | 0.552 | 0.883 | 1753 | 0.323 | 0.777 | 986 | 0.287 | 0.777 | 1644 | 0.437 | 0.777 | 1869 | 0.968 | 0.968 | 468 | 0.433 | 0.777 | 311 | 0.067 | 0.537 | 260 | 0.901 | 0.961 | 1438 | 0.657 | 0.956 | 292 | 0.143 | 0.760 | 1268 | 0.735 | 0.961 |
| 1032 | Afatinib | BIBW2992, Tovok, Gilotrif | 26332 | 1.000 | 1.000 | 9177 | 0.949 | 1.000 | 12491 | 0.188 | 0.500 | 27080.5 | 1.000 | 1.000 | 35463.5 | 0.744 | 1.000 | 32306 | 0.068 | 0.361 | 21774 | 0.785 | 1.000 | 10694.5 | 0.990 | 1.000 | 10597 | 0.422 | 0.843 | 12777 | 0.843 | 1.000 | 14847 | 0.000 | 0.000 | 15741 | 0.125 | 0.413 | 10835 | 0.129 | 0.413 |
| 1033 | Vismodegib | GDC0449, Erivedge | 18083 | 0.390 | 0.780 | 7629 | 0.577 | 0.917 | 14565 | 0.803 | 0.991 | 19216 | 0.512 | 0.910 | 38036 | 0.991 | 0.991 | 32842 | 0.240 | 0.549 | 21957 | 0.870 | 0.991 | 6876 | 0.104 | 0.342 | 8082.5 | 0.026 | 0.211 | 9334 | 0.128 | 0.342 | 29186 | 0.979 | 0.991 | 19345.5 | 0.889 | 0.991 | 7760.5 | 0.000 | 0.004 |
| 1034 | Staurosporine | | 13061 | 0.000 | 0.000 | 6170 | 0.040 | 0.107 | 9902 | 0.002 | 0.012 | 25019 | 0.999 | 0.999 | 37798 | 0.957 | 0.999 | 38284 | 0.893 | 0.999 | 24694 | 0.994 | 0.999 | 9125 | 0.782 | 0.999 | 10446 | 0.364 | 0.831 | 15484 | 0.999 | 0.999 | 22892 | 0.021 | 0.070 | 19385 | 0.780 | 0.999 | 7732 | 0.000 | 0.003 |
| 1036 | PLX-4720 | PLX4720, PLX 4720 | 15303.5 | 0.008 | 0.045 | 7163 | 0.359 | 0.820 | 14156 | 0.757 | 1.000 | 6808 | 0.000 | 0.000 | 33874.5 | 0.607 | 0.971 | 35322.5 | 0.505 | 0.971 | 29923 | 1.000 | 1.000 | 7343 | 0.190 | 0.607 | 13577 | 0.991 | 1.000 | 13320 | 0.934 | 1.000 | 26823 | 0.581 | 0.971 | 22485.5 | 0.999 | 1.000 | 8985 | 0.005 | 0.042 |
| 1037 | BX795 | BX-795 | 937 | 0.062 | 0.330 | 114 | 0.237 | 0.542 | 163 | 0.009 | 0.150 | 2620 | 1.000 | 1.000 | 1129 | 0.600 | 0.877 | 1480 | 0.181 | 0.484 | 1543 | 0.603 | 0.877 | 422 | 0.290 | 0.581 | 262 | 0.026 | 0.211 | 276 | 0.939 | 1.000 | 1561 | 0.849 | 0.970 | 492 | 0.772 | 0.950 | 987 | 0.177 | 0.484 |
| 1038 | NU7441 | KU-57788, NU-7432, NU-7741 | 17139 | 0.208 | 0.665 | 8106 | 0.641 | 1.000 | 15293 | 0.932 | 1.000 | 19843 | 0.769 | 1.000 | 32528 | 0.527 | 0.938 | 32609.5 | 0.285 | 0.760 | 25029 | 1.000 | 1.000 | 9157 | 0.850 | 1.000 | 11414 | 0.915 | 1.000 | 14250 | 0.998 | 1.000 | 25204 | 0.469 | 0.937 | 13739 | 0.007 | 0.055 | 8046 | 0.001 | 0.011 |
| 1039 | SL0101 | SL-0101, SL 0101-1 | 924 | 0.054 | 0.431 | 167 | 0.503 | 0.880 | 440 | 0.608 | 0.880 | 1613 | 0.142 | 0.556 | 1048 | 0.418 | 0.880 | 1723 | 0.581 | 0.880 | 1978 | 0.991 | 0.991 | 557 | 0.715 | 0.880 | 518 | 0.597 | 0.880 | 130 | 0.307 | 0.818 | 1449 | 0.678 | 0.880 | 306 | 0.174 | 0.556 | 963 | 0.145 | 0.556 |
| 1042 | Doramapimod | BIRB-796, BIRB 796 | 19084.5 | 0.465 | 0.996 | 7795 | 0.634 | 0.996 | 14364.5 | 0.748 | 0.998 | 16305 | 0.031 | 0.251 | 39812 | 0.999 | 0.999 | 31483.5 | 0.081 | 0.322 | 22521.5 | 0.951 | 0.999 | 7059.5 | 0.136 | 0.435 | 11005 | 0.894 | 0.999 | 11579 | 0.562 | 0.996 | 26474 | 0.685 | 0.996 | 14832.5 | 0.053 | 0.283 | 6779 | 0.000 | 0.000 |
| 1043 | JNK Inhibitor VIII | | 17406.5 | 0.130 | 0.346 | 8249 | 0.797 | 1.000 | 17674 | 0.999 | 1.000 | 18467 | 0.327 | 0.653 | 35158 | 0.792 | 1.000 | 29428 | 0.010 | 0.077 | 26382 | 1.000 | 1.000 | 6648 | 0.069 | 0.221 | 10585 | 0.815 | 1.000 | 9393 | 0.051 | 0.221 | 22708 | 0.059 | 0.221 | 19478.5 | 0.979 | 1.000 | 7861 | 0.000 | 0.006 |
| 1046 | Wee1 Inhibitor | 681640, Wee1 Inhibitor | 18496 | 0.374 | 0.919 | 7325.5 | 0.430 | 0.919 | 12963 | 0.326 | 0.919 | 20145 | 0.619 | 0.961 | 34125 | 0.565 | 0.961 | 39371 | 0.975 | 0.999 | 18196.5 | 0.236 | 0.919 | 11014 | 0.997 | 0.999 | 4557 | 0.000 | 0.000 | 15089 | 0.999 | 0.999 | 21931 | 0.012 | 0.096 | 17511 | 0.460 | 0.919 | 11153 | 0.210 | 0.919 |
| 1047 | Nutlin-3a (-) | | 18217 | 0.202 | 0.538 | 5324 | 0.011 | 0.037 | 13119 | 0.329 | 0.753 | 13516 | 0.000 | 0.000 | 34336 | 0.528 | 0.768 | 35763 | 0.527 | 0.768 | 29788 | 1.000 | 1.000 | 5451 | 0.003 | 0.012 | 5894 | 0.000 | 0.000 | 13411.5 | 0.933 | 1.000 | 33763 | 1.000 | 1.000 | 22574 | 0.998 | 1.000 | 7475 | 0.000 | 0.000 |
| 1048 | Mirin | | 17708 | 0.796 | 0.979 | 5503 | 0.194 | 0.769 | 8482 | 0.005 | 0.037 | 15970 | 0.266 | 0.769 | 28456 | 0.544 | 0.979 | 33282 | 0.967 | 0.999 | 17007 | 0.782 | 0.979 | 10150 | 0.999 | 0.999 | 3925 | 0.000 | 0.000 | 12475 | 0.999 | 0.999 | 19476 | 0.016 | 0.083 | 15872 | 0.741 | 0.979 | 11445 | 0.691 | 0.979 |
| 1049 | PD173074 | PD-173074, PD 173074 | 18523 | 0.265 | 0.712 | 7222.5 | 0.363 | 0.712 | 11038 | 0.023 | 0.093 | 19449.5 | 0.400 | 0.712 | 34480 | 0.565 | 0.903 | 36978 | 0.745 | 1.000 | 24275 | 0.989 | 1.000 | 4527 | 0.000 | 0.002 | 11828.5 | 0.796 | 1.000 | 14377 | 0.989 | 1.000 | 25994.5 | 0.365 | 0.712 | 24305 | 1.000 | 1.000 | 8200.5 | 0.001 | 0.005 |
| 1050 | ZM447439 | ZM-447439, ZM 447439 | 16646 | 0.131 | 0.419 | 8174 | 0.670 | 0.946 | 11315 | 0.060 | 0.287 | 17337.5 | 0.191 | 0.509 | 38362 | 0.999 | 0.999 | 34871.5 | 0.709 | 0.946 | 16249 | 0.024 | 0.195 | 11155 | 0.999 | 0.999 | 4414.5 | 0.000 | 0.000 | 14755 | 0.999 | 0.999 | 22690.5 | 0.072 | 0.287 | 20147 | 0.969 | 0.999 | 11905 | 0.594 | 0.946 |
| 1051 | Alisertib | MLN8237 | 15098 | 0.006 | 0.038 | 6227 | 0.097 | 0.383 | 10330 | 0.007 | 0.038 | 27560 | 1.000 | 1.000 | 34342.5 | 0.606 | 1.000 | 32932.5 | 0.144 | 0.383 | 17487 | 0.121 | 0.383 | 9328 | 0.858 | 1.000 | 2422 | 0.000 | 0.000 | 14701 | 0.996 | 1.000 | 25419 | 0.302 | 0.605 | 24280 | 1.000 | 1.000 | 10956 | 0.168 | 0.383 |
| 1052 | RO-3306 | | 17436 | 0.127 | 0.290 | 7837 | 0.650 | 1.000 | 11615 | 0.077 | 0.247 | 22539 | 0.983 | 1.000 | 39645.5 | 0.998 | 1.000 | 36640 | 0.855 | 1.000 | 16493.5 | 0.017 | 0.090 | 10405 | 0.985 | 1.000 | 4637 | 0.000 | 0.000 | 15823 | 1.000 | 1.000 | 23520.5 | 0.125 | 0.290 | 13941 | 0.013 | 0.090 | 10310 | 0.076 | 0.247 |
| 1053 | MK-2206 | MK 2206, MK2206 | 19928 | 0.594 | 1.000 | 7215 | 0.357 | 0.951 | 10999 | 0.021 | 0.113 | 22232.5 | 0.928 | 1.000 | 37510 | 0.943 | 1.000 | 36466 | 0.654 | 1.000 | 21027.5 | 0.615 | 1.000 | 9490 | 0.871 | 1.000 | 11258 | 0.631 | 1.000 | 16023 | 1.000 | 1.000 | 28152.5 | 0.786 | 1.000 | 13784.5 | 0.002 | 0.016 | 7309 | 0.000 | 0.001 |
| 1054 | Palbociclib | PD0332991, PD-0332991, PF-00080665-73 | 20283 | 0.678 | 0.904 | 6542 | 0.148 | 0.321 | 10565 | 0.009 | 0.037 | 17751 | 0.092 | 0.246 | 39190 | 0.991 | 1.000 | 35662.5 | 0.509 | 0.814 | 30040 | 1.000 | 1.000 | 8248 | 0.474 | 0.814 | 7989 | 0.008 | 0.037 | 11962 | 0.623 | 0.904 | 30134.5 | 0.968 | 1.000 | 22011 | 0.993 | 1.000 | 7908 | 0.000 | 0.002 |
| 1057 | Dactolisib | NVP-BEZ235, BEZ235 | 12299 | 0.000 | 0.000 | 5423 | 0.017 | 0.068 | 9744 | 0.002 | 0.010 | 18251 | 0.183 | 0.488 | 36573 | 0.905 | 1.000 | 35379.5 | 0.525 | 0.934 | 26774 | 1.000 | 1.000 | 8066 | 0.433 | 0.867 | 13699 | 0.994 | 1.000 | 12689 | 0.842 | 1.000 | 28540.5 | 0.909 | 1.000 | 17354 | 0.412 | 0.867 | 8452 | 0.001 | 0.010 |
| 1058 | Pictilisib | GDC-0941, GDC0941, RG-7621 | 19034 | 0.380 | 0.760 | 6064 | 0.064 | 0.240 | 9308 | 0.001 | 0.008 | 25659 | 1.000 | 1.000 | 34748 | 0.614 | 0.983 | 35234 | 0.439 | 0.781 | 27203 | 1.000 | 1.000 | 10924 | 0.994 | 1.000 | 11731 | 0.772 | 1.000 | 12561 | 0.793 | 1.000 | 23823 | 0.065 | 0.240 | 14556 | 0.009 | 0.074 | 10459 | 0.075 | 0.240 |
| 1059 | AZD8055 | AZD-8055 | 7541.5 | 0.002 | 0.025 | 3666 | 0.012 | 0.063 | 7281 | 0.011 | 0.063 | 7777 | 0.103 | 0.329 | 22235 | 0.411 | 0.738 | 21320.5 | 0.461 | 0.738 | 14438.5 | 1.000 | 1.000 | 4915 | 0.588 | 0.856 | 8265 | 0.965 | 1.000 | 9191 | 0.870 | 1.000 | 19012.5 | 0.994 | 1.000 | 13265 | 0.417 | 0.738 | 4178 | 0.058 | 0.233 |
| 1060 | PD0325901 | PD-0325901, PD 0325901 | 24020 | 0.998 | 1.000 | 9507 | 0.938 | 1.000 | 21480 | 1.000 | 1.000 | 4333 | 0.000 | 0.000 | 37003 | 0.909 | 1.000 | 32380.5 | 0.066 | 0.177 | 34188 | 1.000 | 1.000 | 9367 | 0.844 | 1.000 | 10593 | 0.411 | 0.822 | 7067 | 0.000 | 0.001 | 23020 | 0.025 | 0.079 | 26229.5 | 1.000 | 1.000 | 12490 | 0.562 | 1.000 |
| 1061 | SB590885 | SB-590885 | 21995 | 0.992 | 1.000 | 7446 | 0.384 | 0.878 | 15590.5 | 0.959 | 1.000 | 7663 | 0.000 | 0.000 | 34247 | 0.827 | 1.000 | 34796.5 | 0.697 | 1.000 | 25410 | 1.000 | 1.000 | 9241 | 0.871 | 1.000 | 10609.5 | 0.747 | 1.000 | 8392 | 0.016 | 0.084 | 30446 | 0.998 | 1.000 | 17708.5 | 0.676 | 1.000 | 10636.5 | 0.141 | 0.375 |
| 1062 | Selumetinib | AZD6244, AZD-6244, ARRY-886 | 21919 | 0.987 | 1.000 | 8935 | 0.938 | 1.000 | 20447 | 1.000 | 1.000 | 5191 | 0.000 | 0.000 | 35869 | 0.903 | 1.000 | 32794.5 | 0.233 | 0.621 | 32233 | 1.000 | 1.000 | 8741 | 0.715 | 1.000 | 10036 | 0.408 | 0.763 | 6961 | 0.001 | 0.004 | 22760 | 0.062 | 0.200 | 23360 | 1.000 | 1.000 | 11374 | 0.290 | 0.662 |
| 1067 | CCT007093 | | 16105 | 0.165 | 0.590 | 9189 | 0.939 | 1.000 | 15853.5 | 0.990 | 1.000 | 17663.5 | 0.286 | 0.694 | 33071.5 | 0.696 | 1.000 | 28821 | 0.012 | 0.099 | 25423.5 | 1.000 | 1.000 | 6515 | 0.066 | 0.350 | 11026 | 0.868 | 1.000 | 9781.5 | 0.184 | 0.590 | 27032.5 | 0.864 | 1.000 | 15826.5 | 0.304 | 0.694 | 7683 | 0.000 | 0.005 |
| 1068 | Obatoclax Mesyla | GX15-070MS, Obatoclax, GX15-070 | 10500 | 0.177 | 0.354 | 5409 | 0.382 | 0.556 | 7810 | 0.007 | 0.059 | 6422 | 0.000 | 0.001 | 22410 | 0.077 | 0.245 | 31019.5 | 1.000 | 1.000 | 15211 | 0.985 | 1.000 | 8274 | 0.993 | 1.000 | 5832 | 0.066 | 0.245 | 12674.5 | 1.000 | 1.000 | 15240 | 0.046 | 0.245 | 16312 | 0.995 | 1.000 | 6097 | 0.294 | 0.471 |
| 1069 | EHT-1864 | EHT 1864 | 22075.5 | 0.993 | 1.000 | 8772.5 | 0.852 | 1.000 | 14374 | 0.787 | 1.000 | 23861 | 1.000 | 1.000 | 27278 | 0.006 | 0.061 | 33698 | 0.493 | 0.877 | 19188 | 0.463 | 0.877 | 8702 | 0.726 | 1.000 | 12673 | 0.994 | 1.000 | 9814 | 0.173 | 0.555 | 26297 | 0.710 | 1.000 | 13481 | 0.008 | 0.061 | 11064 | 0.233 | 0.621 |
| 1072 | Avagacestat | BMS-708163, BMS 708163 | 20310 | 0.868 | 0.996 | 9384.5 | 0.939 | 0.996 | 16716 | 0.993 | 0.996 | 16884 | 0.066 | 0.211 | 35157 | 0.809 | 0.996 | 34597 | 0.523 | 0.930 | 24462.5 | 0.996 | 0.996 | 6379 | 0.038 | 0.183 | 10810 | 0.657 | 0.955 | 8420 | 0.013 | 0.183 | 22585 | 0.046 | 0.183 | 17997 | 0.607 | 0.955 | 9942 | 0.041 | 0.183 |
| 1073 | 5-Fluorouracil | 5-FU | 21939 | 0.933 | 1.000 | 6352 | 0.108 | 0.346 | 11828 | 0.079 | 0.318 | 22180 | 0.924 | 1.000 | 34162 | 0.495 | 0.990 | 37253.5 | 0.777 | 1.000 | 26538 | 1.000 | 1.000 | 9275 | 0.822 | 1.000 | 7639 | 0.003 | 0.018 | 12130 | 0.674 | 1.000 | 20745 | 0.001 | 0.006 | 23080 | 0.999 | 1.000 | 10972 | 0.148 | 0.395 |
| 1079 | Dasatinib | BMS-354825-03, BMS-354825, Sprycel | 14879 | 0.002 | 0.010 | 7372 | 0.429 | 0.686 | 16275.5 | 0.976 | 1.000 | 27994 | 1.000 | 1.000 | 28119.5 | 0.002 | 0.010 | 34109 | 0.265 | 0.529 | 34781.5 | 1.000 | 1.000 | 6987 | 0.105 | 0.241 | 14514 | 1.000 | 1.000 | 9535 | 0.051 | 0.136 | 14785.5 | 0.000 | 0.000 | 22623 | 0.999 | 1.000 | 3951 | 0.000 | 0.000 |
| 1080 | Paclitaxel | BMS-181339-01, Taxol, Onxol, Paxene, Praxel, Abraxane | 17415 | 0.152 | 0.498 | 7561 | 0.521 | 0.990 | 13821 | 0.573 | 0.990 | 18273.5 | 0.187 | 0.498 | 33456.5 | 0.428 | 0.979 | 39474 | 0.976 | 0.996 | 18170 | 0.170 | 0.498 | 10729 | 0.992 | 0.996 | 11838 | 0.821 | 0.996 | 7796 | 0.001 | 0.010 | 23217 | 0.058 | 0.309 | 11731 | 0.000 | 0.001 | 12555 | 0.619 | 0.990 |
| 1083 | Crizotinib | Xalkori, PF2341066, PF-2341066, PF 2341066 | 20184 | 0.662 | 1.000 | 6031.5 | 0.060 | 0.193 | 7650 | 0.000 | 0.000 | 20828 | 0.728 | 1.000 | 44762.5 | 1.000 | 1.000 | 34999 | 0.397 | 0.906 | 15818 | 0.002 | 0.008 | 8725 | 0.657 | 1.000 | 3845 | 0.000 | 0.000 | 17322 | 1.000 | 1.000 | 27808 | 0.736 | 1.000 | 23718 | 1.000 | 1.000 | 13980 | 0.912 | 1.000 |
| 1084 | Rapamycin | AY-22989, Sirolimus, WY-090217, Torisel, Rapamune | 13210 | 0.000 | 0.001 | 6691 | 0.216 | 0.575 | 12872 | 0.418 | 0.956 | 18038 | 0.169 | 0.575 | 34076 | 0.594 | 0.999 | 32908 | 0.164 | 0.575 | 21526 | 0.894 | 0.999 | 8296 | 0.544 | 0.999 | 12001 | 0.964 | 0.999 | 15268 | 0.999 | 0.999 | 28276 | 0.903 | 0.999 | 16197.5 | 0.181 | 0.575 | 8089 | 0.001 | 0.005 |
| 1085 | Sorafenib | Nexavar, 284461-73-0, BAY 43-9006 | 21956 | 0.969 | 0.972 | 6791.5 | 0.233 | 0.532 | 14223.5 | 0.687 | 0.972 | 18096 | 0.158 | 0.507 | 37424.5 | 0.957 | 0.972 | 38691 | 0.947 | 0.972 | 17064.5 | 0.050 | 0.264 | 9281 | 0.844 | 0.972 | 5925.5 | 0.000 | 0.000 | 10481 | 0.208 | 0.532 | 29608 | 0.972 | 0.972 | 13541.5 | 0.003 | 0.024 | 12899 | 0.718 | 0.972 |
| 1086 | BI-2536 | | 16833 | 0.219 | 0.438 | 4820 | 0.005 | 0.029 | 9403 | 0.004 | 0.029 | 23433 | 1.000 | 1.000 | 31350 | 0.386 | 0.686 | 35541 | 0.894 | 1.000 | 16299 | 0.091 | 0.234 | 10754 | 0.999 | 1.000 | 4167 | 0.000 | 0.000 | 14041 | 0.999 | 1.000 | 21280 | 0.089 | 0.234 | 13285 | 0.096 | 0.234 | 15631 | 0.999 | 1.000 |
| 1088 | Irinotecan | Camptosar, (+)-Irinotecan, Irinotecanum, irinotecan hydrochloride | 16749 | 0.065 | 0.208 | 4627 | 0.002 | 0.008 | 7283 | 0.000 | 0.000 | 21958.5 | 0.911 | 1.000 | 32454 | 0.225 | 0.601 | 39199 | 0.960 | 1.000 | 15673 | 0.003 | 0.011 | 8678 | 0.649 | 1.000 | 5133 | 0.000 | 0.000 | 14759 | 0.996 | 1.000 | 27157 | 0.626 | 1.000 | 25600.5 | 1.000 | 1.000 | 12276 | 0.511 | 1.000 |
| 1089 | Oxaliplatin | Eloxatin | 18073 | 0.181 | 0.581 | 6947 | 0.265 | 0.658 | 13929 | 0.571 | 1.000 | 22474 | 0.949 | 1.000 | 35677.5 | 0.768 | 1.000 | 40807.5 | 0.993 | 1.000 | 16132 | 0.004 | 0.019 | 9794 | 0.924 | 1.000 | 7194 | 0.001 | 0.008 | 10877.5 | 0.288 | 0.658 | 28297 | 0.814 | 1.000 | 23994.5 | 1.000 | 1.000 | 9521 | 0.015 | 0.058 |
| 1091 | BMS-536924 | BMS 536924 | 21188 | 0.916 | 1.000 | 6581 | 0.172 | 0.307 | 8914 | 0.000 | 0.003 | 26159 | 1.000 | 1.000 | 37253 | 0.949 | 1.000 | 29454 | 0.003 | 0.017 | 28110 | 1.000 | 1.000 | 6512 | 0.047 | 0.107 | 8481 | 0.026 | 0.071 | 8939 | 0.019 | 0.064 | 21047 | 0.003 | 0.017 | 24983 | 1.000 | 1.000 | 9604 | 0.020 | 0.064 |

| DRUG_ID | DRUG_NAME | SYNONYMS | large_intestine_MWU_STAT | large_intestine_PVAL | large_intestine_PVAL_ADJ | thyroid_MWU_STAT | thyroid_PVAL | thyroid_PVAL_ADJ | digestive_system_MWU_STAT | digestive_system_PVAL | digestive_system_PVAL_ADJ | digestive_system_PVAL2 |
|---|---|---|---|---|---|---|---|---|---|---|---|---|
| 1003 | Camptothecin | Camptothecine, (+)-Camptothecin | 18985.5 | 0.956 | 1.000 | 3983 | 0.022 | 0.070 | 20110 | 0.975 | 1.000 | 1.000 |
| 1004 | Vinblastine | Velban | 10848 | 0.311 | 0.710 | 4335 | 0.603 | 0.961 | 12052 | 0.458 | 0.890 | 0.890 |
| 1005 | Cisplatin | cis-Diammineplatinum(II) dichloride, Platinol, CIS-DDP | 13572 | 0.924 | 1.000 | 4888 | 0.820 | 1.000 | 14889.5 | 0.981 | 1.000 | 1.000 |
| 1006 | Cytarabine | Ara-Cytidine, Arabinosyl Cytosine, U-19920 | 10188 | 0.127 | 0.403 | 4404 | 0.635 | 0.899 | 12884 | 0.727 | 0.899 | 0.899 |
| 1007 | Docetaxel | RP-56976, Taxotere | 18132 | 0.917 | 1.000 | 5450 | 0.379 | 1.000 | 18125.5 | 0.742 | 1.000 | 1.000 |
| 1008 | Methotrexate | Abitrexate, Amethopterin, Rheumatrex, Trexall, Folex | 11191 | 0.002 | 0.019 | 4096 | 0.062 | 0.199 | 13880 | 0.202 | 0.379 | 0.379 |
| 1009 | Tretinoin | ATRA, Vesanoid, Renova, Atralin, Tretin-X, Avita | 693 | 0.996 | 0.996 | 8 | 0.061 | 0.243 | 496 | 0.789 | 0.944 | 0.944 |
| 1010 | Gefitinib | ZD-1839, Iressa | 14567 | 0.222 | 0.712 | 6548 | 0.848 | 1.000 | 16409.5 | 0.342 | 0.781 | 0.781 |
| 1011 | Navitoclax | ABT-263, ABT263, ABT 263 | 10995 | 0.000 | 0.000 | 5778 | 0.529 | 1.000 | 14229 | 0.025 | 0.080 | 0.080 |
| 1012 | Vorinostat | Zolinza, SAHA, suberanilohydroxamic acid, suberoylanilide hydroxamic acid, MK-0683 | 12219 | 0.008 | 0.030 | 5368 | 0.359 | 0.718 | 14332.5 | 0.034 | 0.109 | 0.109 |
| 1013 | Nilotinib | Tasigna, AMN 107 | 17331 | 0.885 | 1.000 | 6307 | 0.775 | 1.000 | 17484 | 0.631 | 1.000 | 1.000 |
| 1014 | Refametinib | RDEA119, BAY-86-9766, BAY 869766 | 7314 | 0.000 | 0.000 | 3681 | 0.011 | 0.045 | 14339 | 0.069 | 0.157 | 0.157 |
| 1016 | Temsirolimus | CCI-779, Torisel | 18724.5 | 0.993 | 0.998 | 5729 | 0.589 | 0.998 | 15810.5 | 0.371 | 0.847 | 0.847 |
| 1017 | Olaparib | AZD2281, KU0059436, Lynparza | 20756 | 0.998 | 1.000 | 4126 | 0.032 | 0.104 | 17324 | 0.545 | 0.969 | 0.969 |
| 1018 | Veliparib | ABT-888, ABT888, ABT 888 | 15428 | 0.386 | 0.618 | 5116 | 0.277 | 0.616 | 14340 | 0.067 | 0.358 | 0.358 |
| 1019 | Bosutinib | SKI-606, Bosulif | 14144 | 0.116 | 0.373 | 5481 | 0.436 | 0.872 | 15391 | 0.220 | 0.586 | 0.586 |
| 1020 | Lenalidomide | CDC-501, CC-5013, Revlimid | 15801.5 | 0.489 | 0.919 | 4452 | 0.084 | 0.279 | 14679.5 | 0.103 | 0.279 | 0.279 |
| 1021 | Axitinib | AG-13736, Inlyta | 17368 | 0.827 | 1.000 | 6101 | 0.690 | 1.000 | 15068 | 0.092 | 0.245 | 0.245 |
| 1022 | AZD7762 | AZD-7762, AZD 7762 | 17735 | 0.820 | 1.000 | 5162 | 0.268 | 0.646 | 15140 | 0.095 | 0.305 | 0.305 |
| 1023 | GW441756 | GW 441756 | 15162.5 | 0.317 | 0.563 | 4279.5 | 0.057 | 0.228 | 15727.5 | 0.222 | 0.563 | 0.563 |
| 1024 | Lestaurtinib | CEP-701, SP-924, SPM-924, A-154475, KT-555 | 13401 | 0.044 | 0.236 | 4836 | 0.180 | 0.506 | 16913 | 0.615 | 0.999 | 0.999 |
| 1025 | SB216763 | SB-216763, SB 216763 | 15210.5 | 0.237 | 0.379 | 4083 | 0.033 | 0.107 | 15040.5 | 0.101 | 0.247 | 0.247 |
| 1026 | Tanespimycin | 17-AAG, BMS-722782 | 12589 | 0.011 | 0.059 | 5525 | 0.457 | 0.812 | 17514 | 0.755 | 1.000 | 1.000 |
| 1029 | Motesanib | AMG-706, AMG 706, AMG706 | 16879 | 0.768 | 0.980 | 4806 | 0.169 | 0.541 | 13797 | 0.031 | 0.163 | 0.163 |
| 1030 | KU-55933 | KU55933 | 14294.5 | 0.132 | 0.353 | 5244 | 0.324 | 0.647 | 14503.5 | 0.052 | 0.209 | 0.209 |
| 1031 | Elesclomol | STA-4783 | 537 | 0.877 | 0.961 | 131 | 0.781 | 0.961 | 180 | 0.015 | 0.241 | 0.241 |
| 1032 | Afatinib | BIBW2992, Tovok, Gilotrif | 13603 | 0.026 | 0.205 | 7197 | 0.958 | 1.000 | 16017.5 | 0.227 | 0.520 | 0.520 |
| 1033 | Vismodegib | GDC0449, Erivedge | 16313 | 0.631 | 0.917 | 4476 | 0.089 | 0.342 | 14743.5 | 0.111 | 0.342 | 0.342 |
| 1034 | Staurosporine | | 16391 | 0.472 | 0.945 | 3986 | 0.022 | 0.070 | 18032 | 0.716 | 0.999 | 0.999 |
| 1036 | PLX-4720 | PLX4720, PLX 4720 | 18831 | 0.956 | 1.000 | 4109 | 0.033 | 0.133 | 16111 | 0.266 | 0.709 | 0.709 |
| 1037 | BX795 | BX-795 | 485 | 0.753 | 0.950 | 29 | 0.178 | 0.484 | 387 | 0.420 | 0.746 | 0.746 |
| 1038 | NU7441 | KU-57788, NU-7432, NU-7741 | 13735.5 | 0.079 | 0.316 | 5500 | 0.462 | 0.937 | 13489 | 0.022 | 0.116 | 0.116 |
| 1039 | SL0101 | SL-0101, SL 0101-1 | 558 | 0.913 | 0.974 | 0 | 0.006 | 0.095 | 513 | 0.826 | 0.944 | 0.944 |
| 1042 | Doramapimod | BIRB-796, BIRB 796 | 18187 | 0.947 | 0.999 | 5351 | 0.370 | 0.987 | 17188 | 0.579 | 0.996 | 0.996 |
| 1043 | JNK Inhibitor VIII | | 17406 | 0.865 | 1.000 | 5112 | 0.275 | 0.629 | 16429 | 0.387 | 0.688 | 0.688 |
| 1046 | Wee1 Inhibitor | 681640, Wee1 Inhibitor | 16375 | 0.700 | 0.961 | 6524 | 0.844 | 0.999 | 17854 | 0.721 | 0.961 | 0.961 |
| 1047 | Nutlin-3a (-) | | 17066 | 0.653 | 0.870 | 5672 | 0.477 | 0.768 | 17019.5 | 0.458 | 0.768 | 0.768 |
| 1048 | Mirin | | 15336 | 0.597 | 0.979 | 4231 | 0.336 | 0.769 | 13831 | 0.336 | 0.769 | 0.769 |
| 1049 | PD173074 | PD-173074, PD 173074 | 17733.5 | 0.867 | 1.000 | 5379 | 0.348 | 0.712 | 13345 | 0.005 | 0.028 | 0.028 |
| 1050 | ZM447439 | ZM-447439, ZM 447439 | 14659 | 0.228 | 0.522 | 5533 | 0.481 | 0.856 | 16142 | 0.436 | 0.856 | 0.856 |
| 1051 | Alisertib | MLN8237 | 17347 | 0.763 | 1.000 | 6212 | 0.740 | 1.000 | 17949 | 0.742 | 1.000 | 1.000 |
| 1052 | RO-3306 | | 15104 | 0.292 | 0.584 | 6360 | 0.805 | 1.000 | 17514.5 | 0.663 | 1.000 | 1.000 |
| 1053 | MK-2206 | MK 2206, MK2206 | 15108 | 0.172 | 0.551 | 7032 | 0.936 | 1.000 | 14854 | 0.060 | 0.239 | 0.239 |
| 1054 | Palbociclib | PD0332991, PD-0332991, PF-00080665-73 | 10400.5 | 0.000 | 0.000 | 4866 | 0.160 | 0.321 | 14836.5 | 0.058 | 0.187 | 0.187 |
| 1057 | Dactolisib | NVP-BEZ235, BEZ235 | 18298 | 0.969 | 1.000 | 6424 | 0.811 | 1.000 | 15380 | 0.135 | 0.431 | 0.431 |
| 1058 | Pictilisib | GDC-0941, GDC0941, RG-7621 | 14383 | 0.114 | 0.261 | 6673 | 0.867 | 1.000 | 15296 | 0.106 | 0.261 | 0.261 |
| 1059 | AZD8055 | AZD-8055 | 14128.5 | 0.929 | 1.000 | 4113 | 0.460 | 0.738 | 10209.5 | 0.131 | 0.350 | 0.350 |
| 1060 | PD0325901 | PD-0325901, PD 0325901 | 7269 | 0.000 | 0.000 | 3729 | 0.010 | 0.042 | 15742 | 0.168 | 0.384 | 0.384 |
| 1061 | SB590885 | SB-590885 | 12121.5 | 0.005 | 0.044 | 4116 | 0.041 | 0.148 | 13953 | 0.046 | 0.148 | 0.148 |
| 1062 | Selumetinib | AZD6244, AZD-6244, ARRY-886 | 7558 | 0.000 | 0.000 | 4011 | 0.029 | 0.115 | 16255.5 | 0.429 | 0.763 | 0.763 |
| 1067 | CCT007093 | | 15249 | 0.502 | 0.803 | 4849 | 0.347 | 0.694 | 15996.5 | 0.433 | 0.770 | 0.770 |
| 1068 | Obatoclax Mesyla | GX15-070MS, Obatoclax, GX15-070 | 11966 | 0.146 | 0.334 | 3532 | 0.100 | 0.266 | 12434 | 0.255 | 0.453 | 0.453 |
| 1069 | EHT-1864 | EHT 1864 | 14060 | 0.121 | 0.485 | 5153 | 0.309 | 0.706 | 13881 | 0.042 | 0.222 | 0.222 |
| 1072 | Avagacestat | BMS-708163, BMS 708163 | 15142 | 0.296 | 0.765 | 5386 | 0.383 | 0.765 | 16473 | 0.381 | 0.765 | 0.765 |
| 1073 | 5-Fluorouracil | 5-FU | 11069.5 | 0.000 | 0.002 | 6744 | 0.882 | 1.000 | 16482 | 0.321 | 0.733 | 0.733 |
| 1079 | Dasatinib | BMS-354825-03, BMS-354825, Sprycel | 19979.5 | 0.997 | 1.000 | 3622 | 0.008 | 0.025 | 16654.5 | 0.380 | 0.676 | 0.676 |
| 1080 | Paclitaxel | BMS-181339-01, Taxol, Onxol, Paxene, Praxel, Abraxane | 19411 | 0.996 | 0.996 | 7699 | 0.991 | 0.996 | 20199 | 0.984 | 0.996 | 0.996 |
| 1083 | Crizotinib | Xalkori, PF2341066, PF-2341066, PF 2341066 | 9521 | 0.000 | 0.000 | 6380 | 0.780 | 1.000 | 16089 | 0.237 | 0.633 | 0.633 |
| 1084 | Rapamycin | AY-22989, Sirolimus, WY-090217, Torisel, Rapamune | 18105 | 0.981 | 0.999 | 6584 | 0.946 | 0.999 | 17508.5 | 0.743 | 0.999 | 0.999 |
| 1085 | Sorafenib | Nexavar, 284461-73-0, BAY 43-9006 | 15726 | 0.521 | 0.972 | 6215 | 0.738 | 0.972 | 15422 | 0.141 | 0.507 | 0.507 |
| 1086 | BI-2536 | | 13336 | 0.102 | 0.234 | 6779 | 0.944 | 1.000 | 17555 | 0.856 | 1.000 | 1.000 |
| 1088 | Irinotecan | Camptosar, (+)-Irinotecan, Irinotecanum, irinotecan hydrochloride | 20009 | 0.993 | 1.000 | 5443 | 0.383 | 0.874 | 19220 | 0.922 | 1.000 | 1.000 |
| 1089 | Oxaliplatin | Eloxatin | 8863 | 0.000 | 0.000 | 7522 | 0.982 | 1.000 | 17558 | 0.607 | 1.000 | 1.000 |
| 1091 | BMS-536924 | BMS 536924 | 13486 | 0.064 | 0.127 | 4975 | 0.208 | 0.333 | 17660 | 0.669 | 0.973 | 0.973 |

Suppl. Table 8

| DRUG_ID | DRUG_NAME | SYNONYMS | nervous_system_MWU_STAT | nervous_system_PVAL | nervous_system_PVAL_ADJ | soft_tissue_MWU_STAT | soft_tissue_PVAL | soft_tissue_PVAL_ADJ | bone_MWU_STAT | bone_PVAL | bone_PVAL_ADJ | skin_MWU_STAT | skin_PVAL | skin_PVAL_ADJ | urogenital_system_MWU_STAT | urogenital_system_PVAL | urogenital_system_PVAL_ADJ | lung_NSCLC_MWU_STAT | lung_NSCLC_PVAL | lung_NSCLC_PVAL_ADJ | lung_SCLC_MWU_STAT | lung_SCLC_PVAL | lung_SCLC_PVAL_ADJ | lung_MWU_STAT | lung_PVAL | lung_PVAL_ADJ | neuroblastoma_MWU_STAT | neuroblastoma_PVAL | neuroblastoma_PVAL_ADJ | pancreas_MWU_STAT | pancreas_PVAL | pancreas_PVAL_ADJ | aero_dig_tract_MWU_STAT | aero_dig_tract_PVAL | aero_dig_tract_PVAL_ADJ | breast_MWU_STAT | breast_PVAL | breast_PVAL_ADJ | kidney_MWU_STAT | kidney_PVAL | kidney_PVAL_ADJ |
|---|---|---|---|---|---|---|---|---|---|---|---|---|---|---|---|---|---|---|---|---|---|---|---|---|---|---|---|---|---|---|---|---|---|---|---|---|---|---|---|---|---|
| 1093 | GSK1904529A | GSK-1904529A, GSK 1904529A | 21647 | 0.945 | 1.000 | 6337.5 | 0.110 | 0.352 | 12971 | 0.304 | 0.705 | 26141.5 | 1.000 | 1.000 | 29930.5 | 0.025 | 0.201 | 35022 | 0.420 | 0.746 | 22287 | 0.909 | 1.000 | 7890 | 0.352 | 0.705 | 11460 | 0.707 | 0.942 | 9704 | 0.067 | 0.269 | 27397.5 | 0.674 | 0.942 | 17121 | 0.330 | 0.705 | 14029 | 0.922 | 1.000 |
| 1096 | Tozasertib | MK 0457,MK-0457,MK-045, VX-680 VX 680 VX-68 | 1468 | 0.348 | 0.697 | 65 | 0.051 | 0.203 | 257 | 0.027 | 0.145 | 2136 | 0.179 | 0.476 | 1736 | 0.947 | 1.000 | 1814 | 0.104 | 0.334 | 1131 | 0.001 | 0.011 | 625 | 0.528 | 0.753 | 130 | 0.000 | 0.002 | 234 | 0.614 | 0.755 | 1777 | 0.565 | 0.753 | 5691 | 1.000 | 1.000 | 1699 | 0.843 | 0.963 |
| 1129 | PF-4708671 | PF 4708671, PF4708671 | 1479 | 0.366 | 0.590 | 196 | 0.442 | 0.590 | 528 | 0.540 | 0.664 | 2973.5 | 0.975 | 0.978 | 1291 | 0.346 | 0.590 | 1747 | 0.067 | 0.535 | 1790 | 0.323 | 0.590 | 528 | 0.279 | 0.590 | 593 | 0.442 | 0.590 | 181 | 0.374 | 0.590 | 2236 | 0.978 | 0.978 | 4144.5 | 0.785 | 0.897 | 1306 | 0.237 | 0.590 |
| 1131 | PRIMA-1MET | Prima 1MET, APR-246 | 15378 | 0.202 | 0.404 | 3493 | 0.001 | 0.004 | 13711 | 0.969 | 1.000 | 13223 | 0.005 | 0.021 | 29138 | 0.685 | 1.000 | 35192 | 0.998 | 1.000 | 14562.5 | 0.159 | 0.364 | 8282 | 0.866 | 1.000 | 4999 | 0.000 | 0.002 | 9916 | 0.738 | 1.000 | 30583 | 1.000 | 1.000 | 13676 | 0.159 | 0.364 | 6001 | 0.000 | 0.000 |
| 1133 | Serdemetan | JNJ-26854165 | 20640 | 0.920 | 0.973 | 6907 | 0.296 | 0.676 | 10526.5 | 0.013 | 0.070 | 14867 | 0.003 | 0.049 | 36824 | 0.973 | 0.973 | 34499.5 | 0.544 | 0.973 | 18400 | 0.141 | 0.452 | 7192 | 0.173 | 0.462 | 8329 | 0.043 | 0.174 | 11747 | 0.829 | 0.973 | 26782 | 0.763 | 0.973 | 20084.5 | 0.959 | 0.973 | 9213 | 0.011 | 0.070 |
| 1149 | TW 37 | TW37, TW-37 | 2842 | 0.431 | 0.944 | 1380 | 0.532 | 0.944 | 2335 | 0.070 | 0.373 | 3032 | 0.931 | 0.944 | 5738 | 0.648 | 0.944 | 5378 | 0.556 | 0.944 | 2242 | 0.022 | 0.175 | 1948 | 0.944 | 0.944 | 1434 | 0.120 | 0.478 | 2148 | 0.747 | 0.944 | 4496 | 0.931 | 0.944 | 2945 | 0.536 | 0.944 | 1560 | 0.776 | 0.944 |
| 1168 | Erlotinib | Tarceva, RG-1415, CP-358774, OSI-774, Ro-508231, R-1415 | 21788.5 | 0.961 | 1.000 | 10279 | 0.998 | 1.000 | 16644 | 0.990 | 1.000 | 20997 | 0.794 | 1.000 | 33825 | 0.498 | 0.996 | 31195 | 0.029 | 0.153 | 28146.5 | 1.000 | 1.000 | 7542 | 0.249 | 0.765 | 12826 | 0.961 | 1.000 | 9759.5 | 0.080 | 0.318 | 13170 | 0.000 | 0.000 | 21933.5 | 0.998 | 1.000 | 6753 | 0.000 | 0.000 |
| 1170 | CCT-018159 | CCT018159, CCT 018159 | 855 | 0.024 | 0.191 | 147 | 0.391 | 0.915 | 488 | 0.761 | 0.937 | 1980 | 0.702 | 0.937 | 1029 | 0.376 | 0.915 | 1551 | 0.279 | 0.915 | 1897 | 0.976 | 0.976 | 692 | 0.958 | 0.976 | 493 | 0.515 | 0.915 | 210 | 0.725 | 0.937 | 937 | 0.016 | 0.191 | 342 | 0.271 | 0.915 | 1221 | 0.643 | 0.937 |
| 1175 | Rucaparib | PF-01367338, AG-014699, AG-14447, AG-14699 | 19200 | 0.679 | 0.894 | 6609 | 0.199 | 0.638 | 11182 | 0.043 | 0.342 | 19930 | 0.693 | 0.894 | 34460.5 | 0.782 | 0.894 | 34525 | 0.549 | 0.878 | 18487.5 | 0.154 | 0.615 | 7540 | 0.275 | 0.671 | 6934 | 0.002 | 0.027 | 10011 | 0.294 | 0.671 | 29926.5 | 0.993 | 0.993 | 19905 | 0.947 | 0.993 | 10640 | 0.128 | 0.615 |
| 1177 | Niraparib | MK-4827, MK4827 | 15539 | 0.236 | 0.659 | 4727 | 0.039 | 0.156 | 6369 | 0.000 | 0.000 | 17989 | 0.789 | 1.000 | 28454 | 0.543 | 0.869 | 31284 | 0.781 | 1.000 | 13253 | 0.026 | 0.137 | 6651 | 0.258 | 0.659 | 3711 | 0.000 | 0.000 | 10349 | 0.853 | 1.000 | 26990 | 0.992 | 1.000 | 20707 | 1.000 | 1.000 | 10245 | 0.288 | 0.659 |
| 1179 | MK-1775 | AZD1775, Adavosertib | 22534 | 0.971 | 1.000 | 6442 | 0.128 | 0.292 | 13523 | 0.450 | 0.800 | 25681 | 1.000 | 1.000 | 31697 | 0.123 | 0.292 | 36638 | 0.692 | 1.000 | 17381.5 | 0.027 | 0.109 | 11574 | 0.999 | 1.000 | 4498 | 0.000 | 0.000 | 14817 | 0.996 | 1.000 | 22812 | 0.020 | 0.105 | 24630 | 1.000 | 1.000 | 11316 | 0.223 | 0.447 |
| 1180 | Dinaciclib | SCH727965, SCH 727965, SCH-727-965, MK-7965 | 16619 | 0.527 | 0.999 | 5786 | 0.300 | 0.932 | 9366 | 0.019 | 0.100 | 12269 | 0.000 | 0.003 | 30017 | 0.838 | 0.999 | 33571 | 0.978 | 0.999 | 15591 | 0.408 | 0.932 | 9323 | 0.988 | 0.999 | 8719 | 0.798 | 0.999 | 12442 | 0.999 | 0.999 | 17294 | 0.000 | 0.003 | 14639 | 0.398 | 0.932 | 12064 | 0.855 | 0.999 |
| 1190 | Gemcitabine | Gemzar, LY-188011 | 16280 | 0.024 | 0.096 | 4577 | 0.001 | 0.011 | 9558 | 0.002 | 0.012 | 18181 | 0.164 | 0.327 | 34449 | 0.597 | 0.955 | 39832 | 0.982 | 1.000 | 17900 | 0.124 | 0.327 | 9635 | 0.908 | 1.000 | 8698 | 0.038 | 0.122 | 12712 | 0.840 | 1.000 | 24626 | 0.157 | 0.327 | 26876 | 1.000 | 1.000 | 7716 | 0.000 | 0.003 |
| 1191 | Bortezomib | PS-341, LDP-341, Velcade | 13399 | 0.000 | 0.001 | 6946 | 0.286 | 0.573 | 14755 | 0.817 | 1.000 | 12465 | 0.000 | 0.000 | 35708.5 | 0.822 | 1.000 | 46898.5 | 1.000 | 1.000 | 26636 | 1.000 | 1.000 | 8015 | 0.418 | 0.743 | 8811 | 0.050 | 0.160 | 12845.5 | 0.874 | 1.000 | 24955 | 0.273 | 0.573 | 12404 | 0.000 | 0.001 | 8390 | 0.001 | 0.005 |
| 1192 | GSK269962A | GSK 269962A | 997 | 0.008 | 0.133 | 99 | 0.115 | 0.368 | 408 | 0.221 | 0.555 | 3643 | 1.000 | 1.000 | 1278 | 0.324 | 0.577 | 1787 | 0.088 | 0.368 | 1756 | 0.277 | 0.555 | 631 | 0.543 | 0.749 | 424 | 0.101 | 0.368 | 246 | 0.662 | 0.815 | 2124 | 0.941 | 1.000 | 4270 | 0.870 | 0.995 | 1029 | 0.027 | 0.213 |
| 1194 | SB505124 | SB 505124, SB505124 | 1720 | 0.787 | 1.000 | 109 | 0.143 | 0.763 | 630 | 0.817 | 1.000 | 2709 | 0.884 | 1.000 | 1360 | 0.505 | 1.000 | 2523 | 0.927 | 1.000 | 1715 | 0.260 | 1.000 | 654 | 0.628 | 1.000 | 326 | 0.026 | 0.211 | 272 | 0.769 | 1.000 | 2476 | 0.999 | 1.000 | 665 | 0.879 | 1.000 | 2218 | 1.000 | 1.000 |
| 1199 | Tamoxifen | Nolvadex, Soltamox, Zynoplex, ICI-46474, Kessar | 23114 | 0.996 | 0.996 | 7064 | 0.325 | 0.520 | 11693.5 | 0.077 | 0.308 | 23659 | 0.994 | 0.996 | 34629 | 0.648 | 0.943 | 36701 | 0.753 | 0.996 | 23331 | 0.989 | 0.996 | 7736 | 0.313 | 0.520 | 10097 | 0.287 | 0.520 | 10663 | 0.253 | 0.520 | 25248 | 0.321 | 0.520 | 14735 | 0.025 | 0.201 | 13524 | 0.859 | 0.996 |
| 1200 | Fulvestrant | Faslodex, ICI-182780 | 17996.5 | 0.245 | 0.653 | 8718.5 | 0.886 | 1.000 | 13686 | 0.521 | 0.834 | 25311.5 | 1.000 | 1.000 | 34671.5 | 0.637 | 0.927 | 30414 | 0.010 | 0.055 | 26344.5 | 1.000 | 1.000 | 5448 | 0.003 | 0.026 | 11878.5 | 0.822 | 1.000 | 6542 | 0.000 | 0.000 | 30682 | 0.988 | 1.000 | 14958.5 | 0.033 | 0.132 | 12060 | 0.453 | 0.805 |
| 1237 | EPZ004777 | EPZ-004777 | 19785.5 | 0.661 | 0.934 | 8070 | 0.701 | 0.934 | 11773.5 | 0.079 | 0.420 | 19126.5 | 0.337 | 0.635 | 33757 | 0.447 | 0.715 | 33883.5 | 0.231 | 0.625 | 23155 | 0.970 | 0.973 | 7347.5 | 0.183 | 0.625 | 10384.5 | 0.357 | 0.635 | 9591 | 0.056 | 0.420 | 28254 | 0.819 | 0.973 | 19922 | 0.920 | 0.973 | 10439 | 0.076 | 0.420 |
| 1239 | YK-4-279 | YK 4-279 | 16609 | 0.061 | 0.196 | 6136 | 0.081 | 0.216 | 10262 | 0.006 | 0.048 | 21620 | 0.887 | 1.000 | 34502 | 0.625 | 1.000 | 38787.5 | 0.952 | 1.000 | 18507 | 0.229 | 0.523 | 12654 | 1.000 | 1.000 | 5271 | 0.000 | 0.000 | 14968 | 0.998 | 1.000 | 22929 | 0.042 | 0.196 | 20102 | 0.945 | 1.000 | 13530 | 0.860 | 1.000 |
| 1243 | Piperlongumine | Piplartine | 1213 | 0.478 | 0.955 | 27 | 0.015 | 0.118 | 577 | 0.939 | 0.993 | 1286 | 0.006 | 0.101 | 982 | 0.279 | 0.745 | 2210 | 0.993 | 0.993 | 1229 | 0.102 | 0.498 | 657 | 0.922 | 0.993 | 404 | 0.241 | 0.745 | 283 | 0.952 | 0.993 | 1530 | 0.809 | 0.993 | 492 | 0.772 | 0.993 | 946 | 0.125 | 0.498 |
| 1248 | Daporinad | APO866, FK866, FK866 | 7701 | 0.409 | 1.000 | 1952 | 0.093 | 0.372 | 2871 | 0.000 | 0.000 | 11752 | 1.000 | 1.000 | 11777 | 0.605 | 1.000 | 14967 | 0.977 | 1.000 | 5072 | 0.000 | 0.000 | 4030 | 0.752 | 1.000 | 1846 | 0.000 | 0.002 | 4221 | 0.728 | 1.000 | 9393 | 0.469 | 1.000 | 7654 | 0.996 | 1.000 | 6132 | 0.869 | 1.000 |
| 1249 | BMS-345541 | | 18494 | 0.920 | 0.998 | 6555 | 0.644 | 0.937 | 9579 | 0.081 | 0.291 | 16934 | 0.529 | 0.937 | 30766 | 0.919 | 0.998 | 29570 | 0.456 | 0.937 | 13585 | 0.045 | 0.291 | 8983 | 0.970 | 0.998 | 6083 | 0.007 | 0.108 | 9804 | 0.707 | 0.943 | 22905 | 0.476 | 0.937 | 18932 | 0.998 | 0.998 | 9347 | 0.091 | 0.291 |
| 1250 | AZ960 | AZ 960, AZ-960 | 11083 | 0.000 | 0.000 | 5429 | 0.172 | 0.392 | 8223 | 0.002 | 0.010 | 22303 | 1.000 | 1.000 | 32224 | 0.985 | 1.000 | 29148 | 0.359 | 0.639 | 13990 | 0.079 | 0.252 | 6044 | 0.096 | 0.256 | 3883.5 | 0.000 | 0.000 | 12888 | 1.000 | 1.000 | 21778 | 0.223 | 0.446 | 19225 | 0.999 | 1.000 | 7434 | 0.001 | 0.007 |
| 1259 | Talazoparib | BMN-673, BMN 973 | 14123.5 | 0.001 | 0.005 | 6183 | 0.090 | 0.249 | 8124 | 0.000 | 0.000 | 21132 | 0.822 | 1.000 | 34240.5 | 0.587 | 1.000 | 38621.5 | 0.947 | 1.000 | 16448 | 0.022 | 0.086 | 8314 | 0.534 | 1.000 | 6203 | 0.000 | 0.000 | 13879 | 0.977 | 1.000 | 28753 | 0.929 | 1.000 | 24859.5 | 1.000 | 1.000 | 10509 | 0.093 | 0.249 |
| 1268 | XAV939 | NVP-XAV939, XAV-939, XAV 939 | 14062 | 0.038 | 0.152 | 7824 | 0.967 | 1.000 | 14169 | 0.988 | 1.000 | 19020 | 0.937 | 1.000 | 24590.5 | 0.023 | 0.143 | 24036.5 | 0.001 | 0.017 | 21723 | 1.000 | 1.000 | 5834 | 0.063 | 0.201 | 9155.5 | 0.713 | 1.000 | 8398.5 | 0.209 | 0.558 | 22930.5 | 0.474 | 0.842 | 16794 | 0.910 | 1.000 | 8686 | 0.027 | 0.143 |
| 1372 | Trametinib | GSK1120212, Mekinist | 23374 | 0.992 | 1.000 | 8552 | 0.839 | 1.000 | 21091.5 | 1.000 | 1.000 | 4657 | 0.000 | 0.000 | 37647 | 0.952 | 1.000 | 31769.5 | 0.039 | 0.088 | 35403 | 1.000 | 1.000 | 9262 | 0.822 | 1.000 | 11133 | 0.596 | 1.000 | 9001 | 0.018 | 0.069 | 22878 | 0.022 | 0.069 | 25560 | 1.000 | 1.000 | 12370 | 0.530 | 1.000 |
| 1373 | Dabrafenib | GSK2118436, Tafinlar | 20760.5 | 0.923 | 1.000 | 6475 | 0.157 | 0.494 | 14571.5 | 0.865 | 1.000 | 7284 | 0.000 | 0.000 | 37452.5 | 0.978 | 1.000 | 32695 | 0.140 | 0.494 | 26846 | 1.000 | 1.000 | 7938 | 0.405 | 0.720 | 12728.5 | 0.960 | 1.000 | 11008 | 0.375 | 0.720 | 28465 | 0.890 | 1.000 | 22419 | 0.999 | 1.000 | 7516 | 0.000 | 0.001 |
| 1375 | Temozolomide | Temodar, Temodal, M-39831, SCH 52365 | 16245 | 0.020 | 0.082 | 7200 | 0.358 | 0.716 | 14691 | 0.778 | 0.958 | 16042 | 0.009 | 0.066 | 34472.5 | 0.573 | 0.868 | 35189 | 0.441 | 0.783 | 21455 | 0.723 | 0.958 | 5803.5 | 0.008 | 0.066 | 12861 | 0.957 | 1.000 | 9593 | 0.055 | 0.176 | 27053 | 0.597 | 0.868 | 28058.5 | 1.000 | 1.000 | 9432 | 0.012 | 0.066 |
| 1378 | Bleomycin (50 uM) | | 6924 | 0.001 | 0.011 | 4239 | 0.082 | 0.264 | 6978 | 0.007 | 0.054 | 9204 | 0.755 | 1.000 | 19455 | 0.036 | 0.190 | 22217 | 0.740 | 1.000 | 11889 | 0.923 | 1.000 | 5040 | 0.701 | 1.000 | 5142 | 0.062 | 0.250 | 9101 | 0.937 | 1.000 | 14157 | 0.186 | 0.453 | 17507.5 | 1.000 | 1.000 | 4811 | 0.274 | 0.548 |
| 1401 | AZD5438 | SN1057898678 | 9405 | 0.523 | 0.888 | 3589 | 0.076 | 0.392 | 7750 | 0.185 | 0.492 | 6785 | 0.045 | 0.392 | 20140 | 0.555 | 0.888 | 22619.5 | 0.945 | 0.999 | 10041 | 0.766 | 0.972 | 7588 | 0.999 | 0.999 | 4969 | 0.291 | 0.665 | 8459 | 0.924 | 0.999 | 12831.5 | 0.052 | 0.392 | 12231 | 0.790 | 0.972 | 4777 | 0.122 | 0.392 |
| 1428 | IAP_5620 | SN1043546339 | 9601 | 0.618 | 0.824 | 3403 | 0.045 | 0.317 | 9877 | 0.929 | 1.000 | 9684 | 0.941 | 1.000 | 20151.5 | 0.581 | 0.824 | 18101 | 0.060 | 0.317 | 11067.5 | 0.967 | 1.000 | 5078 | 0.354 | 0.809 | 7838 | 1.000 | 1.000 | 6774 | 0.294 | 0.809 | 14172 | 0.321 | 0.809 | 8380.5 | 0.008 | 0.133 | 5663.5 | 0.524 | 0.824 |
| 1441 | AZD2014 | SN1103949359 | 8712 | 0.252 | 0.550 | 4607 | 0.538 | 0.825 | 7211 | 0.068 | 0.309 | 7138 | 0.097 | 0.309 | 19988 | 0.512 | 0.825 | 19528 | 0.275 | 0.550 | 12282 | 0.999 | 0.999 | 4277 | 0.066 | 0.309 | 6737 | 0.971 | 0.999 | 8659 | 0.953 | 0.999 | 16580 | 0.933 | 0.999 | 9962 | 0.089 | 0.309 | 4802 | 0.129 | 0.345 |
| 1449 | AZD1208 | SN1028350371 | 9695.5 | 0.648 | 0.965 | 5310 | 0.878 | 0.965 | 9259.5 | 0.780 | 0.965 | 9051 | 0.800 | 0.965 | 22197.5 | 0.965 | 0.965 | 19129.5 | 0.197 | 0.524 | 7539.5 | 0.030 | 0.240 | 3083 | 0.001 | 0.013 | 4180 | 0.051 | 0.275 | 6989.5 | 0.382 | 0.854 | 16346.5 | 0.908 | 0.965 | 13184 | 0.959 | 0.965 | 4801.5 | 0.132 | 0.512 |
| 1463 | AZD1332 | SN1061387896 | 8526 | 0.194 | 0.613 | 4466 | 0.454 | 0.726 | 6994 | 0.042 | 0.338 | 11211 | 0.999 | 0.999 | 18916 | 0.230 | 0.613 | 20410 | 0.511 | 0.743 | 10383 | 0.860 | 0.999 | 2428 | 0.000 | 0.000 | 4999.5 | 0.306 | 0.640 | 6406 | 0.157 | 0.613 | 14221.5 | 0.320 | 0.640 | 14533 | 0.999 | 0.999 | 4646 | 0.090 | 0.481 |
| 1494 | SN-38 | 7-Ethyl-10-Hydroxy-Camptothecin, SN 38 | 15634 | 0.016 | 0.043 | 6919 | 0.280 | 0.623 | 10412 | 0.009 | 0.028 | 15251.5 | 0.003 | 0.015 | 35451 | 0.866 | 1.000 | 40039.5 | 0.989 | 1.000 | 15324 | 0.001 | 0.008 | 8520 | 0.616 | 1.000 | 6494 | 0.001 | 0.008 | 14843 | 0.998 | 1.000 | 25126 | 0.311 | 0.623 | 23302.5 | 1.000 | 1.000 | 9066 | 0.007 | 0.027 |
| 1502 | Bicalutamide | ICI-176334, Casodex, Cosudex, ICI 176334 | 903 | 0.043 | 0.663 | 76 | 0.107 | 0.663 | 384 | 0.409 | 0.665 | 1983 | 0.706 | 0.807 | 1060 | 0.444 | 0.665 | 1467 | 0.166 | 0.664 | 1488 | 0.497 | 0.665 | 553 | 0.704 | 0.807 | 487 | 0.495 | 0.665 | 134 | 0.326 | 0.665 | 1864 | 0.995 | 0.995 | 363 | 0.338 | 0.665 | 1154 | 0.499 | 0.665 |
| 1507 | Ruxolitinib | INCB-18424, Ruxolitinib Phosphate, Jakafi | 15605.5 | 0.230 | 0.664 | 6384 | 0.547 | 0.875 | 11297 | 0.401 | 0.712 | 22120 | 1.000 | 1.000 | 27302.5 | 0.269 | 0.664 | 20508.5 | 0.000 | 0.000 | 20613 | 1.000 | 1.000 | 5701.5 | 0.044 | 0.233 | 9256 | 0.729 | 1.000 | 14824 | 1.000 | 1.000 | 22736 | 0.394 | 0.712 | 18290 | 0.991 | 1.000 | 8039 | 0.006 | 0.044 |
| 1510 | Linsitinib | OSI-906, ASP-7487 | 28528 | 1.000 | 1.000 | 6704 | 0.191 | 0.412 | 7339 | 0.000 | 0.000 | 25923 | 1.000 | 1.000 | 41230 | 1.000 | 1.000 | 29982.5 | 0.005 | 0.019 | 17920.5 | 0.056 | 0.148 | 6394 | 0.033 | 0.105 | 4706 | 0.000 | 0.000 | 10559 | 0.206 | 0.412 | 28571 | 0.851 | 1.000 | 22734.5 | 0.999 | 1.000 | 13948 | 0.908 | 1.000 |
| 1511 | Epirubicin | Ellence, Farmorubicin, IMI-28 | 15256 | 0.008 | 0.032 | 5626 | 0.028 | 0.089 | 9231.5 | 0.001 | 0.004 | 12381 | 0.000 | 0.000 | 36161 | 0.868 | 1.000 | 40946 | 0.996 | 1.000 | 19263 | 0.394 | 0.645 | 7594 | 0.266 | 0.607 | 7513 | 0.003 | 0.015 | 16163 | 1.000 | 1.000 | 25651.5 | 0.403 | 0.645 | 24019 | 1.000 | 1.000 | 10463 | 0.086 | 0.228 |
| 1512 | Cyclophosphamid | Cytoxan, Neosar, Cyclophosphamid, Procytox, Cyclophosphane | 19293.5 | 0.574 | 1.000 | 8408.5 | 0.823 | 1.000 | 14321.5 | 0.718 | 1.000 | 21553.5 | 0.883 | 1.000 | 30481 | 0.056 | 0.215 | 32007.5 | 0.067 | 0.215 | 21841.5 | 0.915 | 1.000 | 6127 | 0.021 | 0.110 | 12438.5 | 0.925 | 1.000 | 8891.5 | 0.017 | 0.110 | 32416 | 1.000 | 1.000 | 17415 | 0.434 | 0.869 | 10750 | 0.195 | 0.446 |
| 1529 | Pevonedistat | MLN4924, MLN 4924, MLN-4924 | 18205.5 | 0.311 | 0.712 | 6282 | 0.109 | 0.349 | 10183 | 0.005 | 0.041 | 19228 | 0.399 | 0.798 | 36368 | 0.897 | 0.967 | 37942 | 0.907 | 0.967 | 17018 | 0.105 | 0.349 | 9316.5 | 0.858 | 0.967 | 7772 | 0.006 | 0.041 | 12347 | 0.773 | 0.967 | 21601 | 0.008 | 0.041 | 22670 | 1.000 | 1.000 | 12231.5 | 0.528 | 0.939 |
| 1549 | Sapitinib | AZD8931 | 25031.5 | 1.000 | 1.000 | 10711 | 0.999 | 1.000 | 18669.5 | 1.000 | 1.000 | 27346.5 | 1.000 | 1.000 | 31459.5 | 0.106 | 0.241 | 29028 | 0.002 | 0.010 | 31515 | 1.000 | 1.000 | 10054 | 0.955 | 1.000 | 14232 | 0.998 | 1.000 | 9223.5 | 0.029 | 0.092 | 9649 | 0.000 | 0.000 | 13291 | 0.001 | 0.010 | 8874 | 0.004 | 0.014 |
| 1553 | Uprosertib | GSK2141795, GSK2141795C, GSK-2141795 | 16298 | 0.408 | 0.653 | 7217 | 0.863 | 0.999 | 10808 | 0.257 | 0.653 | 16567 | 0.399 | 0.653 | 27566 | 0.327 | 0.653 | 29501 | 0.402 | 0.653 | 18708.5 | 0.975 | 0.999 | 8989 | 0.966 | 0.999 | 11876 | 0.999 | 0.999 | 12416 | 0.999 | 0.999 | 20585 | 0.062 | 0.492 | 11519 | 0.001 | 0.009 | 9674 | 0.136 | 0.550 |
| 1557 | LCL161 | | 14635 | 0.086 | 0.344 | 7197 | 0.865 | 1.000 | 14237 | 0.990 | 1.000 | 14316 | 0.036 | 0.194 | 28240 | 0.497 | 1.000 | 23181.5 | 0.000 | 0.003 | 22251 | 1.000 | 1.000 | 7624 | 0.654 | 1.000 | 11293.5 | 0.996 | 1.000 | 9337 | 0.534 | 1.000 | 19219 | 0.010 | 0.084 | 15756 | 0.713 | 1.000 | 10140 | 0.258 | 0.687 |
| 1558 | Lapatinib | Tykerb, Tyverb | 23386 | 0.993 | 1.000 | 10200 | 0.996 | 1.000 | 16200 | 0.971 | 1.000 | 22398 | 0.946 | 1.000 | 38975 | 0.990 | 1.000 | 36262 | 0.637 | 1.000 | 27444 | 1.000 | 1.000 | 10033 | 0.953 | 1.000 | 11661 | 0.758 | 1.000 | 14513 | 0.992 | 1.000 | 11354 | 0.000 | 0.000 | 11460 | 0.000 | 0.000 | 5993 | 0.000 | 0.000 |
| 1559 | Luminespib | AUY922, VER-52296,NVP-AUY922, AUY | 12684 | 0.000 | 0.000 | 4017 | 0.001 | 0.002 | 9317 | 0.001 | 0.002 | 19893 | 0.530 | 1.000 | 36030 | 0.835 | 1.000 | 40112 | 0.986 | 1.000 | 26357 | 1.000 | 1.000 | 9853 | 0.937 | 1.000 | 9602 | 0.155 | 0.413 | 15021 | 0.998 | 1.000 | 19496 | 0.000 | 0.001 | 18025 | 0.569 | 1.000 | 14310 | 0.951 | 1.000 |
| 1560 | Alpelisib | BYL719, BYL-719, NVP-BYL719 | 22372 | 0.980 | 0.993 | 8301.5 | 0.777 | 0.993 | 10795 | 0.016 | 0.084 | 23724 | 0.993 | 0.993 | 37823 | 0.964 | 0.993 | 34114.5 | 0.265 | 0.728 | 23545 | 0.983 | 0.993 | 9787 | 0.927 | 0.993 | 10169 | 0.292 | 0.728 | 13126 | 0.904 | 0.993 | 18080 | 0.000 | 0.000 | 12723 | 0.000 | 0.003 | 11705 | 0.335 | 0.728 |
| 1561 | Taselisib | GDC-0032, GDC0032, RG7604 | 20624.5 | 0.756 | 1.000 | 7119 | 0.325 | 1.000 | 10207 | 0.004 | 0.036 | 23725 | 0.992 | 1.000 | 35538 | 0.747 | 1.000 | 36456 | 0.661 | 1.000 | 26989 | 1.000 | 1.000 | 11874 | 1.000 | 1.000 | 10684 | 0.446 | 1.000 | 14480 | 0.991 | 1.000 | 13850.5 | 0.000 | 0.000 | 14703 | 0.020 | 0.105 | 12054 | 0.430 | 1.000 |
| 1563 | EPZ5676 | EPZ-5676, Pinometostat | 16659.5 | 0.060 | 0.191 | 7984 | 0.674 | 1.000 | 15254.5 | 0.889 | 1.000 | 17500 | 0.110 | 0.295 | 36956.5 | 0.922 | 1.000 | 31426 | 0.031 | 0.128 | 27297.5 | 1.000 | 1.000 | 7174 | 0.144 | 0.330 | 11533.5 | 0.732 | 1.000 | 8989.5 | 0.019 | 0.128 | 25331.5 | 0.260 | 0.520 | 21413 | 0.992 | 1.000 | 9405.5 | 0.012 | 0.128 |
| 1564 | SCH772984 | CS-1421 | 24245 | 0.999 | 1.000 | 8837 | 0.904 | 1.000 | 20746 | 1.000 | 1.000 | 5722 | 0.000 | 0.000 | 36947 | 0.917 | 1.000 | 31584.5 | 0.035 | 0.094 | 34898 | 1.000 | 1.000 | 9320 | 0.842 | 1.000 | 11395 | 0.688 | 1.000 | 6433 | 0.000 | 0.000 | 22584 | 0.016 | 0.058 | 23893.5 | 1.000 | 1.000 | 13898 | 0.906 | 1.000 |
| 1576 | IWP-2 | Wnt Inhibitor IWP-2 | 15340 | 0.194 | 0.495 | 7418 | 0.913 | 0.999 | 12885.5 | 0.877 | 0.999 | 18369 | 0.858 | 0.999 | 26817.5 | 0.216 | 0.495 | 24689 | 0.003 | 0.025 | 20252.5 | 0.999 | 0.999 | 7350 | 0.541 | 0.999 | 8910.5 | 0.626 | 0.999 | 6105.5 | 0.001 | 0.022 | 24268 | 0.771 | 0.999 | 17518.5 | 0.970 | 0.999 | 9971.5 | 0.212 | 0.495 |
| 1578 | Leflunomide | Sulol, SU-101, HWA-486, Arava | 18216 | 0.883 | 0.996 | 4915 | 0.061 | 0.202 | 10964 | 0.316 | 0.631 | 20662.5 | 0.996 | 0.996 | 25441.5 | 0.062 | 0.202 | 26042 | 0.022 | 0.178 | 18496.5 | 0.969 | 0.996 | 8190 | 0.843 | 0.996 | 7856 | 0.235 | 0.536 | 8318 | 0.188 | 0.501 | 26090.5 | 0.967 | 0.996 | 16607 | 0.885 | 0.996 | 10668 | 0.426 | 0.758 |
| 1580 | GSK2801 | 1-(7-(2-methylsulfonyl-phenyl)-4-propoxy-1-aza-bicyclo[4.3.0]nona-2,4,6,8-tetraen-9-yl)-ethanone | 10263 | 0.826 | 0.998 | 5592.5 | 0.941 | 0.998 | 9086 | 0.601 | 0.962 | 10469 | 0.980 | 0.998 | 19819 | 0.452 | 0.804 | 17622.5 | 0.024 | 0.193 | 11606 | 0.990 | 0.998 | 5236.5 | 0.423 | 0.804 | 6725.5 | 0.969 | 0.998 | 5917 | 0.055 | 0.291 | 13922.5 | 0.177 | 0.404 | 14403.5 | 0.998 | 0.998 | 4907 | 0.159 | 0.404 |
| 1581 | Bromosporine | ethyl N-[6-(3-methanesulfonamido-4-methylphenyl)-3-methyl-[1,2,4]triazolo[4,3-b]pyridazin-8-yl]carbamate | 7834 | 0.056 | 0.507 | 4120 | 0.262 | 0.699 | 9305 | 0.789 | 0.968 | 8262 | 0.368 | 0.751 | 18058.5 | 0.088 | 0.507 | 20045 | 0.409 | 0.751 | 10488 | 0.883 | 0.968 | 4702.5 | 0.179 | 0.573 | 6204 | 0.876 | 0.968 | 7095 | 0.425 | 0.751 | 16687 | 0.916 | 0.968 | 13335.5 | 0.968 | 0.968 | 5583 | 0.470 | 0.751 |
| 1582 | SGC-CBP30 | 8-(3-chloro-4-methoxy-phenethyl)-4-(3,5-dimethyl-isoxazol-4-yl)-9-(2-(morpholin-4-yl)-propyl)-7,9-diaza-bicyclo[4.3.0]nona-1(6),2,4,7-tetraene | 9803 | 0.683 | 0.910 | 4611 | 0.540 | 0.910 | 8712.5 | 0.562 | 0.910 | 9848.5 | 0.919 | 0.974 | 19535.5 | 0.384 | 0.878 | 18149 | 0.057 | 0.328 | 10742 | 0.928 | 0.974 | 5117 | 0.365 | 0.878 | 5680 | 0.667 | 0.910 | 6550 | 0.201 | 0.645 | 15443.5 | 0.637 | 0.910 | 12756 | 0.904 | 0.974 | 4496 | 0.062 | 0.328 |

| DRUG_ID | DRUG_NAME | SYNONYMS | large_intestine_MWU_STAT | large_intestine_PVAL | large_intestine_PVAL_ADJ | thyroid_MWU_STAT | thyroid_PVAL | thyroid_PVAL_ADJ | digestive_system_MWU_STAT | digestive_system_PVAL | digestive_system_PVAL_ADJ | digestive_system_PVAL2 |
|---|---|---|---|---|---|---|---|---|---|---|---|---|
| 1093 | GSK1904529A | GSK-1904529A, GSK 1904529A | 16483.5 | 0.516 | 0.826 | 4178 | 0.038 | 0.204 | 13619.5 | 0.010 | 0.152 | 0.152 |
| 1096 | Tozasertib | MK 0457,MK-0457,MK-045, VX-680 VX 680 VX-68 | 426 | 0.261 | 0.598 | 105 | 0.502 | 0.753 | 481 | 0.405 | 0.720 | 0.720 |
| 1129 | PF-4708671 | PF 4708671, PF4708671 | 456 | 0.332 | 0.590 | 63 | 0.248 | 0.590 | 263 | 0.031 | 0.498 | 0.498 |
| 1131 | PRIMA-1MET | Prima 1MET, APR-246 | 17056.5 | 0.938 | 1.000 | 3122 | 0.027 | 0.085 | 14816 | 0.630 | 1.000 | 1.000 |
| 1133 | Serdemetan | JNJ-26854165 | 16690.5 | 0.731 | 0.973 | 6699 | 0.901 | 0.973 | 18380 | 0.904 | 0.973 | 0.973 |
| 1149 | TW 37 | TW37, TW-37 | 2594 | 0.011 | 0.175 | 1009 | 0.410 | 0.944 | 3504 | 0.876 | 0.944 | 0.944 |
| 1168 | Erlotinib | Tarceva, RG-1415, CP-358774, OSI-774, Ro-508231, R-1415 | 14853 | 0.287 | 0.765 | 6952 | 0.933 | 1.000 | 16682 | 0.411 | 0.940 | 0.940 |
| 1170 | CCT-018159 | CCT018159, CCT 018159 | 572 | 0.933 | 0.976 | 79 | 0.473 | 0.915 | 394 | 0.445 | 0.915 | 0.915 |
| 1175 | Rucaparib | PF-01367338, AG-014699, AG-14447, AG-14699 | 15521 | 0.417 | 0.774 | 5471.5 | 0.435 | 0.774 | 17438 | 0.743 | 0.894 | 0.894 |
| 1177 | Niraparib | MK-4827, MK4827 | 18260 | 0.993 | 1.000 | 4282 | 0.361 | 0.722 | 14339 | 0.487 | 0.865 | 0.865 |
| 1179 | MK-1775 | AZD1775, Adavosertib | 11842 | 0.001 | 0.006 | 6884.5 | 0.913 | 1.000 | 15391 | 0.119 | 0.292 | 0.292 |
| 1180 | Dinaciclib | SCH727965, SCH 727965, SCH-727-965, MK-7965 | 16998 | 0.934 | 0.999 | 5404 | 0.881 | 0.999 | 12340 | 0.093 | 0.372 | 0.372 |
| 1190 | Gemcitabine | Gemzar, LY-188011 | 17331 | 0.816 | 1.000 | 5803 | 0.555 | 0.955 | 20580 | 0.991 | 1.000 | 1.000 |
| 1191 | Bortezomib | PS-341, LDP-341, Velcade | 17191.5 | 0.865 | 1.000 | 4772 | 0.148 | 0.396 | 18430 | 0.888 | 1.000 | 1.000 |
| 1192 | GSK269962A | GSK 269962A | 492 | 0.436 | 0.697 | 55 | 0.265 | 0.555 | 536 | 0.561 | 0.749 | 0.749 |
| 1194 | SB505124 | SB 505124, SB505124 | 1254 | 0.000 | 0.000 | 142 | 0.687 | 1.000 | 510 | 0.509 | 1.000 | 1.000 |
| 1199 | Tamoxifen | Nolvadex, Soltamox, Zynoplex, ICI-46474, Kessar | 11585 | 0.002 | 0.033 | 4857 | 0.171 | 0.520 | 14709.5 | 0.060 | 0.308 | 0.308 |
| 1200 | Fulvestrant | Faslodex, ICI-182780 | 15310 | 0.393 | 0.787 | 5355 | 0.350 | 0.787 | 15382.5 | 0.129 | 0.413 | 0.413 |
| 1237 | EPZ004777 | EPZ-004777 | 19243 | 0.973 | 0.973 | 5077.5 | 0.234 | 0.625 | 16409.5 | 0.319 | 0.635 | 0.635 |
| 1239 | YK-4-279 | YK 4-279 | 13430 | 0.059 | 0.196 | 5862 | 0.589 | 1.000 | 18176.5 | 0.784 | 1.000 | 1.000 |
| 1243 | Piperlongumine | Piplartine | 534 | 0.872 | 0.993 | 138 | 0.822 | 0.993 | 379 | 0.392 | 0.896 | 0.896 |
| 1248 | Daporinad | APO866, FK866, FK866 | 6336 | 0.669 | 1.000 | 2037 | 0.671 | 1.000 | 5734 | 0.448 | 1.000 | 1.000 |
| 1249 | BMS-345541 |  | 13684 | 0.165 | 0.439 | 4796 | 0.638 | 0.937 | 12454 | 0.072 | 0.291 | 0.291 |
| 1250 | AZ960 | AZ 960, AZ-960 | 18717 | 0.997 | 1.000 | 4517 | 0.486 | 0.777 | 16262.5 | 0.925 | 1.000 | 1.000 |
| 1259 | Talazoparib | BMN-673, BMN 973 | 16413 | 0.710 | 1.000 | 5125 | 0.265 | 0.605 | 19694 | 0.981 | 1.000 | 1.000 |
| 1268 | XAV939 | NVP-XAV939, XAV-939, XAV 939 | 18175.5 | 0.991 | 1.000 | 4085 | 0.267 | 0.583 | 13665 | 0.291 | 0.583 | 0.583 |
| 1372 | Trametinib | GSK1120212, Mekinist | 6766 | 0.000 | 0.000 | 3804 | 0.013 | 0.069 | 14393 | 0.032 | 0.085 | 0.085 |
| 1373 | Dabrafenib | GSK2118436, Tafinlar | 14606.5 | 0.185 | 0.494 | 3206 | 0.002 | 0.011 | 15811.5 | 0.304 | 0.694 | 0.694 |
| 1375 | Temozolomide | Temodar, Temodal, M-39831, SCH 52365 | 18804.5 | 0.969 | 1.000 | 5114 | 0.245 | 0.560 | 14915 | 0.068 | 0.182 | 0.182 |
| 1378 | Bleomycin (50 uM) |  | 11804.5 | 0.619 | 1.000 | 3564 | 0.198 | 0.453 | 13807 | 0.985 | 1.000 | 1.000 |
| 1401 | AZD5438 | SN1057898678 | 11826 | 0.480 | 0.888 | 3887 | 0.784 | 0.972 | 9785 | 0.098 | 0.392 | 0.392 |
| 1428 | IAP_5620 | SN1043546339 | 12077 | 0.589 | 0.824 | 3447 | 0.519 | 0.824 | 10284 | 0.219 | 0.809 | 0.809 |
| 1441 | AZD2014 | SN1103949359 | 13452 | 0.928 | 0.999 | 3529 | 0.567 | 0.825 | 9219 | 0.033 | 0.309 | 0.309 |
| 1449 | AZD1208 | SN1028350371 | 10788.5 | 0.160 | 0.512 | 3703 | 0.685 | 0.965 | 10919.5 | 0.427 | 0.854 | 0.854 |
| 1463 | AZD1332 | SN1061387896 | 13797 | 0.963 | 0.999 | 3972 | 0.825 | 0.999 | 10774 | 0.366 | 0.651 | 0.651 |
| 1494 | SN-38 | 7-Ethyl-10-Hydroxy-Camptothecin, SN 38 | 19294 | 0.990 | 1.000 | 6107 | 0.702 | 1.000 | 20171.5 | 0.985 | 1.000 | 1.000 |
| 1502 | Bicalutamide | ICI-176334, Casodex, Cosudex, ICI 176334 | 579 | 0.941 | 0.995 | 20 | 0.124 | 0.663 | 372 | 0.368 | 0.665 | 0.665 |
| 1507 | Ruxolitinib | INCB-18424, Ruxolitinib Phosphate, Jakafi | 16352 | 0.825 | 1.000 | 4161 | 0.290 | 0.664 | 12699 | 0.089 | 0.357 | 0.357 |
| 1510 | Linsitinib | OSI-906, ASP-7487 | 10431 | 0.000 | 0.000 | 5856 | 0.565 | 0.904 | 16834.5 | 0.415 | 0.738 | 0.738 |
| 1511 | Epirubicin | Ellence, Farmorubicin, IMI-28 | 18786 | 0.986 | 1.000 | 5414 | 0.382 | 0.645 | 18878 | 0.896 | 1.000 | 1.000 |
| 1512 | Cyclophosphami | Cytoxan, Neosar, Cyclophosphamid, Procytox, Cyclophosphane | 17827.5 | 0.939 | 1.000 | 3164 | 0.002 | 0.027 | 15671.5 | 0.186 | 0.446 | 0.446 |
| 1529 | Pevonedistat | MLN4924, MLN 4924, MLN-4924 | 17450 | 0.903 | 0.967 | 4951 | 0.204 | 0.545 | 18348.5 | 0.826 | 0.967 | 0.967 |
| 1549 | Sapitinib | AZD8931 | 14358 | 0.076 | 0.203 | 7205.5 | 0.959 | 1.000 | 16342 | 0.298 | 0.596 | 0.596 |
| 1553 | Uprosertib | GSK2141795, GSK2141795C, GSK-2141795 | 13792.5 | 0.170 | 0.550 | 6428 | 0.994 | 0.999 | 13205 | 0.172 | 0.550 | 0.550 |
| 1557 | LCL161 |  | 17429 | 0.965 | 1.000 | 5006 | 0.736 | 1.000 | 13287 | 0.201 | 0.643 | 0.643 |
| 1558 | Lapatinib | Tykerb, Tyverb | 13852 | 0.059 | 0.238 | 7449 | 0.979 | 1.000 | 15749 | 0.177 | 0.566 | 0.566 |
| 1559 | Luminespib | AUY922, VER-52296,NVP-AUY922, AUY | 18640 | 0.938 | 1.000 | 4727 | 0.130 | 0.413 | 17380 | 0.579 | 1.000 | 1.000 |
| 1560 | Alpelisib | BYL719, BYL-719, NVP-BYL719 | 15917.5 | 0.364 | 0.728 | 7709 | 0.990 | 0.993 | 16327.5 | 0.300 | 0.728 | 0.728 |
| 1561 | Taselisib | GDC-0032, GDC0032, RG7604 | 14041 | 0.048 | 0.193 | 7790 | 0.992 | 1.000 | 17526 | 0.598 | 1.000 | 1.000 |
| 1563 | EPZ5676 | EPZ-5676, Pinometostat | 18203 | 0.892 | 1.000 | 6040 | 0.658 | 1.000 | 14331 | 0.032 | 0.128 | 0.128 |
| 1564 | SCH772984 | CS-1421 | 7482 | 0.000 | 0.000 | 4246 | 0.045 | 0.103 | 13988 | 0.018 | 0.058 | 0.058 |
| 1576 | IWP-2 | Wnt Inhibitor IWP-2 | 16356 | 0.844 | 0.999 | 3911 | 0.195 | 0.495 | 12986.5 | 0.143 | 0.495 | 0.495 |
| 1578 | Leflunomide | Sulol, SU-101, HWA-486, Arava | 15395 | 0.614 | 0.983 | 2903 | 0.013 | 0.178 | 12384 | 0.063 | 0.202 | 0.202 |
| 1580 | GSK2801 | 1-(7-(2-methylsulfonyl-phenyl)-4-propoxy-1-aza-bicyclo[4.3.0]nona-2,4,6,8-tetraen-9-yl)-ethanone | 10867 | 0.169 | 0.404 | 2707 | 0.104 | 0.404 | 8085 | 0.006 | 0.096 | 0.096 |
| 1581 | Bromosporine | ethyl N-[6-(3-methanesulfonamido-4-methylphenyl)-3-methyl-[1,2,4]triazolo[4,3-b]pyridazin-8-yl]carbamate | 12863.5 | 0.820 | 0.968 | 2771 | 0.127 | 0.507 | 9322 | 0.100 | 0.507 | 0.507 |
| 1582 | SGC-CBP30 | 8-(3-chloro-4-methoxy-phenethyl)-4-(3,5-dimethyl-isoxazol-4-yl)-9-(2-(morpholin-4-yl)-propyl)-7,9-diaza-bicyclo[4.3.0]nona-1(6),2,4,7-tetraene | 13976 | 0.974 | 0.974 | 2831 | 0.150 | 0.598 | 8425 | 0.015 | 0.244 | 0.244 |

| DRUG_ID | DRUG_NAME | SYNONYMS | nervous_system_MWU_STAT | nervous_system_PVAL | nervous_system_PVAL_ADJ | soft_tissue_MWU_STAT | soft_tissue_PVAL | soft_tissue_PVAL_ADJ | bone_MWU_STAT | bone_PVAL | bone_PVAL_ADJ | skin_MWU_STAT | skin_PVAL | skin_PVAL_ADJ | urogenital_system_MWU_STAT | urogenital_system_PVAL | urogenital_system_PVAL_ADJ | lung_NSCLC_MWU_STAT | lung_NSCLC_PVAL | lung_NSCLC_PVAL_ADJ | lung_SCLC_MWU_STAT | lung_SCLC_PVAL | lung_SCLC_PVAL_ADJ | lung_MWU_STAT | lung_PVAL | lung_PVAL_ADJ | neuroblastoma_MWU_STAT | neuroblastoma_PVAL | neuroblastoma_PVAL_ADJ | pancreas_MWU_STAT | pancreas_PVAL | pancreas_PVAL_ADJ | aero_dig_tract_MWU_STAT | aero_dig_tract_PVAL | aero_dig_tract_PVAL_ADJ | breast_MWU_STAT | breast_PVAL | breast_PVAL_ADJ | kidney_MWU_STAT | kidney_PVAL | kidney_PVAL_ADJ |
|---|---|---|---|---|---|---|---|---|---|---|---|---|---|---|---|---|---|---|---|---|---|---|---|---|---|---|---|---|---|---|---|---|---|---|---|---|---|---|---|---|---|
| 1583 | GSK-LSD1 | N-((1R,2S)-2-phenylcyclopropyl)piperidin-4-amine | 8888.5 | 0.315 | 0.592 | 4489 | 0.467 | 0.680 | 10194 | 0.962 | 0.998 | 9874 | 0.923 | 0.998 | 19146.5 | 0.283 | 0.592 | 18468.5 | 0.089 | 0.308 | 12076.5 | 0.998 | 0.998 | 4297 | 0.070 | 0.308 | 6710 | 0.968 | 0.998 | 7071 | 0.414 | 0.663 | 13989 | 0.197 | 0.526 | 14502 | 0.998 | 0.998 | 4419 | 0.050 | 0.308 |
| 1584 | BDOCA000347a | 4-bromo-N-(6-methoxy-1,3-dimethyl-2-oxo-2,3-dihydro-1H-benzo[d]imidazol-5-yl)-2-methylbenzenesulfonamide | 9529 | 0.567 | 0.958 | 4839 | 0.667 | 0.970 | 11399 | 0.997 | 0.999 | 11384 | 0.999 | 0.999 | 17889 | 0.067 | 0.267 | 17947 | 0.040 | 0.267 | 11980 | 0.997 | 0.999 | 5219 | 0.414 | 0.917 | 6904 | 0.982 | 0.999 | 6399 | 0.152 | 0.404 | 13066 | 0.051 | 0.267 | 10172 | 0.122 | 0.390 | 5572.5 | 0.459 | 0.917 |
| 1585 | BDF00022089a | | 8952.5 | 0.333 | 0.665 | 5424 | 0.905 | 1.000 | 10350.5 | 0.947 | 1.000 | 10320.5 | 0.971 | 1.000 | 20089 | 0.529 | 0.847 | 16221 | 0.001 | 0.024 | 13101.5 | 1.000 | 1.000 | 4260 | 0.062 | 0.238 | 7171 | 0.993 | 1.000 | 5278 | 0.009 | 0.073 | 14367 | 0.287 | 0.657 | 13315 | 0.965 | 1.000 | 4507 | 0.062 | 0.238 |
| 1586 | BDILV000379a | 4-(4-(1-methyl-1H-pyrazol-4-yl)-1-(2-(1-methyl-1H-pyrazol-4-yl)ethyl)-4,5-dihydro-1H-imidazol-5-yl)benzonitrile | 9698 | 0.636 | 0.999 | 4695 | 0.585 | 0.999 | 10038 | 0.900 | 0.999 | 9306.5 | 0.784 | 0.999 | 20173.5 | 0.553 | 0.999 | 17162.5 | 0.011 | 0.147 | 12395 | 0.999 | 0.999 | 4182 | 0.050 | 0.265 | 7699 | 0.999 | 0.999 | 5508 | 0.018 | 0.147 | 13635.5 | 0.122 | 0.391 | 12789.5 | 0.906 | 0.999 | 4757 | 0.115 | 0.391 |
| 1593 | Entinostat | MS-275 | 20841 | 0.999 | 1.000 | 6372 | 0.542 | 1.000 | 9188 | 0.011 | 0.044 | 11571 | 0.000 | 0.000 | 33150 | 0.995 | 1.000 | 30731 | 0.646 | 1.000 | 7023 | 0.000 | 0.000 | 8071 | 0.796 | 1.000 | 3336 | 0.000 | 0.000 | 12944 | 1.000 | 1.000 | 29207 | 1.000 | 1.000 | 16779 | 0.705 | 1.000 | 12581 | 0.924 | 1.000 |
| 1594 | OSI-027 | A-1065-5 | 18463 | 0.347 | 0.740 | 7202 | 0.370 | 0.740 | 12363 | 0.172 | 0.740 | 16265 | 0.015 | 0.117 | 32808.5 | 0.343 | 0.740 | 35548 | 0.537 | 0.781 | 25025.5 | 0.998 | 1.000 | 8685.5 | 0.660 | 0.880 | 10913 | 0.770 | 0.948 | 10787 | 0.279 | 0.740 | 29309 | 0.937 | 1.000 | 11668 | 0.000 | 0.000 | 11115.5 | 0.192 | 0.740 |
| 1598 | LGK974 | LGK-974 | 24458 | 1.000 | 1.000 | 7625 | 0.535 | 0.855 | 17242 | 0.997 | 1.000 | 29082 | 1.000 | 1.000 | 29367 | 0.014 | 0.044 | 30353 | 0.009 | 0.036 | 18617 | 0.235 | 0.626 | 9561 | 0.894 | 1.000 | 12172 | 0.877 | 1.000 | 8448 | 0.006 | 0.032 | 26260.5 | 0.444 | 0.788 | 13048 | 0.001 | 0.015 | 14304 | 0.951 | 1.000 |
| 1613 | VE-822 | VE 822, VE822, Berzosertib | 18513.5 | 0.916 | 0.993 | 6786.5 | 0.731 | 0.993 | 9486 | 0.023 | 0.091 | 19730 | 0.962 | 0.993 | 27535 | 0.338 | 0.902 | 29590 | 0.440 | 0.993 | 18435 | 0.964 | 0.993 | 9505 | 0.992 | 0.993 | 5397 | 0.001 | 0.012 | 11824 | 0.993 | 0.993 | 20027.5 | 0.033 | 0.105 | 16809.5 | 0.908 | 0.993 | 7709 | 0.003 | 0.021 |
| 1614 | WZ4003 | WZ 4003 | 18807.5 | 0.944 | 0.999 | 5930 | 0.353 | 0.808 | 7763 | 0.000 | 0.004 | 14902 | 0.055 | 0.294 | 29262.5 | 0.700 | 0.933 | 28020 | 0.163 | 0.559 | 20096 | 0.999 | 0.999 | 7043.5 | 0.406 | 0.808 | 8038 | 0.289 | 0.772 | 9140 | 0.454 | 0.808 | 23823 | 0.674 | 0.933 | 18281.5 | 0.992 | 0.999 | 11255 | 0.624 | 0.933 |
| 1615 | CZC24832 | GTPL6653 | 14105 | 0.039 | 0.162 | 7632 | 0.945 | 1.000 | 11439 | 0.360 | 0.720 | 18448 | 0.811 | 1.000 | 25107 | 0.041 | 0.162 | 26106 | 0.023 | 0.162 | 22099 | 1.000 | 1.000 | 7296 | 0.513 | 0.912 | 10518 | 0.971 | 1.000 | 7666.5 | 0.064 | 0.172 | 25109 | 0.890 | 1.000 | 16154 | 0.800 | 1.000 | 8571 | 0.021 | 0.162 |
| 1617 | AZD5582 | GTPL7710, AOB5560 | 18400 | 0.985 | 1.000 | 3423 | 0.002 | 0.014 | 7716 | 0.000 | 0.005 | 16291 | 0.307 | 0.546 | 25963 | 0.133 | 0.355 | 28473.5 | 0.284 | 0.546 | 12621 | 0.010 | 0.040 | 9118 | 0.982 | 1.000 | 11646 | 0.999 | 1.000 | 11495 | 0.987 | 1.000 | 21418 | 0.196 | 0.447 | 10495 | 0.004 | 0.019 | 16339 | 1.000 | 1.000 |
| 1618 | GSK2606414 | AK175551 | 17624.5 | 0.784 | 1.000 | 6131 | 0.451 | 0.902 | 9542 | 0.027 | 0.143 | 19041 | 0.910 | 1.000 | 24334 | 0.024 | 0.143 | 29286 | 0.397 | 0.902 | 20357 | 1.000 | 1.000 | 7506.5 | 0.611 | 0.995 | 10755 | 0.985 | 1.000 | 9900 | 0.738 | 1.000 | 20699 | 0.082 | 0.326 | 18155 | 0.991 | 1.000 | 10088 | 0.247 | 0.659 |
| 1620 | PFI3 | PFI-3, PFI 3, AOB2221 | 14305 | 0.053 | 0.212 | 6760 | 0.721 | 0.961 | 11658 | 0.431 | 0.766 | 21001.5 | 0.996 | 1.000 | 27415 | 0.315 | 0.629 | 24602 | 0.003 | 0.041 | 21618.5 | 1.000 | 1.000 | 6301 | 0.150 | 0.399 | 11893.5 | 0.999 | 1.000 | 6989.5 | 0.015 | 0.081 | 23943 | 0.700 | 0.961 | 16702 | 0.894 | 1.000 | 9698.5 | 0.147 | 0.399 |
| 1621 | PCI-34051 | AK298746 | 17065.5 | 0.637 | 0.926 | 5059 | 0.083 | 0.304 | 9816.5 | 0.042 | 0.226 | 16294 | 0.267 | 0.533 | 28661 | 0.578 | 0.925 | 31068.5 | 0.737 | 0.982 | 15420 | 0.348 | 0.618 | 6515.5 | 0.210 | 0.480 | 3059 | 0.000 | 0.000 | 10943.5 | 0.946 | 1.000 | 29164 | 1.000 | 1.000 | 18611.5 | 0.996 | 1.000 | 9403 | 0.095 | 0.304 |
| 1622 | Wnt-C59 | AOB3540, AK161197 | 19224.5 | 0.970 | 0.996 | 7094 | 0.835 | 0.996 | 13430.5 | 0.906 | 0.996 | 19688 | 0.960 | 0.996 | 26001.5 | 0.105 | 0.280 | 25260.5 | 0.007 | 0.058 | 19565 | 0.996 | 0.996 | 7196 | 0.470 | 0.996 | 9049 | 0.672 | 0.996 | 7771 | 0.078 | 0.276 | 24765 | 0.846 | 0.996 | 13205 | 0.086 | 0.276 | 12112 | 0.857 | 0.996 |
| 1624 | I-BET-762 | GSK525762A | 15584 | 0.246 | 0.703 | 6000 | 0.388 | 0.703 | 12977 | 0.893 | 0.979 | 14946 | 0.061 | 0.484 | 26758.5 | 0.207 | 0.703 | 29748 | 0.483 | 0.703 | 18480 | 0.968 | 0.979 | 6566 | 0.229 | 0.703 | 9424 | 0.796 | 0.979 | 10485 | 0.881 | 0.979 | 22938 | 0.476 | 0.703 | 17715.5 | 0.979 | 0.979 | 10641 | 0.417 | 0.703 |
| 1625 | RVX-208 | 1044870-39-4, Apabetalone | 14605 | 0.083 | 0.265 | 5893 | 0.341 | 0.607 | 13228 | 0.927 | 1.000 | 15789 | 0.169 | 0.450 | 25260 | 0.051 | 0.250 | 26940 | 0.063 | 0.250 | 22733 | 1.000 | 1.000 | 6819 | 0.319 | 0.607 | 9851 | 0.895 | 1.000 | 9137 | 0.458 | 0.721 | 23022 | 0.496 | 0.721 | 17609 | 0.974 | 1.000 | 8446 | 0.016 | 0.250 |
| 1626 | OTX015 | Birabresib, OTX-015, MK-8628 | 14634 | 0.123 | 0.658 | 5693 | 0.261 | 0.668 | 12302 | 0.649 | 0.944 | 15077 | 0.072 | 0.579 | 27058.5 | 0.257 | 0.668 | 30416 | 0.623 | 0.944 | 17942 | 0.928 | 0.989 | 7796 | 0.720 | 0.960 | 9412 | 0.793 | 0.976 | 10389.5 | 0.862 | 0.985 | 22548 | 0.383 | 0.681 | 19114 | 0.999 | 0.999 | 10392 | 0.334 | 0.668 |
| 1627 | GSK343 | GTPL8240, AOB3680, AK175558 | 14356 | 0.057 | 0.230 | 7194 | 0.862 | 1.000 | 14276.5 | 0.979 | 1.000 | 17286 | 0.528 | 0.845 | 25534.5 | 0.066 | 0.230 | 24844 | 0.004 | 0.061 | 23121.5 | 1.000 | 1.000 | 6866.5 | 0.334 | 0.594 | 10874.5 | 0.988 | 1.000 | 7725 | 0.072 | 0.230 | 22089 | 0.276 | 0.552 | 17957 | 0.986 | 1.000 | 9377 | 0.091 | 0.243 |
| 1629 | ML323 | GTPL7898, AOB2313, | 20112 | 0.994 | 0.999 | 7045.5 | 0.820 | 0.999 | 14562 | 0.988 | 0.999 | 21801 | 0.999 | 0.999 | 24683.5 | 0.024 | 0.195 | 28587.5 | 0.247 | 0.791 | 17941 | 0.925 | 0.999 | 7718.5 | 0.687 | 0.999 | 9956 | 0.910 | 0.999 | 8812 | 0.335 | 0.892 | 20785 | 0.083 | 0.334 | 12688 | 0.040 | 0.213 | 10929 | 0.512 | 0.999 |
| 1630 | Entospletinib | GS-9973 | 12731 | 0.003 | 0.025 | 6858 | 0.758 | 1.000 | 10271.5 | 0.090 | 0.206 | 18224 | 0.766 | 1.000 | 26994 | 0.238 | 0.477 | 26440 | 0.034 | 0.091 | 24280.5 | 1.000 | 1.000 | 6902 | 0.348 | 0.619 | 11944 | 1.000 | 1.000 | 10385 | 0.858 | 1.000 | 19565 | 0.017 | 0.068 | 20242 | 1.000 | 1.000 | 7113 | 0.000 | 0.007 |
| 1631 | PRT062607 | P505-15, PRT-2607, BIIB-057 | 19200.5 | 0.969 | 1.000 | 6077 | 0.418 | 0.843 | 8323 | 0.001 | 0.012 | 11885 | 0.000 | 0.002 | 32277 | 0.985 | 1.000 | 27218 | 0.079 | 0.254 | 15871 | 0.474 | 0.843 | 6149 | 0.115 | 0.306 | 8722 | 0.548 | 0.877 | 10218 | 0.819 | 1.000 | 22938 | 0.467 | 0.843 | 21481 | 1.000 | 1.000 | 8795 | 0.033 | 0.153 |
| 1632 | Ribociclib | LEE011, NVP-LEE011, LEE011-BBA | 17220.5 | 0.127 | 0.328 | 8107 | 0.732 | 0.892 | 10129 | 0.005 | 0.040 | 19762.5 | 0.525 | 0.742 | 35106 | 0.781 | 0.892 | 34241 | 0.327 | 0.655 | 29529.5 | 1.000 | 1.000 | 8129 | 0.462 | 0.739 | 7225 | 0.007 | 0.040 | 11626 | 0.556 | 0.742 | 33253.5 | 1.000 | 1.000 | 15701.5 | 0.138 | 0.328 | 9116.5 | 0.007 | 0.040 |
| 1634 | AGI-6780 | AK162157 | 20387.5 | 0.997 | 1.000 | 6271 | 0.512 | 0.819 | 10352 | 0.104 | 0.332 | 23365 | 1.000 | 1.000 | 25761 | 0.087 | 0.332 | 26662 | 0.046 | 0.247 | 15967.5 | 0.509 | 0.819 | 7777 | 0.713 | 0.951 | 8958 | 0.643 | 0.936 | 8341 | 0.194 | 0.498 | 24458.5 | 0.804 | 0.990 | 12177.5 | 0.028 | 0.227 | 13547 | 0.991 | 1.000 |
| 1635 | Picolinici-acid | Picolinate | 15925 | 0.328 | 0.691 | 6680.5 | 0.693 | 1.000 | 13746 | 0.945 | 1.000 | 21423 | 0.999 | 1.000 | 23899 | 0.009 | 0.047 | 27517 | 0.110 | 0.437 | 21564 | 1.000 | 1.000 | 6958 | 0.375 | 0.691 | 11050 | 0.993 | 1.000 | 6173 | 0.002 | 0.015 | 23350.5 | 0.575 | 0.919 | 16231.5 | 0.877 | 1.000 | 10558 | 0.389 | 0.691 |
| 1706 | AZD5153 | SN1045566955, AZD5153 | 9302 | 0.480 | 0.916 | 4071 | 0.239 | 0.637 | 9165 | 0.742 | 0.916 | 7536 | 0.195 | 0.624 | 18679 | 0.182 | 0.624 | 20643 | 0.577 | 0.916 | 9698 | 0.642 | 0.916 | 5564 | 0.607 | 0.916 | 5124 | 0.368 | 0.842 | 7967 | 0.802 | 0.916 | 16536 | 0.928 | 0.985 | 13672 | 0.985 | 0.985 | 6258 | 0.798 | 0.916 |
| 1708 | CDK9_5576 | SN1047226266, CDK9_5576 | 9404 | 0.530 | 0.771 | 4223 | 0.319 | 0.771 | 6474 | 0.021 | 0.237 | 6692 | 0.037 | 0.237 | 19386.5 | 0.354 | 0.771 | 23597 | 0.990 | 0.992 | 9383.5 | 0.521 | 0.771 | 7051 | 0.990 | 0.992 | 4313 | 0.074 | 0.237 | 9269 | 0.992 | 0.992 | 13020 | 0.074 | 0.237 | 11293 | 0.474 | 0.771 | 5895 | 0.640 | 0.853 |
| 1709 | CDK9_5038 | SN1047483750, CDK9_5038 | 9327 | 0.498 | 0.798 | 3575 | 0.074 | 0.230 | 5522 | 0.001 | 0.016 | 6518 | 0.024 | 0.145 | 19903 | 0.499 | 0.798 | 22982 | 0.970 | 1.000 | 8033 | 0.086 | 0.230 | 6879 | 0.982 | 1.000 | 4959 | 0.291 | 0.623 | 9814 | 0.999 | 1.000 | 12450 | 0.027 | 0.145 | 10844 | 0.312 | 0.623 | 6356 | 0.837 | 1.000 |
| 1712 | Eg5_9814 | SN1047613775, Eg5_9814 | 9592 | 0.614 | 0.894 | 3647 | 0.092 | 0.422 | 7611 | 0.154 | 0.422 | 7732 | 0.271 | 0.542 | 21202 | 0.831 | 1.000 | 20328 | 0.511 | 0.894 | 8095 | 0.100 | 0.422 | 7711.5 | 1.000 | 1.000 | 3287 | 0.002 | 0.034 | 9325 | 0.994 | 1.000 | 13529 | 0.158 | 0.422 | 10264 | 0.153 | 0.422 | 7221 | 0.985 | 1.000 |
| 1713 | ERK_2440 | SN1051032892, ERK_2440 | 8884 | 0.313 | 0.715 | 4328 | 0.373 | 0.746 | 12065 | 1.000 | 1.000 | 2492 | 0.000 | 0.000 | 21904 | 0.920 | 1.000 | 21085.5 | 0.694 | 1.000 | 13485.5 | 1.000 | 1.000 | 5363 | 0.498 | 0.807 | 5375 | 0.504 | 0.807 | 6605 | 0.220 | 0.608 | 16073 | 0.859 | 1.000 | 14731 | 0.999 | 1.000 | 5087 | 0.228 | 0.608 |
| 1714 | ERK_6604 | SN1047587618, ERK_6604 | 11133 | 0.969 | 1.000 | 5362 | 0.891 | 1.000 | 12710 | 1.000 | 1.000 | 2747 | 0.000 | 0.000 | 21818 | 0.910 | 1.000 | 17597.5 | 0.025 | 0.065 | 13350 | 1.000 | 1.000 | 4831 | 0.229 | 0.524 | 6008 | 0.812 | 1.000 | 5223.5 | 0.008 | 0.032 | 14047 | 0.270 | 0.540 | 16379 | 1.000 | 1.000 | 7154 | 0.979 | 1.000 |
| 1716 | IRAK4_4710 | SN1050852374, IRAK4_4710 | 9452 | 0.543 | 0.788 | 5373 | 0.894 | 0.998 | 9568.5 | 0.862 | 0.998 | 10611 | 0.995 | 0.998 | 18289.5 | 0.117 | 0.469 | 20451 | 0.523 | 0.788 | 9569.5 | 0.591 | 0.788 | 3856 | 0.018 | 0.294 | 5469 | 0.556 | 0.788 | 6156.5 | 0.096 | 0.469 | 13764 | 0.198 | 0.633 | 14441.5 | 0.998 | 0.998 | 5193 | 0.274 | 0.729 |
| 1718 | JAK1_8709 | SN1049200060, JAK1_8709 | 10467 | 0.879 | 0.946 | 4859 | 0.682 | 0.946 | 6106 | 0.004 | 0.028 | 8375 | 0.529 | 0.946 | 19769 | 0.449 | 0.946 | 19027 | 0.170 | 0.681 | 9857 | 0.702 | 0.946 | 3269 | 0.002 | 0.028 | 4833 | 0.230 | 0.737 | 7996 | 0.811 | 0.946 | 14529 | 0.416 | 0.946 | 14766 | 0.999 | 0.999 | 5867 | 0.620 | 0.946 |
| 1720 | AZD5991 | SN1049446612, AZD5991 | 12208 | 0.999 | 0.999 | 5246 | 0.861 | 0.999 | 6282 | 0.007 | 0.035 | 8189 | 0.577 | 0.999 | 20477 | 0.669 | 0.999 | 22735 | 0.959 | 0.999 | 7486 | 0.027 | 0.109 | 6312 | 0.909 | 0.999 | 3332 | 0.003 | 0.021 | 9637 | 0.998 | 0.999 | 12869 | 0.060 | 0.192 | 6316 | 0.000 | 0.000 | 7466 | 0.994 | 0.999 |
| 1730 | PAK_5339 | SN1046290829, PAK_5339 | 11685 | 0.993 | 0.993 | 4737 | 0.619 | 0.993 | 7907 | 0.239 | 0.636 | 9311.5 | 0.871 | 0.993 | 17475 | 0.040 | 0.163 | 22268.5 | 0.938 | 0.993 | 10444 | 0.878 | 0.993 | 5638 | 0.651 | 0.993 | 4104 | 0.041 | 0.163 | 9090 | 0.986 | 0.993 | 12515 | 0.031 | 0.163 | 11818 | 0.668 | 0.993 | 4418 | 0.051 | 0.163 |
| 1732 | TAF1_5496 | SN1050393042, TAF1_5496 | 11617 | 0.992 | 1.000 | 4724 | 0.612 | 1.000 | 10811 | 0.993 | 1.000 | 11593 | 1.000 | 1.000 | 17585 | 0.064 | 0.204 | 20747 | 0.616 | 1.000 | 6380 | 0.001 | 0.016 | 6439 | 0.933 | 1.000 | 5591 | 0.627 | 1.000 | 6472 | 0.180 | 0.480 | 15392 | 0.702 | 1.000 | 9185 | 0.019 | 0.103 | 6616 | 0.909 | 1.000 |
| 1733 | ULK1_4989 | SN1049257073, ULK1_4989 | 7586 | 0.033 | 0.107 | 3058 | 0.013 | 0.052 | 4482 | 0.000 | 0.000 | 10554 | 0.994 | 1.000 | 18331 | 0.129 | 0.343 | 23893 | 0.994 | 1.000 | 9569 | 0.598 | 1.000 | 6238 | 0.889 | 1.000 | 6050 | 0.915 | 1.000 | 9563 | 0.997 | 1.000 | 11995 | 0.011 | 0.052 | 7986 | 0.001 | 0.005 | 6985 | 0.966 | 1.000 |
| 1734 | VSP34_8731 | SN1049998399, VSP34_8731 | 10704.5 | 0.922 | 1.000 | 3270 | 0.028 | 0.190 | 8475 | 0.458 | 0.814 | 8990 | 0.775 | 1.000 | 19552 | 0.388 | 0.777 | 19941.5 | 0.380 | 0.777 | 12549 | 1.000 | 1.000 | 6448 | 0.932 | 1.000 | 6680 | 0.965 | 1.000 | 8014 | 0.817 | 1.000 | 12547 | 0.031 | 0.190 | 10757 | 0.277 | 0.777 | 4628 | 0.086 | 0.345 |
| 1736 | Selumetinib | SN1103949345, AZD1480 | 10860 | 0.943 | 1.000 | 5997 | 0.986 | 1.000 | 13764 | 1.000 | 1.000 | 2148 | 0.000 | 0.000 | 22462 | 0.964 | 1.000 | 18301.5 | 0.071 | 0.227 | 15070 | 1.000 | 1.000 | 5757 | 0.705 | 1.000 | 5701 | 0.678 | 1.000 | 4203 | 0.000 | 0.001 | 13865.5 | 0.222 | 0.508 | 15140 | 1.000 | 1.000 | 5822 | 0.597 | 1.000 |
| 1738 | IGF1R_3801 | SN1051640224, IGF1R_3801 | 8990 | 0.353 | 0.807 | 3446 | 0.050 | 0.198 | 4088 | 0.000 | 0.000 | 12103 | 1.000 | 1.000 | 22530 | 0.968 | 1.000 | 20070 | 0.415 | 0.831 | 8766 | 0.271 | 0.721 | 4126 | 0.043 | 0.198 | 1884 | 0.000 | 0.000 | 7819 | 0.749 | 1.000 | 13520 | 0.146 | 0.468 | 15048 | 1.000 | 1.000 | 6148 | 0.754 | 1.000 |
| 1739 | JAK_8517 | SN1066590414, JAK_8517 | 7552 | 0.030 | 0.120 | 4181.5 | 0.293 | 0.670 | 4907 | 0.000 | 0.000 | 11726 | 1.000 | 1.000 | 22017.5 | 0.931 | 1.000 | 19298 | 0.223 | 0.595 | 8347 | 0.147 | 0.470 | 3600 | 0.007 | 0.039 | 3055 | 0.001 | 0.006 | 7517 | 0.623 | 1.000 | 15303 | 0.667 | 1.000 | 13896 | 0.992 | 1.000 | 6132 | 0.747 | 1.000 |
| 1776 | GSK2256098C | | 8812.5 | 0.281 | 0.643 | 4480 | 0.457 | 0.853 | 9644 | 0.804 | 0.919 | 9191.5 | 0.746 | 0.919 | 20264.5 | 0.579 | 0.919 | 18294.5 | 0.067 | 0.298 | 12590.5 | 1.000 | 1.000 | 4005 | 0.029 | 0.298 | 5972 | 0.794 | 0.919 | 6233.5 | 0.110 | 0.298 | 14986.5 | 0.480 | 0.853 | 13557.5 | 0.979 | 1.000 | 4510 | 0.063 | 0.298 |
| 1777 | GSK2276186C | | 9134 | 0.404 | 0.923 | 5086 | 0.789 | 0.986 | 9072.5 | 0.596 | 0.986 | 10387 | 0.975 | 0.986 | 18540 | 0.150 | 0.601 | 19163 | 0.188 | 0.603 | 11204 | 0.973 | 0.986 | 3875 | 0.019 | 0.102 | 6234 | 0.882 | 0.986 | 7521.5 | 0.619 | 0.986 | 14664 | 0.376 | 0.923 | 13713 | 0.986 | 0.986 | 3585.5 | 0.003 | 0.026 |
| 1778 | GSK2110183B | | 9150.5 | 0.411 | 0.926 | 4871.5 | 0.684 | 0.998 | 8628 | 0.407 | 0.926 | 9994.5 | 0.938 | 0.998 | 20400.5 | 0.617 | 0.998 | 20279.5 | 0.463 | 0.926 | 10682 | 0.915 | 0.998 | 6009 | 0.809 | 0.998 | 7488 | 0.998 | 0.998 | 9604 | 0.997 | 0.998 | 13280 | 0.072 | 0.301 | 7996 | 0.001 | 0.010 | 4582 | 0.075 | 0.301 |
| 1779 | GSK626616AC | | 9122 | 0.399 | 0.710 | 4305 | 0.356 | 0.710 | 9828.5 | 0.854 | 0.995 | 9975 | 0.935 | 0.995 | 19366 | 0.328 | 0.710 | 17320.5 | 0.014 | 0.229 | 11002.5 | 0.956 | 0.995 | 4420 | 0.093 | 0.374 | 6224 | 0.879 | 0.995 | 5882 | 0.050 | 0.322 | 15450.5 | 0.630 | 0.995 | 14101 | 0.995 | 0.995 | 4497 | 0.060 | 0.322 |
| 1780 | GSK3337463A | | 11193.5 | 0.972 | 0.972 | 4167 | 0.282 | 0.752 | 10308 | 0.942 | 0.972 | 9700 | 0.888 | 0.972 | 19866 | 0.466 | 0.931 | 17222.5 | 0.012 | 0.191 | 9939 | 0.725 | 0.972 | 4549.5 | 0.127 | 0.508 | 6097 | 0.840 | 0.972 | 6725 | 0.260 | 0.752 | 13530.5 | 0.105 | 0.508 | 12943 | 0.928 | 0.972 | 5353 | 0.345 | 0.788 |
| 1782 | GSK2830371A | | 10357 | 0.850 | 0.984 | 4532 | 0.488 | 0.722 | 9940 | 0.880 | 0.984 | 9889.5 | 0.923 | 0.984 | 19601 | 0.391 | 0.696 | 19609 | 0.285 | 0.570 | 8699 | 0.243 | 0.570 | 4104 | 0.040 | 0.559 | 4593 | 0.140 | 0.559 | 6318.5 | 0.130 | 0.559 | 15762 | 0.722 | 0.963 | 14582 | 0.999 | 0.999 | 5096.5 | 0.228 | 0.570 |
| 1783 | LMB_AB1 | | 10762.5 | 0.928 | 0.981 | 4153 | 0.275 | 0.622 | 9850 | 0.860 | 0.981 | 9544 | 0.852 | 0.981 | 18885 | 0.215 | 0.622 | 17078 | 0.009 | 0.146 | 10011 | 0.750 | 0.981 | 5012 | 0.307 | 0.622 | 6597 | 0.954 | 0.981 | 5773 | 0.038 | 0.305 | 15123.5 | 0.525 | 0.933 | 13596 | 0.981 | 0.981 | 5283 | 0.311 | 0.622 |
| 1784 | LMB_AB2 | | 10061.5 | 0.766 | 0.993 | 4679 | 0.576 | 0.993 | 10228 | 0.931 | 0.993 | 9100.5 | 0.713 | 0.993 | 18687 | 0.176 | 0.564 | 16853 | 0.006 | 0.094 | 11511.5 | 0.988 | 0.993 | 4655 | 0.160 | 0.564 | 6856 | 0.979 | 0.993 | 5612 | 0.025 | 0.198 | 14552 | 0.341 | 0.683 | 13994 | 0.993 | 0.993 | 5083 | 0.223 | 0.594 |
| 1785 | LMB_AB3 | | 10855.5 | 0.940 | 0.962 | 5288 | 0.865 | 0.962 | 9830 | 0.855 | 0.962 | 8730 | 0.562 | 0.962 | 20325.5 | 0.596 | 0.962 | 18518 | 0.090 | 0.480 | 10492 | 0.880 | 0.962 | 2956.5 | 0.000 | 0.007 | 6660 | 0.962 | 0.962 | 6961 | 0.358 | 0.962 | 15795.5 | 0.731 | 0.962 | 11979 | 0.708 | 0.962 | 5477 | 0.408 | 0.962 |
| 1786 | AZD4547 | 1035270-39-3 | 18163 | 0.280 | 0.754 | 8321 | 0.789 | 0.958 | 11601 | 0.064 | 0.257 | 21917.5 | 0.913 | 0.958 | 35992 | 0.837 | 0.958 | 35823 | 0.588 | 0.958 | 22270 | 0.943 | 0.958 | 4063 | 0.000 | 0.000 | 10109 | 0.283 | 0.754 | 13606 | 0.958 | 0.958 | 29093 | 0.944 | 0.958 | 19679 | 0.900 | 0.958 | 8703.5 | 0.003 | 0.014 |
| 1799 | Ibrutinib | PCI-32765, CRA-032765, PCI-32765-00 | 19224 | 0.974 | 1.000 | 8191 | 0.989 | 1.000 | 14972.5 | 0.996 | 1.000 | 19752 | 0.968 | 1.000 | 28373.5 | 0.546 | 1.000 | 25957 | 0.022 | 0.118 | 21485.5 | 1.000 | 1.000 | 7387 | 0.566 | 1.000 | 10692 | 0.983 | 1.000 | 11741 | 0.992 | 1.000 | 10259 | 0.000 | 0.000 | 11735 | 0.038 | 0.151 | 6549 | 0.000 | 0.001 |
| 1802 | Zoledronate | Zoledronic acid, Zometa, Reclast | 8983 | 0.344 | 0.633 | 4520 | 0.481 | 0.770 | 7672.5 | 0.103 | 0.330 | 10704.5 | 0.990 | 0.997 | 19471 | 0.356 | 0.633 | 17608.5 | 0.024 | 0.188 | 11576.5 | 0.990 | 0.997 | 4606.5 | 0.144 | 0.384 | 5626 | 0.635 | 0.923 | 5448 | 0.015 | 0.188 | 16359.5 | 0.860 | 0.997 | 14331 | 0.997 | 0.997 | 4554.5 | 0.070 | 0.330 |
| 1803 | Acetalax | Oxyphenisatin acetate | 12514 | 1.000 | 1.000 | 3915 | 0.170 | 0.453 | 9778 | 0.842 | 1.000 | 13131 | 1.000 | 1.000 | 16711 | 0.010 | 0.038 | 21715 | 0.822 | 1.000 | 15174 | 1.000 | 1.000 | 5403 | 0.515 | 1.000 | 8791 | 1.000 | 1.000 | 6303 | 0.126 | 0.404 | 9929 | 0.000 | 0.000 | 5024 | 0.000 | 0.000 | 6358 | 0.830 | 1.000 |
| 1804 | Acetalax | Oxyphenisatin acetate | 11745 | 0.994 | 1.000 | 4146 | 0.272 | 0.621 | 9024 | 0.575 | 1.000 | 12520 | 1.000 | 1.000 | 17810 | 0.060 | 0.208 | 22505 | 0.931 | 1.000 | 15374 | 1.000 | 1.000 | 5995 | 0.803 | 1.000 | 8879 | 1.000 | 1.000 | 5991 | 0.065 | 0.208 | 10424 | 0.000 | 0.001 | 4897 | 0.000 | 0.000 | 6270 | 0.798 | 1.000 |
| 1806 | Oxaliplatin | | 9432 | 0.527 | 0.924 | 5185 | 0.829 | 0.947 | 9477.5 | 0.751 | 0.924 | 9176 | 0.740 | 0.924 | 20003.5 | 0.505 | 0.924 | 22556 | 0.936 | 0.989 | 8058 | 0.086 | 0.588 | 5403 | 0.515 | 0.924 | 4888 | 0.250 | 0.800 | 7536 | 0.625 | 0.924 | 15504 | 0.647 | 0.924 | 13831 | 0.989 | 0.989 | 4739 | 0.110 | 0.588 |
| 1808 | Topotecan | | 8735 | 0.255 | 0.509 | 3624 | 0.082 | 0.322 | 5478 | 0.000 | 0.002 | 8926 | 0.645 | 1.000 | 17445 | 0.034 | 0.184 | 22644 | 0.943 | 1.000 | 8656.5 | 0.229 | 0.509 | 6073 | 0.832 | 1.000 | 2870.5 | 0.000 | 0.002 | 8617 | 0.946 | 1.000 | 14223 | 0.248 | 0.509 | 15349.5 | 1.000 | 1.000 | 5232 | 0.287 | 0.510 |
| 1809 | Teniposide | Vumon | 8417 | 0.160 | 0.367 | 3213.5 | 0.022 | 0.090 | 5912 | 0.001 | 0.006 | 9223 | 0.756 | 1.000 | 17559 | 0.041 | 0.132 | 22316 | 0.911 | 1.000 | 7057 | 0.008 | 0.042 | 6248 | 0.885 | 1.000 | 2835 | 0.000 | 0.004 | 8342 | 0.899 | 1.000 | 15270 | 0.573 | 1.000 | 15276 | 1.000 | 1.000 | 5930 | 0.647 | 1.000 |
| 1810 | Mitoxantrone | | 8504 | 0.184 | 0.576 | 3083 | 0.014 | 0.074 | 6015 | 0.001 | 0.009 | 8858 | 0.617 | 0.947 | 18007 | 0.078 | 0.313 | 21192.5 | 0.710 | 0.947 | 8615 | 0.216 | 0.576 | 4961 | 0.283 | 0.647 | 2548 | 0.000 | 0.001 | 8230 | 0.874 | 0.998 | 16403 | 0.868 | 0.998 | 15615 | 1.000 | 1.000 | 5943 | 0.654 | 0.947 |
| 1811 | Dactinomycin | Actinomycin D | 7878.5 | 0.060 | 0.291 | 3763 | 0.118 | 0.379 | 8923 | 0.532 | 0.710 | 7653 | 0.154 | 0.411 | 18774 | 0.193 | 0.441 | 21571.5 | 0.794 | 0.908 | 13695 | 1.000 | 1.000 | 5789 | 0.715 | 0.880 | 4911 | 0.260 | 0.521 | 7059.5 | 0.403 | 0.645 | 14427.5 | 0.305 | 0.541 | 15196 | 1.000 | 1.000 | 5544.5 | 0.444 | 0.646 |
| 1812 | Bleomycin | | 7974 | 0.073 | 0.372 | 3242 | 0.025 | 0.372 | 7851.5 | 0.142 | 0.372 | 8605 | 0.508 | 0.688 | 18156 | 0.095 | 0.372 | 18882.5 | 0.139 | 0.372 | 12974 | 1.000 | 1.000 | 4988 | 0.296 | 0.526 | 4664 | 0.163 | 0.372 | 8001 | 0.808 | 0.994 | 13235 | 0.067 | 0.372 | 16240 | 1.000 | 1.000 | 5636 | 0.493 | 0.688 |
| 1813 | Fludarabine | Fludara | 11307 | 0.979 | 0.993 | 3608 | 0.079 | 0.349 | 7008 | 0.024 | 0.191 | 6415 | 0.009 | 0.142 | 22168 | 0.941 | 0.993 | 18495 | 0.087 | 0.349 | 11715 | 0.993 | 0.993 | 5195 | 0.401 | 0.802 | 5766 | 0.705 | 0.939 | 6749 | 0.270 | 0.802 | 15984 | 0.780 | 0.960 | 11749.5 | 0.629 | 0.915 | 5613 | 0.480 | 0.829 |
| 1814 | Nelarabine | Atriance, Arranon | 10000.5 | 0.752 | 0.999 | 5357 | 0.889 | 0.999 | 9490 | 0.761 | 0.999 | 9664 | 0.884 | 0.999 | 19040 | 0.309 | 0.627 | 16775 | 0.005 | 0.049 | 12206 | 0.999 | 0.999 | 3557 | 0.006 | 0.049 | 7194 | 0.994 | 0.999 | 5980 | 0.065 | 0.347 | 14464.5 | 0.323 | 0.627 | 13841 | 0.990 | 0.999 | 5359 | 0.353 | 0.627 |
| 1815 | Dacarbazine | DTIC | 10020 | 0.753 | 0.976 | 5868 | 0.976 | 0.976 | 8368.5 | 0.303 | 0.606 | 10217.5 | 0.963 | 0.976 | 17769 | 0.056 | 0.390 | 18027.5 | 0.046 | 0.390 | 11011.5 | 0.957 | 0.976 | 6234 | 0.882 | 0.976 | 6785 | 0.974 | 0.976 | 6749 | 0.270 | 0.606 | 14403.5 | 0.298 | 0.606 | 13472 | 0.975 | 0.976 | 5359 | 0.348 | 0.618 |
| 1816 | Fulvestrant | | 12390 | 0.999 | 0.999 | 5295 | 0.868 | 0.992 | 8843 | 0.498 | 0.885 | 9057 | 0.696 | 0.951 | 22195 | 0.943 | 0.999 | 18996.5 | 0.158 | 0.537 | 9741 | 0.652 | 0.951 | 4501 | 0.113 | 0.537 | 5918 | 0.773 | 0.951 | 5616 | 0.025 | 0.401 | 14284.5 | 0.265 | 0.537 | 10605.5 | 0.225 | 0.537 | 6114 | 0.734 | 0.951 |
| 1817 | Romidepsin | Istodax, FK228, Depsipeptide | 7431 | 0.022 | 0.110 | 5380 | 0.893 | 1.000 | 5975 | 0.001 | 0.008 | 5483 | 0.000 | 0.006 | 22101 | 0.935 | 1.000 | 20089.5 | 0.410 | 0.729 | 8440 | 0.166 | 0.444 | 4827.5 | 0.224 | 0.513 | 6572 | 0.950 | 1.000 | 10078 | 1.000 | 1.000 | 12720 | 0.027 | 0.110 | 13220 | 0.957 | 1.000 | 7375 | 0.990 | 1.000 |

| DRUG_ID | DRUG_NAME | SYNONYMS | large_intestine_MWU_STAT | large_intestine_PVAL | large_intestine_PVAL_ADJ | thyroid_MWU_STAT | thyroid_PVAL | thyroid_PVAL_ADJ | digestive_system_MWU_STAT | digestive_system_PVAL | digestive_system_PVAL_ADJ | digestive_system_PVAL2 |
|---|---|---|---|---|---|---|---|---|---|---|---|---|
| 1583 | GSK-LSD1 | N-((1R,2S)-2-phenylcyclopropyl)piperidin-4-amine | 11415 | 0.333 | 0.592 | 2678 | 0.096 | 0.308 | 8458 | 0.017 | 0.265 | 0.265 |
| 1584 | BDOCA000347a | 4-bromo-N-(6-methoxy-1,3-dimethyl-2-oxo-2,3-dihydro-1H-benzo[d]imidazol-5-yl)-2-methylbenzenesulfonamide | 12781.5 | 0.793 | 0.999 | 3582 | 0.598 | 0.958 | 8566 | 0.021 | 0.267 | 0.267 |
| 1585 | BDF00022089a | | 11680.5 | 0.419 | 0.745 | 3018 | 0.235 | 0.625 | 9173.5 | 0.074 | 0.238 | 0.238 |
| 1586 | BDILV000379a | 4-(4-(1-methyl-1H-pyrazol-4-yl)-1-(2-(1-methyl-1H-pyrazol-4-yl)ethyl)-4,5-dihydro-1H-imidazol-5-yl)benzonitrile | 12525.5 | 0.719 | 0.999 | 3044 | 0.249 | 0.568 | 9620 | 0.157 | 0.419 | 0.419 |
| 1593 | Entinostat | MS-275 | 15290 | 0.557 | 1.000 | 5044 | 0.740 | 1.000 | 13694 | 0.362 | 1.000 | 1.000 |
| 1594 | OSI-027 | A-1065-5 | 21985 | 1.000 | 1.000 | 5631 | 0.475 | 0.775 | 17002 | 0.484 | 0.775 | 0.775 |
| 1598 | LGK974 | LGK-974 | 16069 | 0.410 | 0.788 | 5548 | 0.433 | 0.788 | 13345.5 | 0.006 | 0.032 | 0.032 |
| 1613 | VE-822 | VE 822, VE822, Berzosertib | 16193 | 0.808 | 0.993 | 4934 | 0.699 | 0.993 | 11343 | 0.017 | 0.090 | 0.090 |
| 1614 | WZ4003 | WZ 4003 | 16897 | 0.919 | 0.999 | 3861 | 0.175 | 0.559 | 10698 | 0.004 | 0.035 | 0.035 |
| 1615 | CZC24832 | GTPL6653 | 17365.5 | 0.960 | 1.000 | 4234 | 0.334 | 0.720 | 11968 | 0.051 | 0.162 | 0.162 |
| 1617 | AZD5582 | GTPL7710, AOB5560 | 18195 | 0.994 | 1.000 | 5690 | 0.948 | 1.000 | 11861.5 | 0.052 | 0.167 | 0.167 |
| 1618 | GSK2606414 | AK175551 | 15398 | 0.622 | 0.995 | 3968 | 0.219 | 0.659 | 9747 | 0.000 | 0.007 | 0.007 |
| 1620 | PFI3 | PFI-3, PFI 3, AOB2221 | 15781.5 | 0.713 | 0.961 | 3962.5 | 0.213 | 0.486 | 11186.5 | 0.012 | 0.081 | 0.081 |
| 1621 | PCI-34051 | AK298746 | 17907 | 0.984 | 1.000 | 3894 | 0.187 | 0.480 | 10936 | 0.007 | 0.059 | 0.059 |
| 1622 | Wnt-C59 | AOB3540, AK161197 | 16789.5 | 0.906 | 0.996 | 3538 | 0.085 | 0.276 | 9128.5 | 0.000 | 0.001 | 0.001 |
| 1624 | I-BET-762 | GSK525762A | 14476 | 0.345 | 0.703 | 2972 | 0.016 | 0.260 | 13429 | 0.310 | 0.703 | 0.703 |
| 1625 | RVX-208 | 1044870-39-4, Apabetalone | 17333 | 0.959 | 1.000 | 3258 | 0.040 | 0.250 | 13217 | 0.255 | 0.583 | 0.583 |
| 1626 | OTX015 | Birabresib, OTX-015, MK-8628 | 14425 | 0.331 | 0.668 | 2918 | 0.014 | 0.216 | 13038 | 0.213 | 0.668 | 0.668 |
| 1627 | GSK343 | GTPL8240, AOB3680, AK175558 | 16679.5 | 0.891 | 1.000 | 3156 | 0.029 | 0.230 | 12481 | 0.107 | 0.244 | 0.244 |
| 1629 | ML323 | GTPL7898, AOB2313, | 15108 | 0.523 | 0.999 | 4674 | 0.567 | 0.999 | 8415 | 0.000 | 0.000 | 0.000 |
| 1630 | Entospletinib | GS-9973 | 17733 | 0.978 | 1.000 | 3098 | 0.024 | 0.078 | 11037 | 0.009 | 0.049 | 0.049 |
| 1631 | PRT062607 | P505-15, PRT-2607, BIIB-057 | 17669 | 0.976 | 1.000 | 5196 | 0.810 | 1.000 | 11798.5 | 0.038 | 0.153 | 0.153 |
| 1632 | Ribociclib | LEE011, NVP-LEE011, LEE011-BBA | 12654 | 0.010 | 0.041 | 5408 | 0.383 | 0.681 | 15416 | 0.143 | 0.328 | 0.328 |
| 1634 | AGI-6780 | AK162157 | 14564 | 0.370 | 0.740 | 3970 | 0.218 | 0.498 | 10591 | 0.004 | 0.057 | 0.057 |
| 1635 | Picolinici-acid | Picolinate | 13543 | 0.136 | 0.437 | 4202.5 | 0.322 | 0.691 | 10329 | 0.002 | 0.015 | 0.015 |
| 1706 | AZD5153 | SN1045566955, AZD5153 | 10788 | 0.155 | 0.624 | 2207 | 0.017 | 0.274 | 9473 | 0.056 | 0.446 | 0.446 |
| 1708 | CDK9_5576 | SN1047226266, CDK9_5576 | 13340 | 0.916 | 0.992 | 3392 | 0.477 | 0.771 | 9397 | 0.050 | 0.237 | 0.237 |
| 1709 | CDK9_5038 | SN1047483750, CDK9_5038 | 15999 | 1.000 | 1.000 | 3616 | 0.629 | 0.915 | 9353 | 0.046 | 0.182 | 0.182 |
| 1712 | Eg5_9814 | SN1047613775, Eg5_9814 | 11819.5 | 0.586 | 0.894 | 3759 | 0.722 | 0.963 | 10456 | 0.271 | 0.542 | 0.542 |
| 1713 | ERK_2440 | SN1051032892, ERK_2440 | 7943 | 0.000 | 0.001 | 1907 | 0.004 | 0.022 | 9355 | 0.044 | 0.176 | 0.176 |
| 1714 | ERK_6604 | SN1047587618, ERK_6604 | 7473 | 0.000 | 0.000 | 1975 | 0.006 | 0.031 | 8875 | 0.015 | 0.049 | 0.049 |
| 1716 | IRAK4_4710 | SN1050852374, IRAK4_4710 | 11442 | 0.342 | 0.782 | 3287.5 | 0.402 | 0.788 | 9759 | 0.094 | 0.469 | 0.469 |
| 1718 | JAK1_8709 | SN1049200060, JAK1_8709 | 13181 | 0.887 | 0.946 | 4026 | 0.848 | 0.946 | 9756 | 0.093 | 0.498 | 0.498 |
| 1720 | AZD5991 | SN1049446612, AZD5991 | 13210 | 0.900 | 0.999 | 3686 | 0.839 | 0.999 | 10107 | 0.172 | 0.458 | 0.458 |
| 1730 | PAK_5339 | SN1046290829, PAK_5339 | 12503 | 0.726 | 0.993 | 3620 | 0.632 | 0.993 | 8646 | 0.009 | 0.142 | 0.142 |
| 1732 | TAF1_5496 | SN1050393042, TAF1_5496 | 9274 | 0.008 | 0.065 | 4359 | 0.947 | 1.000 | 9393 | 0.049 | 0.197 | 0.197 |
| 1733 | ULK1_4989 | SN1049257073, ULK1_4989 | 15768 | 1.000 | 1.000 | 3662 | 0.659 | 1.000 | 10398 | 0.247 | 0.564 | 0.564 |
| 1734 | VSP34_8731 | SN1049998399, VSP34_8731 | 11432 | 0.339 | 0.777 | 3444 | 0.509 | 0.814 | 9251 | 0.036 | 0.190 | 0.190 |
| 1736 | Selumetinib | SN1103949345, AZD1480 | 5984 | 0.000 | 0.000 | 1864 | 0.003 | 0.013 | 9744 | 0.092 | 0.244 | 0.244 |
| 1738 | IGF1R_3801 | SN1051640224, IGF1R_3801 | 12212 | 0.621 | 0.994 | 4788 | 0.990 | 1.000 | 11145 | 0.505 | 0.898 | 0.898 |
| 1739 | JAK_8517 | SN1066590414, JAK_8517 | 13352 | 0.914 | 1.000 | 4409 | 0.954 | 1.000 | 11390 | 0.598 | 1.000 | 1.000 |
| 1776 | GSK2256098C | | 12568 | 0.732 | 0.919 | 2713 | 0.106 | 0.298 | 9406 | 0.112 | 0.298 | 0.298 |
| 1777 | GSK2276186C | | 12963.5 | 0.838 | 0.986 | 4216 | 0.910 | 0.986 | 7870 | 0.003 | 0.026 | 0.026 |
| 1778 | GSK2110183B | | 10286 | 0.067 | 0.301 | 4246 | 0.919 | 0.998 | 9731.5 | 0.185 | 0.591 | 0.591 |
| 1779 | GSK626616AC | | 12955.5 | 0.836 | 0.995 | 3252 | 0.374 | 0.710 | 9527.5 | 0.136 | 0.436 | 0.436 |
| 1780 | GSK3337463A | | 13222 | 0.890 | 0.972 | 3479 | 0.529 | 0.940 | 8934 | 0.047 | 0.372 | 0.372 |
| 1782 | GSK2830371A | | 11892 | 0.496 | 0.722 | 3076 | 0.266 | 0.570 | 9177.5 | 0.075 | 0.559 | 0.559 |
| 1783 | LMB_AB1 | | 13062 | 0.859 | 0.981 | 2733 | 0.112 | 0.598 | 9766 | 0.194 | 0.622 | 0.622 |
| 1784 | LMB_AB2 | | 13086.5 | 0.864 | 0.993 | 3122 | 0.293 | 0.670 | 9148 | 0.071 | 0.378 | 0.378 |
| 1785 | LMB_AB3 | | 10805.5 | 0.155 | 0.619 | 3434 | 0.498 | 0.962 | 9121.5 | 0.067 | 0.480 | 0.480 |
| 1786 | AZD4547 | 1035270-39-3 | 11180 | 0.000 | 0.001 | 6600 | 0.857 | 0.958 | 17681 | 0.662 | 0.958 | 0.958 |
| 1799 | Ibrutinib | PCI-32765, CRA-032765, PCI-32765-00 | 16821.5 | 0.919 | 1.000 | 6340 | 0.993 | 1.000 | 12334 | 0.095 | 0.302 | 0.302 |
| 1802 | Zoledronate | Zoledronic acid, Zometa, Reclast | 13251 | 0.895 | 0.997 | 3176 | 0.326 | 0.633 | 9340.5 | 0.100 | 0.330 | 0.330 |
| 1803 | Acetalax | Oxyphenisatin acetate | 8269 | 0.000 | 0.002 | 3875 | 0.775 | 1.000 | 10339 | 0.381 | 0.872 | 0.872 |
| 1804 | Acetalax | Oxyphenisatin acetate | 8136 | 0.000 | 0.001 | 4302 | 0.932 | 1.000 | 9211 | 0.080 | 0.212 | 0.212 |
| 1806 | Oxaliplatin | | 8094 | 0.000 | 0.003 | 3675 | 0.659 | 0.924 | 9671 | 0.169 | 0.678 | 0.678 |
| 1808 | Topotecan | | 14987.5 | 0.998 | 1.000 | 2697 | 0.101 | 0.322 | 11671 | 0.842 | 1.000 | 1.000 |
| 1809 | Teniposide | Vumon | 15963 | 1.000 | 1.000 | 2791 | 0.132 | 0.353 | 10876.5 | 0.589 | 1.000 | 1.000 |
| 1810 | Mitoxantrone | | 15546.5 | 1.000 | 1.000 | 3177 | 0.327 | 0.653 | 10531 | 0.455 | 0.809 | 0.809 |
| 1811 | Dactinomycin | Actinomycin D | 10328 | 0.072 | 0.291 | 2554 | 0.064 | 0.291 | 9161.5 | 0.073 | 0.291 | 0.291 |
| 1812 | Bleomycin | | 13115 | 0.870 | 0.994 | 2978 | 0.214 | 0.428 | 10687 | 0.516 | 0.688 | 0.688 |
| 1813 | Fludarabine | Fludara | 11951 | 0.518 | 0.829 | 3137 | 0.302 | 0.802 | 10368.5 | 0.393 | 0.802 | 0.802 |
| 1814 | Nelarabine | Atriance, Arranon | 11321.5 | 0.302 | 0.627 | 3044 | 0.251 | 0.627 | 9429.5 | 0.120 | 0.479 | 0.479 |
| 1815 | Dacarbazine | DTIC | 10504.5 | 0.098 | 0.390 | 2669.5 | 0.093 | 0.390 | 9770.5 | 0.195 | 0.606 | 0.606 |
| 1816 | Fulvestrant | | 11172 | 0.249 | 0.537 | 3080 | 0.269 | 0.537 | 9420.5 | 0.115 | 0.537 | 0.537 |
| 1817 | Romidepsin | Istodax, FK228, Depsipeptide | 14710.5 | 0.995 | 1.000 | 2501.5 | 0.053 | 0.170 | 10325 | 0.376 | 0.729 | 0.729 |

| DRUG_ID | DRUG_NAME | SYNONYMS | nervous_system_MWU_STAT | nervous_system_PVAL | nervous_system_PVAL_ADJ | soft_tissue_MWU_STAT | soft_tissue_PVAL | soft_tissue_PVAL_ADJ | bone_MWU_STAT | bone_PVAL | bone_PVAL_ADJ | skin_MWU_STAT | skin_PVAL | skin_PVAL_ADJ | urogenital_system_MWU_STAT | urogenital_system_PVAL | urogenital_system_PVAL_ADJ | lung_NSCLC_MWU_STAT | lung_NSCLC_PVAL | lung_NSCLC_PVAL_ADJ | lung_SCLC_MWU_STAT | lung_SCLC_PVAL | lung_SCLC_PVAL_ADJ | lung_MWU_STAT | lung_PVAL | lung_PVAL_ADJ | neuroblastoma_MWU_STAT | neuroblastoma_PVAL | neuroblastoma_PVAL_ADJ | pancreas_MWU_STAT | pancreas_PVAL | pancreas_PVAL_ADJ | aero_dig_tract_MWU_STAT | aero_dig_tract_PVAL | aero_dig_tract_PVAL_ADJ | breast_MWU_STAT | breast_PVAL | breast_PVAL_ADJ | kidney_MWU_STAT | kidney_PVAL | kidney_PVAL_ADJ |
|---|---|---|---|---|---|---|---|---|---|---|---|---|---|---|---|---|---|---|---|---|---|---|---|---|---|---|---|---|---|---|---|---|---|---|---|---|---|---|---|---|---|
| 1819 | Docetaxel | RP-56976, Taxotere | 6721 | 0.070 | 0.279 | 3618 | 0.159 | 0.508 | 6179 | 0.032 | 0.253 | 7388 | 0.788 | 1.000 | 17443 | 0.372 | 0.744 | 19198 | 0.887 | 1.000 | 7548 | 0.642 | 0.973 | 7482 | 1.000 | 1.000 | 3477 | 0.056 | 0.279 | 8270 | 0.989 | 1.000 | 10741 | 0.010 | 0.167 | 10300 | 0.669 | 0.973 | 5895 | 0.959 | 1.000 |
| 1820 | 123829 | | 8065 | 0.087 | 0.278 | 6097 | 0.990 | 1.000 | 8012 | 0.184 | 0.395 | 9402 | 0.813 | 1.000 | 18798 | 0.197 | 0.395 | 18798 | 0.127 | 0.338 | 11903 | 0.996 | 1.000 | 7464 | 0.998 | 1.000 | 8719 | 1.000 | 1.000 | 7908 | 0.777 | 1.000 | 18149 | 0.995 | 1.000 | 11093 | 0.385 | 0.684 | 4598 | 0.078 | 0.278 |
| 1821 | 765771 | | 11196 | 0.972 | 0.999 | 5597 | 0.942 | 0.999 | 11085 | 0.992 | 0.999 | 8664 | 0.534 | 0.854 | 17833 | 0.062 | 0.246 | 15485.5 | 0.000 | 0.004 | 12281 | 0.999 | 0.999 | 4113 | 0.041 | 0.228 | 7215.5 | 0.994 | 0.999 | 6207.5 | 0.104 | 0.334 | 14462.5 | 0.315 | 0.719 | 13679 | 0.985 | 0.999 | 4371.5 | 0.043 | 0.228 |
| 1824 | 123138 | | 8933 | 0.325 | 0.650 | 5696 | 0.957 | 1.000 | 9847 | 0.859 | 1.000 | 9146 | 0.729 | 1.000 | 20834.5 | 0.728 | 1.000 | 18186 | 0.057 | 0.366 | 13290.5 | 1.000 | 1.000 | 2351.5 | 0.000 | 0.000 | 6518.5 | 0.942 | 1.000 | 6313 | 0.129 | 0.366 | 15325.5 | 0.591 | 1.000 | 13100.5 | 0.946 | 1.000 | 4838 | 0.137 | 0.366 |
| 1825 | Podophyllotoxin bromide | | 8000 | 0.077 | 0.245 | 3519 | 0.061 | 0.245 | 6821 | 0.015 | 0.117 | 8990 | 0.670 | 0.968 | 20825 | 0.726 | 0.968 | 22662 | 0.945 | 1.000 | 9306 | 0.475 | 0.950 | 7242 | 0.995 | 1.000 | 2685 | 0.000 | 0.002 | 10072 | 1.000 | 1.000 | 13568 | 0.111 | 0.296 | 11697 | 0.610 | 0.968 | 6820 | 0.942 | 1.000 |
| 1826 | 50869 | | 10320 | 0.841 | 0.994 | 5349 | 0.884 | 0.994 | 7360 | 0.055 | 0.292 | 10297 | 0.969 | 0.994 | 19897.5 | 0.475 | 0.994 | 21361 | 0.749 | 0.994 | 10361.5 | 0.851 | 0.994 | 6449 | 0.931 | 0.994 | 4008.5 | 0.030 | 0.236 | 7531 | 0.623 | 0.994 | 10472.5 | 0.000 | 0.001 | 12901 | 0.922 | 0.994 | 7517 | 0.994 | 0.994 |
| 1827 | Dihydrorotenone | | 11463 | 0.986 | 0.993 | 4726 | 0.603 | 0.993 | 10869 | 0.985 | 0.993 | 10859 | 0.993 | 0.993 | 17446 | 0.035 | 0.276 | 20917 | 0.640 | 0.993 | 8226 | 0.117 | 0.311 | 6120 | 0.848 | 0.993 | 6553 | 0.948 | 0.993 | 6615 | 0.220 | 0.502 | 13569 | 0.111 | 0.311 | 9897 | 0.077 | 0.307 | 6923 | 0.957 | 0.993 |
| 1828 | 720427 | | 9616.5 | 0.603 | 0.965 | 5611 | 0.944 | 0.996 | 9953 | 0.883 | 0.996 | 9484 | 0.836 | 0.996 | 18868 | 0.212 | 0.655 | 17375 | 0.016 | 0.252 | 11233 | 0.974 | 0.996 | 4394 | 0.088 | 0.467 | 7317 | 0.996 | 0.996 | 5726.5 | 0.034 | 0.270 | 14293.5 | 0.267 | 0.655 | 13226.5 | 0.958 | 0.996 | 5437 | 0.387 | 0.718 |
| 1829 | 667880 | | 10637 | 0.908 | 0.987 | 4103 | 0.251 | 0.668 | 8423 | 0.324 | 0.740 | 9501 | 0.841 | 0.987 | 18566 | 0.155 | 0.619 | 22241 | 0.902 | 0.987 | 5833 | 0.000 | 0.002 | 5485 | 0.560 | 0.987 | 2998 | 0.001 | 0.004 | 8120 | 0.844 | 0.987 | 16624 | 0.903 | 0.987 | 9349 | 0.026 | 0.138 | 7295 | 0.987 | 0.987 |
| 1830 | Gallibiscoquinazole | | 8377 | 0.150 | 0.802 | 5814 | 0.971 | 0.971 | 8399 | 0.315 | 0.839 | 9019 | 0.682 | 0.896 | 21085.5 | 0.784 | 0.896 | 20456.5 | 0.513 | 0.896 | 10371.5 | 0.853 | 0.910 | 5003 | 0.303 | 0.839 | 4982 | 0.293 | 0.839 | 5631 | 0.026 | 0.328 | 15961 | 0.774 | 0.896 | 11662 | 0.597 | 0.896 | 6072.5 | 0.715 | 0.896 |
| 1831 | L-Oxonoreleagnine | | 8963 | 0.337 | 0.673 | 5081 | 0.786 | 1.000 | 10214.5 | 0.929 | 1.000 | 8653 | 0.529 | 0.859 | 20306.5 | 0.591 | 0.859 | 19381 | 0.233 | 0.537 | 13259.5 | 1.000 | 1.000 | 3686 | 0.010 | 0.079 | 6776 | 0.973 | 1.000 | 5128 | 0.006 | 0.079 | 13696 | 0.132 | 0.537 | 12655.5 | 0.882 | 1.000 | 5112.5 | 0.235 | 0.537 |
| 1832 | 729189 | | 8155 | 0.103 | 0.411 | 5787 | 0.968 | 1.000 | 9336 | 0.700 | 1.000 | 10576 | 0.985 | 1.000 | 20666 | 0.687 | 1.000 | 19375.5 | 0.232 | 0.478 | 12566.5 | 1.000 | 1.000 | 3782.5 | 0.014 | 0.111 | 7507 | 0.998 | 1.000 | 5826 | 0.044 | 0.233 | 12014 | 0.006 | 0.099 | 12704 | 0.891 | 1.000 | 5397 | 0.367 | 0.652 |
| 1833 | 741909 | | 8721.5 | 0.250 | 0.667 | 4917 | 0.708 | 1.000 | 7042 | 0.026 | 0.139 | 10617 | 0.987 | 1.000 | 19664 | 0.409 | 0.817 | 20571 | 0.545 | 0.872 | 13051 | 1.000 | 1.000 | 2557 | 0.000 | 0.001 | 6868 | 0.980 | 1.000 | 7253 | 0.494 | 0.872 | 17516.5 | 0.979 | 1.000 | 10933 | 0.328 | 0.750 | 3218 | 0.001 | 0.004 |
| 1834 | 743380 | | 8616 | 0.216 | 0.692 | 5337 | 0.881 | 1.000 | 7702 | 0.109 | 0.581 | 10083 | 0.949 | 1.000 | 20014.5 | 0.508 | 0.813 | 21618 | 0.804 | 1.000 | 12850.5 | 1.000 | 1.000 | 2527 | 0.000 | 0.001 | 6898 | 0.982 | 1.000 | 7030 | 0.389 | 0.779 | 15809 | 0.735 | 1.000 | 11024 | 0.360 | 0.779 | 3405 | 0.001 | 0.010 |
| 1835 | Elephantin | | 9641 | 0.613 | 0.791 | 2743 | 0.003 | 0.027 | 8841 | 0.497 | 0.723 | 7760 | 0.183 | 0.419 | 19502 | 0.364 | 0.648 | 27910 | 1.000 | 1.000 | 7451 | 0.023 | 0.117 | 5641 | 0.642 | 0.791 | 3156 | 0.001 | 0.017 | 8287 | 0.887 | 1.000 | 17260.5 | 0.966 | 1.000 | 10551 | 0.210 | 0.420 | 4869.5 | 0.147 | 0.392 |
| 1836 | 150412 | | 10283 | 0.831 | 0.987 | 4509.5 | 0.475 | 0.987 | 10619 | 0.972 | 0.987 | 10493 | 0.981 | 0.987 | 20648.5 | 0.682 | 0.987 | 20393.5 | 0.495 | 0.987 | 10471.5 | 0.876 | 0.987 | 4985 | 0.295 | 0.785 | 6981.5 | 0.987 | 0.987 | 7397.5 | 0.562 | 0.987 | 12256 | 0.011 | 0.146 | 10644 | 0.236 | 0.756 | 6177 | 0.761 | 0.987 |
| 1838 | Sinularin | | 9114 | 0.396 | 0.725 | 4578 | 0.516 | 0.733 | 11122 | 0.993 | 0.993 | 9076 | 0.704 | 0.866 | 20162.5 | 0.550 | 0.733 | 21758 | 0.830 | 0.948 | 11371 | 0.982 | 0.993 | 4741 | 0.190 | 0.571 | 5208 | 0.408 | 0.725 | 5823 | 0.043 | 0.571 | 14914 | 0.456 | 0.730 | 10690 | 0.250 | 0.571 | 5066.5 | 0.216 | 0.571 |
| 1839 | 615590 | | 8394 | 0.155 | 0.360 | 5517 | 0.926 | 0.993 | 10660.5 | 0.974 | 0.993 | 9934 | 0.929 | 0.993 | 18514 | 0.146 | 0.360 | 18365.5 | 0.074 | 0.360 | 11610 | 0.991 | 0.993 | 4746 | 0.192 | 0.384 | 7107 | 0.992 | 0.993 | 6013 | 0.068 | 0.360 | 13388 | 0.086 | 0.360 | 13990.5 | 0.993 | 0.993 | 4903.5 | 0.158 | 0.360 |
| 1840 | 630600 | | 7760 | 0.047 | 0.326 | 4023 | 0.214 | 0.489 | 6476 | 0.005 | 0.086 | 7423 | 0.102 | 0.326 | 19048 | 0.251 | 0.502 | 25414 | 1.000 | 1.000 | 10134 | 0.789 | 0.911 | 7092 | 0.991 | 1.000 | 4540 | 0.124 | 0.331 | 7508 | 0.613 | 0.911 | 13435 | 0.092 | 0.326 | 11969 | 0.704 | 0.911 | 6268 | 0.797 | 0.911 |
| 1843 | LMP744 | MJ-III-65 | 8976 | 0.342 | 0.695 | 3871 | 0.154 | 0.492 | 5069 | 0.000 | 0.000 | 9798 | 0.907 | 1.000 | 19467.5 | 0.355 | 0.695 | 22618 | 0.941 | 1.000 | 6744 | 0.003 | 0.016 | 6486 | 0.937 | 1.000 | 3153 | 0.001 | 0.009 | 8703.5 | 0.956 | 1.000 | 14289 | 0.266 | 0.695 | 15311 | 1.000 | 1.000 | 5651 | 0.501 | 0.801 |
| 1844 | 776928 | | 9107 | 0.393 | 0.827 | 4756 | 0.620 | 0.827 | 6253 | 0.003 | 0.042 | 7612 | 0.144 | 0.575 | 22346 | 0.954 | 0.976 | 21667 | 0.813 | 0.976 | 7448 | 0.022 | 0.161 | 5367 | 0.495 | 0.827 | 4929 | 0.268 | 0.827 | 8934 | 0.976 | 0.976 | 15150 | 0.534 | 0.827 | 11584 | 0.568 | 0.827 | 5803 | 0.582 | 0.827 |
| 1845 | Schweinfurthin A | | 6087 | 0.000 | 0.002 | 3336 | 0.034 | 0.091 | 3384 | 0.000 | 0.000 | 8713 | 0.555 | 0.987 | 20949 | 0.755 | 1.000 | 22864 | 0.959 | 1.000 | 7130 | 0.010 | 0.031 | 6468 | 0.934 | 1.000 | 3410 | 0.003 | 0.018 | 10025 | 0.999 | 1.000 | 18366 | 0.997 | 1.000 | 13969 | 0.992 | 1.000 | 3784 | 0.006 | 0.024 |
| 1847 | BEN | | 11342 | 0.981 | 1.000 | 3662 | 0.091 | 0.252 | 6070 | 0.001 | 0.011 | 10227 | 0.963 | 1.000 | 21182 | 0.804 | 1.000 | 20234 | 0.450 | 0.900 | 6781 | 0.003 | 0.018 | 5703 | 0.674 | 1.000 | 4425 | 0.095 | 0.252 | 7764 | 0.723 | 1.000 | 16005 | 0.785 | 1.000 | 16166 | 1.000 | 1.000 | 3312 | 0.001 | 0.011 |
| 1849 | Sabutoclax | BI-97C1 | 17611 | 0.770 | 0.998 | 4549 | 0.024 | 0.128 | 7923 | 0.000 | 0.005 | 17776 | 0.660 | 0.998 | 27358 | 0.304 | 0.972 | 32237 | 0.895 | 0.998 | 13482 | 0.036 | 0.144 | 8068 | 0.806 | 0.998 | 5361 | 0.001 | 0.005 | 12337 | 0.998 | 0.998 | 22582 | 0.383 | 0.998 | 15328 | 0.588 | 0.998 | 12275 | 0.887 | 0.998 |
| 1852 | LY2109761 | 5XE | 13861.5 | 0.027 | 0.142 | 6528.5 | 0.625 | 0.960 | 12221 | 0.618 | 0.960 | 15491.5 | 0.118 | 0.378 | 30482 | 0.883 | 0.980 | 31855 | 0.853 | 0.980 | 20688.5 | 1.000 | 1.000 | 5929 | 0.075 | 0.301 | 9016.5 | 0.660 | 0.960 | 6751.5 | 0.008 | 0.135 | 19625 | 0.019 | 0.142 | 16895.5 | 0.919 | 0.980 | 9908.5 | 0.193 | 0.515 |
| 1853 | OF-1 | | 18958.5 | 0.955 | 0.999 | 6108 | 0.433 | 0.769 | 14879 | 0.994 | 0.999 | 21726.5 | 0.999 | 0.999 | 23551.5 | 0.005 | 0.077 | 28015 | 0.162 | 0.518 | 16985.5 | 0.772 | 0.999 | 7707 | 0.683 | 0.999 | 10749 | 0.983 | 0.999 | 7060 | 0.018 | 0.144 | 22028.5 | 0.264 | 0.703 | 12616.5 | 0.036 | 0.190 | 10673 | 0.423 | 0.769 |
| 1854 | MN-64 | | 15509 | 0.224 | 0.717 | 6788 | 0.732 | 1.000 | 16136.5 | 1.000 | 1.000 | 18587.5 | 0.837 | 1.000 | 28995 | 0.647 | 1.000 | 25282.5 | 0.007 | 0.040 | 20734 | 1.000 | 1.000 | 3561 | 0.000 | 0.001 | 10678 | 0.980 | 1.000 | 9606.5 | 0.629 | 1.000 | 20502.5 | 0.060 | 0.241 | 14334 | 0.302 | 0.804 | 7024 | 0.000 | 0.003 |
| 1855 | KRAS (G12C) Inhi | GTPL8020 | 17463.5 | 0.737 | 0.976 | 5404 | 0.162 | 0.519 | 9795 | 0.041 | 0.242 | 18798 | 0.870 | 0.976 | 27268 | 0.287 | 0.656 | 31332 | 0.780 | 0.976 | 15101 | 0.267 | 0.656 | 8305 | 0.868 | 0.976 | 7026 | 0.060 | 0.242 | 10187 | 0.811 | 0.976 | 20162 | 0.039 | 0.242 | 15234.5 | 0.561 | 0.976 | 13150 | 0.976 | 0.976 |
| 1862 | MG-132 | LLL cpd, MG 132, MG132 | 16105 | 0.015 | 0.119 | 6657 | 0.100 | 0.269 | 12595 | 0.198 | 0.397 | 18807 | 0.247 | 0.438 | 32084.5 | 0.154 | 0.351 | 45725 | 1.000 | 1.000 | 25704.5 | 0.999 | 1.000 | 9572 | 0.884 | 1.000 | 7558 | 0.003 | 0.042 | 13782 | 0.963 | 1.000 | 25668 | 0.287 | 0.460 | 16258 | 0.101 | 0.269 | 13607 | 0.848 | 1.000 |
| 1866 | BDP-00009066 | | 10923 | 0.615 | 0.824 | 5160.5 | 0.406 | 0.721 | 7977 | 0.057 | 0.653 | 10323 | 0.918 | 0.980 | 22107 | 0.326 | 0.653 | 22585 | 0.618 | 0.824 | 12250 | 0.942 | 0.980 | 4128.5 | 0.183 | 0.653 | 7329 | 0.753 | 0.927 | 8485 | 0.520 | 0.824 | 14440 | 0.144 | 0.653 | 14785 | 0.980 | 0.980 | 4735 | 0.207 | 0.653 |
| 1873 | Buparlisib | BKM120, NVP-BKM120 | 17820 | 0.220 | 0.565 | 6827 | 0.244 | 0.565 | 10160 | 0.005 | 0.039 | 24160 | 0.998 | 0.998 | 34778 | 0.674 | 0.981 | 35644.5 | 0.574 | 0.919 | 22353 | 0.953 | 0.998 | 10839 | 0.994 | 0.998 | 6954 | 0.001 | 0.010 | 14100 | 0.985 | 0.998 | 24586 | 0.204 | 0.565 | 16820 | 0.282 | 0.565 | 13041.5 | 0.755 | 0.998 |
| 1908 | Ulixertinib | BVD-523, VRT752271 | 25999 | 1.000 | 1.000 | 8788 | 0.903 | 1.000 | 18228 | 1.000 | 1.000 | 7549 | 0.000 | 0.000 | 36979 | 0.934 | 1.000 | 32378 | 0.089 | 0.204 | 29767.5 | 1.000 | 1.000 | 9125 | 0.805 | 1.000 | 8638 | 0.036 | 0.143 | 7846 | 0.001 | 0.008 | 25068 | 0.287 | 0.574 | 23385 | 1.000 | 1.000 | 14090 | 0.937 | 1.000 |
| 1909 | Venetoclax | ABT-199, Veneclexta, GDC-0199 | 21282 | 0.925 | 0.999 | 8732 | 0.893 | 0.999 | 10310 | 0.007 | 0.036 | 24172 | 0.998 | 0.999 | 36190 | 0.871 | 0.999 | 36393.5 | 0.705 | 0.999 | 11565 | 0.000 | 0.000 | 8003 | 0.410 | 0.819 | 5017 | 0.000 | 0.000 | 9521 | 0.054 | 0.217 | 31534 | 0.998 | 0.999 | 22198.5 | 0.999 | 0.999 | 11510 | 0.297 | 0.679 |
| 1910 | ABT737 | | 23286 | 0.997 | 1.000 | 8648 | 0.879 | 1.000 | 11058 | 0.029 | 0.155 | 25195 | 1.000 | 1.000 | 38150 | 0.982 | 1.000 | 35593 | 0.575 | 1.000 | 7074 | 0.000 | 0.000 | 9540 | 0.899 | 1.000 | 5532 | 0.000 | 0.000 | 13954 | 0.980 | 1.000 | 28558 | 0.914 | 1.000 | 18222 | 0.644 | 1.000 | 12035 | 0.460 | 1.000 |
| 1911 | Dactinomycin | Actinomycin D, Cosmegen, Lyovac, Meractinomycin, Ac-De, ActD | 15950 | 0.039 | 0.104 | 5047 | 0.008 | 0.032 | 10417 | 0.013 | 0.040 | 14986 | 0.005 | 0.026 | 30655 | 0.250 | 0.572 | 40051 | 0.997 | 1.000 | 19323 | 0.743 | 1.000 | 11035 | 0.998 | 1.000 | 5744 | 0.001 | 0.012 | 15399 | 1.000 | 1.000 | 20857 | 0.004 | 0.026 | 20439 | 0.982 | 1.000 | 14023 | 0.955 | 1.000 |
| 1912 | Afuresertib | GSK2110183, GSK2110183C | 16804 | 0.078 | 0.311 | 8558 | 0.857 | 1.000 | 12415 | 0.189 | 0.505 | 19980 | 0.572 | 1.000 | 35137 | 0.734 | 1.000 | 38133 | 0.913 | 1.000 | 22485 | 0.960 | 1.000 | 9988 | 0.955 | 1.000 | 11977 | 0.850 | 1.000 | 16010 | 1.000 | 1.000 | 24420 | 0.180 | 0.505 | 12792 | 0.001 | 0.009 | 9374 | 0.013 | 0.101 |
| 1913 | AGI-5198 | IDH-C35 | 19172.5 | 0.537 | 0.794 | 7911.5 | 0.659 | 0.811 | 14119 | 0.659 | 0.811 | 22418 | 0.957 | 0.997 | 33064 | 0.356 | 0.743 | 35490 | 0.546 | 0.794 | 24129 | 0.997 | 0.997 | 6546 | 0.050 | 0.267 | 10123.5 | 0.295 | 0.743 | 8593.5 | 0.009 | 0.072 | 29886 | 0.981 | 0.997 | 17193 | 0.371 | 0.743 | 8584 | 0.002 | 0.033 |
| 1915 | AZD3759 | | 20951 | 0.821 | 1.000 | 11291 | 1.000 | 1.000 | 18324 | 1.000 | 1.000 | 22311 | 0.939 | 1.000 | 31747 | 0.133 | 0.354 | 32524 | 0.082 | 0.262 | 30110.5 | 1.000 | 1.000 | 8943 | 0.735 | 1.000 | 12823 | 0.954 | 1.000 | 9691.5 | 0.065 | 0.258 | 14491 | 0.000 | 0.000 | 20750 | 0.974 | 1.000 | 5885 | 0.000 | 0.000 |
| 1916 | AZD5363 | | 14636 | 0.003 | 0.020 | 7033 | 0.314 | 0.728 | 10948 | 0.023 | 0.093 | 20282 | 0.645 | 1.000 | 34166 | 0.563 | 1.000 | 36607 | 0.739 | 1.000 | 23605 | 0.993 | 1.000 | 9973 | 0.954 | 1.000 | 14024 | 0.997 | 1.000 | 16249 | 1.000 | 1.000 | 25235.5 | 0.319 | 0.728 | 12488 | 0.000 | 0.005 | 9578.5 | 0.019 | 0.093 |
| 1917 | AZD6738 | AZD 6738, AZD-6738 | 17664 | 0.192 | 0.614 | 6827 | 0.244 | 0.635 | 9607 | 0.001 | 0.011 | 22144 | 0.938 | 0.998 | 34117 | 0.554 | 0.985 | 37755 | 0.881 | 0.998 | 22560 | 0.964 | 0.998 | 11240 | 0.998 | 0.998 | 5911 | 0.000 | 0.000 | 14582 | 0.995 | 0.998 | 24350 | 0.170 | 0.614 | 19086 | 0.823 | 0.998 | 9447 | 0.015 | 0.078 |
| 1918 | AZD8186 | AZD 8186, AZD-8186 | 14356 | 0.001 | 0.011 | 7668 | 0.564 | 1.000 | 13179.5 | 0.381 | 1.000 | 20939 | 0.784 | 1.000 | 34536.5 | 0.632 | 1.000 | 32580 | 0.105 | 0.422 | 28250.5 | 1.000 | 1.000 | 10384 | 0.981 | 1.000 | 13045.5 | 0.974 | 1.000 | 10900 | 0.320 | 1.000 | 26484.5 | 0.583 | 1.000 | 15535 | 0.077 | 0.409 | 5553 | 0.000 | 0.000 |
| 1919 | Osimertinib | AZD9291, AZD 9291, AZD-9291, Tagrisso, Mereletinib | 23212.5 | 0.996 | 1.000 | 9323 | 0.967 | 1.000 | 15524.5 | 0.930 | 1.000 | 27304 | 1.000 | 1.000 | 32788.5 | 0.308 | 0.704 | 33505 | 0.207 | 0.552 | 26689 | 1.000 | 1.000 | 8419 | 0.570 | 1.000 | 9519 | 0.146 | 0.501 | 12155.5 | 0.714 | 1.000 | 13435 | 0.000 | 0.000 | 16167 | 0.157 | 0.501 | 8251 | 0.001 | 0.007 |
| 1922 | Cediranib | AZD2171, AZD 2171, AZD-2171, Recentin | 17717 | 0.201 | 0.460 | 8176 | 0.751 | 0.994 | 10027 | 0.004 | 0.019 | 22848 | 0.976 | 0.994 | 34111 | 0.553 | 0.983 | 39060.5 | 0.963 | 0.994 | 23261 | 0.988 | 0.994 | 7010 | 0.116 | 0.372 | 7544 | 0.003 | 0.019 | 13892.5 | 0.977 | 0.994 | 23902 | 0.115 | 0.372 | 21524 | 0.994 | 0.994 | 8031 | 0.000 | 0.007 |
| 1924 | Ipatasertib | GDC0068, GDC 0068, GDC-0068 | 18787.5 | 0.439 | 1.000 | 7690 | 0.573 | 1.000 | 11328 | 0.044 | 0.177 | 23705.5 | 0.994 | 1.000 | 34189 | 0.567 | 1.000 | 36741 | 0.759 | 1.000 | 22210.5 | 0.943 | 1.000 | 9517.5 | 0.893 | 1.000 | 14693 | 1.000 | 1.000 | 13669 | 0.965 | 1.000 | 24195 | 0.149 | 0.478 | 10925 | 0.000 | 0.000 | 9898 | 0.034 | 0.177 |
| 1925 | GDC0810 | GDC-0810, GDC 0810 | 19585 | 0.639 | 0.840 | 8688 | 0.885 | 1.000 | 13062 | 0.348 | 0.741 | 26049.5 | 1.000 | 1.000 | 33640.5 | 0.463 | 0.741 | 33237.5 | 0.173 | 0.461 | 25905 | 1.000 | 1.000 | 8035 | 0.422 | 0.741 | 11064 | 0.602 | 0.840 | 8339 | 0.005 | 0.080 | 25738.5 | 0.422 | 0.741 | 15578 | 0.081 | 0.284 | 12770 | 0.682 | 0.840 |
| 1926 | GNE-317 | GNE317, GNE 317 | 16417 | 0.140 | 0.447 | 5657 | 0.037 | 0.147 | 9460 | 0.001 | 0.022 | 20376 | 0.806 | 0.976 | 33047 | 0.519 | 0.976 | 34243 | 0.588 | 0.976 | 26407 | 1.000 | 1.000 | 9217 | 0.863 | 0.976 | 10797 | 0.690 | 0.976 | 12667 | 0.876 | 0.976 | 25431 | 0.587 | 0.976 | 13709 | 0.011 | 0.090 | 9392 | 0.018 | 0.095 |
| 1927 | GSK2578215A | GSK 2578215A, GSK-2578215A | 18299 | 0.321 | 0.571 | 7556.5 | 0.519 | 0.755 | 11807 | 0.090 | 0.403 | 22621.5 | 0.967 | 0.995 | 33308 | 0.400 | 0.641 | 32975 | 0.143 | 0.403 | 23760 | 0.995 | 0.995 | 7305 | 0.182 | 0.417 | 9544.5 | 0.151 | 0.403 | 10868 | 0.310 | 0.571 | 27333 | 0.750 | 0.995 | 20716.5 | 0.978 | 0.995 | 10857 | 0.146 | 0.403 |
| 1928 | I-BRD9 | GSK602, GSK 602, GSK-602 | 21009 | 0.897 | 0.912 | 7046 | 0.319 | 0.679 | 9888 | 0.003 | 0.042 | 21480.5 | 0.870 | 0.912 | 33031.5 | 0.350 | 0.679 | 36040 | 0.646 | 0.912 | 19211.5 | 0.382 | 0.679 | 7121 | 0.139 | 0.582 | 10092 | 0.286 | 0.679 | 12662 | 0.837 | 0.912 | 28468 | 0.902 | 0.912 | 18494.5 | 0.703 | 0.912 | 11040 | 0.182 | 0.582 |
| 1930 | Telomerase Inhibi | MST-312, MST 312, MST312 | 15108 | 0.006 | 0.049 | 6178 | 0.088 | 0.278 | 14156 | 0.669 | 1.000 | 17687 | 0.104 | 0.278 | 30983 | 0.086 | 0.278 | 41580 | 0.999 | 1.000 | 16909 | 0.040 | 0.215 | 11696 | 1.000 | 1.000 | 3865 | 0.000 | 0.000 | 15003 | 0.998 | 1.000 | 25846 | 0.445 | 0.890 | 19689 | 0.908 | 1.000 | 12943 | 0.730 | 1.000 |
| 1931 | MIRA-1 | MIRA 1, MIRA1, NSC19630, NSC-19630, NSC 19630 | 16998.5 | 0.058 | 0.276 | 7013 | 0.290 | 0.547 | 13519 | 0.455 | 0.661 | 18878 | 0.277 | 0.547 | 37328.5 | 0.939 | 0.999 | 33439 | 0.167 | 0.535 | 22592.5 | 0.901 | 0.999 | 7560.5 | 0.240 | 0.547 | 8266 | 0.015 | 0.130 | 10933.5 | 0.308 | 0.547 | 32264 | 0.999 | 0.999 | 19463 | 0.862 | 0.999 | 9563 | 0.016 | 0.130 |
| 1932 | NVP-ADW742 | NVP ADW742, NVPADW742 | 23726 | 0.999 | 1.000 | 5888 | 0.050 | 0.158 | 6286 | 0.000 | 0.000 | 15431 | 0.004 | 0.020 | 42250 | 1.000 | 1.000 | 36004.5 | 0.639 | 1.000 | 18269 | 0.186 | 0.496 | 8601 | 0.638 | 1.000 | 6292 | 0.000 | 0.001 | 12439 | 0.788 | 1.000 | 26452 | 0.576 | 1.000 | 22540 | 0.999 | 1.000 | 13203 | 0.794 | 1.000 |
| 1933 | P22077 | P 22077, P-22077 | 21993.5 | 0.971 | 0.998 | 7291 | 0.412 | 0.725 | 16191 | 0.976 | 0.998 | 19501.5 | 0.453 | 0.725 | 33142.5 | 0.370 | 0.725 | 38007 | 0.903 | 0.998 | 14908 | 0.001 | 0.021 | 9036 | 0.781 | 0.998 | 8142.5 | 0.013 | 0.105 | 12701 | 0.845 | 0.998 | 31321.5 | 0.998 | 0.998 | 14853 | 0.030 | 0.120 | 9695 | 0.024 | 0.120 |
| 1936 | Savolitinib | AZD6094, Volitinib, AZD-6094, AZD 6094 | 16034 | 0.028 | 0.112 | 8261 | 0.778 | 1.000 | 15834 | 0.956 | 1.000 | 23233 | 0.987 | 1.000 | 33956 | 0.523 | 1.000 | 30244.5 | 0.009 | 0.051 | 25208.5 | 1.000 | 1.000 | 8057.5 | 0.430 | 1.000 | 11124 | 0.622 | 1.000 | 7986 | 0.002 | 0.033 | 27198.5 | 0.726 | 1.000 | 18806 | 0.771 | 1.000 | 13467 | 0.849 | 1.000 |
| 1939 | UMI-77 | UMI 77, UMI-77 | 10793 | 0.567 | 0.824 | 4918 | 0.285 | 0.761 | 6654 | 0.002 | 0.022 | 6337 | 0.003 | 0.022 | 24727 | 0.898 | 0.999 | 22106 | 0.493 | 0.824 | 10432 | 0.429 | 0.824 | 6041 | 0.967 | 0.999 | 5630 | 0.087 | 0.461 | 8527 | 0.538 | 0.824 | 14620 | 0.179 | 0.714 | 12088 | 0.366 | 0.824 | 7678 | 0.999 | 0.999 |
| 1940 | WIKI4 | WIKI-4, WIKI 4 | 18546 | 0.379 | 0.759 | 8589 | 0.864 | 1.000 | 17431.5 | 0.998 | 1.000 | 25003 | 1.000 | 1.000 | 29566 | 0.020 | 0.082 | 29701 | 0.005 | 0.037 | 26689.5 | 1.000 | 1.000 | 6715 | 0.069 | 0.222 | 10558.5 | 0.433 | 0.769 | 8697 | 0.011 | 0.060 | 28709 | 0.923 | 1.000 | 18719 | 0.753 | 1.000 | 6938 | 0.000 | 0.000 |
| 1941 | Sepantronium brc | YM155, YM-155, YM 155 | 15528 | 0.013 | 0.056 | 5353 | 0.014 | 0.056 | 11494.5 | 0.057 | 0.153 | 16993 | 0.045 | 0.144 | 27083 | 0.001 | 0.010 | 34355.5 | 0.338 | 0.695 | 15085 | 0.002 | 0.015 | 10177 | 0.970 | 1.000 | 14090 | 0.998 | 1.000 | 13474 | 0.951 | 1.000 | 29977 | 0.983 | 1.000 | 18890 | 0.787 | 1.000 | 13612 | 0.874 | 1.000 |
| 1996 | MIM1 | MIM 1, MIM-1 | 9170 | 0.082 | 0.318 | 5302 | 0.481 | 0.843 | 8040 | 0.065 | 0.318 | 9181 | 0.579 | 0.843 | 21290 | 0.162 | 0.432 | 23193 | 0.760 | 0.993 | 12165 | 0.931 | 0.993 | 4329 | 0.268 | 0.537 | 5297 | 0.039 | 0.311 | 7247 | 0.100 | 0.318 | 18883 | 0.993 | 0.993 | 14853 | 0.983 | 0.993 | 4752 | 0.214 | 0.489 |
| 1997 | WEHI-539 | WEHI539, WEHI-539 | 20287 | 0.697 | 0.998 | 9284 | 0.960 | 0.998 | 12297.5 | 0.154 | 0.434 | 24543 | 0.998 | 0.998 | 32631.5 | 0.252 | 0.575 | 35763.5 | 0.557 | 0.990 | 15650 | 0.002 | 0.025 | 10455 | 0.982 | 0.998 | 7766 | 0.005 | 0.040 | 14636 | 0.994 | 0.998 | 29443 | 0.940 | 0.998 | 15551.5 | 0.103 | 0.414 | 9412 | 0.012 | 0.065 |
| 1998 | BPD-00008900 | | 9368 | 0.115 | 0.307 | 4257 | 0.072 | 0.307 | 9965 | 0.668 | 0.839 | 10285 | 0.912 | 1.000 | 22786 | 0.498 | 0.839 | 20329.5 | 0.115 | 0.307 | 14428.5 | 1.000 | 1.000 | 3078.5 | 0.008 | 0.063 | 6931.5 | 0.581 | 0.839 | 8449 | 0.505 | 0.839 | 16388 | 0.682 | 0.839 | 15610 | 0.997 | 1.000 | 3253 | 0.002 | 0.038 |
| 1999 | N25720-51-A1 | | 11337.5 | 0.761 | 0.891 | 5105 | 0.381 | 0.767 | 9537 | 0.507 | 0.767 | 8857 | 0.451 | 0.767 | 20112 | 0.055 | 0.460 | 24513.5 | 0.947 | 0.985 | 10780 | 0.569 | 0.767 | 4797 | 0.526 | 0.767 | 7390 | 0.780 | 0.891 | 10440.5 | 0.985 | 0.985 | 14032 | 0.086 | 0.460 | 12477.5 | 0.509 | 0.767 | 5142 | 0.401 | 0.767 |
| 2000 | N27922-53-1 | | 10258 | 0.365 | 0.729 | 5253.5 | 0.455 | 0.809 | 11088.5 | 0.942 | 0.995 | 10311 | 0.916 | 0.995 | 22929 | 0.535 | 0.857 | 20648 | 0.161 | 0.474 | 13269 | 0.995 | 0.995 | 3170.5 | 0.011 | 0.181 | 7906 | 0.916 | 0.995 | 7251 | 0.100 | 0.401 | 16525 | 0.719 | 0.995 | 12072.5 | 0.361 | 0.729 | 4274.5 | 0.075 | 0.400 |
| 2002 | N30652-18-1 | | 12749 | 0.980 | 0.989 | 5253.5 | 0.455 | 0.809 | 10984.5 | 0.929 | 0.989 | 10199 | 0.897 | 0.989 | 22815.5 | 0.506 | 0.809 | 19937.5 | 0.072 | 0.345 | 12990 | 0.989 | 0.989 | 3320.5 | 0.019 | 0.311 | 6363 | 0.316 | 0.722 | 7260 | 0.102 | 0.345 | 14218 | 0.108 | 0.345 | 13673.5 | 0.856 | 0.989 | 4551 | 0.144 | 0.383 |
| 2003 | N29087-69-1 | | 11339.5 | 0.756 | 1.000 | 5863 | 0.762 | 1.000 | 10564.5 | 0.851 | 1.000 | 10205 | 0.898 | 1.000 | 22532 | 0.432 | 0.788 | 19678 | 0.051 | 0.250 | 14489 | 1.000 | 1.000 | 2747.5 | 0.002 | 0.031 | 6439 | 0.349 | 0.788 | 6281.5 | 0.010 | 0.080 | 15799.5 | 0.503 | 0.806 | 15718.5 | 0.998 | 1.000 | 4204.5 | 0.062 | 0.250 |
| 2010 | HKMTI-1-005 | | 10523 | 0.595 | 0.952 | 5081 | 0.382 | 0.765 | 6810 | 0.007 | 0.027 | 5508 | 0.000 | 0.003 | 28662 | 1.000 | 1.000 | 23327 | 0.819 | 1.000 | 7669 | 0.001 | 0.011 | 5660 | 0.911 | 1.000 | 5940 | 0.174 | 0.422 | 11075 | 0.998 | 1.000 | 17976 | 0.947 | 1.000 | 14475 | 0.947 | 1.000 | 4643 | 0.185 | 0.422 |
| 2011 | ICL-SIRT078 | | 11849.5 | 0.938 | 0.966 | 5762 | 0.736 | 0.966 | 9310 | 0.546 | 0.966 | 7281 | 0.042 | 0.209 | 25384 | 0.966 | 0.966 | 22951.5 | 0.746 | 0.966 | 10781 | 0.489 | 0.966 | 5917 | 0.957 | 0.966 | 6000 | 0.193 | 0.605 | 8817 | 0.682 | 0.966 | 14969 | 0.227 | 0.605 | 13563.5 | 0.792 | 0.966 | 3398 | 0.005 | 0.075 |
| 2037 | GSK343 | | 11388 | 0.862 | 0.964 | 5429 | 0.570 | 0.892 | 9477 | 0.613 | 0.892 | 5235 | 0.000 | 0.001 | 27975.5 | 1.000 | 1.000 | 21227 | 0.310 | 0.729 | 12176.5 | 0.904 | 0.964 | 4402 | 0.319 | 0.729 | 6843 | 0.561 | 0.892 | 9040 | 0.763 | 0.964 | 14371 | 0.111 | 0.445 | 13669.5 | 0.817 | 0.964 | 3780 | 0.019 | 0.148 |
| 2038 | UNC0638 | | 13164.5 | 0.997 | 0.997 | 5597.5 | 0.695 | 0.997 | 9140 | 0.419 | 0.976 | 9244 | 0.681 | 0.997 | 24261 | 0.949 | 0.997 | 18654 | 0.072 | 0.230 | 7171 | 0.001 | 0.012 | 4921 | 0.647 | 0.997 | 6474 | 0.427 | 0.976 | 8177.5 | 0.582 | 0.997 | 17454 | 0.947 | 0.997 | 13169.5 | 0.947 | 0.997 | 3744 | 0.020 | 0.107 |
| 2039 | AGK2 | | 18366 | 0.709 | 0.872 | 6925 | 0.484 | 0.774 | 10631 | 0.038 | 0.304 | 22784 | 0.997 | 0.999 | 33222.5 | 0.621 | 0.828 | 33437.5 | 0.450 | 0.774 | 23106.5 | 0.999 | 0.999 | 4728 | 0.003 | 0.055 | 10871.5 | 0.836 | 0.956 | 10092 | 0.180 | 0.481 | 25038.5 | 0.558 | 0.811 | 16551 | 0.406 | 0.774 | 10547 | 0.140 | 0.449 |
| 2040 | Foretinib | GSK1363089, XL-880, EXEL-2880, GSK089 | 16079 | 0.057 | 0.151 | 5709 | 0.038 | 0.151 | 6793 | 0.000 | 0.000 | 21567 | 0.905 | 1.000 | 41338 | 1.000 | 1.000 | 35856 | 0.679 | 0.984 | 19841 | 0.666 | 0.984 | 8032 | 0.449 | 0.799 | 5643 | 0.000 | 0.000 | 13829.5 | 0.980 | 1.000 | 27305 | 0.738 | 0.984 | 23488 | 1.000 | 1.000 | 11752 | 0.402 | 0.799 |

| DRUG_ID | DRUG_NAME | SYNONYMS | large_intestine_MWU_STAT | large_intestine_PVAL | large_intestine_PVAL_ADJ | thyroid_MWU_STAT | thyroid_PVAL | thyroid_PVAL_ADJ | digestive_system_MWU_STAT | digestive_system_PVAL | digestive_system_PVAL_ADJ | digestive_system_PVAL2 |
|---|---|---|---|---|---|---|---|---|---|---|---|---|
| 1819 | Docetaxel | RP-56976, Taxotere | 10810 | 0.498 | 0.886 | 2889 | 0.280 | 0.669 | 8911 | 0.293 | 0.669 | 0.669 |
| 1820 | 123829 | | 8054 | 0.000 | 0.003 | 1829 | 0.003 | 0.022 | 8340 | 0.012 | 0.063 | 0.063 |
| 1821 | 765771 | | 11735.5 | 0.439 | 0.780 | 2983 | 0.216 | 0.577 | 10320 | 0.374 | 0.749 | 0.749 |
| 1824 | 123138 | | 10417 | 0.084 | 0.366 | 2960 | 0.205 | 0.468 | 9472 | 0.125 | 0.366 | 0.366 |
| 1825 | Podophyllotoxin bromide | | 12076 | 0.564 | 0.968 | 2575 | 0.068 | 0.245 | 10371 | 0.394 | 0.899 | 0.899 |
| 1826 | 50869 | | 10378 | 0.079 | 0.315 | 3197 | 0.339 | 0.904 | 9829 | 0.211 | 0.676 | 0.676 |
| 1827 | Dihydrorotenone | | 9563 | 0.015 | 0.241 | 4475 | 0.963 | 0.993 | 9008 | 0.054 | 0.288 | 0.288 |
| 1828 | 720427 | | 11294 | 0.287 | 0.655 | 3002 | 0.226 | 0.655 | 10398 | 0.404 | 0.718 | 0.718 |
| 1829 | 667880 | | 14317 | 0.987 | 0.987 | 3942 | 0.808 | 0.987 | 9795 | 0.202 | 0.646 | 0.646 |
| 1830 | Gallibiscoquinazole | | 10026 | 0.041 | 0.328 | 3403 | 0.476 | 0.896 | 10966 | 0.623 | 0.896 | 0.896 |
| 1831 | L-Oxonoreleagnine | | 11093 | 0.227 | 0.537 | 3558 | 0.582 | 0.859 | 9665.5 | 0.168 | 0.537 | 0.537 |
| 1832 | 729189 | | 10873 | 0.170 | 0.453 | 3026.5 | 0.239 | 0.478 | 9637 | 0.161 | 0.453 | 0.453 |
| 1833 | 741909 | | 10831 | 0.160 | 0.513 | 3877 | 0.776 | 1.000 | 9592 | 0.151 | 0.513 | 0.513 |
| 1834 | 743380 | | 11240 | 0.270 | 0.719 | 3398 | 0.473 | 0.813 | 9677 | 0.171 | 0.683 | 0.683 |
| 1835 | Elephantin | | 10656 | 0.124 | 0.392 | 2342 | 0.029 | 0.117 | 10618 | 0.489 | 0.723 | 0.723 |
| 1836 | 150412 | | 10273 | 0.065 | 0.281 | 2583 | 0.070 | 0.281 | 8514 | 0.018 | 0.146 | 0.146 |
| 1838 | Sinularin | | 10923 | 0.182 | 0.571 | 2900 | 0.177 | 0.571 | 9782 | 0.198 | 0.571 | 0.571 |
| 1839 | 615590 | | 11204 | 0.259 | 0.460 | 3315 | 0.416 | 0.666 | 9567 | 0.145 | 0.360 | 0.360 |
| 1840 | 630600 | | 12786 | 0.794 | 0.911 | 2670 | 0.093 | 0.326 | 10683 | 0.514 | 0.911 | 0.911 |
| 1843 | LMP744 | MJ-III-65 | 14045 | 0.977 | 1.000 | 2683 | 0.096 | 0.386 | 10364 | 0.391 | 0.695 | 0.695 |
| 1844 | 776928 | | 13935 | 0.970 | 0.976 | 3608 | 0.616 | 0.827 | 8730 | 0.030 | 0.161 | 0.161 |
| 1845 | Schweinfurthin A | | 15557 | 1.000 | 1.000 | 3291 | 0.400 | 0.800 | 9896 | 0.231 | 0.528 | 0.528 |
| 1847 | BEN | | 10742 | 0.141 | 0.322 | 4343 | 0.941 | 1.000 | 9271 | 0.089 | 0.252 | 0.252 |
| 1849 | Sabutoclax | BI-97C1 | 17477 | 0.966 | 0.998 | 5375 | 0.869 | 0.998 | 14079 | 0.496 | 0.998 | 0.998 |
| 1852 | LY2109761 | 5XE | 15809.5 | 0.720 | 0.960 | 4398 | 0.418 | 0.956 | 14356.5 | 0.581 | 0.960 | 0.960 |
| 1853 | OF-1 | | 14510 | 0.349 | 0.769 | 5801 | 0.956 | 0.999 | 12449 | 0.102 | 0.410 | 0.410 |
| 1854 | MN-64 | | 16486.5 | 0.862 | 1.000 | 4698 | 0.580 | 1.000 | 14895 | 0.733 | 1.000 | 1.000 |
| 1855 | KRAS (G12C) Inhi | GTPL8020 | 17679 | 0.976 | 0.976 | 4830 | 0.648 | 0.976 | 12083 | 0.061 | 0.242 | 0.242 |
| 1862 | MG-132 | LLL cpd, MG 132, MG132 | 14257 | 0.062 | 0.247 | 4161 | 0.035 | 0.184 | 16606 | 0.345 | 0.502 | 0.502 |
| 1866 | BDP-00009066 | | 10917.5 | 0.321 | 0.653 | 3845 | 0.305 | 0.653 | 11411.5 | 0.237 | 0.653 | 0.653 |
| 1873 | Buparlisib | BKM120, NVP-BKM120 | 14741 | 0.261 | 0.565 | 5288 | 0.328 | 0.582 | 15234 | 0.115 | 0.565 | 0.565 |
| 1908 | Ulixertinib | BVD-523, VRT752271 | 6579 | 0.000 | 0.000 | 4251 | 0.048 | 0.152 | 14675.5 | 0.057 | 0.152 | 0.152 |
| 1909 | Venetoclax | ABT-199, Veneclexta, GDC-0199 | 15999 | 0.597 | 0.999 | 4772 | 0.147 | 0.469 | 16147 | 0.279 | 0.679 | 0.679 |
| 1910 | ABT737 | | 14389 | 0.191 | 0.649 | 5953 | 0.634 | 1.000 | 15435 | 0.203 | 0.649 | 0.649 |
| 1911 | Dactinomycin | Actinomycin D, Cosmegen, Lyovac, Meractinomycin, Ac-De, ActD | 17094 | 0.895 | 1.000 | 4786 | 0.305 | 0.609 | 17683 | 0.754 | 1.000 | 1.000 |
| 1912 | Afuresertib | GSK2110183, GSK2110183C | 12737 | 0.020 | 0.107 | 6211 | 0.737 | 1.000 | 16325 | 0.321 | 0.733 | 0.733 |
| 1913 | AGI-5198 | IDH-C35 | 15599 | 0.485 | 0.794 | 4686 | 0.125 | 0.499 | 15831 | 0.212 | 0.679 | 0.679 |
| 1915 | AZD3759 | | 13751 | 0.052 | 0.258 | 6086 | 0.670 | 1.000 | 17250 | 0.532 | 1.000 | 1.000 |
| 1916 | AZD5363 | | 15361 | 0.419 | 0.838 | 6930 | 0.930 | 1.000 | 16230 | 0.298 | 0.728 | 0.728 |
| 1917 | AZD6738 | AZD 6738, AZD-6738 | 14815.5 | 0.278 | 0.635 | 6444 | 0.818 | 0.998 | 16796.5 | 0.442 | 0.883 | 0.883 |
| 1918 | AZD8186 | AZD 8186, AZD-8186 | 16563 | 0.740 | 1.000 | 6564 | 0.853 | 1.000 | 16807.5 | 0.445 | 1.000 | 1.000 |
| 1919 | Osimertinib | AZD9291, AZD 9291, AZD-9291, Tagrisso, Mereletinib | 13424 | 0.058 | 0.311 | 6866 | 0.919 | 1.000 | 16763 | 0.433 | 0.866 | 0.866 |
| 1922 | Cediranib | AZD2171, AZD 2171, AZD-2171, Recentin | 14745.5 | 0.262 | 0.523 | 6051 | 0.672 | 0.994 | 15445.5 | 0.145 | 0.386 | 0.386 |
| 1924 | Ipatasertib | GDC0068, GDC 0068, GDC-0068 | 12817.5 | 0.023 | 0.177 | 6773 | 0.902 | 1.000 | 16206.5 | 0.293 | 0.781 | 0.781 |
| 1925 | GDC0810 | GDC-0810, GDC 0810 | 12745 | 0.020 | 0.108 | 3891 | 0.019 | 0.108 | 15018 | 0.089 | 0.284 | 0.284 |
| 1926 | GNE-317 | GNE317, GNE 317 | 16125 | 0.700 | 0.976 | 6731 | 0.915 | 0.976 | 15334 | 0.230 | 0.614 | 0.614 |
| 1927 | GSK2578215A | GSK 2578215A, GSK-2578215A | 18026 | 0.953 | 0.995 | 4623 | 0.110 | 0.403 | 13746 | 0.014 | 0.218 | 0.218 |
| 1928 | I-BRD9 | GSK602, GSK 602, GSK-602 | 17569 | 0.912 | 0.912 | 4801 | 0.155 | 0.582 | 15392 | 0.137 | 0.582 | 0.582 |
| 1930 | Telomerase Inhibi | MST-312, MST 312, MST312 | 17665 | 0.922 | 1.000 | 5135 | 0.266 | 0.607 | 18903 | 0.899 | 1.000 | 1.000 |
| 1931 | MIRA-1 | MIRA 1, MIRA1, NSC19630, NSC-19630, NSC 19630 | 18069.5 | 0.913 | 0.999 | 4432.5 | 0.069 | 0.276 | 16608.5 | 0.363 | 0.581 | 0.581 |
| 1932 | NVP-ADW742 | NVP ADW742, NVPADW742 | 12320 | 0.009 | 0.038 | 7217.5 | 0.965 | 1.000 | 16427 | 0.346 | 0.791 | 0.791 |
| 1933 | P22077 | P 22077, P-22077 | 15463 | 0.447 | 0.725 | 4964.5 | 0.205 | 0.655 | 16135 | 0.276 | 0.725 | 0.725 |
| 1936 | Savolitinib | AZD6094, Volitinib, AZD-6094, AZD 6094 | 15573.5 | 0.478 | 1.000 | 4814 | 0.158 | 0.506 | 13548.5 | 0.010 | 0.051 | 0.051 |
| 1939 | UMI-77 | UMI 77, UMI-77 | 13186 | 0.949 | 0.999 | 3731 | 0.247 | 0.761 | 13934 | 0.939 | 0.999 | 0.999 |
| 1940 | WIKI4 | WIKI-4, WIKI 4 | 17211.5 | 0.864 | 1.000 | 4748 | 0.140 | 0.359 | 15524 | 0.157 | 0.359 | 0.359 |
| 1941 | Sepantronium brc | YM155, YM-155, YM 155 | 22429 | 1.000 | 1.000 | 5335 | 0.348 | 0.695 | 19470 | 0.952 | 1.000 | 1.000 |
| 1996 | MIM1 | MIM 1, MIM-1 | 11613 | 0.571 | 0.843 | 2618 | 0.008 | 0.135 | 13469 | 0.870 | 0.993 | 0.993 |
| 1997 | WEHI-539 | WEHI539, WEHI-539 | 17312 | 0.803 | 0.998 | 5525 | 0.419 | 0.839 | 15645 | 0.163 | 0.434 | 0.434 |
| 1998 | BPD-00008900 | | 11566 | 0.554 | 0.839 | 4030 | 0.410 | 0.839 | 10677 | 0.085 | 0.307 | 0.307 |
| 1999 | N25720-51-A1 | | 11603 | 0.575 | 0.767 | 3156 | 0.060 | 0.460 | 11598 | 0.298 | 0.767 | 0.767 |
| 2000 | N27922-53-1 | | 12764 | 0.894 | 0.995 | 3575.5 | 0.178 | 0.474 | 10106 | 0.030 | 0.237 | 0.237 |
| 2002 | N30652-18-1 | | 12420 | 0.823 | 0.989 | 3981 | 0.381 | 0.762 | 10686 | 0.086 | 0.345 | 0.345 |
| 2003 | N29087-69-1 | | 10725.5 | 0.260 | 0.694 | 4085 | 0.443 | 0.788 | 10730 | 0.092 | 0.295 | 0.295 |
| 2010 | HKMTI-1-005 | | 7891 | 0.005 | 0.027 | 4208 | 0.535 | 0.952 | 9647 | 0.036 | 0.114 | 0.114 |
| 2011 | ICL-SIRT078 | | 8604.5 | 0.030 | 0.209 | 4665 | 0.786 | 0.966 | 9842 | 0.052 | 0.209 | 0.209 |
| 2037 | GSK343 | | 10754.5 | 0.580 | 0.892 | 3801 | 0.295 | 0.729 | 9526 | 0.028 | 0.148 | 0.148 |
| 2038 | UNC0638 | | 9735.5 | 0.056 | 0.223 | 5100.5 | 0.942 | 0.997 | 8874 | 0.015 | 0.107 | 0.107 |
| 2039 | AGK2 | | 14037 | 0.136 | 0.449 | 5106.5 | 0.307 | 0.701 | 14474 | 0.073 | 0.391 | 0.391 |
| 2040 | Foretinib | GSK1363089, XL-880, EXEL-2880, GSK089 | 12233 | 0.049 | 0.151 | 5379.5 | 0.389 | 0.799 | 13416 | 0.016 | 0.087 | 0.087 |

Suppl. Table 8

| DRUG_ID | DRUG_NAME | SYNONYMS | nervous_system_MWU_STAT | nervous_system_PVAL | nervous_system_PVAL_ADJ | soft_tissue_MWU_STAT | soft_tissue_PVAL | soft_tissue_PVAL_ADJ | bone_MWU_STAT | bone_PVAL | bone_PVAL_ADJ | skin_MWU_STAT | skin_PVAL | skin_PVAL_ADJ | urogenital_system_MWU_STAT | urogenital_system_PVAL | urogenital_system_PVAL_ADJ | lung_NSCLC_MWU_STAT | lung_NSCLC_PVAL | lung_NSCLC_PVAL_ADJ | lung_SCLC_MWU_STAT | lung_SCLC_PVAL | lung_SCLC_PVAL_ADJ | lung_MWU_STAT | lung_PVAL | lung_PVAL_ADJ | neuroblastoma_MWU_STAT | neuroblastoma_PVAL | neuroblastoma_PVAL_ADJ | pancreas_MWU_STAT | pancreas_PVAL | pancreas_PVAL_ADJ | aero_dig_tract_MWU_STAT | aero_dig_tract_PVAL | aero_dig_tract_PVAL_ADJ | breast_MWU_STAT | breast_PVAL | breast_PVAL_ADJ | kidney_MWU_STAT | kidney_PVAL | kidney_PVAL_ADJ |
|---|---|---|---|---|---|---|---|---|---|---|---|---|---|---|---|---|---|---|---|---|---|---|---|---|---|---|---|---|---|---|---|---|---|---|---|---|---|---|---|---|---|
| 2043 | BIBR-1532 | | 17787 | 0.319 | 0.574 | 7743 | 0.622 | 0.889 | 13866 | 0.722 | 0.889 | 20215 | 0.676 | 0.889 | 32422 | 0.304 | 0.574 | 34098.5 | 0.357 | 0.574 | 24657 | 1.000 | 1.000 | 7523 | 0.266 | 0.574 | 9806 | 0.233 | 0.574 | 9628 | 0.075 | 0.403 | 28615.5 | 0.912 | 0.973 | 16648 | 0.359 | 0.574 | 10281 | 0.076 | 0.403 |
| 2044 | Pyridostatin | | 17143 | 0.195 | 0.780 | 6933 | 0.308 | 0.822 | 7738 | 0.000 | 0.000 | 24559 | 0.999 | 1.000 | 34208 | 0.656 | 0.954 | 34786 | 0.556 | 0.954 | 13801 | 0.000 | 0.003 | 9017 | 0.806 | 1.000 | 10364 | 0.410 | 0.938 | 11441 | 0.537 | 0.954 | 22806 | 0.037 | 0.195 | 19398 | 0.937 | 1.000 | 15699 | 1.000 | 1.000 |
| 2045 | AMG-319 | AMG319 | 14940.5 | 0.010 | 0.080 | 6856 | 0.275 | 0.549 | 12223.5 | 0.248 | 0.549 | 18730 | 0.313 | 0.557 | 33911 | 0.582 | 0.847 | 31654.5 | 0.063 | 0.254 | 22418 | 0.981 | 1.000 | 7934.5 | 0.412 | 0.659 | 11961.5 | 0.868 | 1.000 | 9341 | 0.047 | 0.249 | 33238 | 1.000 | 1.000 | 20102.5 | 0.976 | 1.000 | 7067 | 0.000 | 0.000 |
| 2046 | MK-8776 | SCH900776 | 19115 | 0.653 | 0.982 | 5310 | 0.014 | 0.058 | 9473 | 0.002 | 0.020 | 21147.5 | 0.852 | 0.982 | 37835.5 | 0.982 | 0.982 | 34516 | 0.433 | 0.982 | 19625.5 | 0.615 | 0.982 | 8253 | 0.536 | 0.982 | 5489 | 0.000 | 0.000 | 12998.5 | 0.913 | 0.982 | 29639 | 0.972 | 0.982 | 19471.5 | 0.939 | 0.982 | 9281 | 0.012 | 0.058 |
| 2047 | Ulixertinib | BVD-523, VRT752271 | 12819 | 0.994 | 1.000 | 5893 | 0.791 | 1.000 | 11890 | 0.997 | 1.000 | 4783 | 0.000 | 0.000 | 24562 | 0.900 | 1.000 | 21844.5 | 0.468 | 1.000 | 14415 | 1.000 | 1.000 | 4784 | 0.533 | 1.000 | 4694 | 0.007 | 0.029 | 7189 | 0.098 | 0.261 | 16628 | 0.711 | 1.000 | 16285 | 0.999 | 1.000 | 5618 | 0.668 | 1.000 |
| 2048 | Vinorelbine | vinorelbine tartrate, Navelbine, Exelbine | 16979 | 0.106 | 0.243 | 5825 | 0.046 | 0.123 | 8736 | 0.000 | 0.005 | 19879 | 0.575 | 1.000 | 35244 | 0.781 | 1.000 | 37134 | 0.839 | 1.000 | 16310 | 0.019 | 0.082 | 12116 | 1.000 | 1.000 | 7681 | 0.021 | 0.082 | 16385 | 1.000 | 1.000 | 21778 | 0.011 | 0.082 | 19311 | 0.874 | 1.000 | 16551 | 1.000 | 1.000 |
| 2055 | Mycophenolic acid | | 7710 | 0.002 | 0.014 | 4069 | 0.050 | 0.134 | 6591 | 0.004 | 0.021 | 9313 | 0.682 | 0.910 | 22206 | 0.437 | 0.776 | 27584 | 1.000 | 1.000 | 8214 | 0.042 | 0.134 | 6091 | 0.979 | 1.000 | 2430 | 0.000 | 0.000 | 11698 | 1.000 | 1.000 | 15829 | 0.585 | 0.910 | 16531 | 1.000 | 1.000 | 4046 | 0.047 | 0.134 |
| 2057 | Remodelin | | 3428 | 0.993 | 0.998 | 1237 | 0.383 | 0.876 | 2181 | 0.843 | 0.998 | 3244 | 0.998 | 0.998 | 5180 | 0.104 | 0.332 | 5991 | 0.666 | 0.998 | 1470 | 0.045 | 0.241 | 1162 | 0.903 | 0.998 | 1559 | 0.450 | 0.900 | 1726 | 0.187 | 0.499 | 5232 | 0.989 | 0.998 | 2355 | 0.009 | 0.072 | 1431 | 0.676 | 0.998 |
| 2096 | VX-11e | VX11e, VX11e | 23179 | 0.995 | 1.000 | 9185 | 0.962 | 1.000 | 17935 | 1.000 | 1.000 | 7850 | 0.000 | 0.000 | 37904 | 0.983 | 1.000 | 29760 | 0.007 | 0.038 | 29323 | 1.000 | 1.000 | 10326 | 0.982 | 1.000 | 9689 | 0.530 | 0.942 | 9627 | 0.073 | 0.252 | 23306 | 0.079 | 0.252 | 23427 | 1.000 | 1.000 | 11230 | 0.247 | 0.566 |
| 2106 | Uprosertib | GSK2141795, GSK2141795C, GSK-2141795 | 9923 | 0.223 | 0.509 | 5185 | 0.461 | 0.751 | 8091 | 0.141 | 0.377 | 6434 | 0.005 | 0.028 | 25741 | 0.991 | 1.000 | 22855 | 0.771 | 1.000 | 9845 | 0.469 | 0.751 | 6642 | 0.998 | 1.000 | 6788 | 0.810 | 1.000 | 12630 | 1.000 | 1.000 | 13864 | 0.091 | 0.364 | 9396 | 0.005 | 0.028 | 4457 | 0.136 | 0.377 |
| 2107 | LJI308 | | 21182 | 0.898 | 0.994 | 8128 | 0.757 | 0.994 | 15839 | 0.982 | 0.994 | 11784 | 0.000 | 0.000 | 38167 | 0.987 | 0.994 | 30421 | 0.016 | 0.125 | 20253.5 | 0.678 | 0.994 | 8051.5 | 0.453 | 0.994 | 7866 | 0.032 | 0.126 | 12059 | 0.714 | 0.994 | 27262 | 0.777 | 0.994 | 20949 | 0.994 | 0.994 | 9611 | 0.024 | 0.125 |
| 2109 | AZ6102 | | 18204 | 0.258 | 0.515 | 6334 | 0.125 | 0.399 | 12981.5 | 0.451 | 0.722 | 19086 | 0.384 | 0.683 | 32189 | 0.251 | 0.515 | 36087.5 | 0.700 | 0.999 | 24726 | 0.999 | 0.999 | 10391 | 0.985 | 0.999 | 7413 | 0.012 | 0.093 | 14531 | 0.995 | 0.999 | 29952.5 | 0.987 | 0.999 | 19887 | 0.939 | 0.999 | 5746 | 0.000 | 0.000 |
| 2110 | GSK591 | EPZ015866, GSK3203591 | 17263.5 | 0.108 | 0.347 | 8943.5 | 0.937 | 0.999 | 14492 | 0.855 | 0.999 | 16994 | 0.054 | 0.291 | 37706.5 | 0.985 | 0.999 | 31992 | 0.082 | 0.329 | 24675 | 0.999 | 0.999 | 7832.5 | 0.370 | 0.739 | 7486 | 0.014 | 0.112 | 11744.5 | 0.620 | 0.999 | 20611 | 0.002 | 0.030 | 19328 | 0.884 | 0.999 | 11333 | 0.274 | 0.658 |
| 2111 | VE821 | VE 821, VE-821 | 20244 | 0.754 | 0.994 | 7027 | 0.336 | 0.672 | 9670 | 0.004 | 0.038 | 17778 | 0.138 | 0.367 | 36227 | 0.929 | 0.994 | 37425 | 0.911 | 0.994 | 21373 | 0.883 | 0.994 | 10697 | 0.994 | 0.994 | 8583 | 0.113 | 0.360 | 13631 | 0.970 | 0.994 | 24334 | 0.204 | 0.466 | 20443 | 0.975 | 0.994 | 8842 | 0.005 | 0.038 |
| 2112 | VTP-A | | 11710 | 0.826 | 1.000 | 5602 | 0.679 | 1.000 | 10597 | 0.936 | 1.000 | 9506.5 | 0.751 | 1.000 | 21975.5 | 0.377 | 0.755 | 19109.5 | 0.035 | 0.186 | 12634.5 | 0.996 | 1.000 | 4277 | 0.271 | 0.620 | 8705 | 1.000 | 1.000 | 6257.5 | 0.012 | 0.171 | 14266 | 0.151 | 0.463 | 13383.5 | 0.835 | 1.000 | 4243 | 0.080 | 0.321 |
| 2113 | VTP-B | | 10972 | 0.590 | 0.859 | 6030 | 0.853 | 0.975 | 9012.5 | 0.449 | 0.813 | 9780.5 | 0.834 | 0.975 | 22461.5 | 0.504 | 0.813 | 20504.5 | 0.191 | 0.551 | 12908 | 0.998 | 0.998 | 4437.5 | 0.354 | 0.809 | 6126.5 | 0.508 | 0.813 | 6395 | 0.018 | 0.273 | 14537 | 0.207 | 0.551 | 14688 | 0.984 | 0.998 | 4075 | 0.051 | 0.273 |
| 2145 | PBD-288 | | 9648 | 0.193 | 0.514 | 4321 | 0.091 | 0.290 | 6527 | 0.001 | 0.011 | 9180 | 0.605 | 0.917 | 24068 | 0.866 | 1.000 | 25343 | 0.988 | 1.000 | 8130 | 0.010 | 0.052 | 7060 | 1.000 | 1.000 | 2206 | 0.000 | 0.000 | 12439 | 1.000 | 1.000 | 14910 | 0.272 | 0.622 | 13941 | 0.949 | 1.000 | 5366 | 0.536 | 0.917 |
| 2148 | POMHEX | | 15063.5 | 1.000 | 1.000 | 5450 | 0.581 | 0.929 | 8378 | 0.132 | 0.409 | 9102 | 0.573 | 0.929 | 22970 | 0.647 | 0.941 | 21469 | 0.370 | 0.845 | 6557.5 | 0.000 | 0.000 | 5313 | 0.801 | 1.000 | 2749.5 | 0.000 | 0.000 | 10240 | 0.978 | 1.000 | 20826 | 1.000 | 1.000 | 12293.5 | 0.560 | 0.929 | 2980 | 0.001 | 0.004 |
| 2149 | CT7033-2 | | 12176 | 0.943 | 0.988 | 4385 | 0.106 | 0.339 | 5815 | 0.000 | 0.002 | 8006 | 0.166 | 0.444 | 23717 | 0.809 | 0.988 | 23865 | 0.899 | 0.988 | 8994 | 0.067 | 0.267 | 6301 | 0.988 | 0.988 | 4400 | 0.013 | 0.106 | 9862 | 0.946 | 0.988 | 13576 | 0.048 | 0.256 | 14485 | 0.983 | 0.988 | 6060 | 0.850 | 0.988 |
| 2154 | GSK-LSD1-2HCl | | 9929 | 0.276 | 0.589 | 4734 | 0.220 | 0.589 | 9191 | 0.388 | 0.620 | 10149 | 0.900 | 0.987 | 24840 | 0.947 | 0.987 | 21795.5 | 0.455 | 0.661 | 11977 | 0.917 | 0.987 | 3767 | 0.083 | 0.441 | 7903.5 | 0.987 | 0.987 | 6456 | 0.018 | 0.157 | 15303 | 0.384 | 0.620 | 13605 | 0.909 | 0.987 | 4904 | 0.294 | 0.589 |
| 2156 | 5-azacytidine | | 8702 | 0.037 | 0.278 | 4370 | 0.102 | 0.327 | 11159.5 | 0.958 | 0.978 | 7493 | 0.066 | 0.278 | 22344 | 0.485 | 0.775 | 24744 | 0.969 | 0.978 | 12399 | 0.964 | 0.978 | 6095 | 0.976 | 0.978 | 4496 | 0.018 | 0.278 | 10243 | 0.978 | 0.978 | 14810.5 | 0.247 | 0.489 | 12953 | 0.772 | 0.978 | 4631 | 0.180 | 0.413 |
| 2157 | A-366 | | 11364 | 0.786 | 0.898 | 4936 | 0.309 | 0.695 | 7412 | 0.018 | 0.193 | 9259 | 0.636 | 0.853 | 24118.5 | 0.873 | 0.931 | 22201 | 0.563 | 0.853 | 10913.5 | 0.639 | 0.853 | 4202.5 | 0.224 | 0.695 | 5934 | 0.391 | 0.695 | 6863 | 0.050 | 0.266 | 15295.5 | 0.382 | 0.695 | 15718 | 0.999 | 0.999 | 3853 | 0.024 | 0.193 |
| 2158 | CPI-637 | | 10016.5 | 0.305 | 0.610 | 4068 | 0.046 | 0.214 | 8348.5 | 0.126 | 0.403 | 8032 | 0.173 | 0.462 | 19321 | 0.020 | 0.214 | 22609 | 0.667 | 0.999 | 13845.5 | 0.999 | 0.999 | 5221 | 0.762 | 0.999 | 5617 | 0.249 | 0.569 | 10010.5 | 0.962 | 0.999 | 16540.5 | 0.754 | 0.999 | 13539.5 | 0.899 | 0.999 | 4119 | 0.053 | 0.214 |
| 2159 | UNC0379 | | 10612 | 0.526 | 0.765 | 5002 | 0.342 | 0.765 | 8434 | 0.145 | 0.765 | 10533 | 0.954 | 0.997 | 25053 | 0.961 | 0.997 | 21798.5 | 0.455 | 0.765 | 8057 | 0.008 | 0.064 | 4269 | 0.254 | 0.765 | 5550 | 0.223 | 0.765 | 8194 | 0.420 | 0.765 | 16374 | 0.711 | 0.939 | 15116 | 0.997 | 0.997 | 5329.5 | 0.516 | 0.765 |
| 2169 | AZD6482 | AZD 6482, AZD-6482, AK-55409 | 1128 | 0.035 | 0.186 | 148 | 0.256 | 0.790 | 413 | 0.236 | 0.790 | 2398 | 0.509 | 0.999 | 1254 | 0.296 | 0.790 | 2314 | 0.725 | 0.999 | 2715 | 0.999 | 0.999 | 932 | 0.987 | 0.999 | 801 | 0.899 | 0.999 | 318 | 0.888 | 0.999 | 2058 | 0.910 | 0.999 | 2927 | 0.010 | 0.082 | 764 | 0.001 | 0.017 |
| 2170 | AT13148 | | 13685 | 0.000 | 0.002 | 6444 | 0.250 | 0.571 | 12238 | 0.247 | 0.571 | 22605 | 0.974 | 1.000 | 34160.5 | 0.619 | 0.954 | 36539 | 0.777 | 1.000 | 20016 | 0.622 | 0.954 | 9675 | 0.929 | 1.000 | 7962 | 0.038 | 0.203 | 16674 | 1.000 | 1.000 | 30045.5 | 0.989 | 1.000 | 15340 | 0.071 | 0.283 | 7726 | 0.000 | 0.002 |
| 2171 | BMS-754807 | BMS-754807 | 17949 | 0.439 | 0.879 | 6374 | 0.137 | 0.365 | 7669 | 0.000 | 0.000 | 27217 | 1.000 | 1.000 | 41305.5 | 1.000 | 1.000 | 30505 | 0.034 | 0.109 | 15763.5 | 0.003 | 0.012 | 7853 | 0.381 | 0.871 | 3073 | 0.000 | 0.000 | 11998 | 0.875 | 1.000 | 27994 | 0.914 | 1.000 | 23443 | 1.000 | 1.000 | 14065 | 0.947 | 1.000 |
| 2172 | JQ1 | JQ-1, (+)-JQ-1 | 18254 | 0.433 | 0.824 | 6867 | 0.278 | 0.824 | 15267 | 0.917 | 0.960 | 20204 | 0.746 | 0.960 | 31016 | 0.144 | 0.577 | 34398 | 0.515 | 0.824 | 19415 | 0.323 | 0.824 | 9173 | 0.839 | 0.960 | 8695 | 0.080 | 0.428 | 11890 | 0.855 | 0.960 | 25542 | 0.503 | 0.824 | 20129 | 0.960 | 0.960 | 12067 | 0.502 | 0.824 |
| 2173 | PFI-1 | | 19380 | 0.602 | 0.918 | 6550 | 0.198 | 0.528 | 14642 | 0.846 | 1.000 | 19363 | 0.678 | 0.918 | 26285.5 | 0.001 | 0.008 | 33907.5 | 0.649 | 0.918 | 25035.5 | 1.000 | 1.000 | 7533.5 | 0.293 | 0.669 | 9678 | 0.688 | 0.918 | 8689.5 | 0.030 | 0.158 | 26107.5 | 0.680 | 0.918 | 20900 | 0.996 | 1.000 | 14298 | 0.972 | 1.000 |
| 2174 | IOX2 | IOX-2, IOX 2, AK176060 | 635 | 0.001 | 0.013 | 184 | 0.598 | 0.865 | 361 | 0.331 | 0.793 | 2150 | 0.898 | 0.992 | 1040 | 0.400 | 0.800 | 1752 | 0.632 | 0.865 | 1359 | 0.263 | 0.793 | 441 | 0.347 | 0.793 | 589 | 0.799 | 0.983 | 63 | 0.074 | 0.298 | 1834 | 0.992 | 0.992 | 452 | 0.649 | 0.865 | 1138 | 0.464 | 0.824 |
| 2175 | CHIR-99021 | CT 99021, CHIR99021, CHIR 99021 | 811 | 0.013 | 0.107 | 124.5 | 0.271 | 0.628 | 486 | 0.761 | 0.970 | 2213 | 0.939 | 0.998 | 1225 | 0.789 | 0.970 | 1676 | 0.495 | 0.851 | 2075 | 0.998 | 0.998 | 371 | 0.159 | 0.628 | 418 | 0.275 | 0.628 | 172 | 0.532 | 0.851 | 1666 | 0.941 | 0.998 | 334 | 0.242 | 0.628 | 411 | 0.000 | 0.000 |
| 2177 | SGC0946 | | 20689.5 | 0.865 | 1.000 | 7629 | 0.606 | 1.000 | 16606 | 0.994 | 1.000 | 21172 | 0.949 | 1.000 | 30226 | 0.099 | 0.291 | 29570 | 0.041 | 0.164 | 26079 | 1.000 | 1.000 | 6841 | 0.109 | 0.291 | 10424.5 | 0.882 | 1.000 | 6678 | 0.000 | 0.002 | 27840 | 0.925 | 1.000 | 18685 | 0.873 | 1.000 | 10594 | 0.136 | 0.310 |
| 2359 | GSK2830371 | None | 11559 | 0.916 | 0.971 | 6048 | 0.865 | 0.971 | 9877 | 0.699 | 0.971 | 10316.5 | 0.971 | 0.971 | 23122 | 0.744 | 0.971 | 19501 | 0.112 | 0.448 | 8878.5 | 0.064 | 0.420 | 4003 | 0.162 | 0.518 | 5763.5 | 0.225 | 0.599 | 6502 | 0.079 | 0.420 | 16401 | 0.822 | 0.971 | 13677 | 0.900 | 0.971 | 4156.5 | 0.067 | 0.420 |
| 2360 | THR-101 | WIMM synthesis | 11004 | 0.796 | 0.909 | 6303.5 | 0.927 | 0.975 | 11278 | 0.975 | 0.975 | 9109.5 | 0.722 | 0.888 | 22835 | 0.678 | 0.888 | 20035.5 | 0.197 | 0.790 | 10669 | 0.591 | 0.888 | 4473 | 0.382 | 0.872 | 6825 | 0.716 | 0.888 | 6628 | 0.102 | 0.790 | 14664 | 0.315 | 0.872 | 12702 | 0.657 | 0.888 | 4981 | 0.357 | 0.872 |
| 2361 | THR-102 | WIMM synthesis | 12035 | 0.968 | 0.988 | 5515 | 0.644 | 0.932 | 9775 | 0.661 | 0.932 | 6573 | 0.016 | 0.254 | 23183.5 | 0.757 | 0.932 | 20846.5 | 0.384 | 0.682 | 11084 | 0.738 | 0.932 | 4203 | 0.243 | 0.556 | 5385.5 | 0.109 | 0.350 | 6399 | 0.063 | 0.302 | 17720.5 | 0.977 | 0.988 | 14757.5 | 0.988 | 0.988 | 4465 | 0.141 | 0.375 |
| 2362 | THR-103 | WIMM synthesis | 11837 | 0.951 | 0.996 | 5433 | 0.601 | 0.802 | 8996.5 | 0.349 | 0.789 | 3688 | 0.000 | 0.000 | 26152.5 | 0.996 | 0.996 | 22917.5 | 0.869 | 0.996 | 10335 | 0.462 | 0.800 | 4205 | 0.244 | 0.652 | 5746 | 0.218 | 0.652 | 7857 | 0.549 | 0.800 | 15415.5 | 0.550 | 0.800 | 14959 | 0.993 | 0.996 | 4235 | 0.082 | 0.435 |
| 2438 | ascorbate (vitami | back-up solution from YWKim | 10026.5 | 0.358 | 0.717 | 4248 | 0.045 | 0.381 | 9907 | 0.722 | 0.888 | 7622 | 0.162 | 0.381 | 22409 | 0.702 | 0.888 | 22599 | 0.832 | 0.951 | 12588 | 0.997 | 0.997 | 4527 | 0.421 | 0.748 | 5185 | 0.128 | 0.381 | 6531 | 0.051 | 0.381 | 13704 | 0.080 | 0.381 | 14730 | 0.994 | 0.997 | 4348 | 0.112 | 0.381 |
| 2439 | glutathione | G6013, sigma | 9606.5 | 0.219 | 0.570 | 5723 | 0.613 | 0.818 | 12008 | 0.997 | 0.999 | 6845 | 0.034 | 0.254 | 20664.5 | 0.256 | 0.570 | 21571.5 | 0.602 | 0.818 | 12909 | 0.999 | 0.999 | 4688 | 0.514 | 0.818 | 6622 | 0.761 | 0.937 | 6205.5 | 0.023 | 0.254 | 14741 | 0.285 | 0.570 | 13842 | 0.958 | 0.999 | 4022 | 0.048 | 0.254 |
| 2498 | alpha-lipoic acid | aLA | 11646 | 0.757 | 0.903 | 6930 | 0.991 | 0.991 | 10816 | 0.937 | 0.991 | 7678 | 0.177 | 0.903 | 21571 | 0.427 | 0.903 | 17566.5 | 0.009 | 0.148 | 10668 | 0.789 | 0.903 | 4760 | 0.556 | 0.903 | 6367.5 | 0.776 | 0.903 | 7421 | 0.261 | 0.903 | 15358 | 0.472 | 0.903 | 12835 | 0.790 | 0.903 | 4832 | 0.292 | 0.903 |
| 2499 | N-acetyl cysteine | NAC | 10427 | 0.319 | 0.728 | 5458 | 0.625 | 0.833 | 11815 | 0.995 | 0.995 | 8603 | 0.530 | 0.833 | 20248.5 | 0.140 | 0.728 | 20559 | 0.386 | 0.772 | 12394 | 0.994 | 0.995 | 5228.5 | 0.796 | 0.980 | 5942 | 0.579 | 0.833 | 6686.5 | 0.072 | 0.575 | 14397.5 | 0.199 | 0.728 | 14286 | 0.984 | 0.995 | 4743 | 0.251 | 0.728 |

| DRUG_ID | DRUG_NAME | SYNONYMS | large_intestine_MWU_STAT | large_intestine_PVAL | large_intestine_PVAL_ADJ | thyroid_MWU_STAT | thyroid_PVAL | thyroid_PVAL_ADJ | digestive_system_MWU_STAT | digestive_system_PVAL | digestive_system_PVAL_ADJ | digestive_system_PVAL2 |
|---|---|---|---|---|---|---|---|---|---|---|---|---|
| 2043 | BIBR-1532 | | 15731 | 0.821 | 0.939 | 4119 | 0.038 | 0.403 | 15121 | 0.169 | 0.574 | 0.574 |
| 2044 | Pyridostatin | | 17595 | 0.990 | 1.000 | 5865 | 0.622 | 0.954 | 15520 | 0.256 | 0.820 | 0.820 |
| 2045 | AMG-319 | AMG319 | 17467.5 | 0.986 | 1.000 | 5052 | 0.252 | 0.549 | 15363.5 | 0.214 | 0.549 | 0.549 |
| 2046 | MK-8776 | SCH900776 | 15299.5 | 0.727 | 0.982 | 5825 | 0.597 | 0.982 | 14982 | 0.146 | 0.466 | 0.466 |
| 2047 | Ulixertinib | BVD-523, VRT752271 | 6091.5 | 0.000 | 0.000 | 2615 | 0.009 | 0.029 | 8984 | 0.008 | 0.029 | 0.029 |
| 2048 | Vinorelbine | vinorelbine tartrate, Navelbine, Exelbine | 13431 | 0.041 | 0.123 | 5876 | 0.769 | 1.000 | 17144 | 0.650 | 1.000 | 1.000 |
| 2055 | Mycophenolic acid | | 9380 | 0.062 | 0.142 | 3185 | 0.148 | 0.296 | 11917 | 0.644 | 0.910 | 0.910 |
| 2057 | Remodelin | | 1731 | 0.005 | 0.072 | 1125 | 0.869 | 0.998 | 2281 | 0.080 | 0.320 | 0.320 |
| 2096 | VX-11e | VX11e, VX11e | 6246 | 0.000 | 0.000 | 4854 | 0.312 | 0.623 | 15066 | 0.156 | 0.415 | 0.415 |
| 2106 | Uprosertib | GSK2141795, GSK2141795C, GSK-2141795 | 8074 | 0.003 | 0.028 | 5321 | 0.992 | 1.000 | 11032 | 0.332 | 0.663 | 0.663 |
| 2107 | LJI308 | | 16662 | 0.856 | 0.994 | 5094 | 0.421 | 0.994 | 15600 | 0.260 | 0.831 | 0.831 |
| 2109 | AZ6102 | | 13073.5 | 0.063 | 0.274 | 4648 | 0.227 | 0.515 | 14396 | 0.069 | 0.274 | 0.274 |
| 2110 | GSK591 | EPZ015866, GSK3203591 | 16189 | 0.765 | 0.999 | 4798 | 0.288 | 0.658 | 17592.5 | 0.766 | 0.999 | 0.999 |
| 2111 | VE821 | VE 821, VE-821 | 12084 | 0.013 | 0.072 | 5676 | 0.699 | 0.994 | 14288 | 0.063 | 0.250 | 0.250 |
| 2112 | VTP-A | | 10009 | 0.174 | 0.463 | 2575.5 | 0.021 | 0.171 | 11942.5 | 0.652 | 1.000 | 1.000 |
| 2113 | VTP-B | | 11436 | 0.662 | 0.882 | 2792 | 0.047 | 0.273 | 10638 | 0.206 | 0.551 | 0.551 |
| 2145 | PBD-288 | | 10112 | 0.085 | 0.290 | 4367 | 0.631 | 0.917 | 11524 | 0.378 | 0.756 | 0.756 |
| 2148 | POMHEX | | 9886 | 0.057 | 0.227 | 5120 | 0.933 | 1.000 | 10744 | 0.153 | 0.409 | 0.409 |
| 2149 | CT7033-2 | | 11356 | 0.410 | 0.881 | 4443 | 0.674 | 0.988 | 11701 | 0.441 | 0.881 | 0.881 |
| 2154 | GSK-LSD1-2HCl | | 10596 | 0.177 | 0.589 | 2813 | 0.020 | 0.157 | 11179 | 0.266 | 0.589 | 0.589 |
| 2156 | 5-azacytidine | | 9999 | 0.070 | 0.278 | 3494 | 0.156 | 0.413 | 11209 | 0.275 | 0.489 | 0.489 |
| 2157 | A-366 | | 10965 | 0.278 | 0.695 | 3612 | 0.203 | 0.695 | 12495 | 0.717 | 0.883 | 0.883 |
| 2158 | CPI-637 | | 11847 | 0.589 | 0.999 | 2937 | 0.031 | 0.214 | 13070 | 0.864 | 0.999 | 0.999 |
| 2159 | UNC0379 | | 8956 | 0.007 | 0.064 | 4615 | 0.763 | 0.939 | 11249 | 0.287 | 0.765 | 0.765 |
| 2169 | AZD6482 | AZD 6482, AZD-6482, AK-55409 | 704 | 0.921 | 0.999 | 136 | 0.652 | 0.999 | 601 | 0.741 | 0.999 | 0.999 |
| 2170 | AT13148 | | 15240 | 0.516 | 0.954 | 5585 | 0.656 | 0.954 | 14965 | 0.140 | 0.447 | 0.447 |
| 2171 | BMS-754807 | BMS-754807 | 8782 | 0.000 | 0.000 | 6979 | 0.946 | 1.000 | 17482 | 0.748 | 1.000 | 1.000 |
| 2172 | JQ1 | JQ-1, (+)-JQ-1 | 13297 | 0.037 | 0.428 | 4281 | 0.057 | 0.428 | 17950 | 0.839 | 0.960 | 0.960 |
| 2173 | PFI-1 | | 13804.5 | 0.091 | 0.290 | 2985 | 0.001 | 0.008 | 14427.5 | 0.090 | 0.290 | 0.290 |
| 2174 | IOX2 | IOX-2, IOX 2, AK176060 | 656 | 0.991 | 0.992 | 10 | 0.065 | 0.298 | 223 | 0.042 | 0.298 | 0.298 |
| 2175 | CHIR-99021 | CT 99021, CHIR99021, CHIR 99021 | 479 | 0.740 | 0.970 | 35 | 0.160 | 0.628 | 390.5 | 0.430 | 0.851 | 0.851 |
| 2177 | SGC0946 | | 15378 | 0.414 | 0.827 | 3747 | 0.015 | 0.081 | 11428 | 0.000 | 0.002 | 0.002 |
| 2359 | GSK2830371 | None | 11315 | 0.439 | 0.781 | 3837 | 0.336 | 0.672 | 10883 | 0.289 | 0.661 | 0.661 |
| 2360 | THR-101 | WIMM synthesis | 11324 | 0.442 | 0.885 | 3444.5 | 0.151 | 0.790 | 9564 | 0.037 | 0.594 | 0.594 |
| 2361 | THR-102 | WIMM synthesis | 11130.5 | 0.373 | 0.682 | 3187 | 0.076 | 0.302 | 9580 | 0.038 | 0.302 | 0.302 |
| 2362 | THR-103 | WIMM synthesis | 10336 | 0.143 | 0.573 | 3937 | 0.395 | 0.789 | 9790 | 0.058 | 0.435 | 0.435 |
| 2438 | ascorbate (vitami | back-up solution from YWKim | 11487 | 0.518 | 0.754 | 3475 | 0.167 | 0.381 | 11421.5 | 0.494 | 0.754 | 0.754 |
| 2439 | glutathione | G6013, sigma | 10812.5 | 0.279 | 0.570 | 3859.5 | 0.357 | 0.634 | 10688 | 0.241 | 0.570 | 0.570 |
| 2498 | alpha-lipoic acid | aLA | 12059 | 0.719 | 0.903 | 3919 | 0.392 | 0.903 | 10341 | 0.152 | 0.903 | 0.903 |
| 2499 | N-acetyl cysteine | NAC | 8871 | 0.008 | 0.131 | 3766 | 0.304 | 0.728 | 11343 | 0.464 | 0.826 | 0.826 |

Suppl. Table 9

| broad_id | name | colorectal_MWU_STAT | colorectal_PVAL | colorectal_PVAL_ADJ | skin_MWU_STAT | skin_PVAL | skin_PVAL_ADJ | urinary_tract_MWU_STAT | urinary_tract_PVAL | urinary_tract_PVAL_ADJ | lung_MWU_STAT | lung_PVAL | lung_PVAL_ADJ | ovary_MWU_STAT | ovary_PVAL | ovary_PVAL_ADJ | breast_MWU_STAT | breast_PVAL | breast_PVAL_ADJ | pancreas_MWU_STAT | pancreas_PVAL | pancreas_PVAL_ADJ | nervous_system_MWU_STAT | central_nervous_system_PVAL | nervous_system_PVAL_ADJ | soft_tissue_MWU_STAT | soft_tissue_PVAL | soft_tissue_PVAL_ADJ | kidney_MWU_STAT | kidney_PVAL | kidney_PVAL_ADJ | gastric_MWU_STAT | gastric_PVAL | gastric_PVAL_ADJ | bone_MWU_STAT | bone_PVAL | bone_PVAL_ADJ | prostate_MWU_STAT | prostate_PVAL | prostate_PVAL_ADJ | al_nervous_system_MWU_STAT | peripheral_nervous_system_PVAL |
|---|---|---|---|---|---|---|---|---|---|---|---|---|---|---|---|---|---|---|---|---|---|---|---|---|---|---|---|---|---|---|---|---|---|---|---|---|---|---|---|---|---|---|
| BRD-A00758722-001-04-9 | noretynodrel | 1258 | 0.720 | 0.858 | 2798 | 0.860 | 0.864 | 1276.5 | 0.228 | 1.000 | 5625.5 | 0.864 | 0.864 | 1822 | 0.730 | 0.858 | 440 | 0.064 | 0.192 | 1803 | 0.553 | 0.830 | 1717 | 0.735 | 0.858 | 584 | 0.256 | 0.651 | 822.5 | 0.540 | 0.830 | 821 | 0.310 | 0.651 | 421 | 0.372 | 0.710 | 287 | 0.295 | 0.651 | 115 | 0.023 |
| BRD-A03506276-001-01-5 | XL888 | 5959 | 0.877 | 0.996 | 9483 | 0.996 | 0.996 | 3424 | 0.019 | 0.199 | 15323 | 0.249 | 0.655 | 4543 | 0.046 | 0.325 | 4742 | 0.815 | 0.996 | 6385 | 0.384 | 0.806 | 7103 | 0.846 | 0.996 | 1940 | 0.284 | 0.662 | 4525 | 0.958 | 0.996 | 2284 | 0.012 | 0.199 | 2119 | 0.141 | 0.590 | 1100 | 0.799 | 0.996 | 1345 | 0.802 |
| BRD-A03932035-004-04-3 | methscopolamine | 245.5 | 0.839 | 0.839 | 496.5 | 0.600 | 1.000 | 343.5 | 0.098 | 0.938 | 1186 | 0.184 | 0.938 | 561 | 0.268 | 0.938 | 366 | 0.957 | 1.000 | 778 | 0.898 | 1.000 | 500.5 | 0.236 | 0.938 | 92 | 0.797 | 1.000 | 273 | 0.159 | 0.938 | 232 | 0.781 | 1.000 | 109.5 | 0.906 | 1.000 | 148 | 0.685 | 1.000 | 134 | 0.582 |
| BRD-A05906449-004-01-1 | lappaconite | 3245 | 0.337 | 0.807 | 5644 | 0.792 | 0.965 | 2492 | 0.214 | 0.748 | 9002 | 0.070 | 0.661 | 3062 | 0.126 | 0.661 | 3049 | 0.720 | 0.965 | 4710 | 0.982 | 0.982 | 4556 | 0.858 | 0.965 | 1483 | 0.670 | 0.965 | 2677 | 0.973 | 0.982 | 2036 | 0.346 | 0.807 | 1112 | 0.873 | 0.965 | 505 | 0.472 | 0.902 | 501 | 0.176 |
| BRD-A06352418-001-25-9 | terfenadine | 407 | 0.133 | 1.000 | 1254.5 | 0.413 | 0.712 | 825.5 | 0.948 | 0.948 | 2524.5 | 0.748 | 0.924 | 808.5 | 0.443 | 0.712 | 604 | 0.465 | 0.712 | 665 | 0.649 | 0.852 | 1015.5 | 0.615 | 0.852 | 243 | 0.897 | 0.942 | 354.5 | 0.158 | 0.712 | 313 | 0.222 | 0.712 | 196.5 | 0.303 | 0.712 | 37 | 0.032 | 0.221 | 231 | 0.861 |
| BRD-A06352508-001-03-7 | SB-218078 | 1233 | 0.154 | 0.404 | 2732 | 0.272 | 0.572 | 1388 | 0.108 | 0.384 | 6449 | 0.896 | 0.990 | 2041 | 0.128 | 0.384 | 1261 | 0.882 | 0.990 | 2063 | 0.779 | 0.990 | 2439 | 0.793 | 0.990 | 515 | 0.029 | 0.203 | 774 | 0.020 | 0.203 | 691 | 0.023 | 0.203 | 490 | 0.186 | 0.433 | 373 | 0.859 | 0.990 | 590 | 0.941 |
| BRD-A06426627-001-03-5 | diclazuril | 2735 | 0.280 | 0.882 | 5656 | 0.859 | 0.983 | 2477 | 0.198 | 0.882 | 9408 | 0.230 | 0.882 | 3310 | 0.108 | 0.882 | 2915 | 0.738 | 0.969 | 4643 | 0.889 | 0.983 | 4700 | 0.953 | 0.993 | 1632 | 0.634 | 0.969 | 2185 | 0.683 | 0.969 | 1890 | 0.538 | 0.942 | 1114 | 0.192 | 0.447 | 243 | 0.699 | 0.969 | 642 | 0.410 |
| BRD-A06627858-236-03-0 | monensin | 3282 | 0.424 | 0.891 | 5783 | 0.043 | 0.432 | 4549 | 0.538 | 0.941 | 15835.5 | 0.933 | 0.980 | 4298 | 0.062 | 0.432 | 3270.5 | 0.415 | 0.891 | 5591 | 0.729 | 0.957 | 5148 | 0.166 | 0.662 | 1209 | 0.108 | 0.567 | 2870 | 0.252 | 0.662 | 3440 | 0.708 | 0.957 | 1581 | 0.249 | 0.662 | 941 | 0.695 | 0.957 | 1042 | 0.530 |
| BRD-A07986123-001-02-8 | galeterone | 2971.5 | 0.863 | 0.919 | 4724.5 | 0.919 | 0.919 | 2319 | 0.545 | 0.818 | 9663 | 0.730 | 0.919 | 1807 | 0.009 | 0.062 | 2315.5 | 0.249 | 0.746 | 3438.5 | 0.870 | 0.919 | 3927 | 0.890 | 0.919 | 1316 | 0.824 | 0.919 | 1888 | 0.379 | 0.750 | 1802 | 0.162 | 1.000 | 557 | 0.133 | 0.559 | 775 | 0.786 | 0.919 | 222 | 0.234 |
| BRD-A08545410-003-07-8 | doxycycline | 2063 | 0.830 | 0.860 | 3106 | 0.860 | 0.860 | 1212 | 0.202 | 0.860 | 6485 | 0.739 | 0.860 | 2300 | 0.399 | 0.860 | 762 | 0.219 | 0.860 | 3036 | 0.708 | 0.860 | 2159 | 0.800 | 0.860 | 460 | 0.012 | 0.126 | 1247 | 0.624 | 0.860 | 1518 | 0.456 | 0.860 | 332 | 0.008 | 0.126 | 526 | 0.486 | 0.860 | 511 | 0.795 |
| BRD-A08545410-311-03-4 | doxycycline | 1803 | 0.926 | 0.990 | 2653 | 0.827 | 0.965 | 1046 | 0.753 | 0.947 | 5524 | 0.964 | 0.990 | 1257 | 0.002 | 0.010 | 1403 | 0.414 | 1.000 | 1551 | 0.485 | 0.783 | 1685 | 0.692 | 0.947 | 504 | 0.128 | 0.335 | 973 | 0.397 | 0.754 | 889 | 0.249 | 1.000 | 782 | 0.990 | 0.990 | 372 | 0.572 | 0.858 | 330 | 0.431 |
| BRD-A08840375-001-01-3 | CID-5458317 | 4405 | 0.968 | 0.989 | 5431 | 0.341 | 0.717 | 2576 | 0.340 | 0.717 | 13153 | 0.989 | 0.989 | 3145 | 0.010 | 0.079 | 2760 | 0.516 | 0.877 | 3581 | 0.884 | 0.989 | 4275 | 0.699 | 0.917 | 873 | 0.011 | 0.079 | 2326 | 0.755 | 0.933 | 2327 | 0.585 | 0.877 | 875 | 0.004 | 0.076 | 387 | 0.581 | 0.877 | 308 | 0.110 |
| BRD-A10039652-001-01-2 | tipranavir | 1151 | 0.975 | 0.975 | 1038 | 0.168 | 1.000 | 944 | 0.562 | 0.909 | 2776 | 0.523 | 0.909 | 1036 | 0.026 | 0.108 | 577 | 0.899 | 0.975 | 1079 | 0.516 | 0.909 | 796 | 0.231 | 0.807 | 222 | 0.326 | 0.909 | 409 | 0.727 | 0.975 | 410 | 0.029 | 0.613 | 209 | 0.678 | 0.975 | 244 | 0.828 | 0.975 | 287 | 0.938 |
| BRD-A10070317-003-28-3 | propranolol | 398 | 0.949 | 0.969 | 430 | 0.390 | 1.000 | 240 | 0.792 | 0.969 | 667 | 0.165 | 1.000 | 268 | 0.208 | 1.000 | 192 | 0.490 | 1.000 | 171 | 0.347 | 1.000 | 308 | 0.859 | 0.969 | 90 | 0.035 | 0.074 | 102 | 0.534 | 0.863 | 122 | 0.717 | 0.969 | 87 | 0.002 | 0.005 | 50 | 0.507 | 0.863 | 1 | 0.048 |
| BRD-A10977446-001-12-1 | carvedilol | 883 | 0.929 | 0.997 | 1406.5 | 0.729 | 0.956 | 1086 | 0.854 | 0.997 | 2189 | 0.053 | 1.000 | 1238 | 0.702 | 0.956 | 886 | 0.808 | 0.997 | 987 | 0.675 | 0.956 | 1458.5 | 0.958 | 0.997 | 114 | 0.048 | 0.144 | 55 | 0.262 | 0.551 | 888.5 | 0.442 | 0.844 | 63 | 0.061 | 0.160 | 190 | 0.585 | 0.956 | 29 | 0.022 |
| BRD-A11815689-001-01-1 | proscillaridin-a | 4535 | 1.000 | 1.000 | 5898 | 0.982 | 1.000 | 2795 | 0.460 | 0.744 | 8030 | 0.022 | 0.234 | 2980 | 0.055 | 0.306 | 2888 | 0.991 | 1.000 | 3413 | 0.367 | 0.744 | 3187 | 0.418 | 0.744 | 1206 | 0.600 | 0.841 | 1867 | 0.429 | 0.744 | 3078 | 0.971 | 1.000 | 489 | 0.062 | 0.306 | 308 | 0.138 | 0.482 | 53 | 0.014 |
| BRD-A12230535-001-06-7 | nutlin-3 | 121 | 0.009 | 0.115 | 475 | 0.011 | 0.115 | 447 | 0.768 | 0.989 | 1443.5 | 0.989 | 0.989 | 497.5 | 0.533 | 0.933 | 274.5 | 0.777 | 0.989 | 417.5 | 0.976 | 0.989 | 390 | 0.282 | 0.738 | 138.5 | 0.074 | 0.260 | 112 | 0.007 | 0.035 | 283.5 | 0.244 | 0.731 | 198 | 0.316 | 0.738 | 131.5 | 0.642 | 0.989 | 9 | 0.058 |
| BRD-A13133631-001-04-0 | fluorometholone | 1479.5 | 0.328 | 0.765 | 4016.5 | 0.939 | 0.998 | 2163 | 0.727 | 0.954 | 6579 | 0.565 | 0.912 | 1519 | 0.001 | 0.004 | 2148.5 | 0.995 | 0.998 | 2294 | 0.198 | 0.571 | 2255 | 0.818 | 0.990 | 787 | 0.218 | 0.571 | 1547 | 0.998 | 0.998 | 714 | 0.006 | 0.124 | 668.5 | 0.464 | 0.886 | 143 | 0.522 | 0.912 | 71 | 0.035 |
| BRD-A14344385-001-03-2 | IDRA-21 | 435 | 0.165 | 0.480 | 904.5 | 0.802 | 0.901 | 821.5 | 0.612 | 0.857 | 2837 | 0.662 | 0.869 | 812.5 | 0.399 | 0.698 | 422 | 0.309 | 0.591 | 843.5 | 0.183 | 0.480 | 932 | 0.850 | 0.901 | 516.5 | 0.860 | 0.901 | 630 | 0.294 | 0.591 | 130 | 0.083 | 0.917 | 230 | 0.433 | 0.699 |  |  |  | 1 | 0.045 |
| BRD-A17883755-001-06-1 | lenalidomide | 911 | 0.890 | 0.991 | 1193.5 | 0.102 | 0.598 | 583 | 0.299 | 0.785 | 2729.5 | 0.897 | 0.991 | 1023 | 0.806 | 0.991 | 733 | 0.171 | 0.668 | 941.5 | 0.638 | 0.980 | 1067 | 0.741 | 0.980 | 69 | 0.374 | 0.815 | 331 | 0.217 | 0.668 | 353 | 0.114 | 0.598 | 307.5 | 0.388 | 0.815 | 210 | 0.719 | 0.980 | 257 | 0.519 |
| BRD-A18696154-003-22-6 | idazoxan | 803.5 | 0.896 | 0.967 | 2134.5 | 0.795 | 0.967 | 1142 | 0.390 | 1.000 | 4115.5 | 0.685 | 0.967 | 1575 | 0.666 | 0.967 | 1243 | 0.967 | 0.967 | 1349 | 0.766 | 0.967 | 1233 | 0.246 | 0.506 | 432 | 0.265 | 0.506 | 1139 | 0.753 | 0.967 | 643 | 0.026 | 0.283 | 115 | 0.030 | 0.090 | 333.5 | 0.925 | 0.967 | 98 | 0.098 |
| BRD-A22707317-001-10-8 | SB-205384 | 821.5 | 0.236 | 1.000 | 1763.5 | 0.987 | 0.987 | 1327.5 | 0.712 | 0.967 | 2507 | 0.011 | 0.238 | 1710 | 0.933 | 0.987 | 569 | 0.240 | 0.631 | 1266 | 0.600 | 0.913 | 832.5 | 0.853 | 0.987 | 298 | 0.069 | 0.243 | 662.5 | 0.467 | 0.863 | 998.5 | 0.608 | 0.913 | 98 | 0.493 | 0.863 | 283 | 0.853 | 0.987 | 249 | 0.737 |
| BRD-A23072235-050-18-5 | pheniramine | 818.5 | 0.026 | 0.386 | 2586 | 0.736 | 0.910 | 1099 | 0.268 | 0.564 | 4344 | 0.897 | 0.974 | 1410.5 | 0.207 | 0.544 | 1052 | 0.974 | 0.974 | 1749 | 0.721 | 0.910 | 1405.5 | 0.642 | 0.910 | 237 | 0.065 | 0.227 | 638 | 0.956 | 0.974 | 761 | 0.172 | 0.516 | 235 | 0.592 | 0.910 |  |  |  | 13 | 0.011 |
| BRD-A25004090-001-08-4 | erastin | 2056 | 0.982 | 0.982 | 2755 | 0.909 | 1.000 | 1448 | 0.591 | 1.000 | 5483 | 0.170 | 0.446 | 1670 | 0.041 | 0.216 | 1114 | 0.442 | 0.928 | 2832 | 0.884 | 1.000 | 2393 | 0.140 | 0.437 | 548 | 0.012 | 0.216 | 1008 | 0.026 | 0.216 | 1776 | 0.940 | 1.000 | 465 | 0.146 | 0.437 | 183 | 0.246 | 0.575 | 73 | 0.041 |
| BRD-A25569250-001-03-3 | KI-16425 | 298 | 0.126 | 1.000 | 600 | 0.647 | 0.850 | 210 | 0.145 | 1.000 | 1776 | 0.872 | 0.951 | 329 | 0.211 | 1.000 | 308 | 0.320 | 1.000 | 436 | 0.853 | 0.951 | 285 | 0.464 | 0.812 | 113 | 0.039 | 0.103 | 4 | 0.056 | 0.130 | 362 | 0.568 | 0.918 |  |  |  | 72 | 0.633 | 0.850 | 73 | 0.174 |
| BRD-A25608658-001-01-1 | narasin | 6193 | 0.993 | 0.993 | 6635 | 0.218 | 0.555 | 5181 | 0.928 | 0.993 | 16140 | 0.704 | 0.993 | 5736 | 0.401 | 0.762 | 4737 | 0.926 | 0.993 | 7457 | 0.960 | 0.993 | 5823 | 0.175 | 0.526 | 1113 | 0.002 | 0.032 | 2116 | 0.006 | 0.063 | 3240 | 0.264 | 0.555 | 2060 | 0.645 | 0.993 | 533 | 0.094 | 0.328 | 364 | 0.023 |
| BRD-A25687296-300-04-3 | emetine | 5586 | 0.482 | 0.870 | 7669 | 0.538 | 0.870 | 5146 | 0.592 | 0.888 | 15574 | 0.090 | 0.378 | 5020 | 0.070 | 0.370 | 6243 | 0.999 | 0.999 | 9240 | 0.999 | 0.999 | 7990 | 0.909 | 0.999 | 2387 | 0.262 | 0.611 | 4483 | 0.915 | 0.999 | 2389 | 0.014 | 0.165 | 2877 | 0.672 | 0.941 | 485 | 0.056 | 0.370 | 1129 | 0.496 |
| BRD-A26032986-050-02-1 | zaldaride | 127 | 0.312 | 1.000 | 370 | 0.126 | 1.000 | 466 | 0.829 | 0.929 | 1009 | 0.529 | 0.845 | 254 | 0.278 | 1.000 | 169 | 0.625 | 0.875 | 308 | 0.563 | 0.845 | 329 | 0.885 | 0.929 | 80 | 0.021 | 0.062 | 95 | 0.134 | 0.312 | 154 | 0.512 | 0.845 | 105 | 0.180 | 0.378 | 47 | 0.441 | 0.842 | 1 | 0.047 |
| BRD-A27376179-001-01-3 | lestaurtinib | 4548 | 0.642 | 0.959 | 3993 | 0.003 | 0.057 | 4096 | 0.694 | 0.959 | 13274 | 0.917 | 0.959 | 3871 | 0.114 | 0.477 | 3273 | 0.873 | 0.959 | 6018 | 0.959 | 0.959 | 5423 | 0.656 | 0.959 | 1533 | 0.522 | 0.959 | 2716 | 0.601 | 0.959 | 1738 | 0.040 | 0.282 | 1079 | 0.030 | 0.282 | 735 | 0.786 | 0.959 | 445 | 0.259 |
| BRD-A27883417-300-01-6 | alexidine | 3112 | 0.685 | 0.899 | 6550 | 0.999 | 0.999 | 2333 | 0.375 | 0.788 | 9421 | 0.256 | 1.000 | 3425 | 0.130 | 0.907 | 2709 | 0.985 | 0.999 | 4188 | 0.566 | 0.899 | 4750 | 0.844 | 0.999 | 1408 | 0.602 | 0.899 | 1957 | 0.093 | 0.279 | 1559 | 0.106 | 0.907 | 915 | 0.164 | 0.431 | 388 | 0.632 | 0.899 | 564 | 0.941 |
| BRD-A28301258-001-03-0 | salinomycin | 2702 | 0.902 | 1.000 | 2347 | 0.044 | 0.460 | 1898 | 0.461 | 0.875 | 5847 | 0.599 | 0.967 | 2375 | 0.466 | 0.875 | 1630 | 0.905 | 0.983 | 2307 | 0.802 | 1.000 | 2493 | 0.153 | 0.801 | 822 | 0.305 | 0.875 | 1516 | 0.273 | 0.875 | 1034 | 0.461 | 0.875 | 525 | 0.816 | 1.000 | 116 | 0.083 | 0.581 | 667 | 0.500 |
| BRD-A28467416-002-01-0 | PKI-179 | 6139 | 0.990 | 0.990 | 9374 | 0.973 | 0.990 | 5092 | 0.721 | 0.947 | 17539 | 0.681 | 0.947 | 4164 | 0.004 | 0.056 | 5244 | 0.887 | 0.990 | 8361 | 0.976 | 0.990 | 6517 | 0.380 | 0.798 | 1796 | 0.029 | 0.138 | 3688 | 0.646 | 0.947 | 2354 | 0.005 | 0.056 | 2384 | 0.279 | 0.734 | 547 | 0.093 | 0.326 | 1275 | 0.698 |
| BRD-A28970875-300-04-9 | puromycin | 4278 | 0.967 | 1.000 | 6334 | 0.982 | 1.000 | 1825 | 0.024 | 0.248 | 10054 | 0.112 | 0.262 | 3794 | 0.397 | 0.695 | 2733 | 0.451 | 0.729 | 5815 | 0.992 | 1.000 | 4380 | 0.826 | 1.000 | 950 | 0.019 | 0.066 | 3950 | 1.000 | 1.000 | 1642 | 0.007 | 0.153 | 1299 | 0.031 | 0.093 | 396 | 0.222 | 0.423 | 351 | 0.155 |
| BRD-A30886737-001-02-9 | florfenicol | 251 | 0.688 | 0.688 | 1078 | 0.992 | 1.000 | 443 | 0.795 | 1.000 | 797 | 0.093 | 0.486 | 531 | 0.886 | 1.000 | 214 | 0.208 | 0.398 | 775 | 0.979 | 1.000 | 278 | 0.533 | 0.896 | 138 | 0.102 | 0.268 | 197 | 0.147 | 0.308 | 241 | 0.631 | 0.896 |  |  |  | 19 | 0.128 | 0.298 | 0 | 0.042 |
| BRD-A31095847-001-02-3 | mozavaptan | 2695 | 0.862 | 0.935 | 3391 | 0.388 | 0.926 | 2346 | 0.250 | 0.875 | 7392 | 0.449 | 0.926 | 3054 | 0.563 | 0.926 | 2557 | 0.755 | 0.926 | 2502 | 0.702 | 0.926 | 4180 | 0.972 | 0.972 | 1193 | 0.733 | 0.926 | 2010 | 0.794 | 0.926 | 1609 | 0.681 | 0.926 | 568 | 0.018 | 0.141 | 716 | 0.732 | 0.926 | 749 | 0.494 |
| BRD-A35588707-001-05-5 | teniposide | 6727 | 0.934 | 0.987 | 6691 | 0.098 | 0.342 | 4667 | 0.331 | 0.870 | 14995 | 0.081 | 0.342 | 4939 | 0.040 | 0.278 | 5118 | 0.987 | 0.987 | 8194 | 0.981 | 0.987 | 8217 | 0.982 | 0.987 | 1288 | 0.001 | 0.026 | 4196 | 0.916 | 0.987 | 2794 | 0.088 | 0.342 | 2806 | 0.645 | 0.987 | 1007 | 0.660 | 0.987 | 1032 | 0.388 |
| BRD-A36057565-001-01-0 | bruceantin | 5198 | 0.626 | 0.995 | 6427 | 0.051 | 0.356 | 4665 | 0.539 | 0.995 | 15548 | 0.210 | 0.646 | 5590 | 0.197 | 0.646 | 5735 | 0.973 | 0.995 | 7025 | 0.944 | 0.995 | 6854 | 0.673 | 0.995 | 1656 | 0.045 | 0.356 | 2872 | 0.964 | 0.995 | 3005 | 0.215 | 0.646 | 1725 | 0.728 | 0.995 | 784 | 0.367 | 0.856 | 1252 | 0.711 |
| BRD-A36267905-003-18-4 | buphenine | 292 | 0.922 | 1.000 | 402.5 | 0.431 | 0.836 | 318 | 0.243 | 0.808 | 1148.5 | 0.783 | 1.000 | 366.5 | 0.774 | 1.000 | 400.5 | 0.683 | 1.000 | 508.5 | 0.295 | 0.808 | 324.5 | 0.272 | 0.808 |  |  |  | 103.5 | 0.438 | 0.836 | 292 | 0.922 | 0.922 | 146 | 0.358 | 0.836 | 85.5 | 0.266 | 0.808 | 3 | 0.028 |
| BRD-A36331462-001-02-1 | oprozomib | 5090 | 0.779 | 0.909 | 4469 | 0.013 | 0.264 | 5017 | 0.954 | 0.989 | 13973 | 0.914 | 0.989 | 5005 | 0.731 | 0.909 | 2280 | 0.088 | 0.461 | 5018 | 0.204 | 0.612 | 4805 | 0.180 | 0.612 | 1804 | 0.773 | 0.909 | 2659 | 0.190 | 0.612 | 2050 | 0.030 | 0.264 | 1715 | 0.459 | 0.894 | 519 | 0.364 | 0.850 | 512 | 0.352 |
| BRD-A38350138-001-04-9 | ketoconazole | 529 | 0.743 | 0.975 | 709 | 0.704 | 0.975 | 296 | 0.377 | 0.917 | 1055 | 0.746 | 0.870 | 461 | 0.167 | 0.612 | 226 | 0.553 | 0.975 | 545 | 0.621 | 0.975 | 259 | 0.442 | 0.928 | 123 | 0.229 | 0.688 | 58 | 0.126 | 0.612 | 200 | 0.393 | 0.917 | 24 | 0.030 | 0.381 | 101 | 0.919 | 0.965 | 25 | 0.175 |
| BRD-A39390670-236-10-7 | rabeprazole | 123 | 0.665 | 0.936 | 603 | 0.781 | 0.911 | 135 | 0.349 | 0.916 | 1038 | 0.857 | 0.947 | 516 | 0.183 | 0.747 | 223 | 0.613 | 0.936 | 321 | 0.188 | 0.747 | 433 | 0.672 | 0.936 | 28 | 0.213 | 0.747 |  |  |  | 394 | 0.713 | 0.936 | 55 | 0.127 | 0.747 | 134 | 0.342 | 0.916 | 59 | 0.584 |
| BRD-A39935389-001-05-9 | teroxirone | 2243 | 0.699 | 0.968 | 3093 | 0.379 | 0.956 | 1513 | 0.281 | 0.956 | 7059 | 0.150 | 0.956 | 2527 | 0.299 | 0.956 | 1844 | 0.371 | 0.956 | 2801 | 0.721 | 0.968 | 2659 | 0.830 | 0.968 | 562 | 0.076 | 0.956 | 2177 | 0.959 | 0.976 | 1343 | 0.400 | 0.956 | 1146 | 0.746 | 0.968 | 634 | 0.632 | 0.968 | 253 | 0.383 |
| BRD-A41304429-001-14-8 | practolol | 1199 | 0.128 | 0.714 | 2152.5 | 0.215 | 0.752 | 1826 | 0.975 | 0.987 | 5741 | 0.902 | 0.987 | 1498 | 0.137 | 0.714 | 1325 | 0.615 | 0.760 | 2059 | 0.586 | 0.760 | 1503 | 0.240 | 0.504 | 161 | 0.052 | 0.137 | 1625 | 0.987 | 0.987 | 1207 | 0.409 | 0.760 | 455.5 | 0.454 | 0.760 | 235 | 0.490 | 0.760 | 214 | 0.915 |
| BRD-A42699921-001-03-6 | teriflunomide | 921.5 | 0.926 | 0.989 | 1889.5 | 0.545 | 0.831 | 685 | 0.248 | 0.734 | 2824.5 | 0.280 | 0.734 | 1477.5 | 0.846 | 1.000 | 554 | 0.554 | 0.831 | 967.5 | 0.525 | 0.831 | 1078 | 0.269 | 0.734 | 136 | 0.017 | 0.349 | 442 | 0.475 | 0.831 | 774 | 0.271 | 0.734 | 354.5 | 0.989 | 0.989 | 78 | 0.400 | 0.831 | 19 | 0.087 |
| BRD-A42746051-001-02-9 | melengestrol-acetate | 377 | 0.535 | 0.864 | 323 | 0.799 | 0.917 | 356 | 0.423 | 1.000 | 1154 | 0.254 | 1.000 | 378 | 0.143 | 1.000 | 112 | 0.146 | 1.000 | 343 | 0.356 | 1.000 | 532 | 0.985 | 0.985 | 1 | 0.030 | 0.070 | 234 | 0.919 | 0.964 | 80 | 0.228 | 1.000 | 79 | 0.791 | 0.917 |  |  |  | 43 | 0.333 |
| BRD-A43671941-003-11-9 | oxprenolol | 1562 | 0.829 | 0.829 | 2401 | 0.148 | 0.707 | 2579.5 | 0.991 | 1.000 | 5472.5 | 0.286 | 0.751 | 1650 | 0.186 | 0.707 | 1358 | 0.882 | 1.000 | 2190 | 0.726 | 1.000 | 1580.5 | 0.222 | 0.707 | 593 | 0.429 | 0.819 | 928 | 0.211 | 0.707 | 1127.5 | 0.759 | 1.000 | 394.5 | 0.236 | 0.707 | 546 | 0.625 | 1.000 | 242 | 0.039 |
| BRD-A44090213-001-25-6 | indoprofen | 909 | 0.232 | 0.938 | 1913 | 0.024 | 0.504 | 1731 | 0.918 | 0.997 | 4818 | 0.155 | 0.938 | 1870 | 0.176 | 0.938 | 1498 | 0.997 | 0.997 | 1622 | 0.826 | 0.997 | 2128 | 0.616 | 0.997 | 937 | 0.901 | 0.997 | 1268 | 0.665 | 0.997 | 1054 | 0.486 | 0.927 | 668 | 0.908 | 0.997 | 538 | 0.925 | 0.997 | 46 | 0.024 |
| BRD-A44500574-001-01-8 | FLI-06 | 1329 | 0.934 | 0.980 | 2487 | 0.308 | 0.892 | 1235 | 0.487 | 0.929 | 4317 | 0.130 | 0.680 | 2001 | 0.811 | 0.978 | 661 | 0.425 | 0.892 | 1859 | 0.885 | 0.978 | 2212 | 0.806 | 0.978 | 581 | 0.105 | 0.680 | 832 | 0.013 | 0.133 | 903 | 0.709 | 0.978 | 529 | 0.162 | 0.680 | 91 | 0.349 | 0.892 | 55 | 0.006 |
| BRD-A45498368-001-07-1 | WYE-125132 | 4069 | 0.970 | 0.988 | 5543 | 0.862 | 0.988 | 2761 | 0.357 | 1.000 | 9469 | 0.192 | 1.000 | 3562 | 0.170 | 1.000 | 2641 | 0.256 | 1.000 | 5280 | 0.884 | 0.988 | 5339 | 0.904 | 0.988 | 1209 | 0.115 | 0.268 | 2004 | 0.407 | 0.763 | 2079 | 0.490 | 0.763 | 863 | 0.216 | 0.454 | 716 | 0.509 | 0.763 | 936 | 0.861 |
| BRD-A48237631-001-04-8 | mitomycin-c | 4365 | 0.297 | 0.900 | 6618 | 0.702 | 0.985 | 4185 | 0.295 | 0.900 | 11128 | 0.001 | 0.015 | 4043 | 0.082 | 0.857 | 4041 | 0.837 | 0.985 | 6404 | 0.911 | 0.985 | 7351 | 0.987 | 0.987 | 1632 | 0.152 | 0.900 | 3833 | 0.920 | 0.985 | 2608 | 0.203 | 0.900 | 2032 | 0.519 | 0.985 | 551 | 0.386 | 0.900 | 1099 | 0.877 |
| BRD-A48300215-001-04-8 | ajmaline | 425 | 0.848 | 0.940 | 638 | 0.850 | 0.940 | 518 | 0.707 | 0.940 | 1862 | 0.516 | 0.910 | 603 | 0.563 | 0.910 | 442 | 0.111 | 0.582 | 265 | 0.029 | 0.304 | 997 | 0.989 | 0.989 | 369 | 0.896 | 0.940 | 447 | 0.693 | 0.940 | 101 | 0.017 | 0.304 | 274 | 0.847 | 0.940 | 72 | 0.535 | 0.910 | 126 | 0.426 |
| BRD-A49035384-003-28-9 | methacycline | 841 | 0.976 | 0.976 | 1811 | 0.458 | 0.991 | 1117 | 0.708 | 1.000 | 3003 | 0.295 | 0.991 | 1478 | 0.424 | 0.991 | 1475 | 0.838 | 1.000 | 1000 | 0.657 | 1.000 | 1259 | 0.472 | 0.991 | 357 | 0.060 | 0.421 | 401 | 0.875 | 1.000 | 556 | 0.736 | 1.000 | 217 | 0.616 | 1.000 | 50 | 0.036 | 0.378 | 24 | 0.098 |
| BRD-A49358627-001-14-6 | ciprofibrate | 396 | 0.109 | 0.363 | 748 | 0.842 | 0.977 | 233 | 0.194 | 0.506 | 1207.5 | 0.559 | 0.839 | 450 | 0.121 | 0.363 | 339.5 | 0.681 | 0.953 | 693 | 0.848 | 0.977 | 726 | 0.971 | 0.977 | 214.5 | 0.343 | 0.600 | 166 | 0.797 | 0.977 | 135 | 0.065 | 1.000 | 97 | 0.082 | 0.343 | 51 | 0.381 | 0.615 | 9 | 0.072 |
| BRD-A49765801-001-04-1 | fludroxycortide | 717 | 0.493 | 0.740 | 1794 | 0.800 | 0.938 | 825 | 0.350 | 1.000 | 3223 | 0.894 | 0.968 | 1214 | 0.341 | 0.659 | 215 | 0.078 | 0.329 | 1115 | 0.184 | 0.483 | 1295 | 0.992 | 0.992 | 243 | 0.124 | 0.371 | 826 | 0.352 | 0.659 | 630 | 0.136 | 1.000 | 294 | 0.922 | 0.968 | 96 | 0.121 | 0.371 | 251 | 0.804 |
| BRD-A50049929-001-01-9 | oxazepam | 39 | 0.331 | 1.000 | 502 | 0.397 | 1.000 | 33 | 0.050 | 1.000 | 1129.5 | 0.497 | 0.678 | 500 | 0.971 | 0.971 | 198 | 0.451 | 0.678 | 213 | 0.270 | 0.810 | 305.5 | 0.516 | 0.678 | 206.5 | 0.832 | 0.951 | 321 | 0.360 | 0.678 | 196.5 | 0.441 | 0.678 |  |  |  | 89 | 0.361 | 0.947 | 69 | 0.207 |
| BRD-A50311610-300-10-4 | meclizine | 1055 | 0.380 | 0.873 | 2270 | 0.978 | 0.978 | 716 | 0.129 | 0.675 | 3865 | 0.437 | 0.873 | 1491 | 0.659 | 0.899 | 485 | 0.024 | 0.499 | 1077 | 0.075 | 0.588 | 1330 | 0.715 | 0.899 | 623 | 0.499 | 0.873 | 750 | 0.563 | 0.899 | 1100 | 0.836 | 0.899 | 576 | 0.649 | 0.899 | 176 | 0.348 | 0.873 | 177 | 0.351 |
| BRD-A51571244-001-01-9 | hypoestoxide | 3005 | 0.622 | 0.965 | 5389 | 0.888 | 0.997 | 1909 | 0.278 | 0.850 | 9651 | 0.438 | 0.854 | 3415 | 0.324 | 0.850 | 2760 | 0.257 | 0.850 | 2614 | 0.050 | 0.523 | 4230 | 0.981 | 0.997 | 1148 | 0.735 | 0.965 | 1939 | 0.308 | 0.850 | 1825 | 0.203 | 0.850 | 1600 | 0.958 | 0.997 | 393 | 0.656 | 0.965 | 556 | 0.619 |
| BRD-A51964809-003-12-6 | halofantrine | 449 | 0.808 | 0.943 | 718 | 0.977 | 0.977 | 208 | 0.770 | 0.943 | 1006 | 0.106 | 1.000 | 425 | 0.489 | 0.789 | 410 | 0.421 | 0.736 | 534 | 0.733 | 0.943 | 393 | 0.575 | 0.862 | 179 | 0.931 | 0.977 | 77 | 0.215 | 0.501 | 311 | 0.420 | 0.736 | 179 | 0.931 | 0.977 |  |  |  | 69 | 0.166 |
| BRD-A53134341-001-03-2 | Ro-106-9920 | 4690 | 0.839 | 0.881 | 5970 | 0.211 | 0.627 | 3445 | 0.051 | 0.627 | 13644 | 0.239 | 0.627 | 4942 | 0.654 | 0.808 | 4526 | 0.753 | 0.879 | 6223 | 0.830 | 0.881 | 5504 | 0.654 | 0.808 | 2272 | 0.540 | 0.808 | 5298 | 1.000 | 1.000 | 2193 | 0.156 | 0.627 | 2551 | 0.616 | 0.808 | 319 | 0.078 | 0.627 | 952 | 0.389 |
| BRD-A53951045-004-01-6 | butylscopolamine-bromide | 538 | 0.118 | 1.000 | 1358 | 0.228 | 0.479 | 915.5 | 0.574 | 0.861 | 2866 | 0.876 | 0.956 | 1730.5 | 0.911 | 0.956 | 616.5 | 0.715 | 0.956 | 1396.5 | 0.830 | 0.956 | 1625.5 | 0.983 | 0.983 | 270 | 0.186 | 0.479 | 382 | 0.561 | 0.861 | 480 | 0.144 | 0.431 | 64 | 0.057 | 0.238 | 273 | 0.877 | 0.956 | 54 | 0.009 |
| BRD-A54927599-001-16-9 | KF-38789 | 3518 | 0.841 | 0.939 | 6234 | 0.809 | 0.939 | 3214 | 0.470 | 0.939 | 11264 | 0.149 | 0.784 | 3677 | 0.045 | 0.477 | 3146 | 0.842 | 0.939 | 4764 | 0.739 | 0.939 | 5400 | 0.717 | 0.939 | 2001 | 0.850 | 0.939 | 2743 | 0.513 | 0.939 | 2448 | 0.396 | 0.939 | 1719 | 0.567 | 0.939 | 290 | 0.298 | 0.893 | 457 | 0.093 |
| BRD-A55424491-001-19-9 | methotrexate | 1729 | 0.033 | 0.209 | 4209 | 0.925 | 0.994 | 1871 | 0.040 | 0.209 | 6710 | 0.140 | 0.490 | 3495 | 0.905 | 0.994 | 1168 | 0.540 | 0.961 | 2785 | 0.549 | 0.961 | 3515 | 0.994 | 0.994 | 591 | 0.096 | 0.401 | 2068 | 0.968 | 0.994 | 1032 | 0.004 | 0.085 | 681 | 0.719 | 0.961 | 292 | 0.502 | 0.961 | 533 | 0.984 |
| BRD-A55484088-050-02-5 | BNTX | 1778 | 0.058 | 0.203 | 3271 | 0.009 | 0.198 | 1722 | 0.280 | 0.588 | 8551 | 0.944 | 0.995 | 2934 | 0.441 | 0.772 | 2181 | 0.974 | 0.995 | 3515 | 0.503 | 0.812 | 3520 | 0.995 | 0.995 | 752 | 0.246 | 0.588 | 1815 | 0.889 | 0.995 | 1281 | 0.054 | 0.379 | 547 | 0.729 | 0.986 | 363 | 0.275 | 0.588 | 209 | 0.054 |
| BRD-A55913614-316-09-6 | primaquine | 710 | 0.618 | 0.811 | 1307 | 0.603 | 0.811 | 389 | 0.273 | 1.000 | 2706.5 | 0.889 | 0.917 | 801 | 0.670 | 0.827 | 111 | 0.015 | 0.156 | 1160.5 | 0.917 | 0.917 | 864.5 | 0.814 | 0.917 |  |  |  | 222 | 0.849 | 1.000 | 176 | 0.006 | 0.116 | 52 | 0.054 | 0.125 | 250 | 0.591 | 1.000 | 20 | 0.102 |
| BRD-A55962179-001-22-1 | omeprazole | 501 | 0.758 | 0.995 | 749 | 0.183 | 0.594 | 818 | 0.933 | 1.000 | 1593.5 | 0.099 | 0.490 | 1225.5 | 0.989 | 1.000 | 496 | 0.498 | 0.936 | 480 | 0.117 | 0.490 | 267 | 0.396 | 0.831 | 246 | 0.305 | 0.800 | 279 | 0.198 | 0.594 | 248 | 0.041 | 0.490 | 68 | 0.098 | 0.490 | 225 | 0.535 | 0.936 | 19 | 0.103 |
| BRD-A56085258-001-01-8 | LGX818 | 187.5 | 0.666 | 1.000 | 128.5 | 0.000 | 0.000 | 590 | 0.977 | 1.000 | 1608 | 1.000 | 1.000 | 230.5 | 0.284 | 1.000 | 193.5 | 0.706 | 1.000 | 226.5 | 0.265 | 1.000 | 184.5 | 0.296 | 1.000 | 215 | 0.827 | 1.000 | 165.5 | 0.507 | 1.000 | 344 | 0.980 | 0.980 | 174 | 0.926 | 1.000 | 87.5 | 0.848 | 1.000 |  |  |
| BRD-A57798112-001-03-7 | PD-318088 | 1476 | 0.004 | 0.044 | 2043 | 0.000 | 0.000 | 1511.5 | 0.122 | 0.638 | 7744.5 | 0.993 | 0.999 | 2934 | 0.962 | 0.999 | 2172.5 | 0.982 | 0.999 | 2838 | 0.078 | 0.274 | 2056 | 0.952 | 0.999 | 695.5 | 0.264 | 0.682 | 1299.5 | 0.951 | 0.999 | 1081 | 0.287 | 0.682 | 585 | 0.892 | 0.999 | 200 | 0.778 | 0.999 | 534.5 | 0.463 |
| BRD-A57886255-001-01-1 | golgicide-a | 603 | 0.160 | 1.000 | 1567 | 0.068 | 1.000 | 792 | 0.097 | 1.000 | 3200 | 0.599 | 0.972 | 1376 | 0.653 | 0.972 | 675 | 0.293 | 0.615 | 1124 | 0.687 | 0.972 | 1381 | 0.233 | 0.612 | 408 | 0.279 | 0.615 | 940 | 0.964 | 0.972 | 693 | 0.169 | 0.508 | 389 | 0.089 | 0.310 | 156 | 0.789 | 0.972 | 257 | 0.364 |
| BRD-A58193911-003-18-7 | oxyphencyclimine | 3783 | 0.993 | 0.993 | 4040 | 0.870 | 0.993 | 2298 | 0.025 | 0.200 | 9889.5 | 0.655 | 0.930 | 3184.5 | 0.210 | 1.000 | 2169.5 | 0.444 | 1.000 | 4421.5 | 0.748 | 0.981 | 2864 | 0.548 | 1.000 | 1372 | 0.394 | 1.000 | 1907 | 0.989 | 0.993 | 1404 | 0.251 | 1.000 | 1115 | 0.255 | 0.486 | 504 | 0.903 | 0.993 | 561.5 | 0.664 |
| BRD-A60594020-034-01-1 | 8-hydroxy-PIPAT | 1083 | 0.599 | 0.824 | 1638 | 0.550 | 0.824 | 1557 | 0.417 | 1.000 | 4425 | 0.952 | 0.952 | 1420 | 0.667 | 0.824 | 971 | 0.372 | 0.650 | 1882 | 0.769 | 0.850 | 1520 | 0.359 | 0.650 | 356 | 0.288 | 0.605 | 698 | 0.667 | 0.824 | 567 | 0.153 | 1.000 | 250 | 0.257 | 0.605 | 107 | 0.112 | 0.362 | 111 | 0.121 |
| BRD-A60786380-412-01-5 | sodium-stibogluconate | 469.5 | 0.816 | 0.941 | 1368 | 0.608 | 1.000 | 774.5 | 0.056 | 0.391 | 3308.5 | 0.680 | 1.000 | 916 | 0.110 | 0.580 | 1105 | 0.581 | 1.000 | 722 | 0.147 | 0.617 | 1083 | 0.712 | 0.921 | 704 | 0.673 | 1.000 | 468 | 0.520 | 1.000 | 524.5 | 0.697 | 0.921 | 278 | 0.492 | 1.000 | 403 | 0.904 | 0.941 | 161 | 0.366 |
| BRD-A63546914-003-02-0 | Ro-04-5595 | 614 | 0.694 | 0.909 | 317.5 | 0.811 | 0.909 | 496 | 0.870 | 0.909 | 982 | 0.378 | 0.881 | 496 | 0.870 | 0.909 | 314.5 | 0.083 | 0.436 | 665.5 | 0.909 | 0.909 | 497 | 0.265 | 0.829 | 75.5 | 0.079 | 0.436 | 65.5 | 0.658 | 1.000 | 197.5 | 0.235 | 0.829 | 193.5 | 0.753 | 0.909 | 193.5 | 0.753 | 0.909 | 65.5 | 0.658 |
| BRD-A64064900-001-02-3 | sangivamycin | 336 | 0.195 | 0.743 | 1092 | 0.076 | 0.743 | 744 | 0.255 | 0.755 | 2614 | 0.320 | 0.755 | 710 | 0.349 | 0.755 | 392 | 0.155 | 0.743 | 1020 | 0.360 | 0.755 | 1770 | 0.989 | 0.989 | 115 | 0.212 | 0.743 | 873 | 0.899 | 0.989 | 506 | 0.721 | 0.989 | 565 | 0.863 | 0.989 | 167 | 0.443 | 0.846 | 135 | 0.077 |
| BRD-A64092382-003-18-3 | mexiletine | 3847 | 0.258 | 0.899 | 3928 | 0.002 | 0.040 | 2675 | 0.021 | 0.216 | 12311 | 0.464 | 0.935 | 4528 | 0.503 | 0.935 | 3567 | 0.826 | 0.994 | 6473 | 0.994 | 0.994 | 4412 | 0.090 | 0.376 | 2123 | 0.753 | 0.994 | 2646 | 0.534 | 0.935 | 1973 | 0.060 | 0.423 | 1790 | 0.941 | 0.994 | 980 | 0.985 | 0.994 | 1482 | 0.985 |
| BRD-A64242993-001-05-6 | lanatoside-c | 4836 | 0.980 | 1.000 | 8068 | 1.000 | 1.000 | 2798 | 0.076 | 0.442 | 9330 | 0.000 | 0.001 | 4058 | 0.206 | 0.721 | 4252 | 0.980 | 1.000 | 5392 | 0.978 | 1.000 | 5446 | 0.786 | 0.970 | 1487 | 0.268 | 0.750 | 2812 | 0.701 | 0.970 | 2711 | 0.772 | 0.970 | 1982 | 0.419 | 0.750 | 427 | 0.067 | 0.442 | 635 | 0.105 |
| BRD-A64553394-001-04-4 | DPI-201106 | 2012 | 0.443 | 0.665 | 4861 | 0.572 | 0.751 | 2064 | 0.336 | 0.665 | 8736 | 0.304 | 0.665 | 3283 | 0.288 | 0.665 | 2377 | 0.517 | 0.723 | 4678 | 1.000 | 1.000 | 4640 | 0.930 | 1.000 | 1383 | 0.962 | 1.000 | 2723 | 0.928 | 1.000 | 930 | 0.004 | 0.085 | 592 | 0.381 | 0.665 | 207 | 0.044 | 0.309 | 492 | 0.067 |
| BRD-A66419424-001-02-4 | ixazomib-citrate | 4387 | 0.531 | 0.932 | 6043 | 0.051 | 0.407 | 3525 | 0.199 | 0.696 | 15363 | 0.843 | 0.932 | 4627 | 0.696 | 0.932 | 4609 | 0.947 | 0.947 | 6724 | 0.926 | 0.947 | 4975 | 0.035 | 0.407 | 2116 | 0.586 | 0.932 | 2574 | 0.168 | 0.696 | 2635 | 0.058 | 0.407 | 2603 | 0.826 | 0.932 | 873 | 0.576 | 0.932 | 725 | 0.343 |
| BRD-A67748489-001-10-0 | K-858 | 1383.5 | 0.781 | 0.965 | 1788.5 | 0.593 | 0.887 | 1266 | 0.252 | 0.662 | 4163 | 0.143 | 0.600 | 1781 | 0.299 | 0.697 | 1466.5 | 0.876 | 0.970 | 1987.5 | 0.924 | 0.970 | 2395.5 | 0.908 | 0.970 | 501 | 0.634 | 0.887 | 1533 | 0.979 | 0.979 | 526 | 0.175 | 0.613 | 3 | 0.047 | 0.600 | 223 | 0.473 | 0.825 | 459 | 0.510 |
| BRD-A68083442-001-01-6 | pranidipine | 169.5 | 0.762 | 1.000 | 383.5 | 0.248 | 1.000 | 381.5 | 0.863 | 0.880 | 919.5 | 0.191 | 1.000 | 168.5 | 0.028 | 0.597 | 177.5 | 0.302 | 1.000 | 277.5 | 0.821 | 0.880 | 432 | 0.880 | 0.880 | 57.5 | 0.665 | 1.000 |  |  |  | 224 | 0.794 | 0.880 | 114 | 0.721 | 1.000 | 57.5 | 0.665 | 1.000 | 2 | 0.003 |
| BRD-A68631409-001-05-4 | evodiamine | 2170 | 0.165 | 0.868 | 4834 | 0.983 | 0.999 | 1634 | 0.038 | 0.267 | 6754 | 0.027 | 0.267 | 2041 | 0.011 | 0.233 | 3154 | 0.905 | 0.999 | 3468 | 0.835 | 0.999 | 3590 | 0.720 | 0.999 | 998 | 0.632 | 0.999 | 2380 | 0.999 | 0.999 | 1621 | 0.440 | 0.839 | 926 | 0.267 | 0.560 | 645 | 0.874 | 0.999 | 476 | 0.071 |
| BRD-A68969091-001-12-7 | echinomycin | 7127 | 0.995 | 0.999 | 6300 | 0.014 | 0.065 | 3571 | 0.020 | 0.069 | 19744 | 0.972 | 0.999 | 5509 | 0.209 | 0.628 | 4482 | 0.409 | 0.795 | 9249 | 0.998 | 0.999 | 7836 | 0.853 | 0.999 | 1424 | 0.003 | 0.034 | 5199 | 0.999 | 0.999 | 2611 | 0.016 | 0.065 | 2701 | 0.507 | 0.795 | 761 | 0.289 | 0.734 | 1095 | 0.442 |
| BRD-A70083328-001-23-5 | secnidazole | 339 | 0.360 | 0.851 | 1127 | 0.117 | 0.822 | 1016 | 0.856 | 1.000 | 2214 | 0.621 | 1.000 | 677 | 0.365 | 0.851 | 609 | 0.934 | 1.000 | 306 | 0.244 | 0.851 | 862 | 0.688 | 1.000 | 172 | 0.235 | 0.851 | 381 | 0.526 | 0.949 | 451 | 0.781 | 1.000 | 123 | 0.320 | 0.851 | 232 | 0.896 | 1.000 | 40 | 0.037 |
| BRD-A70649075-008-30-8 | sulconazole | 818 | 0.395 | 0.862 | 1678 | 0.905 | 0.905 | 697 | 0.549 | 0.869 | 3456 | 0.124 | 1.000 | 1536 | 0.743 | 0.869 | 735 | 0.646 | 0.869 | 1356 | 0.590 | 0.869 | 1599 | 0.827 | 0.869 | 712 | 0.818 | 0.869 | 356 | 0.149 | 0.523 | 380 | 0.073 | 1.000 | 84 | 0.395 | 0.862 | 180 | 0.411 | 0.862 | 234 | 0.668 |
| BRD-A72703248-003-04-9 | SKF-96365 | 3269 | 0.880 | 0.969 | 4322 | 0.834 | 0.969 | 3104 | 0.644 | 0.966 | 11668 | 0.829 | 0.969 | 4139 | 0.604 | 0.966 | 2694 | 0.400 | 0.840 | 3504 | 0.223 | 0.587 | 4481 | 0.951 | 0.969 | 1104 | 0.082 | 0.371 | 1820 | 0.085 | 0.371 | 1401 | 0.088 | 0.371 | 1024 | 0.123 | 0.431 | 545 | 0.567 | 0.966 | 834 | 0.969 |
| BRD-A72711497-236-03-8 | lasalocid | 2934 | 0.543 | 0.877 | 5340 | 0.600 | 0.888 | 3157 | 0.451 | 1.000 | 10104 | 0.421 | 1.000 | 3568 | 0.285 | 1.000 | 2734 | 0.506 | 1.000 | 5449 | 0.904 | 0.989 | 3155 | 0.043 | 0.099 | 2332 | 0.968 | 0.989 | 1481 | 0.008 | 0.020 | 1641 | 0.022 | 0.470 | 744 | 0.283 | 0.588 | 599 | 0.308 | 0.588 | 990 | 0.680 |
| BRD-A72716251-001-01-5 | peruvoside | 6424 | 0.999 | 0.999 | 9069 | 0.999 | 0.999 | 4097 | 0.251 | 0.785 | 12266 | 0.002 | 0.045 | 5163 | 0.336 | 0.785 | 5194 | 0.950 | 0.999 | 5695 | 0.551 | 0.827 | 5652 | 0.306 | 0.785 | 2314 | 0.714 | 0.938 | 2372 | 0.060 | 0.422 | 3270 | 0.473 | 0.827 | 2190 | 0.395 | 0.827 | 863 | 0.521 | 0.827 | 1308 | 0.811 |
| BRD-A73427433-240-01-3 | adaprev | 809 | 0.718 | 0.870 | 1635 | 0.755 | 0.870 | 1104 | 0.189 | 1.000 | 4392 | 0.941 | 0.941 | 1233 | 0.078 | 0.328 | 763 | 0.096 | 0.338 | 1263 | 0.787 | 0.870 | 1332 | 0.159 | 0.477 | 408 | 0.475 | 0.870 | 1035 | 0.901 | 0.941 | 463 | 0.345 | 1.000 | 188 | 0.402 | 0.870 | 191 | 0.415 | 0.870 | 203 | 0.470 |
| BRD-A73871920-236-03-6 | nanchangmycin | 5729 | 0.936 | 0.970 | 6517 | 0.312 | 0.656 | 4287 | 0.518 | 0.970 | 16186 | 0.913 | 0.970 | 4632 | 0.065 | 0.344 | 4776 | 0.905 | 0.970 | 6565.5 | 0.970 | 0.970 | 5230 | 0.052 | 0.344 | 2263 | 0.678 | 0.970 | 2348 | 0.057 | 0.344 | 3055 | 0.195 | 0.511 | 2191 | 0.802 | 0.970 | 574 | 0.135 | 0.404 | 686 | 0.090 |
| BRD-A74391928-051-01-3 | ibutilide | 555.5 | 0.935 | 0.937 | 1115 | 0.901 | 0.937 | 532 | 0.117 | 1.000 | 2603.5 | 0.740 | 0.937 | 807 | 0.077 | 1.000 | 307.5 | 0.449 | 0.786 | 920 | 0.355 | 0.786 | 871 | 0.399 | 0.786 | 426.5 | 0.611 | 0.856 | 661 | 0.819 | 0.937 | 509.5 | 0.374 | 0.786 | 165.5 | 0.524 | 0.813 | 86 | 0.542 | 0.813 | 115 | 0.243 |
| BRD-A74914197-001-02-9 | pralatrexate | 1872 | 0.642 | 0.913 | 2938.5 | 0.803 | 0.913 | 1076 | 0.054 | 0.680 | 4661.5 | 0.529 | 1.000 | 1505 | 0.086 | 0.680 | 1444.5 | 0.980 | 0.984 | 2224 | 0.984 | 0.984 | 1771 | 0.793 | 0.913 | 929 | 0.076 | 0.144 | 826 | 0.312 | 0.546 | 1551 | 0.801 | 0.913 | 435.5 | 0.059 | 0.123 | 162 | 0.740 | 0.913 | 213 | 0.031 |
| BRD-A75850590-213-01-0 | lomefloxacin | 696 | 0.470 | 0.822 | 1997 | 0.989 | 0.989 | 638 | 0.164 | 0.822 | 2896 | 0.427 | 0.822 | 852 | 0.446 | 0.822 | 1008 | 0.435 | 0.822 | 822 | 0.777 | 0.893 | 1051 | 0.360 | 0.822 | 255 | 0.816 | 0.893 | 842 | 0.626 | 0.877 | 850 | 0.829 | 0.893 | 437 | 0.225 | 0.822 | 126 | 0.686 | 0.893 | 266 | 0.850 |
| BRD-A75975749-001-01-4 | bafetinib | 2209.5 | 0.151 | 0.710 | 1951 | 0.000 | 0.000 | 1420 | 0.338 | 0.710 | 7492 | 0.967 | 0.999 | 3078 | 0.852 | 0.999 | 587 | 0.251 | 0.710 | 2479 | 0.108 | 0.710 | 2978 | 0.779 | 0.999 | 717 | 0.040 | 0.416 | 2198 | 0.999 | 0.999 | 1635 | 0.817 | 0.999 | 1112 | 0.900 | 0.999 | 285.5 | 0.955 | 0.999 | 560 | 0.472 |
| BRD-A78391468-001-02-8 | prednisolone-hemisuccinate | 533 | 0.634 | 1.000 | 671.5 | 0.282 | 1.000 | 311.5 | 0.439 | 1.000 | 1694.5 | 0.875 | 1.000 | 369.5 | 0.001 | 0.019 | 507 | 0.516 | 1.000 | 766.5 | 0.632 | 1.000 | 541 | 0.668 | 1.000 | 86.5 | 0.744 | 1.000 | 340 | 0.903 | 1.000 | 172 | 0.821 | 1.000 | 353 | 0.674 | 1.000 | 24 | 0.090 | 0.472 | 15 | 0.052 |
| BRD-A79981887-003-16-4 | midodrine | 236.5 | 0.506 | 0.802 | 376 | 0.205 | 1.000 | 382 | 0.223 | 1.000 | 1396.5 | 0.690 | 0.905 | 480 | 0.029 | 0.607 | 556 | 0.955 | 0.972 | 420.5 | 0.358 | 1.000 | 639.5 | 0.972 | 0.972 | 46 | 0.014 | 0.033 | 84 | 0.236 | 0.414 | 100 | 0.347 | 1.000 | 127.5 | 0.564 | 0.802 | 26 | 0.166 | 0.318 | 57 | 0.103 |
| BRD-A81129465-001-03-8 | fenoprofen | 1075 | 0.325 | 0.683 | 2408 | 0.811 | 0.946 | 1577 | 0.610 | 0.915 | 4878 | 0.153 | 0.683 | 2155 | 0.780 | 0.946 | 1047 | 0.281 | 0.683 | 1641 | 0.377 | 0.719 | 2754 | 0.903 | 0.957 | 395 | 0.276 | 0.683 | 594 | 0.101 | 0.683 | 1114 | 0.599 | 0.915 | 457 | 0.187 | 0.683 | 255 | 0.198 | 0.683 | 556 | 0.696 |
| BRD-A81513827-001-03-6 | saxagliptin | 898 | 0.270 | 0.606 | 1966 | 0.892 | 0.982 | 926 | 0.177 | 0.464 | 3917 | 0.289 | 0.606 | 1667 | 0.363 | 0.693 | 664 | 0.978 | 0.982 | 1506 | 0.695 | 0.982 | 2090 | 0.833 | 0.982 | 723 | 0.933 | 0.982 | 933 | 0.908 | 0.982 | 569 | 0.172 | 0.464 | 371 | 0.137 | 0.464 | 287 | 0.811 | 0.982 | 123 | 0.156 |

| broad_id | name | al_nervous_system_PVAL_ADJ | fibroblast_MWU_STAT | fibroblast_PVAL | fibroblast_PVAL_ADJ | bile_duct_MWU_STAT | bile_duct_PVAL | bile_duct_PVAL_ADJ | thyroid_MWU_STAT | thyroid_PVAL | thyroid_PVAL_ADJ | upper_aerodigestive_MWU_STAT | upper_aerodigestive_PVAL | upper_aerodigestive_PVAL_ADJ | uterus_MWU_STAT | uterus_PVAL | uterus_PVAL_ADJ | liver_MWU_STAT | liver_PVAL | liver_PVAL_ADJ | esophagus_MWU_STAT | esophagus_PVAL | esophagus_PVAL_ADJ |
|---|---|---|---|---|---|---|---|---|---|---|---|---|---|---|---|---|---|---|---|---|---|---|---|
| BRD-A00758722-001-04-9 | noretynodrel | 0.080 | | | | 363 | 0.542 | 0.830 | 463 | 0.083 | 1.000 | 1308 | 0.793 | 0.864 | 859 | 0.097 | 1.000 | 1148.5 | 0.729 | 0.858 | 1420 | 0.286 | 1.000 |
| BRD-A03506276-001-01-5 | XL888 | 0.996 | 80 | 0.182 | 0.639 | 2182 | 0.974 | 0.996 | 2480 | 0.781 | 0.996 | 3423 | 0.073 | 0.382 | 4176 | 0.740 | 0.996 | 3166 | 0.458 | 0.875 | 4597 | 0.235 | 0.655 |
| BRD-A03932035-004-04-3 | methscopolamine | 0.940 | | | | | | | 360 | 0.502 | 1.000 | 267.5 | 0.632 | 1.000 | 294 | 0.761 | 1.000 | 496.5 | 0.600 | 1.000 | 324.5 | 0.163 | 0.938 |
| BRD-A05906449-004-01-1 | lappaconite | 0.737 | 267 | 0.825 | 0.965 | 446 | 0.114 | 0.661 | 1298 | 0.655 | 0.965 | 2854 | 0.263 | 0.788 | 1607 | 0.055 | 0.661 | 1449 | 0.399 | 0.839 | 3232 | 0.736 | 0.965 |
| BRD-A06352418-001-25-9 | terfenadine | 0.942 | | | | 126 | 0.306 | 0.712 | 77 | 0.474 | 0.712 | 431 | 0.373 | 0.712 | 219 | 0.407 | 0.712 | 735.5 | 0.823 | 0.942 | 746 | 0.288 | 0.712 |
| BRD-A06352508-001-03-7 | SB-218078 | 0.990 | 26 | 0.085 | 0.384 | 920 | 0.953 | 0.990 | 442 | 0.119 | 0.384 | 2073 | 0.790 | 0.990 | 1615 | 0.837 | 0.990 | 1345 | 0.990 | 0.990 | 1558 | 0.600 | 0.990 |
| BRD-A06426627-001-03-5 | diclazuril | 0.799 | | | | 1163 | 0.993 | 0.993 | 1208 | 0.294 | 0.882 | 2348 | 0.016 | 0.332 | 2376 | 0.233 | 0.882 | 1630 | 0.419 | 0.799 | 3405 | 0.847 | 0.983 |
| BRD-A06627858-236-03-0 | monensin | 0.941 | 120 | 0.236 | 0.662 | 1548 | 0.868 | 0.959 | 1698 | 0.588 | 0.950 | 5424 | 0.992 | 0.992 | 3964 | 0.790 | 0.959 | 2240 | 0.022 | 0.432 | 4465 | 0.843 | 0.959 |
| BRD-A07986123-001-02-8 | galeterone | 0.746 | | | | 729 | 0.393 | 0.750 | 911 | 0.451 | 0.789 | 2129 | 0.333 | 0.750 | 1423 | 0.542 | 0.818 | 969 | 0.022 | 0.118 | 2110 | 0.313 | 0.750 |
| BRD-A08545410-003-07-8 | doxycycline | 0.860 | 63 | 0.181 | 0.860 | 521 | 0.473 | 0.860 | 1051 | 0.740 | 0.860 | 1865 | 0.291 | 0.860 | 1445 | 0.533 | 0.860 | 1049 | 0.503 | 0.860 | 1569 | 0.357 | 0.860 |
| BRD-A08545410-311-03-4 | doxycycline | 0.754 | | | | | | | 813 | 0.766 | 0.947 | 919 | 0.031 | 0.660 | 1058 | 0.203 | 1.000 | 617 | 0.147 | 0.343 | 1410 | 0.425 | 0.754 |
| BRD-A08840375-001-01-3 | CID-5458317 | 0.399 | 42 | 0.120 | 0.399 | 348 | 0.477 | 0.877 | 1077 | 0.133 | 0.399 | 2686 | 0.299 | 0.717 | 2129 | 0.065 | 0.341 | 2693 | 0.958 | 0.989 | 3109 | 0.686 | 0.917 |
| BRD-A10039652-001-01-2 | tipranavir | 0.975 | | | | 105 | 0.165 | 1.000 | 89 | 0.508 | 0.909 | 847 | 0.519 | 0.909 | 813 | 0.946 | 0.975 | 443 | 0.826 | 0.975 | 839 | 0.499 | 0.909 |
| BRD-A10070317-003-28-3 | propranolol | 0.091 | | | | 91 | 0.923 | 0.969 | 414 | 0.969 | 0.969 | 345 | 0.808 | 0.969 | 11 | 0.093 | 1.000 | 351 | 0.960 | 0.969 | 348 | 0.603 | 1.000 |
| BRD-A10977446-001-12-1 | carvedilol | 0.076 | | | | 502.5 | 0.997 | 0.997 | 297.5 | 0.663 | 0.956 | 711.5 | 0.111 | 1.000 | 358 | 0.263 | 1.000 | 288 | 0.105 | 0.245 | 596 | 0.287 | 1.000 |
| BRD-A11815689-001-01-1 | proscillaridin-a | 0.234 | | | | 702 | 0.296 | 0.744 | 678 | 0.260 | 0.744 | 2254 | 0.524 | 0.786 | 2268 | 0.710 | 0.932 | 1656 | 0.358 | 0.744 | 2123 | 0.073 | 0.306 |
| BRD-A12230535-001-06-7 | nutlin-3 | 0.245 | | | | 85.5 | 0.811 | 0.989 | 170 | 0.892 | 0.989 | 433.5 | 0.905 | 0.989 | 229 | 0.510 | 0.933 | 117.5 | 0.518 | 0.933 | 380 | 0.923 | 0.989 |
| BRD-A13133631-001-04-0 | fluorometholone | 0.124 | | | | 735 | 0.611 | 0.917 | 583 | 0.285 | 1.000 | 2245 | 0.377 | 0.792 | 1646 | 0.239 | 1.000 | 1181 | 0.668 | 0.935 | 2690 | 0.849 | 0.990 |
| BRD-A14344385-001-03-2 | IDRA-21 | 0.191 | | | | | | | 17 | 0.087 | 0.917 | 615.5 | 0.259 | 0.591 | 628.5 | 0.901 | 0.901 | 478.5 | 0.765 | 0.901 | 967 | 0.607 | 0.857 |
| BRD-A17883755-001-06-1 | lenalidomide | 0.908 | | | | 118 | 0.747 | 0.980 | 268 | 0.087 | 0.598 | 622 | 0.223 | 0.668 | 657.5 | 0.506 | 0.908 | 66 | 0.070 | 0.598 | 1049.5 | 0.942 | 0.942 |
| BRD-A18696154-003-22-6 | idazoxan | 0.257 | | | | 283.5 | 0.390 | 1.000 | 459 | 0.142 | 0.994 | 723.5 | 0.027 | 0.283 | 899.5 | 0.874 | 0.967 | 411 | 0.217 | 0.506 | 1261.5 | 0.618 | 0.967 |
| BRD-A22707317-001-10-8 | SB-205384 | 0.967 | | | | 289.5 | 0.478 | 0.863 | 760 | 0.984 | 0.987 | 1161 | 0.250 | 1.000 | 1009 | 0.436 | 1.000 | 597.5 | 0.144 | 0.433 | 1107 | 0.458 | 0.863 |
| BRD-A23072235-050-18-5 | pheniramine | 0.059 | | | | 159 | 0.265 | 0.564 | 381 | 0.712 | 0.910 | 1255 | 0.037 | 0.386 | 1089 | 0.927 | 0.974 | 538.5 | 0.526 | 0.910 | 1571 | 0.595 | 0.910 |
| BRD-A25004090-001-08-4 | erastin | 0.216 | | | | 31 | 0.121 | 0.437 | 769 | 0.491 | 0.936 | 2243 | 0.962 | 0.982 | 1441 | 0.766 | 1.000 | 1177 | 0.761 | 1.000 | 2229 | 0.896 | 1.000 |
| BRD-A25569250-001-03-3 | KI-16425 | 0.366 | | | | 8 | 0.070 | 1.000 | 327 | 0.906 | 0.951 | 149 | 0.724 | 0.894 | 357 | 0.310 | 1.000 | 280 | 0.212 | 0.405 | 516 | 0.998 | 0.998 |
| BRD-A25608658-001-01-1 | narasin | 0.163 | 31 | 0.070 | 0.328 | 1608 | 0.851 | 0.993 | 1860 | 0.435 | 0.762 | 4785 | 0.776 | 0.993 | 3394 | 0.241 | 0.555 | 2845 | 0.080 | 0.328 | 5203 | 0.870 | 0.993 |
| BRD-A25687296-300-04-3 | emetine | 0.870 | 354 | 0.775 | 0.999 | 2044 | 0.908 | 0.999 | 1888 | 0.198 | 0.611 | 4207 | 0.260 | 0.611 | 3313 | 0.016 | 0.165 | 3131 | 0.218 | 0.611 | 5032 | 0.519 | 0.870 |
| BRD-A26032986-050-02-1 | zaldaride | 0.125 | | | | | | | 270 | 0.875 | 0.929 | 310 | 0.819 | 0.929 | 115 | 0.235 | 1.000 | 307 | 0.962 | 0.962 | 457 | 0.800 | 0.929 |
| BRD-A27376179-001-01-3 | lestaurtinib | 0.883 | 41 | 0.109 | 0.477 | 1298 | 0.914 | 0.959 | 1606 | 0.612 | 0.959 | 3406 | 0.294 | 0.883 | 3213 | 0.408 | 0.959 | 2565 | 0.796 | 0.959 | 3750 | 0.426 | 0.959 |
| BRD-A27883417-300-01-6 | alexidine | 0.999 | | | | 395 | 0.649 | 0.899 | 969 | 0.221 | 1.000 | 1392 | 0.001 | 0.017 | 2804 | 0.920 | 0.999 | 1656 | 0.495 | 0.899 | 2666 | 0.271 | 0.631 |
| BRD-A28301258-001-03-0 | salinomycin | 0.875 | | | | 690 | 0.983 | 0.983 | 501 | 0.413 | 0.875 | 1618 | 0.256 | 0.875 | 1886 | 0.768 | 1.000 | 833 | 0.025 | 0.460 | 2158 | 0.957 | 0.983 |
| BRD-A28467416-002-01-0 | PKI-179 | 0.947 | 121 | 0.269 | 0.734 | 1685 | 0.639 | 0.947 | 2642 | 0.852 | 0.990 | 4128 | 0.362 | 0.798 | 3426 | 0.029 | 0.138 | 2367 | 0.033 | 0.138 | 5087 | 0.719 | 0.947 |
| BRD-A28970875-300-04-9 | puromycin | 0.326 | | | | 879 | 0.770 | 1.000 | 1475 | 0.568 | 0.852 | 2469 | 0.069 | 0.404 | 2345 | 0.077 | 0.404 | 2580 | 0.993 | 1.000 | 2996 | 0.098 | 0.257 |
| BRD-A30886737-001-02-9 | florfenicol | 0.141 | | | | | | | 296 | 0.632 | 0.896 | 205 | 0.022 | 0.456 | 186 | 0.640 | 0.896 | 112 | 0.047 | 0.141 | 155 | 0.422 | 1.000 |
| BRD-A31095847-001-02-3 | mozavaptan | 0.926 | 89 | 0.294 | 0.882 | 997 | 0.890 | 0.935 | 1053 | 0.758 | 0.926 | 2036 | 0.205 | 0.875 | 1811 | 0.236 | 0.875 | 684 | 0.019 | 0.141 | 1380 | 0.020 | 0.141 |
| BRD-A35588707-001-05-5 | teniposide | 0.906 | 360 | 0.799 | 0.987 | 1397 | 0.572 | 0.987 | 2255 | 0.544 | 0.987 | 4717 | 0.629 | 0.987 | 3480 | 0.037 | 0.278 | 2831 | 0.313 | 0.870 | 5973 | 0.911 | 0.987 |
| BRD-A36057565-001-01-0 | bruceantin | 0.995 | 378 | 0.859 | 0.995 | 1238 | 0.695 | 0.995 | 2807 | 0.989 | 0.995 | 4160 | 0.331 | 0.856 | 3561 | 0.068 | 0.359 | 4448 | 0.995 | 0.995 | 3745 | 0.019 | 0.356 |
| BRD-A36267905-003-18-4 | buphenine | 0.594 | | | | 74.5 | 0.763 | 1.000 | 148 | 0.842 | 1.000 | 422 | 0.535 | 0.936 | 177 | 0.237 | 0.808 | 105.5 | 0.110 | 0.808 | 378 | 0.308 | 0.808 |
| BRD-A36331462-001-02-1 | oprozomib | 0.850 | 14 | 0.038 | 0.264 | 872 | 0.639 | 0.909 | 1601 | 0.550 | 0.894 | 3824 | 0.535 | 0.894 | 3897 | 0.728 | 0.909 | 3441 | 0.989 | 0.989 | 4027 | 0.553 | 0.894 |
| BRD-A38350138-001-04-9 | ketoconazole | 0.612 | | | | 99 | 0.909 | 0.965 | 102 | 0.036 | 0.381 | 112 | 0.174 | 0.612 | 251 | 0.699 | 0.975 | 307 | 0.700 | 0.975 | 349 | 0.980 | 0.980 |
| BRD-A39390670-236-10-7 | rabeprazole | 0.936 | | | | 93 | 0.395 | 0.923 | 7 | 0.070 | 0.747 | 225 | 0.626 | 0.936 | 286 | 0.211 | 0.747 | 108 | 0.532 | 0.936 | 406 | 0.762 | 0.762 |
| BRD-A39935389-001-05-9 | teroxirone | 0.670 | | | | 459 | 0.912 | 0.976 | 1083 | 0.976 | 0.976 | 1875 | 0.410 | 0.956 | 1671 | 0.317 | 0.956 | 1013 | 0.065 | 0.125 | 2635 | 0.811 | 0.968 |
| BRD-A41304429-001-14-8 | practolol | 0.987 | | | | 241 | 0.170 | 0.714 | 239 | 0.166 | 0.714 | 842 | 0.167 | 0.714 | 1484 | 0.515 | 0.760 | 466 | 0.215 | 0.503 | 1541 | 0.607 | 0.760 |
| BRD-A42699921-001-03-6 | teriflunomide | 0.611 | | | | 344.5 | 0.984 | 0.989 | 369 | 0.528 | 0.831 | 489 | 0.070 | 0.611 | 450 | 0.251 | 0.734 | 608 | 0.910 | 0.989 | 956.5 | 0.854 | 1.000 |
| BRD-A42746051-001-02-9 | melengestrol-acetate | 0.700 | | | | 175 | 0.583 | 0.875 | 79 | 0.791 | 0.917 | 513 | 0.303 | 1.000 | 272 | 0.829 | 0.917 | 268 | 0.491 | 0.937 | 348 | 0.654 | 0.915 |
| BRD-A43671941-003-11-9 | oxprenolol | 0.707 | | | | 845.5 | 0.915 | 1.000 | 799 | 0.862 | 1.000 | 1341 | 0.328 | 0.766 | 1050 | 0.405 | 0.819 | 783 | 0.070 | 0.707 | 1801 | 0.819 | 1.000 |
| BRD-A44090213-001-25-6 | indoprofen | 0.051 | | | | 293 | 0.694 | 0.997 | 379 | 0.233 | 0.938 | 1560 | 0.268 | 0.938 | 1288 | 0.324 | 0.971 | 354 | 0.005 | 0.011 | 2307 | 0.996 | 0.997 |
| BRD-A44500574-001-01-8 | FLI-06 | 0.129 | 102 | 0.410 | 0.892 | 659 | 0.996 | 0.996 | 575 | 0.239 | 0.837 | 1093 | 0.824 | 0.978 | 747 | 0.865 | 0.978 | 937 | 0.545 | 0.954 | 2284 | 0.772 | 0.978 |
| BRD-A45498368-001-07-1 | WYE-125132 | 0.988 | | | | 1065 | 0.506 | 0.763 | 2479 | 0.988 | 0.988 | 3339 | 0.590 | 0.825 | 2185 | 0.041 | 0.854 | 961 | 0.008 | 0.020 | 3035 | 0.461 | 1.000 |
| BRD-A48237631-001-04-8 | mitomycin-c | 0.985 | 223 | 0.542 | 0.985 | 1109 | 0.354 | 0.900 | 1713 | 0.821 | 0.985 | 3935 | 0.779 | 0.985 | 3852 | 0.726 | 0.985 | 3251 | 0.938 | 0.985 | 3942 | 0.372 | 0.900 |
| BRD-A48300215-001-04-8 | ajmaline | 0.910 | 35 | 0.201 | 0.582 | 107 | 0.832 | 0.940 | 85 | 0.181 | 0.582 | 439 | 0.222 | 0.582 | 147 | 0.206 | 0.582 | 411 | 0.552 | 0.910 | 298 | 0.148 | 0.582 |
| BRD-A49035384-003-28-9 | methacycline | 0.517 | | | | 500 | 0.985 | 1.000 | 404 | 0.881 | 1.000 | 994 | 0.450 | 0.991 | 390 | 0.003 | 0.070 | 341 | 0.139 | 0.582 | 1226 | 0.740 | 1.000 |
| BRD-A49358627-001-14-6 | ciprofibrate | 0.343 | | | | 207.5 | 0.950 | 0.977 | | | | 462.5 | 0.273 | 0.574 | 239.5 | 0.217 | 0.506 | 534 | 0.977 | 0.977 | 377 | 0.339 | 0.600 |
| BRD-A49765801-001-04-1 | fludroxycortide | 0.938 | | | | 114 | 0.651 | 0.854 | 331 | 0.147 | 1.000 | 957 | 0.464 | 0.740 | 837 | 0.580 | 0.811 | 226 | 0.293 | 0.659 | 918 | 0.377 | 0.659 |
| BRD-A50049929-001-01-9 | oxazepam | 0.724 | | | | 179.5 | 0.958 | 0.971 | 123 | 0.665 | 0.821 | 253 | 0.491 | 0.678 | 260 | 0.282 | 1.000 | 80 | 0.077 | 0.162 | 382.5 | 0.861 | 0.951 |
| BRD-A50311610-300-10-4 | meclizine | 0.873 | 180 | 0.846 | 0.899 | 166 | 0.084 | 0.588 | 545 | 0.304 | 0.873 | 1204 | 0.319 | 0.873 | 849 | 0.777 | 0.899 | 719 | 0.488 | 0.873 | 1536 | 0.856 | 0.899 |
| BRD-A51571244-001-01-9 | hypoestoxide | 0.965 | 292 | 0.867 | 0.997 | 1626 | 0.997 | 0.997 | 934 | 0.400 | 0.854 | 2196 | 0.271 | 0.850 | 2946 | 0.711 | 0.965 | 1756 | 0.448 | 0.854 | 1853 | 0.005 | 0.115 |
| BRD-A51964809-003-12-6 | halofantrine | 0.499 | | | | | | | 137 | 0.707 | 0.943 | 170 | 0.207 | 1.000 | 221 | 0.222 | 1.000 | 178 | 0.245 | 0.514 | 295 | 0.158 | 1.000 |
| BRD-A53134341-001-03-2 | Ro-106-9920 | 0.808 | 271 | 0.652 | 0.808 | 1122 | 0.631 | 0.808 | 1749 | 0.404 | 0.808 | 3880 | 0.199 | 0.627 | 3413 | 0.237 | 0.627 | 2586 | 0.175 | 0.627 | 4105 | 0.461 | 0.808 |
| BRD-A53951045-004-01-6 | butylscopolamine-bromide | 0.045 | | | | 265.5 | 0.457 | 0.800 | 202 | 0.215 | 0.479 | 1029 | 0.455 | 0.800 | 479 | 0.021 | 0.445 | 541 | 0.122 | 0.427 | 999 | 0.754 | 0.956 |
| BRD-A54927599-001-16-9 | KF-38789 | 0.654 | 290 | 0.768 | 0.939 | 1055 | 0.916 | 0.962 | 2476 | 0.968 | 0.968 | 3109 | 0.252 | 0.883 | 2020 | 0.020 | 0.414 | 2586 | 0.848 | 0.939 | 3092 | 0.241 | 0.883 |
| BRD-A55424491-001-19-9 | methotrexate | 0.994 | | | | 428 | 0.192 | 0.576 | 688 | 0.431 | 0.961 | 2152 | 0.732 | 0.961 | 1506 | 0.628 | 0.961 | 1578 | 0.726 | 0.961 | 2258 | 0.259 | 0.681 |
| BRD-A55484088-050-02-5 | BNTX | 0.203 | | | | 527 | 0.332 | 0.633 | 1203 | 0.751 | 0.986 | 1737 | 0.021 | 0.223 | 2200 | 0.686 | 0.986 | 1235 | 0.187 | 0.561 | 3416 | 0.968 | 0.995 |
| BRD-A55913614-316-09-6 | primaquine | 0.214 | | | | 72 | 0.448 | 1.000 | 229 | 0.192 | 1.000 | 657.5 | 0.463 | 1.000 | 210 | 0.044 | 0.306 | 138 | 0.906 | 0.917 | 979 | 0.593 | 1.000 |
| BRD-A55962179-001-22-1 | omeprazole | 0.490 | | | | 234 | 0.583 | 0.942 | 334.5 | 0.701 | 0.982 | 393 | 0.638 | 0.957 | 844 | 0.395 | 0.831 | 320 | 0.915 | 1.000 | 763.5 | 0.992 | 0.992 |
| BRD-A56085258-001-01-8 | LGX818 | | | | | 132.5 | 0.692 | 1.000 | 136.5 | 0.094 | 0.986 | 180.5 | 0.618 | 1.000 | 384.5 | 0.569 | 1.000 | 115.5 | 0.545 | 1.000 | 352.5 | 0.887 | 1.000 |
| BRD-A57798112-001-03-7 | PD-318088 | 0.883 | | | | 350.5 | 0.325 | 0.682 | 295 | 0.056 | 0.391 | 1956 | 0.319 | 0.682 | 1023 | 0.633 | 0.999 | 1233.5 | 0.532 | 0.931 | 3255 | 0.999 | 0.999 |
| BRD-A57886255-001-01-1 | golgicide-a | 0.695 | | | | | | | 363 | 0.749 | 0.972 | 1352 | 0.883 | 0.972 | 1067 | 0.972 | 0.972 | 381 | 0.803 | 0.972 | 1361 | 0.892 | 0.972 |
| BRD-A58193911-003-18-7 | oxyphencyclimine | 0.930 | | | | 659 | 0.512 | 1.000 | 1197 | 0.831 | 0.993 | 1987 | 0.018 | 0.200 | 1423 | 0.029 | 0.200 | 1578.5 | 0.281 | 0.492 | 3380 | 0.616 | 0.930 |
| BRD-A60594020-034-01-1 | 8-hydroxy-PIPAT | 0.362 | | | | 420 | 0.822 | 0.863 | 509 | 0.749 | 0.850 | 1276 | 0.155 | 1.000 | 598 | 0.202 | 1.000 | 738 | 0.288 | 0.605 | 1399 | 0.462 | 0.747 |
| BRD-A60786380-412-01-5 | sodium-stibogluconate | 0.549 | | | | 541 | 0.941 | 0.941 | 111 | 0.008 | 0.174 | 1363 | 0.867 | 0.941 | 639 | 0.746 | 0.921 | 397 | 0.044 | 0.391 | 1066 | 0.679 | 1.000 |
| BRD-A63546914-003-02-0 | Ro-04-5595 | 0.909 | | | | 65.5 | 0.658 | 1.000 | 70.5 | 0.056 | 0.436 | 250 | 0.040 | 0.436 | 202.5 | 0.276 | 0.829 | 265 | 0.368 | 0.881 | 256 | 0.785 | 0.909 |
| BRD-A64064900-001-02-3 | sangivamycin | 0.743 | 144 | 0.806 | 0.989 | 624 | 0.947 | 0.989 | 361 | 0.851 | 0.989 | 735 | 0.123 | 0.743 | 1022 | 0.533 | 0.933 | 336 | 0.777 | 0.989 | 468 | 0.603 | 0.974 |
| BRD-A64092382-003-18-3 | mexiletine | 0.994 | | | | 1171 | 0.966 | 0.994 | 1658 | 0.449 | 0.935 | 3077 | 0.300 | 0.899 | 2747 | 0.460 | 0.935 | 2833 | 0.845 | 0.994 | 4242 | 0.880 | 0.994 |
| BRD-A64242993-001-05-6 | lanatoside-c | 0.442 | 184 | 0.481 | 0.776 | 1016 | 0.603 | 0.904 | 1632 | 0.428 | 0.750 | 2491 | 0.398 | 0.750 | 2080 | 0.103 | 0.442 | 2408 | 0.322 | 0.750 | 3482 | 0.365 | 0.750 |
| BRD-A64553394-001-04-4 | DPI-201106 | 0.309 | 23 | 0.073 | 0.309 | 1008 | 0.817 | 1.000 | 1501 | 0.370 | 0.665 | 2564 | 0.401 | 0.665 | 2297 | 0.429 | 0.665 | 1337 | 0.352 | 0.665 | 1972 | 0.074 | 0.309 |
| BRD-A66419424-001-02-4 | ixazomib-citrate | 0.932 | 204 | 0.492 | 0.932 | 1288 | 0.827 | 0.932 | 1872 | 0.755 | 0.932 | 4063 | 0.709 | 0.932 | 3550 | 0.766 | 0.932 | 2751 | 0.615 | 0.932 | 3995 | 0.104 | 0.548 |
| BRD-A67748489-001-10-0 | K-858 | 0.825 | 20 | 0.078 | 0.600 | 217 | 0.447 | 0.825 | 462 | 0.092 | 0.600 | 1425 | 0.121 | 0.600 | 1028 | 0.761 | 0.965 | 994.5 | 0.483 | 0.825 | 852 | 0.222 | 0.662 |
| BRD-A68083442-001-01-6 | pranidipine | 0.006 | | | | 114 | 0.721 | 1.000 | 57.5 | 0.665 | 1.000 | 277.5 | 0.821 | 0.880 | 175.5 | 0.282 | 1.000 | 277.5 | 0.821 | 0.880 | 375.5 | 0.201 | 1.000 |
| BRD-A68631409-001-05-4 | evodiamine | 0.212 | | | | 626 | 0.847 | 0.999 | 750 | 0.215 | 0.905 | 3415 | 0.945 | 0.999 | 1753 | 0.263 | 0.560 | 1776 | 0.818 | 0.999 | 2135 | 0.241 | 0.560 |
| BRD-A68969091-001-12-7 | echinomycin | 0.795 | 144 | 0.315 | 0.734 | 2123 | 0.936 | 0.999 | 2286 | 0.530 | 0.795 | 2994 | 0.003 | 0.034 | 3112 | 0.006 | 0.040 | 3811 | 0.821 | 0.999 | 5242 | 0.499 | 0.795 |
| BRD-A70083328-001-23-5 | secnidazole | 0.604 | | | | 6 | 0.058 | 0.604 | 372 | 0.490 | 0.949 | 586 | 0.156 | 0.822 | 385 | 0.542 | 0.949 | 272 | 0.718 | 1.000 | 629 | 0.832 | 0.832 |
| BRD-A70649075-008-30-8 | sulconazole | 0.869 | | | | 348 | 0.697 | 0.869 | 476 | 0.220 | 1.000 | 917 | 0.248 | 1.000 | 889 | 0.767 | 0.869 | 532 | 0.354 | 0.862 | 1094 | 0.604 | 0.869 |
| BRD-A72703248-003-04-9 | SKF-96365 | 0.969 | 169 | 0.491 | 0.938 | 1275 | 0.850 | 0.969 | 396 | 0.077 | 0.371 | 1760 | 0.028 | 0.371 | 1747 | 0.386 | 0.840 | 1202 | 0.145 | 0.434 | 3651 | 0.705 | 0.969 |
| BRD-A72711497-236-03-8 | lasalocid | 0.888 | | | | 779 | 0.635 | 0.888 | 1000 | 0.419 | 1.000 | 2791 | 0.719 | 0.888 | 3843 | 0.989 | 0.989 | 2090 | 0.523 | 0.877 | 3649 | 0.842 | 0.982 |
| BRD-A72716251-001-01-5 | peruvoside | 0.946 | 22 | 0.054 | 0.422 | 1506 | 0.780 | 0.946 | 2093 | 0.498 | 0.827 | 3823 | 0.311 | 0.785 | 4338 | 0.677 | 0.938 | 2290 | 0.309 | 0.785 | 4299 | 0.261 | 0.785 |
| BRD-A73427433-240-01-3 | adaprev | 0.870 | | | | 368 | 0.699 | 0.870 | 800 | 0.699 | 0.870 | 988 | 0.077 | 1.000 | 848 | 0.576 | 0.870 | 861 | 0.605 | 0.870 | 1578 | 0.518 | 0.870 |
| BRD-A73871920-236-03-6 | nanchangmycin | 0.362 | 57 | 0.104 | 0.362 | 1500 | 0.786 | 0.970 | 1678 | 0.290 | 0.656 | 4494 | 0.663 | 0.970 | 3917.5 | 0.678 | 0.970 | 2018 | 0.002 | 0.034 | 4912 | 0.881 | 0.970 |
| BRD-A74391928-051-01-3 | ibutilide | 0.729 | | | | 365.5 | 0.937 | 0.937 | 399 | 0.802 | 0.937 | 695 | 0.168 | 1.000 | 347 | 0.316 | 1.000 | 151 | 0.437 | 0.786 | 489.5 | 0.314 | 1.000 |
| BRD-A74914197-001-02-9 | pralatrexate | 0.072 | | | | 48 | 0.153 | 0.680 | 399 | 0.114 | 0.680 | 1098.5 | 0.826 | 0.913 | 607.5 | 0.567 | 1.000 | 967 | 0.813 | 0.913 | 1127 | 0.162 | 0.680 |
| BRD-A75850590-213-01-0 | lomefloxacin | 0.893 | | | | 213 | 0.269 | 0.822 | 338 | 0.415 | 0.822 | 831 | 0.125 | 0.822 | 202 | 0.068 | 0.822 | 336 | 0.174 | 0.822 | 893 | 0.545 | 0.817 |
| BRD-A75975749-001-01-4 | bafetinib | 0.902 | 228 | 0.842 | 0.999 | 580 | 0.239 | 0.710 | 1013 | 0.312 | 0.710 | 2770 | 0.987 | 0.999 | 1585 | 0.233 | 0.710 | 1241 | 0.272 | 0.710 | 2388 | 0.994 | 0.999 |
| BRD-A78391468-001-02-8 | prednisolone-hemisuccinate | 0.361 | | | | 86.5 | 0.744 | 1.000 | 118.5 | 0.384 | 1.000 | 209 | 0.002 | 0.019 | 504 | 0.945 | 1.000 | 172 | 0.328 | 1.000 | 664 | 0.969 | 0.969 |
| BRD-A79981887-003-16-4 | midodrine | 0.217 | | | | 111 | 0.432 | 1.000 | 90 | 0.275 | 1.000 | 396.5 | 0.912 | 0.972 | 363 | 0.573 | 0.802 | 409.5 | 0.936 | 0.972 | 429.5 | 0.835 | 0.972 |
| BRD-A81129465-001-03-8 | fenoprofen | 0.946 | 181 | 0.743 | 0.946 | 674 | 0.911 | 0.957 | 411 | 0.314 | 0.683 | 1047 | 0.472 | 0.827 | 979 | 0.040 | 0.683 | 489 | 0.246 | 0.683 | 2491 | 0.992 | 0.992 |
| BRD-A81513827-001-03-6 | saxagliptin | 0.464 | | | | 393 | 0.982 | 0.982 | 332 | 0.566 | 0.982 | 956 | 0.025 | 0.523 | 1304 | 0.936 | 0.982 | 190 | 0.031 | 0.163 | 593 | 0.092 | 0.970 |

| broad_id | name | colorectal_MWU_STAT | colorectal_PVAL | colorectal_PVAL_ADJ | skin_MWU_STAT | skin_PVAL | skin_PVAL_ADJ | urinary_tract_MWU_STAT | urinary_tract_PVAL | urinary_tract_PVAL_ADJ | lung_MWU_STAT | lung_PVAL | lung_PVAL_ADJ | ovary_MWU_STAT | ovary_PVAL | ovary_PVAL_ADJ | breast_MWU_STAT | breast_PVAL | breast_PVAL_ADJ | pancreas_MWU_STAT | pancreas_PVAL | pancreas_PVAL_ADJ | nervous_system_MWU_STAT | central_nervous_system_PVAL | nervous_system_PVAL_ADJ | soft_tissue_MWU_STAT | soft_tissue_PVAL | soft_tissue_PVAL_ADJ | kidney_MWU_STAT | kidney_PVAL | kidney_PVAL_ADJ | gastric_MWU_STAT | gastric_PVAL | gastric_PVAL_ADJ | bone_MWU_STAT | bone_PVAL | bone_PVAL_ADJ | prostate_MWU_STAT | prostate_PVAL | prostate_PVAL_ADJ | al_nervous_system_MWU_STAT | peripheral_nervous_system_PVAL |
|---|---|---|---|---|---|---|---|---|---|---|---|---|---|---|---|---|---|---|---|---|---|---|---|---|---|---|---|---|---|---|---|---|---|---|---|---|---|---|---|---|---|---|
| BRD-A83650191-004-02-8 | A61603 | 1860 | 0.228 | 0.704 | 3719.5 | 0.801 | 0.914 | 2508 | 0.898 | 0.914 | 6917 | 0.304 | 0.720 | 3278.5 | 0.763 | 0.914 | 1817.5 | 0.822 | 0.914 | 3391 | 0.914 | 0.914 | 2824 | 0.580 | 0.870 | 254 | 0.106 | 0.597 | 1073 | 0.377 | 0.720 | 1079 | 0.096 | 0.597 | 656.5 | 0.155 | 0.651 | 248 | 0.356 | 0.720 | 558.5 | 0.446 |
| BRD-A92630576-050-24-1 | trimebutine | 476 | 0.586 | 0.843 | 1005 | 0.360 | 0.688 | 607 | 0.178 | 0.936 | 2009 | 0.240 | 1.000 | 452 | 0.052 | 0.936 | 545 | 0.952 | 0.966 | 1145 | 0.966 | 0.966 | 1292 | 0.914 | 0.966 | 171 | 0.602 | 0.843 | 426 | 0.680 | 0.892 | 265 | 0.325 | 0.688 | 411 | 0.881 | 0.966 | 24 | 0.118 | 0.309 | 1 | 0.045 |
| BRD-A92872453-001-01-8 | N-acetylmannosamine | 1106.5 | 0.756 | 0.997 | 2156.5 | 0.997 | 0.997 | 1228.5 | 0.494 | 0.875 | 2196 | 0.058 | 0.535 | 1368 | 0.319 | 0.743 | 1162.5 | 0.697 | 0.997 | 1049.5 | 0.300 | 0.743 | 814 | 0.500 | 0.875 | 393.5 | 0.885 | 0.997 | 346.5 | 0.980 | 0.997 | 784 | 0.843 | 0.997 | 197.5 | 0.191 | 0.743 | 116 | 0.178 | 0.743 | 219 | 0.259 |
| BRD-A93236127-001-04-5 | digitoxin | 3378 | 0.994 | 0.997 | 5448 | 0.997 | 0.997 | 2174 | 0.117 | 0.822 | 6233 | 0.003 | 0.066 | 3581 | 0.770 | 0.997 | 2385 | 0.995 | 0.997 | 3796 | 0.897 | 0.997 | 2104 | 0.084 | 0.173 | 732 | 0.216 | 0.348 | 1915 | 0.678 | 0.997 | 2286 | 0.759 | 0.997 | 386 | 0.110 | 0.192 | 247 | 0.091 | 0.173 | 78 | 0.034 |
| BRD-A93477898-001-04-2 | PETCM | 185 | 0.711 | 0.996 | 661.5 | 0.687 | 1.000 | 434 | 0.471 | 1.000 | 958.5 | 0.391 | 1.000 | 271 | 0.335 | 1.000 | 235.5 | 0.389 | 1.000 | 289 | 0.427 | 1.000 | 556 | 0.778 | 0.860 | 67 | 0.681 | 1.000 | 121 | 0.082 | 0.577 | 75 | 0.064 | 0.577 | | | | 111 | 0.559 | 1.000 | 99.5 | 0.936 |
| BRD-A94756469-001-04-7 | digoxin | 5973 | 0.988 | 0.998 | 9138 | 0.998 | 0.998 | 4464 | 0.628 | 0.879 | 11998 | 0.003 | 0.056 | 5430 | 0.283 | 0.850 | 4898 | 0.971 | 0.998 | 6397 | 0.892 | 0.998 | 5140 | 0.061 | 0.345 | 2048 | 0.142 | 0.497 | 3204 | 0.584 | 0.879 | 3550 | 0.921 | 0.998 | 2375 | 0.575 | 0.879 | 679 | 0.571 | 0.879 | 622 | 0.057 |
| BRD-A95121829-001-08-9 | ginkgolide-a | 552 | 0.728 | 0.955 | 851.5 | 0.466 | 1.000 | 551.5 | 0.300 | 1.000 | 1826.5 | 0.197 | 1.000 | 868.5 | 0.512 | 1.000 | 219 | 0.528 | 1.000 | 426.5 | 0.528 | 1.000 | 754.5 | 0.720 | 0.955 | 384 | 0.886 | 0.970 | 321 | 0.369 | 1.000 | 561 | 0.533 | 1.000 | 18 | 0.100 | 0.161 | 115 | 0.847 | 0.970 | 26 | 0.024 |
| BRD-A97739905-001-26-5 | ketoprofen | 422 | 0.903 | 0.927 | 718 | 0.444 | 1.000 | 450 | 0.828 | 0.927 | 1100 | 0.590 | 0.825 | 532 | 0.391 | 1.000 | 29 | 0.208 | 1.000 | 308 | 0.436 | 1.000 | 293 | 0.629 | 0.825 | 246 | 0.927 | 0.927 | 154 | 0.426 | 0.812 | 130 | 0.269 | 1.000 | 137 | 0.718 | 0.887 | 167 | 0.891 | 0.927 | 130 | 0.269 |
| BRD-A98299281-050-09-6 | hydroxytacrine-maleate-(r,s) | 453 | 0.088 | 0.928 | 421 | 0.000 | 0.000 | 620 | 0.854 | 0.957 | 1454 | 0.918 | 0.957 | 206 | 0.662 | 0.957 | 167 | 0.842 | 0.957 | 802 | 0.878 | 0.957 | 617 | 0.656 | 0.957 | | | | 117 | 0.440 | 0.957 | 96 | 0.263 | 0.689 | 91 | 0.226 | 0.793 | | | | 70 | 0.108 |
| BRD-K00259736-001-16-4 | colchicine | 4160 | 0.884 | 0.928 | 5071 | 0.839 | 0.928 | 2670 | 0.087 | 0.648 | 11625 | 0.578 | 0.928 | 4087 | 0.277 | 0.648 | 2222 | 0.278 | 0.648 | 4154 | 0.794 | 0.928 | 4749 | 0.754 | 0.928 | 1453 | 0.167 | 0.648 | 1661 | 0.787 | 0.928 | 2141 | 0.354 | 0.675 | 1259 | 0.139 | 0.648 | 427 | 0.275 | 0.648 | 392 | 0.060 |
| BRD-K00317371-001-06-1 | RITA | 2130 | 0.020 | 0.413 | 3813 | 0.929 | 0.989 | 2105 | 0.931 | 0.989 | 8463 | 0.455 | 0.868 | 3124 | 0.416 | 0.868 | 1791 | 0.176 | 0.412 | 4144 | 0.940 | 0.989 | 3423 | 0.947 | 0.989 | 1652 | 0.955 | 0.989 | 1303 | 0.003 | 0.016 | 1337 | 0.062 | 0.217 | 1095 | 0.151 | 0.397 | 548 | 0.700 | 0.989 | 492 | 0.566 |
| BRD-K00494077-066-01-9 | trovafloxacin | 656 | 0.561 | 0.751 | 1006 | 0.022 | 0.463 | 763.5 | 0.444 | 1.000 | 2086 | 0.552 | 0.751 | 1031 | 0.572 | 0.751 | 677.5 | 0.408 | 1.000 | 947.5 | 0.719 | 0.839 | 1091 | 0.393 | 0.988 | 307 | 0.423 | 0.988 | 749 | 0.999 | 0.999 | 240 | 0.061 | 0.643 | 87 | 0.537 | 0.751 | 139 | 0.879 | 0.923 | | |
| BRD-K00535541-001-06-3 | 9-aminoacridine | 2080 | 0.231 | 0.692 | 4180 | 0.934 | 0.981 | 2538 | 0.720 | 0.969 | 7368 | 0.137 | 0.576 | 2559 | 0.125 | 0.576 | 2802 | 0.989 | 0.989 | 2792 | 0.550 | 0.888 | 3545 | 0.840 | 0.969 | 843 | 0.188 | 0.656 | 2160 | 0.780 | 0.969 | 1514 | 0.194 | 1.000 | 815 | 0.341 | 0.718 | 640 | 0.877 | 0.969 | 245 | 0.323 |
| BRD-K00634421-001-01-0 | brivaracetam | 3579 | 0.981 | 0.999 | 4917 | 0.714 | 0.999 | 2387 | 0.014 | 0.096 | 10297 | 0.108 | 0.453 | 4709 | 0.924 | 0.999 | 2997 | 0.594 | 0.999 | 4168 | 0.842 | 0.999 | 5334 | 0.953 | 0.999 | 1363 | 0.250 | 0.749 | 2244 | 0.999 | 0.999 | 1929 | 0.104 | 0.453 | 1851 | 0.820 | 0.999 | 602 | 0.645 | 0.999 | 451 | 0.334 |
| BRD-K01507359-001-22-9 | rifampin | 876 | 0.954 | 0.995 | 1756 | 0.033 | 0.301 | 1256 | 0.409 | 0.858 | 4439 | 0.939 | 1.000 | 986 | 0.014 | 0.301 | 914 | 0.290 | 0.858 | 1858 | 0.774 | 1.000 | 1277 | 0.292 | 0.858 | 209 | 0.043 | 0.301 | 950 | 0.561 | 0.982 | 983 | 0.946 | 1.000 | 114 | 0.552 | 0.982 | 182 | 0.367 | 0.858 | 346 | 0.611 |
| BRD-K01614093-001-02-6 | dihydromyricetin | 621 | 0.612 | 0.857 | 843 | 0.121 | 0.548 | 662 | 0.496 | 1.000 | 2282 | 0.135 | 0.548 | 860 | 0.604 | 1.000 | 1038 | 0.978 | 0.978 | 1154 | 0.946 | 0.978 | 1134 | 0.532 | 1.000 | 191 | 0.225 | 0.675 | 524 | 0.157 | 0.548 | 357 | 0.112 | 0.548 | 466 | 0.899 | 0.978 | 182 | 0.559 | 0.839 | 2 | 0.046 |
| BRD-K01669786-001-02-2 | triapine | 2358 | 0.992 | 0.992 | 4095.5 | 0.661 | 1.000 | 1571 | 0.214 | 0.747 | 6608 | 0.166 | 0.697 | 1480 | 0.000 | 0.007 | 1881.5 | 0.767 | 1.000 | 2937 | 0.965 | 1.000 | 2722 | 0.111 | 0.697 | 496 | 0.321 | 0.790 | 1096 | 0.008 | 0.087 | 2056 | 0.800 | 1.000 | 861 | 0.273 | 0.790 | 376 | 0.770 | 1.000 | 504 | 0.339 |
| BRD-K01683783-001-02-6 | TAK-901 | 4941 | 0.382 | 0.729 | 8956 | 0.972 | 1.000 | 3844 | 0.186 | 0.487 | 16143 | 0.543 | 0.950 | 5448 | 0.251 | 0.528 | 4157 | 0.781 | 1.000 | 7092 | 0.891 | 1.000 | 6678 | 0.732 | 1.000 | 1172 | 0.002 | 0.051 | 3810 | 1.000 | 1.000 | 2097 | 0.014 | 0.146 | 1601 | 0.091 | 0.384 | 1408 | 0.988 | 1.000 | 335 | 0.079 |
| BRD-K01773560-001-02-4 | UNC2250 | 856.5 | 0.874 | 0.966 | 1300.5 | 0.848 | 0.966 | 335 | 0.035 | 0.294 | 2014.5 | 0.483 | 0.845 | 923.5 | 0.211 | 0.740 | 697 | 0.976 | 0.976 | 753 | 0.837 | 1.000 | 869.5 | 0.378 | 0.845 | 541 | 0.462 | 0.845 | 252.5 | 0.550 | 0.854 | 422.5 | 0.139 | 0.700 | 165.5 | 0.167 | 0.700 | 75 | 0.444 | 0.845 | 90 | 0.028 |
| BRD-K02113016-001-15-4 | olaparib | 263.5 | 0.333 | 0.635 | 741.5 | 0.946 | 0.976 | 353.5 | 0.271 | 1.000 | 1049 | 0.315 | 1.000 | 645 | 0.264 | 1.000 | 191.5 | 0.245 | 1.000 | 416 | 0.770 | 0.976 | 529.5 | 0.549 | 0.886 | 134 | 0.218 | 0.510 | 377.5 | 0.976 | 0.976 | 92 | 0.017 | 0.355 | 122.5 | 0.012 | 0.032 | 41 | 0.285 | 0.598 | 242 | 0.844 |
| BRD-K02130563-001-11-4 | panobinostat | 4379 | 0.687 | 0.872 | 5896 | 0.180 | 0.539 | 3720 | 0.328 | 0.689 | 13718 | 0.643 | 0.872 | 4820 | 0.381 | 0.690 | 4278 | 0.966 | 0.994 | 5237 | 0.994 | 0.994 | 5776 | 0.913 | 0.994 | 1683 | 0.241 | 0.634 | 3583 | 0.984 | 0.994 | 2063 | 0.144 | 0.504 | 2114 | 0.692 | 0.872 | 342 | 0.025 | 0.174 | 440 | 0.016 |
| BRD-K02241333-001-02-7 | CHIR-124 | 2674 | 0.277 | 0.582 | 4198 | 0.574 | 0.861 | 2555 | 0.189 | 0.499 | 9836 | 0.877 | 0.998 | 2650 | 0.026 | 0.439 | 2894 | 0.951 | 0.998 | 3986 | 0.520 | 0.861 | 3499 | 0.105 | 0.439 | 1334 | 0.730 | 0.959 | 2413 | 0.190 | 0.499 | 1265 | 0.100 | 0.439 | 1110 | 0.407 | 0.778 | 737 | 0.905 | 0.998 | 537 | 0.085 |
| BRD-K02407574-001-09-7 | parbendazole | 3973 | 0.034 | 0.239 | 7698 | 0.913 | 0.999 | 3367 | 0.023 | 0.239 | 15260 | 0.376 | 0.719 | 4757 | 0.133 | 0.466 | 5191 | 0.992 | 0.999 | 7288 | 0.988 | 0.999 | 6485 | 0.763 | 0.999 | 1957 | 0.194 | 0.499 | 4810 | 0.999 | 0.999 | 3091 | 0.647 | 0.970 | 2175 | 0.371 | 0.719 | 829 | 0.462 | 0.809 | 790 | 0.166 |
| BRD-K02594908-001-16-4 | sulfanilamide | 504 | 0.754 | 0.930 | 1536 | 0.585 | 0.930 | 676 | 0.740 | 0.930 | 2076.5 | 0.142 | 0.995 | 961 | 0.290 | 1.000 | 750 | 0.886 | 0.930 | 1350.5 | 0.882 | 0.930 | 1468.5 | 0.990 | 0.990 | 271 | 0.555 | 0.930 | 911.5 | 0.859 | 0.930 | 511 | 0.505 | 0.930 | 71 | 0.376 | 0.878 | 167 | 0.464 | 0.930 | 131 | 0.807 |
| BRD-K02764365-001-13-6 | methiazole | 2118 | 0.393 | 0.770 | 3878 | 0.978 | 0.985 | 1554.5 | 0.050 | 0.353 | 6522 | 0.440 | 0.770 | 3012 | 0.544 | 0.817 | 2379 | 0.920 | 0.985 | 2427.5 | 0.606 | 0.848 | 3864.5 | 0.949 | 0.985 | 948 | 0.412 | 0.770 | 1902.5 | 0.903 | 0.985 | 1401 | 0.487 | 0.787 | 617 | 0.297 | 0.770 | 576 | 0.837 | 0.985 | 410 | 0.162 |
| BRD-K02898090-001-02-7 | kifunensine | 268 | 0.792 | 1.000 | 863.5 | 0.048 | 0.338 | 798 | 0.696 | 1.000 | 1578.5 | 0.499 | 1.000 | 872.5 | 0.725 | 1.000 | 707 | 0.985 | 1.000 | 670.5 | 0.996 | 1.000 | 618.5 | 0.194 | 0.876 | 231.5 | 0.276 | 0.966 | 256 | 0.000 | 0.010 | 214 | 0.209 | 0.876 | 3 | 0.048 | 0.338 | 119.5 | 0.883 | 1.000 | 153.5 | 0.583 |
| BRD-K03063480-001-06-6 | PF-477736 | 4570 | 0.748 | 0.924 | 6499.5 | 0.616 | 0.924 | 4387 | 0.854 | 0.924 | 13572 | 0.684 | 0.924 | 4323 | 0.148 | 0.444 | 4431.5 | 0.873 | 0.924 | 6350 | 0.880 | 0.924 | 4755 | 0.277 | 0.727 | 1505 | 0.100 | 0.357 | 2614 | 0.740 | 0.924 | 2310 | 0.084 | 0.357 | 1340.5 | 0.005 | 0.057 | 819 | 0.858 | 0.924 | 669 | 0.102 |
| BRD-K03109492-001-02-2 | NSC-663284 | 2050 | 0.302 | 0.752 | 3861 | 0.100 | 0.468 | 2012 | 0.156 | 0.468 | 8308.5 | 0.668 | 0.949 | 3224 | 0.534 | 0.949 | 1643 | 0.683 | 0.949 | 2474 | 0.322 | 0.752 | 3892 | 0.914 | 0.949 | 682 | 0.141 | 0.468 | 1974 | 0.861 | 0.949 | 1302 | 0.115 | 0.468 | 887 | 0.445 | 0.934 | 546 | 0.703 | 0.949 | 277 | 0.116 |
| BRD-K03243820-001-23-8 | trifluridine | 3060 | 0.003 | 0.053 | 7332 | 0.804 | 0.989 | 3415 | 0.403 | 0.886 | 11873.5 | 0.036 | 0.253 | 5253 | 0.231 | 0.795 | 4636 | 0.987 | 0.989 | 5256 | 0.548 | 0.886 | 6936 | 0.912 | 0.989 | 912 | 0.005 | 0.058 | 3063 | 0.871 | 0.989 | 2346 | 0.161 | 0.676 | 1253 | 0.057 | 0.302 | 817 | 0.506 | 0.886 | 1006 | 0.487 |
| BRD-K03273112-001-01-8 | 1-azakenpaullone | 759 | 0.815 | 1.000 | 1194 | 0.453 | 0.733 | 750 | 0.395 | 0.733 | 2431 | 0.442 | 0.733 | 1104 | 0.405 | 0.733 | 532 | 0.880 | 1.000 | 995 | 0.917 | 1.000 | 795 | 0.330 | 0.733 | 278 | 0.302 | 0.733 | 417 | 0.826 | 1.000 | 351 | 0.124 | 0.733 | 157 | 0.450 | 0.733 | 208 | 0.734 | 1.000 | 8 | 0.010 |
| BRD-K03289018-001-02-7 | CCT137690 | 3799 | 0.869 | 0.960 | 6295 | 0.952 | 0.998 | 1848 | 0.001 | 0.013 | 10509 | 0.311 | 0.906 | 3904 | 0.345 | 0.906 | 4363.5 | 0.998 | 0.998 | 3680 | 0.549 | 0.911 | 4307 | 0.658 | 0.921 | 1145 | 0.071 | 0.495 | 2750.5 | 0.770 | 0.960 | 1522 | 0.107 | 0.560 | 574 | 0.002 | 0.020 | 573 | 0.563 | 0.911 | 662 | 0.154 |
| BRD-K03390685-001-01-7 | cobimetinib | 1318 | 0.002 | 0.026 | 1319 | 0.000 | 0.000 | 1659 | 0.150 | 0.526 | 8037 | 1.000 | 1.000 | 2217 | 0.897 | 1.000 | 2407 | 0.992 | 1.000 | 2478 | 0.267 | 0.700 | 1646 | 0.753 | 1.000 | 1006 | 0.836 | 1.000 | 1182 | 0.677 | 1.000 | 1189 | 0.142 | 0.526 | 1402 | 0.999 | 1.000 | 464 | 0.952 | 1.000 | 194 | 0.235 |
| BRD-K03406345-001-27-8 | azacitidine | 1308 | 0.390 | 0.898 | 1728 | 0.452 | 0.898 | 1419 | 0.751 | 0.898 | 4695 | 0.804 | 0.898 | 1625 | 0.439 | 0.898 | 810 | 0.097 | 0.693 | 1867 | 0.663 | 0.898 | 1342 | 0.293 | 0.898 | 973 | 0.898 | 0.898 | 707 | 0.868 | 0.898 | 518 | 0.009 | 0.192 | 313 | 0.842 | 0.898 | 235 | 0.559 | 0.898 | 134 | 0.168 |
| BRD-K03449891-001-08-6 | foretinib | 1305.5 | 0.969 | 0.984 | 2753.5 | 0.915 | 0.984 | 1115 | 0.028 | 0.233 | 4568.5 | 0.232 | 0.646 | 1868 | 0.426 | 0.812 | 825.5 | 0.751 | 0.984 | 2683.5 | 0.874 | 0.984 | 1382 | 0.107 | 0.375 | 507 | 0.041 | 0.233 | 879 | 0.378 | 0.795 | 580 | 0.090 | 0.375 | 930 | 0.277 | 0.646 | 452 | 0.984 | 0.984 | 350 | 0.866 |
| BRD-K03549949-001-01-0 | XL388 | 2638 | 0.953 | 1.000 | 3791 | 0.049 | 0.341 | 1771 | 0.463 | 0.850 | 9130 | 0.898 | 1.000 | 2032 | 0.011 | 0.241 | 1856 | 0.392 | 0.850 | 3482 | 0.763 | 1.000 | 3880 | 0.751 | 1.000 | 642 | 0.107 | 0.561 | 1091 | 0.027 | 0.289 | 1511 | 0.173 | 0.606 | 329 | 0.204 | 0.611 | 399 | 0.349 | 0.850 | 298 | 0.464 |
| BRD-K03601870-001-01-2 | ixabepilone | 6298 | 0.910 | 0.998 | 9483 | 0.990 | 0.998 | 3705 | 0.054 | 0.375 | 14148 | 0.017 | 0.356 | 4959 | 0.091 | 0.433 | 5770 | 0.997 | 0.998 | 7250 | 0.886 | 0.998 | 6783 | 0.599 | 0.898 | 2042 | 0.103 | 0.433 | 4905 | 0.998 | 0.998 | 3033 | 0.514 | 0.898 | 2337 | 0.277 | 0.668 | 777 | 0.347 | 0.668 | 986 | 0.350 |
| BRD-K03765900-001-01-9 | XL-647 | 1084 | 0.924 | 1.000 | 2853 | 1.000 | 1.000 | 1054 | 0.017 | 0.116 | 5304 | 0.906 | 1.000 | 1478 | 0.030 | 0.157 | 1207 | 0.391 | 1.000 | 1638 | 0.162 | 0.682 | 2178 | 1.000 | 1.000 | 682 | 0.719 | 1.000 | 1396 | 0.862 | 1.000 | 1281 | 0.519 | 0.927 | 767 | 0.982 | 1.000 | | | | 810 | 0.992 |
| BRD-K04111260-001-10-0 | raclopride | 70 | 0.036 | 0.378 | 1001 | 0.538 | 1.000 | 337.5 | 0.439 | 1.000 | 729.5 | 0.002 | 0.036 | 491 | 0.627 | 1.000 | 185 | 0.536 | 1.000 | 355 | 0.290 | 1.000 | 951.5 | 0.993 | 1.000 | 73 | 0.638 | 1.000 | 202 | 0.306 | 1.000 | 203.5 | 0.653 | 0.915 | 111.5 | 0.428 | 1.000 | 100 | 0.868 | 1.000 | 33 | 0.219 |
| BRD-K04394237-001-03-7 | drospirenone | 636 | 0.150 | 0.450 | 1346 | 0.003 | 0.072 | 1044 | 0.389 | 0.908 | 4517.5 | 0.948 | 0.987 | 1067 | 0.018 | 0.075 | 628 | 0.536 | 0.987 | 1655.5 | 0.895 | 0.987 | 850 | 0.103 | 0.360 | 709 | 0.739 | 0.987 | 839 | 0.936 | 0.987 | 625 | 0.008 | 0.084 | 265.5 | 0.746 | 0.987 | 268 | 0.339 | 0.891 | 270 | 0.763 |
| BRD-K04548931-003-16-5 | epirubicin | 2938 | 0.869 | 1.000 | 3564 | 0.206 | 0.529 | 2135 | 0.219 | 0.529 | 6889 | 0.211 | 0.529 | 2926 | 0.369 | 0.704 | 1628 | 0.555 | 0.896 | 4193 | 0.994 | 1.000 | 3319 | 0.962 | 1.000 | 541 | 0.053 | 0.445 | 2507 | 0.996 | 1.000 | 1051 | 0.004 | 0.082 | 587 | 0.490 | 0.857 | 528 | 0.701 | 0.981 | 154 | 0.136 |
| BRD-K04568635-300-01-4 | octenidine | 659 | 0.157 | 0.549 | 1242 | 0.493 | 0.863 | 903 | 0.859 | 0.913 | 3219 | 0.586 | 0.913 | 923 | 0.344 | 0.801 | 1196 | 0.869 | 0.913 | 1293 | 0.988 | 0.988 | 1483 | 0.775 | 0.913 | 138 | 0.013 | 0.094 | 1140 | 0.791 | 0.913 | 426 | 0.021 | 0.449 | 913 | 0.710 | 0.913 | 87 | 0.108 | 0.454 | 131 | 0.688 |
| BRD-K04623885-001-07-5 | BIBR-1532 | 825 | 0.623 | 0.959 | 2318 | 0.172 | 0.603 | 1934 | 0.882 | 0.959 | 5969 | 0.762 | 0.959 | 2096 | 0.030 | 0.317 | 607 | 0.003 | 0.058 | 2567 | 0.792 | 0.959 | 2767 | 0.646 | 0.959 | 767 | 0.499 | 0.959 | 1669 | 0.959 | 0.959 | 1517 | 0.955 | 0.959 | 930 | 0.812 | 0.959 | 377 | 0.466 | 0.959 | 368 | 0.845 |
| BRD-K04701033-001-03-9 | VE-822 | 4484 | 0.072 | 0.304 | 9230 | 0.967 | 0.997 | 3997 | 0.091 | 0.319 | 18309 | 0.662 | 0.984 | 4983 | 0.035 | 0.197 | 5740 | 0.973 | 0.997 | 8382 | 0.984 | 0.997 | 9076 | 0.997 | 0.997 | 2507 | 0.737 | 0.984 | 3501 | 0.317 | 0.666 | 2003 | 0.001 | 0.011 | 1506 | 0.038 | 0.197 | 1257 | 0.902 | 0.997 | 852 | 0.174 |
| BRD-K04833372-001-06-2 | GSK1904529A | 615 | 0.381 | 0.885 | 851.5 | 0.028 | 0.289 | 211.5 | 0.041 | 0.289 | 2368 | 0.975 | 0.975 | 890 | 0.908 | 0.965 | 589 | 0.755 | 0.965 | 560 | 0.107 | 0.331 | 969 | 0.913 | 0.965 | 27 | 0.121 | 0.331 | 394 | 0.594 | 0.965 | 422 | 0.199 | 1.000 | 394.5 | 0.874 | 0.965 | 116.5 | 0.833 | 0.965 | 28 | 0.126 |
| BRD-K04923131-001-15-4 | GSK-3-inhibitor-IX | 565 | 0.879 | 1.000 | 1067 | 0.666 | 1.000 | 506 | 0.159 | 0.476 | 1747 | 0.945 | 1.000 | 416 | 0.039 | 0.302 | 333 | 0.283 | 0.594 | 739 | 0.985 | 1.000 | 270 | 0.013 | 0.274 | 0 | 0.043 | 0.302 | 291 | 0.153 | 0.476 | 386 | 0.270 | 0.594 | 160 | 0.093 | 0.476 | 71 | 0.546 | 0.882 | 93 | 0.235 |
| BRD-K04993719-001-01-7 | LY2334737 | 2897 | 0.599 | 0.899 | 3872 | 0.507 | 0.819 | 2877 | 0.428 | 0.808 | 9791 | 0.462 | 0.808 | 3366 | 0.146 | 0.614 | 3706 | 0.966 | 0.966 | 4817 | 0.953 | 0.966 | 4286 | 0.814 | 0.943 | 826 | 0.010 | 0.104 | 2650 | 0.813 | 0.943 | 1173 | 0.123 | 0.614 | 1285 | 0.408 | 0.808 | 596 | 0.336 | 0.808 | 220 | 0.008 |
| BRD-K05104363-001-10-0 | PD-184352 | 739.5 | 0.027 | 0.236 | 983 | 0.000 | 0.000 | 1013.5 | 0.686 | 0.997 | 2817.5 | 0.922 | 0.997 | 931 | 0.708 | 0.997 | 948 | 0.975 | 0.997 | 1829.5 | 0.488 | 0.997 | 1277.5 | 0.925 | 0.997 | 164 | 0.355 | 0.997 | 484 | 0.523 | 0.997 | 928.5 | 0.312 | 0.997 | 57 | 0.042 | 0.236 | 302.5 | 0.916 | 0.997 | 235 | 0.700 |
| BRD-K05444225-001-01-7 | E7449 | 737 | 0.658 | 0.992 | 1168 | 0.545 | 0.992 | 712 | 0.374 | 0.840 | 1841 | 0.007 | 0.153 | 589.5 | 0.262 | 0.840 | 580 | 0.906 | 0.992 | 1017.5 | 0.992 | 0.992 | 1310 | 0.386 | 0.840 | 582.5 | 0.910 | 0.992 | 876.5 | 0.400 | 0.840 | 387.5 | 0.149 | 0.785 | 171 | 0.150 | 0.785 | 388 | 0.925 | 0.992 | 326 | 0.767 |
| BRD-K05445342-001-05-3 | WP1066 | 4653 | 0.106 | 0.886 | 6425 | 0.152 | 0.886 | 4428 | 0.625 | 0.886 | 17306 | 0.730 | 0.886 | 5541 | 0.438 | 0.886 | 5018 | 0.838 | 0.890 | 5970 | 0.259 | 0.886 | 5693 | 0.217 | 0.886 | 2591 | 0.686 | 0.886 | 2884 | 0.139 | 0.886 | 3480 | 0.699 | 0.886 | 2579 | 0.483 | 0.886 | 782 | 0.354 | 0.886 | 1197 | 0.630 |
| BRD-K05524748-003-04-4 | dronedarone | 2914 | 0.543 | 0.899 | 5521 | 0.889 | 0.937 | 1589 | 0.013 | 0.265 | 9750 | 0.562 | 0.899 | 3926 | 0.645 | 0.899 | 3132 | 0.931 | 0.937 | 2976 | 0.215 | 0.621 | 3980 | 0.937 | 0.937 | 1361 | 0.184 | 0.621 | 1389 | 0.209 | 0.621 | 1416 | 0.054 | 0.564 | 1242 | 0.632 | 0.899 | 404 | 0.685 | 0.899 | 672 | 0.838 |
| BRD-K05638300-001-01-2 | sevoflurane | 564 | 0.925 | 0.970 | 600.5 | 0.059 | 0.625 | 763 | 0.933 | 0.970 | 1109.5 | 0.249 | 0.915 | 403 | 0.403 | 1.000 | 155 | 0.029 | 0.599 | 534.5 | 0.970 | 0.970 | 404 | 0.217 | 0.915 | 184.5 | 0.475 | 1.000 | 397.5 | 0.861 | 0.970 | 377.5 | 0.540 | 0.873 | 330 | 0.865 | 0.970 | 196.5 | 0.911 | 0.970 | 26 | 0.155 |
| BRD-K05804044-001-06-0 | AZ-628 | 790 | 0.037 | 0.323 | 741.5 | 0.000 | 0.000 | 1458 | 0.948 | 0.987 | 3836 | 0.987 | 0.987 | 1129.5 | 0.664 | 1.000 | 725.5 | 0.613 | 1.000 | 1663 | 0.654 | 1.000 | 957 | 0.494 | 1.000 | 129 | 0.046 | 0.323 | 641 | 0.876 | 0.987 | 594.5 | 0.787 | 0.987 | 505 | 0.980 | 0.987 | 253 | 0.743 | 0.987 | 177 | 0.392 |
| BRD-K05804044-001-18-5 | AZ-628 | 561 | 0.073 | 0.383 | 618.5 | 0.000 | 0.000 | 1012.5 | 0.484 | 0.912 | 3769 | 1.000 | 1.000 | 633.5 | 0.565 | 0.912 | 1477.5 | 0.996 | 1.000 | 688 | 0.479 | 0.912 | 1259 | 0.974 | 1.000 | 200 | 0.016 | 0.165 | 992 | 0.982 | 1.000 | 347.5 | 0.468 | 0.912 | 433.5 | 0.971 | 1.000 | 145.5 | 0.862 | 1.000 | 92 | 0.111 |
| BRD-K06152156-001-01-8 | methoxyflurane | 1616 | 0.379 | 0.711 | 2923 | 0.406 | 0.711 | 1747.5 | 0.250 | 0.704 | 6329 | 0.645 | 0.903 | 2072 | 0.082 | 0.704 | 1443 | 0.177 | 0.704 | 2858 | 0.953 | 0.953 | 3366 | 0.900 | 0.953 | 850 | 0.337 | 0.707 | 1377 | 0.584 | 0.876 | 1073 | 0.072 | 0.704 | 706.5 | 0.302 | 0.704 | 295 | 0.580 | 0.876 | 153 | 0.143 |
| BRD-K06294530-001-01-5 | UK-383367 | 46 | 0.070 | 0.738 | 671 | 0.726 | 1.000 | 521 | 0.913 | 1.000 | 1093 | 0.052 | 0.738 | 416 | 0.356 | 0.948 | 90 | 0.809 | 1.000 | 373 | 0.633 | 1.000 | 510 | 0.338 | 0.948 | 101 | 0.361 | 0.948 | 229 | 0.809 | 1.000 | 440 | 0.873 | 0.873 | 141 | 0.277 | 0.948 | 79 | 0.211 | 0.948 | | |
| BRD-K06426971-001-02-7 | ryuvidine | 3983 | 0.966 | 0.966 | 3839 | 0.001 | 0.024 | 2599 | 0.551 | 0.878 | 11562 | 0.911 | 0.966 | 4076 | 0.585 | 0.878 | 3085 | 0.697 | 0.966 | 4482 | 0.854 | 0.966 | 4062 | 0.238 | 0.787 | 714 | 0.246 | 0.787 | 2686 | 0.964 | 0.966 | 2215 | 0.492 | 0.867 | 887 | 0.112 | 0.588 | 699 | 0.495 | 0.867 | 349 | 0.009 |
| BRD-K06519765-065-01-6 | vinblastine | 5177 | 0.896 | 1.000 | 7482 | 0.921 | 1.000 | 3376 | 0.052 | 0.386 | 15487 | 0.337 | 0.644 | 5200 | 0.483 | 0.725 | 4244 | 0.860 | 1.000 | 7730 | 1.000 | 1.000 | 6066 | 0.254 | 0.534 | 869 | 0.001 | 0.017 | 4639 | 0.997 | 1.000 | 2595 | 0.075 | 0.386 | 2254 | 0.462 | 0.725 | 516 | 0.093 | 0.386 | 751 | 0.137 |
| BRD-K06543683-066-02-1 | bisindolylmaleimide-ix | 810 | 0.001 | 0.013 | 4825 | 0.936 | 0.990 | 2623 | 0.466 | 0.979 | 7456 | 0.214 | 0.627 | 3282 | 0.579 | 0.990 | 2162 | 0.583 | 0.990 | 3996 | 0.990 | 0.990 | 3501 | 0.769 | 0.990 | 1349 | 0.693 | 0.990 | 1561 | 0.910 | 0.990 | 1357 | 0.072 | 0.380 | 1074 | 0.753 | 0.990 | 638 | 0.860 | 0.990 | 692 | 0.354 |
| BRD-K06593056-001-04-8 | LE-135 | 603 | 0.401 | 0.791 | 2874 | 0.928 | 0.974 | 1327 | 0.310 | 1.000 | 5173 | 0.994 | 0.994 | 1698 | 0.457 | 0.800 | 899 | 0.023 | 0.237 | 1201 | 0.414 | 0.791 | 1835 | 0.797 | 0.935 | 673 | 0.833 | 0.935 | 96 | 0.004 | 0.010 | 862 | 0.760 | 0.935 | 496 | 0.385 | 0.791 | 462 | 0.596 | 0.893 | 280 | 0.757 |
| BRD-K06749501-001-02-1 | GSK1838705A | 467 | 0.328 | 0.811 | 795 | 0.071 | 0.735 | 497 | 0.661 | 1.000 | 1264 | 0.163 | 0.747 | 777 | 0.749 | 1.000 | 381 | 0.996 | 1.000 | 1065 | 0.992 | 1.000 | 343 | 0.562 | 0.983 | 201 | 0.884 | 1.000 | 219 | 0.276 | 0.811 | 370 | 0.909 | 0.909 | 52 | 0.347 | 0.811 | 31 | 0.178 | 0.747 | 40 | 0.009 |
| BRD-K06814349-304-01-9 | fosbretabulin | 1797 | 0.017 | 0.175 | 3530 | 0.013 | 0.175 | 1935 | 0.412 | 0.721 | 9117 | 0.667 | 0.998 | 3396 | 0.341 | 0.721 | 3261 | 0.997 | 0.999 | 3692 | 0.952 | 0.999 | 3598 | 0.761 | 0.998 | 873 | 0.371 | 0.721 | 2451 | 0.999 | 0.999 | 1893 | 0.220 | 0.678 | 846 | 0.058 | 0.242 | 436 | 0.813 | 0.999 | 511 | 0.251 |
| BRD-K06814349-304-02-7 | fosbretabulin | 898.5 | 0.636 | 0.953 | 1280.5 | 0.824 | 0.995 | 918 | 0.294 | 0.953 | 2275 | 0.010 | 0.107 | 1248 | 0.450 | 0.953 | 562.5 | 0.480 | 0.953 | 1204.5 | 0.986 | 0.995 | 1339 | 0.776 | 0.995 | 455 | 0.430 | 0.953 | 1266.5 | 0.995 | 0.995 | 723.5 | 0.429 | 0.953 | 308 | 0.008 | 0.107 | 298 | 0.537 | 0.953 | 322 | 0.633 |
| BRD-K06854232-001-19-9 | AM-580 | 1127 | 0.407 | 0.841 | 2496 | 0.818 | 0.971 | 1174 | 0.050 | 0.521 | 4577 | 0.745 | 0.971 | 1448 | 0.302 | 0.841 | 664 | 0.054 | 0.521 | 2125 | 0.917 | 0.971 | 1786 | 0.676 | 0.971 | 218 | 0.480 | 0.841 | 959 | 0.877 | 0.971 | 789 | 0.170 | 0.714 | 222 | 0.956 | 0.971 | 310 | 0.420 | 0.841 | 327 | 0.480 |
| BRD-K06858286-001-01-3 | cabazitaxel | 6269 | 0.983 | 0.999 | 8506.5 | 0.954 | 0.999 | 2909 | 0.002 | 0.039 | 14674 | 0.109 | 0.503 | 4822 | 0.120 | 0.503 | 5891 | 0.996 | 0.999 | 6286 | 0.689 | 0.965 | 6856 | 0.580 | 0.869 | 2322 | 0.286 | 0.619 | 4395 | 0.999 | 0.999 | 3119 | 0.444 | 0.717 | 2393 | 0.344 | 0.619 | 764 | 0.336 | 0.619 | 967 | 0.336 |
| BRD-K07202345-001-19-4 | frentizole | 1127.5 | 0.602 | 0.983 | 1773 | 0.480 | 1.000 | 1159 | 0.663 | 0.983 | 2817.5 | 0.015 | 0.309 | 1379 | 0.259 | 1.000 | 932 | 0.388 | 1.000 | 1595.5 | 0.983 | 0.983 | 1527.5 | 0.793 | 0.983 | 559 | 0.902 | 0.983 | 928 | 0.929 | 0.983 | 622 | 0.063 | 0.664 | 0 | 0.043 | 0.101 | 225 | 0.217 | 0.415 | 110 | 0.131 |
| BRD-K07265709-003-05-6 | dexrazoxane | 2897 | 0.913 | 0.999 | 5224 | 0.765 | 0.999 | 2094 | 0.202 | 0.505 | 9264 | 0.185 | 0.505 | 3592 | 0.754 | 0.999 | 2986 | 0.945 | 0.999 | 4349 | 0.981 | 0.999 | 3767 | 0.409 | 0.660 | 932 | 0.004 | 0.075 | 1459 | 0.999 | 0.999 | 1901 | 0.288 | 0.550 | 665 | 0.088 | 0.505 | 218 | 0.216 | 0.505 | 253 | 0.075 |
| BRD-K07310275-001-02-5 | PF-04691502 | 5159 | 0.767 | 0.983 | 6556 | 0.406 | 0.852 | 3928 | 0.214 | 0.749 | 14816 | 0.552 | 0.983 | 4393 | 0.116 | 0.608 | 3358 | 0.611 | 0.983 | 7035 | 0.942 | 0.983 | 5444 | 0.369 | 0.852 | 1202 | 0.098 | 0.608 | 3663 | 0.823 | 0.983 | 2121 | 0.010 | 0.213 | 2316 | 0.763 | 0.983 | 1122 | 0.883 | 0.983 | 1219 | 0.748 |
| BRD-K07442505-001-03-6 | BAM7 | 186 | 0.794 | 0.853 | 393 | 0.810 | 0.853 | 239 | 0.793 | 0.853 | 773 | 0.055 | 0.851 | 239 | 0.480 | 0.853 | 105 | 0.557 | 0.853 | 348 | 0.591 | 0.853 | 418 | 0.500 | 0.853 | 216 | 0.650 | 0.853 | 19 | 0.126 | 0.851 | 50 | 0.097 | 0.851 | 147 | 0.491 | 0.853 | 115 | 0.658 | 0.853 | 40 | 0.356 |
| BRD-K07487750-001-03-3 | WHI-P154 | 414 | 0.921 | 0.933 | 703 | 0.720 | 1.000 | 522 | 0.281 | 1.000 | 1097 | 0.061 | 0.428 | 744 | 0.828 | 0.933 | 511 | 0.842 | 0.933 | 597 | 0.742 | 0.933 | 247 | 0.230 | 0.603 | 274 | 0.659 | 0.933 | 477 | 0.506 | 0.933 | 200 | 0.028 | 0.290 | 215 | 0.687 | 0.933 | | | | 100 | 0.856 |
| BRD-K07572174-001-39-4 | curcumin | 2189 | 0.207 | 0.395 | 4556 | 0.954 | 1.000 | 1962 | 0.136 | 0.388 | 10555 | 1.000 | 1.000 | 2706 | 0.045 | 0.362 | 1631 | 0.177 | 0.395 | 3629 | 0.796 | 1.000 | 2822 | 0.204 | 0.395 | 419 | 0.148 | 0.388 | 1581 | 0.934 | 1.000 | 1211 | 0.071 | 0.372 | 409 | 0.009 | 0.193 | 703 | 0.941 | 1.000 | 523 | 0.319 |
| BRD-K07612980-001-05-3 | sparfloxacin | 928 | 0.637 | 1.000 | 1729 | 0.358 | 0.939 | 855 | 0.281 | 0.939 | 2989 | 0.104 | 0.925 | 1484 | 0.147 | 0.925 | 921 | 0.942 | 1.000 | 1440 | 0.722 | 1.000 | 1322 | 0.357 | 0.939 | 229 | 0.246 | 0.939 | 679 | 0.492 | 0.990 | 736 | 0.971 | 0.971 | 96 | 0.104 | 0.925 | 235 | 0.667 | 1.000 | 123 | 0.176 |
| BRD-K07736136-001-04-6 | VX-702 | 433 | 0.709 | 0.952 | 370 | 0.845 | 0.968 | 133 | 0.040 | 0.749 | 871.5 | 0.125 | 0.749 | 547.5 | 0.908 | 0.968 | 123.5 | 0.689 | 0.952 | 248 | 0.112 | 0.749 | 389 | 0.725 | 0.952 | 203.5 | 0.837 | 0.968 | 127 | 0.312 | 0.654 | 266.5 | 0.601 | 0.952 | 96.5 | 0.932 | 0.968 | 181.5 | 0.968 | 0.968 | 122.5 | 0.681 |
| BRD-K07762753-001-04-4 | aminopurvalanol-a | 1418 | 0.699 | 0.863 | 3088.5 | 1.000 | 1.000 | 1557 | 0.248 | 0.579 | 5277.5 | 0.245 | 0.579 | 1749.5 | 0.143 | 0.579 | 915 | 0.419 | 0.800 | 1739.5 | 0.500 | 0.863 | 2029.5 | 0.745 | 0.869 | 1156.5 | 0.659 | 0.863 | 522 | 0.044 | 0.232 | 764 | 0.029 | 0.603 | | | | 210 | 0.366 | 0.768 | 697 | 0.932 |
| BRD-K07881437-001-04-6 | danusertib | 6404 | 0.839 | 0.998 | 9278 | 0.991 | 0.998 | 3516 | 0.010 | 0.105 | 16594 | 0.403 | 0.817 | 4571 | 0.020 | 0.105 | 5337 | 0.985 | 0.998 | 7668 | 0.904 | 0.998 | 7241 | 0.761 | 0.998 | 1627 | 0.011 | 0.105 | 4732 | 0.998 | 0.998 | 2420 | 0.018 | 0.105 | 2191 | 0.155 | 0.465 | 1310 | 0.942 | 0.998 | 1065 | 0.428 |
| BRD-K07955840-001-02-3 | PF-05212384 | 7930 | 0.999 | 1.000 | 9316 | 0.910 | 1.000 | 4790 | 0.340 | 0.648 | 17025 | 0.321 | 0.648 | 6033 | 0.264 | 0.616 | 3638 | 0.025 | 0.173 | 8022 | 0.899 | 1.000 | 7399 | 0.762 | 1.000 | 1870 | 0.082 | 0.245 | 6041 | 1.000 | 1.000 | 2575 | 0.013 | 0.134 | 1575 | 0.051 | 0.213 | 510 | 0.065 | 0.229 | 909 | 0.491 |
| BRD-K07972848-001-02-0 | GSK2126458 | 5333 | 0.886 | 0.990 | 5799 | 0.464 | 0.886 | 3116 | 0.026 | 0.182 | 13353 | 0.755 | 0.990 | 4032 | 0.129 | 0.677 | 2811 | 0.445 | 0.886 | 5200 | 0.299 | 0.886 | 6733 | 0.990 | 0.990 | 1463 | 0.384 | 0.886 | 3539 | 0.937 | 0.990 | 1720 | 0.004 | 0.044 | 2131 | 0.868 | 0.990 | 530 | 0.384 | 0.886 | 531 | 0.386 |
| BRD-K08078237-001-19-8 | tetrandrine | 1110 | 0.024 | 0.169 | 3310 | 0.792 | 0.973 | 1196 | 0.104 | 0.397 | 5643 | 0.749 | 0.973 | 2801 | 0.508 | 0.820 | 1830 | 0.880 | 0.973 | 2843 | 0.877 | 0.973 | 3335 | 0.970 | 0.995 | 463 | 0.351 | 0.662 | 953 | 0.185 | 0.406 | 663 | 0.005 | 0.103 | 105 | 0.081 | 0.397 | 370 | 0.164 | 0.406 | 335 | 0.113 |
| BRD-K08109215-001-06-4 | I-BET-762 | 932 | 0.407 | 0.760 | 1365 | 0.424 | 0.760 | 610 | 0.580 | 0.811 | 2277 | 0.509 | 0.763 | 960 | 0.310 | 0.760 | 505 | 0.774 | 0.957 | 1520 | 0.997 | 0.997 | 883 | 0.833 | 0.957 | 152 | 0.007 | 0.150 | 577 | 0.729 | 0.957 | 478 | 0.416 | 0.760 | 448 | 0.322 | 0.760 | 254 | 0.491 | 0.763 | 184 | 0.201 |
| BRD-K08248804-001-01-8 | LCL-161 | 1330.5 | 0.334 | 0.779 | 2846 | 0.043 | 0.308 | 1921 | 0.330 | 0.779 | 6505 | 0.585 | 0.999 | 1711 | 0.226 | 0.779 | 1448 | 0.192 | 0.779 | 2422 | 0.783 | 0.999 | 2008 | 0.008 | 0.176 | 501 | 0.059 | 0.308 | 900 | 0.990 | 0.999 | 1235 | 0.051 | 0.308 | 1002.5 | 0.972 | 0.999 | 513 | 0.788 | 0.999 | 265 | 0.950 |
| BRD-K08303368-001-10-0 | chlorpyrifos | 343.5 | 0.344 | 0.698 | 831.5 | 0.798 | 0.931 | 557.5 | 0.500 | 0.774 | 1296.5 | 0.169 | 0.698 | 797 | 0.553 | 0.774 | 52 | 0.073 | 0.698 | 573 | 0.361 | 0.698 | 730.5 | 0.516 | 0.774 | 185.5 | 0.446 | 0.774 | 435.5 | 0.991 | 0.991 | 169 | 0.349 | 0.698 | 95 | 0.255 | 0.698 | 48 | 0.324 | 0.698 | 112 | 0.366 |
| BRD-K08542803-001-02-3 | gambogic-acid | 2011 | 0.678 | 0.966 | 4304 | 0.006 | 0.040 | 2585 | 0.099 | 0.411 | 13323 | 1.000 | 1.000 | 3348 | 0.294 | 0.617 | 2288 | 0.286 | 0.617 | 3217 | 0.920 | 0.966 | 3034 | 0.005 | 0.040 | 730 | 0.004 | 0.040 | 2785 | 0.621 | 0.966 | 2416 | 0.875 | 0.966 | 1314 | 0.049 | 0.258 | 710 | 0.857 | 0.966 | 472 | 0.145 |
| BRD-K08542803-001-03-1 | gambogic-acid | 3865 | 0.978 | 0.999 | 5001 | 0.309 | 0.590 | 2454 | 0.164 | 0.435 | 12310 | 0.999 | 0.999 | 2928 | 0.062 | 0.326 | 3823 | 0.972 | 0.999 | 5714 | 0.998 | 0.999 | 2710 | 0.001 | 0.022 | 1388 | 0.299 | 0.590 | 1400 | 0.012 | 0.131 | 2093 | 0.373 | 0.592 | 723 | 0.103 | 0.432 | 474 | 0.789 | 0.999 | 642 | 0.163 |
| BRD-K08547377-001-04-4 | irinotecan | 1466.5 | 0.473 | 0.834 | 3244.5 | 0.861 | 0.990 | 1889 | 0.076 | 0.399 | 5415.5 | 0.070 | 0.399 | 1455 | 0.003 | 0.066 | 1924.5 | 0.952 | 0.990 | 3001 | 0.990 | 0.990 | 2917 | 0.951 | 0.990 | 930 | 0.238 | 0.555 | 1433 | 0.934 | 0.990 | 1620.5 | 0.336 | 1.000 | 406 | 0.017 | 0.050 | 371 | 0.372 | 0.782 | 437 | 0.235 |
| BRD-K08547377-394-03-5 | irinotecan | 3349 | 0.505 | 0.884 | 5563 | 0.998 | 0.998 | 2073 | 0.072 | 0.378 | 8236 | 0.261 | 0.783 | 2483 | 0.022 | 0.153 | 1843 | 0.450 | 0.884 | 4003 | 0.939 | 0.998 | 3417.5 | 0.569 | 0.884 | 1331 | 0.589 | 0.884 | 1957 | 0.901 | 0.998 | 1660.5 | 0.243 | 0.783 | 191 | 0.000 | 0.008 | 614 | 0.788 | 0.998 | 138 | 0.017 |
| BRD-K08619838-300-05-7 | tremorine | 2481 | 0.689 | 0.904 | 3862 | 0.708 | 0.904 | 2020 | 0.113 | 0.792 | 7803 | 0.300 | 1.000 | 1672 | 0.066 | 0.690 | 2596.5 | 0.988 | 0.989 | 3010 | 0.785 | 0.916 | 4614 | 0.989 | 0.989 | 1104 | 0.834 | 0.922 | 1376 | 0.101 | 0.236 | 1642 | 0.350 | 1.000 | 806 | 0.153 | 0.320 | 413 | 0.397 | 0.759 | 433 | 0.449 |
| BRD-K08703257-001-12-1 | 3-amino-benzamide | 196.5 | 0.749 | 0.937 | 337.5 | 0.861 | 0.937 | 323 | 0.325 | 1.000 | 907 | 0.247 | 1.000 | 606 | 0.866 | 0.937 | 293 | 0.206 | 0.540 | 493.5 | 0.839 | 0.937 | 375 | 0.194 | 0.540 | 172.5 | 0.927 | 0.937 | 175 | 0.280 | 0.588 | 184 | 0.330 | 0.630 | 137.5 | 0.749 | 0.937 | 79 | 0.785 | 0.937 | 25 | 0.180 |
| BRD-K08799216-001-05-3 | pelitinib | 1812 | 0.865 | 1.000 | 2912 | 0.978 | 1.000 | 1116 | 0.049 | 0.345 | 5466.5 | 0.667 | 1.000 | 1882 | 0.094 | 0.491 | 1523 | 0.821 | 1.000 | 1418.5 | 0.842 | 1.000 | 2244 | 1.000 | 1.000 | 813 | 0.429 | 1.000 | 726 | 0.494 | 1.000 | 1206 | 0.330 | 0.991 | | | | 222 | 0.915 | 1.000 | 688 | 0.918 |
| BRD-K08893438-001-04-9 | CCG-50014 | 2354 | 0.043 | 0.310 | 6818 | 0.997 | 0.997 | 3731 | 0.857 | 0.997 | 8776 | 0.044 | 0.310 | 3633 | 0.293 | 0.800 | 3142 | 0.543 | 0.853 | 4872 | 0.904 | 0.997 | 4568 | 0.315 | 0.800 | 1157 | 0.381 | 0.800 | 2550 | 0.777 | 0.997 | 2046 | 0.155 | 0.652 | 1377 | 0.449 | 0.822 | 447 | 0.316 | 0.800 | 375 | 0.011 |
| BRD-K09090949-001-01-9 | CP-945,598 | 1257 | 0.979 | 0.979 | 1279 | 0.408 | 1.000 | 752 | 0.179 | 0.557 | 3194 | 0.809 | 0.979 | 1130 | 0.026 | 0.275 | 1229 | 0.634 | 1.000 | 518 | 0.186 | 0.557 | 1058 | 0.025 | 0.275 | 60 | 0.049 | 0.342 | 1006 | 0.961 | 0.979 | 579 | 0.155 | 0.557 | | | | 376 | 0.835 | 0.979 | 55 | 0.235 |
| BRD-K09372874-001-01-0 | merimepodib | 2285.5 | 0.631 | 0.883 | 4664 | 0.929 | 0.990 | 2343.5 | 0.384 | 1.000 | 8055.5 | 0.756 | 0.984 | 2438 | 0.140 | 1.000 | 2365.5 | 0.846 | 0.987 | 2673 | 0.598 | 0.883 | 3643 | 0.955 | 0.990 | 655 | 0.015 | 0.046 | 1855 | 0.797 | 0.984 | 1103 | 0.002 | 0.040 | 361 | 0.085 | 0.198 | 371 | 0.296 | 0.621 | 43 | 0.018 |
| BRD-K09426783-300-01-0 | TAS-103 | 4018 | 0.380 | 0.984 | 7298 | 0.909 | 0.984 | 3900 | 0.563 | 0.984 | 12291 | 0.131 | 0.467 | 3657 | 0.098 | 0.467 | 3624 | 0.881 | 0.984 | 6074 | 0.921 | 0.984 | 6519 | 0.938 | 0.984 | 1154 | 0.004 | 0.046 | 2754 | 0.708 | 0.984 | 1544 | 0.003 | 0.046 | 1039 | 0.133 | 0.467 | 1064 | 0.872 | 0.984 | 1040 | 0.572 |
| BRD-K09443272-001-03-3 | PHA-680632 | 3935 | 0.269 | 0.628 | 8926 | 0.996 | 1.000 | 3299 | 0.037 | 0.189 | 12801 | 0.149 | 0.448 | 3702 | 0.088 | 0.307 | 4607 | 0.962 | 1.000 | 5635 | 0.783 | 1.000 | 7693 | 0.998 | 1.000 | 1208 | 0.045 | 0.189 | 4288 | 1.000 | 1.000 | 2017 | 0.003 | 0.030 | 1989 | 0.872 | 1.000 | 963 | 0.733 | 1.000 | 1117 | 0.651 |
| BRD-K09549677-300-03-4 | mibefradil | 12 | 0.014 | 0.196 | 423 | 0.019 | 0.196 | 433 | 0.354 | 0.930 | 884 | 0.473 | 0.938 | 299 | 0.663 | 0.938 | 249 | 0.938 | 0.938 | 509 | 0.673 | 0.938 | 603 | 0.663 | 0.938 | 363 | 0.909 | 0.938 | 80 | 0.062 | 0.324 | 142 | 0.341 | 0.930 | 92 | 0.340 | 0.930 | 93 | 0.880 | 0.938 | 50 | 0.089 |
| BRD-K09877029-001-01-2 | VS-4718 | 933 | 0.700 | 0.919 | 1218.5 | 0.381 | 0.802 | 795.5 | 0.382 | 0.802 | 3356.5 | 0.425 | 0.812 | 1411 | 0.587 | 0.880 | 1024 | 0.993 | 0.993 | 1170 | 0.102 | 0.355 | 1754 | 0.920 | 0.972 | 745.5 | 0.896 | 0.972 | 530 | 0.656 | 0.919 | 409.5 | 0.117 | 1.000 | 591 | 0.554 | 0.880 | | | | 164 | 0.098 |
| BRD-K09951645-001-06-8 | dabrafenib | 382.5 | 0.436 | 1.000 | 302 | 0.000 | 0.000 | 625.5 | 0.494 | 1.000 | 2906 | 0.972 | 1.000 | 798.5 | 0.376 | 1.000 | 333 | 0.866 | 1.000 | 974.5 | 0.915 | 1.000 | 1038.5 | 0.777 | 1.000 | 385.5 | 0.752 | 1.000 | 317.5 | 0.217 | 1.000 | 415 | 0.983 | 0.983 | 118 | 0.779 | 1.000 | | | | 217 | 0.793 |
| BRD-K10843433-001-22-7 | phenylbutazone | 187 | 0.416 | 0.879 | 536 | 0.437 | 0.879 | 94 | 0.924 | 0.924 | 1021 | 0.888 | 0.924 | 383 | 0.894 | 0.924 | 385.5 | 0.727 | 0.924 | 323 | 0.230 | 0.847 | 427.5 | 0.502 | 0.879 | 118 | 0.652 | 0.924 | 90 | 0.118 | 0.826 | 202.5 | 0.841 | 0.924 | 29 | 0.009 | 0.187 | 37 | 0.320 | 0.879 | 114 | 0.236 |

| broad_id | name | al_nervous_system_PVAL_ADJ | fibroblast_MWU_STAT | fibroblast_PVAL | fibroblast_PVAL_ADJ | bile_duct_MWU_STAT | bile_duct_PVAL | bile_duct_PVAL_ADJ | thyroid_MWU_STAT | thyroid_PVAL | thyroid_PVAL_ADJ | upper_aerodigestive_MWU_STAT | upper_aerodigestive_PVAL | upper_aerodigestive_PVAL_ADJ | uterus_MWU_STAT | uterus_PVAL | uterus_PVAL_ADJ | liver_MWU_STAT | liver_PVAL | liver_PVAL_ADJ | esophagus_MWU_STAT | esophagus_PVAL | esophagus_PVAL_ADJ |
|---|---|---|---|---|---|---|---|---|---|---|---|---|---|---|---|---|---|---|---|---|---|---|---|
| BRD-A83650191-004-02-8 | A61603 | 0.781 | 112 | 0.343 | 0.720 | 377 | 0.114 | 0.597 | 1438.5 | 0.889 | 0.914 | 1991.5 | 0.235 | 0.704 | 2548.5 | 0.563 | 0.870 | 1133 | 0.061 | 0.597 | 1955 | 0.658 | 0.914 |
| BRD-A92630576-050-24-1 | trimebutine | 0.134 | | | | 35 | 0.173 | 0.936 | 267 | 0.333 | 0.688 | 511 | 0.454 | 0.795 | 575 | 0.870 | 0.966 | 311 | 0.525 | 0.843 | 590 | 0.148 | 0.936 |
| BRD-A92872453-001-01-8 | N-acetylmannosamine | 0.743 | 181.5 | 0.945 | 0.997 | 147 | 0.833 | 0.997 | 272 | 0.466 | 0.875 | 698 | 0.004 | 0.074 | 362 | 0.076 | 0.535 | 214 | 0.242 | 0.743 | 1531.5 | 0.935 | 0.997 |
| BRD-A93236127-001-04-5 | digitoxin | 0.079 | | | | 627 | 0.542 | 1.000 | 817 | 0.616 | 1.000 | 2209 | 0.924 | 0.997 | 1739 | 0.643 | 1.000 | 1087 | 0.030 | 0.078 | 1946 | 0.035 | 0.369 |
| BRD-A93477898-001-04-2 | PETCM | 1.000 | | | | | | | 18 | 0.132 | 0.692 | 130 | 0.033 | 0.577 | 262 | 0.544 | 1.000 | | | | 653 | 0.992 | 0.992 |
| BRD-A94756469-001-04-7 | digoxin | 0.345 | 197 | 0.462 | 0.879 | 1342 | 0.594 | 0.879 | 2102 | 0.508 | 0.879 | 4817 | 0.726 | 0.953 | 3160 | 0.082 | 0.345 | 2756 | 0.068 | 0.345 | 4144 | 0.540 | 0.879 |
| BRD-A95121829-001-08-9 | ginkgolide-a | 0.042 | | | | 268 | 0.982 | 0.982 | 74 | 0.121 | 1.000 | 249 | 0.046 | 0.969 | 466 | 0.890 | 0.970 | 485 | 0.924 | 0.970 | 523 | 0.633 | 0.949 |
| BRD-A97739905-001-26-5 | ketoprofen | 0.434 | | | | | | | 199 | 0.162 | 1.000 | 213 | 0.215 | 1.000 | 169 | 0.225 | 1.000 | 98 | 0.117 | 0.246 | 283 | 0.575 | 0.825 |
| BRD-A98299281-050-09-6 | hydroxytacrine-maleate-(r,s) | 0.569 | | | | 180 | 0.470 | 0.957 | 84 | 0.763 | 0.957 | 506 | 0.659 | 0.957 | 330 | 0.923 | 0.957 | 167 | 0.842 | 0.957 | 740 | 0.957 | 0.957 |
| BRD-K00259736-001-16-4 | colchicine | 0.648 | 92 | 0.255 | 0.648 | 1495 | 0.950 | 0.950 | 1396 | 0.699 | 0.928 | 2947 | 0.351 | 0.675 | 2318 | 0.129 | 0.648 | 2690 | 0.864 | 0.928 | 3340 | 0.414 | 0.724 |
| BRD-K00317371-001-06-1 | RITA | 0.929 | | | | 390 | 0.102 | 0.305 | 810 | 0.575 | 0.929 | 2167 | 0.989 | 0.989 | 1567 | 0.054 | 0.571 | 1080 | 0.024 | 0.100 | 3446 | 0.982 | 0.989 |
| BRD-K00494077-066-01-9 | trovafloxacin | | | | | | | | 482 | 0.767 | 0.847 | 933.5 | 0.688 | 0.839 | 697 | 0.462 | 1.000 | 300 | 0.395 | 0.751 | 561 | 0.505 | 0.751 |
| BRD-K00535541-001-06-3 | 9-aminoacridine | 0.718 | | | | 639 | 0.876 | 0.969 | 956 | 0.342 | 0.718 | 2168 | 0.466 | 0.815 | 1955 | 0.724 | 0.969 | 933 | 0.022 | 0.156 | 2515 | 0.398 | 0.760 |
| BRD-K00634421-001-01-0 | brivaracetam | 0.876 | 240 | 0.673 | 0.999 | 491 | 0.414 | 0.967 | 1476 | 0.602 | 0.999 | 2092 | 0.005 | 0.080 | 2545 | 0.198 | 0.694 | 2331 | 0.993 | 0.999 | 2155 | 0.008 | 0.080 |
| BRD-K01507359-001-22-9 | rifampin | 0.987 | | | | 592 | 0.995 | 0.995 | 721 | 0.927 | 1.000 | 872 | 0.392 | 0.858 | 839 | 0.173 | 0.725 | 644 | 0.148 | 0.725 | 979 | 0.817 | 1.000 |
| BRD-K01614093-001-02-6 | dihydromyricetin | 0.075 | | | | 136 | 0.081 | 0.548 | 159 | 0.130 | 0.548 | 1011 | 0.593 | 1.000 | 668 | 0.741 | 0.916 | 385 | 0.677 | 0.889 | 1112 | 0.910 | 0.978 |
| BRD-K01669786-001-02-2 | triapine | 0.790 | | | | 604 | 0.883 | 1.000 | 390 | 0.136 | 0.697 | 2828 | 0.973 | 1.000 | 1543 | 0.865 | 1.000 | 1693 | 0.713 | 1.000 | 2394 | 0.931 | 1.000 |
| BRD-K01683783-001-02-6 | TAK-901 | 0.384 | 59 | 0.138 | 0.447 | 1444 | 0.904 | 1.000 | 2002 | 0.591 | 0.954 | 3569 | 0.045 | 0.316 | 3578 | 0.149 | 0.447 | 3631 | 0.930 | 1.000 | 4277 | 0.216 | 0.505 |
| BRD-K01773560-001-02-4 | UNC2250 | 0.294 | | | | 457.5 | 0.931 | 0.976 | 105 | 0.042 | 0.294 | 496 | 0.569 | 0.854 | 655.5 | 0.367 | 0.845 | 295.5 | 0.388 | 0.845 | 1219.5 | 0.837 | 1.000 |
| BRD-K02113016-001-15-4 | olaparib | 0.976 | | | | 206 | 0.956 | 0.976 | 80 | 0.201 | 1.000 | 322.5 | 0.884 | 0.976 | 639.5 | 0.743 | 0.976 | 229 | 0.786 | 0.976 | 403.5 | 0.476 | 0.833 |
| BRD-K02130563-001-11-4 | panobinostat | 0.164 | 115 | 0.292 | 0.682 | 931 | 0.427 | 0.690 | 807 | 0.009 | 0.164 | 4040 | 0.706 | 0.872 | 2676 | 0.061 | 0.322 | 2367 | 0.127 | 0.504 | 2949 | 0.413 | 0.690 |
| BRD-K02241333-001-02-7 | CHIR-124 | 0.439 | 60 | 0.178 | 0.499 | 878 | 0.563 | 0.861 | 1910 | 0.920 | 0.998 | 3866 | 1.000 | 1.000 | 2786 | 0.250 | 0.582 | 1529 | 0.090 | 0.439 | 3159 | 0.725 | 0.959 |
| BRD-K02407574-001-09-7 | parbendazole | 0.498 | 20 | 0.049 | 0.256 | 1913 | 0.984 | 0.999 | 2186 | 0.583 | 0.942 | 2633 | 0.003 | 0.070 | 3306 | 0.214 | 0.499 | 3177 | 0.714 | 0.999 | 4034 | 0.129 | 0.466 |
| BRD-K02594908-001-16-4 | sulfanilamide | 0.930 | | | | 110 | 0.044 | 0.465 | 244 | 0.432 | 1.000 | 933 | 0.746 | 0.930 | 560 | 0.214 | 1.000 | 305 | 0.015 | 0.040 | 644 | 0.023 | 0.465 |
| BRD-K02764365-001-13-6 | methiazole | 0.485 | | | | 748 | 0.985 | 0.985 | 770.5 | 0.335 | 1.000 | 1785 | 0.025 | 0.262 | 963 | 0.017 | 0.262 | 1202 | 0.388 | 0.770 | 2100.5 | 0.146 | 0.769 |
| BRD-K02898090-001-02-7 | kifunensine | 1.000 | | | | | | | 343.5 | 0.484 | 1.000 | 599 | 0.689 | 1.000 | 297 | 0.586 | 1.000 | 197.5 | 0.424 | 1.000 | 660 | 0.957 | 0.957 |
| BRD-K03063480-001-06-6 | PF-477736 | 0.357 | 2 | 0.044 | 0.308 | 1381 | 0.742 | 0.924 | 1825 | 0.550 | 0.924 | 4315 | 0.984 | 0.984 | 1751 | 0.000 | 0.004 | 2773.5 | 0.707 | 0.924 | 5105 | 0.831 | 0.924 |
| BRD-K03109492-001-02-2 | NSC-663284 | 0.468 | 6 | 0.049 | 0.468 | 765 | 0.798 | 0.949 | 1245 | 0.775 | 0.949 | 2481 | 0.627 | 0.949 | 1305 | 0.023 | 0.468 | 2095.5 | 0.933 | 0.949 | 3406 | 0.949 | 0.949 |
| BRD-K03243820-001-23-8 | trifluridine | 0.886 | 130 | 0.265 | 0.795 | 1236 | 0.799 | 0.989 | 1720 | 0.401 | 0.886 | 3918 | 0.507 | 0.886 | 3594 | 0.825 | 0.989 | 3900.5 | 0.980 | 0.989 | 5730 | 0.989 | 0.989 |
| BRD-K03273112-001-01-8 | 1-azakenpaullone | 0.215 | | | | 176 | 0.561 | 0.842 | 245 | 0.194 | 0.733 | 392 | 0.086 | 0.733 | 459 | 0.410 | 0.733 | 327 | 0.226 | 0.733 | 1201 | 0.985 | 0.985 |
| BRD-K03289018-001-02-7 | CCT137690 | 0.645 | 199 | 0.564 | 0.911 | 996 | 0.655 | 0.921 | 1743 | 0.456 | 0.911 | 3915 | 0.839 | 0.960 | 2795 | 0.344 | 0.906 | 2253 | 0.456 | 0.911 | 3599 | 0.857 | 0.960 |
| BRD-K03390685-001-01-7 | cobimetinib | 0.700 | | | | 352 | 0.332 | 0.775 | 816 | 0.501 | 1.000 | 1657 | 0.019 | 0.133 | 2380 | 0.998 | 1.000 | 1515 | 0.572 | 1.000 | 2059 | 0.884 | 1.000 |
| BRD-K03406345-001-27-8 | azacitidine | 0.796 | 140 | 0.673 | 0.898 | 555 | 0.816 | 0.898 | 431 | 0.815 | 0.898 | 812 | 0.099 | 0.693 | 1001 | 0.571 | 0.898 | 832 | 0.873 | 0.898 | 1354 | 0.190 | 0.796 |
| BRD-K03449891-001-08-6 | foretinib | 0.984 | 77 | 0.268 | 0.646 | 72 | 0.044 | 0.233 | 790 | 0.896 | 0.984 | 2093 | 0.927 | 0.984 | 857.5 | 0.033 | 0.233 | 1510 | 0.944 | 0.984 | 1355 | 0.636 | 0.984 |
| BRD-K03549949-001-01-0 | XL388 | 0.850 | | | | 1179 | 0.881 | 1.000 | 1261 | 0.995 | 0.995 | 2355 | 0.486 | 0.850 | 1741 | 0.615 | 0.994 | 1371 | 0.167 | 0.606 | 3428 | 0.955 | 0.995 |
| BRD-K03601870-001-01-2 | ixabepilone | 0.668 | 261 | 0.589 | 0.898 | 1248 | 0.693 | 0.970 | 1837 | 0.201 | 0.668 | 3667 | 0.244 | 0.668 | 3976 | 0.313 | 0.668 | 3198 | 0.804 | 0.998 | 3862 | 0.050 | 0.375 |
| BRD-K03765900-001-01-9 | XL-647 | 1.000 | | | | 367 | 0.530 | 0.927 | 391 | 0.608 | 0.982 | 790 | 0.000 | 0.001 | 1297 | 0.218 | 0.764 | 810 | 0.124 | 0.261 | 1402 | 0.007 | 0.076 |
| BRD-K04111260-001-10-0 | raclopride | 0.328 | | | | | | | 181 | 0.510 | 1.000 | 834 | 0.925 | 1.000 | 296.5 | 0.973 | 1.000 | 271 | 0.381 | 1.000 | 387 | 0.666 | 0.666 |
| BRD-K04394237-001-03-7 | drospirenone | 0.987 | | | | 325 | 0.551 | 0.987 | 556 | 0.881 | 0.987 | 1486 | 0.975 | 0.987 | 663 | 0.035 | 0.245 | 754.5 | 0.962 | 0.987 | 1652 | 0.987 | 0.987 |
| BRD-K04548931-003-16-5 | epirubicin | 0.529 | | | | 650 | 0.628 | 0.942 | 566 | 0.193 | 0.529 | 1735 | 0.335 | 0.704 | 1038 | 0.064 | 0.445 | 1117 | 0.227 | 0.529 | 2991 | 0.898 | 0.898 |
| BRD-K04568635-300-01-4 | octenidine | 0.913 | | | | 332 | 0.351 | 0.801 | 170 | 0.403 | 0.801 | 708 | 0.240 | 0.720 | 790 | 0.420 | 0.801 | 172 | 0.031 | 0.165 | 888 | 0.655 | 0.913 |
| BRD-K04623885-001-07-5 | BIBR-1532 | 0.959 | 41 | 0.120 | 0.532 | 197 | 0.812 | 0.959 | 666 | 0.127 | 0.532 | 1612 | 0.327 | 0.959 | 1067 | 0.102 | 0.532 | 1066 | 0.374 | 0.959 | 1967 | 0.799 | 0.959 |
| BRD-K04701033-001-03-9 | VE-822 | 0.458 | 257 | 0.562 | 0.984 | 1184 | 0.299 | 0.666 | 2390 | 0.637 | 0.984 | 4775 | 0.617 | 0.984 | 3503 | 0.033 | 0.197 | 3894 | 0.749 | 0.984 | 4362 | 0.147 | 0.440 |
| BRD-K04833372-001-06-2 | GSK1904529A | 0.331 | | | | 352.5 | 0.738 | 0.965 | 165.5 | 0.594 | 0.965 | 168 | 0.000 | 0.004 | 492.5 | 0.422 | 0.885 | 364.5 | 0.783 | 0.965 | 823 | 0.919 | 0.965 |
| BRD-K04923131-001-15-4 | GSK-3-inhibitor-IX | 0.594 | | | | 239 | 0.974 | 1.000 | 67 | 0.505 | 0.882 | 399 | 0.964 | 1.000 | 129 | 0.152 | 0.476 | 169 | 0.334 | 0.639 | 645 | 0.901 | 0.901 |
| BRD-K04993719-001-01-7 | LY2334737 | 0.104 | 115 | 0.335 | 0.808 | 701 | 0.853 | 0.943 | 1306 | 0.115 | 0.614 | 2502 | 0.362 | 0.808 | 2801 | 0.660 | 0.913 | 1839 | 0.696 | 0.913 | 3207 | 0.452 | 0.808 |
| BRD-K05104363-001-10-0 | PD-184352 | 0.997 | 169.5 | 0.902 | 0.997 | 333.5 | 0.678 | 0.997 | 207 | 0.056 | 0.236 | 869 | 0.051 | 0.236 | 1060.5 | 0.997 | 0.997 | 1022 | 0.959 | 0.997 | 1298 | 0.983 | 0.997 |
| BRD-K05444225-001-01-7 | E7449 | 0.992 | 112 | 0.681 | 0.992 | 474 | 0.895 | 0.992 | 385.5 | 0.340 | 0.840 | 941 | 0.375 | 0.840 | 627.5 | 0.819 | 0.992 | 230.5 | 0.037 | 0.384 | 1103 | 0.954 | 0.992 |
| BRD-K05445342-001-05-3 | WP1066 | 0.886 | 77 | 0.175 | 0.886 | 1423 | 0.375 | 0.886 | 2451 | 0.900 | 0.900 | 4486 | 0.663 | 0.886 | 5042 | 0.848 | 0.890 | 3788 | 0.760 | 0.886 | 4748 | 0.431 | 0.886 |
| BRD-K05524748-003-04-4 | dronedarone | 0.937 | 56 | 0.169 | 0.621 | 901 | 0.626 | 0.899 | 1252 | 0.409 | 0.899 | 2267 | 0.124 | 0.621 | 2150 | 0.237 | 0.621 | 2439 | 0.930 | 0.937 | 2450 | 0.538 | 0.899 |
| BRD-K05638300-001-01-2 | sevoflurane | 0.271 | | | | 182.5 | 0.859 | 0.970 | 148 | 0.261 | 0.915 | 537.5 | 0.090 | 0.628 | 316 | 0.532 | 1.000 | 173 | 0.815 | 0.970 | 247.5 | 0.489 | 1.000 |
| BRD-K05804044-001-06-0 | AZ-628 | 0.686 | | | | 754.5 | 0.979 | 0.987 | 537.5 | 0.171 | 0.895 | 1516.5 | 0.922 | 0.987 | 929 | 0.639 | 1.000 | 160 | 0.085 | 0.161 | 1072.5 | 0.967 | 0.987 |
| BRD-K05804044-001-18-5 | AZ-628 | 0.459 | 145.5 | 0.862 | 1.000 | 106 | 0.153 | 0.459 | 191 | 0.052 | 0.361 | 961.5 | 0.552 | 0.912 | 399.5 | 0.149 | 0.459 | 606 | 0.928 | 1.000 | 358 | 0.228 | 0.598 |
| BRD-K06152156-001-01-8 | methoxyflurane | 0.704 | 82 | 0.246 | 0.704 | 529.5 | 0.813 | 0.949 | 336 | 0.301 | 0.704 | 2148 | 0.746 | 0.922 | 1613 | 0.734 | 0.922 | 787 | 0.103 | 0.704 | 2274.5 | 0.939 | 0.953 |
| BRD-K06294530-001-01-5 | UK-383367 | | | | | 53 | 0.426 | 0.968 | 370 | 0.853 | 1.000 | 348 | 0.777 | 1.000 | 367 | 0.189 | 0.948 | 282 | 0.461 | 0.968 | 434 | 0.857 | 1.000 |
| BRD-K06426971-001-02-7 | ryuvidine | 0.093 | 96 | 0.273 | 0.787 | 494 | 0.429 | 0.867 | 950 | 0.353 | 0.823 | 3672 | 0.765 | 0.966 | 2189 | 0.300 | 0.787 | 1130 | 0.032 | 0.221 | 3675 | 0.867 | 0.966 |
| BRD-K06519765-065-01-6 | vinblastine | 0.386 | 73 | 0.173 | 0.403 | 1372 | 0.870 | 1.000 | 1350 | 0.073 | 0.386 | 3513 | 0.147 | 0.386 | 3315 | 0.140 | 0.386 | 3302 | 0.668 | 0.936 | 4103 | 0.379 | 0.664 |
| BRD-K06543683-066-02-1 | bisindolylmaleimide-ix | 0.826 | | | | 938 | 0.963 | 0.990 | 915 | 0.116 | 0.489 | 2125 | 0.239 | 0.627 | 1346 | 0.029 | 0.309 | 1275 | 0.198 | 0.627 | 2919 | 0.761 | 0.990 |
| BRD-K06593056-001-04-8 | LE-135 | 0.935 | 124 | 0.593 | 0.893 | | | | 361 | 0.034 | 0.238 | 1082 | 0.372 | 1.000 | 819 | 0.009 | 0.190 | 309 | 0.846 | 0.935 | 703 | 0.207 | 1.000 |
| BRD-K06749501-001-02-1 | GSK1838705A | 0.190 | | | | 105 | 0.829 | 1.000 | 181 | 0.387 | 0.813 | 635 | 0.105 | 0.735 | 226 | 0.307 | 0.811 | 195 | 0.468 | 0.893 | 686 | 0.832 | 1.000 |
| BRD-K06814349-304-01-9 | fosbretabulin | 0.678 | | | | 429 | 0.380 | 0.721 | 757 | 0.744 | 0.998 | 1906 | 0.068 | 0.475 | 1513 | 0.258 | 0.678 | 2042 | 0.952 | 0.999 | 3146 | 0.486 | 0.786 |
| BRD-K06814349-304-02-7 | fosbretabulin | 0.953 | 135 | 0.749 | 0.995 | 360 | 0.982 | 0.995 | 577.5 | 0.285 | 0.953 | 557 | 0.107 | 0.752 | 1139.5 | 0.575 | 0.953 | 921 | 0.860 | 0.995 | 941 | 0.339 | 0.953 |
| BRD-K06854232-001-19-9 | AM-580 | 0.841 | 186 | 0.874 | 0.971 | 95 | 0.400 | 0.841 | 687 | 0.971 | 0.971 | 1395 | 0.074 | 0.521 | 625 | 0.421 | 0.841 | 302 | 0.140 | 0.714 | 1419 | 0.572 | 0.924 |
| BRD-K06858286-001-01-3 | cabazitaxel | 0.619 | 160 | 0.321 | 0.619 | 1588 | 0.823 | 0.999 | 1992 | 0.346 | 0.619 | 4186 | 0.354 | 0.619 | 3051 | 0.079 | 0.503 | 3780.5 | 0.782 | 0.999 | 3713 | 0.064 | 0.503 |
| BRD-K07202345-001-19-4 | frentizole | 0.275 | | | | 358.5 | 0.966 | 0.983 | 341 | 0.108 | 0.758 | 1336 | 0.796 | 0.983 | 1032 | 0.923 | 0.983 | 197 | 0.039 | 0.101 | 1496.5 | 0.871 | 0.983 |
| BRD-K07265709-003-05-6 | dexrazoxane | 0.505 | 68 | 0.206 | 0.505 | 1340 | 0.993 | 0.999 | 726 | 0.328 | 0.574 | 2525 | 0.206 | 0.505 | 2078 | 0.190 | 0.505 | 1424 | 0.664 | 0.996 | 3027 | 0.265 | 0.550 |
| BRD-K07310275-001-02-5 | PF-04691502 | 0.983 | 289 | 0.692 | 0.983 | 776 | 0.174 | 0.730 | 2844 | 0.983 | 0.983 | 3482 | 0.269 | 0.791 | 3347 | 0.301 | 0.791 | 1773 | 0.053 | 0.556 | 5367 | 0.925 | 0.983 |
| BRD-K07442505-001-03-6 | BAM7 | 0.853 | 74 | 0.813 | 0.853 | 110 | 0.609 | 0.853 | 143 | 0.162 | 0.851 | 358 | 0.220 | 0.853 | 74 | 0.813 | 0.853 | 230 | 0.741 | 0.853 | 401 | 0.961 | 0.961 |
| BRD-K07487750-001-03-3 | WHI-P154 | 0.933 | 100 | 0.856 | 0.933 | 199 | 0.933 | 0.933 | 100 | 0.856 | 0.933 | 154 | 0.095 | 0.498 | 50 | 0.013 | 0.268 | 154 | 0.723 | 0.933 | 476 | 0.732 | 0.933 |
| BRD-K07572174-001-39-4 | curcumin | 0.558 | 15 | 0.052 | 0.362 | 431 | 0.433 | 0.650 | 972 | 0.853 | 1.000 | 2260 | 0.405 | 0.650 | 1290 | 0.116 | 0.388 | 1208 | 0.738 | 1.000 | 2179 | 0.116 | 0.388 |
| BRD-K07612980-001-05-3 | sparfloxacin | 0.925 | | | | 404 | 0.854 | 1.000 | 525 | 0.601 | 1.000 | 853 | 0.465 | 0.990 | 1106 | 0.792 | 1.000 | 594 | 0.518 | 0.990 | 1290 | 0.898 | 1.000 |
| BRD-K07736136-001-04-6 | VX-702 | 0.952 | | | | 49 | 0.467 | 1.000 | 57 | 0.143 | 0.749 | 233.5 | 0.192 | 0.806 | 52 | 0.507 | 1.000 | 99 | 0.467 | 0.892 | 343.5 | 0.505 | 1.000 |
| BRD-K07762753-001-04-4 | aminopurvalanol-a | 1.000 | | | | 390 | 0.579 | 0.863 | 416 | 0.659 | 0.863 | 1842 | 0.227 | 0.579 | 1153.5 | 0.954 | 1.000 | 1170.5 | 0.683 | 0.863 | 1380 | 0.167 | 0.579 |
| BRD-K07881437-001-04-6 | danusertib | 0.817 | 93 | 0.208 | 0.545 | 1475 | 0.660 | 0.998 | 2479 | 0.741 | 0.998 | 3626 | 0.060 | 0.253 | 3765 | 0.094 | 0.330 | 3893 | 0.780 | 0.998 | 4721 | 0.357 | 0.817 |
| BRD-K07955840-001-02-3 | PF-05212384 | 0.859 | 294 | 0.639 | 0.958 | 1670 | 0.820 | 1.000 | 2366 | 0.601 | 0.958 | 3383 | 0.009 | 0.134 | 3410 | 0.042 | 0.213 | 4610 | 0.977 | 1.000 | 4308 | 0.118 | 0.309 |
| BRD-K07972848-001-02-0 | GSK2126458 | 0.886 | 235 | 0.593 | 0.990 | 1141 | 0.728 | 0.990 | 2187 | 0.975 | 0.990 | 2075 | 0.000 | 0.007 | 4382 | 0.945 | 0.990 | 2838 | 0.795 | 0.990 | 3454 | 0.163 | 0.684 |
| BRD-K08078237-001-19-8 | tetrandrine | 0.397 | | | | 448 | 0.650 | 0.973 | 1407 | 0.995 | 0.995 | 1521 | 0.193 | 0.406 | 1682 | 0.869 | 0.973 | 926 | 0.155 | 0.406 | 1560 | 0.378 | 0.662 |
| BRD-K08109215-001-06-4 | I-BET-762 | 0.743 | 152 | 0.907 | 0.957 | 34 | 0.148 | 0.743 | 334 | 0.212 | 0.743 | 794 | 0.434 | 0.760 | 571 | 0.912 | 0.957 | 306 | 0.146 | 0.743 | 605 | 0.174 | 0.743 |
| BRD-K08248804-001-01-8 | LCL-161 | 0.999 | 146 | 0.553 | 0.999 | 816.5 | 0.962 | 0.999 | 941 | 0.940 | 0.999 | 1663 | 0.302 | 0.779 | 1248 | 0.938 | 0.999 | 1528.5 | 0.634 | 0.999 | 2854 | 0.999 | 0.999 |
| BRD-K08303368-001-10-0 | chlorpyrifos | 0.698 | 16 | 0.098 | 0.698 | 242 | 0.981 | 0.991 | 228 | 0.348 | 0.698 | 552.5 | 0.692 | 0.855 | 222 | 0.662 | 0.855 | 191.5 | 0.192 | 0.698 | 350.5 | 0.890 | 0.984 |
| BRD-K08542803-001-02-3 | gambogic-acid | 0.436 | 190 | 0.547 | 0.966 | 446 | 0.117 | 0.411 | 1612 | 0.609 | 0.966 | 3543 | 0.719 | 0.966 | 2530 | 0.236 | 0.617 | 2579 | 0.866 | 0.966 | 2307 | 0.797 | 0.966 |
| BRD-K08542803-001-03-1 | gambogic-acid | 0.435 | 118 | 0.337 | 0.590 | 352 | 0.504 | 0.706 | 924 | 0.020 | 0.137 | 3165 | 0.880 | 0.999 | 2159 | 0.166 | 0.435 | 1775 | 0.218 | 0.509 | 3057 | 0.395 | 0.592 |
| BRD-K08547377-001-04-4 | irinotecan | 0.555 | | | | 552.5 | 0.835 | 0.990 | 722 | 0.051 | 0.399 | 2105.5 | 0.482 | 0.834 | 2007 | 0.516 | 0.834 | 1602 | 0.837 | 0.990 | 2824.5 | 0.963 | 0.990 |
| BRD-K08547377-394-03-5 | irinotecan | 0.153 | 139 | 0.403 | 0.884 | 416 | 0.757 | 0.998 | 1047 | 0.390 | 0.884 | 2976 | 0.560 | 0.884 | 1934 | 0.132 | 0.555 | 2119.5 | 0.967 | 0.998 | 3113.5 | 0.811 | 0.998 |
| BRD-K08619838-300-05-7 | tremorine | 0.786 | | | | 546.5 | 0.732 | 0.904 | 505 | 0.291 | 1.000 | 1950 | 0.250 | 1.000 | 920 | 0.057 | 0.690 | 1876 | 0.654 | 0.904 | 2581 | 0.498 | 0.805 |
| BRD-K08703257-001-12-1 | 3-amino-benzamide | 0.540 | 102.5 | 0.937 | 0.937 | | | | 121 | 0.619 | 0.937 | 234 | 0.640 | 0.937 | 228 | 0.313 | 1.000 | 260 | 0.249 | 0.580 | 267 | 0.279 | 1.000 |
| BRD-K08799216-001-05-3 | pelitinib | 1.000 | 244.5 | 0.954 | 1.000 | 287 | 0.255 | 1.000 | 471 | 0.796 | 1.000 | 853 | 0.000 | 0.000 | 1829.5 | 0.609 | 1.000 | 1335 | 0.545 | 1.000 | 1151 | 0.005 | 0.055 |
| BRD-K08893438-001-04-9 | CCG-50014 | 0.238 | 292 | 0.809 | 0.997 | 581 | 0.093 | 0.490 | 1471 | 0.353 | 0.800 | 2572 | 0.470 | 0.822 | 3170 | 0.568 | 0.853 | 2131 | 0.722 | 0.997 | 4220 | 0.961 | 0.997 |
| BRD-K09090949-001-01-9 | CP-945,598 | 0.449 | | | | 336 | 0.970 | 0.979 | 558 | 0.951 | 0.979 | 882 | 0.145 | 0.557 | 831 | 0.979 | 0.979 | 427 | 0.372 | 0.651 | 1287 | 0.739 | 1.000 |
| BRD-K09372874-001-01-0 | merimepodib | 0.048 | | | | 259 | 0.363 | 1.000 | 1579.5 | 0.990 | 0.990 | 1928.5 | 0.230 | 1.000 | 1697.5 | 0.421 | 0.679 | 989 | 0.408 | 0.679 | 3155.5 | 0.415 | 0.679 |
| BRD-K09426783-300-01-0 | TAS-103 | 0.984 | 308 | 0.769 | 0.984 | 1407 | 0.777 | 0.984 | 1908 | 0.844 | 0.984 | 4960 | 0.988 | 0.988 | 2452 | 0.019 | 0.130 | 2743 | 0.359 | 0.984 | 4176 | 0.497 | 0.984 |
| BRD-K09443272-001-03-3 | PHA-680632 | 1.000 | 269 | 0.659 | 1.000 | 1555 | 0.880 | 1.000 | 1987 | 0.871 | 1.000 | 3815 | 0.306 | 0.642 | 1219 | 0.000 | 0.004 | 2412 | 0.205 | 0.537 | 3451 | 0.035 | 0.189 |
| BRD-K09549677-300-03-4 | mibefradil | 0.375 | | | | 93 | 0.880 | 0.938 | 236 | 0.904 | 0.938 | 387 | 0.580 | 0.938 | 408 | 0.874 | 0.938 | 226 | 0.554 | 0.938 | 283 | 0.576 | 0.938 |
| BRD-K09877029-001-01-2 | VS-4718 | 0.685 | | | | 179.5 | 0.926 | 0.972 | 307 | 0.567 | 0.880 | 866 | 0.197 | 0.516 | 912.5 | 0.841 | 0.972 | 313 | 0.092 | 0.355 | 886.5 | 0.121 | 0.364 |
| BRD-K09951645-001-06-8 | dabrafenib | 1.000 | | | | 235.5 | 0.466 | 1.000 | 126 | 0.077 | 0.626 | 863.5 | 0.729 | 1.000 | 515 | 0.089 | 0.626 | 565.5 | 0.944 | 1.000 | 1098 | 0.951 | 1.000 |
| BRD-K10843433-001-22-7 | phenylbutazone | 0.847 | 30 | 0.242 | 0.847 | 39 | 0.345 | 0.879 | 147.5 | 0.473 | 0.879 | 177 | 0.693 | 0.924 | 152 | 0.024 | 0.255 | 111 | 0.589 | 0.924 | 325 | 0.882 | 0.924 |

Suppl. Table 9

| broad_id | name | colorectal_MWU_STAT | colorectal_PVAL | colorectal_PVAL_ADJ | skin_MWU_STAT | skin_PVAL | skin_PVAL_ADJ | urinary_tract_MWU_STAT | urinary_tract_PVAL | urinary_tract_PVAL_ADJ | lung_MWU_STAT | lung_PVAL | lung_PVAL_ADJ | ovary_MWU_STAT | ovary_PVAL | ovary_PVAL_ADJ | breast_MWU_STAT | breast_PVAL | breast_PVAL_ADJ | pancreas_MWU_STAT | pancreas_PVAL | pancreas_PVAL_ADJ | nervous_system_MWU_STAT | central_nervous_system_PVAL | nervous_system_PVAL_ADJ | soft_tissue_MWU_STAT | soft_tissue_PVAL | soft_tissue_PVAL_ADJ | kidney_MWU_STAT | kidney_PVAL | kidney_PVAL_ADJ | gastric_MWU_STAT | gastric_PVAL | gastric_PVAL_ADJ | bone_MWU_STAT | bone_PVAL | bone_PVAL_ADJ | prostate_MWU_STAT | prostate_PVAL | prostate_PVAL_ADJ | al_nervous_system_MWU_STAT | peripheral_nervous_system_PVAL |
|---|---|---|---|---|---|---|---|---|---|---|---|---|---|---|---|---|---|---|---|---|---|---|---|---|---|---|---|---|---|---|---|---|---|---|---|---|---|---|---|---|---|---|
| BRD-K11071038-001-01-7 | ICI-162846 | 159 | 0.647 | 0.978 | 450 | 0.738 | 0.816 | 450 | 0.738 | 0.816 | 883 | 0.082 | 0.864 | 444 | 0.158 | 0.978 | 210 | 0.666 | 0.978 | 357 | 0.713 | 0.816 | 210 | 0.666 | 0.978 | 107 | 0.625 | 0.978 | 210 | 0.666 | 0.978 | 309 | 0.699 | 0.978 | 107 | 0.625 | 0.978 | 107 | 0.625 | 0.978 | | |
| BRD-K11073688-001-01-6 | CPI-1189 | 448 | 0.078 | 0.824 | 880 | 0.675 | 0.993 | 826 | 0.852 | 0.993 | 2116 | 0.371 | 0.811 | 426 | 0.272 | 0.811 | 277 | 0.451 | 0.811 | 400 | 0.402 | 0.811 | 724 | 0.416 | 0.811 | 292 | 0.846 | 0.993 | 193 | 0.041 | 0.214 | 252 | 0.132 | 0.556 | 264 | 0.739 | 0.993 | 153 | 0.550 | 0.888 | 195 | 0.795 |
| BRD-K11196887-001-22-0 | norfloxacin | 565 | 0.218 | 1.000 | 1283 | 0.957 | 0.957 | 804 | 0.627 | 0.885 | 2868 | 0.843 | 0.885 | 1012 | 0.042 | 0.219 | 649 | 0.426 | 0.745 | 1129 | 0.792 | 0.885 | 750 | 0.701 | 0.885 | 90 | 0.026 | 0.184 | 473 | 0.650 | 0.885 | 715 | 0.224 | 0.745 | 234 | 0.136 | 0.569 | 148 | 0.357 | 0.745 | 227 | 0.352 |
| BRD-K11215326-001-03-7 | tazemetostat | 477 | 0.162 | 0.954 | 1592 | 0.580 | 0.954 | 891.5 | 0.578 | 0.954 | 2643.5 | 0.446 | 0.954 | 1444 | 0.837 | 0.954 | 216 | 0.283 | 0.954 | 1037 | 0.864 | 0.954 | 1308 | 0.325 | 0.954 | 349 | 0.080 | 0.837 | 283 | 0.243 | 0.954 | 735.5 | 0.817 | 0.954 | 192.5 | 0.202 | 0.954 | 133 | 0.794 | 0.954 | 279 | 0.548 |
| BRD-K11267252-001-04-4 | alectinib | 1136 | 0.709 | 0.940 | 3185 | 0.612 | 0.940 | 1781 | 0.858 | 0.940 | 5283.5 | 0.340 | 0.843 | 2432 | 0.640 | 0.940 | 1488 | 0.828 | 0.940 | 2416.5 | 0.102 | 0.536 | 2152 | 0.580 | 0.940 | 804 | 0.322 | 0.843 | 1063 | 0.361 | 0.843 | 1300 | 0.979 | 0.979 | 435 | 0.034 | 0.368 | 353 | 0.806 | 0.940 | 244 | 0.035 |
| BRD-K11433652-001-17-0 | aspirin | 154 | 0.185 | 0.870 | 130 | 0.099 | 0.870 | 38 | 0.060 | 0.870 | 1023 | 0.221 | 0.870 | 406 | 0.176 | 0.870 | 99 | 0.926 | 0.955 | 420 | 0.784 | 0.955 | 707 | 0.955 | 0.955 | 5 | 0.011 | 0.025 | 170 | 0.585 | 0.955 | 243 | 0.706 | 0.955 | 239 | 0.683 | 0.955 | 161 | 0.891 | 0.955 | 73 | 0.733 |
| BRD-K11510141-001-01-8 | teprenone | 482 | 0.048 | 1.000 | 1682.5 | 0.552 | 0.831 | 1043.5 | 0.594 | 0.831 | 2902.5 | 0.309 | 0.811 | 1178 | 0.134 | 0.561 | 625 | 0.230 | 0.689 | 1569 | 0.942 | 0.942 | 1516 | 0.905 | 0.942 | 654 | 0.756 | 0.882 | 632.5 | 0.451 | 0.831 | 801.5 | 0.656 | 0.831 | 207 | 0.210 | 0.689 | 25 | 0.103 | 0.540 | 174 | 0.417 |
| BRD-K11630072-001-13-2 | carmofur | 618.5 | 0.316 | 0.603 | 1175.5 | 0.247 | 0.576 | 729 | 0.225 | 0.576 | 3068.5 | 0.867 | 0.981 | 516 | 0.007 | 0.035 | 1084 | 0.981 | 0.981 | 1185 | 0.937 | 0.981 | 918 | 0.836 | 0.981 | 576 | 0.219 | 0.576 | 742.5 | 0.431 | 0.754 | 521.5 | 0.124 | 0.521 | 214 | 0.284 | 0.596 | 239 | 0.800 | 0.981 | 149 | 0.876 |
| BRD-K11636097-001-03-1 | JNJ-7706621 | 2388 | 0.551 | 0.999 | 4379 | 0.902 | 0.999 | 3230 | 0.952 | 0.999 | 6277 | 0.014 | 0.297 | 2628 | 0.055 | 0.440 | 2705 | 0.999 | 0.999 | 3387 | 0.953 | 0.999 | 3661 | 0.717 | 0.999 | 640 | 0.111 | 0.440 | 1548 | 0.576 | 0.999 | 1411 | 0.382 | 0.891 | 595 | 0.076 | 0.440 | 439 | 0.837 | 0.999 | 415 | 0.143 |
| BRD-K11663430-001-02-3 | pyroxamide | 1025 | 0.127 | 0.885 | 4186 | 0.996 | 0.996 | 1649 | 0.253 | 0.885 | 5455 | 0.789 | 0.873 | 1322 | 0.032 | 0.662 | 603 | 0.107 | 0.885 | 1645.5 | 0.545 | 0.817 | 2154 | 0.779 | 0.873 | 946 | 0.497 | 0.803 | 708 | 0.250 | 0.556 | 1247 | 0.265 | 0.556 | 379 | 0.080 | 0.210 | 441 | 0.740 | 0.873 | 95 | 0.354 |
| BRD-K11851476-003-01-7 | pardoprunox | 3273 | 0.716 | 0.998 | 6202 | 0.900 | 0.998 | 2198 | 0.013 | 0.089 | 11674 | 0.259 | 0.776 | 4606 | 0.794 | 0.998 | 2295 | 0.092 | 0.386 | 4669 | 0.901 | 0.998 | 4926 | 0.783 | 0.998 | 1552 | 0.218 | 0.761 | 2284 | 0.998 | 0.998 | 1333 | 0.012 | 0.089 | 2265 | 0.782 | 0.998 | 1195 | 0.985 | 0.998 | 819 | 0.643 |
| BRD-K11927976-050-02-9 | ER-27319 | 3490 | 0.120 | 0.431 | 7602 | 0.825 | 0.900 | 3843 | 0.578 | 0.900 | 15430 | 0.872 | 0.900 | 4060 | 0.050 | 0.362 | 4779 | 0.810 | 0.900 | 5775 | 0.631 | 0.900 | 5711 | 0.701 | 0.900 | 1582 | 0.123 | 0.431 | 2913 | 0.770 | 0.900 | 2135 | 0.031 | 0.362 | 1122 | 0.069 | 0.364 | 1000 | 0.771 | 0.900 | 602 | 0.189 |
| BRD-K11973162-003-01-8 | tipiracil | 1759 | 0.088 | 0.920 | 3690.5 | 0.086 | 0.920 | 1858 | 0.251 | 0.586 | 9607 | 0.933 | 0.950 | 2706 | 0.300 | 0.604 | 1741 | 0.432 | 0.697 | 3551 | 0.950 | 0.950 | 3849.5 | 0.629 | 0.778 | 821 | 0.612 | 0.778 | 2195 | 0.906 | 0.950 | 1714 | 0.242 | 0.586 | 38 | 0.016 | 0.086 | 153 | 0.110 | 0.454 | 314 | 0.171 |
| BRD-K12068470-001-02-5 | LY2608204 | 1651 | 0.412 | 0.666 | 1420 | 0.002 | 0.033 | 1391 | 0.337 | 1.000 | 5276 | 0.592 | 0.888 | 1896 | 0.984 | 0.984 | 1531 | 0.249 | 1.000 | 2055 | 0.811 | 0.984 | 2910 | 0.970 | 0.984 | 398 | 0.291 | 0.666 | 679 | 0.216 | 0.567 | 488 | 0.252 | 1.000 | 207 | 0.371 | 0.666 | 235 | 0.971 | 0.984 | 665 | 0.670 |
| BRD-K12184916-001-15-4 | NVP-BEZ235 | 2321 | 0.770 | 0.967 | 4703 | 0.974 | 0.981 | 2095 | 0.691 | 0.967 | 8310.5 | 0.783 | 0.967 | 2673 | 0.272 | 0.683 | 2305 | 0.185 | 0.556 | 2751 | 0.621 | 0.967 | 3629.5 | 0.551 | 0.964 | 1166 | 0.876 | 0.981 | 2042 | 0.456 | 0.899 | 1063 | 0.057 | 0.427 | 1072 | 0.061 | 0.427 | 157 | 0.116 | 0.556 | 782 | 0.981 |
| BRD-K12184916-001-19-6 | NVP-BEZ235 | 671 | 0.998 | 0.998 | 1798.5 | 0.936 | 0.983 | 438 | 0.223 | 1.000 | 2149.5 | 0.646 | 1.000 | 652 | 0.143 | 1.000 | 772 | 0.778 | 0.908 | 500 | 0.879 | 0.972 | 692.5 | 0.770 | 0.908 | 224 | 0.005 | 0.012 | 413 | 0.621 | 0.908 | 578.5 | 0.665 | 0.908 | 274 | 0.355 | 0.678 | 97 | 0.662 | 0.908 | 139 | 0.399 |
| BRD-K12244279-001-05-8 | MEK1-2-inhibitor | 183 | 0.012 | 0.261 | 756 | 0.137 | 0.843 | 504 | 0.459 | 1.000 | 1934 | 0.933 | 0.960 | 634 | 0.498 | 1.000 | 493 | 0.851 | 0.940 | 534 | 0.203 | 0.852 | 633 | 0.960 | 0.960 | 297 | 0.651 | 0.912 | 104 | 0.766 | 0.913 | 584 | 0.732 | 0.913 | 181 | 0.783 | 0.913 | | | | 61 | 0.106 |
| BRD-K12251893-065-04-7 | vincristine | 4316.5 | 0.977 | 0.999 | 4030 | 0.263 | 0.553 | 1893 | 0.159 | 0.418 | 11457 | 0.604 | 0.846 | 4174.5 | 0.261 | 0.553 | 3377.5 | 0.750 | 0.984 | 5349.5 | 0.996 | 0.999 | 3697 | 0.348 | 0.664 | 1327 | 0.094 | 0.283 | 2814.5 | 0.999 | 0.999 | 1431 | 0.012 | 0.224 | 1000 | 0.079 | 0.278 | 574 | 0.583 | 0.846 | 86 | 0.032 |
| BRD-K12343256-001-08-9 | trametinib | 2614 | 0.002 | 0.021 | 2001 | 0.000 | 0.000 | 3189 | 0.321 | 0.674 | 12190 | 0.993 | 0.999 | 4761.5 | 0.931 | 0.999 | 3196 | 0.979 | 0.999 | 3471 | 0.007 | 0.024 | 4557.5 | 0.999 | 0.999 | 1070 | 0.273 | 0.636 | 2808 | 0.996 | 0.999 | 1861 | 0.071 | 0.213 | 1039.5 | 0.943 | 0.999 | 873.5 | 0.970 | 0.999 | 742 | 0.265 |
| BRD-K12343256-001-14-7 | trametinib | 972 | 0.000 | 0.002 | 1540 | 0.000 | 0.000 | 1175 | 0.060 | 0.313 | 7219 | 1.000 | 1.000 | 1268 | 0.911 | 1.000 | 2057 | 0.991 | 1.000 | 2261 | 0.064 | 0.193 | 1528 | 0.972 | 1.000 | 1068 | 0.645 | 1.000 | 1447 | 0.939 | 1.000 | 1094 | 0.277 | 0.727 | 690 | 0.993 | 1.000 | 662 | 0.987 | 1.000 | 338 | 0.373 |
| BRD-K12502280-001-08-0 | TG-101348 | 568 | 0.357 | 1.000 | 1189 | 0.913 | 0.972 | 925 | 0.736 | 0.967 | 2051 | 0.621 | 0.932 | 802 | 0.042 | 0.145 | 400 | 0.275 | 0.721 | 671 | 0.856 | 0.972 | 1116 | 0.972 | 0.972 | 95 | 0.009 | 0.045 | 783 | 0.565 | 0.913 | 374 | 0.431 | 0.913 | 254 | 0.091 | 0.272 | 215 | 0.800 | 0.972 | 284 | 0.716 |
| BRD-K12539581-001-23-7 | nocodazole | 5244 | 0.684 | 0.993 | 8449 | 0.976 | 0.993 | 3771 | 0.085 | 0.355 | 15589 | 0.353 | 0.742 | 4830 | 0.083 | 0.355 | 4979 | 0.861 | 0.993 | 6987 | 0.974 | 0.993 | 7531 | 0.966 | 0.993 | 2090 | 0.271 | 0.711 | 4354 | 0.993 | 0.993 | 2760 | 0.504 | 0.882 | 2133 | 0.167 | 0.502 | 1036 | 0.744 | 0.993 | 552 | 0.029 |
| BRD-K12787259-001-04-3 | CX-5461 | 2467.5 | 0.837 | 0.977 | 3907 | 0.783 | 0.968 | 2168.5 | 0.241 | 0.506 | 7215.5 | 0.232 | 0.506 | 2815 | 0.645 | 0.968 | 1566 | 0.160 | 0.506 | 3952 | 0.996 | 1.000 | 3473.5 | 0.755 | 0.968 | 1311 | 0.496 | 0.868 | 1809 | 1.000 | 1.000 | 1461 | 0.043 | 0.447 | 55 | 0.138 | 0.506 | 366 | 0.296 | 0.565 | 850 | 0.931 |
| BRD-K12867552-001-04-7 | PCI-24781 | 2674 | 0.053 | 0.251 | 5164 | 0.249 | 0.522 | 3607 | 0.649 | 0.916 | 11430 | 0.605 | 0.916 | 3332 | 0.036 | 0.251 | 3546 | 0.930 | 0.999 | 5472 | 0.974 | 0.999 | 6321 | 0.999 | 0.999 | 1631 | 0.454 | 0.794 | 3680 | 0.988 | 0.999 | 2242 | 0.060 | 0.251 | 1158 | 0.058 | 0.251 | 478 | 0.110 | 0.325 | 532 | 0.049 |
| BRD-K12885236-001-02-7 | dapivirine | 1560 | 0.724 | 0.989 | 3396 | 0.971 | 0.989 | 2703 | 0.867 | 0.989 | 6211.5 | 0.066 | 0.316 | 2497 | 0.066 | 0.316 | 1517 | 0.467 | 0.892 | 4108.5 | 0.989 | 0.989 | 3667 | 0.969 | 0.989 | 462 | 0.088 | 0.316 | 1135 | 0.915 | 0.989 | 1413 | 0.181 | 0.522 | 512 | 0.047 | 0.316 | 540 | 0.773 | 0.989 | 399 | 0.824 |
| BRD-K13044802-213-20-7 | ciclopirox | 1311 | 0.458 | 0.802 | 2900.5 | 0.660 | 0.955 | 1060.5 | 0.025 | 0.524 | 6270 | 0.779 | 0.968 | 1786 | 0.130 | 0.457 | 1683.5 | 0.979 | 0.979 | 1806 | 0.682 | 0.955 | 2118.5 | 0.242 | 0.638 | 332 | 0.364 | 0.802 | 951 | 0.437 | 0.802 | 839 | 0.112 | 1.000 | 239.5 | 0.939 | 0.979 | 53 | 0.027 | 0.115 | 288 | 0.243 |
| BRD-K13049116-001-04-0 | BMS-754807 | 568 | 0.336 | 0.863 | 825 | 0.823 | 0.993 | 627 | 0.079 | 0.416 | 2123 | 0.065 | 0.416 | 954 | 0.772 | 0.993 | 820 | 0.934 | 0.993 | 321 | 0.010 | 0.217 | 1131 | 0.972 | 0.993 | 635 | 0.758 | 0.993 | 852 | 0.993 | 0.993 | 472 | 0.764 | 0.993 | 214 | 0.370 | 0.863 | 150 | 0.928 | 0.993 | 81 | 0.113 |
| BRD-K13049116-001-05-7 | BMS-754807 | 1345.5 | 0.190 | 0.884 | 2093 | 0.738 | 0.990 | 1167 | 0.610 | 0.990 | 4436 | 0.003 | 0.070 | 2326 | 0.849 | 0.990 | 2312 | 0.990 | 0.990 | 2641.5 | 0.353 | 0.961 | 1827.5 | 0.540 | 0.990 | 799.5 | 0.362 | 0.961 | 951 | 0.874 | 0.990 | 822.5 | 0.211 | 0.884 | 418 | 0.033 | 0.229 | 237.5 | 0.934 | 0.990 | 243 | 0.010 |
| BRD-K13050541-001-01-9 | acivicin | 1223 | 0.433 | 1.000 | 1734 | 0.953 | 0.976 | 1323 | 0.615 | 1.000 | 4247 | 0.286 | 1.000 | 1775 | 0.552 | 1.000 | 917 | 0.057 | 0.425 | 1180 | 0.061 | 0.425 | 1645 | 0.651 | 1.000 | 511 | 0.717 | 1.000 | 865 | 0.746 | 0.976 | 520 | 0.739 | 0.976 | 284 | 0.034 | 0.060 | 363 | 0.938 | 0.976 | 373 | 0.650 |
| BRD-K13060017-001-01-3 | UNC0631 | 4703 | 0.897 | 0.999 | 3737 | 0.000 | 0.000 | 3629 | 0.326 | 0.790 | 15452 | 0.916 | 0.999 | 4365 | 0.604 | 0.914 | 5410 | 0.999 | 0.999 | 5462 | 0.612 | 0.914 | 5290 | 0.203 | 0.790 | 1198 | 0.046 | 0.344 | 3970 | 0.990 | 0.999 | 3208 | 0.732 | 0.961 | 1351 | 0.236 | 0.790 | 416 | 0.049 | 0.344 | 991 | 0.484 |
| BRD-K13087974-001-02-5 | CGP-52411 | 1023 | 0.947 | 0.947 | 1666.5 | 0.164 | 0.690 | 824 | 0.239 | 0.755 | 4040.5 | 0.628 | 0.903 | 1455 | 0.499 | 0.903 | 1599 | 0.731 | 0.903 | 2117.5 | 0.930 | 1.000 | 1709 | 0.314 | 0.755 | 113 | 0.119 | 0.690 | 888 | 0.939 | 1.000 | 716 | 0.100 | 0.690 | 276 | 0.319 | 0.755 | 89 | 0.074 | 0.690 | 187 | 0.360 |
| BRD-K13094524-001-07-5 | PFI-1 | 2867 | 0.616 | 0.786 | 4232 | 0.463 | 0.767 | 1941 | 0.253 | 1.000 | 9066.5 | 0.749 | 0.873 | 3489 | 0.291 | 0.654 | 2591 | 0.636 | 0.786 | 2836 | 0.308 | 0.654 | 4296 | 0.973 | 0.973 | 1273 | 0.302 | 0.654 | 1847 | 0.940 | 0.973 | 1536 | 0.071 | 1.000 | 1041 | 0.391 | 0.746 | 548 | 0.312 | 0.654 | 489 | 0.209 |
| BRD-K13112821-001-17-4 | artemisinin | 375 | 0.598 | 0.757 | 1072.5 | 0.721 | 0.757 | 328 | 0.073 | 0.621 | 1902 | 0.718 | 0.757 | 524 | 0.227 | 0.621 | 101 | 0.237 | 0.621 | 710 | 0.594 | 0.757 | 679 | 0.498 | 0.757 | 100 | 0.231 | 0.621 | 399 | 0.694 | 0.757 | 280 | 0.219 | 0.621 | 17 | 0.016 | 0.344 | 16 | 0.091 | 0.621 | 185 | 0.344 |
| BRD-K13167095-001-02-9 | AR-42 | 3725 | 0.779 | 0.962 | 6021 | 0.747 | 0.962 | 2439 | 0.596 | 0.894 | 10799 | 0.190 | 0.505 | 3908 | 0.192 | 0.505 | 3869 | 0.972 | 0.990 | 5333 | 0.987 | 0.990 | 6121 | 0.990 | 0.990 | 1317 | 0.155 | 0.505 | 2875 | 0.916 | 0.990 | 1916 | 0.064 | 0.270 | 835 | 0.057 | 0.270 | 441 | 0.279 | 0.597 | 547 | 0.060 |
| BRD-K13169950-001-14-4 | NSC-3852 | 515 | 0.175 | 0.472 | 914 | 0.180 | 0.472 | 302 | 0.247 | 0.519 | 3540 | 0.629 | 0.943 | 1158 | 0.114 | 0.400 | 1072 | 0.984 | 0.984 | 1261 | 0.926 | 0.984 | 1832 | 0.937 | 0.984 | 292 | 0.075 | 0.315 | 628 | 0.438 | 0.708 | 536 | 0.428 | 0.708 | 139 | 0.065 | 0.315 | 153 | 0.849 | 0.984 | 66 | 0.057 |
| BRD-K13183738-317-10-2 | pentamidine | 672 | 0.976 | 0.976 | 658 | 0.115 | 1.000 | 428 | 0.608 | 1.000 | 1379 | 0.060 | 1.000 | 399 | 0.264 | 1.000 | 603 | 0.751 | 0.928 | 826 | 0.668 | 0.928 | 695 | 0.457 | 0.799 | 162 | 0.268 | 0.625 | 209 | 0.523 | 0.845 | 183 | 0.376 | 1.000 | 320 | 0.951 | 0.976 | 64 | 0.451 | 0.799 | 117 | 0.359 |
| BRD-K13387373-004-14-5 | thonzonium | 3076 | 0.260 | 0.779 | 5604 | 0.590 | 0.968 | 2693 | 0.343 | 0.902 | 12700 | 0.834 | 0.968 | 3906 | 0.191 | 0.739 | 3355 | 0.877 | 0.968 | 4722 | 0.755 | 0.968 | 5131 | 0.796 | 0.968 | 643 | 0.003 | 0.027 | 2279 | 0.431 | 0.968 | 2201 | 0.211 | 0.739 | 1200 | 0.640 | 0.968 | 769 | 0.866 | 0.968 | 880 | 0.736 |
| BRD-K13390322-001-06-3 | AT-7519 | 3411 | 0.469 | 0.984 | 7312.5 | 0.995 | 1.000 | 2271 | 0.006 | 0.064 | 12114 | 0.248 | 0.744 | 3435 | 0.089 | 0.465 | 3686 | 0.905 | 1.000 | 5384 | 0.777 | 1.000 | 3948 | 0.363 | 0.847 | 1563 | 0.195 | 0.682 | 3243 | 1.000 | 1.000 | 1864 | 0.086 | 0.465 | 2552.5 | 0.922 | 1.000 | 844 | 0.926 | 1.000 | 447.5 | 0.669 |
| BRD-K13409143-001-01-9 | isopropyl-myristate | 703 | 0.779 | 0.932 | 1516 | 0.255 | 0.701 | 812 | 0.096 | 0.701 | 3468.5 | 0.442 | 0.888 | 1253 | 0.843 | 0.932 | 918 | 0.932 | 0.932 | 1299 | 0.894 | 0.932 | 1602.5 | 0.800 | 0.932 | 519 | 0.301 | 0.701 | 586 | 0.251 | 0.701 | 524 | 0.857 | 0.932 | 387 | 0.199 | 0.701 | 203 | 0.507 | 0.888 | 44 | 0.029 |
| BRD-K13514097-001-05-3 | everolimus | 1031 | 0.998 | 0.998 | 1311 | 0.711 | 0.978 | 750 | 0.217 | 1.000 | 3192 | 0.932 | 0.978 | 1131 | 0.225 | 0.429 | 371 | 0.853 | 0.978 | 1171 | 0.760 | 0.978 | 1196 | 0.213 | 0.429 | 65 | 0.012 | 0.050 | 543 | 0.778 | 0.978 | 486 | 0.340 | 1.000 | 561 | 0.156 | 0.364 | 134 | 0.065 | 0.214 | 33 | 0.136 |
| BRD-K13566078-001-02-3 | BMS-345541 | 730 | 0.981 | 0.981 | 1390.5 | 0.439 | 0.838 | 662 | 0.300 | 0.700 | 3355 | 0.954 | 1.000 | 1067.5 | 0.171 | 0.513 | 1024 | 0.727 | 1.000 | 1388.5 | 0.862 | 1.000 | 1604 | 0.284 | 0.700 | 160 | 0.024 | 0.395 | 829.5 | 0.710 | 1.000 | 669.5 | 0.542 | 0.949 | 203.5 | 0.056 | 0.395 | | | | 38 | 0.150 |
| BRD-K13571841-001-14-4 | pepstatin | 745 | 0.138 | 0.711 | 1752 | 0.911 | 0.953 | 773 | 0.173 | 0.711 | 4261.5 | 0.177 | 0.711 | 1428 | 0.203 | 0.711 | 1358.5 | 0.942 | 0.953 | 1114 | 0.432 | 0.881 | 1606 | 0.953 | 0.953 | 727 | 0.711 | 0.953 | 1128.5 | 0.656 | 0.953 | 834 | 0.470 | 0.881 | 326 | 0.504 | 0.881 | 448.5 | 0.870 | 0.953 | 133 | 0.040 |
| BRD-K13606314-001-01-3 | P5091 | 1518 | 0.841 | 0.930 | 2076 | 0.368 | 0.900 | 1362 | 0.285 | 0.900 | 4184 | 0.312 | 0.900 | 1663 | 0.756 | 0.900 | 1120 | 0.559 | 0.900 | 1447 | 0.093 | 0.900 | 1569 | 0.772 | 0.900 | 522 | 0.405 | 0.900 | 875 | 0.724 | 0.900 | 1163 | 0.641 | 0.900 | 182 | 0.320 | 0.900 | | | | 206 | 0.124 |
| BRD-K13646352-001-03-8 | midostaurin | 6412 | 0.970 | 0.992 | 8665 | 0.991 | 0.992 | 4712 | 0.548 | 0.923 | 16072 | 0.410 | 0.862 | 5159 | 0.156 | 0.655 | 4833 | 0.927 | 0.992 | 8492 | 0.992 | 0.992 | 5257 | 0.028 | 0.148 | 1703 | 0.021 | 0.148 | 1959 | 0.002 | 0.027 | 2904 | 0.261 | 0.750 | 1206 | 0.003 | 0.027 | 894 | 0.846 | 0.992 | 931 | 0.286 |
| BRD-K13662825-001-07-5 | dinaciclib | 6749 | 0.999 | 1.000 | 7445 | 0.695 | 1.000 | 2963 | 0.012 | 0.125 | 13615 | 0.163 | 0.490 | 4991 | 0.537 | 0.939 | 4285 | 0.821 | 1.000 | 8083 | 1.000 | 1.000 | 6243 | 0.819 | 1.000 | 1707 | 0.086 | 0.302 | 3438 | 0.909 | 1.000 | 2780 | 0.289 | 0.758 | 2212 | 0.463 | 0.939 | 924 | 0.919 | 1.000 | 373 | 0.028 |
| BRD-K13756951-001-01-4 | bitopertin | 1647 | 0.591 | 0.913 | 2504.5 | 0.497 | 0.913 | 1549 | 0.609 | 0.913 | 5269 | 0.415 | 0.913 | 1956 | 0.841 | 0.922 | 1190 | 0.762 | 0.922 | 1327 | 0.264 | 0.913 | 1708 | 0.922 | 0.922 | 550 | 0.070 | 0.913 | 505 | 0.292 | 0.913 | 1095 | 0.391 | 0.913 | 534 | 0.360 | 0.913 | 374 | 0.916 | 0.922 | 258.5 | 0.581 |
| BRD-K13810148-311-03-2 | givinostat | 4069 | 0.190 | 0.388 | 7862 | 0.865 | 1.000 | 4480 | 0.440 | 0.660 | 12899 | 0.136 | 0.371 | 4173 | 0.244 | 0.427 | 4916 | 0.958 | 1.000 | 8078 | 1.000 | 1.000 | 8230 | 1.000 | 1.000 | 1704 | 0.085 | 0.357 | 3733 | 0.977 | 1.000 | 2697 | 0.131 | 0.371 | 950 | 0.006 | 0.132 | 313 | 0.014 | 0.147 | 637 | 0.071 |
| BRD-K13949769-001-01-7 | tenovin-6 | 2711 | 0.191 | 0.555 | 5634 | 0.983 | 0.999 | 2417 | 0.108 | 0.531 | 9783.5 | 0.411 | 0.780 | 2873 | 0.204 | 0.555 | 3799.5 | 0.995 | 0.999 | 3476 | 0.329 | 0.691 | 3807 | 0.483 | 0.780 | 1072 | 0.071 | 0.499 | 3297 | 0.999 | 0.999 | 1427 | 0.049 | 0.499 | 668 | 0.211 | 0.555 | 394 | 0.250 | 0.583 | 314 | 0.126 |
| BRD-K14109347-001-03-4 | LY2603618 | 1486 | 0.651 | 0.911 | 1441 | 0.412 | 0.722 | 1051 | 0.059 | 0.622 | 4141 | 0.773 | 0.983 | 1383 | 0.471 | 0.760 | 1654.5 | 0.873 | 0.983 | 1681 | 0.896 | 0.983 | 1281.5 | 0.299 | 0.658 | 135 | 0.189 | 0.658 | 453.5 | 0.308 | 0.658 | 471.5 | 0.020 | 0.419 | 445.5 | 0.584 | 0.876 | 539 | 0.982 | 0.983 | 294 | 0.152 |
| BRD-K14560436-001-01-4 | TG-02 | 3743 | 0.062 | 0.519 | 7681 | 0.760 | 0.949 | 3801 | 0.458 | 0.873 | 15632 | 0.574 | 0.873 | 5811 | 0.431 | 0.873 | 5101 | 0.941 | 0.969 | 5533 | 0.273 | 0.819 | 7335 | 0.969 | 0.969 | 1448 | 0.051 | 0.519 | 3171 | 0.582 | 0.873 | 1935 | 0.098 | 0.519 | 1751 | 0.371 | 0.873 | 1066 | 0.814 | 0.949 | 552 | 0.123 |
| BRD-K14618467-001-04-6 | IKK-16 | 1209 | 0.003 | 0.070 | 3766 | 0.276 | 0.984 | 2185 | 0.482 | 0.984 | 7228 | 0.747 | 0.984 | 1592 | 0.014 | 0.100 | 1985 | 0.560 | 0.984 | 3104 | 0.861 | 0.984 | 2571 | 0.486 | 0.984 | 737 | 0.125 | 0.655 | 1675 | 0.502 | 0.984 | 1239 | 0.674 | 0.984 | 567 | 0.485 | 0.984 | 178 | 0.615 | 0.984 | 574 | 0.829 |
| BRD-K14796088-003-26-8 | berberine | 3879 | 0.923 | 0.928 | 5047 | 0.560 | 0.928 | 4169 | 0.902 | 0.928 | 11082 | 0.719 | 0.928 | 4026 | 0.261 | 0.913 | 1860 | 0.014 | 0.096 | 5271 | 0.928 | 0.928 | 5057 | 0.915 | 0.928 | 896 | 0.013 | 0.096 | 1972 | 0.748 | 0.928 | 1554 | 0.028 | 0.146 | 629 | 0.347 | 0.928 | 589 | 0.608 | 0.928 | 343 | 0.458 |
| BRD-K14991967-001-02-6 | GSK650394 | 842 | 0.427 | 0.815 | 2867 | 0.982 | 0.982 | 1726 | 0.560 | 0.892 | 4524 | 0.656 | 0.892 | 1736 | 0.191 | 0.501 | 370 | 0.102 | 0.306 | 1751 | 0.981 | 0.982 | 1981 | 0.877 | 0.982 | 156 | 0.013 | 0.055 | 849 | 0.679 | 0.892 | 597 | 0.008 | 0.171 | 356 | 0.246 | 0.574 | 143 | 0.679 | 0.892 | 116 | 0.521 |
| BRD-K15025317-001-15-1 | BAY-11-7082 | 745 | 0.016 | 0.342 | 3267 | 0.978 | 0.984 | 1593 | 0.984 | 0.984 | 5406 | 0.606 | 0.978 | 2248 | 0.341 | 0.716 | 1144 | 0.543 | 0.951 | 2784 | 0.817 | 0.978 | 2128 | 0.081 | 0.244 | 589 | 0.400 | 0.764 | 1337 | 0.838 | 0.978 | 1009 | 0.307 | 0.716 | 74 | 0.044 | 0.152 | 329 | 0.745 | 0.978 | 101 | 0.011 |
| BRD-K15108141-001-06-6 | gemcitabine | 2528 | 0.165 | 0.624 | 4862 | 0.926 | 0.994 | 1312 | 0.037 | 0.624 | 8920 | 0.178 | 0.624 | 2512 | 0.382 | 0.775 | 3159 | 0.990 | 0.994 | 5208 | 0.994 | 0.994 | 2592 | 0.072 | 0.624 | 704 | 0.324 | 0.759 | 1506 | 0.119 | 0.624 | 1081 | 0.445 | 0.775 | 1150 | 0.325 | 0.759 | 453 | 0.161 | 0.624 | 503 | 0.544 |
| BRD-K15179879-001-03-2 | carfilzomib | 5105 | 0.618 | 0.864 | 6715 | 0.045 | 0.473 | 4142 | 0.167 | 0.637 | 19619 | 0.989 | 0.999 | 5554 | 0.291 | 0.637 | 4774 | 0.672 | 0.864 | 7075 | 0.699 | 0.864 | 6386 | 0.239 | 0.637 | 1784 | 0.151 | 0.637 | 5015 | 0.999 | 0.999 | 2891 | 0.125 | 0.637 | 2039 | 0.552 | 0.864 | 755 | 0.303 | 0.637 | 949 | 0.582 |
| BRD-K15409150-001-05-8 | penfluridol | 4552 | 0.961 | 0.961 | 5493 | 0.360 | 0.630 | 2674 | 0.039 | 0.397 | 11632 | 0.246 | 0.607 | 3686 | 0.031 | 0.397 | 3817 | 0.848 | 0.961 | 5255 | 0.903 | 0.961 | 6014 | 0.914 | 0.961 | 1401 | 0.177 | 0.607 | 3700 | 0.948 | 0.961 | 1799 | 0.224 | 0.607 | 1495 | 0.260 | 0.607 | 680 | 0.350 | 0.630 | 663 | 0.323 |
| BRD-K15424032-001-03-6 | SB-228357 | 323 | 0.175 | 0.408 | 531.5 | 0.998 | 0.998 | 434 | 0.639 | 0.823 | 1139 | 0.666 | 0.823 | 455.5 | 0.526 | 0.790 | 200 | 0.779 | 0.909 | 322 | 0.571 | 0.800 | 172 | 0.273 | 0.573 | 263.5 | 0.523 | 0.790 | 583.5 | 0.930 | 0.980 | 123 | 0.024 | 0.504 | | | | 54 | 0.506 | 0.790 | 173 | 0.933 |
| BRD-K15567136-003-27-2 | papaverine | 1423 | 0.718 | 0.887 | 3177 | 0.918 | 0.962 | 1466 | 0.404 | 1.000 | 5920 | 0.510 | 1.000 | 2281 | 0.285 | 1.000 | 2132 | 0.962 | 0.962 | 2455 | 0.483 | 1.000 | 2957 | 0.929 | 0.962 | 464 | 0.004 | 0.083 | 568 | 0.609 | 1.000 | 1236 | 0.630 | 1.000 | 540 | 0.097 | 0.170 | 376 | 0.442 | 0.663 | 138 | 0.122 |
| BRD-K15600710-066-04-1 | obatoclax | 2664 | 0.567 | 0.870 | 4773 | 0.692 | 0.870 | 2241 | 0.754 | 0.870 | 9747 | 0.942 | 0.942 | 3082 | 0.415 | 0.870 | 2322 | 0.229 | 0.870 | 2899 | 0.787 | 0.870 | 3880 | 0.841 | 0.883 | 1066 | 0.691 | 0.870 | 1869 | 0.195 | 0.870 | 2075 | 0.573 | 0.870 | 1006 | 0.345 | 0.870 | 316 | 0.486 | 0.870 | 667 | 0.561 |
| BRD-K15600710-066-05-8 | obatoclax | 2727.5 | 0.746 | 0.975 | 3479 | 0.098 | 0.410 | 2543 | 0.409 | 0.866 | 7737 | 0.757 | 0.975 | 3850.5 | 0.840 | 0.975 | 1579.5 | 0.413 | 0.866 | 4733.5 | 1.000 | 1.000 | 3016 | 0.928 | 0.975 | 1072 | 0.294 | 0.772 | 2354 | 0.910 | 0.975 | 981 | 0.031 | 0.220 | 1267.5 | 0.595 | 0.975 | 474 | 0.906 | 0.975 | 845 | 0.662 |
| BRD-K15898725-001-01-7 | maytansinol-isobutyrate | 1663 | 0.604 | 0.936 | 1215 | 0.669 | 0.936 | 1160 | 0.381 | 0.728 | 3110 | 0.219 | 0.574 | 1305 | 0.474 | 0.830 | 1065 | 0.909 | 0.955 | 2214 | 0.983 | 0.983 | 1653 | 0.201 | 0.574 | 335 | 0.085 | 0.574 | 1515 | 0.815 | 0.951 | 677 | 0.373 | 0.728 | 406 | 0.196 | 0.574 | 153 | 0.064 | 0.574 | 258 | 0.295 |
| BRD-K16136380-001-01-7 | pyrithione-zinc | 4874 | 0.954 | 0.975 | 6001 | 0.864 | 0.975 | 2879 | 0.278 | 0.649 | 12515 | 0.604 | 0.975 | 4090 | 0.050 | 0.362 | 3645 | 0.623 | 0.975 | 5765 | 0.947 | 0.975 | 4591 | 0.097 | 0.362 | 1752 | 0.562 | 0.975 | 2367 | 0.160 | 0.446 | 2115 | 0.056 | 0.362 | 1357 | 0.026 | 0.362 | 949 | 0.975 | 0.975 | 660 | 0.661 |
| BRD-K16180792-001-01-7 | bis(maltolato)oxovanadium(IV) | 554 | 0.996 | 0.999 | 873 | 0.000 | 0.003 | 600 | 0.695 | 1.000 | 1747 | 0.305 | 1.000 | 877 | 0.103 | 0.720 | 719 | 0.933 | 0.999 | 1103 | 0.999 | 0.999 | 1062 | 0.580 | 1.000 | 317 | 0.843 | 0.999 | 520 | 0.690 | 1.000 | 277 | 0.694 | 1.000 | 216 | 0.399 | 1.000 | 153 | 0.943 | 0.999 | 248 | 0.914 |
| BRD-K16189898-003-03-3 | CHIR-99021 | 108.5 | 0.490 | 0.910 | 306 | 0.430 | 1.000 | 228.5 | 0.086 | 0.907 | 1183 | 0.231 | 1.000 | 467 | 0.520 | 0.910 | 148 | 0.842 | 0.922 | 321 | 0.816 | 0.922 | 526 | 0.368 | 0.904 | 98.5 | 0.387 | 0.904 | 238.5 | 0.665 | 0.922 | 260.5 | 0.457 | 1.000 | | | | 74.5 | 0.763 | 0.922 | 9 | 0.046 |
| BRD-K16478699-001-09-2 | PLX-4720 | 2554 | 0.963 | 0.992 | 2012 | 0.000 | 0.000 | 2039.5 | 0.151 | 0.795 | 9601 | 0.697 | 0.992 | 3302 | 0.310 | 0.923 | 2160 | 0.547 | 0.992 | 3065.5 | 0.840 | 0.992 | 2829 | 0.643 | 0.992 | 1834.5 | 0.990 | 0.992 | 1847 | 0.992 | 0.992 | 1811 | 0.476 | 0.992 | 919.5 | 0.246 | 0.860 | 369 | 0.663 | 0.992 | 211 | 0.212 |
| BRD-K16551357-001-03-2 | evacetrapib | 773 | 0.110 | 0.460 | 1707 | 0.851 | 0.903 | 1500 | 0.932 | 0.932 | 4489 | 0.860 | 0.903 | 1003 | 0.497 | 0.869 | 1008 | 0.711 | 0.878 | 1502 | 0.557 | 0.878 | 1397 | 0.698 | 0.878 | 347 | 0.291 | 0.680 | 582 | 0.204 | 0.612 | 759 | 0.613 | 0.878 | | | | 365 | 0.695 | 0.878 | 5 | 0.051 |
| BRD-K16730910-001-07-3 | regorafenib | 727.5 | 0.879 | 1.000 | 1248.5 | 0.158 | 0.665 | 785 | 0.827 | 1.000 | 2791 | 0.409 | 1.000 | 1275 | 0.829 | 1.000 | 556 | 0.898 | 1.000 | 1101.5 | 0.913 | 1.000 | 876 | 0.838 | 1.000 | 97 | 0.034 | 0.226 | 242 | 0.449 | 1.000 | 405.5 | 0.012 | 0.226 | | | | 374 | 0.922 | 1.000 | 170 | 0.503 |
| BRD-K17061841-001-03-5 | stattic | 3360 | 0.351 | 1.000 | 4378 | 0.005 | 0.069 | 3732 | 0.522 | 1.000 | 11883 | 0.607 | 0.959 | 3707 | 0.256 | 1.000 | 3789 | 0.959 | 0.959 | 4427 | 0.550 | 0.959 | 4913 | 0.550 | 0.959 | 1582 | 0.241 | 0.722 | 2325 | 0.662 | 0.959 | 2576 | 0.557 | 0.959 | 1227 | 0.895 | 0.959 | 535 | 0.458 | 0.959 | 942 | 0.820 |
| BRD-K17068645-003-04-2 | bendamustine | 391.5 | 0.377 | 0.660 | 484.5 | 0.130 | 0.680 | 522 | 0.965 | 0.993 | 949.5 | 0.453 | 0.732 | 423 | 0.734 | 0.964 | 327 | 0.302 | 0.944 | 281.5 | 0.309 | 0.944 | 284 | 0.064 | 0.602 | 132 | 0.680 | 0.952 | 126.5 | 0.248 | 0.474 | 75 | 0.055 | 0.602 | 105.5 | 0.938 | 0.993 | 190 | 0.958 | 0.993 | 93 | 0.101 |
| BRD-K17075857-001-18-4 | chloroxine | 2930 | 0.945 | 0.989 | 3171 | 0.022 | 0.247 | 2495 | 0.959 | 0.989 | 7426 | 0.035 | 0.247 | 2310 | 0.031 | 0.247 | 1452 | 0.128 | 0.539 | 3965 | 0.989 | 0.989 | 3214 | 0.524 | 0.885 | 1009 | 0.403 | 0.885 | 1360 | 0.496 | 0.885 | 1742 | 0.616 | 0.885 | 924 | 0.512 | 0.885 | 608 | 0.483 | 0.885 | 676 | 0.632 |
| BRD-K17203476-001-01-9 | LY2874455 | 5238 | 0.962 | 0.996 | 7616 | 0.992 | 0.996 | 3689 | 0.138 | 0.415 | 14068 | 0.353 | 0.741 | 3245 | 0.055 | 0.287 | 4856 | 0.996 | 0.996 | 6607 | 0.980 | 0.996 | 5192 | 0.202 | 0.472 | 820 | 0.003 | 0.026 | 2977 | 0.823 | 0.996 | 2471 | 0.135 | 0.415 | 989 | 0.001 | 0.019 | 928 | 0.936 | 0.996 | 628 | 0.074 |
| BRD-K17610631-001-03-3 | indisulam | 1794 | 0.422 | 0.885 | 2184 | 0.997 | 0.997 | 1703.5 | 0.772 | 0.997 | 4614 | 0.421 | 0.885 | 1441.5 | 0.067 | 0.200 | 1736.5 | 0.917 | 0.997 | 2544 | 0.955 | 0.997 | 2039.5 | 0.471 | 0.899 | 298 | 0.024 | 0.083 | 1226 | 0.775 | 0.997 | 1100 | 0.557 | 0.900 | 345.5 | 0.013 | 0.054 | 358.5 | 0.876 | 0.997 | 146 | 0.002 |
| BRD-K17702546-001-03-2 | PD-168393 | 2572 | 0.738 | 1.000 | 5831 | 1.000 | 1.000 | 2770 | 0.203 | 0.825 | 11457.5 | 0.718 | 1.000 | 3639.5 | 0.134 | 0.705 | 2257 | 0.021 | 0.146 | 4096 | 0.548 | 1.000 | 4878 | 0.993 | 1.000 | 1157 | 0.705 | 1.000 | 1571 | 0.561 | 1.000 | 1993 | 0.303 | 0.908 | 742 | 0.903 | 1.000 | 392 | 0.236 | 0.825 | 276 | 0.847 |
| BRD-K17705806-003-04-4 | JTC-801 | 1163 | 0.553 | 0.829 | 3378 | 0.968 | 0.968 | 1074 | 0.050 | 0.528 | 5429.5 | 0.434 | 1.000 | 2423 | 0.517 | 1.000 | 1944.5 | 0.334 | 1.000 | 2390 | 0.929 | 0.968 | 2380 | 0.846 | 0.967 | 684 | 0.330 | 0.533 | 1662 | 0.745 | 0.921 | 1181 | 0.215 | 1.000 | 617 | 0.210 | 0.422 | 191 | 0.263 | 0.460 | 105 | 0.074 |
| BRD-K17743125-001-08-4 | belinostat | 1618 | 0.428 | 0.847 | 4169 | 0.629 | 0.870 | 2565 | 0.535 | 0.847 | 8619 | 0.681 | 0.870 | 2395 | 0.022 | 0.234 | 1991.5 | 0.365 | 0.847 | 2990.5 | 0.954 | 0.992 | 3248 | 0.992 | 0.992 | 1582 | 0.381 | 0.847 | 2285 | 0.704 | 0.870 | 1096.5 | 0.149 | 0.627 | 952 | 0.022 | 0.234 | 565.5 | 0.379 | 0.847 | 481 | 0.534 |
| BRD-K17849083-001-32-9 | tranilast | 2865 | 0.393 | 0.735 | 4966 | 0.423 | 0.735 | 2546 | 0.249 | 0.870 | 9082 | 0.221 | 0.870 | 3832 | 0.327 | 0.982 | 3549 | 0.914 | 0.999 | 5627 | 1.000 | 1.000 | 4691.5 | 0.952 | 0.999 | 1163 | 0.110 | 0.257 | 1720 | 0.738 | 0.912 | 1675 | 0.035 | 0.600 | 1574.5 | 0.560 | 0.735 | 527 | 0.519 | 0.735 | 637 | 0.397 |
| BRD-K18157228-001-01-7 | fdcyd | 2490 | 0.006 | 0.126 | 6882 | 1.000 | 1.000 | 3036 | 0.361 | 0.690 | 12850 | 0.679 | 0.891 | 4113 | 0.325 | 0.682 | 3579 | 0.899 | 0.974 | 4756 | 0.928 | 0.974 | 4336 | 0.268 | 0.682 | 1412 | 0.116 | 0.654 | 2605 | 0.592 | 0.848 | 1594 | 0.256 | 0.682 | 1025 | 0.187 | 0.654 | 682 | 0.401 | 0.703 | 628 | 0.311 |
| BRD-K18163608-001-05-6 | HA14-1 | 1001.5 | 0.748 | 0.924 | 970.5 | 0.012 | 0.253 | 815 | 0.901 | 0.941 | 2898 | 0.491 | 0.924 | 1581 | 0.920 | 0.941 | 777 | 0.649 | 0.924 | 660 | 0.576 | 0.924 | 1017 | 0.153 | 0.536 | 317 | 0.666 | 0.924 | 716 | 0.722 | 0.924 | 623 | 0.472 | 0.924 | 16 | 0.078 | 0.327 | 116 | 0.182 | 0.546 | 68 | 0.013 |
| BRD-K18375018-376-02-3 | spermine | 435 | 0.964 | 0.986 | 629 | 0.241 | 0.871 | 232 | 0.082 | 0.861 | 1176 | 0.300 | 0.871 | 469 | 0.188 | 0.871 | 364 | 0.325 | 0.871 | 506 | 0.683 | 0.991 | 618 | 0.690 | 0.991 | 85 | 0.239 | 0.871 | 413 | 0.935 | 0.986 | 89 | 0.789 | 0.991 | 127 | 0.552 | 0.991 | 69 | 0.595 | 0.991 | 153 | 0.332 |
| BRD-K18424115-003-01-3 | TG-100572 | 1512 | 0.830 | 0.871 | 2791 | 0.758 | 0.871 | 1544 | 0.241 | 0.696 | 6001 | 0.982 | 0.982 | 1461 | 0.085 | 0.696 | 1379.5 | 0.295 | 0.696 | 2016 | 0.598 | 0.837 | 2115 | 0.318 | 0.696 | 310 | 0.110 | 0.696 | 923.5 | 0.681 | 0.842 | 1021 | 0.236 | 0.696 | 399 | 0.099 | 0.696 | 475 | 0.818 | 0.871 | 420 | 0.676 |
| BRD-K18518344-001-15-9 | digitoxigenin | 6029.5 | 1.000 | 1.000 | 8822 | 0.987 | 1.000 | 4021 | 0.275 | 0.697 | 13258 | 0.003 | 0.072 | 5452 | 0.344 | 0.697 | 5495 | 0.949 | 1.000 | 7372 | 0.947 | 1.000 | 5806.5 | 0.169 | 0.592 | 1731 | 0.156 | 0.592 | 2550 | 0.103 | 0.557 | 3968 | 0.955 | 1.000 | 1964 | 0.337 | 0.697 | 469 | 0.204 | 0.612 | 1054 | 0.467 |
| BRD-K18619710-001-14-4 | digoxigenin | 2873 | 0.955 | 0.986 | 5519 | 0.981 | 0.986 | 2256 | 0.201 | 0.603 | 7613 | 0.004 | 0.080 | 4302 | 0.585 | 0.986 | 1591 | 0.029 | 0.206 | 4028 | 0.714 | 0.986 | 4037 | 0.720 | 0.986 | 971 | 0.236 | 0.621 | 1310 | 0.140 | 0.586 | 1456 | 0.273 | 0.638 | 783 | 0.024 | 0.206 | 634 | 0.774 | 0.986 | 476 | 0.169 |
| BRD-K18787491-001-08-6 | U-0126 | 444 | 0.249 | 0.735 | 1604.5 | 0.041 | 0.851 | 1033.5 | 0.698 | 0.915 | 2514 | 0.796 | 0.929 | 1127.5 | 0.251 | 0.735 | 1067 | 0.762 | 0.929 | 1228.5 | 0.596 | 0.834 | 612 | 0.490 | 0.735 | 320.5 | 0.362 | 0.735 | 368 | 0.256 | 0.735 | 612 | 0.135 | 1.000 | 340 | 0.984 | 0.984 | 157 | 0.376 | 0.735 | 84 | 0.451 |
| BRD-K18834913-001-01-1 | LB42708 | 5171 | 0.831 | 0.918 | 5841 | 0.141 | 1.000 | 3572 | 0.167 | 1.000 | 14497 | 0.448 | 1.000 | 5024 | 0.522 | 0.777 | 3406 | 0.555 | 0.777 | 5314 | 0.478 | 0.777 | 5003.5 | 0.508 | 0.777 | 1757 | 0.256 | 0.672 | 3456 | 0.981 | 0.995 | 2222 | 0.197 | 1.000 | 1885 | 0.798 | 0.918 | 638 | 0.231 | 0.672 | 791 | 0.465 |
| BRD-K18961567-001-01-1 | LY2801653 | 213 | 0.232 | 1.000 | 1563.5 | 0.768 | 0.955 | 1205 | 0.586 | 1.000 | 2807.5 | 0.063 | 1.000 | 1091.5 | 0.127 | 1.000 | 674.5 | 0.997 | 0.997 | 941.5 | 0.910 | 0.955 | 1023.5 | 0.235 | 0.549 | 456 | 0.093 | 0.243 | 300 | 0.027 | 0.082 | 735.5 | 0.730 | 0.955 | 813.5 | 0.876 | 0.955 | | | | 91 | 0.471 |
| BRD-K19034817-001-01-4 | pacritinib | 2483 | 0.001 | 0.009 | 7254 | 0.926 | 1.000 | 3712 | 0.462 | 0.809 | 12844 | 0.390 | 0.787 | 3016 | 0.008 | 0.059 | 4601 | 0.999 | 1.000 | 5835 | 0.932 | 1.000 | 7298 | 1.000 | 1.000 | 753 | 0.000 | 0.009 | 2813 | 0.630 | 0.926 | 2417 | 0.271 | 0.711 | 1752 | 0.165 | 0.494 | 530 | 0.139 | 0.494 | 629 | 0.088 |
| BRD-K19061412-001-02-4 | mubritinib | 2422 | 0.438 | 0.938 | 5458 | 0.932 | 0.980 | 1394 | 0.136 | 0.538 | 9714 | 0.914 | 0.980 | 3212 | 0.266 | 0.799 | 1807 | 0.015 | 0.178 | 2994 | 0.309 | 0.812 | 4475 | 0.972 | 0.980 | 606 | 0.062 | 0.435 | 1621 | 0.758 | 0.980 | 2012 | 0.474 | 0.938 | 1148 | 0.551 | 0.963 | 382 | 0.664 | 0.980 | 810 | 0.980 |
| BRD-K19085898-001-01-8 | darapladib | 280 | 0.377 | 0.756 | 893 | 0.854 | 0.944 | 1067 | 0.989 | 0.989 | 1960 | 0.086 | 0.986 | 712 | 0.135 | 0.986 | 312 | 0.243 | 0.986 | 1174 | 0.972 | 0.989 | 905 | 0.561 | 0.785 | 161 | 0.188 | 0.494 | 773 | 0.769 | 0.897 | 357 | 0.184 | 0.986 | 43 | 0.277 | 0.646 | 71 | 0.453 | 0.768 | 166 | 0.558 |
| BRD-K19295594-015-14-3 | gossypol | 4972 | 0.657 | 0.835 | 6595 | 0.428 | 0.835 | 3577 | 0.475 | 0.835 | 13732 | 0.367 | 0.835 | 4121 | 0.049 | 0.518 | 3802 | 0.502 | 0.835 | 6000 | 0.602 | 0.835 | 7936 | 0.998 | 0.998 | 1302 | 0.025 | 0.518 | 2898 | 0.247 | 0.740 | 3051 | 0.687 | 0.835 | 1428 | 0.123 | 0.708 | 649 | 0.234 | 0.740 | 764 | 0.163 |
| BRD-K19318328-001-01-3 | FR-139317 | 1296 | 0.380 | 0.725 | 3698.5 | 0.873 | 0.934 | 1869 | 0.308 | 0.702 | 6425 | 0.334 | 0.702 | 2262 | 0.138 | 0.413 | 1513 | 0.868 | 0.934 | 2933 | 0.879 | 0.934 | 3564.5 | 0.965 | 0.965 | 571 | 0.092 | 0.322 | 1093 | 0.061 | 0.258 | 607 | 0.015 | 0.245 | 499 | 0.046 | 0.258 | 585 | 0.874 | 0.934 | 336 | 0.023 |
| BRD-K19333160-001-01-3 | RKI-1447 | 970 | 0.941 | 0.941 | 1644 | 0.979 | 1.000 | 1081 | 0.516 | 0.903 | 2777 | 0.335 | 0.704 | 1667 | 0.897 | 1.000 | 436 | 0.716 | 1.000 | 803 | 0.275 | 0.704 | 1607 | 0.454 | 0.866 | 424 | 0.164 | 0.572 | 717 | 0.135 | 0.572 | 481 | 0.129 | 0.572 | 458 | 0.780 | 1.000 | 183 | 0.150 | 0.572 | 129 | 0.222 |
| BRD-K19540840-001-09-4 | saracatinib | 1048 | 0.999 | 1.000 | 1920 | 1.000 | 1.000 | 596 | 0.002 | 0.049 | 2821 | 0.686 | 1.000 | 759 | 0.005 | 0.049 | 490 | 0.634 | 1.000 | 1414 | 0.871 | 1.000 | 1096 | 0.911 | 1.000 | 517 | 0.929 | 1.000 | 768 | 0.136 | 0.260 | 474 | 0.856 | 1.000 | 244 | 0.793 | 1.000 | 270 | 0.878 | 1.000 | | |
| BRD-K19687926-379-07-4 | lapatinib | 1990 | 0.116 | 0.348 | 4257 | 1.000 | 1.000 | 1533.5 | 0.006 | 0.064 | 8196.5 | 0.966 | 1.000 | 3411.5 | 0.476 | 0.908 | 1582 | 0.009 | 0.037 | 3552 | 0.379 | 0.795 | 2527.5 | 0.997 | 1.000 | 1329 | 0.983 | 1.000 | 1505 | 0.778 | 1.000 | 1224 | 0.018 | 0.063 | 396 | 0.790 | 1.000 | 396 | 0.790 | 1.000 | 854 | 0.936 |
| BRD-K19761926-001-03-6 | eplerenone | 2360 | 0.613 | 0.823 | 3643 | 0.365 | 1.000 | 2264 | 0.660 | 0.823 | 8046 | 0.842 | 0.931 | 2162 | 0.024 | 0.501 | 1580 | 0.958 | 0.958 | 2202 | 0.179 | 1.000 | 2975 | 0.309 | 1.000 | 1200 | 0.936 | 0.958 | 1527 | 0.621 | 0.823 | 2127 | 0.811 | 0.931 | 506 | 0.319 | 0.559 | 431 | 0.469 | 0.757 | 92 | 0.311 |
| BRD-K20285085-074-04-5 | R406 | 2337 | 0.987 | 0.998 | 2996 | 0.998 | 0.998 | 1346 | 0.010 | 0.210 | 6556.5 | 0.176 | 0.481 | 2284.5 | 0.108 | 0.376 | 1866 | 0.954 | 0.998 | 3562 | 0.932 | 0.998 | 3302 | 0.791 | 0.998 | 671.5 | 0.067 | 0.353 | 1425.5 | 0.091 | 0.376 | 1311 | 0.043 | 0.300 | 944 | 0.036 | 0.300 | 572 | 0.823 | 0.998 | 123 | 0.393 |

| broad_id | name | al_nervous_system_PVAL_ADJ | fibroblast_MWU_STAT | fibroblast_PVAL | fibroblast_PVAL_ADJ | bile_duct_MWU_STAT | bile_duct_PVAL | bile_duct_PVAL_ADJ | thyroid_MWU_STAT | thyroid_PVAL | thyroid_PVAL_ADJ | upper_aerodigestive_MWU_STAT | upper_aerodigestive_PVAL | upper_aerodigestive_PVAL_ADJ | uterus_MWU_STAT | uterus_PVAL | uterus_PVAL_ADJ | liver_MWU_STAT | liver_PVAL | liver_PVAL_ADJ | esophagus_MWU_STAT | esophagus_PVAL | esophagus_PVAL_ADJ |
|---|---|---|---|---|---|---|---|---|---|---|---|---|---|---|---|---|---|---|---|---|---|---|---|
| BRD-K11071038-001-01-7 | ICI-162846 | | | | | 107 | 0.625 | 0.978 | 107 | 0.625 | 0.978 | 157 | 0.012 | 0.259 | 159 | 0.647 | 0.978 | 159 | 0.647 | 0.978 | 309 | 0.699 | 0.699 |
| BRD-K11073688-001-01-6 | CPI-1189 | 0.993 | | | | 141 | 0.944 | 0.993 | 235 | 0.267 | 0.811 | 280 | 0.079 | 0.824 | 416 | 0.463 | 0.811 | 398 | 0.993 | 0.993 | 803 | 0.981 | 0.993 |
| BRD-K11196887-001-22-0 | norfloxacin | 0.745 | | | | 225 | 0.758 | 0.885 | 305 | 0.693 | 0.885 | 872 | 0.403 | 0.745 | 489 | 0.424 | 0.745 | 530 | 0.813 | 0.885 | 936 | 0.377 | 0.745 |
| BRD-K11215326-001-03-7 | tazemetostat | 0.954 | | | | 117 | 0.696 | 0.954 | 478 | 0.622 | 0.954 | 728 | 0.373 | 0.954 | 683 | 0.698 | 0.954 | 359 | 0.032 | 0.673 | 1142 | 0.884 | 0.884 |
| BRD-K11267252-001-04-4 | alectinib | 0.368 | 18 | 0.069 | 0.484 | 619 | 0.749 | 0.940 | 778 | 0.277 | 0.843 | 2218 | 0.895 | 0.940 | 1027 | 0.746 | 0.940 | 1157 | 0.526 | 0.940 | 1765 | 0.142 | 0.595 |
| BRD-K11433652-001-17-0 | aspirin | 0.955 | | | | 132 | 0.718 | 0.955 | 80 | 0.274 | 0.870 | 356 | 0.290 | 0.870 | 153 | 0.854 | 0.955 | 263 | 0.520 | 0.955 | 443 | 0.858 | 0.955 |
| BRD-K11510141-001-01-8 | teprenone | 0.831 | | | | 118 | 0.657 | 0.831 | 427 | 0.673 | 0.831 | 1017 | 0.877 | 0.942 | 633.5 | 0.247 | 1.000 | 470 | 0.502 | 0.831 | 822 | 0.487 | 0.831 |
| BRD-K11630072-001-13-2 | carmofur | 0.981 | | | | | | | 211 | 0.677 | 0.981 | 541 | 0.154 | 0.540 | 1200.5 | 0.948 | 0.981 | 516 | 0.755 | 0.981 | 554 | 0.032 | 0.677 |
| BRD-K11636097-001-03-1 | JNJ-7706621 | 0.440 | 207 | 0.673 | 0.999 | 930 | 0.800 | 0.999 | 1405 | 0.931 | 0.999 | 2232 | 0.147 | 0.440 | 1911 | 0.484 | 0.999 | 862 | 0.087 | 0.440 | 2485 | 0.358 | 0.891 |
| BRD-K11663430-001-02-3 | pyroxamide | 0.676 | | | | 365 | 0.890 | 0.935 | 445 | 0.751 | 0.873 | 1132 | 0.251 | 0.885 | 896 | 0.214 | 0.885 | 693.5 | 0.447 | 0.782 | 1818 | 0.783 | 0.873 |
| BRD-K11851476-003-01-7 | pardoprunox | 0.998 | 155 | 0.418 | 0.982 | 1298 | 0.942 | 0.998 | 1939 | 0.644 | 0.998 | 2774 | 0.054 | 0.282 | 3772 | 0.902 | 0.998 | 1398 | 0.421 | 0.982 | 2580 | 0.010 | 0.089 |
| BRD-K11927976-050-02-9 | ER-27319 | 0.568 | 325 | 0.789 | 0.900 | 1017 | 0.496 | 0.900 | 1788 | 0.690 | 0.900 | 4799 | 0.900 | 0.900 | 2915 | 0.052 | 0.362 | 2839 | 0.542 | 0.900 | 4148 | 0.524 | 0.900 |
| BRD-K11973162-003-01-8 | tipiracil | 0.512 | | | | 510 | 0.623 | 0.778 | 726 | 0.423 | 0.697 | 3078 | 0.946 | 0.950 | 1785 | 0.317 | 0.604 | 1319 | 0.130 | 0.454 | 2256 | 0.543 | 0.778 |
| BRD-K12068470-001-02-5 | LY2608204 | 0.923 | | | | 50 | 0.210 | 1.000 | 638 | 0.870 | 0.984 | 1626 | 0.703 | 0.923 | 1194 | 0.360 | 1.000 | 764 | 0.371 | 0.666 | 1637 | 0.392 | 0.666 |
| BRD-K12184916-001-15-4 | NVP-BEZ235 | 0.981 | 148 | 0.471 | 0.899 | 1231 | 0.927 | 0.981 | 1239 | 0.772 | 0.967 | 2023 | 0.168 | 0.556 | 1899 | 0.293 | 0.683 | 1073 | 0.141 | 0.556 | 1665 | 0.051 | 0.427 |
| BRD-K12184916-001-19-6 | NVP-BEZ235 | 0.698 | | | | | | | 116 | 0.269 | 1.000 | 815 | 0.705 | 0.908 | 565 | 0.392 | 1.000 | 253 | 0.100 | 0.210 | 172 | 0.001 | 0.028 |
| BRD-K12244279-001-05-8 | MEK1-2-inhibitor | 0.203 | | | | 21 | 0.161 | 0.843 | 247 | 0.410 | 1.000 | 273 | 0.038 | 0.398 | 214 | 0.574 | 1.000 | 219 | 0.601 | 0.902 | 600 | 0.584 | 0.902 |
| BRD-K12251893-065-04-7 | vincristine | 0.224 | 22 | 0.065 | 0.274 | 1756.5 | 0.997 | 0.999 | 1569 | 0.481 | 0.814 | 2544.5 | 0.028 | 0.224 | 1978 | 0.060 | 0.274 | 2400.5 | 0.806 | 0.996 | 3574 | 0.504 | 0.814 |
| BRD-K12343256-001-08-9 | trametinib | 0.636 | | | | 710 | 0.497 | 0.949 | 506 | 0.050 | 0.260 | 2307 | 0.017 | 0.122 | 3634.5 | 0.996 | 0.999 | 3247 | 0.888 | 0.999 | 4543.5 | 0.983 | 0.999 |
| BRD-K12343256-001-14-7 | trametinib | 0.871 | | | | 345 | 0.140 | 0.586 | 524 | 0.556 | 1.000 | 1736 | 0.048 | 0.313 | 1043 | 0.814 | 1.000 | 1400 | 0.772 | 1.000 | 1378 | 0.742 | 1.000 |
| BRD-K12502280-001-08-0 | TG-101348 | 0.967 | | | | 160 | 0.956 | 0.972 | 199 | 0.314 | 1.000 | 692 | 0.512 | 0.913 | 393 | 0.504 | 0.913 | 135 | 0.026 | 0.108 | 768 | 0.524 | 0.913 |
| BRD-K12539581-001-23-7 | nocodazole | 0.206 | 151 | 0.348 | 0.742 | 1942 | 0.904 | 0.993 | 2092 | 0.457 | 0.873 | 3066 | 0.005 | 0.099 | 2685 | 0.019 | 0.202 | 2838 | 0.573 | 0.926 | 4252 | 0.121 | 0.424 |
| BRD-K12787259-001-04-3 | CX-5461 | 1.000 | | | | 549 | 0.753 | 0.968 | 998 | 0.238 | 0.506 | 1161 | 0.010 | 0.217 | 1596 | 0.097 | 0.681 | 1441 | 0.690 | 0.968 | 2101 | 0.185 | 0.506 |
| BRD-K12867552-001-04-7 | PCI-24781 | 0.251 | 46 | 0.124 | 0.325 | 1148 | 0.801 | 0.989 | 1798 | 0.654 | 0.916 | 3399 | 0.225 | 0.522 | 3047 | 0.771 | 0.989 | 2157 | 0.279 | 0.533 | 3633 | 0.110 | 0.325 |
| BRD-K12885236-001-02-7 | dapivirine | 0.989 | 32 | 0.090 | 0.316 | 557 | 0.461 | 0.892 | 823 | 0.439 | 0.892 | 2251 | 0.716 | 0.989 | 1571 | 0.546 | 0.955 | 1432 | 0.199 | 0.522 | 1803 | 0.014 | 0.291 |
| BRD-K13044802-213-20-7 | ciclopirox | 0.638 | | | | 506 | 0.848 | 0.968 | 1040.5 | 0.822 | 0.968 | 1809.5 | 0.370 | 1.000 | 1611 | 0.876 | 0.968 | 797 | 0.618 | 0.955 | 1634.5 | 0.433 | 0.802 |
| BRD-K13049116-001-04-0 | BMS-754807 | 0.476 | 103 | 0.679 | 0.993 | 275 | 0.956 | 0.993 | 48 | 0.242 | 0.727 | 462 | 0.045 | 0.416 | 523 | 0.668 | 0.993 | 543 | 0.480 | 0.993 | 390 | 0.232 | 0.727 |
| BRD-K13049116-001-05-7 | BMS-754807 | 0.102 | 175 | 0.745 | 0.990 | 421.5 | 0.948 | 0.990 | 831.5 | 0.898 | 0.990 | 1814 | 0.366 | 0.961 | 1595 | 0.948 | 0.990 | 1068.5 | 0.852 | 0.990 | 1929.5 | 0.818 | 0.990 |
| BRD-K13050541-001-01-9 | acivicin | 0.975 | 196 | 0.887 | 0.976 | 373 | 0.318 | 0.513 | | | | 538 | 0.013 | 0.274 | 1268 | 0.516 | 1.000 | 1080 | 0.976 | 0.976 | 1835 | 0.881 | 0.976 |
| BRD-K13060017-001-01-3 | UNC0631 | 0.914 | 258 | 0.638 | 0.914 | 1484 | 0.836 | 0.999 | 1641 | 0.339 | 0.790 | 3451 | 0.326 | 0.790 | 3872 | 0.653 | 0.914 | 1559 | 0.461 | 0.914 | 3651 | 0.142 | 0.748 |
| BRD-K13087974-001-02-5 | CGP-52411 | 0.755 | | | | 234 | 0.564 | 0.903 | 616 | 0.699 | 0.903 | 1132 | 0.157 | 0.690 | 1486 | 0.552 | 0.903 | 676.5 | 0.326 | 0.755 | 1675 | 0.696 | 0.903 |
| BRD-K13094524-001-07-5 | PFI-1 | 0.654 | | | | 638 | 0.835 | 0.922 | 1129 | 0.533 | 0.786 | 2585.5 | 0.475 | 0.767 | 1689 | 0.158 | 1.000 | 1495 | 0.611 | 0.786 | 2420 | 0.308 | 0.654 |
| BRD-K13112821-001-17-4 | artemisinin | 0.723 | 65 | 0.434 | 0.757 | 30 | 0.159 | 0.621 | 237 | 0.288 | 0.672 | 803 | 0.675 | 0.757 | 435.5 | 0.563 | 0.757 | 229.5 | 0.585 | 0.757 | 1186.5 | 0.987 | 0.987 |
| BRD-K13167095-001-02-9 | AR-42 | 0.270 | 212 | 0.570 | 0.894 | 809 | 0.324 | 0.618 | 1740 | 0.407 | 0.712 | 3110 | 0.284 | 0.597 | 1972 | 0.040 | 0.270 | 1806 | 0.036 | 0.270 | 4194 | 0.744 | 0.962 |
| BRD-K13169950-001-14-4 | NSC-3852 | 0.315 | | | | 333 | 0.693 | 0.971 | 569 | 0.287 | 0.548 | 759 | 0.773 | 0.984 | 1164 | 0.206 | 0.480 | 1131 | 0.968 | 0.984 | 400 | 0.117 | 1.000 |
| BRD-K13183738-317-10-2 | pentamidine | 0.755 | | | | 247 | 0.973 | 0.976 | 289 | 0.887 | 0.976 | 467 | 0.508 | 1.000 | 536 | 0.752 | 0.928 | 68 | 0.024 | 0.064 | 569 | 0.645 | 0.928 |
| BRD-K13387373-004-14-5 | thonzonium | 0.968 | 12 | 0.035 | 0.229 | 1136 | 0.968 | 0.968 | 799 | 0.044 | 0.229 | 4254 | 0.936 | 0.968 | 1429 | 0.000 | 0.000 | 2769 | 0.735 | 0.968 | 4205 | 0.921 | 0.968 |
| BRD-K13390322-001-06-3 | AT-7519 | 1.000 | 250 | 0.707 | 1.000 | 1160 | 0.548 | 1.000 | 1467 | 0.897 | 1.000 | 2040 | 0.001 | 0.011 | 3238 | 0.327 | 0.847 | 2792.5 | 0.936 | 1.000 | 3309 | 0.164 | 0.682 |
| BRD-K13409143-001-01-9 | isopropyl-myristate | 0.609 | 115 | 0.594 | 0.932 | 107 | 0.128 | 0.701 | 630.5 | 0.847 | 0.932 | 1033 | 0.275 | 0.701 | 1002.5 | 0.766 | 0.932 | 1149 | 0.495 | 0.888 | 1105 | 0.146 | 0.701 |
| BRD-K13514097-001-05-3 | everolimus | 0.356 | | | | 401 | 0.915 | 0.978 | 251 | 0.814 | 0.978 | 1099 | 0.908 | 0.978 | 228 | 0.008 | 0.171 | 637 | 0.071 | 0.214 | 1224 | 0.386 | 0.675 |
| BRD-K13566078-001-02-3 | BMS-345541 | 0.513 | | | | 259 | 0.390 | 0.818 | 610.5 | 0.981 | 1.000 | 1035.5 | 0.134 | 0.513 | 494 | 0.136 | 0.513 | 330.5 | 0.042 | 0.395 | 1389.5 | 0.740 | 1.000 |
| BRD-K13571841-001-14-4 | pepstatin | 0.711 | 31 | 0.110 | 0.711 | 430.5 | 0.832 | 0.953 | 522.5 | 0.765 | 0.953 | 1346.5 | 0.340 | 0.881 | 638 | 0.491 | 0.881 | 640.5 | 0.264 | 0.792 | 1477 | 0.561 | 0.907 |
| BRD-K13606314-001-01-3 | P5091 | 0.900 | 156 | 0.743 | 0.900 | 357 | 0.570 | 0.900 | 470 | 0.568 | 0.900 | 1552 | 0.749 | 0.900 | 713 | 0.182 | 0.900 | 820 | 0.197 | 0.900 | 1721 | 0.920 | 0.920 |
| BRD-K13646352-001-03-8 | midostaurin | 0.750 | 434 | 0.980 | 0.992 | 1371 | 0.571 | 0.923 | 2049 | 0.382 | 0.862 | 3747 | 0.190 | 0.665 | 4738 | 0.813 | 0.992 | 3262 | 0.537 | 0.923 | 5423 | 0.735 | 0.992 |
| BRD-K13662825-001-07-5 | dinaciclib | 0.146 | 401 | 0.957 | 1.000 | 635 | 0.070 | 0.294 | 1618 | 0.703 | 1.000 | 2744 | 0.002 | 0.034 | 2425 | 0.021 | 0.145 | 2649 | 0.504 | 0.939 | 4350 | 0.352 | 0.821 |
| BRD-K13756951-001-01-4 | bitopertin | 0.913 | 60 | 0.196 | 0.913 | 204 | 0.888 | 0.922 | 814 | 0.742 | 0.922 | 1455 | 0.457 | 0.913 | 604 | 0.119 | 0.913 | 651 | 0.655 | 0.917 | 1864 | 0.201 | 0.913 |
| BRD-K13810148-311-03-2 | givinostat | 0.357 | 390 | 0.931 | 1.000 | 1284 | 0.556 | 0.778 | 1523 | 0.360 | 0.581 | 3730 | 0.199 | 0.388 | 3199 | 0.203 | 0.388 | 2362 | 0.141 | 0.371 | 3775 | 0.081 | 0.357 |
| BRD-K13949769-001-01-7 | tenovin-6 | 0.531 | 225 | 0.709 | 0.876 | 581 | 0.662 | 0.869 | 1407 | 0.814 | 0.949 | 2415 | 0.057 | 0.499 | 2284 | 0.480 | 0.780 | 1681 | 0.521 | 0.782 | 3384 | 0.640 | 0.869 |
| BRD-K14109347-001-03-4 | LY2603618 | 0.658 | 60 | 0.228 | 0.658 | 383 | 0.383 | 0.722 | 811 | 0.983 | 0.983 | 1064.5 | 0.920 | 0.983 | 657 | 0.165 | 0.658 | 360 | 0.313 | 0.658 | 1102 | 0.172 | 0.658 |
| BRD-K14560436-001-01-4 | TG-02 | 0.519 | 400 | 0.944 | 0.969 | 709 | 0.110 | 0.519 | 2119 | 0.751 | 0.949 | 4287 | 0.795 | 0.949 | 3155 | 0.160 | 0.559 | 2941 | 0.388 | 0.873 | 4087 | 0.528 | 0.873 |
| BRD-K14618467-001-04-6 | IKK-16 | 0.984 | 209 | 0.722 | 0.984 | 845 | 0.752 | 0.984 | 804 | 0.405 | 0.984 | 2806 | 0.922 | 0.984 | 908 | 0.011 | 0.100 | 1968 | 0.984 | 0.984 | 2566 | 0.941 | 0.984 |
| BRD-K14796088-003-26-8 | berberine | 0.928 | 324 | 0.900 | 0.928 | 815 | 0.682 | 0.928 | 1472 | 0.799 | 0.928 | 1912 | 0.001 | 0.027 | 3147 | 0.829 | 0.928 | 2072 | 0.176 | 0.739 | 3458 | 0.543 | 0.928 |
| BRD-K14991967-001-02-6 | GSK650394 | 0.892 | | | | 217 | 0.945 | 0.982 | 878 | 0.737 | 0.911 | 1391 | 0.338 | 1.000 | 899 | 0.336 | 1.000 | 675 | 0.057 | 0.199 | 1315 | 0.371 | 0.780 |
| BRD-K15025317-001-15-1 | BAY-11-7082 | 0.047 | | | | 625 | 0.779 | 0.978 | 1330 | 0.944 | 0.984 | 1706 | 0.214 | 1.000 | 1244 | 0.044 | 0.461 | 877 | 0.281 | 0.716 | 1912 | 0.774 | 0.978 |
| BRD-K15108141-001-06-6 | gemcitabine | 0.775 | 170 | 0.525 | 0.775 | 1084 | 0.908 | 0.994 | 831 | 0.553 | 0.775 | 2637 | 0.512 | 0.775 | 2498 | 0.819 | 0.994 | 1766 | 0.210 | 0.631 | 2577 | 0.605 | 0.794 |
| BRD-K15179879-001-03-2 | carfilzomib | 0.864 | 112 | 0.251 | 0.637 | 1591 | 0.786 | 0.917 | 2133 | 0.430 | 0.753 | 4214 | 0.199 | 0.637 | 4136 | 0.380 | 0.726 | 4030 | 0.937 | 0.999 | 3864 | 0.021 | 0.448 |
| BRD-K15409150-001-05-8 | penfluridol | 0.630 | 210 | 0.542 | 0.876 | 1363 | 0.946 | 0.961 | 1301 | 0.243 | 0.607 | 2914 | 0.057 | 0.397 | 3849 | 0.935 | 0.961 | 2985 | 0.913 | 0.961 | 3884 | 0.177 | 0.607 |
| BRD-K15424032-001-03-6 | SB-228357 | 0.980 | | | | 48 | 0.093 | 0.675 | 19 | 0.136 | 0.675 | 226 | 0.136 | 0.675 | 105 | 0.161 | 0.675 | 17 | 0.022 | 0.057 | 161 | 0.518 | 0.790 |
| BRD-K15567136-003-27-2 | papaverine | 0.198 | | | | 751 | 0.714 | 0.887 | 581 | 0.141 | 0.984 | 1527 | 0.193 | 1.000 | 951 | 0.013 | 0.141 | 1331 | 0.915 | 0.962 | 2272 | 0.684 | 0.887 |
| BRD-K15600710-066-04-1 | obatoclax | 0.870 | 74 | 0.232 | 0.870 | 604 | 0.771 | 0.870 | 1032 | 0.637 | 0.870 | 1941 | 0.018 | 0.194 | 1940 | 0.411 | 0.870 | 1683 | 0.283 | 0.870 | 1767 | 0.013 | 0.194 |
| BRD-K15600710-066-05-8 | obatoclax | 0.975 | 77 | 0.195 | 0.672 | 465 | 0.894 | 0.975 | 946 | 0.563 | 0.975 | 1503 | 0.001 | 0.024 | 1823 | 0.224 | 0.672 | 1134 | 0.095 | 0.410 | 1604.5 | 0.016 | 0.170 |
| BRD-K15898725-001-01-7 | maytansinol-isobutyrate | 0.688 | 29 | 0.139 | 0.574 | 699 | 0.890 | 0.955 | 393 | 0.171 | 0.574 | 647 | 0.802 | 0.951 | 821 | 0.140 | 0.574 | 1248 | 0.725 | 0.951 | 1006 | 0.659 | 0.936 |
| BRD-K16136380-001-01-7 | pyrithione-zinc | 0.975 | 39 | 0.103 | 0.362 | 385 | 0.170 | 0.446 | 1073 | 0.402 | 0.843 | 4451 | 0.946 | 0.975 | 3403 | 0.840 | 0.975 | 2190 | 0.079 | 0.362 | 3417 | 0.727 | 0.975 |
| BRD-K16180792-001-01-7 | bis(maltolato)oxovanadium(IV) | 0.999 | | | | 29 | 0.139 | 0.727 | 310 | 0.485 | 1.000 | 398 | 0.821 | 0.999 | 654 | 0.062 | 0.646 | 687 | 0.510 | 1.000 | 453 | 0.732 | 0.999 |
| BRD-K16189898-003-03-3 | CHIR-99021 | 0.138 | | | | 148 | 0.842 | 0.922 | 220.5 | 0.890 | 0.922 | 465 | 0.746 | 0.922 | 174 | 0.060 | 0.907 | 292 | 0.922 | 0.922 | 247.5 | 0.724 | 0.922 |
| BRD-K16478699-001-09-2 | PLX-4720 | 0.860 | 35 | 0.090 | 0.632 | 763 | 0.775 | 0.992 | 879 | 0.405 | 0.945 | 2132.5 | 0.352 | 0.923 | 1455 | 0.024 | 0.248 | 1714 | 0.887 | 0.992 | 3751.5 | 0.991 | 0.992 |
| BRD-K16551357-001-03-2 | evacetrapib | 0.358 | | | | 402 | 0.806 | 0.903 | 512 | 0.493 | 0.869 | 1128 | 0.375 | 0.787 | 1162 | 0.280 | 0.680 | 561 | 0.168 | 0.590 | 878 | 0.012 | 0.245 |
| BRD-K16730910-001-07-3 | regorafenib | 1.000 | | | | 86 | 0.508 | 1.000 | 329.5 | 0.212 | 0.742 | 573.5 | 0.738 | 1.000 | 356 | 0.043 | 0.226 | 257 | 0.022 | 0.226 | 910 | 0.889 | 0.889 |
| BRD-K17061841-001-03-5 | stattic | 0.959 | | | | 1258 | 0.917 | 0.959 | 1751 | 0.826 | 0.959 | 1814 | 0.007 | 0.069 | 2960 | 0.590 | 0.959 | 2783 | 0.873 | 0.959 | 3899 | 0.778 | 0.959 |
| BRD-K17068645-003-04-2 | bendamustine | 0.211 | | | | 188.5 | 0.955 | 0.993 | 94 | 0.360 | 0.944 | 291 | 0.353 | 0.944 | 11 | 0.086 | 0.602 | 247 | 0.677 | 0.952 | 639.5 | 0.993 | 0.993 |
| BRD-K17075857-001-18-4 | chloroxine | 0.885 | 13 | 0.057 | 0.302 | 852.5 | 0.907 | 0.989 | 1720 | 0.910 | 0.989 | 2417 | 0.978 | 0.989 | 2341.5 | 0.632 | 0.885 | 1230 | 0.309 | 0.885 | 2290 | 0.573 | 0.885 |
| BRD-K17203476-001-01-9 | LY2874455 | 0.309 | 5 | 0.015 | 0.102 | 1516 | 0.626 | 0.939 | 1520 | 0.394 | 0.751 | 3422 | 0.173 | 0.455 | 3617 | 0.565 | 0.912 | 2950 | 0.483 | 0.845 | 4725 | 0.683 | 0.956 |
| BRD-K17610631-001-03-3 | indisulam | 0.008 | | | | 808 | 0.990 | 0.997 | 551 | 0.938 | 0.997 | 2071 | 0.515 | 0.900 | 1039 | 0.058 | 1.000 | 560 | 0.168 | 0.442 | 1516 | 0.197 | 1.000 |
| BRD-K17702546-001-03-2 | PD-168393 | 1.000 | 202 | 0.614 | 1.000 | 1065 | 0.824 | 1.000 | 2057 | 0.993 | 1.000 | 1408 | 0.000 | 0.002 | 2382 | 0.381 | 1.000 | 1973 | 0.643 | 1.000 | 1949 | 0.000 | 0.005 |
| BRD-K17705806-003-04-4 | JTC-801 | 0.174 | | | | 216 | 0.875 | 0.967 | 522 | 0.523 | 1.000 | 1573 | 0.048 | 0.528 | 1192 | 0.604 | 0.845 | 737 | 0.221 | 0.422 | 2265 | 0.738 | 0.921 |
| BRD-K17743125-001-08-4 | belinostat | 0.847 | 152 | 0.480 | 0.847 | 541 | 0.124 | 0.627 | 1269 | 0.565 | 0.847 | 2487 | 0.873 | 0.965 | 2044.5 | 0.276 | 0.847 | 1585 | 0.120 | 0.627 | 2792.5 | 0.762 | 0.889 |
| BRD-K17849083-001-32-9 | tranilast | 0.735 | | | | 907 | 0.861 | 0.999 | 918 | 0.057 | 0.600 | 2318 | 0.107 | 0.736 | 2238 | 0.140 | 0.736 | 2065 | 0.550 | 0.735 | 2995 | 0.516 | 0.735 |
| BRD-K18157228-001-01-7 | fdcyd | 0.682 | 58 | 0.158 | 0.654 | 1102 | 0.771 | 0.953 | 1550 | 0.826 | 0.964 | 2322 | 0.103 | 0.654 | 2458 | 0.445 | 0.719 | 2442 | 0.606 | 0.848 | 3164 | 0.134 | 0.654 |
| BRD-K18163608-001-05-6 | HA14-1 | 0.067 | | | | 305 | 0.941 | 0.941 | 887 | 0.867 | 0.941 | 795 | 0.475 | 0.924 | 885 | 0.489 | 0.924 | 773 | 0.639 | 0.924 | 927 | 0.396 | 1.000 |
| BRD-K18375018-376-02-3 | spermine | 0.871 | | | | 230 | 0.986 | 0.986 | 163 | 0.802 | 0.991 | 297 | 0.513 | 0.991 | 176 | 0.065 | 0.861 | 214 | 0.714 | 0.991 | 430 | 0.605 | 0.991 |
| BRD-K18424115-003-01-3 | TG-100572 | 0.842 | 91 | 0.331 | 0.696 | 255 | 0.546 | 0.819 | 588 | 0.222 | 0.696 | 1126 | 0.404 | 0.707 | 1108 | 0.373 | 0.707 | 632 | 0.306 | 0.696 | 1429 | 0.542 | 0.819 |
| BRD-K18518344-001-15-9 | digitoxigenin | 0.708 | 60 | 0.106 | 0.557 | 1268 | 0.472 | 0.708 | 2028 | 0.398 | 0.697 | 4541 | 0.756 | 0.992 | 4316 | 0.612 | 0.857 | 1904 | 0.029 | 0.310 | 4572 | 0.382 | 0.697 |
| BRD-K18619710-001-14-4 | digoxigenin | 0.591 | 39 | 0.118 | 0.586 | 1131 | 0.913 | 0.986 | 1087 | 0.393 | 0.825 | 3231 | 0.986 | 0.986 | 2755 | 0.816 | 0.986 | 2252 | 0.483 | 0.922 | 3791 | 0.967 | 0.986 |
| BRD-K18787491-001-08-6 | U-0126 | 0.735 | | | | 340 | 0.984 | 0.984 | 201 | 0.226 | 1.000 | 1020 | 0.488 | 0.735 | 597 | 0.446 | 0.735 | 408.5 | 0.379 | 0.735 | 1262 | 0.969 | 0.984 |
| BRD-K18834913-001-01-1 | LB42708 | 0.777 | | | | 838 | 0.545 | 0.777 | 1689 | 0.597 | 0.784 | 3103 | 0.016 | 0.328 | 4563.5 | 0.995 | 0.995 | 3357 | 0.671 | 0.829 | 4293 | 0.519 | 0.777 |
| BRD-K18961567-001-01-1 | LY2801653 | 0.900 | | | | 354.5 | 0.773 | 0.955 | 486.5 | 0.844 | 0.955 | 1099.5 | 0.867 | 0.955 | 738 | 0.506 | 1.000 | 733.5 | 0.291 | 0.611 | 1326 | 0.800 | 0.955 |
| BRD-K19034817-001-01-4 | pacritinib | 0.464 | 330 | 0.836 | 1.000 | 919 | 0.412 | 0.787 | 1654 | 0.832 | 1.000 | 3389 | 0.347 | 0.787 | 2757 | 0.157 | 0.494 | 2466 | 0.661 | 0.926 | 4232 | 0.590 | 0.926 |
| BRD-K19061412-001-02-4 | mubritinib | 0.980 | 282 | 0.868 | 0.980 | 1192 | 0.841 | 0.980 | 874 | 0.631 | 0.980 | 2174 | 0.116 | 0.538 | 1968 | 0.154 | 0.538 | 1204 | 0.017 | 0.178 | 2763 | 0.491 | 0.938 |
| BRD-K19085898-001-01-8 | darapladib | 0.785 | | | | 117 | 0.282 | 0.986 | 437 | 0.693 | 0.856 | 798 | 0.648 | 0.850 | 295 | 0.194 | 0.986 | 357 | 0.396 | 0.756 | 524 | 0.475 | 0.768 |
| BRD-K19295594-015-14-3 | gossypol | 0.708 | 298 | 0.714 | 0.835 | 1270 | 0.542 | 0.835 | 1904 | 0.355 | 0.835 | 4576 | 0.866 | 0.910 | 4093 | 0.715 | 0.835 | 2407 | 0.169 | 0.708 | 5194 | 0.785 | 0.868 |
| BRD-K19318328-001-01-3 | FR-139317 | 0.245 | 11 | 0.056 | 0.258 | 481 | 0.660 | 0.934 | 994.5 | 0.311 | 0.702 | 2071.5 | 0.720 | 0.934 | 2004 | 0.792 | 0.934 | 1526 | 0.723 | 0.934 | 2817 | 0.890 | 0.934 |
| BRD-K19333160-001-01-3 | RKI-1447 | 0.666 | 16 | 0.077 | 0.572 | | | | 319 | 0.309 | 0.704 | 986 | 0.684 | 1.000 | 577 | 0.010 | 0.214 | 168 | 0.909 | 1.000 | 1220 | 0.902 | 1.000 |
| BRD-K19540840-001-09-4 | saracatinib | | | | | 3 | 0.048 | 0.216 | 169 | 0.927 | 1.000 | 974 | 0.032 | 0.216 | 547 | 0.293 | 1.000 | 315 | 0.127 | 0.260 | 1151 | 0.051 | 0.216 |
| BRD-K19687926-379-07-4 | lapatinib | 1.000 | | | | 671 | 0.366 | 0.795 | 723 | 0.777 | 1.000 | 1467 | 0.001 | 0.028 | 2499 | 0.866 | 1.000 | 1163 | 0.284 | 0.747 | 1929 | 0.010 | 0.071 |
| BRD-K19761926-001-03-6 | eplerenone | 0.559 | | | | 196 | 0.220 | 1.000 | 1193 | 0.548 | 1.000 | 2237 | 0.206 | 1.000 | 2068 | 0.592 | 1.000 | 1269 | 0.666 | 0.823 | 2635 | 0.340 | 1.000 |
| BRD-K20285085-074-04-5 | R406 | 0.747 | 100 | 0.294 | 0.617 | 871 | 0.785 | 0.998 | 614 | 0.283 | 0.617 | 2033 | 0.427 | 0.747 | 1431 | 0.183 | 0.481 | 2062 | 0.785 | 0.998 | 3195 | 0.948 | 0.998 |

Suppl. Table 9

| broad_id | name | colorectal_MWU_STAT | colorectal_PVAL | colorectal_PVAL_ADJ | skin_MWU_STAT | skin_PVAL | skin_PVAL_ADJ | urinary_tract_MWU_STAT | urinary_tract_PVAL | urinary_tract_PVAL_ADJ | lung_MWU_STAT | lung_PVAL | lung_PVAL_ADJ | ovary_MWU_STAT | ovary_PVAL | ovary_PVAL_ADJ | breast_MWU_STAT | breast_PVAL | breast_PVAL_ADJ | pancreas_MWU_STAT | pancreas_PVAL | pancreas_PVAL_ADJ | nervous_system_MWU_STAT | central_nervous_system_PVAL | nervous_system_PVAL_ADJ | soft_tissue_MWU_STAT | soft_tissue_PVAL | soft_tissue_PVAL_ADJ | kidney_MWU_STAT | kidney_PVAL | kidney_PVAL_ADJ | gastric_MWU_STAT | gastric_PVAL | gastric_PVAL_ADJ | bone_MWU_STAT | bone_PVAL | bone_PVAL_ADJ | prostate_MWU_STAT | prostate_PVAL | prostate_PVAL_ADJ | al_nervous_system_MWU_STAT | peripheral_nervous_system_PVAL |
|---|---|---|---|---|---|---|---|---|---|---|---|---|---|---|---|---|---|---|---|---|---|---|---|---|---|---|---|---|---|---|---|---|---|---|---|---|---|---|---|---|---|---|
| BRD-K20290250-001-02-1 | BIX-02188 | 736.5 | 0.794 | 1.000 | 1598.5 | 0.792 | 1.000 | 485 | 0.067 | 0.351 | 3576.5 | 0.886 | 1.000 | 1044 | 0.514 | 0.968 | 641 | 0.553 | 0.968 | 1038 | 0.335 | 0.703 | 1116 | 0.213 | 0.558 | 414 | 0.067 | 0.351 | 539 | 0.953 | 1.000 | 333 | 0.162 | 0.558 | 17 | 0.014 | 0.294 | 240 | 0.777 | 1.000 | 206 | 0.622 |
| BRD-K20463415-300-01-4 | LY2606368 | 6138 | 0.969 | 0.998 | 7447 | 0.585 | 0.905 | 3901.5 | 0.064 | 0.446 | 17220.5 | 0.603 | 0.905 | 4428 | 0.106 | 0.446 | 3878 | 0.830 | 0.989 | 6507 | 0.694 | 0.911 | 6545 | 0.501 | 0.877 | 1460 | 0.101 | 0.446 | 2947 | 0.305 | 0.724 | 3501 | 0.421 | 0.804 | 3021 | 0.848 | 0.989 | 1309 | 0.957 | 0.998 | 305 | 0.003 |
| BRD-K20468903-001-01-6 | BNC105 | 5781 | 0.892 | 0.994 | 8429 | 0.987 | 0.994 | 3577 | 0.058 | 0.304 | 15103 | 0.251 | 0.479 | 5164 | 0.460 | 0.690 | 3994 | 0.842 | 0.994 | 6551 | 0.994 | 0.994 | 6112 | 0.370 | 0.598 | 1637 | 0.121 | 0.363 | 4604 | 0.987 | 0.994 | 2311 | 0.099 | 0.346 | 1866 | 0.282 | 0.493 | 456 | 0.057 | 0.304 | 779 | 0.161 |
| BRD-K20482099-001-09-4 | rutin | 1171 | 0.971 | 0.971 | 1982.5 | 0.935 | 0.971 | 942.5 | 0.084 | 0.351 | 4157 | 0.759 | 0.971 | 1268.5 | 0.039 | 0.273 | 1223 | 0.827 | 0.971 | 2108.5 | 0.946 | 0.971 | 1612.5 | 0.145 | 0.381 | 404.5 | 0.180 | 0.420 | 558 | 0.305 | 0.637 | 312 | 0.057 | 1.000 | 924 | 0.872 | 0.971 | 199 | 0.426 | 0.746 | 41 | 0.143 |
| BRD-K20545304-001-01-6 | bardoxolone | 1584 | 0.797 | 0.982 | 2744 | 0.740 | 0.982 | 1114 | 0.029 | 0.161 | 6518 | 0.729 | 0.982 | 830 | 0.005 | 0.095 | 1788 | 0.870 | 0.982 | 3232 | 0.982 | 0.982 | 2920 | 0.973 | 0.982 | 549 | 0.120 | 0.328 | 1724 | 0.976 | 0.982 | 915 | 0.545 | 0.982 | 682 | 0.591 | 0.982 | 145 | 0.031 | 0.161 | 293 | 0.072 |
| BRD-K20605374-001-05-4 | NSC-697923 | 4279 | 0.720 | 0.979 | 5189 | 0.020 | 0.291 | 3498 | 0.550 | 0.878 | 13781 | 0.930 | 0.980 | 4530 | 0.280 | 0.589 | 4546 | 0.769 | 0.979 | 6271 | 0.932 | 0.980 | 6772 | 0.996 | 0.996 | 1553 | 0.274 | 0.589 | 2486 | 0.472 | 0.826 | 2162 | 0.102 | 0.429 | 1979 | 0.336 | 0.642 | 764 | 0.792 | 0.979 | 752 | 0.186 |
| BRD-K20655524-003-12-0 | mefexamide | 5262 | 0.993 | 1.000 | 6559 | 0.176 | 0.528 | 3468 | 0.166 | 0.528 | 12715 | 0.017 | 0.279 | 5391 | 0.793 | 0.980 | 4283 | 0.718 | 0.953 | 6174 | 0.930 | 1.000 | 6017 | 0.726 | 0.953 | 2009 | 0.281 | 0.656 | 4787 | 1.000 | 1.000 | 2145 | 0.062 | 0.327 | 1564 | 0.221 | 0.579 | 289 | 0.058 | 0.327 | 562 | 0.141 |
| BRD-K20722021-001-02-1 | CEP-32496 | 548 | 0.555 | 0.996 | 882 | 0.000 | 0.000 | 1309 | 0.868 | 0.996 | 3032 | 0.979 | 0.996 | 1493 | 0.996 | 0.996 | 731 | 0.943 | 0.996 | 1213 | 0.856 | 0.996 | 876 | 0.341 | 0.894 | 358 | 0.087 | 0.260 | 633 | 0.791 | 0.996 | 288 | 0.089 | 0.621 | 164 | 0.412 | 0.961 | 185 | 0.524 | 0.996 | 221 | 0.708 |
| BRD-K20958582-001-01-4 | TH-302 | 2508.5 | 0.793 | 0.985 | 2748 | 0.392 | 0.892 | 2105 | 0.482 | 0.892 | 7129.5 | 0.678 | 0.985 | 2116 | 0.138 | 0.483 | 1637 | 0.718 | 0.985 | 3958 | 0.999 | 1.000 | 2225 | 0.477 | 0.892 | 506 | 0.384 | 0.892 | 1007 | 0.084 | 0.483 | 602 | 0.015 | 0.161 | 71 | 0.001 | 0.029 | 486 | 0.965 | 1.000 | 420 | 0.510 |
| BRD-K21025364-001-05-6 | NVP-TAE226 | 4596 | 0.209 | 0.619 | 7793 | 0.662 | 0.987 | 4143 | 0.260 | 0.619 | 14535 | 0.325 | 0.683 | 6186 | 0.701 | 0.987 | 5483 | 0.998 | 0.998 | 5897 | 0.230 | 0.619 | 7956 | 0.989 | 0.998 | 2181 | 0.773 | 0.987 | 3460 | 0.878 | 0.998 | 1795 | 0.018 | 0.191 | 2252 | 0.265 | 0.619 | 1100 | 0.986 | 0.998 | 582 | 0.040 |
| BRD-K21283037-001-21-5 | riluzole | 309.5 | 0.820 | 0.906 | 492 | 0.577 | 1.000 | 337.5 | 0.910 | 0.910 | 751.5 | 0.209 | 1.000 | 439.5 | 0.350 | 1.000 | 197.5 | 0.809 | 0.906 | 390.5 | 0.734 | 0.906 | 536.5 | 0.888 | 0.910 |  |  |  | 268 | 0.611 | 1.000 | 282.5 | 0.692 | 1.000 |  |  |  | 67 | 0.698 | 0.906 | 34 | 0.280 |
| BRD-K21361524-001-01-1 | selinexor | 7069 | 0.968 | 0.995 | 8799 | 0.660 | 0.866 | 4342 | 0.096 | 0.484 | 18962 | 0.558 | 0.836 | 6555 | 0.451 | 0.789 | 4585 | 0.274 | 0.614 | 7797 | 0.840 | 0.995 | 6794 | 0.269 | 0.614 | 2033 | 0.129 | 0.484 | 5307 | 0.995 | 0.995 | 3619 | 0.321 | 0.614 | 2257 | 0.138 | 0.484 | 967 | 0.542 | 0.836 | 1289 | 0.649 |
| BRD-K21396683-001-04-8 | marimastat | 3453 | 0.219 | 0.765 | 6593 | 0.571 | 0.922 | 3176 | 0.093 | 0.514 | 13531 | 0.293 | 0.880 | 4419 | 0.122 | 0.514 | 3737 | 0.824 | 1.000 | 5465 | 0.398 | 0.922 | 6085 | 0.912 | 1.000 | 1947 | 0.470 | 0.922 | 3940 | 1.000 | 1.000 | 2077 | 0.063 | 0.514 | 1644 | 0.567 | 0.922 | 960 | 0.961 | 1.000 | 1001 | 0.502 |
| BRD-K21670884-001-01-9 | cariporide | 683 | 0.348 | 0.915 | 1708 | 0.579 | 0.925 | 575 | 0.139 | 0.730 | 2660 | 0.068 | 0.712 | 1517 | 0.899 | 0.974 | 1359 | 0.922 | 0.974 | 1045 | 0.766 | 0.974 | 1234 | 0.617 | 0.925 | 495 | 0.574 | 0.925 | 1099 | 0.948 | 0.974 | 454 | 0.031 | 0.655 | 216 | 0.070 | 0.211 | 330 | 0.960 | 0.974 | 5 | 0.052 |
| BRD-K21673112-003-01-3 | BF2.649 | 409 | 0.899 | 0.947 | 615 | 0.724 | 0.947 | 326 | 0.812 | 0.947 | 980 | 0.124 | 0.652 | 335 | 0.057 | 0.485 | 157 | 0.872 | 0.947 | 412 | 0.906 | 0.947 | 441 | 0.202 | 0.530 | 310 | 0.947 | 0.947 | 210 | 0.448 | 0.941 | 156 | 0.432 | 1.000 |  |  |  | 99 | 0.389 | 0.908 | 8 | 0.051 |
| BRD-K21680192-300-14-4 | mitoxantrone | 4940 | 0.943 | 0.943 | 7099 | 0.931 | 0.943 | 3117 | 0.099 | 0.558 | 12694 | 0.330 | 0.677 | 4195 | 0.311 | 0.677 | 4095 | 0.928 | 0.943 | 5149 | 0.896 | 0.943 | 5033 | 0.536 | 0.866 | 1566 | 0.303 | 0.677 | 2760 | 0.879 | 0.943 | 1890 | 0.070 | 0.558 | 2481 | 0.675 | 0.886 | 494 | 0.106 | 0.558 | 977 | 0.500 |
| BRD-K21718444-001-06-8 | KW-2449 | 1737 | 0.549 | 0.965 | 3377.5 | 0.643 | 0.976 | 1622 | 0.240 | 1.000 | 6362 | 0.551 | 0.965 | 2799.5 | 0.519 | 1.000 | 1640 | 0.899 | 0.976 | 2691.5 | 0.782 | 0.976 | 2420.5 | 0.764 | 0.976 | 619 | 0.063 | 0.146 | 1886.5 | 0.874 | 0.976 | 712 | 0.019 | 0.408 | 523 | 0.007 | 0.021 | 427 | 0.916 | 0.976 | 124 | 0.019 |
| BRD-K21728777-001-02-3 | AMG900 | 3024 | 0.328 | 0.764 | 6616 | 0.998 | 0.998 | 2490 | 0.048 | 0.169 | 10842 | 0.663 | 0.972 | 3945 | 0.449 | 0.943 | 2581 | 0.924 | 0.998 | 4374 | 0.862 | 0.998 | 3453 | 0.045 | 0.169 | 1123 | 0.028 | 0.169 | 2478 | 0.740 | 0.972 | 1320 | 0.016 | 0.169 | 891 | 0.039 | 0.169 | 642 | 0.731 | 0.972 | 395 | 0.017 |
| BRD-K22025381-003-01-9 | U-18666A | 551.5 | 0.099 | 0.345 | 1000 | 0.786 | 0.970 | 546.5 | 0.342 | 0.652 | 2049 | 0.977 | 0.977 | 349.5 | 0.282 | 0.591 | 450.5 | 0.899 | 0.970 | 355 | 0.571 | 0.970 | 714 | 0.852 | 0.970 | 194.5 | 0.156 | 0.467 | 216.5 | 0.233 | 0.544 | 180 | 0.009 | 0.192 | 335.5 | 0.786 | 0.970 | 196.5 | 0.851 | 0.970 | 385.5 | 0.924 |
| BRD-K22064724-001-01-8 | napabucasin | 4977 | 0.570 | 0.983 | 3485 | 0.000 | 0.000 | 3637 | 0.105 | 0.841 | 14776 | 0.651 | 0.983 | 4118 | 0.157 | 0.841 | 4053 | 0.578 | 0.983 | 6498 | 0.874 | 0.983 | 6654 | 0.918 | 0.983 | 1479 | 0.164 | 0.841 | 3954 | 0.982 | 0.983 | 2744 | 0.621 | 0.983 | 2532 | 0.912 | 0.983 | 465 | 0.240 | 0.841 | 1582 | 0.983 |
| BRD-K22127577-001-03-7 | crenolanib | 5240 | 0.682 | 0.996 | 8310 | 0.996 | 0.996 | 3922 | 0.327 | 0.834 | 13496 | 0.003 | 0.067 | 4990 | 0.196 | 0.602 | 4977 | 0.969 | 0.996 | 6681 | 0.717 | 0.996 | 7012 | 0.690 | 0.996 | 2213 | 0.377 | 0.834 | 2469 | 0.145 | 0.602 | 2218 | 0.055 | 0.289 | 1417 | 0.013 | 0.132 | 648 | 0.201 | 0.602 | 1455 | 0.906 |
| BRD-K22134346-001-24-9 | simvastatin | 1210 | 0.574 | 0.791 | 1958 | 0.270 | 0.791 | 1388 | 0.530 | 0.791 | 4213 | 0.554 | 0.791 | 1555 | 0.480 | 0.791 | 1738 | 0.984 | 0.984 | 1767 | 0.512 | 0.791 | 1702 | 0.182 | 0.753 | 610 | 0.908 | 0.984 | 423 | 0.449 | 0.791 | 692 | 0.074 | 0.753 | 530 | 0.759 | 0.886 | 60 | 0.215 | 0.753 | 379 | 0.667 |
| BRD-K22503835-001-11-0 | scriptaid | 2790 | 0.311 | 0.594 | 5307 | 0.823 | 0.994 | 2972 | 0.880 | 0.994 | 10904.5 | 0.719 | 0.994 | 3419 | 0.053 | 0.159 | 2541 | 0.831 | 0.994 | 4185 | 0.945 | 0.994 | 5483 | 0.994 | 0.994 | 1225 | 0.065 | 0.170 | 2683.5 | 0.993 | 0.994 | 1807 | 0.139 | 1.000 | 503 | 0.016 | 0.081 | 624 | 0.365 | 0.639 | 238 | 0.053 |
| BRD-K23190681-001-01-1 | AV-412 | 740 | 0.862 | 0.990 | 1609 | 0.972 | 0.990 | 882 | 0.047 | 0.330 | 3511 | 0.697 | 0.990 | 1798 | 0.740 | 0.990 | 1337 | 0.941 | 0.990 | 1652 | 0.780 | 0.990 | 1143 | 0.990 | 0.990 | 304 | 0.528 | 0.990 | 621 | 0.842 | 0.990 | 492 | 0.041 | 0.330 | 174 | 0.871 | 0.990 | 21 | 0.109 | 0.383 | 283 | 0.833 |
| BRD-K23204545-001-16-4 | busulfan | 901 | 0.952 | 0.985 | 921 | 0.215 | 0.928 | 782 | 0.600 | 0.742 | 2255 | 0.985 | 0.985 | 604 | 0.268 | 0.928 | 331 | 0.706 | 0.781 | 629 | 0.191 | 0.928 | 308 | 0.309 | 0.928 | 240 | 0.110 | 0.928 | 16 | 0.091 | 0.928 | 498 | 0.531 | 0.742 | 124 | 0.366 | 0.549 | 104 | 0.249 | 0.402 | 42 | 0.043 |
| BRD-K23228615-001-02-8 | GDC-0980 | 2658 | 0.990 | 0.993 | 3953 | 0.988 | 0.993 | 2356 | 0.261 | 0.699 | 7427.5 | 0.712 | 0.959 | 1425 | 0.001 | 0.011 | 1773 | 0.295 | 0.699 | 3011 | 0.797 | 0.959 | 2998 | 0.993 | 0.993 | 804 | 0.031 | 0.216 | 1746 | 0.419 | 0.875 | 759 | 0.007 | 0.071 | 878 | 0.300 | 0.699 | 398 | 0.822 | 0.959 | 438 | 0.518 |
| BRD-K23301018-001-11-7 | broxyquinoline | 1581 | 0.089 | 0.469 | 3196 | 0.016 | 0.284 | 1857 | 0.027 | 0.284 | 6124 | 0.045 | 0.317 | 2777 | 0.606 | 0.987 | 2097 | 0.891 | 0.987 | 3582 | 0.736 | 0.987 | 2526 | 0.477 | 0.987 | 935 | 0.595 | 0.987 | 1715 | 0.987 | 0.987 | 1615 | 0.344 | 0.982 | 741 | 0.803 | 0.987 | 534 | 0.365 | 0.982 | 274 | 0.421 |
| BRD-K23363278-001-02-1 | CYT-997 | 4887 | 0.483 | 0.867 | 8949 | 0.997 | 0.997 | 2508 | 0.033 | 0.228 | 14588 | 0.527 | 0.867 | 5199 | 0.569 | 0.867 | 3957 | 0.762 | 0.889 | 6590 | 0.994 | 0.997 | 5123 | 0.137 | 0.463 | 1327 | 0.030 | 0.228 | 3265 | 0.702 | 0.867 | 2188 | 0.075 | 0.315 | 2116 | 0.219 | 0.463 | 599 | 0.171 | 0.463 | 452 | 0.015 |
| BRD-K23403298-003-01-9 | benzyldimethylhexadecylammonium | 648 | 0.842 | 0.842 | 812.5 | 0.917 | 1.000 | 506 | 0.378 | 1.000 | 1642 | 0.861 | 1.000 | 382.5 | 0.073 | 0.512 | 436.5 | 0.777 | 1.000 | 218 | 0.132 | 0.558 | 823.5 | 0.996 | 1.000 | 134 | 0.197 | 0.348 | 30 | 0.184 | 0.348 | 269 | 0.133 | 0.558 |  |  |  | 32 | 0.199 | 0.348 | 79 | 0.182 |
| BRD-K23677682-003-01-2 | voreloxin | 4441 | 0.501 | 0.912 | 6060 | 0.668 | 0.953 | 3647 | 0.426 | 0.912 | 12188 | 0.218 | 0.807 | 4264 | 0.177 | 0.807 | 3013 | 0.801 | 0.953 | 5710 | 0.907 | 0.953 | 5609 | 0.874 | 0.953 | 1163 | 0.046 | 0.322 | 3096 | 0.565 | 0.912 | 1900 | 0.035 | 0.322 | 799 | 0.009 | 0.184 | 993 | 0.820 | 0.953 | 812 | 0.269 |
| BRD-K23779958-001-05-8 | dichlorvos | 2585 | 0.891 | 0.945 | 2516 | 0.010 | 0.209 | 1834 | 0.265 | 1.000 | 7152 | 0.225 | 1.000 | 3239 | 0.847 | 0.945 | 2192 | 0.685 | 0.899 | 3313 | 0.945 | 0.945 | 3543 | 0.660 | 0.899 | 1359 | 0.915 | 0.945 | 1731 | 0.465 | 0.807 | 1340 | 0.168 | 1.000 | 1222 | 0.337 | 0.807 | 289 | 0.457 | 0.807 | 416 | 0.405 |
| BRD-K23925186-001-02-8 | oridonin | 267 | 0.518 | 0.989 | 1015.5 | 0.031 | 0.278 | 565 | 0.869 | 0.989 | 1630 | 0.807 | 0.989 | 554 | 0.073 | 0.278 | 333 | 0.815 | 0.989 | 490 | 0.857 | 0.989 | 308 | 0.079 | 0.278 | 141 | 0.057 | 0.278 | 390 | 0.948 | 0.989 | 247 | 0.066 | 0.278 | 79 | 0.620 | 0.989 | 56 | 0.390 | 0.989 | 61 | 0.439 |
| BRD-K23984367-001-07-5 | sorafenib | 650 | 0.997 | 0.997 | 564 | 0.709 | 0.992 | 175 | 0.192 | 1.000 | 1289 | 0.151 | 1.000 | 511 | 0.512 | 0.827 | 374 | 0.856 | 0.997 | 546 | 0.826 | 0.997 | 280 | 0.954 | 0.997 | 187 | 0.009 | 0.023 | 83 | 0.230 | 0.538 | 213 | 0.373 | 0.712 | 272 | 0.940 | 0.997 | 48 | 0.366 | 0.712 | 201 | 0.648 |
| BRD-K23984367-075-15-2 | sorafenib | 1210 | 0.860 | 1.000 | 1210 | 0.945 | 1.000 | 620 | 0.070 | 0.395 | 2836 | 0.173 | 0.518 | 1187 | 0.370 | 0.970 | 681 | 0.887 | 1.000 | 890 | 0.128 | 0.448 | 1518 | 0.965 | 1.000 | 165 | 0.127 | 0.448 | 345 | 0.075 | 0.395 | 668 | 0.659 | 1.000 |  |  |  | 298 | 0.630 | 0.945 | 84 | 0.455 |
| BRD-K24187789-001-01-7 | VU0238429 | 3299 | 0.920 | 1.000 | 5410 | 0.930 | 1.000 | 2522 | 0.177 | 0.532 | 9250 | 0.446 | 0.780 | 3283 | 0.315 | 0.701 | 3154 | 0.850 | 1.000 | 4289.5 | 1.000 | 1.000 | 3145.5 | 0.007 | 0.120 | 1350 | 0.316 | 0.701 | 1845 | 0.738 | 1.000 | 1824 | 0.334 | 0.701 | 579 | 0.011 | 0.120 | 874 | 0.850 | 1.000 | 417 | 0.027 |
| BRD-K24556098-001-16-7 | deoxycorticosterone-acetate | 1382 | 0.193 | 0.624 | 3676 | 0.294 | 0.624 | 2376 | 0.262 | 0.624 | 7686 | 0.233 | 0.624 | 3151 | 0.860 | 0.994 | 2602 | 0.641 | 0.841 | 3410 | 0.962 | 0.994 | 4232 | 0.975 | 0.994 | 1067 | 0.297 | 0.624 | 842 | 0.994 | 0.994 | 1432 | 0.061 | 0.688 | 431 | 0.159 | 0.624 | 666 | 0.630 | 0.841 | 570 | 0.416 |
| BRD-K24556407-001-02-3 | ETP-46464 | 4028 | 0.661 | 0.868 | 6596 | 0.942 | 0.996 | 3191 | 0.172 | 0.513 | 11455 | 0.278 | 0.583 | 4171 | 0.072 | 0.477 | 3537 | 0.408 | 0.779 | 6128 | 0.977 | 0.996 | 6119 | 0.996 | 0.996 | 1063 | 0.191 | 0.513 | 2750 | 0.646 | 0.868 | 1524 | 0.030 | 0.477 | 1124 | 0.252 | 0.583 | 505 | 0.129 | 0.513 | 717 | 0.764 |
| BRD-K24576554-001-04-8 | AT-9283 | 1424 | 0.021 | 0.161 | 2888 | 0.997 | 0.997 | 1287 | 0.015 | 0.161 | 6958 | 0.897 | 0.997 | 2308 | 0.266 | 0.699 | 1491 | 0.771 | 0.997 | 2885 | 0.765 | 0.997 | 1813 | 0.221 | 0.699 | 520 | 0.023 | 0.161 | 1748 | 0.593 | 0.958 | 828 | 0.153 | 0.641 | 793 | 0.261 | 0.699 | 249 | 0.919 | 0.997 | 371 | 0.412 |
| BRD-K24593301-001-02-3 | AZD3463 | 3879 | 0.488 | 0.747 | 6231 | 0.755 | 0.933 | 2328 | 0.011 | 0.112 | 11481 | 0.435 | 0.747 | 4195 | 0.478 | 0.747 | 3501 | 0.992 | 0.999 | 4413 | 0.204 | 0.713 | 5985.5 | 0.999 | 0.999 | 1006 | 0.159 | 0.713 | 2388 | 0.530 | 0.747 | 2567 | 0.533 | 0.747 | 1616 | 0.063 | 0.439 | 839 | 0.932 | 0.999 | 559 | 0.197 |
| BRD-K24603946-001-02-1 | 5-hydroxymethyl-tolterodine | 1343 | 0.448 | 0.830 | 2996 | 0.816 | 0.969 | 1545 | 0.284 | 0.830 | 4743 | 0.409 | 0.830 | 1850 | 0.574 | 0.862 | 1288 | 0.541 | 0.862 | 2181 | 0.969 | 0.969 | 1563 | 0.460 | 0.830 | 809 | 0.923 | 0.969 | 1310 | 0.764 | 0.969 | 803 | 0.466 | 0.830 | 101 | 0.081 | 0.388 | 542 | 0.939 | 0.969 | 73 | 0.046 |
| BRD-K24715592-406-02-1 | ethacridine-lactate-monohydrate | 1106 | 0.759 | 0.971 | 2963 | 1.000 | 1.000 | 1369 | 0.820 | 0.971 | 4766 | 0.833 | 0.971 | 1462 | 0.202 | 0.471 | 451 | 0.001 | 0.007 | 2216 | 0.909 | 1.000 | 1736 | 0.024 | 0.114 | 374 | 0.111 | 0.334 | 661 | 0.956 | 1.000 | 571 | 0.051 | 1.000 | 160 | 0.062 | 0.216 | 401 | 0.730 | 0.971 | 306 | 0.819 |
| BRD-K24844714-001-24-5 | 5-fluorouracil | 1214.5 | 0.321 | 0.937 | 1944 | 0.670 | 0.937 | 1729 | 0.502 | 0.937 | 4221.5 | 0.380 | 0.937 | 2259.5 | 0.801 | 0.937 | 1148 | 0.912 | 0.937 | 1969.5 | 0.914 | 0.937 | 1902.5 | 0.466 | 0.937 | 696.5 | 0.547 | 0.937 | 934.5 | 0.583 | 0.937 | 851 | 0.106 | 0.721 | 201 | 0.901 | 0.937 | 297 | 0.763 | 0.937 | 154 | 0.720 |
| BRD-K24869513-001-02-1 | CNX-2006 | 246 | 0.671 | 0.918 | 663 | 0.948 | 0.987 | 1062 | 0.697 | 0.918 | 1510 | 0.181 | 0.946 | 532 | 0.247 | 0.946 | 237 | 0.036 | 0.374 | 1019 | 0.938 | 0.987 | 572 | 0.785 | 0.969 | 377 | 0.606 | 0.918 | 398 | 0.690 | 0.918 | 256 | 0.053 | 0.374 | 345 | 0.470 | 0.918 | 113 | 0.835 | 0.974 | 94 | 0.699 |
| BRD-K25224017-001-25-0 | pirenperone | 4584 | 0.771 | 0.899 | 6100 | 0.497 | 0.820 | 2453 | 0.008 | 0.167 | 13333 | 0.293 | 0.683 | 3926 | 0.045 | 0.314 | 4042 | 0.884 | 0.977 | 7434 | 1.000 | 1.000 | 5379 | 0.698 | 0.896 | 1667 | 0.604 | 0.896 | 3175 | 0.726 | 0.896 | 2731 | 0.508 | 0.820 | 1579 | 0.499 | 0.820 | 936 | 0.718 | 0.896 | 587 | 0.189 |
| BRD-K25244359-066-03-4 | apatinib | 603.5 | 0.790 | 0.790 | 613 | 0.628 | 1.000 | 504.5 | 0.881 | 1.000 | 1381.5 | 0.369 | 1.000 | 508 | 0.700 | 0.918 | 592.5 | 0.559 | 1.000 | 528 | 0.340 | 1.000 | 429.5 | 0.403 | 1.000 | 138.5 | 0.052 | 0.730 | 337 | 0.280 | 1.000 | 592 | 0.557 | 1.000 | 224 | 0.635 | 0.889 | 83 | 0.178 | 0.287 |  |  |
| BRD-K25279140-001-01-5 | MK-8745 | 2956 | 0.817 | 0.977 | 5709 | 0.977 | 0.977 | 3245 | 0.524 | 0.977 | 7135 | 0.001 | 0.029 | 2556 | 0.111 | 0.388 | 2645 | 0.926 | 0.977 | 4510 | 0.967 | 0.977 | 4784 | 0.685 | 0.977 | 1360 | 0.077 | 0.383 | 2111 | 0.816 | 0.977 | 1672 | 0.091 | 0.383 | 585 | 0.039 | 0.383 | 609 | 0.723 | 0.977 | 513 | 0.067 |
| BRD-K25340465-001-11-1 | OSI-930 | 1823 | 0.266 | 0.642 | 2304 | 0.468 | 0.818 | 1690 | 0.139 | 0.596 | 4998 | 0.262 | 0.642 | 1980 | 0.215 | 0.642 | 1578 | 0.704 | 0.985 | 1429 | 0.306 | 0.642 | 2358 | 0.995 | 0.995 | 325 | 0.117 | 0.596 | 1351 | 0.966 | 0.995 | 1026 | 0.100 | 0.596 | 858 | 0.275 | 0.642 | 144 | 0.600 | 0.969 | 561 | 0.669 |
| BRD-K25412176-001-01-9 | GSK2110183 | 2098 | 0.172 | 0.601 | 4944.5 | 0.997 | 0.999 | 2206.5 | 0.262 | 0.687 | 8711 | 0.999 | 0.999 | 1940 | 0.020 | 0.085 | 1540 | 0.006 | 0.029 | 2699.5 | 0.778 | 0.961 | 2720 | 0.386 | 0.728 | 1124.5 | 0.411 | 0.728 | 1869 | 0.480 | 0.728 | 1461 | 0.085 | 0.895 | 469 | 0.230 | 0.687 | 508 | 0.304 | 0.708 | 139 | 0.448 |
| BRD-K25630527-001-03-8 | LY2784544 | 767 | 0.124 | 0.346 | 957 | 0.987 | 0.992 | 754 | 0.760 | 0.992 | 2192 | 0.014 | 0.075 | 1043 | 0.363 | 0.786 | 659 | 0.733 | 0.992 | 884 | 0.951 | 0.992 | 1480 | 0.890 | 0.992 | 423 | 0.806 | 0.992 | 222 | 0.759 | 0.992 | 385 | 0.024 | 0.501 | 224 | 0.374 | 0.786 |  |  |  | 36 | 0.025 |
| BRD-K25835157-001-01-4 | PRT062070 | 446.5 | 0.897 | 0.941 | 139.5 | 0.068 | 1.000 | 387.5 | 0.431 | 0.755 | 1366.5 | 0.855 | 0.941 | 525.5 | 0.207 | 0.621 | 265.5 | 0.941 | 0.941 | 522.5 | 0.924 | 0.941 | 650.5 | 0.303 | 0.636 | 226 | 0.178 | 0.621 | 215 | 0.375 | 0.716 | 242 | 0.243 | 0.636 | 24 | 0.019 | 0.098 | 89.5 | 0.817 | 0.941 | 93 | 0.282 |
| BRD-K26011976-001-01-0 | licochalcone-a | 154 | 0.079 | 0.826 | 825.5 | 0.929 | 0.953 | 122 | 0.286 | 0.668 | 845 | 0.417 | 0.795 | 97 | 0.011 | 0.059 | 364 | 0.837 | 0.953 | 475 | 0.868 | 0.953 | 282 | 0.919 | 0.953 | 121.5 | 0.682 | 0.953 | 172 | 0.953 | 0.953 | 93 | 0.130 | 0.341 | 17 | 0.129 | 0.450 | 160 | 0.917 | 0.953 | 44 | 0.087 |
| BRD-K26026438-001-01-0 | tucatinib | 283 | 0.575 | 0.886 | 398 | 0.997 | 0.997 | 282 | 0.146 | 1.000 | 1053.5 | 0.892 | 0.997 | 437 | 0.217 | 0.650 | 410 | 0.145 | 0.509 | 193 | 0.143 | 0.509 | 264 | 0.962 | 0.997 | 207 | 0.982 | 0.997 | 327 | 0.789 | 0.997 | 258 | 0.212 | 1.000 | 76 | 0.735 | 0.997 | 63 | 0.591 | 0.886 |  |  |
| BRD-K26253904-001-01-5 | NMS-873 | 3927 | 0.768 | 0.849 | 6447 | 0.590 | 0.984 | 2733 | 0.149 | 0.970 | 11781 | 0.185 | 0.970 | 4893 | 0.568 | 0.984 | 4021 | 0.707 | 0.984 | 5905 | 0.667 | 0.984 | 5916 | 0.674 | 0.984 | 1074 | 0.025 | 0.312 | 3327 | 0.865 | 0.865 | 2110 | 0.315 | 0.984 | 1243 | 0.354 | 0.984 | 560 | 0.448 | 0.984 | 908 | 0.400 |
| BRD-K26530649-001-01-6 | carboxypyridine-disulfide | 400 | 0.151 | 0.631 | 357 | 0.365 | 0.852 | 589 | 0.563 | 0.857 | 1790 | 0.481 | 0.857 | 651 | 0.571 | 0.857 | 381 | 0.734 | 0.967 | 768 | 0.991 | 0.991 | 582 | 0.538 | 0.857 | 500 | 0.879 | 0.972 | 402 | 0.807 | 0.967 | 365 | 0.198 | 0.631 | 201 | 0.210 | 0.631 | 132 | 0.951 | 0.991 | 125 | 0.439 |
| BRD-K26531771-001-02-2 | EPZ-5676 | 687 | 0.058 | 1.000 | 2131 | 0.999 | 0.999 | 1440 | 0.824 | 0.999 | 3173 | 0.037 | 0.250 | 1266 | 0.371 | 0.779 | 385 | 0.791 | 0.999 | 1248 | 0.501 | 0.896 | 1567 | 0.596 | 0.938 | 861 | 0.968 | 0.999 | 723 | 0.956 | 0.999 | 619 | 0.066 | 0.276 | 281 | 0.048 | 0.250 | 228 | 0.233 | 0.698 | 165 | 0.327 |
| BRD-K26603252-003-04-9 | PD-153035 | 1006.5 | 0.877 | 0.980 | 1294 | 0.309 | 0.590 | 623 | 0.008 | 0.079 | 3503.5 | 0.914 | 0.980 | 1135 | 0.793 | 0.980 | 993.5 | 0.957 | 0.980 | 1573.5 | 0.954 | 0.980 | 1177.5 | 0.946 | 0.980 | 360 | 0.451 | 0.789 | 510 | 0.645 | 0.980 | 155 | 0.023 | 0.160 | 348.5 | 0.980 | 0.980 |  |  |  | 51 | 0.215 |
| BRD-K26667523-001-02-5 | TAK-733 | 2593 | 0.000 | 0.004 | 3065 | 0.000 | 0.000 | 4175 | 0.581 | 1.000 | 14006 | 0.982 | 1.000 | 5279 | 0.946 | 1.000 | 3608 | 0.965 | 1.000 | 3818 | 0.002 | 0.012 | 7189 | 1.000 | 1.000 | 1979 | 0.578 | 1.000 | 3079 | 0.996 | 1.000 | 2374 | 0.257 | 0.900 | 1362 | 0.942 | 1.000 | 1019 | 0.987 | 1.000 | 724 | 0.410 |
| BRD-K26712974-001-01-5 | VX-765 | 1119 | 0.227 | 0.812 | 2434 | 0.716 | 0.885 | 1877 | 0.706 | 0.885 | 4928 | 0.135 | 0.812 | 1840 | 0.501 | 0.885 | 540 | 0.351 | 0.885 | 2001 | 0.928 | 0.928 | 2568 | 0.629 | 0.885 | 568 | 0.076 | 0.796 | 907 | 0.646 | 0.885 | 1103 | 0.570 | 0.885 | 267 | 0.064 | 0.796 | 526 | 0.902 | 0.928 | 267 | 0.224 |
| BRD-K26756394-001-02-2 | VE-821 | 300 | 0.307 | 0.538 | 612 | 0.547 | 0.841 | 475 | 0.128 | 1.000 | 1279 | 0.916 | 0.961 | 538 | 0.288 | 0.865 | 191 | 0.102 | 0.357 | 188 | 0.095 | 0.357 | 476 | 0.819 | 0.938 | 253 | 0.624 | 0.841 | 291 | 0.120 | 0.253 | 341 | 0.271 | 1.000 | 282 | 0.771 | 0.938 |  |  |  |  |  |
| BRD-K26818574-305-04-3 | BIX-01294 | 2628 | 0.078 | 0.458 | 3092 | 0.000 | 0.000 | 2360 | 0.087 | 0.458 | 12297 | 0.961 | 0.998 | 4177 | 0.355 | 0.771 | 3282 | 0.957 | 0.998 | 3346 | 0.128 | 0.465 | 5692 | 0.998 | 0.998 | 1086 | 0.133 | 0.465 | 3390 | 0.994 | 0.998 | 2476 | 0.542 | 0.876 | 1276 | 0.789 | 0.998 | 764 | 0.889 | 0.998 | 681 | 0.436 |
| BRD-K26823213-001-02-9 | GW-842166 | 202 | 0.977 | 0.977 | 562.5 | 0.072 | 0.708 | 508.5 | 0.841 | 0.977 | 821 | 0.142 | 0.708 | 398.5 | 0.407 | 1.000 | 361 | 0.255 | 0.766 | 453 | 0.956 | 0.977 | 559 | 0.835 | 0.977 | 63 | 0.150 | 0.708 | 117 | 0.197 | 0.708 | 375 | 0.758 | 0.977 | 181.5 | 0.940 | 0.977 | 73 | 0.705 | 0.977 | 44 | 0.362 |
| BRD-K26838195-075-04-9 | AST-1306 | 817 | 0.063 | 0.266 | 1346 | 0.999 | 0.999 | 1050 | 0.131 | 0.387 | 4189 | 0.974 | 0.999 | 945 | 0.524 | 0.846 | 1031 | 0.341 | 0.651 | 764 | 0.166 | 0.387 | 1634 | 0.997 | 0.999 | 396 | 0.254 | 0.532 | 730 | 0.874 | 0.999 | 675 | 0.154 | 0.387 | 161 | 0.868 | 0.999 | 139 | 0.767 | 0.999 | 9 | 0.060 |
| BRD-K26997899-001-02-2 | D-64131 | 2720 | 0.221 | 1.000 | 4076 | 0.891 | 0.935 | 1999 | 0.271 | 0.633 | 7527 | 0.185 | 0.633 | 2960 | 0.436 | 0.763 | 1935 | 0.846 | 0.935 | 2953 | 0.818 | 0.935 | 2993 | 0.343 | 0.694 | 1179 | 0.259 | 0.633 | 2510 | 0.999 | 0.999 | 1832 | 0.747 | 0.935 | 471 | 0.075 | 0.392 | 580 | 0.775 | 0.935 | 297 | 0.155 |
| BRD-K27182532-001-02-3 | metatinib | 2621 | 0.995 | 0.995 | 4285 | 0.876 | 0.986 | 1898 | 0.040 | 0.555 | 8325 | 0.490 | 0.986 | 3018 | 0.130 | 0.555 | 2201 | 0.764 | 0.986 | 3636 | 0.956 | 0.995 | 2953 | 0.249 | 0.653 | 633 | 0.239 | 0.653 | 1759 | 0.790 | 0.986 | 2033 | 0.579 | 0.986 | 682 | 0.132 | 0.555 | 403 | 0.338 | 0.788 | 479 | 0.525 |
| BRD-K27184429-300-01-3 | levocetirizine | 404.5 | 0.578 | 0.864 | 839 | 0.143 | 1.000 | 620.5 | 0.773 | 0.864 | 2584 | 0.691 | 0.864 | 1220.5 | 0.531 | 1.000 | 379 | 0.478 | 1.000 | 908 | 0.740 | 0.864 | 798.5 | 0.823 | 0.864 | 63 | 0.003 | 0.009 | 7 | 0.059 | 0.139 | 284 | 0.160 | 1.000 | 196.5 | 0.732 | 0.864 | 26 | 0.126 | 0.265 | 296.5 | 0.788 |
| BRD-K27217864-001-22-3 | butamben | 755 | 0.937 | 0.961 | 1497 | 0.936 | 0.961 | 757 | 0.810 | 0.961 | 2312 | 0.843 | 0.961 | 592 | 0.000 | 0.008 | 543 | 0.437 | 0.961 | 1061 | 0.462 | 0.961 | 994 | 0.641 | 0.961 | 468 | 0.716 | 0.961 | 620 | 0.875 | 0.961 | 397 | 0.224 | 0.781 | 58 | 0.058 | 0.306 | 105 | 0.675 | 0.961 | 29 | 0.132 |
| BRD-K27316855-001-13-3 | calcitriol | 492 | 0.017 | 0.201 | 1572 | 0.826 | 0.957 | 1106 | 0.480 | 0.957 | 3521 | 0.355 | 0.957 | 991 | 0.615 | 0.957 | 444 | 0.691 | 0.957 | 1588 | 0.446 | 0.957 | 1244 | 0.879 | 0.957 | 148 | 0.809 | 0.957 | 1032 | 0.957 | 0.957 | 254 | 0.036 | 0.250 | 402 | 0.868 | 0.957 | 279 | 0.447 | 0.957 | 350 | 0.722 |
| BRD-K28061410-001-06-6 | beta-lapachone | 5329 | 0.345 | 0.871 | 4929 | 0.000 | 0.006 | 4365 | 0.159 | 0.724 | 19048 | 0.904 | 0.949 | 7569 | 0.996 | 0.996 | 4873 | 0.566 | 0.871 | 7105 | 0.768 | 0.949 | 7918 | 0.901 | 0.949 | 1840 | 0.172 | 0.724 | 3476 | 0.467 | 0.871 | 3190 | 0.408 | 0.871 | 2444 | 0.699 | 0.949 | 959 | 0.581 | 0.871 | 1090 | 0.451 |
| BRD-K28192623-066-08-1 | spiradoline | 3358 | 0.849 | 0.963 | 4809 | 0.576 | 0.935 | 1927 | 0.017 | 0.347 | 9878 | 0.493 | 0.935 | 3060 | 0.227 | 0.680 | 2962 | 0.963 | 0.963 | 4824 | 0.922 | 0.963 | 4139 | 0.738 | 0.935 | 1251 | 0.370 | 0.935 | 2388 | 0.879 | 0.963 | 1545 | 0.042 | 0.446 | 1151 | 0.113 | 0.493 | 319 | 0.436 | 0.935 | 339 | 0.158 |
| BRD-K28352084-001-02-3 | CH5132799 | 3228 | 0.734 | 0.936 | 4981 | 0.989 | 0.991 | 2065 | 0.091 | 0.476 | 8684 | 0.758 | 0.936 | 3242 | 0.362 | 0.752 | 2188 | 0.737 | 0.936 | 3089 | 0.238 | 0.712 | 3905 | 0.990 | 0.991 | 664 | 0.287 | 0.712 | 2033 | 0.991 | 0.991 | 1263 | 0.013 | 0.141 | 751 | 0.441 | 0.752 | 614 | 0.806 | 0.940 | 246 | 0.305 |
| BRD-K28360340-001-08-3 | TW-37 | 4139 | 0.042 | 0.221 | 6978 | 0.302 | 0.804 | 4108 | 0.404 | 0.848 | 14946 | 0.498 | 0.876 | 5630 | 0.662 | 0.876 | 4629 | 0.930 | 0.977 | 6969 | 0.995 | 0.995 | 5773 | 0.306 | 0.804 | 1294 | 0.007 | 0.153 | 3129 | 0.248 | 0.804 | 2446 | 0.170 | 0.716 | 2466 | 0.846 | 0.977 | 951 | 0.667 | 0.876 | 534 | 0.029 |
| BRD-K28383148-001-01-4 | AP1903 | 513 | 0.312 | 0.727 | 607 | 0.058 | 0.404 | 486 | 0.020 | 0.212 | 1762 | 0.753 | 0.975 | 885 | 0.975 | 0.975 | 328 | 0.657 | 0.975 | 860 | 0.573 | 0.975 | 690 | 0.659 | 0.975 | 336 | 0.939 | 0.975 | 297 | 0.845 | 0.975 | 341 | 0.184 | 0.948 | 384 | 0.855 | 0.975 |  |  |  | 4 | 0.051 |
| BRD-K28537285-001-01-8 | chidamide | 1053 | 0.828 | 0.915 | 3738 | 0.998 | 0.998 | 1612 | 0.524 | 0.739 | 4279 | 0.340 | 0.739 | 1340 | 0.255 | 0.739 | 984 | 0.302 | 0.739 | 625 | 0.400 | 0.739 | 925 | 0.598 | 0.739 | 394 | 0.044 | 0.465 | 554 | 0.492 | 0.739 | 1282 | 0.476 | 0.739 | 747 | 0.440 | 0.739 | 385 | 0.664 | 0.774 | 240 | 0.196 |
| BRD-K28578425-001-07-0 | cilostamide | 464 | 0.705 | 0.983 | 622 | 0.766 | 0.983 | 297 | 0.016 | 0.115 | 1597 | 0.475 | 0.866 | 934 | 0.861 | 0.983 | 339 | 0.775 | 0.983 | 835 | 0.797 | 0.983 | 683 | 0.034 | 0.179 | 173 | 0.303 | 0.866 | 174 | 0.308 | 0.866 | 256 | 0.396 | 0.866 | 132 | 0.932 | 0.983 | 132 | 0.932 | 0.983 | 150 | 0.201 |
| BRD-K28822270-001-03-7 | resminostat | 2155 | 0.909 | 0.996 | 2966 | 0.662 | 0.870 | 1956 | 0.284 | 0.595 | 5574 | 0.242 | 0.595 | 1703 | 0.043 | 0.311 | 2010.5 | 0.991 | 0.996 | 2532 | 0.996 | 0.996 | 2222.5 | 0.943 | 0.996 | 735 | 0.079 | 0.311 | 1774 | 0.924 | 0.996 | 1286 | 0.494 | 0.864 | 1230 | 0.612 | 0.864 | 199 | 0.069 | 0.311 | 107 | 0.015 |
| BRD-K28824103-001-02-8 | genipin | 3309 | 0.611 | 0.916 | 5035 | 0.915 | 0.997 | 2331 | 0.395 | 1.000 | 9544 | 0.275 | 1.000 | 3667 | 0.083 | 0.876 | 1505 | 0.175 | 1.000 | 4815 | 1.000 | 1.000 | 4588 | 0.860 | 0.997 | 888 | 0.007 | 0.015 | 1380 | 0.359 | 0.629 | 1710 | 0.225 | 1.000 | 1000 | 0.121 | 0.232 | 615 | 0.735 | 0.997 | 488 | 0.466 |
| BRD-K28907958-001-04-3 | CD-437 | 3951 | 0.066 | 0.430 | 7697 | 0.930 | 0.930 | 3275 | 0.070 | 0.430 | 17003 | 0.878 | 0.930 | 4508 | 0.075 | 0.430 | 5180 | 0.901 | 0.930 | 5930 | 0.598 | 0.930 | 5692 | 0.344 | 0.802 | 1832 | 0.130 | 0.462 | 3260 | 0.792 | 0.930 | 2795 | 0.417 | 0.819 | 2019 | 0.132 | 0.462 | 1082 | 0.825 | 0.930 | 1049 | 0.492 |
| BRD-K28965160-001-01-1 | BVD-523 | 1614 | 0.143 | 0.458 | 1950 | 0.004 | 0.081 | 1233 | 0.506 | 0.993 | 6397 | 0.930 | 0.993 | 2856 | 0.974 | 0.993 | 1329 | 0.841 | 0.993 | 1622 | 0.043 | 0.320 | 1433 | 0.640 | 0.993 | 459 | 0.153 | 0.458 | 1300.5 | 0.987 | 0.993 | 996.5 | 0.072 | 0.380 | 867 | 0.993 | 0.993 | 415 | 0.939 | 0.993 | 140 | 0.138 |
| BRD-K29254801-001-14-7 | thiram | 3425 | 0.947 | 0.968 | 4190 | 0.322 | 0.778 | 2667 | 0.258 | 0.778 | 11060 | 0.699 | 0.918 | 2020 | 0.006 | 0.119 | 1743 | 0.123 | 0.778 | 4026 | 0.654 | 0.918 | 4060 | 0.561 | 0.906 | 1032 | 0.290 | 0.778 | 2816 | 0.918 | 0.968 | 2316 | 0.681 | 0.918 | 939 | 0.027 | 0.282 | 537 | 0.557 | 0.906 | 451 | 0.371 |
| BRD-K29415052-050-04-8 | BGT226 | 6585 | 0.995 | 0.997 | 9930 | 0.997 | 0.997 | 3966 | 0.077 | 0.567 | 15464 | 0.364 | 0.670 | 3984 | 0.081 | 0.567 | 3853 | 0.160 | 0.570 | 6671 | 0.767 | 0.997 | 7011 | 0.645 | 0.903 | 2097 | 0.257 | 0.670 | 2443 | 0.059 | 0.567 | 3101 | 0.161 | 0.570 | 1959 | 0.163 | 0.570 | 1200 | 0.895 | 0.997 | 1466 | 0.903 |
| BRD-K29542628-001-03-9 | 1-naphthyl-PP1 | 879 | 0.416 | 0.873 | 1456.5 | 0.308 | 1.000 | 1344.5 | 0.583 | 0.874 | 4711.5 | 0.967 | 0.967 | 1573 | 0.791 | 0.874 | 1136 | 0.551 | 0.874 | 1350 | 0.886 | 0.930 | 1415 | 0.152 | 0.402 | 372 | 0.706 | 0.874 | 940.5 | 0.551 | 0.874 | 816 | 0.287 | 1.000 | 384.5 | 0.746 | 0.874 |  |  |  | 206 | 0.153 |
| BRD-K29656036-001-02-5 | MK-8245 | 1710 | 0.349 | 0.666 | 2604 | 0.893 | 0.984 | 1957 | 0.699 | 0.957 | 4466 | 0.168 | 1.000 | 1428 | 0.247 | 1.000 | 2027 | 0.985 | 0.985 | 1518 | 0.234 | 0.491 | 1993 | 0.937 | 0.984 | 390 | 0.600 | 0.899 | 857 | 0.806 | 0.957 | 1371 | 0.182 | 1.000 | 214 | 0.114 | 0.300 | 166 | 0.742 | 0.957 | 247 | 0.176 |
| BRD-K29905972-001-06-3 | axitinib | 634.5 | 0.682 | 0.990 | 295.5 | 0.915 | 1.000 | 576.5 | 0.302 | 0.990 | 1713 | 0.686 | 0.990 | 658.5 | 0.377 | 0.990 | 396.5 | 0.950 | 1.000 | 765.5 | 0.988 | 1.000 | 385 | 0.109 | 0.572 | 190.5 | 0.001 | 0.027 | 190 | 0.430 | 0.990 | 436 | 0.883 | 0.883 | 245 | 0.057 | 0.400 | 68 | 0.500 | 0.990 | 379 | 0.705 |
| BRD-K29950728-048-17-4 | clomifene | 944 | 0.002 | 0.032 | 1769 | 0.387 | 0.836 | 1317 | 0.367 | 0.836 | 4734 | 0.987 | 0.997 | 1244 | 0.398 | 0.836 | 1101 | 0.481 | 0.838 | 1057 | 0.066 | 0.347 | 1038 | 0.997 | 0.997 | 659 | 0.602 | 0.859 | 728 | 0.519 | 0.838 | 725 | 0.759 | 0.947 | 210 | 0.955 | 0.997 | 57 | 0.215 | 0.836 | 290 | 0.820 |
| BRD-K29968218-001-03-2 | KX2-391 | 4250 | 0.446 | 0.817 | 6933 | 0.955 | 0.998 | 4300 | 0.615 | 0.888 | 12140 | 0.149 | 0.625 | 4974 | 0.704 | 0.888 | 3747 | 0.747 | 0.888 | 6496.5 | 0.998 | 0.998 | 4237 | 0.141 | 0.625 | 1793 | 0.761 | 0.888 | 2964 | 0.576 | 0.888 | 2514 | 0.335 | 0.817 | 1058 | 0.021 | 0.219 | 711 | 0.366 | 0.817 | 444 | 0.017 |
| BRD-K30431096-004-01-3 | domiphen | 356 | 0.591 | 0.955 | 1239 | 0.892 | 0.958 | 575 | 0.144 | 0.504 | 2937.5 | 0.587 | 1.000 | 968.5 | 0.136 | 0.504 | 632.5 | 0.253 | 0.760 | 1083.5 | 0.901 | 0.958 | 1013.5 | 0.801 | 0.958 | 415 | 0.502 | 1.000 | 529.5 | 0.854 | 0.958 | 228 | 0.044 | 0.504 | 410.5 | 0.784 | 0.958 |  |  |  | 13 | 0.074 |
| BRD-K30480208-001-18-5 | torasemide | 203 | 0.182 | 0.660 | 363 | 0.283 | 0.660 | 227 | 0.118 | 0.617 | 1269 | 0.923 | 1.000 | 378 | 0.088 | 0.614 | 194 | 0.374 | 0.786 | 555 | 0.706 | 1.000 | 146 | 0.040 | 0.546 | 150 | 0.414 | 0.790 | 87 | 0.854 | 1.000 | 419 | 0.922 | 1.000 | 32 | 0.233 | 0.660 |  |  |  | 38 | 0.052 |
| BRD-K30577245-001-05-0 | docetaxel | 1403 | 0.868 | 0.964 | 2891 | 0.910 | 0.964 | 1539.5 | 0.007 | 0.134 | 4748 | 0.029 | 0.205 | 2094 | 0.482 | 0.964 | 2120.5 | 0.964 | 0.964 | 2525 | 0.964 | 0.964 | 2386.5 | 0.700 | 0.964 | 963 | 0.381 | 0.964 | 973.5 | 0.856 | 0.964 | 1686.5 | 0.743 | 0.964 | 784 | 0.276 | 0.964 | 493 | 0.777 | 0.964 | 214 | 0.326 |
| BRD-K30577245-341-01-9 | docetaxel | 7857 | 0.999 | 1.000 | 8609 | 0.805 | 1.000 | 4055 | 0.067 | 0.280 | 15154 | 0.051 | 0.269 | 5911 | 0.453 | 0.792 | 5495 | 0.884 | 1.000 | 7147 | 0.785 | 1.000 | 6824 | 0.522 | 0.844 | 2345 | 0.240 | 0.629 | 5672 | 1.000 | 1.000 | 2817 | 0.164 | 0.576 | 2534 | 0.387 | 0.792 | 851 | 0.423 | 0.792 | 1274 | 0.687 |
| BRD-K30867024-003-03-3 | SB-216641 | 1072 | 0.351 | 0.717 | 2392 | 0.834 | 0.976 | 1185 | 0.556 | 0.898 | 5867 | 0.976 | 0.977 | 966 | 0.001 | 0.004 | 1324 | 0.784 | 0.976 | 2466 | 0.696 | 0.976 | 2337 | 0.977 | 0.977 | 435 | 0.163 | 0.490 | 1168 | 0.335 | 0.717 | 955 | 0.333 | 0.717 | 355 | 0.495 | 0.867 | 394 | 0.938 | 0.977 | 433 | 0.728 |
| BRD-K30933884-001-01-0 | UNBS-5162 | 1498 | 0.512 | 0.901 | 3733 | 0.847 | 0.994 | 2263 | 0.904 | 0.994 | 5852 | 0.154 | 0.485 | 2304 | 0.185 | 0.485 | 1721 | 0.808 | 0.994 | 2774 | 0.994 | 0.994 | 2551 | 0.561 | 0.907 | 670 | 0.031 | 0.214 | 1212 | 0.474 | 0.901 | 933 | 0.007 | 0.137 | 607 | 0.048 | 0.251 | 468 | 0.639 | 0.959 | 559 | 0.509 |
| BRD-K31086665-005-04-6 | NH125 | 2682 | 0.686 | 0.945 | 5234 | 0.148 | 0.779 | 2940.5 | 0.304 | 1.000 | 13055.5 | 0.928 | 0.945 | 3570 | 0.075 | 0.528 | 3529 | 0.793 | 0.945 | 4922 | 0.804 | 0.945 | 5828 | 0.945 | 0.945 | 959 | 0.006 | 0.014 | 2267 | 0.629 | 0.945 | 2092 | 0.274 | 1.000 | 937 | 0.542 | 0.945 | 260 | 0.058 | 0.122 | 836 | 0.939 |
| BRD-K31309378-001-01-5 | lucitanib | 983 | 0.883 | 0.999 | 2219 | 0.999 | 0.999 | 1081 | 0.530 | 0.845 | 3575 | 0.672 | 0.845 | 738 | 0.018 | 0.186 | 1170 | 0.951 | 0.999 | 1264 | 0.532 | 0.845 | 1380 | 0.177 | 0.618 | 173 | 0.006 | 0.133 | 654 | 0.401 | 0.845 | 577 | 0.040 | 0.280 | 279 | 0.409 | 0.845 | 199 | 0.483 | 0.845 | 235 | 0.651 |
| BRD-K31476763-001-01-5 | AZD9668 | 249 | 0.730 | 0.974 | 721 | 0.520 | 1.000 | 383 | 0.206 | 0.721 | 1470 | 0.087 | 0.552 | 576 | 0.070 | 0.552 | 387 | 0.704 | 1.000 | 1061 | 0.984 | 0.984 | 751 | 0.612 | 1.000 | 262 | 0.789 | 0.974 | 447 | 0.670 | 1.000 | 345 | 0.270 | 0.809 | 129 | 0.132 | 0.552 | 121 | 0.900 | 0.984 | 15 | 0.088 |
| BRD-K31491153-003-05-9 | 1-phenylbiguanide | 260 | 0.103 | 0.434 | 905 | 0.363 | 0.847 | 748 | 0.093 | 0.434 | 2439 | 0.658 | 0.860 | 880 | 0.465 | 0.860 | 706 | 0.918 | 0.963 | 971 | 0.837 | 0.925 | 731 | 0.637 | 0.860 | 194 | 0.706 | 0.860 | 442 | 0.696 | 0.860 | 281 | 0.016 | 0.344 | 10 | 0.067 | 0.434 | 196 | 0.716 | 0.860 | 26 | 0.125 |
| BRD-K31812033-003-11-0 | levobunolol | 602.5 | 0.132 | 1.000 | 1171 | 0.505 | 0.873 | 676 | 0.134 | 1.000 | 2939.5 | 0.594 | 0.873 | 1283 | 0.728 | 0.873 | 654 | 0.398 | 0.760 | 1272 | 0.709 | 0.873 | 952 | 0.222 | 0.582 | 304 | 0.316 | 0.738 | 699.5 | 0.748 | 0.873 | 637 | 0.353 | 0.742 | 578.5 | 0.666 | 0.873 |  |  |  | 252 | 0.841 |

| broad_id | name | al_nervous_system_PVAL_ADJ | fibroblast_MWU_STAT | fibroblast_PVAL | fibroblast_PVAL_ADJ | bile_duct_MWU_STAT | bile_duct_PVAL | bile_duct_PVAL_ADJ | thyroid_MWU_STAT | thyroid_PVAL | thyroid_PVAL_ADJ | upper_aerodigestive_MWU_STAT | upper_aerodigestive_PVAL | upper_aerodigestive_PVAL_ADJ | uterus_MWU_STAT | uterus_PVAL | uterus_PVAL_ADJ | liver_MWU_STAT | liver_PVAL | liver_PVAL_ADJ | esophagus_MWU_STAT | esophagus_PVAL | esophagus_PVAL_ADJ |
|---|---|---|---|---|---|---|---|---|---|---|---|---|---|---|---|---|---|---|---|---|---|---|---|
| BRD-K20290250-001-02-1 | BIX-02188 | 1.000 | | | | 10 | 0.063 | 0.351 | 433 | 0.211 | 0.558 | 1207.5 | 0.927 | 1.000 | 835 | 0.240 | 0.561 | 263 | 0.176 | 0.558 | 616 | 0.921 | 0.921 |
| BRD-K20463415-300-01-4 | LY2606368 | 0.059 | 102 | 0.178 | 0.540 | 1674 | 0.679 | 0.911 | 1245 | 0.032 | 0.341 | 6065 | 0.998 | 0.998 | 3888 | 0.180 | 0.540 | 2758 | 0.312 | 0.724 | 4182 | 0.345 | 0.724 |
| BRD-K20468903-001-01-6 | BNC105 | 0.423 | 20 | 0.049 | 0.304 | 1581 | 0.849 | 0.994 | 1276 | 0.049 | 0.304 | 3682 | 0.231 | 0.479 | 3429 | 0.193 | 0.451 | 2476 | 0.093 | 0.346 | 4642 | 0.748 | 0.994 |
| BRD-K20482099-001-09-4 | rutin | 0.381 | | | | 275.5 | 0.334 | 0.637 | 777 | 0.824 | 0.971 | 1344 | 0.696 | 0.971 | 1236.5 | 0.502 | 0.811 | 254 | 0.082 | 0.351 | 902 | 0.123 | 0.381 |
| BRD-K20545304-001-01-6 | bardoxolone | 0.301 | 184 | 0.706 | 0.982 | 635 | 0.779 | 0.982 | 516 | 0.087 | 0.306 | 1004 | 0.027 | 0.161 | 1469 | 0.453 | 0.951 | 1097 | 0.125 | 0.328 | 1148 | 0.318 | 0.741 |
| BRD-K20605374-001-05-4 | NSC-697923 | 0.557 | 20 | 0.053 | 0.291 | 900 | 0.934 | 0.980 | 1212 | 0.055 | 0.291 | 3496 | 0.174 | 0.557 | 3067 | 0.220 | 0.577 | 1827 | 0.047 | 0.291 | 4451 | 0.585 | 0.878 |
| BRD-K20655524-003-12-0 | mefexamide | 0.528 | 295 | 0.708 | 0.953 | 1365 | 0.667 | 0.953 | 2159 | 0.624 | 0.953 | 2960 | 0.027 | 0.279 | 3946 | 0.475 | 0.906 | 3372 | 0.958 | 1.000 | 4353 | 0.369 | 0.774 |
| BRD-K20722021-001-02-1 | CEP-32496 | 0.996 | | | | 161 | 0.910 | 0.996 | 209 | 0.077 | 0.621 | 863 | 0.178 | 0.936 | 377 | 0.280 | 1.000 | 302 | 0.640 | 0.996 | 1401 | 0.766 | 0.996 |
| BRD-K20958582-001-01-4 | TH-302 | 0.892 | | | | 321 | 0.645 | 0.985 | 594 | 0.121 | 0.483 | 2059.5 | 0.179 | 0.536 | 2249.5 | 0.798 | 0.985 | 575 | 0.103 | 0.483 | 2491 | 0.880 | 0.880 |
| BRD-K21025364-001-05-6 | NVP-TAE226 | 0.282 | 306 | 0.711 | 0.987 | 1291 | 0.517 | 0.987 | 1993 | 0.799 | 0.987 | 3142 | 0.018 | 0.191 | 4436 | 0.719 | 0.987 | 2557 | 0.211 | 0.619 | 3819 | 0.113 | 0.594 |
| BRD-K21283037-001-21-5 | riluzole | 0.587 | | | | 141.5 | 0.415 | 1.000 | 57 | 0.140 | 0.982 | 71 | 0.014 | 0.284 | 373.5 | 0.432 | 1.000 | 212 | 0.571 | 1.000 | 72 | 0.059 | 0.615 |
| BRD-K21361524-001-01-1 | selinexor | 0.866 | 77 | 0.165 | 0.495 | 2114 | 0.909 | 0.995 | 1836 | 0.132 | 0.484 | 4145 | 0.096 | 0.484 | 4445 | 0.312 | 0.614 | 4631 | 0.965 | 0.995 | 4634 | 0.124 | 0.484 |
| BRD-K21396683-001-04-8 | marimastat | 0.922 | 285 | 0.706 | 0.989 | 1185 | 0.756 | 0.993 | 978 | 0.090 | 0.514 | 4011 | 0.624 | 0.936 | 2734 | 0.116 | 0.514 | 3314 | 0.910 | 1.000 | 4150 | 0.453 | 0.922 |
| BRD-K21670884-001-01-9 | cariporide | 0.182 | | | | 181 | 0.936 | 0.974 | 474 | 0.506 | 0.925 | 964 | 0.139 | 0.730 | 712 | 0.974 | 0.974 | 619 | 0.405 | 0.925 | 828 | 0.490 | 0.925 |
| BRD-K21673112-003-01-3 | BF2.649 | 0.153 | | | | 157 | 0.872 | 0.947 | 50 | 0.069 | 0.485 | 281 | 0.041 | 0.485 | 310 | 0.947 | 0.947 | 234 | 0.919 | 0.947 | 271 | 0.509 | 0.891 |
| BRD-K21680192-300-14-4 | mitoxantrone | 0.866 | 54 | 0.139 | 0.583 | 1076 | 0.647 | 0.886 | 1759 | 0.317 | 0.677 | 3946 | 0.646 | 0.886 | 2428 | 0.043 | 0.558 | 2695 | 0.355 | 0.677 | 3300 | 0.178 | 0.625 |
| BRD-K21718444-001-06-8 | KW-2449 | 0.050 | | | | 803 | 0.951 | 0.976 | 1348 | 0.976 | 0.976 | 1804 | 0.111 | 1.000 | 1255 | 0.389 | 1.000 | 497 | 0.389 | 0.817 | 1849.5 | 0.700 | 0.976 |
| BRD-K21728777-001-02-3 | AMG900 | 0.169 | 215 | 0.640 | 0.972 | 737 | 0.557 | 0.972 | 743 | 0.107 | 0.321 | 3238 | 0.257 | 0.675 | 2885 | 0.960 | 0.998 | 2122 | 0.552 | 0.972 | 4525 | 0.961 | 0.998 |
| BRD-K22025381-003-01-9 | U-18666A | 0.970 | | | | 159.5 | 0.647 | 0.970 | 33 | 0.178 | 0.467 | 496 | 0.095 | 0.345 | 605.5 | 0.751 | 0.970 | 24 | 0.020 | 0.140 | 214.5 | 0.076 | 0.345 |
| BRD-K22064724-001-01-8 | napabucasin | 0.983 | 309 | 0.747 | 0.983 | 868 | 0.284 | 0.852 | 1715 | 0.816 | 0.983 | 3863 | 0.201 | 0.841 | 3962 | 0.884 | 0.983 | 2917 | 0.889 | 0.983 | 4025 | 0.557 | 0.983 |
| BRD-K22127577-001-03-7 | crenolanib | 0.996 | 13 | 0.032 | 0.224 | 1677 | 0.892 | 0.996 | 1866 | 0.437 | 0.834 | 5021 | 0.876 | 0.996 | 5058 | 0.978 | 0.996 | 3724 | 0.764 | 0.996 | 4613 | 0.401 | 0.834 |
| BRD-K22134346-001-24-9 | simvastatin | 0.823 | 47 | 0.161 | 0.753 | 127 | 0.150 | 0.753 | 572 | 0.301 | 0.791 | 1187 | 0.349 | 0.791 | 1294 | 0.957 | 0.984 | 803 | 0.603 | 0.791 | 602 | 0.176 | 0.753 |
| BRD-K22503835-001-11-0 | scriptaid | 0.159 | | | | 1214 | 0.767 | 0.994 | 1889 | 0.966 | 0.994 | 2461 | 0.291 | 0.594 | 1371 | 0.000 | 0.010 | 1540 | 0.034 | 0.142 | 3034 | 0.266 | 0.594 |
| BRD-K23190681-001-01-1 | AV-412 | 0.990 | | | | 56 | 0.284 | 0.744 | 243 | 0.697 | 0.990 | 547 | 0.000 | 0.000 | 695 | 0.074 | 0.388 | 677 | 0.740 | 0.990 | 1533 | 0.209 | 0.627 |
| BRD-K23204545-001-16-4 | busulfan | 0.075 | | | | 127 | 0.385 | 1.000 | 231 | 0.584 | 0.742 | 481 | 0.469 | 1.000 | 233 | 0.266 | 0.928 | 169 | 0.659 | 0.769 | 802 | 0.479 | 1.000 |
| BRD-K23228615-001-02-8 | GDC-0980 | 0.906 | 175 | 0.642 | 0.959 | 505 | 0.694 | 0.959 | 450 | 0.230 | 0.699 | 2364 | 0.268 | 0.699 | 1643.5 | 0.458 | 0.875 | 906 | 0.168 | 0.699 | 2013 | 0.757 | 0.959 |
| BRD-K23301018-001-11-7 | broxyquinoline | 0.982 | | | | 754 | 0.823 | 0.987 | 1115 | 0.408 | 0.982 | 2693 | 0.889 | 0.987 | 2155 | 0.822 | 0.987 | 2134 | 0.972 | 0.987 | 3097 | 0.938 | 0.987 |
| BRD-K23363278-001-02-1 | CYT-997 | 0.228 | 115 | 0.220 | 0.463 | 1570 | 0.977 | 0.997 | 1264 | 0.054 | 0.284 | 3954 | 0.625 | 0.867 | 3524 | 0.194 | 0.463 | 3246 | 0.687 | 0.867 | 4498 | 0.602 | 0.867 |
| BRD-K23403298-003-01-9 | benzyldimethylhexadecylammonium | 0.348 | | | | 214 | 0.952 | 1.000 | 68 | 0.007 | 0.143 | 190 | 0.023 | 0.240 | 406 | 0.224 | 0.786 | 242 | 0.986 | 1.000 | 402 | 0.399 | 0.645 |
| BRD-K23677682-003-01-2 | voreloxin | 0.807 | 194 | 0.495 | 0.912 | 1432 | 0.966 | 0.966 | 1463 | 0.402 | 0.912 | 3967 | 0.892 | 0.953 | 3075 | 0.257 | 0.807 | 2707 | 0.545 | 0.912 | 4469 | 0.862 | 0.953 |
| BRD-K23779958-001-05-8 | dichlorvos | 0.807 | | | | 488 | 0.592 | 0.889 | 1007 | 0.439 | 1.000 | 2366 | 0.409 | 1.000 | 1802 | 0.741 | 0.915 | 1602 | 0.480 | 0.807 | 2582 | 0.499 | 0.807 |
| BRD-K23925186-001-02-8 | oridonin | 0.989 | 2 | 0.049 | 0.278 | 259.5 | 0.989 | 0.989 | 234 | 0.698 | 0.989 | 775 | 0.887 | 0.989 | 209 | 0.241 | 0.724 | 186 | 0.829 | 0.989 | 518 | 0.913 | 0.989 |
| BRD-K23984367-001-07-5 | sorafenib | 0.973 | | | | | | | 23 | 0.146 | 1.000 | 435 | 0.968 | 0.997 | 392 | 0.456 | 0.798 | 134 | 0.006 | 0.017 | 246 | 0.276 | 1.000 |
| BRD-K23984367-075-15-2 | sorafenib | 0.734 | | | | 236 | 0.777 | 1.000 | 274 | 0.023 | 0.395 | 974 | 0.898 | 1.000 | 336 | 0.066 | 0.395 | 429 | 0.760 | 1.000 | 877 | 0.894 | 0.894 |
| BRD-K24187789-001-01-7 | VU0238429 | 0.173 | 18 | 0.062 | 0.260 | 675 | 0.507 | 0.819 | 1021 | 0.132 | 0.463 | 2838 | 0.442 | 0.780 | 1886 | 0.033 | 0.173 | 2007 | 0.732 | 1.000 | 3309 | 0.969 | 1.000 |
| BRD-K24556098-001-16-7 | deoxycorticosterone-acetate | 0.728 | | | | 397 | 0.348 | 0.664 | 1282 | 0.630 | 0.841 | 1934 | 0.065 | 0.688 | 1871 | 0.283 | 0.624 | 1385 | 0.772 | 0.954 | 2702 | 0.600 | 0.841 |
| BRD-K24556407-001-02-3 | ETP-46464 | 0.892 | 34 | 0.091 | 0.477 | 821 | 0.602 | 0.868 | 1401 | 0.604 | 0.868 | 3807 | 0.758 | 0.892 | 3499 | 0.519 | 0.868 | 2006 | 0.074 | 0.477 | 3396 | 0.195 | 0.513 |
| BRD-K24576554-001-04-8 | AT-9283 | 0.818 | 100 | 0.371 | 0.818 | 484 | 0.715 | 0.997 | 650 | 0.462 | 0.818 | 1414 | 0.467 | 0.818 | 2071 | 0.823 | 0.997 | 921 | 0.056 | 0.293 | 2557 | 0.952 | 0.997 |
| BRD-K24593301-001-02-3 | AZD3463 | 0.713 | 192 | 0.518 | 0.747 | 1121 | 0.517 | 0.747 | 938 | 0.258 | 0.743 | 2129 | 0.008 | 0.112 | 3298 | 0.924 | 0.999 | 1971 | 0.283 | 0.743 | 4022.5 | 0.729 | 0.933 |
| BRD-K24603946-001-02-1 | 5-hydroxymethyl-tolterodine | 0.388 | 115 | 0.474 | 0.830 | 328 | 0.819 | 0.969 | 401 | 0.111 | 0.388 | 1546 | 0.099 | 0.388 | 774 | 0.015 | 0.314 | 303 | 0.109 | 0.388 | 1822 | 0.918 | 0.969 |
| BRD-K24715592-406-02-1 | ethacridine-lactate-monohydrate | 0.971 | | | | 330 | 0.491 | 0.921 | 221 | 0.493 | 0.921 | 1688 | 0.526 | 0.921 | 775 | 0.774 | 0.971 | 354 | 0.027 | 0.114 | 972 | 0.166 | 0.435 |
| BRD-K24844714-001-24-5 | 5-fluorouracil | 0.937 | 170 | 0.794 | 0.937 | 520 | 0.937 | 0.937 | 456.5 | 0.837 | 0.937 | 1394 | 0.099 | 0.721 | 879 | 0.265 | 0.937 | 596 | 0.137 | 0.721 | 1048 | 0.055 | 0.721 |
| BRD-K24869513-001-02-1 | CNX-2006 | 0.918 | | | | 275 | 0.987 | 0.987 | 142 | 0.490 | 1.000 | 775 | 0.270 | 0.946 | 40 | 0.041 | 0.374 | 130 | 0.123 | 0.287 | 919 | 0.647 | 0.918 |
| BRD-K25224017-001-25-0 | pirenperone | 0.530 | 80 | 0.201 | 0.530 | 598 | 0.202 | 0.530 | 1350 | 0.111 | 0.530 | 2782 | 0.040 | 0.314 | 4736 | 0.986 | 1.000 | 2668 | 0.449 | 0.820 | 3906 | 0.196 | 0.530 |
| BRD-K25244359-066-03-4 | apatinib | | | | | 64 | 0.103 | 0.730 | 203.5 | 0.892 | 1.000 | 638.5 | 0.964 | 1.000 | 220 | 0.104 | 0.730 | 308 | 0.187 | 0.982 | 675 | 0.631 | 0.889 |
| BRD-K25279140-001-01-5 | MK-8745 | 0.383 | 242 | 0.767 | 0.977 | 1161 | 0.739 | 0.977 | 1190 | 0.776 | 0.977 | 3007 | 0.589 | 0.977 | 1998 | 0.522 | 0.977 | 2018 | 0.546 | 0.977 | 3653 | 0.844 | 0.977 |
| BRD-K25340465-001-11-1 | OSI-930 | 0.985 | 24 | 0.082 | 0.596 | 462 | 0.756 | 0.987 | 619 | 0.799 | 0.987 | 1076 | 0.142 | 0.596 | 1178 | 0.436 | 0.818 | 1288 | 0.935 | 0.995 | 2195 | 0.932 | 0.995 |
| BRD-K25412176-001-01-9 | GSK2110183 | 0.728 | | | | 933.5 | 0.973 | 0.999 | 1085 | 0.960 | 0.999 | 2551 | 0.485 | 0.728 | 1246 | 0.000 | 0.003 | 1511.5 | 0.767 | 0.961 | 2565 | 0.649 | 0.909 |
| BRD-K25630527-001-03-8 | LY2784544 | 0.103 | | | | 51 | 0.048 | 0.501 | 248 | 0.478 | 0.913 | 978 | 0.992 | 0.992 | 500 | 0.132 | 0.346 | 550 | 0.883 | 0.992 | 871 | 0.656 | 0.992 |
| BRD-K25835157-001-01-4 | PRT062070 | 0.636 | | | | 89.5 | 0.817 | 0.941 | 351 | 0.518 | 0.837 | 400 | 0.934 | 0.941 | 320.5 | 0.655 | 0.941 | 74 | 0.161 | 0.621 | 361.5 | 0.147 | 1.000 |
| BRD-K26011976-001-01-0 | licochalcone-a | 0.364 | | | | 158.5 | 0.912 | 0.953 | 86 | 0.357 | 0.749 | 262 | 0.004 | 0.089 | 178 | 0.700 | 0.953 | | | | 424.5 | 0.867 | 0.953 |
| BRD-K26026438-001-01-0 | tucatinib | | | | | 155.5 | 0.436 | 0.763 | 189 | 0.956 | 0.997 | 383 | 0.185 | 1.000 | 182 | 0.291 | 0.555 | 212 | 0.082 | 0.192 | 502 | 0.285 | 0.555 |
| BRD-K26253904-001-01-5 | NMS-873 | 0.984 | | | | 1223 | 0.831 | 0.865 | 2590 | 0.994 | 1.000 | 2812 | 0.030 | 0.312 | 3434 | 0.535 | 0.984 | 2421 | 0.440 | 0.984 | 4085 | 0.750 | 0.984 |
| BRD-K26530649-001-01-6 | carboxypyridine-disulfide | 0.857 | 2 | 0.049 | 0.257 | | | | 164 | 0.302 | 0.793 | 186 | 0.829 | 0.967 | 261 | 0.031 | 0.644 | 255 | 0.797 | 0.967 | 154 | 0.079 | 0.832 |
| BRD-K26531771-001-02-2 | EPZ-5676 | 0.762 | | | | 205 | 0.512 | 0.896 | 335 | 0.625 | 0.938 | 739 | 0.099 | 0.347 | 999 | 0.977 | 0.999 | 522 | 0.847 | 0.999 | 1369 | 0.266 | 0.698 |
| BRD-K26603252-003-04-9 | PD-153035 | 0.502 | | | | | | | 398 | 0.293 | 0.590 | 680 | 0.000 | 0.007 | 1048 | 0.922 | 0.980 | 444 | 0.203 | 0.502 | 1226 | 0.069 | 0.208 |
| BRD-K26667523-001-02-5 | TAK-733 | 1.000 | 278 | 0.768 | 1.000 | 811 | 0.563 | 1.000 | 714 | 0.017 | 0.087 | 2961 | 0.032 | 0.135 | 3337 | 0.882 | 1.000 | 2917 | 0.419 | 1.000 | 5078 | 0.943 | 1.000 |
| BRD-K26712974-001-01-5 | VX-765 | 0.812 | 76 | 0.313 | 0.885 | 510 | 0.877 | 0.928 | 555 | 0.386 | 0.885 | 1683 | 0.878 | 0.928 | 1123 | 0.232 | 0.812 | 760 | 0.591 | 0.885 | 1698 | 0.613 | 0.885 |
| BRD-K26756394-001-02-2 | VE-821 | | | | | | | | 71 | 0.169 | 1.000 | 381 | 0.986 | 0.986 | 365 | 0.849 | 0.938 | 125 | 0.562 | 0.841 | 427 | 0.641 | 0.841 |
| BRD-K26818574-305-04-3 | BIX-01294 | 0.771 | 149 | 0.414 | 0.771 | 393 | 0.220 | 0.661 | 1371 | 0.441 | 0.771 | 2341 | 0.041 | 0.427 | 2132 | 0.357 | 0.771 | 2390 | 0.792 | 0.998 | 4476 | 0.973 | 0.998 |
| BRD-K26823213-001-02-9 | GW-842166 | 0.584 | | | | | | | 125 | 0.075 | 0.708 | 308.5 | 0.439 | 1.000 | 118 | 0.202 | 0.708 | 232.5 | 0.889 | 0.977 | 317 | 0.747 | 1.000 |
| BRD-K26838195-075-04-9 | AST-1306 | 0.266 | | | | 352 | 0.156 | 0.387 | 344 | 0.969 | 0.999 | 583 | 0.008 | 0.088 | 972 | 0.915 | 0.999 | 622 | 0.387 | 0.677 | 770 | 0.003 | 0.053 |
| BRD-K26997899-001-02-2 | D-64131 | 0.633 | | | | 629 | 0.859 | 0.935 | 831 | 0.364 | 0.694 | 1550 | 0.026 | 0.551 | 2192 | 0.652 | 0.935 | 1940 | 0.850 | 0.935 | 2353 | 0.235 | 0.633 |
| BRD-K27182532-001-02-3 | metatinib | 0.986 | 270 | 0.892 | 0.986 | 800 | 0.835 | 0.986 | 850 | 0.167 | 0.584 | 2152 | 0.080 | 0.555 | 2402 | 0.663 | 0.986 | 1370 | 0.072 | 0.555 | 2829 | 0.652 | 0.986 |
| BRD-K27184429-300-01-3 | levocetirizine | 0.864 | | | | 133 | 0.358 | 1.000 | 338 | 0.623 | 0.864 | 380 | 0.781 | 0.864 | 524.5 | 0.059 | 1.000 | 529.5 | 0.992 | 0.992 | 689 | 0.543 | 0.864 |
| BRD-K27217864-001-22-3 | butamben | 0.555 | 91 | 0.566 | 0.961 | 133 | 0.849 | 0.961 | 394 | 0.961 | 0.961 | 763 | 0.260 | 0.781 | 159 | 0.010 | 0.107 | 177 | 0.058 | 0.306 | 1040 | 0.741 | 0.961 |
| BRD-K27316855-001-13-3 | calcitriol | 0.957 | 51 | 0.205 | 0.957 | 210 | 0.571 | 0.957 | 180 | 0.424 | 0.957 | 930 | 0.019 | 0.201 | 940 | 0.501 | 0.957 | 1119 | 0.945 | 0.957 | 745 | 0.464 | 0.957 |
| BRD-K28061410-001-06-6 | beta-lapachone | 0.871 | 150 | 0.331 | 0.871 | 1928 | 0.845 | 0.949 | 2551 | 0.782 | 0.949 | 4150 | 0.151 | 0.724 | 4623 | 0.538 | 0.871 | 3701 | 0.478 | 0.871 | 3761 | 0.047 | 0.489 |
| BRD-K28192623-066-08-1 | spiradoline | 0.553 | 173 | 0.508 | 0.935 | 793 | 0.717 | 0.935 | 831 | 0.091 | 0.493 | 2304 | 0.117 | 0.493 | 2729 | 0.757 | 0.935 | 2073 | 0.595 | 0.935 | 3190 | 0.734 | 0.935 |
| BRD-K28352084-001-02-3 | CH5132799 | 0.712 | 147 | 0.465 | 0.752 | 93 | 0.005 | 0.099 | 1139 | 0.582 | 0.873 | 2051 | 0.043 | 0.304 | 1910 | 0.149 | 0.521 | 1330 | 0.116 | 0.488 | 2621 | 0.418 | 0.752 |
| BRD-K28360340-001-08-3 | TW-37 | 0.201 | 10 | 0.026 | 0.201 | 1062 | 0.518 | 0.876 | 2176 | 0.798 | 0.977 | 4364 | 0.916 | 0.977 | 3692 | 0.377 | 0.848 | 2846 | 0.650 | 0.876 | 4465 | 0.658 | 0.876 |
| BRD-K28383148-001-01-4 | AP1903 | 0.153 | | | | 42 | 0.226 | 0.948 | 89 | 0.632 | 0.975 | 558 | 0.001 | 0.015 | 539 | 0.843 | 0.975 | 529 | 0.958 | 0.975 | 1194 | 0.937 | 0.975 |
| BRD-K28537285-001-01-8 | chidamide | 0.739 | 0 | 0.043 | 0.465 | 276 | 0.295 | 0.739 | 902 | 0.549 | 0.739 | 1268 | 0.928 | 0.974 | 1067 | 0.077 | 0.536 | 607 | 0.167 | 0.739 | 1335 | 0.570 | 0.739 |
| BRD-K28578425-001-07-0 | cilostamide | 0.843 | | | | 263 | 0.983 | 0.983 | 229 | 0.936 | 0.983 | 601 | 0.495 | 0.866 | 468 | 0.472 | 0.866 | 308 | 0.344 | 0.866 | 676 | 0.354 | 1.000 |
| BRD-K28822270-001-03-7 | resminostat | 0.311 | 147 | 0.572 | 0.864 | 439 | 0.283 | 0.595 | 633 | 0.208 | 0.595 | 1418 | 0.508 | 0.864 | 1504 | 0.089 | 0.311 | 847 | 0.082 | 0.311 | 1488 | 0.617 | 0.864 |
| BRD-K28824103-001-02-8 | genipin | 0.752 | | | | 1304 | 0.898 | 0.997 | 1200 | 0.558 | 1.000 | 2635 | 0.842 | 0.997 | 1778 | 0.018 | 0.371 | 1025 | 0.057 | 0.119 | 3383 | 0.950 | 0.997 |
| BRD-K28907958-001-04-3 | CD-437 | 0.861 | 34 | 0.082 | 0.430 | 1556 | 0.832 | 0.930 | 1611 | 0.666 | 0.930 | 3393 | 0.179 | 0.538 | 4377 | 0.913 | 0.930 | 2414 | 0.429 | 0.819 | 4380 | 0.322 | 0.802 |
| BRD-K28965160-001-01-1 | BVD-523 | 0.458 | 174 | 0.738 | 0.993 | 434 | 0.667 | 0.993 | 545 | 0.046 | 0.320 | 1885 | 0.199 | 0.521 | 1171 | 0.608 | 0.993 | 1114.5 | 0.725 | 0.993 | 1802.5 | 0.657 | 0.993 |
| BRD-K29254801-001-14-7 | thiram | 0.778 | 168 | 0.491 | 0.906 | 476 | 0.158 | 0.778 | 1459 | 0.968 | 0.968 | 3379 | 0.933 | 0.968 | 2190 | 0.866 | 0.968 | 2159 | 0.207 | 0.778 | 2482 | 0.358 | 0.778 |
| BRD-K29415052-050-04-8 | BGT226 | 0.997 | 270 | 0.613 | 0.903 | 1852 | 0.834 | 0.997 | 1704 | 0.466 | 0.752 | 4248 | 0.382 | 0.670 | 3682 | 0.269 | 0.670 | 3448 | 0.375 | 0.670 | 4640 | 0.383 | 0.670 |
| BRD-K29542628-001-03-9 | 1-naphthyl-PP1 | 0.402 | | | | 257 | 0.094 | 0.660 | 724 | 0.757 | 0.874 | 704 | 0.021 | 0.284 | 648 | 0.027 | 0.284 | 659 | 0.602 | 0.874 | 1001.5 | 0.679 | 0.874 |
| BRD-K29656036-001-02-5 | MK-8245 | 0.411 | | | | 135 | 0.143 | 1.000 | 823 | 0.275 | 1.000 | 1769 | 0.432 | 0.756 | 1364 | 0.821 | 0.957 | 463 | 0.073 | 0.218 | 1441 | 0.592 | 0.899 |
| BRD-K29905972-001-06-3 | axitinib | 0.990 | | | | 60 | 0.420 | 0.990 | 288 | 0.897 | 1.000 | 370.5 | 0.670 | 0.990 | 71 | 0.006 | 0.064 | 387 | 0.464 | 0.990 | 579 | 0.707 | 0.990 |
| BRD-K29950728-048-17-4 | clomifene | 0.957 | | | | 164 | 0.296 | 0.836 | 91 | 0.017 | 0.184 | 1551 | 0.614 | 0.859 | 729 | 0.767 | 0.947 | 720 | 0.499 | 0.838 | 1415 | 0.249 | 0.836 |
| BRD-K29968218-001-03-2 | KX2-391 | 0.219 | 189 | 0.467 | 0.817 | 1367 | 0.749 | 0.888 | 1681 | 0.415 | 0.817 | 3396 | 0.332 | 0.817 | 2754 | 0.041 | 0.284 | 2306 | 0.310 | 0.817 | 4682.5 | 0.846 | 0.935 |
| BRD-K30431096-004-01-3 | domiphen | 0.141 | | | | 289 | 0.958 | 0.958 | 415.5 | 0.109 | 0.504 | 416 | 0.110 | 0.504 | 298 | 0.143 | 0.504 | 391.5 | 0.950 | 0.958 | 789.5 | 0.664 | 0.958 |
| BRD-K30480208-001-18-5 | torasemide | 0.546 | | | | 87 | 0.854 | 1.000 | 28 | 0.195 | 0.660 | 384 | 0.602 | 1.000 | 318 | 0.283 | 0.660 | 228 | 0.896 | 1.000 | 461 | 0.977 | 0.977 |
| BRD-K30577245-001-05-0 | docetaxel | 0.964 | 139 | 0.539 | 0.964 | 305 | 0.653 | 0.964 | 963.5 | 0.620 | 0.964 | 1222.5 | 0.061 | 0.322 | 1812 | 0.742 | 0.964 | 1809 | 0.952 | 0.964 | 1134.5 | 0.013 | 0.134 |
| BRD-K30577245-341-01-9 | docetaxel | 1.000 | 16 | 0.037 | 0.261 | 1645 | 0.817 | 1.000 | 1921 | 0.227 | 0.629 | 2940 | 0.003 | 0.049 | 4136 | 0.349 | 0.792 | 4527 | 0.974 | 1.000 | 3548 | 0.005 | 0.049 |
| BRD-K30867024-003-03-3 | SB-216641 | 0.976 | | | | 428 | 0.375 | 0.717 | 173 | 0.066 | 0.798 | 1284 | 0.115 | 0.802 | 1366 | 0.836 | 0.976 | 487 | 0.034 | 0.118 | 1320 | 0.076 | 0.798 |
| BRD-K30933884-001-01-0 | UNBS-5162 | 0.901 | 50 | 0.134 | 0.485 | 819 | 0.948 | 0.994 | 970 | 0.515 | 0.901 | 2135 | 0.911 | 0.994 | 1349 | 0.017 | 0.176 | 1120 | 0.172 | 0.485 | 2910 | 0.993 | 0.994 |
| BRD-K31086665-005-04-6 | NH125 | 0.945 | | | | 1427 | 0.902 | 0.945 | 1129 | 0.059 | 0.528 | 2279 | 0.010 | 0.212 | 2839 | 0.669 | 1.000 | 2143 | 0.686 | 0.945 | 3269 | 0.734 | 0.945 |
| BRD-K31309378-001-01-5 | lucitanib | 0.845 | 20 | 0.084 | 0.441 | 443 | 0.916 | 0.999 | 543 | 0.630 | 0.845 | 1280 | 0.257 | 0.770 | 965 | 0.684 | 0.845 | 417 | 0.105 | 0.441 | 1049 | 0.658 | 0.845 |
| BRD-K31476763-001-01-5 | AZD9668 | 0.132 | | | | 241 | 0.965 | 0.984 | 122 | 0.111 | 0.552 | 155 | 0.019 | 0.395 | 328 | 0.753 | 0.974 | 388 | 0.708 | 1.000 | 845 | 0.984 | 0.984 |
| BRD-K31491153-003-05-9 | 1-phenylbiguanide | 0.437 | | | | 131 | 0.337 | 0.847 | 200 | 0.737 | 0.860 | 902 | 0.356 | 0.847 | 440 | 0.689 | 0.860 | 628 | 0.990 | 0.990 | 582 | 0.409 | 0.859 |
| BRD-K31812033-003-11-0 | levobunolol | 0.902 | | | | | | | 564 | 0.859 | 0.902 | 779 | 0.178 | 1.000 | 306 | 0.670 | 0.873 | 269 | 0.207 | 0.582 | 898 | 0.923 | 0.923 |

Suppl. Table 9

| broad_id | name | colorectal_MWU_STAT | colorectal_PVAL | colorectal_PVAL_ADJ | skin_MWU_STAT | skin_PVAL | skin_PVAL_ADJ | urinary_tract_MWU_STAT | urinary_tract_PVAL | urinary_tract_PVAL_ADJ | lung_MWU_STAT | lung_PVAL | lung_PVAL_ADJ | ovary_MWU_STAT | ovary_PVAL | ovary_PVAL_ADJ | breast_MWU_STAT | breast_PVAL | breast_PVAL_ADJ | pancreas_MWU_STAT | pancreas_PVAL | pancreas_PVAL_ADJ | nervous_system_MWU_STAT | central_nervous_system_PVAL | nervous_system_PVAL_ADJ | soft_tissue_MWU_STAT | soft_tissue_PVAL | soft_tissue_PVAL_ADJ | kidney_MWU_STAT | kidney_PVAL | kidney_PVAL_ADJ | gastric_MWU_STAT | gastric_PVAL | gastric_PVAL_ADJ | bone_MWU_STAT | bone_PVAL | bone_PVAL_ADJ | prostate_MWU_STAT | prostate_PVAL | prostate_PVAL_ADJ | al_nervous_system_MWU_STAT | peripheral_nervous_system_PVAL |
|---|---|---|---|---|---|---|---|---|---|---|---|---|---|---|---|---|---|---|---|---|---|---|---|---|---|---|---|---|---|---|---|---|---|---|---|---|---|---|---|---|---|---|
| BRD-K31866293-001-01-6 | TAK-632 | 3088 | 0.018 | 0.185 | 3490 | 0.000 | 0.000 | 4127 | 0.573 | 0.993 | 14993 | 0.968 | 0.997 | 4620 | 0.662 | 0.993 | 4432 | 0.975 | 0.997 | 5937 | 0.922 | 0.997 | 6525 | 0.928 | 0.997 | 2097 | 0.428 | 0.899 | 3229 | 0.997 | 0.997 | 2485 | 0.086 | 0.362 | 1420 | 0.308 | 0.775 | 510 | 0.332 | 0.775 | 595 | 0.061 |
| BRD-K31928526-001-02-1 | barasertib | 1028 | 0.816 | 1.000 | 1370 | 0.975 | 1.000 | 640 | 0.027 | 0.283 | 2691.5 | 0.509 | 1.000 | 551.5 | 0.045 | 0.315 | 597.5 | 0.799 | 1.000 | 1148.5 | 0.873 | 1.000 | 1109 | 0.356 | 1.000 | 254 | 0.068 | 0.357 | 560 | 0.989 | 1.000 | 189 | 0.018 | 0.283 | 599.5 | 0.341 | 1.000 | | | | 153 | 0.120 |
| BRD-K32372024-001-01-0 | D-7193 | 759.5 | 0.818 | 0.927 | 1709 | 0.414 | 0.870 | 1151.5 | 0.387 | 0.870 | 3517.5 | 0.331 | 0.870 | 1221 | 0.347 | 0.870 | 607 | 0.234 | 0.870 | 1740 | 0.332 | 0.870 | 2021 | 0.744 | 0.927 | 328.5 | 0.229 | 0.870 | 769 | 0.835 | 0.927 | 463 | 0.333 | 0.870 | 468.5 | 0.655 | 0.927 | 489.5 | 0.948 | 0.962 | 505.5 | 0.962 |
| BRD-K32501161-300-06-2 | vanoxerine | 1762 | 0.612 | 0.857 | 3399 | 0.594 | 1.000 | 1804 | 0.339 | 1.000 | 7272 | 0.869 | 0.922 | 1846 | 0.153 | 0.805 | 1544 | 0.477 | 1.000 | 1967 | 0.554 | 1.000 | 2559 | 0.546 | 1.000 | 578 | 0.604 | 1.000 | 1013 | 0.659 | 0.865 | 1241 | 0.224 | 0.811 | 190 | 0.011 | 0.239 | 641 | 0.960 | 0.960 | 152 | 0.031 |
| BRD-K32572029-001-04-4 | orotic-acid | 303 | 0.751 | 0.988 | 513 | 0.269 | 0.807 | 509 | 0.764 | 0.988 | 968 | 0.667 | 1.000 | 272 | 0.026 | 0.450 | 326 | 0.846 | 0.988 | 219 | 0.287 | 0.807 | 410 | 0.347 | 0.807 | 94 | 0.396 | 0.807 | 115 | 0.064 | 0.450 | 124 | 0.267 | 0.807 | 41 | 0.349 | 0.807 | 47 | 0.423 | 0.807 | | |
| BRD-K32730239-001-01-0 | SU3327 | 590.5 | 0.690 | 0.895 | 988 | 0.035 | 0.733 | 795 | 0.640 | 0.895 | 1703.5 | 0.657 | 0.895 | 1007 | 0.217 | 0.895 | 555 | 0.578 | 0.895 | 968 | 0.944 | 0.984 | 758 | 0.538 | 0.895 | 147 | 0.137 | 0.747 | 381 | 0.486 | 0.895 | 217 | 0.417 | 0.895 | 192 | 0.098 | 0.747 | 173 | 0.601 | 0.895 | 149 | 0.142 |
| BRD-K32744045-001-31-2 | disulfiram | 6320 | 0.971 | 0.971 | 7367 | 0.342 | 0.685 | 4117 | 0.091 | 0.382 | 18523 | 0.897 | 0.971 | 5887 | 0.359 | 0.685 | 3705 | 0.141 | 0.491 | 6883.5 | 0.865 | 0.971 | 5949 | 0.091 | 0.382 | 2315 | 0.399 | 0.697 | 4420 | 0.914 | 0.971 | 2950 | 0.079 | 0.382 | 1811 | 0.165 | 0.491 | 1328 | 0.952 | 0.971 | 547 | 0.025 |
| BRD-K32821942-001-21-3 | azathioprine | 666 | 0.979 | 0.979 | 1368 | 0.853 | 0.979 | 440 | 0.736 | 0.965 | 1698 | 0.121 | 0.509 | 995 | 0.333 | 0.777 | 712 | 0.952 | 0.979 | 627 | 0.603 | 0.875 | 832 | 0.398 | 0.792 | 166 | 0.195 | 0.584 | 383 | 0.515 | 0.831 | 443 | 0.235 | 1.000 | 164 | 0.187 | 0.584 | 24 | 0.110 | 0.509 | 18 | 0.013 |
| BRD-K33127281-001-05-6 | adapalene | 274 | 0.919 | 0.940 | 364 | 0.854 | 0.940 | 259 | 0.093 | 0.945 | 873 | 0.020 | 0.415 | 274 | 0.919 | 0.940 | 348 | 0.786 | 0.940 | 551 | 0.940 | 0.940 | 366 | 0.620 | 0.940 | 149 | 0.455 | 0.940 | 77 | 0.231 | 0.606 | 158 | 0.536 | 1.000 | 78 | 0.042 | 0.125 | | | | 70 | 0.759 |
| BRD-K33277808-331-01-3 | ouabain | 5746 | 1.000 | 1.000 | 6700.5 | 1.000 | 1.000 | 3442 | 0.511 | 0.832 | 10977.5 | 0.011 | 0.240 | 4265 | 0.373 | 0.783 | 3951 | 0.936 | 1.000 | 4440 | 0.627 | 0.874 | 3817 | 0.073 | 0.514 | 1698 | 0.329 | 0.768 | 2254 | 0.063 | 0.514 | 2368 | 0.666 | 0.874 | 1517 | 0.538 | 0.832 | 904 | 0.757 | 0.936 | 727 | 0.804 |
| BRD-K33379087-001-07-5 | tivantinib | 751 | 0.499 | 0.952 | 2065.5 | 0.979 | 0.980 | 690 | 0.015 | 0.104 | 2714.5 | 0.180 | 0.631 | 1123 | 0.073 | 0.381 | 783 | 0.796 | 0.980 | 1378 | 0.976 | 0.980 | 1339.5 | 0.627 | 0.980 | 1073.5 | 0.980 | 0.980 | 668 | 0.937 | 0.980 | 607.5 | 0.180 | 0.631 | 363 | 0.428 | 0.898 | 142 | 0.787 | 0.980 | 354 | 0.396 |
| BRD-K33610132-001-02-9 | rociletinib | | | | 788 | 0.999 | 0.999 | 625 | 0.269 | 0.629 | 1802 | 0.743 | 0.977 | 569 | 0.447 | 0.939 | 365 | 0.175 | 0.526 | 871 | 0.795 | 0.977 | 191 | 0.833 | 0.977 | 69 | 0.115 | 0.485 | 524 | 0.727 | 0.977 | 376 | 0.681 | 0.977 | 186 | 0.810 | 0.977 | 125 | 0.921 | 0.977 | 77 | 0.145 |
| BRD-K33622447-066-01-9 | abemaciclib | 1399 | 0.779 | 0.984 | 2311 | 0.812 | 0.984 | 1049 | 0.331 | 0.773 | 3634 | 0.507 | 0.984 | 982 | 0.025 | 0.107 | 1273 | 0.885 | 0.984 | 773 | 0.184 | 0.551 | 1455 | 0.984 | 0.984 | 507 | 0.010 | 0.068 | 1099 | 0.017 | 0.088 | 851 | 0.526 | 1.000 | 421 | 0.213 | 0.559 | 168 | 0.830 | 0.984 | 194 | 0.031 |
| BRD-K33720171-001-01-0 | difluprednate | 2770 | 0.973 | 0.973 | 4212 | 0.890 | 0.966 | 1803 | 0.266 | 0.797 | 7410 | 0.746 | 0.963 | 2204 | 0.029 | 0.483 | 1658 | 0.249 | 0.797 | 2337 | 0.422 | 0.963 | 2872 | 0.826 | 0.963 | 1014 | 0.736 | 0.963 | 1999 | 0.669 | 0.963 | 1638 | 0.064 | 0.483 | 762 | 0.552 | 0.963 | 711 | 0.756 | 0.963 | 807 | 0.642 |
| BRD-K33743361-003-02-6 | ML133 | 1706 | 0.358 | 0.823 | 2825 | 0.968 | 0.981 | 1148 | 0.204 | 1.000 | 4776 | 0.026 | 0.541 | 1898 | 0.470 | 0.823 | 1296 | 0.416 | 0.823 | 2644 | 0.981 | 0.981 | 2084 | 0.840 | 0.981 | 723 | 0.441 | 0.823 | 1485 | 0.867 | 0.981 | 1075 | 0.250 | 1.000 | 509 | 0.236 | 0.708 | 535 | 0.889 | 0.981 | 545 | 0.611 |
| BRD-K33813875-003-01-8 | valnemulin | 2783.5 | 0.369 | 0.896 | 6836 | 0.997 | 0.997 | 2566 | 0.640 | 0.896 | 10188.5 | 0.180 | 0.655 | 3627 | 0.128 | 0.655 | 3198.5 | 0.937 | 0.994 | 4183.5 | 0.487 | 0.896 | 4497 | 0.821 | 0.994 | 1612 | 0.331 | 0.896 | 2668 | 0.947 | 0.994 | 1787.5 | 0.021 | 0.217 | 1443 | 0.543 | 0.896 | 529 | 0.187 | 0.655 | 471 | 0.121 |
| BRD-K33882852-003-02-8 | ZK-93423 | 225 | 0.791 | 0.923 | 732 | 0.752 | 0.923 | 394 | 0.723 | 0.923 | 1432 | 0.588 | 0.923 | 404 | 0.078 | 0.329 | 272 | 0.712 | 0.923 | 556 | 0.862 | 0.923 | 588 | 0.923 | 0.923 | 134 | 0.239 | 0.718 | 217 | 0.405 | 0.851 | 262 | 0.162 | 1.000 | 160 | 0.801 | 0.923 | | | | 1 | 0.047 |
| BRD-K34022604-001-06-6 | clofarabine | 4747 | 0.650 | 0.975 | 4484 | 0.001 | 0.013 | 4087 | 0.588 | 0.950 | 14189 | 0.343 | 0.902 | 5307 | 0.411 | 0.950 | 4375 | 0.942 | 0.978 | 6036 | 0.945 | 0.978 | 5854 | 0.531 | 0.950 | 1765 | 0.118 | 0.495 | 2613 | 0.488 | 0.950 | 2406 | 0.045 | 0.236 | 1031 | 0.012 | 0.086 | 874 | 0.577 | 0.950 | 882 | 0.296 |
| BRD-K34332569-001-01-3 | imidapril | 1573 | 0.840 | 1.000 | 2296.5 | 0.219 | 0.918 | 1448 | 0.324 | 0.918 | 5635 | 0.575 | 0.918 | 2024 | 0.159 | 0.918 | 1068.5 | 0.439 | 0.918 | 1785 | 0.338 | 0.918 | 2185 | 0.839 | 1.000 | 357 | 0.004 | 0.086 | 2020 | 0.961 | 1.000 | 993 | 0.519 | 0.918 | 755 | 0.960 | 1.000 | 281 | 0.612 | 0.918 | 455 | 0.380 |
| BRD-K34851558-001-02-3 | parthenolide-(-) | 1342 | 0.447 | 0.682 | 1975 | 0.216 | 0.908 | 1219 | 0.420 | 0.980 | 4510 | 0.455 | 0.682 | 2289.5 | 0.918 | 0.980 | 1601 | 0.138 | 0.812 | 1804 | 0.803 | 0.980 | 2821 | 0.980 | 0.980 | 205 | 0.103 | 0.812 | 769 | 0.649 | 0.852 | 666 | 0.410 | 0.980 | 473 | 0.499 | 0.698 | 491.5 | 0.870 | 0.980 | 350 | 0.186 |
| BRD-K35329391-334-01-7 | GZD824 | 5296 | 0.980 | 0.990 | 5227 | 0.031 | 0.292 | 4262 | 0.545 | 0.990 | 12679 | 0.382 | 0.990 | 3836 | 0.056 | 0.292 | 4177 | 0.990 | 0.990 | 6145 | 0.879 | 0.990 | 4978 | 0.775 | 0.990 | 1674 | 0.101 | 0.425 | 3392 | 0.736 | 0.990 | 2589 | 0.230 | 0.690 | 1984 | 0.516 | 0.990 | 611 | 0.519 | 0.990 | 1388 | 0.932 |
| BRD-K35520305-001-17-7 | dacarbazine | 1031 | 0.694 | 0.911 | 1361 | 0.863 | 0.917 | 557 | 0.253 | 1.000 | 2373 | 0.371 | 1.000 | 751 | 0.057 | 0.569 | 325 | 0.813 | 0.917 | 858 | 0.651 | 1.000 | 975 | 0.739 | 0.913 | 344 | 0.551 | 1.000 | 272 | 0.098 | 0.569 | 709 | 0.675 | 0.911 | 202 | 0.691 | 0.911 | 140 | 0.873 | 0.917 | 32 | 0.145 |
| BRD-K35960502-001-20-0 | niclosamide | 1086 | 0.743 | 0.857 | 1916.5 | 0.608 | 0.857 | 1297 | 0.736 | 0.857 | 3832.5 | 0.271 | 0.857 | 1690.5 | 0.179 | 0.857 | 1633 | 0.464 | 0.857 | 1861.5 | 0.895 | 0.939 | 1204 | 0.389 | 0.857 | 232 | 0.054 | 0.857 | 866.5 | 0.306 | 0.857 | 796 | 0.595 | 0.857 | 409 | 0.763 | 0.857 | 227 | 0.529 | 0.857 | 451 | 0.542 |
| BRD-K36021395-300-02-4 | ABT-702 | 1013 | 0.499 | 0.901 | 986 | 0.000 | 0.010 | 1122 | 0.567 | 0.901 | 2843 | 0.956 | 0.981 | 1138 | 0.601 | 0.901 | 498 | 0.694 | 0.911 | 1216 | 0.753 | 0.931 | 764 | 0.475 | 0.901 | 465 | 0.587 | 0.901 | 569 | 0.981 | 0.981 | 401 | 0.369 | 1.000 | 514 | 0.940 | 0.981 | 260 | 0.866 | 0.981 | 34 | 0.144 |
| BRD-K36055864-001-19-2 | cycloheximide | 3484.5 | 0.839 | 0.979 | 6076 | 0.968 | 0.997 | 3694 | 0.343 | 0.665 | 12130.5 | 0.584 | 0.817 | 3591 | 0.037 | 0.227 | 3413 | 0.997 | 0.997 | 4633 | 0.443 | 0.665 | 5853.5 | 0.962 | 0.997 | 1617 | 0.265 | 0.665 | 2591 | 0.727 | 0.939 | 2000 | 0.043 | 0.227 | 1548 | 0.375 | 0.665 | 466 | 0.310 | 0.665 | 289 | 0.075 |
| BRD-K36198571-001-02-0 | WAY-170523 | 716 | 0.950 | 0.968 | 1581 | 0.286 | 1.000 | 1413 | 0.343 | 1.000 | 5136 | 0.459 | 1.000 | 1686 | 0.057 | 1.000 | 1824 | 0.635 | 0.889 | 1983 | 0.552 | 1.000 | 2151 | 0.766 | 0.893 | 959 | 0.746 | 0.893 | 875 | 0.084 | 0.195 | 840 | 0.287 | 1.000 | 244 | 0.502 | 0.889 | 380 | 0.552 | 0.889 | 451 | 0.761 |
| BRD-K36222532-001-01-8 | broxaldine | 3023.5 | 0.854 | 0.960 | 4513.5 | 0.582 | 0.861 | 2669 | 0.399 | 0.776 | 9832 | 0.341 | 0.776 | 3109 | 0.063 | 0.657 | 2569 | 0.615 | 0.861 | 4773 | 0.997 | 0.997 | 2545 | 0.029 | 0.606 | 1363 | 0.165 | 0.757 | 1812 | 0.854 | 0.960 | 2206 | 0.743 | 0.960 | 1136.5 | 0.232 | 0.757 | 546 | 0.252 | 0.757 | 891 | 0.584 |
| BRD-K36269323-300-02-6 | asymmetrical-dimethylarginine | 2249.5 | 0.389 | 0.742 | 3256 | 0.093 | 0.393 | 1925 | 0.189 | 0.501 | 6528 | 0.425 | 0.744 | 3068 | 0.715 | 0.938 | 1953 | 0.824 | 0.989 | 3064.5 | 0.902 | 0.989 | 3017 | 0.976 | 0.989 | 970 | 0.703 | 0.938 | 1509 | 0.989 | 0.989 | 2181.5 | 0.619 | 0.929 | 837 | 0.230 | 0.501 | 199 | 0.053 | 0.280 | 88 | 0.045 |
| BRD-K36363294-001-04-2 | I-BET151 | 232 | 0.274 | 1.000 | 594 | 0.679 | 0.838 | 293 | 0.672 | 0.838 | 1066 | 0.647 | 0.838 | 591 | 0.844 | 0.886 | 35 | 0.023 | 0.070 | 195 | 0.108 | 0.261 | 342 | 0.357 | 0.750 | 437 | 0.616 | 0.838 | 362 | 0.758 | 0.884 | 290 | 0.653 | 0.838 | 215 | 0.496 | 0.838 | 1 | 0.021 | 0.070 | 64 | 0.112 |
| BRD-K36386086-001-01-1 | dolutegravir | 1645 | 0.975 | 0.996 | 1349 | 0.536 | 1.000 | 959 | 0.296 | 1.000 | 3107 | 0.194 | 0.814 | 988 | 0.026 | 0.269 | 975 | 0.512 | 1.000 | 1241 | 0.123 | 0.814 | 1786 | 0.984 | 0.996 | 625 | 0.166 | 0.814 | 1221 | 0.996 | 0.996 | 229 | 0.020 | 0.269 | 601 | 0.958 | 0.996 | 416 | 0.871 | 0.996 | 331 | 0.616 |
| BRD-K36529613-001-02-6 | PU-H71 | 3860 | 0.788 | 0.985 | 8336 | 1.000 | 1.000 | 3136 | 0.084 | 0.381 | 13160 | 0.221 | 0.478 | 4943 | 0.520 | 0.911 | 3681 | 0.797 | 0.985 | 7092 | 0.982 | 1.000 | 6025 | 0.750 | 0.985 | 1591 | 0.068 | 0.381 | 3558 | 0.973 | 1.000 | 1648 | 0.002 | 0.045 | 1775 | 0.297 | 0.567 | 646 | 0.583 | 0.931 | 1117 | 0.911 |
| BRD-K36740062-001-06-6 | GSK1070916 | 2703 | 0.466 | 0.739 | 5691 | 0.939 | 0.986 | 1649 | 0.032 | 0.546 | 7874 | 0.198 | 0.720 | 2416 | 0.319 | 0.720 | 2372 | 0.866 | 0.986 | 3316 | 0.647 | 0.906 | 3300 | 0.374 | 0.720 | 768 | 0.087 | 0.609 | 2782 | 1.000 | 1.000 | 1345 | 0.052 | 0.546 | 1024 | 0.377 | 0.720 | 475 | 0.493 | 0.739 | 783 | 0.780 |
| BRD-K36889451-001-01-1 | tetramethylthiuram-monosulfide | 1621 | 0.997 | 0.997 | 2214.5 | 0.365 | 0.856 | 1492 | 0.506 | 0.856 | 4780 | 0.361 | 0.856 | 1631 | 0.048 | 0.493 | 776.5 | 0.094 | 0.493 | 2333.5 | 0.648 | 0.856 | 2272.5 | 0.438 | 0.856 | 788 | 0.683 | 0.856 | 1056 | 0.732 | 0.856 | 750 | 0.074 | 0.493 | 466 | 0.208 | 0.728 | 301 | 0.734 | 0.856 | 57 | 0.184 |
| BRD-K37270826-001-27-6 | mifepristone | 625 | 0.080 | 0.241 | 881 | 0.387 | 0.808 | 875 | 0.757 | 0.937 | 2760 | 0.929 | 0.937 | 954 | 0.071 | 0.241 | 1165 | 0.817 | 0.937 | 1197 | 0.418 | 0.808 | 1167 | 0.820 | 0.937 | 14 | 0.074 | 0.241 | 363 | 0.103 | 0.271 | 477 | 0.072 | 0.755 | 509 | 0.937 | 0.937 | 327 | 0.756 | 0.937 | | |
| BRD-K37312348-001-15-0 | kenpaullone | 266.5 | 0.325 | 0.717 | 589.5 | 0.793 | 0.891 | 426.5 | 0.840 | 0.891 | 1361 | 0.295 | 1.000 | 261.5 | 0.105 | 0.442 | 342.5 | 0.075 | 0.394 | 701 | 0.794 | 0.891 | 274.5 | 0.369 | 0.717 | 244.5 | 0.533 | 0.861 | 89.5 | 0.229 | 0.601 | 27 | 0.133 | 1.000 | 166 | 0.849 | 0.891 | | | | 169 | 0.410 |
| BRD-K37379014-001-02-0 | filanesib | 865 | 0.366 | 1.000 | 1380 | 0.558 | 1.000 | 709 | 0.107 | 0.951 | 3166.5 | 0.218 | 0.951 | 1085 | 0.186 | 0.951 | 609.5 | 0.861 | 0.952 | 916.5 | 0.861 | 0.952 | 1232.5 | 0.601 | 1.000 | 484 | 0.272 | 0.951 | 796.5 | 0.955 | 0.980 | 1005 | 0.846 | 0.952 | 578 | 0.139 | 0.243 | 170 | 0.388 | 0.627 | 274 | 0.445 |
| BRD-K37561857-001-15-5 | zardaverine | 2967 | 0.944 | 0.967 | 3839 | 0.135 | 0.628 | 2749 | 0.553 | 0.967 | 9610 | 0.150 | 0.628 | 4283 | 0.645 | 0.967 | 2290 | 0.264 | 0.734 | 4641 | 0.865 | 0.967 | 4103 | 0.280 | 0.734 | 1384 | 0.738 | 0.967 | 2019 | 0.470 | 0.967 | 2055 | 0.198 | 0.693 | 802 | 0.064 | 0.628 | 694 | 0.824 | 0.967 | 357 | 0.047 |
| BRD-K37618799-001-03-0 | MRS-1220 | 811.5 | 0.943 | 0.958 | 397 | 0.545 | 0.958 | 485.5 | 0.236 | 1.000 | 1430.5 | 0.531 | 0.958 | 833 | 0.771 | 0.958 | 397 | 0.004 | 0.014 | 386.5 | 0.766 | 0.958 | 547.5 | 0.958 | 0.958 | 118.5 | 0.910 | 0.958 | 380 | 0.742 | 0.958 | 464.5 | 0.343 | 0.801 | 118.5 | 0.910 | 0.958 | 29 | 0.166 | 0.437 | | |
| BRD-K37687095-001-06-9 | AZD8330 | 3060 | 0.001 | 0.010 | 2782 | 0.000 | 0.000 | 3509 | 0.073 | 0.384 | 17932 | 1.000 | 1.000 | 6555 | 0.972 | 1.000 | 4253 | 0.967 | 1.000 | 4328 | 0.002 | 0.006 | 6559 | 1.000 | 1.000 | 2135 | 0.627 | 1.000 | 3307 | 0.953 | 1.000 | 2598 | 0.109 | 0.285 | 1760 | 0.969 | 1.000 | 967 | 0.955 | 1.000 | 172 | 0.089 |
| BRD-K37720887-001-07-8 | SB-525334 | 548 | 0.974 | 1.000 | 364 | 0.051 | 0.667 | 494 | 0.753 | 1.000 | 1215 | 0.261 | 1.000 | 486 | 0.195 | 1.000 | 375 | 0.064 | 0.667 | 331 | 0.280 | 1.000 | 517 | 0.825 | 1.000 | 215 | 0.789 | 1.000 | 293 | 0.586 | 1.000 | 75 | 0.705 | 1.000 | 105 | 0.428 | 1.000 | | | | | |
| BRD-K37723606-001-01-5 | dioscin | 1864 | 0.776 | 0.932 | 2745 | 0.307 | 1.000 | 1737 | 0.292 | 1.000 | 6829 | 0.796 | 0.932 | 2199 | 0.124 | 1.000 | 1723 | 0.423 | 0.802 | 2062 | 0.843 | 0.932 | 2281 | 0.111 | 0.293 | 603 | 0.682 | 0.932 | 1124 | 0.965 | 0.974 | 771 | 0.109 | 1.000 | | | | 260 | 0.974 | 0.974 | 347 | 0.758 |
| BRD-K37764012-001-04-1 | PF-03758309 | 3001 | 0.863 | 1.000 | 5090 | 0.816 | 1.000 | 2282 | 0.645 | 0.903 | 7775 | 0.082 | 0.298 | 1557 | 0.085 | 0.298 | 2698 | 0.755 | 0.991 | 2761 | 0.261 | 0.608 | 4278 | 0.975 | 1.000 | 638 | 0.234 | 0.608 | 2904 | 1.000 | 1.000 | 1022 | 0.004 | 0.085 | 627 | 0.080 | 0.298 | 504 | 0.561 | 0.903 | 314 | 0.483 |
| BRD-K37798499-001-27-2 | etoposide | 2930 | 0.803 | 0.992 | 4065 | 0.330 | 0.770 | 2532 | 0.574 | 0.976 | 8635 | 0.193 | 0.578 | 2404 | 0.107 | 0.448 | 3512 | 0.992 | 0.992 | 3331 | 0.933 | 0.992 | 3983 | 0.990 | 0.992 | 1131 | 0.027 | 0.187 | 1738 | 0.877 | 0.992 | 1128 | 0.010 | 0.187 | 888 | 0.076 | 0.397 | 493 | 0.903 | 0.992 | 482 | 0.502 |
| BRD-K37865504-001-04-1 | LY2183240 | 1059 | 0.161 | 0.582 | 2862 | 0.998 | 0.998 | 1139 | 0.260 | 0.582 | 4864 | 0.262 | 0.582 | 2193 | 0.305 | 0.582 | 1367 | 0.449 | 0.674 | 1878 | 0.718 | 0.943 | 1906 | 0.870 | 0.998 | 831 | 0.059 | 0.412 | 1586 | 0.970 | 0.998 | 514 | 0.293 | 0.582 | 117 | 0.022 | 0.267 | 292 | 0.282 | 0.582 | 207 | 0.025 |
| BRD-K37890730-001-15-1 | camptothecin | 1700 | 0.451 | 0.677 | 4641 | 0.967 | 0.991 | 1994 | 0.509 | 0.710 | 5410 | 0.314 | 0.582 | 2223 | 0.226 | 0.582 | 2711 | 0.972 | 0.991 | 2822 | 0.360 | 0.582 | 2164 | 0.979 | 0.991 | 498 | 0.146 | 0.548 | 2410 | 0.991 | 0.991 | 1506 | 0.541 | 0.710 | 105 | 0.337 | 0.582 | 256 | 0.127 | 0.548 | 37 | 0.018 |
| BRD-K38183498-001-02-0 | clonazepam | 4073 | 0.742 | 0.974 | 5561 | 0.037 | 0.386 | 3480 | 0.303 | 0.645 | 12977 | 0.230 | 0.645 | 6071 | 0.868 | 1.000 | 4260 | 0.737 | 0.974 | 7110 | 0.985 | 1.000 | 6817 | 0.920 | 1.000 | 2605 | 0.688 | 0.974 | 3328 | 1.000 | 1.000 | 2434 | 0.113 | 0.591 | 1690 | 0.192 | 0.645 | 779 | 0.438 | 0.836 | 882 | 0.307 |
| BRD-K38305202-001-17-4 | domperidone | 188.5 | 0.122 | 0.853 | 272 | 0.000 | 0.000 | 363.5 | 0.482 | 1.000 | 881.5 | 0.618 | 0.965 | 512 | 0.875 | 0.965 | 85.5 | 0.840 | 0.965 | 623 | 0.758 | 0.965 | 450.5 | 0.662 | 0.965 | 227.5 | 0.579 | 0.965 | 139.5 | 0.756 | 0.965 | 220 | 0.530 | 0.965 | 254.5 | 0.740 | 0.965 | 170 | 0.919 | 0.965 | 33 | 0.237 |
| BRD-K38332599-001-01-3 | uprosertib | 1721 | 0.698 | 0.999 | 2806.5 | 0.634 | 0.999 | 1066 | 0.023 | 0.137 | 5502 | 0.891 | 0.999 | 1844 | 0.026 | 0.137 | 1379 | 0.214 | 0.563 | 3068.5 | 0.999 | 0.999 | 2402 | 0.995 | 0.999 | 977 | 0.264 | 0.615 | 1332.5 | 0.999 | 0.999 | 869.5 | 0.131 | 0.429 | 1081 | 0.857 | 0.999 | 244 | 0.143 | 0.429 | 409 | 0.588 |
| BRD-K38350380-001-01-8 | FK-3311 | 1388 | 0.959 | 0.959 | 3322 | 0.952 | 0.959 | 1477 | 0.690 | 0.888 | 4594 | 0.510 | 0.888 | 1583 | 0.040 | 0.209 | 1040 | 0.728 | 0.888 | 1707 | 0.387 | 0.879 | 1830 | 0.419 | 0.879 | 270 | 0.073 | 0.306 | 937 | 0.166 | 0.581 | 54 | 0.001 | 0.026 | 712 | 0.532 | 0.888 | 174 | 0.260 | 0.781 | 266 | 0.619 |
| BRD-K38527262-300-01-0 | atiprimod | 807 | 0.000 | 0.006 | 2306 | 0.003 | 0.032 | 1106 | 0.584 | 0.985 | 6213 | 0.999 | 0.999 | 1491 | 0.474 | 0.985 | 979 | 0.346 | 0.985 | 1363 | 0.805 | 0.985 | 2458 | 0.754 | 0.985 | 843 | 0.770 | 0.985 | 1065 | 0.038 | 0.263 | 1159 | 0.680 | 0.985 | 921 | 0.980 | 0.999 | 392 | 0.933 | 0.999 | 332 | 0.815 |
| BRD-K38548312-001-01-0 | VER-49009 | 2278 | 0.265 | 0.506 | 5597 | 0.962 | 0.977 | 3191 | 0.679 | 0.950 | 10773 | 0.612 | 0.918 | 2996 | 0.149 | 0.452 | 3543 | 0.904 | 0.977 | 4150 | 0.936 | 0.977 | 4839 | 0.891 | 0.977 | 1122 | 0.382 | 0.669 | 2140 | 0.977 | 0.977 | 1857 | 0.172 | 0.452 | 1192 | 0.124 | 0.452 | 357 | 0.182 | 0.452 | 634 | 0.158 |
| BRD-K38852836-001-02-1 | ganetespib | 4170 | 0.720 | 0.907 | 5780 | 0.856 | 0.907 | 3463 | 0.427 | 0.748 | 11289.5 | 0.365 | 0.748 | 4404 | 0.522 | 0.835 | 2434.5 | 0.409 | 0.748 | 6103.5 | 0.995 | 0.995 | 4241 | 0.290 | 0.748 | 1908 | 0.233 | 0.748 | 2763 | 0.857 | 0.907 | 2707 | 0.822 | 0.907 | 1499 | 0.323 | 0.748 | 603 | 0.596 | 0.835 | 181 | 0.022 |
| BRD-K38860038-001-01-2 | JK-184 | 4360 | 0.199 | 0.487 | 8438 | 0.997 | 0.997 | 3863 | 0.318 | 0.558 | 12870 | 0.138 | 0.487 | 3922 | 0.030 | 0.315 | 4652 | 0.987 | 0.997 | 6952 | 0.972 | 0.997 | 6964 | 0.910 | 0.997 | 1693 | 0.194 | 0.487 | 2852 | 0.724 | 0.997 | 1889 | 0.102 | 0.487 | 1553 | 0.108 | 0.487 | 931 | 0.678 | 0.997 | 799 | 0.209 |
| BRD-K38868394-001-01-3 | MK-0773 | 276 | 0.655 | 0.963 | 687 | 0.963 | 0.963 | 306 | 0.802 | 0.963 | 573 | 0.021 | 0.283 | 537 | 0.585 | 1.000 | 195 | 0.793 | 0.963 | 452 | 0.920 | 0.963 | 435 | 0.717 | 0.963 | 86 | 0.349 | 0.667 | 164 | 0.925 | 0.963 | 349 | 0.766 | 0.963 | 0 | 0.045 | 0.117 | 165 | 0.929 | 0.963 | 2 | 0.051 |
| BRD-K38904395-001-02-3 | SAR131675 | 806 | 0.673 | 0.876 | 647 | 0.220 | 1.000 | 340 | 0.436 | 1.000 | 1536 | 0.069 | 1.000 | 847 | 0.609 | 1.000 | 653 | 0.608 | 1.000 | 763 | 0.885 | 0.900 | 635 | 0.346 | 1.000 | 292 | 0.863 | 0.900 | 434 | 0.299 | 1.000 | 280 | 0.473 | 1.000 | 115 | 0.852 | 0.900 | 34 | 0.172 | 0.278 | 141 | 0.476 |
| BRD-K39171998-001-01-1 | STF-118804 | 3204 | 0.313 | 0.657 | 6095 | 0.422 | 0.683 | 4238 | 0.953 | 0.980 | 13996.5 | 0.942 | 0.980 | 4319 | 0.171 | 0.448 | 3950 | 0.750 | 0.926 | 6796 | 0.980 | 0.980 | 5720 | 0.299 | 0.657 | 1471 | 0.039 | 0.136 | 3385 | 0.946 | 0.980 | 2416 | 0.405 | 0.683 | 1371 | 0.021 | 0.111 | 622 | 0.554 | 0.755 | 534 | 0.039 |
| BRD-K39252998-003-01-5 | atipamezole | 975.5 | 0.306 | 0.848 | 1094 | 0.564 | 0.895 | 502.5 | 0.244 | 0.848 | 2199.5 | 0.079 | 0.359 | 1411 | 0.601 | 0.895 | 951 | 0.771 | 0.895 | 1244 | 0.713 | 0.895 | 917.5 | 0.957 | 0.957 | 299 | 0.323 | 0.848 | 641.5 | 0.853 | 0.895 | 504.5 | 0.474 | 0.895 | 4 | 0.051 | 0.356 | 209 | 0.684 | 0.895 | 196 | 0.617 |
| BRD-K39484304-001-16-5 | triptolide | 4729 | 0.620 | 0.930 | 8900 | 0.999 | 1.000 | 3681 | 0.378 | 0.793 | 11943 | 0.073 | 0.384 | 2822 | 0.005 | 0.101 | 4053 | 0.524 | 0.847 | 4380 | 0.069 | 0.384 | 8051 | 1.000 | 1.000 | 1455 | 0.350 | 0.793 | 3637 | 0.878 | 0.971 | 1666 | 0.017 | 0.182 | 1651 | 0.800 | 0.944 | 629 | 0.239 | 0.717 | 932 | 0.709 |
| BRD-K39558284-001-01-2 | SDZ-WAG-994 | 315 | 0.350 | 0.735 | 442 | 0.444 | 0.718 | 372 | 0.194 | 1.000 | 1384 | 0.803 | 0.934 | 802 | 0.890 | 0.934 | 416 | 0.184 | 0.543 | 407 | 0.759 | 0.934 | 448 | 0.694 | 0.934 | | | | 206 | 0.959 | 0.959 | 161 | 0.142 | 1.000 | 146 | 0.298 | 0.696 | 11 | 0.082 | 0.287 | 79 | 0.207 |
| BRD-K39841531-001-02-1 | TG-101209 | 2353 | 0.842 | 0.960 | 4826 | 0.960 | 0.960 | 2458 | 0.344 | 0.803 | 9154 | 0.547 | 1.000 | 2724 | 0.024 | 0.165 | 2209 | 0.945 | 0.960 | 3263 | 0.822 | 1.000 | 4199 | 0.956 | 0.960 | 554 | 0.003 | 0.039 | 1798 | 0.250 | 0.751 | 1778 | 0.231 | 0.751 | 777 | 0.031 | 0.165 | 665 | 0.873 | 0.960 | 540 | 0.643 |
| BRD-K39987650-001-24-1 | bisacodyl | 647 | 0.510 | 0.816 | 902 | 0.255 | 1.000 | 713 | 0.482 | 1.000 | 2587 | 0.645 | 0.816 | 1164 | 0.521 | 0.816 | 674 | 0.589 | 0.816 | 714 | 0.699 | 0.816 | 980 | 0.276 | 0.695 | 309 | 0.176 | 0.612 | 471 | 0.928 | 0.975 | 228 | 0.046 | 0.968 | 415 | 0.520 | 0.816 | 130 | 0.298 | 0.695 | 335 | 0.848 |
| BRD-K40109029-001-02-9 | SB-505124 | 314 | 0.663 | 0.870 | 1308.5 | 0.922 | 1.000 | 479 | 0.287 | 0.622 | 1914.5 | 0.843 | 1.000 | 742 | 0.360 | 0.622 | 501.5 | 0.357 | 0.622 | 510 | 0.385 | 0.622 | 487.5 | 0.162 | 0.622 | 349.5 | 0.511 | 0.766 | 305 | 0.323 | 0.622 | 179 | 0.327 | 0.622 | 165 | 0.656 | 0.870 | 35 | 0.193 | 0.622 | 131 | 0.132 |
| BRD-K40175214-001-11-6 | torin-1 | 7386 | 0.999 | 0.999 | 9618 | 0.983 | 0.999 | 4094 | 0.127 | 0.332 | 19698 | 0.966 | 0.999 | 4405 | 0.018 | 0.186 | 4882 | 0.564 | 0.911 | 7726 | 0.943 | 0.999 | 5507 | 0.036 | 0.255 | 1726 | 0.109 | 0.332 | 2838 | 0.092 | 0.332 | 2957 | 0.239 | 0.557 | 2552 | 0.397 | 0.695 | 524 | 0.076 | 0.332 | 1543 | 0.918 |
| BRD-K40255344-001-17-6 | tyrphostin-A9 | 3920 | 0.797 | 0.964 | 6825 | 0.996 | 0.996 | 2372 | 0.066 | 0.514 | 10972 | 0.073 | 0.514 | 3320 | 0.024 | 0.504 | 2880 | 0.369 | 0.753 | 4348 | 0.623 | 0.935 | 5717 | 0.996 | 0.996 | 1294 | 0.295 | 0.698 | 2330 | 0.673 | 0.942 | 1972 | 0.299 | 0.698 | 1550 | 0.826 | 0.964 | 402 | 0.216 | 0.698 | 1132 | 0.807 |
| BRD-K40308497-001-01-2 | crizotinib-(S) | 1896 | 0.761 | 0.932 | 2963 | 0.163 | 0.490 | 1388 | 0.033 | 0.345 | 7220 | 0.576 | 0.932 | 2167 | 0.107 | 0.450 | 3334 | 0.998 | 0.998 | 3379 | 0.959 | 0.998 | 2994 | 0.756 | 0.932 | 837 | 0.230 | 0.604 | 1670 | 0.094 | 0.450 | 1011 | 0.024 | 0.345 | 812 | 0.083 | 0.450 | 509 | 0.966 | 0.998 | 507 | 0.677 |
| BRD-K40331046-305-01-5 | CR8-(R) | 4407 | 0.224 | 0.522 | 9307 | 0.990 | 0.990 | 4386 | 0.319 | 0.670 | 18019 | 0.862 | 0.990 | 5328 | 0.142 | 0.483 | 4574 | 0.577 | 0.808 | 6235 | 0.388 | 0.710 | 7858 | 0.961 | 0.990 | 2291 | 0.405 | 0.710 | 3890 | 0.813 | 0.990 | 2199 | 0.018 | 0.371 | 1734 | 0.060 | 0.417 | 1259 | 0.926 | 0.990 | 828 | 0.175 |
| BRD-K40438880-305-06-6 | spermidine | 791 | 0.679 | 0.894 | 2304 | 0.681 | 0.894 | 999 | 0.296 | 0.831 | 4338 | 0.859 | 1.000 | 1738 | 0.631 | 0.894 | 1302 | 0.840 | 1.000 | 1067 | 0.424 | 0.831 | 1154 | 0.258 | 0.831 | 125 | 0.037 | 0.262 | 1108 | 0.993 | 1.000 | 485 | 0.073 | 0.383 | 168 | 0.005 | 0.105 | 95 | 0.435 | 0.831 | 131 | 0.182 |
| BRD-K40496271-396-01-1 | eltrombopag | 1324.5 | 0.992 | 0.995 | 1432 | 0.202 | 0.848 | 1184 | 0.336 | 1.000 | 4335 | 0.926 | 0.995 | 1074.5 | 0.046 | 0.848 | 338 | 0.100 | 0.848 | 1543 | 0.962 | 0.995 | 2036 | 0.716 | 0.939 | 305 | 0.191 | 0.365 | 404 | 0.009 | 0.020 | 598 | 0.471 | 1.000 | 498 | 0.466 | 0.699 | 77 | 0.330 | 0.539 | 51 | 0.034 |
| BRD-K40530731-001-11-6 | hyoscyamine | 1039 | 0.258 | 1.000 | 1846 | 0.103 | 1.000 | 810 | 0.323 | 0.688 | 3674 | 0.468 | 0.820 | 1831 | 0.440 | 0.820 | 169 | 0.022 | 0.076 | 1404 | 0.598 | 0.944 | 1608 | 0.952 | 0.952 | 546 | 0.877 | 0.944 | 257 | 0.328 | 0.688 | 1041 | 0.262 | 0.687 | 27 | 0.010 | 0.041 | 318 | 0.899 | 0.944 | 46 | 0.227 |
| BRD-K40718343-001-02-6 | AEE788 | 454.5 | 0.734 | 0.998 | 661.5 | 0.998 | 0.998 | 243.5 | 0.113 | 0.637 | 1165.5 | 0.602 | 0.981 | 491 | 0.849 | 0.998 | 210.5 | 0.369 | 0.775 | 442.5 | 0.688 | 0.998 | 369.5 | 0.852 | 0.998 | 69 | 0.608 | 0.981 | 242 | 0.549 | 0.981 | 377.5 | 0.219 | 0.511 | 108.5 | 0.920 | 0.998 | | | | | |
| BRD-K41104902-001-01-3 | GSK2656157 | 435.5 | 0.978 | 0.981 | 1445.5 | 0.897 | 0.981 | 1205.5 | 0.904 | 0.981 | 2464.5 | 0.420 | 0.980 | 576 | 0.249 | 0.682 | 800.5 | 0.630 | 1.000 | 967.5 | 0.182 | 0.682 | 1023.5 | 0.967 | 0.981 | 92 | 0.001 | 0.030 | 581 | 0.260 | 0.682 | 802 | 0.823 | 0.981 | 441 | 0.981 | 0.981 | 144 | 0.869 | 0.981 | 205.5 | 0.673 |
| BRD-K41141507-001-16-2 | cyproterone-acetate | 1497 | 0.431 | 0.731 | 2374 | 0.556 | 0.731 | 1294 | 0.159 | 1.000 | 4452 | 0.489 | 0.731 | 1542 | 0.347 | 0.731 | 1800 | 0.938 | 0.944 | 2270 | 0.944 | 0.944 | 1263 | 0.382 | 0.731 | 358 | 0.232 | 0.696 | 1049 | 0.210 | 0.696 | 643 | 0.200 | 1.000 | 461 | 0.504 | 0.731 | 254 | 0.592 | 0.731 | 254 | 0.592 |
| BRD-K41213548-001-02-0 | KW-2478 | 4361 | 0.642 | 0.968 | 5669 | 0.355 | 0.677 | 3453 | 0.150 | 0.449 | 14916 | 0.829 | 0.968 | 5314 | 0.779 | 0.968 | 3891 | 0.703 | 0.968 | 6753 | 0.974 | 1.000 | 6355 | 0.972 | 1.000 | 1491 | 0.042 | 0.297 | 3020.5 | 1.000 | 1.000 | 1595 | 0.004 | 0.047 | 931.5 | 0.071 | 0.371 | 614 | 0.218 | 0.509 | 563 | 0.433 |
| BRD-K41278234-001-03-0 | CUDC-101 | 253 | 0.286 | 0.601 | 512 | 0.837 | 0.925 | 221 | 0.156 | 0.546 | 954 | 0.749 | 0.874 | 127 | 0.110 | 0.546 | 195 | 0.085 | 0.546 | 320 | 0.108 | 0.546 | 626 | 0.989 | 0.993 | | | | 346 | 0.993 | 0.993 | 136 | 0.046 | 0.546 | | | | 94 | 0.430 | 0.626 | 47 | 0.447 |
| BRD-K41337261-001-03-0 | ZM-306416 | 729 | 0.692 | 0.996 | 414 | 0.968 | 0.996 | 845 | 0.236 | 1.000 | 2704.5 | 0.864 | 0.996 | 1517.5 | 0.477 | 0.996 | 554.5 | 0.421 | 0.981 | 992 | 0.736 | 0.996 | 711.5 | 0.996 | 0.996 | 233 | 0.813 | 0.996 | 430 | 0.279 | 0.733 | 553 | 0.104 | 0.725 | 251.5 | 0.875 | 0.996 | 89 | 0.528 | 0.996 | 312.5 | 0.978 |
| BRD-K41599323-001-02-3 | lonafarnib | 1145 | 0.832 | 0.874 | 1626 | 0.986 | 0.986 | 1292.5 | 0.375 | 0.874 | 4113.5 | 0.056 | 0.393 | 1627.5 | 0.611 | 0.874 | 994 | 0.774 | 0.874 | 1815 | 0.324 | 0.874 | 1504.5 | 0.571 | 0.874 | 478 | 0.624 | 0.874 | 868.5 | 0.750 | 0.874 | 377 | 0.012 | 0.254 | 328.5 | 0.496 | 0.874 | 104 | 0.100 | 0.523 | 195 | 0.389 |
| BRD-K41783120-001-02-8 | STA-5326 | 1608.5 | 0.029 | 0.154 | 3358 | 1.000 | 1.000 | 2307.5 | 0.886 | 1.000 | 6735.5 | 0.955 | 1.000 | 2210.5 | 0.173 | 0.399 | 1009 | 0.732 | 1.000 | 1731.5 | 0.827 | 1.000 | 2321 | 0.985 | 1.000 | 1165 | 0.557 | 0.900 | 1039.5 | 0.006 | 0.039 | 745.5 | 0.002 | 0.032 | 829 | 0.190 | 0.399 | 224 | 0.371 | 0.650 | 221 | 0.099 |
| BRD-K41859756-001-06-8 | NVP-AUY922 | 3463 | 0.878 | 0.994 | 6817 | 0.989 | 0.994 | 2929 | 0.124 | 0.523 | 10814 | 0.307 | 0.768 | 4205 | 0.456 | 0.863 | 2206 | 0.591 | 0.886 | 5552 | 0.994 | 0.994 | 3727 | 0.329 | 0.768 | 1289 | 0.010 | 0.216 | 3154 | 0.962 | 0.994 | 2025 | 0.584 | 0.886 | 1387 | 0.242 | 0.757 | 264 | 0.252 | 0.757 | 720 | 0.493 |
| BRD-K42191735-001-05-3 | buparlisib | 4081 | 0.885 | 0.998 | 6731 | 0.973 | 0.998 | 2880 | 0.046 | 0.324 | 11067 | 0.201 | 0.560 | 2747 | 0.267 | 0.560 | 3581 | 0.813 | 0.998 | 5351 | 0.901 | 0.998 | 6543 | 0.998 | 0.998 | 818 | 0.005 | 0.111 | 2850 | 0.993 | 0.998 | 1955 | 0.082 | 0.325 | 1054 | 0.093 | 0.325 | 849 | 0.696 | 0.974 | 654 | 0.357 |
| BRD-K42205652-001-02-7 | MNS-(3,4-Methylenedioxy-nitrostyrene) | 314 | 0.868 | 0.995 | 492.5 | 0.313 | 0.547 | 226 | 0.402 | 0.649 | 1128.5 | 0.995 | 0.995 | 333 | 0.298 | 0.547 | 255 | 0.848 | 0.995 | 388 | 0.924 | 0.995 | 159 | 0.034 | 0.142 | 72 | 0.253 | 0.547 | 367.5 | 0.974 | 0.995 | 17 | 0.023 | 0.488 | 35 | 0.305 | 0.547 | 68 | 0.730 | 0.972 | 23 | 0.032 |
| BRD-K42436189-001-01-2 | AZ20 | 2629 | 0.122 | 0.367 | 4985 | 0.839 | 0.961 | 2131 | 0.091 | 0.637 | 10705 | 0.844 | 0.961 | 2708 | 0.014 | 0.057 | 3084.5 | 0.862 | 0.961 | 5197 | 0.957 | 0.961 | 3375 | 0.322 | 0.687 | 1104 | 0.182 | 0.477 | 2718.5 | 0.849 | 0.961 | 1199 | 0.009 | 0.180 | 948 | 0.376 | 0.717 | 823 | 0.961 | 0.961 | 501 | 0.055 |
| BRD-K42497885-001-01-1 | satraplatin | 483 | 0.251 | 1.000 | 2031.5 | 0.648 | 0.960 | 792.5 | 0.202 | 0.606 | 2697.5 | 0.185 | 0.606 | 1632 | 0.387 | 0.903 | 568.5 | 0.480 | 0.960 | 1341 | 0.880 | 0.960 | 1667.5 | 0.918 | 0.960 | 212 | 0.016 | 0.085 | 844.5 | 0.704 | 0.960 | 739 | 0.254 | 0.666 | 138 | 0.003 | 0.022 | 323.5 | 0.944 | 0.960 | 132 | 0.052 |
| BRD-K42673188-001-01-1 | verubulin | 4860 | 0.645 | 0.903 | 7068 | 0.956 | 1.000 | 3508 | 0.086 | 0.300 | 14855 | 0.194 | 0.408 | 5776 | 0.619 | 0.903 | 4458 | 0.935 | 1.000 | 7828 | 1.000 | 1.000 | 6547 | 0.527 | 0.852 | 1250 | 0.015 | 0.189 | 4308 | 0.951 | 1.000 | 2010 | 0.026 | 0.189 | 1692 | 0.154 | 0.377 | 560 | 0.128 | 0.377 | 562 | 0.036 |
| BRD-K42805893-001-04-9 | osimertinib | 898 | 0.473 | 1.000 | 2479 | 1.000 | 1.000 | 691 | 0.007 | 0.058 | 3039 | 0.714 | 1.000 | 1469 | 0.467 | 1.000 | 453 | 0.089 | 0.313 | 1452 | 0.727 | 1.000 | 875 | 0.943 | 1.000 | 328 | 0.681 | 1.000 | 811 | 0.688 | 1.000 | 612 | 0.211 | 0.633 | 330 | 0.964 | 1.000 | 112 | 0.620 | 1.000 | 136 | 0.770 |
| BRD-K42898655-001-01-8 | temsirolimus | 824 | 1.000 | 1.000 | 234 | 0.309 | 1.000 | 161 | 0.188 | 0.920 | 890 | 0.104 | 0.920 | 341 | 0.088 | 0.920 | 291 | 0.989 | 1.000 | 517 | 0.865 | 1.000 | 290 | 0.347 | 0.662 | 154 | 0.052 | 0.136 | 269 | 0.790 | 0.976 | 210 | 0.453 | 0.793 | 59 | 0.543 | 0.815 | 58 | 0.531 | 0.815 | 33 | 0.243 |
| BRD-K42981054-001-01-9 | mibampator | 1757 | 0.770 | 0.988 | 2991 | 0.531 | 0.988 | 1555 | 0.110 | 0.862 | 5258 | 0.384 | 0.988 | 1395 | 0.037 | 0.775 | 1103 | 0.164 | 0.862 | 1929 | 0.536 | 0.988 | 2402 | 0.823 | 0.988 | 831 | 0.432 | 0.988 | 1239 | 0.894 | 0.988 | 1097 | 0.491 | 0.988 | 104 | 0.382 | 0.988 | 360 | 0.853 | 0.988 | 132 | 0.124 |
| BRD-K43002771-034-01-6 | SR-33805 | 3344 | 0.393 | 0.743 | 4539 | 0.347 | 0.743 | 2708 | 0.488 | 0.743 | 9465 | 0.566 | 0.787 | 3239 | 0.305 | 0.743 | 2029 | 0.258 | 0.743 | 5503 | 0.997 | 0.997 | 3734 | 0.729 | 0.901 | 816 | 0.476 | 0.743 | 2527 | 0.959 | 0.997 | 1235 | 0.040 | 0.514 | 873 | 0.311 | 0.743 | 294 | 0.387 | 0.743 | 365 | 0.219 |
| BRD-K43187018-001-03-3 | GDC-0349 | 6481.5 | 1.000 | 1.000 | 8308.5 | 0.829 | 1.000 | 4320 | 0.329 | 0.865 | 16370 | 0.528 | 0.916 | 4415 | 0.038 | 0.264 | 5141 | 0.911 | 1.000 | 6637 | 0.956 | 1.000 | 6862 | 0.703 | 0.984 | 2251 | 0.611 | 0.916 | 2858 | 0.149 | 0.447 | 2124 | 0.015 | 0.185 | 1765 | 0.080 | 0.279 | 310 | 0.059 | 0.272 | 1528 | 0.943 |
| BRD-K43236057-001-07-1 | polydatin | 3212 | 0.970 | 0.981 | 4841.5 | 0.754 | 0.981 | 2265 | 0.151 | 0.580 | 9266 | 0.437 | 0.706 | 3571 | 0.300 | 0.603 | 2392 | 0.533 | 0.800 | 3027 | 0.127 | 0.580 | 4759 | 0.981 | 0.981 | 845 | 0.127 | 0.580 | 1215 | 0.972 | 0.981 | 1810 | 0.571 | 0.800 | 827 | 0.266 | 0.603 | 981 | 0.959 | 0.981 | 239 | 0.255 |
| BRD-K43389675-001-02-1 | daunorubicin | 4467 | 0.996 | 0.996 | 5224 | 0.587 | 0.996 | 2098 | 0.003 | 0.063 | 10753 | 0.651 | 0.996 | 3543 | 0.139 | 0.485 | 2721 | 0.835 | 0.996 | 3895 | 0.984 | 0.996 | 4808 | 0.967 | 0.996 | 922 | 0.020 | 0.106 | 2486 | 0.905 | 0.996 | 1656 | 0.066 | 0.278 | 386 | 0.015 | 0.106 | 411 | 0.274 | 0.718 | 613 | 0.350 |
| BRD-K43389675-003-20-9 | daunorubicin | 4642 | 0.833 | 0.998 | 6570 | 0.280 | 0.619 | 3731 | 0.345 | 0.619 | 13212 | 0.183 | 0.619 | 4158 | 0.067 | 0.468 | 5466 | 0.998 | 0.998 | 6533 | 0.968 | 0.998 | 6576 | 0.841 | 0.998 | 1318 | 0.011 | 0.199 | 4054 | 0.998 | 0.998 | 2406 | 0.186 | 0.619 | 1636 | 0.152 | 0.619 | 446 | 0.215 | 0.619 | 703 | 0.318 |
| BRD-K43389698-001-09-9 | BMS-387032 | 5842 | 0.658 | 1.000 | 8814 | 0.921 | 1.000 | 4139 | 0.089 | 0.373 | 14929 | 0.046 | 0.245 | 5787 | 0.389 | 0.816 | 5114 | 0.908 | 1.000 | 6519 | 0.271 | 0.666 | 8579 | 0.994 | 1.000 | 2144 | 0.127 | 0.443 | 5887 | 1.000 | 1.000 | 3219 | 0.286 | 0.666 | 3432 | 0.958 | 1.000 | 664 | 0.189 | 0.568 | 1468 | 0.876 |
| BRD-K43449605-001-01-0 | oncrasin-1 | 2418 | 0.135 | 0.453 | 6545 | 0.997 | 0.997 | 2458.5 | 0.264 | 0.504 | 11885 | 0.767 | 0.988 | 3153 | 0.081 | 0.453 | 2379.5 | 0.493 | 0.859 | 3729 | 0.687 | 0.962 | 4475.5 | 0.917 | 0.997 | 1094.5 | 0.576 | 0.865 | 2383.5 | 0.207 | 0.453 | 1937 | 0.216 | 0.453 | 989.5 | 0.198 | 0.453 | 364 | 0.532 | 0.859 | 211 | 0.164 |
| BRD-K43471985-001-03-0 | OTX015 | 1844.5 | 0.770 | 0.770 | 2097.5 | 0.621 | 1.000 | 1371.5 | 0.906 | 1.000 | 3380 | 0.410 | 0.861 | 1579.5 | 0.151 | 0.517 | 1276 | 0.917 | 1.000 | 1793 | 0.965 | 1.000 | 741 | 0.163 | 0.517 | 458 | 0.325 | 0.775 | 391.5 | 0.172 | 0.517 | 812.5 | 0.143 | 0.517 | 111 | 0.027 | 0.517 | | | | 68 | 0.049 |
| BRD-K43578482-001-01-4 | defactinib | 651.5 | 0.498 | 0.805 | 1138.5 | 0.974 | 0.974 | 686 | 0.798 | 0.974 | 2314 | 0.884 | 0.974 | 624 | 0.242 | 0.564 | 414 | 0.193 | 0.564 | 368 | 0.040 | 0.208 | 990 | 0.924 | 0.974 | 425 | 0.706 | 0.974 | 556 | 0.115 | 0.484 | 155 | 0.050 | 0.840 | 59 | 0.351 | 0.614 | 259 | 0.957 | 0.974 | 92 | 0.641 |
| BRD-K43621685-001-02-5 | CP-673451 | 156.5 | 0.758 | 0.933 | 255 | 0.587 | 1.000 | 294.5 | 0.244 | 0.732 | 1622.5 | 0.700 | 1.000 | 641 | 0.397 | 1.000 | 364 | 0.549 | 1.000 | 531 | 0.909 | 0.933 | 574 | 0.113 | 0.732 | 138.5 | 0.075 | 0.732 | 122 | 0.012 | 0.262 | 77 | 0.187 | 0.732 | 70 | 0.151 | 0.732 | 171.5 | 0.841 | 0.933 | 196.5 | 0.933 |

| broad_id | name | al_nervous_system_PVAL_ADJ | fibroblast_MWU_STAT | fibroblast_PVAL | fibroblast_PVAL_ADJ | bile_duct_MWU_STAT | bile_duct_PVAL | bile_duct_PVAL_ADJ | thyroid_MWU_STAT | thyroid_PVAL | thyroid_PVAL_ADJ | upper_aerodigestive_MWU_STAT | upper_aerodigestive_PVAL | upper_aerodigestive_PVAL_ADJ | uterus_MWU_STAT | uterus_PVAL | uterus_PVAL_ADJ | liver_MWU_STAT | liver_PVAL | liver_PVAL_ADJ | esophagus_MWU_STAT | esophagus_PVAL | esophagus_PVAL_ADJ |
|---|---|---|---|---|---|---|---|---|---|---|---|---|---|---|---|---|---|---|---|---|---|---|---|
| BRD-K31866293-001-01-6 | TAK-632 | 0.362 | 124 | 0.309 | 0.775 | 1499 | 0.633 | 0.993 | 1137 | 0.086 | 0.362 | 3503 | 0.249 | 0.775 | 3270 | 0.477 | 0.910 | 3455 | 0.773 | 0.997 | 4657 | 0.797 | 0.997 |
| BRD-K31928526-001-02-1 | barasertib | 0.180 | | | | | | | 26 | 0.116 | 0.489 | 966 | 0.695 | 1.000 | 406.5 | 0.462 | 1.000 | 228 | 0.385 | 1.000 | 792 | 0.953 | 0.953 |
| BRD-K32372024-001-01-0 | D-7193 | 0.962 | | | | 127 | 0.166 | 0.870 | 541 | 0.839 | 0.927 | 1087 | 0.040 | 0.839 | 1366 | 0.771 | 0.927 | 660.5 | 0.598 | 1.000 | 1367 | 0.611 | 0.927 |
| BRD-K32501161-300-06-2 | vanoxerine | 0.047 | | | | 112 | 0.088 | 0.805 | 536 | 0.232 | 0.811 | 1914 | 0.125 | 0.805 | 1453 | 0.722 | 0.891 | 844 | 0.840 | 0.922 | 2533 | 0.878 | 0.922 |
| BRD-K32572029-001-04-4 | orotic-acid | | | | | 127 | 0.690 | 1.000 | 70 | 0.050 | 0.450 | 381 | 0.847 | 0.988 | 126 | 0.279 | 0.807 | 185 | 0.966 | 1.000 | 486 | 0.993 | 0.993 |
| BRD-K32730239-001-01-0 | SU3327 | 0.747 | | | | 105 | 0.718 | 0.794 | 445 | 0.725 | 0.895 | 742 | 0.493 | 0.895 | 592 | 0.464 | 0.895 | 382 | 0.490 | 0.895 | 787 | 0.984 | 0.984 |
| BRD-K32744045-001-31-2 | disulfiram | 0.382 | 162 | 0.315 | 0.685 | 859 | 0.187 | 0.491 | 2959.5 | 0.967 | 0.971 | 4845 | 0.577 | 0.865 | 4342 | 0.516 | 0.833 | 4347 | 0.891 | 0.971 | 3773 | 0.055 | 0.382 |
| BRD-K32821942-001-21-3 | azathioprine | 0.091 | | | | 123 | 0.852 | 0.979 | 287 | 0.415 | 0.792 | 486 | 0.625 | 0.875 | 666 | 0.895 | 0.979 | 448 | 0.483 | 0.831 | 704 | 0.240 | 0.631 |
| BRD-K33127281-001-05-6 | adapalene | 0.940 | | | | 70 | 0.759 | 0.940 | 139 | 0.838 | 0.940 | 232 | 0.135 | 0.945 | 139 | 0.838 | 0.940 | 147 | 0.437 | 0.940 | 413 | 0.621 | 0.940 |
| BRD-K33277808-331-01-3 | ouabain | 0.938 | 57 | 0.114 | 0.601 | 1158 | 0.554 | 0.832 | 1074 | 0.211 | 0.634 | 2222 | 0.200 | 0.634 | 2340 | 0.170 | 0.634 | 1970 | 0.263 | 0.690 | 3794 | 0.520 | 0.832 |
| BRD-K33379087-001-07-5 | tivantinib | 0.898 | | | | 185.5 | 0.941 | 0.980 | 648.5 | 0.718 | 0.980 | 759 | 0.311 | 0.884 | 771 | 0.337 | 0.884 | 969.5 | 0.780 | 0.980 | 258 | 0.001 | 0.018 |
| BRD-K33610132-001-02-9 | rociletinib | 0.509 | 125 | 0.921 | 0.977 | | | | 305 | 0.930 | 0.977 | 526 | 0.007 | 0.075 | 375 | 0.202 | 0.529 | 337 | 0.511 | 0.976 | 476 | 0.037 | 0.196 |
| BRD-K33622447-066-01-9 | abemaciclib | 0.108 | | | | 211 | 0.949 | 0.984 | 813 | 0.878 | 0.984 | 1493 | 0.959 | 0.984 | 1402 | 0.783 | 0.984 | 956 | 0.534 | 0.984 | 894 | 0.622 | 0.984 |
| BRD-K33720171-001-01-0 | difluprednate | 0.963 | 118 | 0.360 | 0.945 | 469 | 0.920 | 0.966 | 378 | 0.108 | 0.483 | 2031 | 0.078 | 0.483 | 1841 | 0.469 | 0.963 | 1374 | 0.115 | 0.483 | 2966 | 0.792 | 0.963 |
| BRD-K33743361-003-02-6 | ML133 | 0.906 | | | | 648 | 0.551 | 0.890 | 710 | 0.694 | 0.910 | 2028 | 0.647 | 0.906 | 1099 | 0.285 | 1.000 | 701 | 0.079 | 0.278 | 1402 | 0.403 | 0.823 |
| BRD-K33813875-003-01-8 | valnemulin | 0.655 | 149 | 0.385 | 0.896 | 1179 | 0.894 | 0.994 | 1919.5 | 0.694 | 0.911 | 2113 | 0.005 | 0.108 | 1990 | 0.576 | 0.896 | 2081 | 0.486 | 0.896 | 3715 | 0.623 | 0.896 |
| BRD-K33882852-003-02-8 | ZK-93423 | 0.246 | | | | 179 | 0.891 | 0.923 | 66 | 0.574 | 0.923 | 281 | 0.014 | 0.293 | 149 | 0.323 | 0.754 | 172 | 0.862 | 0.923 | 299 | 0.293 | 0.754 |
| BRD-K34022604-001-06-6 | clofarabine | 0.889 | 4 | 0.012 | 0.086 | 1585 | 0.882 | 0.978 | 1639 | 0.291 | 0.889 | 4503 | 0.845 | 0.978 | 4648 | 0.902 | 0.978 | 3199 | 0.910 | 0.978 | 5239 | 0.978 | 0.978 |
| BRD-K34332569-001-01-3 | imidapril | 0.918 | 135 | 0.549 | 0.918 | | | | 792 | 0.607 | 0.918 | 1438 | 0.666 | 0.932 | 1271 | 0.063 | 0.663 | 811 | 0.199 | 0.918 | 2031 | 0.964 | 0.964 |
| BRD-K34851558-001-02-3 | parthenolide-(-) | 0.325 | | | | 296 | 0.307 | 0.921 | 527 | 0.146 | 0.812 | 1645 | 0.283 | 0.921 | 935 | 0.155 | 0.812 | 578 | 0.093 | 0.177 | 2121 | 0.949 | 0.980 |
| BRD-K35329391-334-01-7 | GZD824 | 0.990 | 251 | 0.624 | 0.990 | 1357 | 0.713 | 0.990 | 1688 | 0.220 | 0.690 | 2785 | 0.019 | 0.292 | 2802 | 0.043 | 0.292 | 3330 | 0.919 | 0.990 | 4576 | 0.857 | 0.990 |
| BRD-K35520305-001-17-7 | dacarbazine | 0.234 | | | | 82 | 0.108 | 0.569 | 623 | 0.999 | 0.999 | 468 | 0.037 | 0.569 | 493 | 0.505 | 1.000 | 359 | 0.312 | 1.000 | 1160 | 0.651 | 1.000 |
| BRD-K35960502-001-20-0 | niclosamide | 0.857 | 74 | 0.285 | 0.857 | 705 | 0.982 | 0.982 | 339 | 0.534 | 0.857 | 1131.5 | 0.632 | 0.857 | 749 | 0.130 | 0.857 | 707 | 0.197 | 0.857 | 1535 | 0.775 | 0.857 |
| BRD-K36021395-300-02-4 | ABT-702 | 0.377 | | | | 47 | 0.208 | 1.000 | 424 | 0.447 | 1.000 | 1125 | 0.165 | 1.000 | 564 | 0.372 | 1.000 | 445 | 0.264 | 0.616 | 575 | 0.656 | 0.911 |
| BRD-K36055864-001-19-2 | cycloheximide | 0.315 | | | | 505 | 0.387 | 0.665 | 1296 | 0.760 | 0.939 | 3189.5 | 0.331 | 0.665 | 3135 | 0.424 | 0.665 | 2067 | 0.224 | 0.665 | 2876 | 0.039 | 0.826 |
| BRD-K36198571-001-02-0 | WAY-170523 | 0.893 | | | | 429 | 0.968 | 0.968 | 654 | 0.629 | 0.889 | 1875 | 0.835 | 0.923 | 1358 | 0.266 | 1.000 | 1201 | 0.539 | 0.889 | 1780 | 0.571 | 0.889 |
| BRD-K36222532-001-01-8 | broxaldine | 0.861 | 119 | 0.301 | 0.776 | 800.5 | 0.956 | 0.997 | 1274 | 0.407 | 0.776 | 2858 | 0.587 | 0.861 | 2168 | 0.219 | 0.757 | 1879 | 0.224 | 0.757 | 3548 | 0.869 | 0.960 |
| BRD-K36269323-300-02-6 | asymmetrical-dimethylarginine | 0.280 | 5 | 0.048 | 0.280 | 1312.5 | 0.986 | 0.989 | 798 | 0.180 | 0.501 | 1842 | 0.224 | 0.501 | 1607 | 0.238 | 0.501 | 497 | 0.038 | 0.280 | 2533 | 0.567 | 0.915 |
| BRD-K36363294-001-04-2 | I-BET151 | 0.261 | | | | | | | 25 | 0.012 | 0.250 | 381 | 0.841 | 0.886 | 148 | 0.371 | 1.000 | 286 | 0.911 | 0.911 | 166 | 0.526 | 0.838 |
| BRD-K36386086-001-01-1 | dolutegravir | 0.924 | | | | 354 | 0.699 | 0.929 | 247 | 0.708 | 0.929 | 914 | 0.585 | 0.924 | 815 | 0.360 | 1.000 | 639 | 0.085 | 0.162 | 613 | 0.554 | 0.924 |
| BRD-K36529613-001-02-6 | PU-H71 | 1.000 | 91 | 0.228 | 0.478 | 561 | 0.159 | 0.477 | 1531 | 0.115 | 0.401 | 2939 | 0.038 | 0.381 | 3244 | 0.621 | 0.931 | 1991 | 0.183 | 0.478 | 3490 | 0.091 | 0.381 |
| BRD-K36740062-001-06-6 | GSK1070916 | 0.986 | 114 | 0.357 | 0.720 | 1185 | 0.849 | 0.986 | 1089 | 0.479 | 0.739 | 2165 | 0.218 | 0.720 | 1246 | 0.278 | 0.720 | 2434 | 0.902 | 0.986 | 2696 | 0.322 | 0.720 |
| BRD-K36889451-001-01-1 | tetramethylthiuram-monosulfide | 0.728 | 121 | 0.506 | 0.856 | 58 | 0.033 | 0.493 | 1014 | 0.965 | 0.997 | 1173 | 0.525 | 0.856 | 1471 | 0.925 | 0.997 | 869 | 0.602 | 0.856 | 1387 | 0.697 | 0.856 |
| BRD-K37270826-001-27-6 | mifepristone | | | | | 429 | 0.774 | 0.937 | 276 | 0.911 | 0.937 | 739 | 0.640 | 0.937 | 375 | 0.015 | 0.315 | 739 | 0.839 | 0.937 | 1050 | 0.423 | 0.808 |
| BRD-K37312348-001-15-0 | kenpaullone | 0.717 | | | | 166 | 0.849 | 0.891 | 137.5 | 0.204 | 1.000 | 443.5 | 0.406 | 0.717 | 535 | 0.802 | 0.891 | 247.5 | 0.897 | 0.897 | 448.5 | 0.691 | 0.891 |
| BRD-K37379014-001-02-0 | filanesib | 0.667 | | | | 356 | 0.980 | 0.980 | 565.5 | 0.768 | 1.000 | 899.5 | 0.268 | 0.951 | 890.5 | 0.252 | 0.951 | 382.5 | 0.837 | 0.952 | 862.5 | 0.770 | 0.952 |
| BRD-K37561857-001-15-5 | zardaverine | 0.628 | 271 | 0.771 | 0.967 | 1496 | 0.967 | 0.967 | 1155 | 0.097 | 0.628 | 3173 | 0.500 | 0.967 | 3219 | 0.905 | 0.967 | 1770 | 0.569 | 0.967 | 4053 | 0.895 | 0.967 |
| BRD-K37618799-001-03-0 | MRS-1220 | | | | | 75 | 0.587 | 0.958 | 174 | 0.777 | 0.958 | 268.5 | 0.256 | 1.000 | 522.5 | 0.928 | 0.958 | 271 | 0.039 | 0.117 | 606 | 0.153 | 1.000 |
| BRD-K37687095-001-06-9 | AZD8330 | 0.267 | | | | 1386 | 0.457 | 1.000 | 1405 | 0.481 | 1.000 | 2599 | 0.002 | 0.016 | 3872 | 0.975 | 1.000 | 3360 | 0.653 | 1.000 | 5204 | 0.902 | 1.000 |
| BRD-K37720887-001-07-8 | SB-525334 | | | | | 43 | 0.337 | 0.443 | | | | 220 | 0.479 | 1.000 | 235 | 0.875 | 1.000 | 45 | 0.360 | 1.000 | 372 | 0.988 | 0.988 |
| BRD-K37723606-001-01-5 | dioscin | 0.932 | | | | 680 | 0.832 | 0.932 | 472 | 0.364 | 0.764 | 1984 | 0.458 | 0.802 | 1289 | 0.311 | 1.000 | 920 | 0.758 | 0.932 | 2453 | 0.341 | 0.764 |
| BRD-K37764012-001-04-1 | PF-03758309 | 0.903 | | | | 1601 | 0.981 | 1.000 | 958 | 0.518 | 0.903 | 2957 | 0.311 | 0.654 | 1636 | 0.129 | 1.000 | 1975 | 0.642 | 0.903 | 2663 | 0.188 | 0.565 |
| BRD-K37798499-001-27-2 | etoposide | 0.976 | 68 | 0.160 | 0.560 | 834 | 0.986 | 0.992 | 1490 | 0.604 | 0.976 | 2724 | 0.762 | 0.992 | 1883 | 0.027 | 0.187 | 1621 | 0.570 | 0.976 | 1834 | 0.286 | 0.751 |
| BRD-K37865504-001-04-1 | LY2183240 | 0.267 | 111 | 0.441 | 0.674 | 452 | 0.768 | 0.949 | 475 | 0.482 | 0.674 | 1511 | 0.211 | 0.582 | 1402 | 0.336 | 0.587 | 403 | 0.288 | 0.582 | 2154 | 0.982 | 0.998 |
| BRD-K37890730-001-15-1 | camptothecin | 0.184 | 49 | 0.133 | 0.548 | 790 | 0.931 | 0.991 | 368 | 0.003 | 0.058 | 1477 | 0.183 | 0.548 | 1946.5 | 0.300 | 0.582 | 1057.5 | 0.250 | 0.582 | 1917 | 0.163 | 0.548 |
| BRD-K38183498-001-02-0 | clonazepam | 0.645 | 96 | 0.235 | 0.645 | 1347 | 0.665 | 0.974 | 1772 | 0.258 | 0.645 | 2339 | 0.001 | 0.024 | 2807 | 0.066 | 0.459 | 3366 | 0.904 | 1.000 | 4439 | 0.600 | 0.974 |
| BRD-K38305202-001-17-4 | domperidone | 0.828 | | | | 119.5 | 0.592 | 0.965 | 118.5 | 0.583 | 0.965 | 367.5 | 0.752 | 0.965 | 18 | 0.114 | 0.853 | 85.5 | 0.840 | 0.965 | 590 | 0.981 | 0.981 |
| BRD-K38332599-001-01-3 | uprosertib | 0.999 | 117 | 0.444 | 0.932 | 160 | 0.669 | 0.999 | 565 | 0.925 | 0.999 | 1271 | 0.026 | 0.137 | 1012 | 0.002 | 0.043 | 1583 | 0.830 | 0.999 | 1016 | 0.036 | 0.149 |
| BRD-K38350380-001-01-8 | FK-3311 | 0.888 | | | | 567 | 0.761 | 0.888 | 915 | 0.902 | 0.959 | 1526 | 0.760 | 0.888 | 1149 | 0.181 | 1.000 | 1288 | 0.749 | 0.888 | 1294 | 0.386 | 0.879 |
| BRD-K38527262-300-01-0 | atiprimod | 0.985 | 129 | 0.542 | 0.985 | 246 | 0.170 | 0.713 | 522 | 0.613 | 0.985 | 1269 | 0.470 | 0.985 | 1273 | 0.845 | 0.985 | 833 | 0.137 | 0.713 | 1670 | 0.419 | 0.985 |
| BRD-K38548312-001-01-0 | VER-49009 | 0.452 | 67 | 0.194 | 0.452 | 1169 | 0.909 | 0.977 | 1109 | 0.181 | 0.452 | 2576 | 0.024 | 0.452 | 2043 | 0.054 | 0.452 | 1834 | 0.457 | 0.739 | 2710 | 0.247 | 0.506 |
| BRD-K38852836-001-02-1 | ganetespib | 0.463 | 112 | 0.246 | 0.748 | 763 | 0.864 | 0.907 | 1703 | 0.570 | 0.835 | 3227 | 0.155 | 0.748 | 2457.5 | 0.278 | 0.748 | 1865 | 0.102 | 0.748 | 2676 | 0.116 | 0.748 |
| BRD-K38860038-001-01-2 | JK-184 | 0.487 | 259 | 0.630 | 0.997 | 1714 | 0.997 | 0.997 | 1795 | 0.278 | 0.558 | 2165 | 0.001 | 0.020 | 3684 | 0.319 | 0.558 | 3237 | 0.979 | 0.997 | 3480 | 0.194 | 0.487 |
| BRD-K38868394-001-01-3 | MK-0773 | 0.119 | | | | | | | 64 | 0.660 | 0.963 | 298 | 0.027 | 0.283 | 117 | 0.247 | 1.000 | 60 | 0.159 | 0.335 | 259 | 0.302 | 1.000 |
| BRD-K38904395-001-02-3 | SAR131675 | 0.714 | | | | 176 | 0.708 | 0.876 | 193 | 0.126 | 1.000 | 533 | 0.660 | 1.000 | 548 | 0.900 | 0.900 | 162 | 0.620 | 1.000 | 686 | 0.709 | 0.876 |
| BRD-K39171998-001-01-1 | STF-118804 | 0.136 | | | | 1235 | 0.575 | 0.755 | 1531 | 0.455 | 0.683 | 3329 | 0.278 | 1.000 | 3897 | 0.437 | 0.683 | 2084.5 | 0.147 | 0.441 | 3293 | 0.157 | 1.000 |
| BRD-K39252998-003-01-5 | atipamezole | 0.895 | | | | 217.5 | 0.725 | 0.895 | 86 | 0.488 | 0.895 | 1102.5 | 0.746 | 0.895 | 894.5 | 0.824 | 0.895 | 207 | 0.085 | 0.359 | 593.5 | 0.139 | 1.000 |
| BRD-K39484304-001-16-5 | triptolide | 0.944 | 326 | 0.809 | 0.944 | 976 | 0.227 | 0.717 | 1721 | 0.436 | 0.816 | 3810 | 0.756 | 0.944 | 3015 | 0.172 | 0.717 | 2329 | 0.304 | 0.793 | 3977 | 0.466 | 0.816 |
| BRD-K39558284-001-01-2 | SDZ-WAG-994 | 0.543 | | | | 169 | 0.441 | 0.718 | 246 | 0.870 | 0.934 | 452 | 0.486 | 0.728 | 92 | 0.020 | 0.410 | 6 | 0.063 | 0.263 | 606 | 0.831 | 0.934 |
| BRD-K39841531-001-02-1 | TG-101209 | 0.960 | | | | 488 | 0.896 | 0.960 | 788 | 0.095 | 0.400 | 3248 | 0.812 | 1.000 | 1018 | 0.004 | 0.039 | 1706 | 0.302 | 0.792 | 3208 | 0.880 | 0.960 |
| BRD-K39987650-001-24-1 | bisacodyl | 0.938 | | | | 416 | 0.977 | 0.977 | 346 | 0.568 | 0.816 | 695 | 0.254 | 1.000 | 373 | 0.674 | 0.816 | 179 | 0.204 | 0.612 | 690 | 0.635 | 0.816 |
| BRD-K40109029-001-02-9 | SB-505124 | 0.622 | | | | 4 | 0.053 | 0.622 | 276.5 | 0.221 | 0.622 | 602 | 0.166 | 0.622 | 289.5 | 0.864 | 1.000 | 187 | 0.369 | 0.622 | 639.5 | 0.988 | 0.988 |
| BRD-K40175214-001-11-6 | torin-1 | 0.999 | 147 | 0.324 | 0.618 | 2030 | 0.903 | 0.999 | 2458 | 0.706 | 0.988 | 4091 | 0.126 | 0.332 | 4786 | 0.638 | 0.957 | 1829 | 0.001 | 0.011 | 4637 | 0.283 | 0.595 |
| BRD-K40255344-001-17-6 | tyrphostin-A9 | 0.964 | 224 | 0.603 | 0.935 | 1227 | 0.991 | 0.996 | 1552 | 0.395 | 0.753 | 3109 | 0.185 | 0.698 | 2701 | 0.227 | 0.698 | 2093 | 0.259 | 0.698 | 3272 | 0.569 | 0.935 |
| BRD-K40308497-001-01-2 | crizotinib-(S) | 0.932 | 43 | 0.150 | 0.490 | 327 | 0.614 | 0.932 | 838 | 0.725 | 0.932 | 1941 | 0.476 | 0.932 | 1933 | 0.799 | 0.932 | 1028 | 0.542 | 0.932 | 2727 | 0.626 | 0.932 |
| BRD-K40331046-305-01-5 | CR8-(R) | 0.483 | 81 | 0.184 | 0.483 | 1031 | 0.183 | 0.483 | 1968 | 0.502 | 0.808 | 4549 | 0.560 | 0.808 | 3261 | 0.135 | 0.483 | 2419 | 0.049 | 0.417 | 5201 | 0.712 | 0.935 |
| BRD-K40438880-305-06-6 | spermidine | 0.763 | | | | 313 | 0.500 | 0.875 | 554 | 0.608 | 0.894 | 557 | 0.024 | 0.252 | 764 | 0.381 | 0.831 | 561 | 0.351 | 0.831 | 1778 | 0.906 | 0.906 |
| BRD-K40496271-396-01-1 | eltrombopag | 0.071 | | | | 493.5 | 0.962 | 0.995 | 851 | 0.995 | 0.995 | 787 | 0.133 | 0.848 | 816.5 | 0.170 | 0.848 | 639 | 0.334 | 0.539 | 1018 | 0.550 | 0.770 |
| BRD-K40530731-001-11-6 | hyoscyamine | 0.681 | | | | 27 | 0.135 | 1.000 | 468 | 0.694 | 0.944 | 1158 | 0.826 | 0.944 | 1015 | 0.761 | 0.944 | 519 | 0.824 | 0.944 | 1492 | 0.866 | 0.944 |
| BRD-K40718343-001-02-6 | AEE788 | | | | | 33 | 0.040 | 0.425 | 193.5 | 0.937 | 0.998 | 422.5 | 0.004 | 0.084 | 463.5 | 0.989 | 0.998 | 156.5 | 0.041 | 0.108 | 562.5 | 0.121 | 0.637 |
| BRD-K41104902-001-01-3 | GSK2656157 | 0.981 | | | | 250.5 | 0.859 | 0.981 | 111 | 0.191 | 0.682 | 561.5 | 0.007 | 0.069 | 757 | 0.728 | 1.000 | 343.5 | 0.088 | 0.614 | 938.5 | 0.247 | 0.682 |
| BRD-K41141507-001-16-2 | cyproterone-acetate | 0.731 | | | | 376 | 0.274 | 1.000 | 580 | 0.521 | 0.731 | 1101 | 0.850 | 0.944 | 1142 | 0.543 | 0.731 | 539 | 0.009 | 0.037 | 1793 | 0.934 | 0.944 |
| BRD-K41213548-001-02-0 | KW-2478 | 0.759 | 45 | 0.091 | 0.382 | 1151 | 0.735 | 0.968 | 1405 | 0.304 | 0.638 | 2564 | 0.001 | 0.025 | 2759 | 0.140 | 0.449 | 2289 | 0.822 | 0.968 | 3902 | 0.205 | 0.509 |
| BRD-K41278234-001-03-0 | CUDC-101 | 0.626 | | | | 210 | 0.570 | 0.749 | 127 | 0.727 | 0.874 | 137 | 0.145 | 0.546 | 155 | 0.226 | 0.678 | 346 | 0.934 | 1.000 | 132 | 0.356 | 0.575 |
| BRD-K41337261-001-03-0 | ZM-306416 | 0.996 | | | | 399.5 | 0.954 | 0.996 | 148 | 0.895 | 0.996 | 629 | 0.001 | 0.029 | 374 | 0.056 | 0.585 | 411.5 | 0.230 | 0.733 | 1260 | 0.252 | 0.733 |
| BRD-K41599323-001-02-3 | lonafarnib | 0.874 | | | | 166 | 0.806 | 0.874 | 786.5 | 0.809 | 0.874 | 1407 | 0.575 | 0.874 | 1355.5 | 0.661 | 0.874 | 770 | 0.534 | 0.874 | 1283 | 0.532 | 0.874 |
| BRD-K41783120-001-02-8 | STA-5326 | 0.346 | | | | 162 | 0.179 | 0.399 | 524 | 0.858 | 1.000 | 1042 | 0.085 | 0.346 | 1086 | 0.233 | 0.445 | 886 | 0.130 | 0.391 | 1796.5 | 0.760 | 1.000 |
| BRD-K41859756-001-06-8 | NVP-AUY922 | 0.863 | 280 | 0.823 | 0.994 | 829 | 0.692 | 0.908 | 1572 | 0.683 | 0.908 | 2994 | 0.091 | 0.502 | 2688 | 0.394 | 0.827 | 1539 | 0.052 | 0.502 | 2709 | 0.096 | 0.502 |
| BRD-K42191735-001-05-3 | buparlisib | 0.681 | 179 | 0.484 | 0.847 | 592 | 0.258 | 0.560 | 1078 | 0.244 | 0.560 | 2267 | 0.020 | 0.205 | 2508 | 0.677 | 0.974 | 2084 | 0.611 | 0.974 | 3157 | 0.078 | 0.325 |
| BRD-K42205652-001-02-7 | MNS-(3,4-Methylenedioxy-nitrostyrene) | 0.142 | | | | 69 | 0.741 | 0.972 | 197 | 0.510 | 0.765 | 281.5 | 0.252 | 0.547 | 106 | 0.061 | 0.645 | 49 | 0.111 | 0.390 | 173 | 0.142 | 0.427 |
| BRD-K42436189-001-01-2 | AZ20 | 0.194 | | | | 1004 | 0.745 | 0.961 | 1625 | 0.827 | 0.961 | 3693 | 0.959 | 0.961 | 2179 | 0.059 | 0.615 | 1706 | 0.327 | 0.687 | 3135 | 0.655 | 0.961 |
| BRD-K42497885-001-01-1 | satraplatin | 0.217 | | | | 409 | 0.883 | 0.960 | 575.5 | 0.771 | 0.960 | 1289 | 0.817 | 0.960 | 595.5 | 0.558 | 0.960 | 585 | 0.960 | 0.960 | 908.5 | 0.637 | 0.960 |
| BRD-K42673188-001-01-1 | verubulin | 0.189 | 68 | 0.162 | 0.377 | 2151 | 0.983 | 1.000 | 1282 | 0.051 | 0.215 | 3872 | 0.357 | 0.682 | 2872 | 0.028 | 0.189 | 3370 | 0.854 | 1.000 | 4202 | 0.456 | 0.799 |
| BRD-K42805893-001-04-9 | osimertinib | 1.000 | | | | 191 | 0.508 | 1.000 | 345.5 | 0.977 | 1.000 | 673 | 0.002 | 0.045 | 202 | 0.198 | 1.000 | 801.5 | 0.858 | 1.000 | 898 | 0.008 | 0.058 |
| BRD-K42898655-001-01-8 | temsirolimus | 0.511 | | | | 64 | 0.604 | 0.845 | 247 | 0.931 | 1.000 | 369 | 0.734 | 0.964 | 332 | 0.162 | 0.920 | 312 | 0.241 | 0.511 | 168 | 0.219 | 0.920 |
| BRD-K42981054-001-01-9 | mibampator | 0.862 | | | | 155 | 0.654 | 0.988 | 496 | 0.219 | 0.921 | 1724 | 0.566 | 0.988 | 1657 | 0.796 | 0.988 | 1209 | 0.689 | 0.988 | 1954 | 0.848 | 0.848 |
| BRD-K43002771-034-01-6 | SR-33805 | 0.743 | 201 | 0.599 | 0.787 | 663 | 0.495 | 0.743 | 997 | 0.775 | 0.904 | 2205 | 0.049 | 0.514 | 1970 | 0.114 | 0.743 | 1501 | 0.180 | 0.743 | 3607 | 0.922 | 0.997 |
| BRD-K43187018-001-03-3 | GDC-0349 | 1.000 | 251 | 0.603 | 0.916 | 1347 | 0.566 | 0.916 | 2405 | 0.897 | 1.000 | 4504 | 0.451 | 0.916 | 3817 | 0.381 | 0.890 | 2329 | 0.018 | 0.185 | 3520 | 0.065 | 0.272 |
| BRD-K43236057-001-07-1 | polydatin | 0.603 | 54 | 0.124 | 0.580 | 243 | 0.265 | 0.603 | 1369 | 0.395 | 0.692 | 2288 | 0.166 | 0.580 | 1829 | 0.070 | 0.580 | 1476.5 | 0.917 | 0.981 | 2901 | 0.316 | 0.603 |
| BRD-K43389675-001-02-1 | daunorubicin | 0.796 | 159 | 0.456 | 0.870 | 686 | 0.818 | 0.996 | 1446 | 0.379 | 0.796 | 3432 | 0.741 | 0.996 | 2091 | 0.015 | 0.106 | 1502 | 0.261 | 0.718 | 2853 | 0.669 | 0.996 |
| BRD-K43389675-003-20-9 | daunorubicin | 0.619 | 99 | 0.242 | 0.619 | 1827 | 0.898 | 0.998 | 1522 | 0.383 | 0.619 | 3759 | 0.365 | 0.619 | 2694 | 0.019 | 0.199 | 2374 | 0.463 | 0.695 | 4688 | 0.755 | 0.998 |
| BRD-K43389698-001-09-9 | BMS-387032 | 1.000 | 363 | 0.800 | 1.000 | 1389 | 0.552 | 0.965 | 2215 | 0.491 | 0.937 | 3569 | 0.047 | 0.245 | 2981 | 0.004 | 0.078 | 3789 | 0.836 | 1.000 | 4079 | 0.041 | 0.245 |
| BRD-K43449605-001-01-0 | oncrasin-1 | 0.453 | 311 | 0.905 | 0.997 | 630.5 | 0.142 | 0.453 | 1036 | 0.043 | 0.453 | 3389.5 | 0.800 | 0.988 | 2036 | 0.009 | 0.198 | 1569 | 0.168 | 0.453 | 4095 | 0.981 | 0.997 |
| BRD-K43471985-001-03-0 | OTX015 | 0.074 | | | | 79 | 0.332 | 0.775 | 462 | 0.643 | 1.000 | 765 | 0.094 | 0.517 | 881 | 0.123 | 0.517 | 1327.5 | 0.989 | 1.000 | 1338.5 | 0.741 | 1.000 |
| BRD-K43578482-001-01-4 | defactinib | 0.961 | | | | 43 | 0.229 | 0.564 | 54 | 0.310 | 0.604 | 684 | 0.794 | 0.974 | 411 | 0.080 | 0.840 | 257.5 | 0.317 | 0.604 | 590.5 | 0.172 | 0.564 |
| BRD-K43621685-001-02-5 | CP-673451 | 0.933 | | | | 107.5 | 0.905 | 0.933 | 227.5 | 0.777 | 0.933 | 343 | 0.236 | 0.732 | 271.5 | 0.931 | 0.933 | 60 | 0.489 | 1.000 | 564.5 | 0.850 | 0.933 |

Suppl. Table 9

| broad_id | name | colorectal_MWU_STAT | colorectal_PVAL | colorectal_PVAL_ADJ | skin_MWU_STAT | skin_PVAL | skin_PVAL_ADJ | urinary_tract_MWU_STAT | urinary_tract_PVAL | urinary_tract_PVAL_ADJ | lung_MWU_STAT | lung_PVAL | lung_PVAL_ADJ | ovary_MWU_STAT | ovary_PVAL | ovary_PVAL_ADJ | breast_MWU_STAT | breast_PVAL | breast_PVAL_ADJ | pancreas_MWU_STAT | pancreas_PVAL | pancreas_PVAL_ADJ | nervous_system_MWU_STAT | central_nervous_system_PVAL | nervous_system_PVAL_ADJ | soft_tissue_MWU_STAT | soft_tissue_PVAL | soft_tissue_PVAL_ADJ | kidney_MWU_STAT | kidney_PVAL | kidney_PVAL_ADJ | gastric_MWU_STAT | gastric_PVAL | gastric_PVAL_ADJ | bone_MWU_STAT | bone_PVAL | bone_PVAL_ADJ | prostate_MWU_STAT | prostate_PVAL | prostate_PVAL_ADJ | peripheral_nervous_system_MWU_STAT | peripheral_nervous_system_PVAL |
|---|---|---|---|---|---|---|---|---|---|---|---|---|---|---|---|---|---|---|---|---|---|---|---|---|---|---|---|---|---|---|---|---|---|---|---|---|---|---|---|---|---|---|
| BRD-K43644456-001-04-0 | WP1130 | 5222 | 0.653 | 0.998 | 7397 | 0.242 | 0.584 | 4579 | 0.258 | 0.584 | 19121 | 0.894 | 0.998 | 3933 | 0.006 | 0.074 | 6549 | 0.998 | 0.998 | 6542 | 0.695 | 0.998 | 7883 | 0.937 | 0.998 | 2201 | 0.284 | 0.584 | 4287 | 0.852 | 0.998 | 2734 | 0.127 | 0.584 | 1594 | 0.009 | 0.074 | 1190 | 0.856 | 0.998 | 974 | 0.306 |
| BRD-K43905307-001-01-2 | L-798,106 | 449 | 0.175 | 1.000 | 976.5 | 0.413 | 0.820 | 364 | 0.054 | 1.000 | 2249 | 0.509 | 0.820 | 517 | 0.077 | 0.322 | 1122 | 0.867 | 0.955 | 853 | 0.460 | 0.820 | 1061.5 | 0.447 | 0.820 | 588 | 0.996 | 0.996 | 422 | 0.546 | 0.820 | 507 | 0.318 | 0.820 | 477.5 | 0.737 | 0.915 | 98 | 0.618 | 0.865 | 245.5 | 0.191 |
| BRD-K44084986-300-08-4 | Y-27632 | 1542 | 0.769 | 1.000 | 2569 | 0.585 | 0.945 | 1552 | 0.162 | 0.426 | 6012 | 0.872 | 1.000 | 2161 | 0.093 | 0.389 | 1741 | 0.706 | 1.000 | 2746 | 0.970 | 1.000 | 2257 | 0.232 | 0.538 | 450 | 0.138 | 0.426 | 778 | 0.143 | 0.426 | 884 | 0.060 | 0.340 | 113 | 0.015 | 0.287 | 183 | 0.256 | 0.538 | 243 | 0.449 |
| BRD-K44227013-001-06-4 | ponatinib | 2669 | 0.987 | 0.987 | 2958 | 0.081 | 0.840 | 2337 | 0.475 | 0.946 | 7044 | 0.865 | 1.000 | 2586 | 0.161 | 0.840 | 1510 | 0.738 | 1.000 | 2403 | 0.823 | 1.000 | 2256 | 0.380 | 0.946 | 228 | 0.000 | 0.006 | 1476 | 0.495 | 0.946 | 1632 | 0.869 | 1.000 | 601 | 0.316 | 0.946 | 200 | 0.775 | 1.000 | 377 | 0.808 |
| BRD-K44408410-001-17-6 | 2-methoxyestradiol | 597 | 0.731 | 0.963 | 812 | 0.866 | 0.963 | 712 | 0.436 | 0.654 | 2337 | 0.747 | 0.963 | 1097 | 0.301 | 1.000 | 532.5 | 0.994 | 0.994 | 758 | 0.903 | 0.963 | 426 | 0.404 | 0.654 | 195 | 0.109 | 0.255 | 430 | 0.917 | 0.963 | 354 | 0.399 | 0.654 | 173 | 0.070 | 0.184 | 122 | 0.837 | 0.963 | 212 | 0.406 |
| BRD-K44590731-001-01-7 | TU-2100 | 376 | 0.707 | 0.942 | 292 | 0.306 | 1.000 | 409 | 0.214 | 1.000 | 1337 | 0.897 | 0.942 | 390 | 0.015 | 0.061 | 382 | 0.142 | 0.425 | 517 | 0.814 | 0.942 | 481 | 0.691 | 0.942 | 18 | 0.118 | 0.413 | 196 | 0.672 | 0.942 | 158 | 0.152 | 1.000 | 235 | 0.870 | 0.942 | 39 | 0.289 | 0.760 | 102 | 0.395 |
| BRD-K44665581-001-01-8 | bromosporine | 376.5 | 0.088 | 1.000 | 630.5 | 0.533 | 0.747 | 383 | 0.863 | 0.975 | 1298.5 | 0.475 | 0.712 | 504 | 0.450 | 0.712 | 271 | 0.928 | 0.975 | 798 | 0.997 | 0.997 | 521.5 | 0.723 | 0.948 | 74 | 0.041 | 0.143 | 261 | 0.904 | 0.975 | 268 | 0.356 | 0.682 | 156 | 0.342 | 0.682 | 5 | 0.059 | 0.154 | 42 | 0.057 |
| BRD-K44844162-001-01-6 | taselisib | 847.5 | 0.022 | 0.078 | 1962 | 0.994 | 1.000 | 1239.5 | 0.145 | 0.434 | 5559.5 | 1.000 | 1.000 | 2121.5 | 0.574 | 1.000 | 628.5 | 0.012 | 0.057 | 1062 | 0.007 | 0.057 | 1595 | 0.917 | 1.000 | 235 | 0.013 | 0.057 | 542 | 0.466 | 0.994 | 564.5 | 0.005 | 0.057 | 281.5 | 0.732 | 1.000 | 329.5 | 0.873 | 1.000 | 67 | 0.243 |
| BRD-K45293975-001-02-0 | 7-hydroxystaurosporine | 5167 | 0.837 | 0.925 | 5715 | 0.441 | 0.847 | 4441 | 0.742 | 0.917 | 12764 | 0.572 | 0.851 | 3862 | 0.084 | 0.534 | 3642 | 0.966 | 0.991 | 5771 | 0.689 | 0.904 | 4239 | 0.102 | 0.534 | 1379 | 0.302 | 0.847 | 1795 | 0.003 | 0.060 | 2226 | 0.147 | 0.617 | 1644 | 0.392 | 0.847 | 405 | 0.531 | 0.851 | 602 | 0.528 |
| BRD-K45401373-001-20-9 | betulinic-acid | 733 | 0.454 | 0.914 | 759 | 0.110 | 1.000 | 835 | 0.876 | 0.952 | 1763 | 0.081 | 1.000 | 990 | 0.747 | 0.952 | 480 | 0.566 | 0.914 | 1014 | 0.482 | 0.914 | 998 | 0.763 | 0.952 | 145 | 0.129 | 0.300 | 122 | 0.019 | 0.050 | 514 | 0.430 | 1.000 | 163 | 0.534 | 0.914 | 398 | 0.980 | 0.980 | | |
| BRD-K45528773-001-07-2 | M-344 | 5447 | 0.754 | 0.990 | 7457 | 0.571 | 0.856 | 4925 | 0.693 | 0.970 | 15538 | 0.375 | 0.656 | 4671 | 0.071 | 0.214 | 5463 | 0.923 | 1.000 | 8570 | 1.000 | 1.000 | 8116 | 0.986 | 1.000 | 1484 | 0.017 | 0.088 | 4673 | 0.994 | 1.000 | 1751 | 0.004 | 0.064 | 1860 | 0.109 | 0.287 | 332 | 0.014 | 0.088 | 406 | 0.006 |
| BRD-K45551181-001-02-1 | MPI-0479605 | 3291 | 0.188 | 0.498 | 7948 | 0.996 | 0.996 | 3017 | 0.134 | 0.498 | 12904 | 0.616 | 0.957 | 4440 | 0.190 | 0.498 | 3759 | 0.996 | 0.996 | 4968 | 0.638 | 0.957 | 5407 | 0.587 | 0.957 | 2006 | 0.602 | 0.957 | 3175 | 0.895 | 0.996 | 2034 | 0.258 | 0.602 | 853 | 0.015 | 0.162 | 1097 | 0.921 | 0.996 | 626 | 0.087 |
| BRD-K45724504-001-01-6 | anguidine | 4909 | 0.454 | 0.784 | 7193 | 0.467 | 0.784 | 5263 | 0.938 | 0.986 | 14720 | 0.126 | 0.615 | 4431 | 0.021 | 0.438 | 5415 | 0.986 | 0.986 | 6405 | 0.439 | 0.784 | 7670 | 0.976 | 0.986 | 1720 | 0.026 | 0.078 | 4230 | 0.985 | 0.986 | 2758 | 0.191 | 0.502 | 3102 | 0.894 | 0.986 | 667 | 0.219 | 0.510 | 1101 | 0.523 |
| BRD-K46392623-001-02-5 | TG-100713 | 1736.5 | 0.922 | 0.922 | 3572 | 0.997 | 1.000 | 1813 | 0.600 | 1.000 | 5344.5 | 0.186 | 0.639 | 2074.5 | 0.139 | 0.595 | 1526 | 0.532 | 1.000 | 2493 | 0.709 | 1.000 | 2774 | 0.924 | 1.000 | 670 | 0.293 | 0.639 | 715 | 0.075 | 0.527 | 1483 | 0.823 | 1.000 | 322.5 | 0.305 | 0.639 | 288 | 0.219 | 0.639 | 190 | 0.255 |
| BRD-K46690591-001-01-2 | KP-1212 | 1031 | 0.965 | 0.984 | 1385 | 0.283 | 1.000 | 657 | 0.285 | 1.000 | 3253 | 0.357 | 1.000 | 1501 | 0.479 | 1.000 | 352 | 0.002 | 0.036 | 794 | 0.399 | 1.000 | 1566 | 0.930 | 0.984 | 97 | 0.130 | 1.000 | 795 | 0.822 | 0.984 | 580 | 0.302 | 1.000 | 315 | 0.104 | 0.198 | 142 | 0.274 | 0.443 | 101 | 0.140 |
| BRD-K47000838-001-01-6 | cytochalasin-b | 3907 | 0.711 | 0.879 | 7296 | 1.000 | 1.000 | 2279 | 0.024 | 0.332 | 10315 | 0.061 | 0.332 | 3790 | 0.063 | 0.332 | 2806 | 0.653 | 0.879 | 4974 | 0.910 | 1.000 | 4386 | 0.576 | 0.879 | 1400 | 0.248 | 0.692 | 1919 | 0.450 | 0.879 | 2582 | 0.606 | 0.879 | 1227 | 0.241 | 0.692 | 637 | 0.966 | 1.000 | 595 | 0.597 |
| BRD-K47079459-004-04-6 | cetrimonium | 3485 | 0.294 | 0.687 | 7156 | 0.997 | 0.997 | 2494 | 0.174 | 0.731 | 12303 | 0.528 | 0.997 | 2734 | 0.002 | 0.037 | 4450 | 0.966 | 0.997 | 5226 | 0.615 | 0.997 | 4988 | 0.779 | 0.997 | 1337 | 0.070 | 0.184 | 1784 | 0.064 | 0.184 | 1727 | 0.020 | 0.214 | 1278 | 0.725 | 0.997 | 971 | 0.842 | 0.997 | 1003 | 0.875 |
| BRD-K47114202-001-14-6 | chlorogenic-acid | 608 | 0.540 | 1.000 | 676 | 0.337 | 1.000 | 582 | 0.264 | 1.000 | 2222 | 0.574 | 0.751 | 691 | 0.119 | 0.834 | 347 | 0.373 | 1.000 | 1164 | 0.999 | 0.999 | 407 | 0.608 | 0.751 | 119 | 0.076 | 0.834 | 318 | 0.555 | 0.751 | 390 | 0.542 | 0.751 | 254 | 0.105 | 0.183 | 217 | 0.431 | 0.697 | 6 | 0.057 |
| BRD-K47150025-001-05-3 | KI-8751 | 1182.5 | 0.777 | 0.902 | 2505.5 | 0.902 | 0.902 | 1213 | 0.654 | 0.902 | 4067.5 | 0.392 | 0.823 | 2083 | 0.874 | 0.902 | 659.5 | 0.826 | 0.902 | 1270.5 | 0.888 | 0.902 | 1697 | 0.154 | 0.540 | 123 | 0.002 | 0.009 | 1336 | 0.693 | 0.902 | 398 | 0.020 | 0.414 | 301 | 0.050 | 0.209 | 190 | 0.391 | 0.823 | 190 | 0.391 |
| BRD-K47337578-034-02-5 | 4-(4-fluorobenzoyl)-1-(4-phenylbutyl)-piperidine | 430 | 0.568 | 0.884 | 387 | 0.019 | 0.397 | 480 | 0.921 | 0.967 | 1085 | 0.679 | 0.947 | 416 | 0.502 | 1.000 | 62 | 0.589 | 0.884 | 88 | 0.856 | 0.967 | 565 | 0.722 | 0.947 | 261 | 0.967 | 0.967 | 115 | 0.184 | 0.352 | 349 | 0.411 | 1.000 | 87 | 0.304 | 0.532 | | | | 23 | 0.152 |
| BRD-K47598052-001-15-8 | PP-1 | 389 | 0.796 | 1.000 | 337 | 0.817 | 0.925 | 337 | 0.309 | 1.000 | 1176 | 0.468 | 1.000 | 340 | 0.571 | 1.000 | 295 | 0.881 | 0.925 | 480 | 0.728 | 1.000 | 316 | 0.015 | 0.299 | 343 | 0.839 | 0.925 | 61 | 0.557 | 1.000 | 270 | 0.474 | 1.000 | 28 | 0.028 | 0.299 | 88 | 0.841 | 0.925 | 98 | 0.372 |
| BRD-K47832606-001-30-1 | floxuridine | 3090 | 0.002 | 0.037 | 8234 | 0.965 | 0.987 | 3965 | 0.212 | 0.496 | 15741 | 0.769 | 0.987 | 4800 | 0.187 | 0.496 | 4775 | 0.987 | 0.987 | 6033 | 0.868 | 0.987 | 5884 | 0.288 | 0.550 | 2057 | 0.164 | 0.496 | 3438 | 0.900 | 0.987 | 2136 | 0.024 | 0.247 | 1831 | 0.271 | 0.550 | 533 | 0.324 | 0.566 | 478 | 0.070 |
| BRD-K47869605-001-32-0 | podophyllotoxin | 6422 | 0.921 | 0.979 | 8120 | 0.737 | 0.967 | 4343 | 0.391 | 0.684 | 14724 | 0.022 | 0.350 | 5842 | 0.349 | 0.667 | 5503 | 0.957 | 0.979 | 7795 | 0.969 | 0.979 | 7112 | 0.725 | 0.967 | 2180 | 0.291 | 0.667 | 4196 | 0.922 | 0.979 | 3687 | 0.663 | 0.967 | 2098 | 0.229 | 0.667 | 753 | 0.303 | 0.667 | 713 | 0.086 |
| BRD-K47976015-001-01-2 | balapiravir | 687 | 0.779 | 0.905 | 1254 | 0.882 | 0.905 | 496 | 0.105 | 0.743 | 1954 | 0.052 | 0.743 | 748 | 0.221 | 1.000 | 466 | 0.560 | 1.000 | 692.5 | 0.792 | 0.905 | 1052 | 0.895 | 0.905 | | | | 376.5 | 0.972 | 1.000 | 679.5 | 0.760 | 1.000 | 0 | 0.044 | 0.091 | 143 | 0.434 | 0.760 | 6 | 0.057 |
| BRD-K48443249-001-01-0 | CNX-774 | 148 | 0.443 | 0.962 | 422.5 | 0.000 | 0.006 | 200 | 0.206 | 0.865 | 1063 | 0.943 | 0.980 | 375.5 | 0.843 | 0.980 | 130.5 | 0.729 | 0.980 | 508.5 | 0.914 | 0.980 | 512 | 0.618 | 0.980 | 100 | 0.458 | 0.962 | 56 | 0.546 | 0.980 | 101 | 0.147 | 0.773 | 191 | 0.753 | 0.980 | 88.5 | 0.883 | 0.980 | 26 | 0.191 |
| BRD-K48488978-001-05-3 | YM-201636 | 291 | 0.083 | 0.869 | 975.5 | 0.923 | 0.979 | 659.5 | 0.355 | 0.702 | 1813 | 0.728 | 0.979 | 628 | 0.035 | 0.183 | 635.5 | 0.871 | 0.979 | 498.5 | 0.913 | 0.979 | 909.5 | 0.981 | 0.981 | 176 | 0.084 | 0.295 | 520 | 0.079 | 0.295 | 427.5 | 0.111 | 0.333 | 185.5 | 0.310 | 0.702 | 90 | 0.634 | 0.979 | 134 | 0.401 |
| BRD-K48684885-001-04-2 | BAY-11-7085 | 803 | 0.645 | 0.888 | 1905 | 0.377 | 0.753 | 1442 | 0.513 | 0.829 | 4392 | 0.860 | 0.959 | 1127 | 0.090 | 0.533 | 691 | 0.193 | 0.729 | 1770 | 0.928 | 0.975 | 1580.5 | 0.430 | 0.753 | 608 | 0.421 | 0.753 | 913 | 0.654 | 0.888 | 710 | 0.424 | 0.753 | 235 | 0.208 | 0.729 | 373.5 | 0.677 | 0.888 | 53 | 0.033 |
| BRD-K48988407-001-01-2 | cytochlor | 720.5 | 0.771 | 0.980 | 1290 | 0.000 | 0.004 | 898 | 0.078 | 0.422 | 4217.5 | 0.919 | 0.980 | 1839 | 0.891 | 0.980 | 1310.5 | 0.865 | 0.980 | 1177 | 0.664 | 0.980 | 1424 | 0.876 | 0.980 | 466 | 0.366 | 0.955 | 1049 | 0.409 | 0.955 | 571 | 0.080 | 0.422 | | | | 502.5 | 0.967 | 0.980 | 35 | 0.127 |
| BRD-K49027941-001-01-1 | PSB-1115 | 480 | 0.751 | 0.751 | 734.5 | 0.326 | 1.000 | 480 | 0.751 | 0.830 | 1263.5 | 0.605 | 1.000 | 480 | 0.751 | 0.830 | 418.5 | 0.154 | 1.000 | 480 | 0.751 | 0.830 | 363 | 0.126 | 1.000 | 63.5 | 0.605 | 1.000 | 189.5 | 0.000 | 0.004 | 248 | 0.684 | 1.000 | 187.5 | 0.662 | 1.000 | | | | 63.5 | 0.605 |
| BRD-K49049886-001-08-7 | CGS-15943 | 1080.5 | 0.202 | 0.596 | 2648 | 1.000 | 1.000 | 799 | 0.000 | 0.002 | 5676 | 0.931 | 1.000 | 2161 | 0.695 | 1.000 | 1718 | 0.047 | 0.198 | 1245 | 0.831 | 1.000 | 1298.5 | 0.996 | 1.000 | 751.5 | 0.977 | 1.000 | 763 | 0.865 | 1.000 | 1175.5 | 0.194 | 0.596 | 354.5 | 0.878 | 1.000 | 432.5 | 0.732 | 1.000 | 228.5 | 0.940 |
| BRD-K49075727-001-08-5 | nintedanib | 552 | 0.840 | 0.925 | 581 | 0.484 | 0.846 | 385 | 0.905 | 0.925 | 1378 | 0.795 | 0.925 | 569 | 0.221 | 0.515 | 458 | 0.703 | 0.925 | 537 | 0.788 | 0.925 | 401 | 0.166 | 0.498 | 78 | 0.119 | 0.416 | 189 | 0.030 | 0.159 | 159 | 0.291 | 0.610 | 306 | 0.200 | 0.515 | 24 | 0.084 | 0.354 | 79 | 0.722 |
| BRD-K49294207-300-03-8 | BIBU-1361 | 894 | 0.152 | 0.639 | 2106 | 0.233 | 0.730 | 1360 | 0.120 | 0.840 | 3946 | 0.944 | 0.970 | 1210 | 0.724 | 0.970 | 601 | 0.243 | 0.730 | 1813 | 0.637 | 0.970 | 1206 | 0.956 | 0.970 | 636 | 0.970 | 0.970 | 931 | 0.770 | 0.970 | 344 | 0.033 | 0.347 | 191 | 0.423 | 0.970 | 409 | 0.830 | 0.970 | 277 | 0.794 |
| BRD-K49313711-001-05-4 | ribitol | 279 | 0.611 | 0.950 | 611 | 0.613 | 0.950 | 163 | 0.081 | 0.849 | 889.5 | 0.397 | 0.926 | 760.5 | 0.900 | 0.950 | 107 | 0.049 | 0.172 | 219 | 0.561 | 0.950 | 432.5 | 0.950 | 0.950 | 80 | 0.800 | 0.950 | 284 | 0.885 | 0.950 | 6 | 0.065 | 0.849 | 86 | 0.849 | 0.950 | 169.5 | 0.923 | 0.950 | 12 | 0.016 |
| BRD-K49328571-001-15-0 | dasatinib | 1625 | 0.999 | 1.000 | 4016.5 | 1.000 | 1.000 | 1065.5 | 0.010 | 0.055 | 4009 | 0.533 | 0.861 | 832.5 | 0.000 | 0.001 | 1315.5 | 0.820 | 1.000 | 2105 | 0.964 | 1.000 | 1415.5 | 0.800 | 1.000 | 475.5 | 0.253 | 0.760 | 750 | 0.002 | 0.018 | 880 | 0.442 | 0.777 | 1199 | 0.989 | 1.000 | 520 | 0.931 | 1.000 | 483 | 0.879 |
| BRD-K49350383-001-13-7 | thioguanine | 3368 | 0.365 | 0.817 | 4952 | 0.367 | 0.817 | 2726 | 0.082 | 0.432 | 10428 | 0.186 | 0.779 | 4442 | 0.449 | 0.835 | 3769 | 0.957 | 0.999 | 5729 | 0.999 | 0.999 | 5798 | 0.992 | 0.999 | 1072 | 0.014 | 0.095 | 2401 | 0.389 | 0.817 | 2102 | 0.267 | 0.817 | 937 | 0.005 | 0.092 | 610 | 0.611 | 0.870 | 996 | 0.622 |
| BRD-K49350383-001-14-5 | thioguanine | 847 | 0.723 | 0.977 | 874 | 0.617 | 0.977 | 487.5 | 0.440 | 0.924 | 1692.5 | 0.112 | 0.554 | 684.5 | 0.437 | 0.924 | 785.5 | 0.971 | 0.977 | 877.5 | 0.977 | 0.977 | 750 | 0.641 | 0.977 | 120 | 0.003 | 0.049 | 498 | 0.740 | 0.977 | 83 | 0.005 | 0.049 | 232 | 0.224 | 0.784 | 113.5 | 0.830 | 0.977 | 84 | 0.132 |
| BRD-K49371609-003-03-8 | PIK-75 | 4575 | 0.329 | 0.732 | 6866 | 0.792 | 0.906 | 3944 | 0.282 | 0.732 | 15739 | 0.533 | 0.905 | 4856 | 0.124 | 0.520 | 5118 | 0.880 | 0.924 | 6363 | 0.735 | 0.906 | 8468 | 0.999 | 0.999 | 965 | 0.019 | 0.201 | 2665 | 0.661 | 0.906 | 2520 | 0.349 | 0.732 | 2511 | 0.710 | 0.906 | 863 | 0.527 | 0.905 | 686 | 0.093 |
| BRD-K49456274-001-01-9 | pilaralisib | 385 | 0.542 | 0.920 | 1099 | 0.471 | 0.920 | 824 | 0.411 | 0.920 | 2105 | 0.509 | 0.920 | 649 | 0.691 | 0.927 | 403 | 0.614 | 0.920 | 907 | 0.995 | 1.000 | 605 | 0.192 | 0.806 | 118 | 0.019 | 0.403 | 155 | 0.516 | 0.920 | 596 | 0.751 | 0.927 | 106 | 0.230 | 0.807 | 169 | 0.603 | 0.920 | | |
| BRD-K49522529-001-02-9 | alfacalcidol | 827 | 0.262 | 0.799 | 1619 | 0.772 | 0.900 | 804 | 0.112 | 1.000 | 3354 | 0.526 | 0.850 | 1132 | 0.714 | 0.900 | 1199 | 0.668 | 0.900 | 1779 | 0.930 | 0.939 | 1607 | 0.939 | 0.939 | 483 | 0.503 | 0.850 | 651 | 0.708 | 0.900 | 913 | 0.266 | 0.799 | 184 | 0.934 | 0.939 | 199 | 0.517 | 0.850 | 140 | 0.245 |
| BRD-K49669041-001-08-4 | BX-912 | 3800 | 0.874 | 0.968 | 5366 | 0.968 | 0.968 | 3090 | 0.470 | 0.871 | 11444 | 0.425 | 0.871 | 3703 | 0.087 | 0.459 | 2622 | 0.498 | 0.871 | 4717 | 0.945 | 0.968 | 4382 | 0.366 | 0.854 | 998 | 0.027 | 0.459 | 2482 | 0.940 | 0.968 | 1615 | 0.081 | 0.459 | 1556 | 0.059 | 0.459 | 649 | 0.711 | 0.933 | 780 | 0.310 |
| BRD-K49865102-001-08-4 | PD-0325901 | 2977 | 0.005 | 0.051 | 3054 | 0.000 | 0.000 | 3591 | 0.439 | 0.922 | 12503 | 0.990 | 1.000 | 5083 | 0.932 | 1.000 | 3564 | 0.987 | 1.000 | 3318 | 0.002 | 0.007 | 5515 | 1.000 | 1.000 | 1841 | 0.753 | 1.000 | 2263 | 0.918 | 1.000 | 2308 | 0.320 | 0.854 | 1055 | 0.936 | 1.000 | 1257 | 0.993 | 1.000 | 487 | 0.125 |
| BRD-K49954789-001-02-9 | mesna | 635 | 0.129 | 0.905 | 783.5 | 0.000 | 0.000 | 869 | 0.838 | 0.996 | 2360 | 0.780 | 0.996 | 1378 | 0.772 | 0.996 | 433.5 | 0.985 | 0.996 | 898 | 0.224 | 0.941 | 821 | 0.742 | 0.996 | 495 | 0.953 | 0.996 | 108 | 0.685 | 0.996 | 552 | 0.433 | 0.996 | 167.5 | 0.494 | 0.996 | 145.5 | 0.895 | 0.996 | 231 | 0.404 |
| BRD-K49962337-001-02-9 | baicalin | 246 | 0.503 | 0.887 | 691 | 0.830 | 0.948 | 378 | 0.167 | 1.000 | 1569 | 0.857 | 0.948 | 700.5 | 0.945 | 0.948 | 202 | 0.274 | 0.720 | 639.5 | 0.948 | 0.948 | 471 | 0.155 | 0.464 | 74 | 0.623 | 0.948 | 129 | 0.000 | 0.000 | 258 | 0.282 | 1.000 | 41 | 0.053 | 0.185 | 57 | 0.440 | 0.887 | | |
| BRD-K50000283-001-04-4 | PHA-767491 | 1705 | 0.058 | 0.614 | 3528 | 0.543 | 0.878 | 1946 | 0.483 | 0.878 | 7129 | 0.574 | 0.878 | 1733 | 0.034 | 0.614 | 2503 | 0.669 | 0.878 | 2920 | 0.426 | 0.878 | 2714 | 0.343 | 0.878 | 705 | 0.089 | 0.625 | 2199 | 0.961 | 0.961 | 1631 | 0.607 | 0.878 | 762 | 0.311 | 0.878 | 580 | 0.837 | 0.925 | 217 | 0.267 |
| BRD-K50010139-001-01-5 | poziotinib | 403 | 0.809 | 1.000 | 689 | 1.000 | 1.000 | 200 | 0.002 | 0.018 | 1113 | 0.774 | 1.000 | 450 | 0.358 | 1.000 | 84 | 0.260 | 1.000 | 447 | 0.922 | 1.000 | 152 | 0.785 | 1.000 | 309 | 0.895 | 1.000 | 595 | 0.958 | 1.000 | 384 | 0.498 | 0.926 | | | | | | | | |
| BRD-K50010139-001-02-3 | poziotinib | 1428 | 0.101 | 0.304 | 3930 | 1.000 | 1.000 | 1984 | 0.015 | 0.106 | 10658 | 0.991 | 1.000 | 2388 | 0.074 | 0.260 | 1804 | 0.022 | 0.111 | 3195 | 0.834 | 1.000 | 4783 | 1.000 | 1.000 | 1312 | 0.833 | 1.000 | 2478 | 0.943 | 1.000 | 979 | 0.026 | 0.111 | 741 | 0.958 | 1.000 | 276 | 0.385 | 0.736 | 164 | 0.531 |
| BRD-K50140147-001-10-1 | NVP-TAE684 | 1216 | 0.012 | 0.244 | 4006 | 0.979 | 0.986 | 2909 | 0.769 | 0.986 | 7553.5 | 0.343 | 0.720 | 2207 | 0.008 | 0.030 | 2935.5 | 0.982 | 0.986 | 3585 | 0.745 | 0.986 | 3337 | 0.971 | 0.986 | 1222 | 0.517 | 0.904 | 2291 | 0.986 | 0.986 | 1630 | 0.296 | 0.691 | 1232 | 0.165 | 0.495 | 657 | 0.896 | 0.986 | 135 | 0.004 |
| BRD-K50168500-001-07-9 | canertinib | 630 | 0.638 | 0.984 | 1477 | 1.000 | 1.000 | 599 | 0.033 | 0.293 | 2353 | 0.383 | 1.000 | 639 | 0.056 | 0.293 | 801 | 0.656 | 0.984 | 1401 | 0.869 | 1.000 | 705 | 0.995 | 1.000 | 578 | 0.731 | 1.000 | 289 | 0.649 | 0.984 | 693 | 0.584 | 0.984 | 486 | 0.996 | 1.000 | | | | | |
| BRD-K50677762-001-01-5 | oxiracetam | 3377 | 0.865 | 0.999 | 5393 | 0.696 | 0.914 | 3716 | 0.871 | 0.999 | 9872 | 0.509 | 0.764 | 3244 | 0.172 | 0.524 | 2584 | 0.468 | 0.764 | 4360 | 0.999 | 0.999 | 3159 | 0.129 | 0.524 | 756 | 0.018 | 0.297 | 2284 | 0.978 | 0.999 | 1655 | 0.081 | 0.524 | 1168 | 0.268 | 0.704 | 468 | 0.144 | 0.524 | 493 | 0.175 |
| BRD-K50853363-001-02-3 | dienogest | 578.5 | 0.665 | 0.961 | 971.5 | 0.413 | 1.000 | 879 | 0.686 | 0.961 | 1866 | 0.024 | 0.506 | 1137 | 0.503 | 1.000 | 469 | 0.963 | 0.988 | 429 | 0.201 | 1.000 | 997.5 | 0.903 | 0.988 | 173 | 0.226 | 0.474 | 343 | 0.653 | 0.961 | 300 | 0.078 | 0.823 | 159.5 | 0.525 | 0.918 | 222 | 0.449 | 0.858 | 147.5 | 0.936 |
| BRD-K51040301-001-02-1 | cobicistat | 197 | 0.274 | 0.635 | 462 | 0.162 | 1.000 | 486 | 0.383 | 0.730 | 1353 | 0.605 | 0.971 | 878 | 0.804 | 0.971 | 309 | 0.302 | 0.635 | 715 | 0.995 | 0.995 | 410 | 0.924 | 0.971 | 93 | 0.814 | 0.971 | 184 | 0.024 | 0.102 | 290 | 0.230 | 0.603 | 9 | 0.073 | 0.256 | 175 | 0.856 | 0.971 | 20 | 0.126 |
| BRD-K51313569-001-07-8 | palbociclib | 305.5 | 0.624 | 0.820 | 876.5 | 0.412 | 1.000 | 540.5 | 0.291 | 1.000 | 2152.5 | 0.909 | 0.950 | 580.5 | 0.413 | 1.000 | 383.5 | 0.898 | 0.950 | 593.5 | 0.857 | 0.950 | 805.5 | 0.950 | 0.950 | 167 | 0.024 | 0.051 | 492 | 0.015 | 0.035 | 329.5 | 0.424 | 0.685 | 54 | 0.341 | 0.596 | | | | 107 | 0.277 |
| BRD-K51313569-003-03-3 | palbociclib | 467.5 | 0.856 | 0.920 | 787.5 | 0.218 | 0.654 | 466 | 0.356 | 0.831 | 2335.5 | 0.973 | 0.973 | 912 | 0.841 | 0.920 | 783.5 | 0.869 | 0.920 | 648 | 0.516 | 0.920 | 295.5 | 0.085 | 0.368 | 177.5 | 0.082 | 0.368 | 415 | 0.088 | 0.368 | 125 | 0.022 | 0.368 | 281.5 | 0.183 | 0.641 | 124.5 | 0.876 | 0.920 | 338 | 0.692 |
| BRD-K51318897-001-15-3 | fenbendazole | 4312 | 0.766 | 0.946 | 5983 | 0.749 | 0.946 | 3267 | 0.357 | 0.750 | 12619 | 0.612 | 0.918 | 3557 | 0.217 | 0.750 | 3334 | 0.832 | 0.957 | 3843 | 0.417 | 0.750 | 5007 | 0.917 | 0.963 | 1608 | 0.118 | 0.718 | 3161 | 0.976 | 0.976 | 2321 | 0.428 | 0.750 | 1545 | 0.087 | 0.718 | 484 | 0.114 | 0.718 | 956 | 0.526 |
| BRD-K51324732-001-01-0 | meglitinide | 312 | 0.935 | 0.935 | 770 | 0.643 | 1.000 | 453 | 0.664 | 1.000 | 1179 | 0.740 | 1.000 | 429 | 0.206 | 0.758 | 87 | 0.839 | 0.979 | 227 | 0.046 | 0.758 | 130 | 0.276 | 0.758 | 202 | 0.759 | 1.000 | 223 | 0.547 | 1.000 | 169 | 0.084 | 0.758 | 180 | 0.289 | 0.758 | 173 | 0.919 | 1.000 | | |
| BRD-K51350053-048-16-5 | toremifene | 1555 | 0.215 | 0.602 | 3278 | 0.276 | 0.602 | 2460 | 0.810 | 0.978 | 6906 | 0.546 | 0.819 | 3159 | 0.615 | 0.861 | 2357 | 0.838 | 0.978 | 3541 | 0.978 | 0.983 | 2303 | 0.087 | 0.602 | 858 | 0.287 | 0.602 | 891 | 0.835 | 0.978 | 1394 | 0.507 | 0.819 | 618 | 0.903 | 0.983 | 297 | 0.191 | 0.602 | 444 | 0.235 |
| BRD-K51544265-001-04-2 | cabozantinib | 955 | 0.866 | 1.000 | 623.5 | 0.685 | 1.000 | 497.5 | 0.301 | 0.755 | 2257 | 0.323 | 0.755 | 1085.5 | 0.923 | 1.000 | 326 | 0.828 | 1.000 | 1240.5 | 0.825 | 1.000 | 604.5 | 0.003 | 0.069 | 348 | 0.120 | 0.658 | 776.5 | 0.668 | 1.000 | 436 | 0.157 | 0.658 | 351.5 | 0.302 | 0.755 | | | | 148 | 0.910 |
| BRD-K51575138-001-13-2 | TPCA-1 | 462 | 0.437 | 0.835 | 988.5 | 0.831 | 1.000 | 417 | 0.282 | 1.000 | 1741 | 0.723 | 1.000 | 736 | 0.344 | 1.000 | 572.5 | 0.993 | 1.000 | 578 | 0.950 | 1.000 | 899 | 0.781 | 1.000 | 101 | 0.062 | 0.163 | 424.5 | 0.544 | 0.952 | 365 | 0.314 | 1.000 | 127 | 0.120 | 0.280 | 102 | 0.772 | 1.000 | 94 | 0.209 |
| BRD-K51747290-001-13-1 | hydroxyurea | 416 | 0.485 | 0.869 | 745 | 0.514 | 0.869 | 392 | 0.097 | 0.676 | 1379.5 | 0.335 | 1.000 | 701 | 0.538 | 0.869 | 92 | 0.790 | 0.975 | 788 | 0.992 | 0.992 | 555.5 | 0.593 | 0.889 | 122.5 | 0.949 | 0.992 | 302.5 | 0.784 | 0.975 | 247 | 0.095 | 0.676 | 40 | 0.270 | 0.629 | 105 | 0.877 | 0.992 | 80 | 0.194 |
| BRD-K51791723-003-01-7 | P276-00 | 3210 | 0.202 | 0.607 | 6557 | 0.791 | 0.976 | 2818 | 0.049 | 0.518 | 12553 | 0.837 | 0.976 | 2764 | 0.002 | 0.038 | 3319 | 0.915 | 0.976 | 5215 | 0.866 | 0.976 | 6231 | 0.996 | 0.996 | 1664 | 0.152 | 0.582 | 2755 | 0.824 | 0.976 | 1800 | 0.135 | 0.582 | 1644 | 0.270 | 0.630 | 612 | 0.589 | 0.976 | 762 | 0.240 |
| BRD-K51911221-001-01-2 | nemorubicin | 6350 | 0.849 | 0.954 | 8401 | 0.576 | 0.954 | 4777 | 0.597 | 0.954 | 16419 | 0.191 | 0.574 | 4721 | 0.024 | 0.168 | 5870 | 0.954 | 0.954 | 8299 | 0.951 | 0.954 | 7088 | 0.505 | 0.954 | 2070 | 0.174 | 0.574 | 4556 | 0.924 | 0.954 | 2415 | 0.006 | 0.060 | 2235 | 0.154 | 0.574 | 1335 | 0.941 | 0.954 | 379 | 0.004 |
| BRD-K51967704-001-03-6 | BIIB021 | 3067 | 0.601 | 0.976 | 4959 | 0.998 | 0.998 | 2110 | 0.037 | 0.207 | 10658 | 0.907 | 0.998 | 2981 | 0.168 | 0.504 | 1949 | 0.543 | 0.976 | 4209 | 0.934 | 0.998 | 3750 | 0.791 | 0.977 | 1403 | 0.127 | 0.482 | 2303 | 0.959 | 0.998 | 1065 | 0.013 | 0.149 | 1000 | 0.138 | 0.482 | 398 | 0.697 | 0.976 | 596 | 0.737 |
| BRD-K52075715-001-06-7 | oxibendazole | 4243 | 0.971 | 0.980 | 6079 | 0.832 | 0.980 | 2573 | 0.337 | 0.885 | 9439 | 0.023 | 0.181 | 2738 | 0.020 | 0.181 | 2833 | 0.292 | 0.875 | 4465 | 0.946 | 0.980 | 5110 | 0.887 | 0.980 | 1265 | 0.789 | 0.980 | 2400 | 0.660 | 0.980 | 1904 | 0.174 | 0.610 | 854 | 0.084 | 0.442 | 589 | 0.615 | 0.980 | 440 | 0.026 |
| BRD-K52233191-001-02-4 | R547 | 2985 | 0.683 | 0.956 | 4387 | 0.976 | 0.995 | 2382 | 0.372 | 0.791 | 8379 | 0.071 | 0.375 | 2948 | 0.249 | 0.791 | 3089 | 0.943 | 0.995 | 3573 | 0.676 | 0.956 | 4232 | 0.948 | 0.995 | 435 | 0.002 | 0.039 | 2337 | 0.327 | 0.791 | 1301 | 0.311 | 0.791 | 1218 | 0.641 | 0.956 | 539 | 0.286 | 0.791 | 350 | 0.058 |
| BRD-K52256627-300-06-6 | chlorhexidine | 1122 | 0.775 | 1.000 | 2036 | 0.431 | 0.926 | 1535 | 0.827 | 1.000 | 5204 | 0.188 | 0.564 | 1951 | 0.612 | 0.994 | 753 | 0.146 | 0.552 | 2001 | 0.965 | 1.000 | 2303 | 0.375 | 0.926 | 761 | 0.155 | 0.552 | 917 | 0.636 | 0.994 | 937 | 0.441 | 0.926 | 507 | 0.034 | 0.346 | 148 | 0.596 | 0.994 | 108 | 0.015 |
| BRD-K52762805-001-01-2 | epothilone-d | 2114 | 0.332 | 0.862 | 4215 | 0.533 | 0.862 | 1499 | 0.144 | 0.840 | 8519 | 0.532 | 0.862 | 2979 | 0.649 | 0.862 | 2887 | 0.821 | 0.884 | 3150 | 0.675 | 0.862 | 3612 | 0.812 | 0.884 | 1202 | 0.884 | 0.884 | 2143 | 0.698 | 0.862 | 2302 | 0.843 | 0.884 | 557 | 0.013 | 0.268 | 189 | 0.182 | 0.840 | 535 | 0.650 |
| BRD-K52911425-001-09-8 | GDC-0941 | 3316 | 0.793 | 1.000 | 4048 | 0.925 | 1.000 | 2754 | 0.420 | 0.979 | 9030 | 0.677 | 1.000 | 3406 | 0.361 | 0.948 | 2371 | 0.485 | 1.000 | 3408 | 0.108 | 0.379 | 4469 | 0.979 | 1.000 | 598 | 0.005 | 0.095 | 1151 | 0.009 | 0.095 | 1673 | 0.056 | 0.295 | 679 | 0.098 | 0.379 | 208 | 0.183 | 0.550 | 701 | 0.590 |
| BRD-K53414658-001-08-2 | tivozanib | 378 | 0.980 | 0.980 | 432 | 0.998 | 1.000 | 535.5 | 0.996 | 1.000 | 658.5 | 0.049 | 0.514 | 262 | 0.168 | 0.708 | 370.5 | 0.974 | 1.000 | 490.5 | 0.787 | 1.000 | 279 | 0.229 | 0.457 | 281.5 | 0.239 | 0.457 | 213 | 0.314 | 0.550 | 241.5 | 0.482 | 0.778 | 4 | 0.059 | 0.154 | | | | | |
| BRD-K53417444-003-03-1 | OTS167 | 5038 | 0.085 | 0.403 | 8936 | 0.744 | 0.926 | 5035 | 0.434 | 0.702 | 19662 | 0.833 | 0.926 | 5662 | 0.096 | 0.403 | 5750 | 0.907 | 0.952 | 7952 | 0.838 | 0.926 | 6187 | 0.134 | 0.403 | 1923 | 0.088 | 0.403 | 5282 | 0.995 | 0.995 | 3548 | 0.291 | 0.669 | 2757 | 0.700 | 0.926 | 624 | 0.133 | 0.403 | 443 | 0.035 |
| BRD-K53523901-001-04-4 | arctigenin | 406.5 | 0.559 | 0.859 | 1786.5 | 0.928 | 0.958 | 1415 | 0.936 | 0.958 | 3729 | 0.619 | 0.859 | 1247 | 0.268 | 0.704 | 1035 | 0.695 | 0.859 | 1143 | 0.372 | 0.831 | 1751 | 0.958 | 0.958 | 488.5 | 0.258 | 0.704 | 712 | 0.607 | 0.859 | 294 | 0.201 | 1.000 | 400 | 0.537 | 0.859 | 178 | 0.122 | 0.425 | 84 | 0.400 |
| BRD-K53561341-001-07-5 | aurora-a-inhibitor-i | 1725 | 0.126 | 0.347 | 5102 | 0.996 | 0.996 | 2219 | 0.787 | 0.960 | 7559 | 0.083 | 0.289 | 3377 | 0.395 | 0.829 | 2448 | 0.718 | 0.960 | 4082 | 0.916 | 0.996 | 3982 | 0.793 | 0.960 | 686 | 0.051 | 0.213 | 2068 | 0.791 | 0.960 | 1425 | 0.047 | 0.497 | 808 | 0.132 | 0.347 | 440 | 0.823 | 0.960 | 56 | 0.001 |
| BRD-K53581288-001-02-9 | baricitinib | 843 | 0.978 | 0.992 | 754 | 0.977 | 0.992 | 563 | 0.114 | 1.000 | 2557.5 | 0.931 | 0.992 | 694 | 0.795 | 0.992 | 473 | 0.171 | 0.597 | 687 | 0.779 | 0.992 | 917 | 0.318 | 0.556 | 49 | 0.267 | 0.510 | 433 | 0.226 | 0.474 | 488.5 | 0.640 | 0.992 | 19 | 0.003 | 0.009 | | | | 113 | 0.067 |
| BRD-K53665955-001-03-0 | MK-5108 | 3797 | 0.835 | 0.990 | 5736 | 0.978 | 0.990 | 3047 | 0.818 | 0.990 | 9202 | 0.000 | 0.009 | 3207 | 0.095 | 0.662 | 3018 | 0.798 | 0.990 | 6267 | 0.990 | 0.990 | 4173 | 0.376 | 0.756 | 1837 | 0.739 | 0.990 | 2836 | 0.907 | 0.990 | 1692 | 0.046 | 0.485 | 1279 | 0.279 | 0.756 | 620 | 0.295 | 0.756 | 655 | 0.136 |
| BRD-K53734668-003-02-0 | PRT062607 | 3145 | 0.457 | 0.795 | 6335 | 0.966 | 0.980 | 2817 | 0.604 | 0.795 | 9796.5 | 0.085 | 0.356 | 4313 | 0.759 | 0.873 | 2983 | 0.606 | 0.795 | 4472 | 0.420 | 0.795 | 4440 | 0.980 | 0.980 | 674 | 0.022 | 0.151 | 2231 | 0.690 | 0.853 | 1742 | 0.183 | 1.000 | 668 | 0.182 | 0.638 | 273 | 0.075 | 0.356 | 781 | 0.341 |
| BRD-K54095730-001-03-1 | CMPD-1 | 3069 | 0.908 | 0.931 | 4264 | 0.836 | 0.931 | 1954 | 0.290 | 0.869 | 9239 | 0.720 | 0.931 | 2996 | 0.060 | 0.573 | 2962 | 0.929 | 0.931 | 2318 | 0.535 | 0.931 | 3072 | 0.435 | 0.916 | 1187 | 0.219 | 0.869 | 1017 | 0.190 | 0.869 | 1662 | 0.652 | 0.931 | 1385 | 0.480 | 0.916 | 583 | 0.741 | 0.931 | 435 | 0.407 |
| BRD-K54247840-004-01-3 | halofuginone | 5225 | 0.854 | 0.993 | 5552 | 0.011 | 0.120 | 3983 | 0.319 | 1.000 | 13974 | 0.414 | 1.000 | 5436 | 0.316 | 1.000 | 5149 | 0.993 | 0.993 | 6043 | 0.682 | 0.993 | 6177 | 0.543 | 0.950 | 1631 | 0.022 | 0.057 | 4200 | 0.973 | 0.993 | 2503 | 0.204 | 1.000 | 2832 | 0.916 | 0.993 | 885 | 0.877 | 0.993 | 1045 | 0.494 |
| BRD-K54256913-001-08-7 | MK-1775 | 2035 | 0.216 | 0.697 | 2796 | 0.595 | 0.892 | 2309 | 0.232 | 0.697 | 8943 | 0.879 | 0.923 | 2458.5 | 0.158 | 0.697 | 2044 | 0.837 | 0.923 | 2731.5 | 0.526 | 0.892 | 3020.5 | 0.419 | 0.880 | 1430 | 0.654 | 0.916 | 1244 | 0.100 | 0.697 | 1189 | 0.509 | 0.892 | 924 | 0.557 | 0.892 | 509.5 | 0.953 | 0.953 | 324 | 0.057 |
| BRD-K54262262-001-09-0 | monobenzone | 251 | 0.842 | 1.000 | 616.5 | 0.908 | 1.000 | 490 | 0.985 | 1.000 | 765 | 0.143 | 1.000 | 563.5 | 0.624 | 1.000 | 299.5 | 0.968 | 1.000 | 381.5 | 0.761 | 1.000 | 69 | 0.015 | 0.045 | 47 | 0.105 | 0.244 | 71 | 0.065 | 0.172 | 155.5 | 0.918 | 0.918 | 16 | 0.125 | 0.255 | | | | 4 | 0.059 |
| BRD-K54314721-001-10-2 | zolmitriptan | 247 | 0.543 | 0.781 | 678 | 0.695 | 0.781 | 252.5 | 0.283 | 0.742 | 1350 | 0.516 | 0.781 | 740 | 0.722 | 0.781 | 178.5 | 0.484 | 0.781 | 444 | 0.650 | 0.781 | 492 | 0.235 | 0.742 | 327.5 | 0.653 | 0.781 | 48 | 0.356 | 0.781 | 71 | 0.034 | 0.723 | 145.5 | 0.686 | 0.781 | 81 | 0.715 | 0.781 | 9 | 0.070 |
| BRD-K54395039-001-01-7 | PR-619 | 2668 | 0.571 | 0.887 | 4011 | 0.065 | 0.680 | 3405 | 0.676 | 0.887 | 9383 | 0.343 | 1.000 | 3161 | 0.455 | 1.000 | 2770 | 0.671 | 0.887 | 3676 | 0.646 | 0.887 | 3582 | 0.317 | 0.666 | 1452 | 0.255 | 0.595 | 2016 | 0.734 | 0.899 | 1582 | 0.025 | 0.530 | 570 | 0.034 | 0.090 | 682 | 0.517 | 0.887 | 617 | 0.390 |
| BRD-K54606188-001-13-1 | JQ1-(+) | 1136 | 0.370 | 0.657 | 1012 | 0.295 | 1.000 | 571 | 0.760 | 0.869 | 3653 | 0.786 | 0.869 | 1556 | 0.392 | 0.657 | 640 | 0.425 | 0.657 | 1955 | 0.989 | 0.989 | 972 | 0.767 | 0.869 | 208 | 0.015 | 0.052 | 355 | 0.157 | 0.471 | 758 | 0.071 | 0.749 | 523 | 0.627 | 0.823 | 183 | 0.438 | 0.657 | 125 | 0.190 |
| BRD-K54838207-001-03-8 | fudosteine | 349 | 0.885 | 0.990 | 547 | 0.490 | 1.000 | 286 | 0.620 | 1.000 | 927.5 | 0.109 | 0.596 | 399 | 0.007 | 0.137 | 291 | 0.190 | 0.796 | 529 | 0.978 | 0.990 | 476 | 0.385 | 1.000 | 177.5 | 0.937 | 0.990 | 38 | 0.303 | 1.000 | 150 | 0.821 | 1.000 | 242.5 | 0.930 | 0.990 | 43 | 0.360 | 0.581 | 176.5 | 0.934 |
| BRD-K54955827-001-02-2 | niraparib | 483 | 0.361 | 0.690 | 701 | 0.801 | 0.986 | 511 | 0.249 | 1.000 | 2203 | 0.614 | 0.986 | 863 | 0.113 | 0.401 | 498 | 0.662 | 0.986 | 1108 | 0.943 | 0.986 | 1076 | 0.816 | 0.986 | 442 | 0.240 | 0.561 | 279 | 0.975 | 0.986 | 477 | 0.342 | 0.690 | 214 | 0.004 | 0.010 | | | | 198 | 0.114 |
| BRD-K54997624-001-06-0 | alpelisib | 357.5 | 0.237 | 0.553 | 576 | 0.753 | 0.972 | 495 | 0.134 | 1.000 | 2003.5 | 0.884 | 0.972 | 822.5 | 0.486 | 0.863 | 645.5 | 0.537 | 0.867 | 964.5 | 0.692 | 0.969 | 716.5 | 0.997 | 0.997 | 274 | 0.064 | 0.224 | 290 | 0.493 | 0.863 | 383.5 | 0.148 | 1.000 | 49 | 0.052 | 0.218 | | | | 325 | 0.922 |
| BRD-K55187425-236-05-2 | rigosertib | 1625.5 | 0.598 | 0.897 | 3568 | 0.990 | 0.996 | 1562 | 0.197 | 0.522 | 5794.5 | 0.314 | 0.660 | 1184.5 | 0.036 | 0.519 | 1874.5 | 0.958 | 0.996 | 3000 | 0.996 | 0.996 | 2846 | 0.729 | 0.931 | 1063 | 0.302 | 0.660 | 1342 | 0.754 | 0.931 | 1652.5 | 0.456 | 0.805 | 853.5 | 0.178 | 0.522 | 269 | 0.498 | 0.805 | 471 | 0.130 |
| BRD-K55305701-001-01-2 | SC-144 | 1159 | 0.986 | 0.986 | 1411 | 0.886 | 0.986 | 988 | 0.182 | 0.546 | 3568 | 0.297 | 0.779 | 1288.5 | 0.005 | 0.027 | 712 | 0.943 | 0.986 | 1486 | 0.861 | 0.986 | 1181.5 | 0.352 | 0.821 | 286.5 | 0.049 | 0.204 | 539 | 0.060 | 0.210 | 289 | 0.161 | 1.000 | 18 | 0.002 | 0.017 | 301.5 | 0.877 | 0.986 | 252 | 0.715 |
| BRD-K55696337-003-24-4 | topotecan | 6474 | 0.893 | 0.997 | 10022 | 0.994 | 0.997 | 3934 | 0.071 | 0.249 | 16301 | 0.102 | 0.278 | 4575 | 0.008 | 0.097 | 5495 | 0.929 | 0.997 | 9055 | 0.997 | 0.997 | 7382 | 0.762 | 0.997 | 1446 | 0.009 | 0.097 | 4069 | 0.708 | 0.997 | 3492 | 0.301 | 0.632 | 1821 | 0.067 | 0.249 | 1016 | 0.645 | 0.997 | 549 | 0.022 |
| BRD-K56032964-001-02-1 | AP26113 | 1247 | 0.684 | 0.980 | 1838.5 | 0.980 | 0.980 | 1158 | 0.100 | 0.525 | 4239 | 0.060 | 0.447 | 1099.5 | 0.064 | 0.447 | 1251.5 | 0.956 | 0.980 | 2050.5 | 0.216 | 0.651 | 2114 | 0.892 | 0.980 | 175 | 0.794 | 0.980 | 1882.5 | 0.890 | 0.980 | 1176.5 | 0.214 | 0.651 | 472 | 0.231 | 0.651 | 435.5 | 0.980 | 0.980 | 272.5 | 0.248 |
| BRD-K56277358-001-07-5 | MGCD-265 | 2041 | 0.955 | 1.000 | 3795 | 0.997 | 1.000 | 1415 | 0.148 | 0.578 | 6157 | 0.438 | 0.707 | 2596 | 0.542 | 0.813 | 1547 | 0.636 | 0.890 | 3033 | 0.748 | 0.981 | 2405 | 0.085 | 0.578 | 493 | 0.149 | 0.578 | 1297 | 0.269 | 0.578 | 1654 | 0.973 | 0.973 | 391 | 0.053 | 0.578 | 366 | 0.377 | 0.660 | 111 | 0.074 |
| BRD-K56291712-001-01-0 | voxtalisib | 902 | 0.989 | 1.000 | 1125 | 0.722 | 0.967 | 276 | 0.044 | 0.341 | 2133.5 | 0.316 | 0.830 | 628.5 | 0.185 | 0.646 | 422 | 0.626 | 0.967 | 1422 | 0.934 | 1.000 | 711.5 | 0.221 | 0.662 | 72 | 0.004 | 0.080 | 282.5 | 0.049 | 0.341 | 374 | 0.083 | 0.347 | 61 | 0.065 | 0.341 | 93 | 0.616 | 0.967 | 71 | 0.426 |
| BRD-K56301217-001-07-4 | ABT-737 | 315.5 | 0.883 | 1.000 | 778 | 0.139 | 0.488 | 412 | 0.340 | 0.793 | 1236 | 0.766 | 1.000 | 690.5 | 0.781 | 1.000 | 312 | 0.615 | 1.000 | 537.5 | 0.939 | 1.000 | 275 | 0.203 | 0.533 | 151 | 0.746 | 1.000 | 248 | 0.124 | 0.488 | 242 | 0.110 | 0.488 | 5 | 0.059 | 0.488 | | | | 53 | 0.091 |
| BRD-K56334280-001-05-1 | amonafide | 313 | 0.635 | 0.890 | 589.5 | 0.820 | 0.912 | 282 | 0.234 | 1.000 | 1456 | 0.487 | 1.000 | 481.5 | 0.139 | 1.000 | 388.5 | 0.912 | 0.912 | 445 | 0.715 | 0.912 | 421 | 0.834 | 0.912 | 135.5 | 0.077 | 0.163 | 236.5 | 0.530 | 0.855 | 83 | 0.058 | 1.000 | 118.5 | 0.168 | 0.320 | 54 | 0.442 | 0.773 | 5 | 0.057 |
| BRD-K56405753-001-02-4 | MK-2461 | 557 | 0.890 | 0.890 | 799 | 0.864 | 0.955 | 352 | 0.175 | 0.682 | 1600 | 0.701 | 0.955 | 652 | 0.156 | 0.682 | 250 | 0.811 | 0.955 | 623 | 0.730 | 0.955 | 536 | 0.242 | 0.685 | 187 | 0.165 | 0.682 | 250 | 0.195 | 0.682 | 406 | 0.858 | 0.955 | 171 | 0.116 | 0.682 | 71 | 0.560 | 0.938 | 127 | 0.473 |
| BRD-K56707426-001-02-0 | nitarsone | 160 | 0.358 | 0.893 | 798 | 0.716 | 0.950 | 212 | 0.135 | 0.709 | 960 | 0.024 | 0.165 | 411 | 0.284 | 0.893 | 405 | 0.468 | 0.893 | 611 | 0.926 | 0.950 | 501 | 0.422 | 0.893 | 241 | 0.827 | 0.950 | 320 | 0.337 | 0.893 | 194 | 0.092 | 1.000 | 257 | 0.885 | 0.950 | 27 | 0.169 | 0.712 | 107 | 0.896 |
| BRD-K56957086-001-06-3 | dacinostat | 5887 | 0.770 | 0.951 | 8429 | 0.717 | 0.942 | 5119 | 0.703 | 0.942 | 16138 | 0.168 | 0.294 | 5194 | 0.112 | 0.261 | 5569 | 0.951 | 1.000 | 9900 | 1.000 | 1.000 | 9065 | 0.997 | 1.000 | 2182 | 0.138 | 0.263 | 4647 | 0.996 | 1.000 | 2717 | 0.027 | 0.196 | 2720 | 0.539 | 0.809 | 567 | 0.102 | 0.261 | 496 | 0.014 |
| BRD-K56981171-001-01-0 | brigatinib | 1190.5 | 0.984 | 0.984 | 1124.5 | 0.709 | 0.935 | 1413 | 0.237 | 0.830 | 3620 | 0.105 | 0.554 | 1405 | 0.510 | 0.901 | 1126.5 | 0.712 | 0.935 | 1400 | 0.040 | 0.426 | 1414.5 | 0.981 | 0.984 | 221 | 0.543 | 0.901 | 742 | 0.544 | 0.901 | 1028 | 0.970 | 0.984 | 420 | 0.228 | 0.830 | 115 | 0.558 | 0.901 | 464 | 0.336 |

| broad_id | name | al_nervous_system_PVAL_ADJ | fibroblast_MWU_STAT | fibroblast_PVAL | fibroblast_PVAL_ADJ | bile_duct_MWU_STAT | bile_duct_PVAL | bile_duct_PVAL_ADJ | thyroid_MWU_STAT | thyroid_PVAL | thyroid_PVAL_ADJ | upper_aerodigestive_MWU_STAT | upper_aerodigestive_PVAL | upper_aerodigestive_PVAL_ADJ | uterus_MWU_STAT | uterus_PVAL | uterus_PVAL_ADJ | liver_MWU_STAT | liver_PVAL | liver_PVAL_ADJ | esophagus_MWU_STAT | esophagus_PVAL | esophagus_PVAL_ADJ |
|---|---|---|---|---|---|---|---|---|---|---|---|---|---|---|---|---|---|---|---|---|---|---|---|
| BRD-K43644456-001-04-0 | WP1130 | 0.584 | 196 | 0.432 | 0.756 | 2318 | 0.986 | 0.998 | 1746 | 0.249 | 0.584 | 4220 | 0.179 | 0.584 | 3208 | 0.011 | 0.074 | 3386 | 0.259 | 0.584 | 5677 | 0.871 | 0.998 |
| BRD-K43905307-001-01-2 | L-798,106 | 0.668 | | | | 414 | 0.810 | 0.946 | 553 | 0.910 | 0.955 | 710.5 | 0.475 | 0.820 | 309 | 0.174 | 1.000 | 411.5 | 0.507 | 0.820 | 811 | 0.741 | 0.915 |
| BRD-K44084986-300-08-4 | Y-27632 | 0.826 | | | | 91 | 0.065 | 0.340 | 1119 | 0.885 | 1.000 | 2026 | 0.984 | 1.000 | 1580 | 0.472 | 0.826 | 412 | 0.027 | 0.287 | 2158 | 0.878 | 0.878 |
| BRD-K44227013-001-06-4 | ponatinib | 1.000 | | | | 588 | 0.590 | 1.000 | 723 | 0.308 | 0.946 | 1613 | 0.200 | 0.840 | 1291 | 0.124 | 0.840 | 1542 | 0.406 | 0.946 | 1745 | 0.942 | 1.000 |
| BRD-K44408410-001-17-6 | 2-methoxyestradiol | 0.654 | | | | | | | 142 | 0.034 | 0.719 | 547 | 0.185 | 1.000 | 621 | 0.368 | 1.000 | 422.5 | 0.193 | 0.405 | 664 | 0.306 | 1.000 |
| BRD-K44590731-001-01-7 | TU-2100 | 0.864 | | | | 197 | 0.961 | 0.961 | 64 | 0.577 | 0.942 | 227 | 0.509 | 0.942 | 147 | 0.756 | 0.942 | 228 | 0.843 | 0.942 | 415 | 0.411 | 0.864 |
| BRD-K44665581-001-01-8 | bromosporine | 0.154 | | | | | | | 112 | 0.131 | 1.000 | 296 | 0.251 | 1.000 | 427.5 | 0.357 | 0.682 | 108 | 0.906 | 0.975 | 444.5 | 0.423 | 0.712 |
| BRD-K44844162-001-01-6 | taselisib | 0.637 | 201.5 | 0.914 | 1.000 | 270 | 0.690 | 1.000 | 719 | 0.983 | 1.000 | 1197.5 | 0.012 | 0.057 | 990.5 | 0.521 | 0.994 | 1379 | 0.928 | 1.000 | 1785.5 | 0.502 | 0.994 |
| BRD-K45293975-001-02-0 | 7-hydroxystaurosporine | 0.851 | 29 | 0.076 | 0.534 | 907 | 0.398 | 0.847 | 1432 | 0.361 | 0.847 | 3886 | 0.608 | 0.851 | 4720 | 0.991 | 0.991 | 2425 | 0.444 | 0.847 | 4420 | 0.834 | 0.925 |
| BRD-K45401373-001-20-9 | betulinic-acid | | | | | 136 | 0.891 | 0.952 | 137 | 0.376 | 1.000 | 725 | 0.821 | 0.952 | 299 | 0.194 | 1.000 | 693 | 0.906 | 0.952 | 761 | 0.728 | 0.952 |
| BRD-K45528773-001-07-2 | M-344 | 0.064 | 23 | 0.054 | 0.190 | 2042 | 0.937 | 1.000 | 1617 | 0.189 | 0.410 | 4553 | 0.447 | 0.723 | 4221 | 0.357 | 0.656 | 2575 | 0.047 | 0.190 | 4316 | 0.195 | 0.410 |
| BRD-K45551181-001-02-1 | MPI-0479605 | 0.498 | 364 | 0.931 | 0.996 | 1484 | 0.879 | 0.996 | 1998 | 0.789 | 0.996 | 2340 | 0.006 | 0.127 | 3400 | 0.376 | 0.790 | 2048 | 0.148 | 0.498 | 3552 | 0.154 | 0.498 |
| BRD-K45724504-001-01-6 | anguidine | 0.784 | | | | 1673 | 0.884 | 0.986 | 2971 | 0.981 | 0.986 | 3234 | 0.146 | 0.615 | 3470 | 0.097 | 0.615 | 2992 | 0.515 | 0.784 | 4258 | 0.116 | 0.615 |
| BRD-K46392623-001-02-5 | TG-100713 | 0.639 | | | | 158 | 0.039 | 0.405 | 908.5 | 0.941 | 1.000 | 1476 | 0.002 | 0.037 | 682.5 | 0.142 | 0.595 | 1169 | 0.916 | 1.000 | 2082 | 0.641 | 1.000 |
| BRD-K46690591-001-01-2 | KP-1212 | 0.245 | | | | 352 | 0.984 | 0.984 | 595 | 0.973 | 0.984 | 960 | 0.772 | 0.984 | 897 | 0.437 | 1.000 | 414 | 0.610 | 0.916 | 1407 | 0.755 | 0.984 |
| BRD-K47000838-001-01-6 | cytochalasin-b | 0.879 | 70 | 0.193 | 0.692 | 1437 | 0.989 | 1.000 | 948 | 0.050 | 0.332 | 2871 | 0.263 | 0.692 | 2489 | 0.512 | 0.879 | 1795 | 0.507 | 0.879 | 3899 | 0.706 | 0.879 |
| BRD-K47079459-004-04-6 | cetrimonium | 0.997 | | | | 847 | 0.671 | 0.997 | 813 | 0.045 | 0.318 | 2677 | 0.107 | 0.561 | 3544 | 0.750 | 0.997 | 2291 | 0.974 | 0.997 | 3919 | 0.762 | 0.997 |
| BRD-K47114202-001-14-6 | chlorogenic-acid | 0.109 | | | | 149 | 0.466 | 1.000 | 193 | 0.105 | 0.834 | 749 | 0.356 | 1.000 | 607 | 0.759 | 0.881 | 646 | 0.849 | 0.892 | 850 | 0.797 | 0.881 |
| BRD-K47150025-001-05-3 | KI-8751 | 0.823 | | | | 137 | 0.681 | 0.902 | 614.5 | 0.730 | 0.902 | 776 | 0.600 | 0.902 | 588 | 0.185 | 1.000 | 546.5 | 0.835 | 0.902 | 1345.5 | 0.371 | 0.823 |
| BRD-K47337578-034-02-5 | 4-(4-fluorobenzoyl)-1-(4-phenylbutyl)-piperidine | 0.399 | | | | | | | 117 | 0.054 | 0.570 | 315 | 0.258 | 1.000 | 440 | 0.819 | 0.967 | 43 | 0.064 | 0.135 | 420 | 0.916 | 0.967 |
| BRD-K47598052-001-15-8 | PP-1 | 0.558 | | | | 142 | 0.747 | 1.000 | 170 | 0.204 | 1.000 | 313 | 0.093 | 0.648 | 104 | 0.424 | 1.000 | | | | 674 | 0.941 | 0.941 |
| BRD-K47832606-001-30-1 | floxuridine | 0.489 | 107 | 0.198 | 0.496 | 2063 | 0.971 | 0.987 | 2023 | 0.457 | 0.738 | 2999 | 0.102 | 0.496 | 4302 | 0.958 | 0.987 | 2417 | 0.161 | 0.496 | 4861 | 0.790 | 0.987 |
| BRD-K47869605-001-32-0 | podophyllotoxin | 0.527 | 14 | 0.033 | 0.350 | 1968 | 0.979 | 0.979 | 1781 | 0.300 | 0.667 | 3601 | 0.110 | 0.527 | 3836 | 0.125 | 0.527 | 3373 | 0.589 | 0.951 | 4473 | 0.347 | 0.667 |
| BRD-K47976015-001-01-2 | balapiravir | 0.109 | | | | 285 | 0.765 | 0.905 | 262 | 0.665 | 1.000 | 497 | 0.106 | 0.743 | 558.5 | 0.626 | 1.000 | 136 | 0.905 | 0.905 | 517 | 0.486 | 1.000 |
| BRD-K48443249-001-01-0 | CNX-774 | 0.573 | | | | 3 | 0.051 | 0.358 | 198.5 | 0.453 | 0.962 | 182.5 | 0.016 | 0.168 | 213 | 0.866 | 0.980 | 262.5 | 0.980 | 0.980 | 376 | 0.966 | 0.980 |
| BRD-K48488978-001-05-3 | YM-201636 | 0.702 | | | | 3 | 0.050 | 0.869 | 335.5 | 0.932 | 0.979 | 839.5 | 0.833 | 0.979 | 714.5 | 0.712 | 0.979 | 289 | 0.210 | 0.551 | 535 | 0.369 | 0.702 |
| BRD-K48684885-001-04-2 | BAY-11-7085 | 0.533 | 79 | 0.320 | 0.753 | 445 | 0.868 | 0.959 | 787 | 0.834 | 0.959 | 963 | 0.096 | 0.533 | 862 | 0.329 | 0.753 | 931 | 0.998 | 0.998 | 1228 | 0.102 | 0.533 |
| BRD-K48988407-001-01-2 | cytochlor | 0.446 | | | | 203.5 | 0.950 | 0.980 | 459.5 | 0.655 | 0.980 | 1128 | 0.035 | 0.367 | 657.5 | 0.980 | 0.980 | 348 | 0.642 | 0.980 | 1630.5 | 0.632 | 0.980 |
| BRD-K49027941-001-01-1 | PSB-1115 | 1.000 | | | | 126 | 0.637 | 1.000 | 126 | 0.637 | 1.000 | 187.5 | 0.662 | 1.000 | 187.5 | 0.662 | 1.000 | 248 | 0.684 | 1.000 | 423.5 | 0.736 | 1.000 |
| BRD-K49049886-001-08-7 | CGS-15943 | 1.000 | | | | 171 | 0.243 | 0.596 | 126 | 0.534 | 1.000 | 2022.5 | 0.255 | 0.596 | 1245.5 | 0.654 | 1.000 | 963.5 | 0.016 | 0.085 | 1648 | 0.014 | 0.297 |
| BRD-K49075727-001-08-5 | nintedanib | 0.925 | | | | 79 | 0.722 | 0.925 | 79 | 0.124 | 1.000 | 310 | 0.879 | 0.925 | 287 | 0.388 | 0.741 | 459 | 0.925 | 0.925 | 479 | 0.263 | 1.000 |
| BRD-K49294207-300-03-8 | BIBU-1361 | 0.970 | | | | 377 | 0.741 | 0.970 | 571 | 0.909 | 0.970 | 339 | 0.000 | 0.000 | 555 | 0.885 | 0.970 | 834 | 0.785 | 0.970 | 1517 | 0.303 | 0.795 |
| BRD-K49313711-001-05-4 | ribitol | 0.069 | | | | 89 | 0.871 | 0.950 | 65 | 0.643 | 0.950 | 398 | 0.155 | 1.000 | 298.5 | 0.447 | 0.939 | 66 | 0.180 | 0.540 | 372.5 | 0.359 | 0.926 |
| BRD-K49328571-001-15-0 | dasatinib | 1.000 | 107 | 0.444 | 0.777 | 300 | 0.341 | 0.777 | 332 | 0.166 | 0.582 | 1363.5 | 0.003 | 0.022 | 775.5 | 0.112 | 0.471 | 661 | 0.438 | 0.777 | 1250 | 0.350 | 0.777 |
| BRD-K49350383-001-13-7 | thioguanine | 0.870 | 352 | 0.946 | 0.999 | 1113 | 0.519 | 0.838 | 1085 | 0.477 | 0.835 | 3511 | 0.761 | 0.999 | 3807 | 0.964 | 0.999 | 1573 | 0.009 | 0.092 | 3121 | 0.299 | 0.817 |
| BRD-K49350383-001-14-5 | thioguanine | 0.554 | 113.5 | 0.830 | 0.977 | 122.5 | 0.326 | 0.924 | 127.5 | 0.358 | 0.924 | 594.5 | 0.581 | 0.977 | 337.5 | 0.950 | 0.977 | 593.5 | 0.947 | 0.977 | 376 | 0.115 | 0.554 |
| BRD-K49371609-003-03-8 | PIK-75 | 0.486 | 349 | 0.820 | 0.906 | 1007 | 0.078 | 0.486 | 2092 | 0.718 | 0.906 | 2893 | 0.015 | 0.201 | 3831 | 0.329 | 0.732 | 2959 | 0.560 | 0.905 | 3827 | 0.326 | 0.732 |
| BRD-K49456274-001-01-9 | pilaralisib | | 28 | 0.139 | 0.731 | 111 | 0.065 | 0.456 | 274 | 0.727 | 0.927 | 698 | 0.423 | 0.920 | 379 | 0.046 | 0.456 | 491 | 0.881 | 1.000 | 776 | 0.820 | 0.820 |
| BRD-K49522529-001-02-9 | alfacalcidol | 0.799 | | | | 196 | 0.502 | 0.850 | 252 | 0.759 | 0.900 | 583 | 0.003 | 0.059 | 666 | 0.495 | 0.850 | 457 | 0.422 | 0.850 | 777 | 0.178 | 1.000 |
| BRD-K49669041-001-08-4 | BX-912 | 0.854 | 83 | 0.230 | 0.805 | 1149 | 0.609 | 0.898 | 1452 | 0.164 | 0.687 | 2778 | 0.336 | 0.854 | 3078 | 0.756 | 0.934 | 1894 | 0.642 | 0.898 | 3055 | 0.591 | 0.898 |
| BRD-K49865102-001-08-4 | PD-0325901 | 0.438 | | | | 1066 | 0.742 | 1.000 | 1313 | 0.327 | 0.854 | 2403 | 0.009 | 0.064 | 2167 | 0.710 | 1.000 | 2357 | 0.366 | 0.854 | 4629 | 0.979 | 1.000 |
| BRD-K49954789-001-02-9 | mesna | 0.996 | | | | 306 | 0.742 | 0.996 | 423.5 | 0.533 | 0.996 | 671 | 0.092 | 0.905 | 264 | 0.559 | 0.996 | 254.5 | 0.895 | 0.996 | 1475 | 0.996 | 0.996 |
| BRD-K49962337-001-02-9 | baicalin | | | | | 117 | 0.931 | 0.948 | 240 | 0.807 | 0.948 | 576 | 0.665 | 0.948 | 364 | 0.507 | 0.887 | 114 | 0.419 | 0.887 | 431 | 0.942 | 0.948 |
| BRD-K50000283-001-04-4 | PHA-767491 | 0.878 | 143 | 0.486 | 0.878 | 771 | 0.603 | 0.878 | 1003 | 0.751 | 0.893 | 2279 | 0.935 | 0.961 | 1312 | 0.342 | 0.878 | 1756 | 0.765 | 0.893 | 2061 | 0.628 | 0.878 |
| BRD-K50010139-001-01-5 | poziotinib | | | | | 118 | 0.529 | 0.926 | 106 | 0.925 | 1.000 | 267 | 0.034 | 0.238 | 173 | 0.075 | 0.394 | 348 | 0.823 | 1.000 | 404 | 0.002 | 0.018 |
| BRD-K50010139-001-02-3 | poziotinib | 0.929 | 244 | 0.830 | 1.000 | 532 | 0.307 | 0.645 | 1780 | 0.998 | 1.000 | 1327 | 0.000 | 0.000 | 2177 | 0.266 | 0.621 | 1727 | 0.227 | 0.595 | 1260 | 0.000 | 0.000 |
| BRD-K50140147-001-10-1 | NVP-TAE684 | 0.016 | | | | 925 | 0.790 | 0.986 | 1211 | 0.735 | 0.986 | 2183 | 0.110 | 0.848 | 1694 | 0.121 | 0.848 | 1338 | 0.276 | 0.691 | 2153 | 0.422 | 0.806 |
| BRD-K50168500-001-07-9 | canertinib | | | | | 163 | 0.938 | 1.000 | 163 | 0.938 | 1.000 | 566 | 0.000 | 0.008 | 849 | 0.576 | 1.000 | 64 | 0.064 | 0.135 | 1253 | 0.051 | 0.293 |
| BRD-K50677762-001-01-5 | oxiracetam | 0.524 | 122 | 0.312 | 0.728 | 1224 | 0.958 | 0.999 | 1342 | 0.504 | 0.764 | 2838 | 0.416 | 0.764 | 1877 | 0.028 | 0.297 | 1226 | 0.577 | 0.808 | 2556 | 0.439 | 0.764 |
| BRD-K50853363-001-02-3 | dienogest | 0.988 | | | | 253.5 | 0.608 | 1.000 | 406.5 | 0.864 | 0.988 | 204 | 0.126 | 0.880 | 386 | 0.808 | 0.988 | 266 | 0.124 | 0.290 | 970.5 | 0.988 | 0.988 |
| BRD-K51040301-001-02-1 | cobicistat | 0.378 | | | | 106 | 0.896 | 0.971 | 231 | 0.792 | 0.971 | 251 | 0.044 | 0.915 | 222 | 0.748 | 0.971 | 338 | 0.699 | 0.971 | 237 | 0.223 | 1.000 |
| BRD-K51313569-001-07-8 | palbociclib | 0.832 | | | | 49 | 0.298 | 1.000 | 229.5 | 0.602 | 0.820 | 528 | 0.676 | 0.835 | 347.5 | 0.502 | 0.753 | 303.5 | 0.317 | 0.596 | 499 | 0.348 | 1.000 |
| BRD-K51313569-003-03-3 | palbociclib | 0.920 | 124.5 | 0.876 | 0.920 | 124.5 | 0.876 | 0.920 | 259.5 | 0.336 | 0.831 | 285.5 | 0.071 | 0.368 | 267 | 0.725 | 0.920 | 826 | 0.462 | 0.920 | 518 | 0.539 | 0.920 |
| BRD-K51318897-001-15-3 | fenbendazole | 0.850 | 329 | 0.866 | 0.957 | 1042 | 0.658 | 0.921 | 1400 | 0.385 | 0.750 | 3328 | 0.407 | 0.750 | 2974 | 0.261 | 0.750 | 2329 | 0.277 | 0.750 | 2926 | 0.137 | 0.718 |
| BRD-K51324732-001-01-0 | meglitinide | | | | | 94 | 0.886 | 1.000 | 179 | 0.614 | 1.000 | 164 | 0.210 | 0.758 | 194 | 0.152 | 0.758 | 37 | 0.289 | 0.758 | 440 | 0.810 | 1.000 |
| BRD-K51350053-048-16-5 | toremifene | 0.602 | 20 | 0.073 | 0.602 | 106 | 0.372 | 0.709 | 825 | 0.238 | 0.602 | 1413 | 0.536 | 0.819 | 1864 | 0.024 | 0.513 | 776 | 0.175 | 0.602 | 2715 | 0.983 | 0.983 |
| BRD-K51544265-001-04-2 | cabozantinib | 1.000 | | | | 72 | 0.086 | 0.658 | 355 | 0.314 | 0.755 | 822.5 | 0.914 | 1.000 | 665 | 0.575 | 1.000 | 288 | 0.133 | 0.658 | 1015 | 0.935 | 0.935 |
| BRD-K51575138-001-13-2 | TPCA-1 | 0.438 | | | | | | | 232 | 0.616 | 0.994 | 453 | 0.003 | 0.058 | 329 | 0.422 | 1.000 | | | | 825.5 | 0.597 | 0.597 |
| BRD-K51747290-001-13-1 | hydroxyurea | 0.510 | | | | | | | 268 | 0.325 | 1.000 | 260 | 0.046 | 0.676 | 128 | 0.527 | 0.869 | 207 | 0.945 | 0.992 | 394 | 0.640 | 0.896 |
| BRD-K51791723-003-01-7 | P276-00 | 0.629 | 353 | 0.929 | 0.976 | 1197 | 0.849 | 0.976 | 1474 | 0.713 | 0.976 | 3145 | 0.166 | 0.582 | 2738 | 0.123 | 0.582 | 2114 | 0.587 | 0.976 | 3611 | 0.366 | 0.768 |
| BRD-K51911221-001-01-2 | nemorubicin | 0.060 | 44 | 0.097 | 0.511 | 1863 | 0.935 | 0.954 | 2101 | 0.352 | 0.822 | 5377 | 0.810 | 0.954 | 4366 | 0.329 | 0.822 | 3781 | 0.658 | 0.954 | 5355 | 0.689 | 0.954 |
| BRD-K51967704-001-03-6 | BIIB021 | 0.976 | 227 | 0.743 | 0.976 | 630 | 0.451 | 0.947 | 1060 | 0.644 | 0.976 | 2551 | 0.380 | 0.886 | 1935 | 0.209 | 0.549 | 1081 | 0.039 | 0.207 | 1826 | 0.014 | 0.149 |
| BRD-K52075715-001-06-7 | oxibendazole | 0.181 | 210 | 0.588 | 0.980 | 1126 | 0.980 | 0.980 | 1540 | 0.678 | 0.980 | 2641 | 0.160 | 0.610 | 2591 | 0.824 | 0.980 | 1927 | 0.516 | 0.980 | 3680 | 0.847 | 0.980 |
| BRD-K52233191-001-02-4 | R547 | 0.375 | 192 | 0.587 | 0.956 | 810 | 0.795 | 0.995 | 1333 | 0.995 | 0.995 | 1948 | 0.377 | 0.791 | 2424 | 0.577 | 0.956 | 2205 | 0.826 | 0.995 | 2202 | 0.033 | 0.342 |
| BRD-K52256627-300-06-6 | chlorhexidine | 0.314 | | | | 241 | 0.158 | 0.552 | 1043 | 0.963 | 1.000 | 2072 | 0.945 | 1.000 | 1762 | 0.663 | 0.994 | 538 | 0.049 | 0.346 | 2328 | 0.792 | 0.792 |
| BRD-K52762805-001-01-2 | epothilone-d | 0.862 | 132 | 0.420 | 0.862 | 1019 | 0.648 | 0.862 | 961 | 0.550 | 0.862 | 1973 | 0.200 | 0.840 | 2077 | 0.177 | 0.840 | 1513 | 0.285 | 0.862 | 2299 | 0.381 | 0.862 |
| BRD-K52911425-001-09-8 | GDC-0941 | 1.000 | | | | 878 | 0.875 | 1.000 | 770 | 0.722 | 1.000 | 1964 | 0.014 | 0.100 | 3208 | 0.963 | 1.000 | 2332 | 0.771 | 1.000 | 3963 | 0.946 | 0.946 |
| BRD-K53414658-001-08-2 | tivozanib | | | | | 106 | 0.552 | 0.828 | 49 | 0.024 | 0.510 | 120 | 0.680 | 0.951 | 182 | 0.169 | 0.708 | 113 | 0.237 | 0.457 | 133 | 0.135 | 0.708 |
| BRD-K53417444-003-03-1 | OTS167 | 0.403 | 60 | 0.130 | 0.403 | 1498 | 0.369 | 0.702 | 2099 | 0.319 | 0.669 | 4633 | 0.318 | 0.669 | 4365 | 0.410 | 0.702 | 3691 | 0.544 | 0.817 | 5808 | 0.759 | 0.926 |
| BRD-K53523901-001-04-4 | arctigenin | 0.831 | | | | 379.5 | 0.808 | 0.942 | 371.5 | 0.187 | 1.000 | 643.5 | 0.680 | 0.859 | 551 | 0.036 | 0.756 | 147 | 0.001 | 0.004 | 1090 | 0.435 | 0.831 |
| BRD-K53561341-001-07-5 | aurora-a-inhibitor-i | 0.006 | | | | 51 | 0.165 | 0.385 | 974 | 0.976 | 0.996 | 2711 | 0.816 | 0.960 | 688 | 0.016 | 0.337 | 1776 | 0.447 | 0.853 | 2874 | 0.565 | 0.960 |
| BRD-K53581288-001-02-9 | baricitinib | 0.282 | | | | 394.5 | 0.984 | 0.992 | 534.5 | 0.783 | 0.992 | 301 | 0.000 | 0.002 | 617 | 0.992 | 0.992 | 4 | 0.009 | 0.022 | 548 | 0.363 | 1.000 |
| BRD-K53665955-001-03-0 | MK-5108 | 0.716 | 118 | 0.266 | 0.756 | 1062 | 0.937 | 0.990 | 1849 | 0.905 | 0.990 | 3950 | 0.904 | 0.990 | 3173 | 0.347 | 0.756 | 2063 | 0.396 | 0.756 | 3382 | 0.263 | 0.756 |
| BRD-K53734668-003-02-0 | PRT062607 | 0.795 | | | | 926 | 0.589 | 0.795 | 1230 | 0.519 | 0.795 | 2720 | 0.356 | 0.795 | 2863 | 0.790 | 0.873 | 1671 | 0.434 | 0.795 | 3171.5 | 0.345 | 0.795 |
| BRD-K54095730-001-03-1 | CMPD-1 | 0.916 | 84 | 0.266 | 0.869 | 907 | 0.931 | 0.931 | 626 | 0.082 | 0.573 | 1801 | 0.008 | 0.171 | 2211 | 0.581 | 0.931 | 2194 | 0.865 | 0.931 | 2667 | 0.447 | 0.916 |
| BRD-K54247840-004-01-3 | halofuginone | 0.942 | | | | 1636 | 0.701 | 0.993 | 2230 | 0.837 | 0.993 | 2825 | 0.002 | 0.044 | 4843 | 0.990 | 0.993 | 2850 | 0.191 | 0.445 | 4005 | 0.467 | 0.942 |
| BRD-K54256913-001-08-7 | MK-1775 | 0.602 | 105 | 0.300 | 0.788 | 405 | 0.381 | 0.880 | 1439.5 | 0.852 | 0.923 | 1921 | 0.861 | 0.923 | 2175 | 0.227 | 0.697 | 1000.5 | 0.043 | 0.602 | 2974 | 0.866 | 0.923 |
| BRD-K54262262-001-09-0 | monobenzone | 0.089 | | | | | | | 110 | 0.610 | 1.000 | 101 | 0.181 | 1.000 | 208.5 | 0.893 | 1.000 | 91 | 0.133 | 0.255 | 325 | 0.146 | 1.000 |
| BRD-K54314721-001-10-2 | zolmitriptan | 0.293 | | | | | | | 274 | 0.690 | 0.781 | 342.5 | 0.232 | 1.000 | 468.5 | 0.744 | 0.781 | 337.5 | 0.921 | 0.921 | 451.5 | 0.261 | 0.742 |
| BRD-K54395039-001-01-7 | PR-619 | 0.745 | | | | 1006 | 0.771 | 0.899 | 1314 | 0.475 | 1.000 | 2333 | 0.549 | 1.000 | 3146 | 0.970 | 0.970 | 2011 | 0.962 | 0.970 | 3759 | 0.901 | 0.970 |
| BRD-K54606188-001-13-1 | JQ1-(+) | 0.500 | | | | 7 | 0.055 | 0.749 | 267 | 0.400 | 0.657 | 1163 | 0.422 | 0.657 | 759 | 0.293 | 1.000 | 790 | 0.940 | 0.987 | 1155 | 0.582 | 0.814 |
| BRD-K54838207-001-03-8 | fudosteine | 0.990 | | | | | | | 47 | 0.407 | 1.000 | 48 | 0.090 | 0.596 | 54 | 0.113 | 0.596 | 214 | 0.990 | 0.990 | 495 | 0.830 | 0.990 |
| BRD-K54955827-001-02-2 | niraparib | 0.401 | | | | 118 | 0.815 | 0.986 | 317 | 0.867 | 0.986 | 692 | 0.986 | 0.986 | 217 | 0.051 | 1.000 | 283 | 0.751 | 0.986 | 501 | 0.224 | 1.000 |
| BRD-K54997624-001-06-0 | alpelisib | 0.972 | 88 | 0.628 | 0.942 | 101 | 0.221 | 0.553 | 360 | 0.790 | 0.972 | 532.5 | 0.046 | 0.971 | 497 | 0.276 | 0.580 | 581 | 0.926 | 0.972 | 279 | 0.190 | 0.553 |
| BRD-K55187425-236-05-2 | rigosertib | 0.522 | 171 | 0.677 | 0.931 | 549.5 | 0.861 | 0.996 | 870.5 | 0.199 | 0.522 | 1014 | 0.049 | 0.519 | 1401 | 0.154 | 0.522 | 1427.5 | 0.488 | 0.805 | 1423.5 | 0.174 | 0.522 |
| BRD-K55305701-001-01-2 | SC-144 | 0.986 | | | | 277.5 | 0.808 | 0.986 | 864 | 0.856 | 0.986 | 1232 | 0.790 | 0.986 | 1206.5 | 0.749 | 0.986 | 1078.5 | 0.957 | 0.986 | 1278.5 | 0.535 | 0.986 |
| BRD-K55696337-003-24-4 | topotecan | 0.115 | 54 | 0.119 | 0.278 | 2333 | 0.984 | 0.997 | 1388 | 0.019 | 0.115 | 4549 | 0.595 | 0.997 | 3694 | 0.110 | 0.278 | 4219 | 0.796 | 0.997 | 5299 | 0.780 | 0.997 |
| BRD-K56032964-001-02-1 | AP26113 | 0.651 | 114 | 0.474 | 0.980 | 134 | 0.589 | 0.980 | 346 | 0.475 | 0.980 | 1444 | 0.054 | 0.447 | 1314.5 | 0.916 | 0.980 | 949 | 0.777 | 0.980 | 2204.5 | 0.778 | 0.980 |
| BRD-K56277358-001-07-5 | MGCD-265 | 0.578 | | | | 327 | 0.275 | 0.578 | 812 | 0.268 | 0.578 | 1698 | 0.321 | 0.612 | 1213 | 0.173 | 0.578 | 1373 | 0.218 | 0.578 | 2084 | 0.805 | 0.995 |
| BRD-K56291712-001-01-0 | voxtalisib | 0.895 | | | | 134.5 | 0.886 | 1.000 | 268.5 | 0.658 | 0.967 | 843.5 | 0.737 | 0.967 | 368.5 | 0.418 | 0.895 | 330.5 | 0.574 | 0.967 | 1204.5 | 0.995 | 0.995 |
| BRD-K56301217-001-07-4 | ABT-737 | 0.488 | | | | | | | 157.5 | 0.135 | 0.488 | 364.5 | 0.836 | 1.000 | 318 | 0.185 | 0.533 | 111.5 | 0.932 | 1.000 | 433 | 0.969 | 0.969 |
| BRD-K56334280-001-05-1 | amonafide | 0.133 | | | | | | | 101.5 | 0.893 | 0.912 | 362.5 | 0.601 | 1.000 | 145.5 | 0.308 | 1.000 | 251 | 0.614 | 0.890 | 484.5 | 0.845 | 0.912 |
| BRD-K56405753-001-02-4 | MK-2461 | 0.938 | | | | 42 | 0.261 | 0.685 | 264 | 0.536 | 0.938 | 243 | 0.172 | 0.682 | 201 | 0.541 | 0.938 | 141 | 0.581 | 0.938 | 560 | 0.726 | 0.955 |
| BRD-K56707426-001-02-0 | nitarsone | 0.950 | | | | 121 | 0.950 | 0.950 | 157 | 0.751 | 0.950 | 646 | 0.742 | 0.950 | 346 | 0.721 | 0.950 | 116 | 0.449 | 0.893 | 522 | 0.880 | 0.950 |
| BRD-K56957086-001-06-3 | dacinostat | 0.196 | 36 | 0.081 | 0.261 | 1424 | 0.332 | 0.536 | 1696 | 0.095 | 0.261 | 3656 | 0.058 | 0.261 | 3925 | 0.132 | 0.263 | 2358 | 0.028 | 0.196 | 4361 | 0.090 | 0.261 |
| BRD-K56981171-001-01-0 | brigatinib | 0.901 | 122 | 0.602 | 0.903 | 187 | 0.389 | 0.901 | 228 | 0.059 | 0.426 | 1295 | 0.061 | 0.426 | 918.5 | 0.976 | 0.984 | 972.5 | 0.939 | 0.984 | 1294 | 0.478 | 0.901 |

Suppl. Table 9

| broad_id | name | colorectal_MWU_STAT | colorectal_PVAL | colorectal_PVAL_ADJ | skin_MWU_STAT | skin_PVAL | skin_PVAL_ADJ | urinary_tract_MWU_STAT | urinary_tract_PVAL | urinary_tract_PVAL_ADJ | lung_MWU_STAT | lung_PVAL | lung_PVAL_ADJ | ovary_MWU_STAT | ovary_PVAL | ovary_PVAL_ADJ | breast_MWU_STAT | breast_PVAL | breast_PVAL_ADJ | pancreas_MWU_STAT | pancreas_PVAL | pancreas_PVAL_ADJ | nervous_system_MWU_STAT | central_nervous_system_PVAL | nervous_system_PVAL_ADJ | soft_tissue_MWU_STAT | soft_tissue_PVAL | soft_tissue_PVAL_ADJ | kidney_MWU_STAT | kidney_PVAL | kidney_PVAL_ADJ | gastric_MWU_STAT | gastric_PVAL | gastric_PVAL_ADJ | bone_MWU_STAT | bone_PVAL | bone_PVAL_ADJ | prostate_MWU_STAT | prostate_PVAL | prostate_PVAL_ADJ | al_nervous_system_MWU_STAT | peripheral_nervous_system_PVAL |
|---|---|---|---|---|---|---|---|---|---|---|---|---|---|---|---|---|---|---|---|---|---|---|---|---|---|---|---|---|---|---|---|---|---|---|---|---|---|---|---|---|---|---|
| BRD-K56981171-001-02-8 | brigatinib | 627 | 0.808 | 0.893 | 670 | 0.977 | 0.977 | 972 | 0.592 | 0.822 | 2166 | 0.626 | 0.822 | 681 | 0.350 | 0.799 | 998 | 0.971 | 0.977 | 598 | 0.080 | 0.335 | 287 | 0.418 | 0.799 | 224 | 0.467 | 0.817 | 177 | 0.246 | 0.720 | 382 | 0.259 | 0.720 | 117 | 0.073 | 0.335 | 198 | 0.752 | 0.877 | 255 | 0.623 |
| BRD-K57080016-001-15-9 | selumetinib | 715.5 | 0.002 | 0.019 | 841.5 | 0.000 | 0.000 | 749 | 0.380 | 0.983 | 3105.5 | 0.983 | 0.983 | 1196 | 0.975 | 0.983 | 921.5 | 0.926 | 0.983 | 1219 | 0.242 | 0.845 | 1225 | 0.941 | 0.983 | 495 | 0.898 | 0.983 | 870 | 0.964 | 0.983 | 95 | 0.023 | 0.123 | 448.5 | 0.974 | 0.983 | 389 | 0.899 | 0.983 | 166.5 | 0.409 |
| BRD-K57136142-001-02-4 | CCT129202 | 1391 | 0.960 | 1.000 | 3212 | 0.999 | 1.000 | 1699.5 | 0.377 | 0.659 | 4635 | 0.798 | 1.000 | 1364 | 0.201 | 0.477 | 1451 | 0.649 | 0.909 | 1137.5 | 0.041 | 0.277 | 2762 | 0.941 | 1.000 | 158 | 0.050 | 0.277 | 520 | 0.053 | 0.277 | 1071 | 0.208 | 0.477 | 452 | 0.193 | 0.477 | 345 | 0.472 | 0.707 | 263 | 0.066 |
| BRD-K57169635-001-04-5 | dacomitinib | 1655 | 0.312 | 0.595 | 3945 | 0.999 | 1.000 | 1527 | 0.004 | 0.026 | 8583 | 0.943 | 1.000 | 2040 | 0.118 | 0.498 | 2059 | 0.342 | 0.599 | 2118 | 0.411 | 0.664 | 4089 | 1.000 | 1.000 | 1210 | 0.839 | 1.000 | 2235 | 0.929 | 1.000 | 1162 | 0.181 | 0.507 | 749 | 0.850 | 1.000 | | | | 473 | 0.610 |
| BRD-K57284066-001-01-6 | Ro-9187 | 421 | 0.039 | 0.684 | 1474 | 0.853 | 0.919 | 910 | 0.521 | 0.781 | 2776 | 0.735 | 0.902 | 1128.5 | 0.504 | 0.781 | 782 | 0.392 | 0.781 | 933 | 0.757 | 0.902 | 732 | 0.276 | 0.781 | 190 | 0.065 | 0.684 | 640 | 0.422 | 0.781 | 249 | 0.461 | 0.781 | 163 | 0.447 | 0.781 | 155 | 0.912 | 0.919 | 133 | 0.290 |
| BRD-K58049875-001-04-7 | thiostrepton | 2044 | 0.919 | 0.981 | 1934 | 0.403 | 0.941 | 1573 | 0.521 | 0.981 | 5750 | 0.367 | 0.941 | 1573 | 0.069 | 0.488 | 1944 | 0.568 | 0.981 | 1671 | 0.819 | 0.981 | 2286 | 0.832 | 0.981 | 1123 | 0.759 | 0.981 | 1483 | 0.963 | 0.981 | 1229 | 0.890 | 0.981 | 444 | 0.139 | 0.488 | 127 | 0.508 | 0.981 | 472 | 0.780 |
| BRD-K58122457-001-02-2 | hesperadin | 2845 | 0.677 | 0.991 | 4410 | 0.898 | 0.991 | 1795 | 0.026 | 0.242 | 10048 | 0.415 | 0.872 | 2495 | 0.069 | 0.291 | 3212.5 | 0.968 | 0.991 | 2602 | 0.293 | 0.688 | 3912 | 0.799 | 0.991 | 835 | 0.044 | 0.242 | 2143 | 0.965 | 0.991 | 1288 | 0.144 | 0.504 | 388 | 0.006 | 0.121 | 1106 | 0.991 | 0.991 | 360 | 0.213 |
| BRD-K58148589-001-03-6 | betamethasone-dipropionate | 456 | 0.582 | 0.889 | 704 | 0.498 | 1.000 | 485 | 0.111 | 0.780 | 2046 | 0.807 | 0.889 | 992 | 0.739 | 0.889 | 232 | 0.236 | 0.990 | 646 | 0.096 | 0.780 | 435 | 0.771 | 0.889 | 216 | 0.470 | 0.889 | 318 | 0.131 | 0.306 | 311 | 0.043 | 0.780 | 160 | 0.575 | 0.889 | 152 | 0.523 | 0.889 | 167 | 0.620 |
| BRD-K58262659-001-09-7 | doxifluridine | 297 | 0.428 | 0.835 | 702 | 0.205 | 1.000 | 520 | 0.131 | 1.000 | 1670 | 0.827 | 0.918 | 690.5 | 0.787 | 0.918 | 23 | 0.137 | 0.481 | 768.5 | 0.831 | 0.918 | 296 | 0.011 | 0.030 | 110 | 0.370 | 0.835 | 380 | 0.795 | 0.918 | 409 | 0.661 | 0.918 | | | | 124 | 0.945 | 0.945 | 114 | 0.909 |
| BRD-K58435339-001-03-0 | AT13387 | 1668.5 | 0.194 | 0.601 | 4874 | 0.956 | 0.986 | 2844 | 0.327 | 0.601 | 8283.5 | 0.039 | 0.462 | 2923 | 0.106 | 0.558 | 2443 | 0.831 | 0.986 | 5377 | 0.982 | 0.986 | 3872.5 | 0.895 | 0.986 | 1394 | 0.103 | 0.558 | 2385 | 0.967 | 0.986 | 1566 | 0.233 | 0.601 | 2055 | 0.914 | 0.986 | 417 | 0.307 | 0.601 | 398 | 0.269 |
| BRD-K58466253-003-14-0 | palmatine-chloride | 748 | 0.945 | 0.945 | 693 | 0.809 | 0.940 | 379 | 0.089 | 0.469 | 1244 | 0.862 | 0.940 | 421 | 0.074 | 0.469 | 145 | 0.748 | 0.940 | 310 | 0.049 | 0.469 | 490 | 0.895 | 0.940 | 76 | 0.084 | 0.196 | 216 | 0.792 | 0.940 | 296 | 0.474 | 0.904 | 145 | 0.748 | 0.940 | | | | 73 | 0.684 |
| BRD-K58529924-001-01-5 | ONC201 | 359 | 0.195 | 0.744 | 775 | 0.153 | 0.744 | 92 | 0.009 | 0.179 | 2094.5 | 0.507 | 0.930 | 933.5 | 0.664 | 0.930 | 753.5 | 0.554 | 0.930 | 763.5 | 0.583 | 0.930 | 788.5 | 0.651 | 0.930 | 332.5 | 0.319 | 0.744 | 481.5 | 0.847 | 0.971 | 434 | 0.220 | 0.744 | 193.5 | 0.317 | 0.744 | 44 | 0.041 | 0.435 | 312 | 0.853 |
| BRD-K58550667-001-08-7 | FK-866 | 3220 | 0.166 | 0.438 | 6228 | 0.974 | 0.995 | 3391 | 0.902 | 0.995 | 10755 | 0.750 | 0.995 | 4137 | 0.167 | 0.438 | 3130 | 0.450 | 0.788 | 4751 | 0.930 | 0.995 | 5303 | 0.885 | 0.995 | 1221 | 0.015 | 0.139 | 3114 | 0.995 | 0.995 | 2800 | 0.618 | 0.951 | 1262 | 0.020 | 0.139 | 1072 | 0.942 | 0.995 | 284 | 0.017 |
| BRD-K58770988-001-01-8 | NMS-E973 | 6943 | 0.999 | 0.999 | 7634 | 0.464 | 0.827 | 4107 | 0.476 | 0.827 | 15018 | 0.290 | 0.762 | 6005 | 0.692 | 0.908 | 4736 | 0.762 | 0.942 | 6983 | 0.991 | 0.999 | 6098 | 0.413 | 0.827 | 1568 | 0.012 | 0.107 | 4138 | 0.995 | 0.999 | 3056 | 0.429 | 0.827 | 1430 | 0.039 | 0.206 | 486 | 0.069 | 0.243 | 1140 | 0.593 |
| BRD-K58772419-001-07-0 | AZD6482 | 5404 | 0.899 | 0.993 | 6682 | 0.337 | 0.885 | 3712 | 0.469 | 0.984 | 12624 | 0.020 | 0.419 | 5601 | 0.735 | 0.993 | 3457 | 0.720 | 0.993 | 6515 | 0.965 | 0.994 | 4600 | 0.150 | 0.655 | 2559 | 0.809 | 0.993 | 2927 | 0.292 | 0.877 | 2532 | 0.156 | 0.655 | 1782 | 0.455 | 0.984 | 702 | 0.653 | 0.993 | 307 | 0.071 |
| BRD-K59146805-001-03-7 | AZ3146 | 1528 | 0.401 | 0.763 | 2387 | 0.783 | 0.983 | 835 | 0.049 | 0.510 | 3648 | 0.098 | 0.685 | 2376 | 0.968 | 0.983 | 1435 | 0.958 | 0.983 | 1564 | 0.892 | 0.983 | 1163 | 0.084 | 0.253 | 442 | 0.082 | 0.253 | 1194 | 0.960 | 0.983 | 531 | 0.436 | 0.763 | 409 | 0.742 | 0.983 | 388 | 0.961 | 0.983 | 204 | 0.412 |
| BRD-K59317601-001-05-5 | MLN0128 | 4935 | 1.000 | 1.000 | 4917 | 0.610 | 0.754 | 3038 | 0.419 | 0.734 | 9620 | 0.389 | 0.734 | 2470 | 0.003 | 0.069 | 2764 | 0.452 | 0.734 | 3153 | 0.172 | 0.722 | 4943 | 0.999 | 1.000 | 1161 | 0.111 | 0.722 | 2293 | 0.296 | 0.734 | 1362 | 0.148 | 0.722 | 1512 | 0.483 | 0.734 | 619 | 0.366 | 0.734 | 585 | 0.652 |
| BRD-K59325863-001-02-8 | delanzomib | 5766 | 0.730 | 0.973 | 7471 | 0.207 | 0.844 | 5174 | 0.753 | 0.973 | 18906 | 0.928 | 0.975 | 6557 | 0.880 | 0.973 | 5349 | 0.835 | 0.973 | 5695 | 0.041 | 0.285 | 4579 | 0.001 | 0.011 | 2405 | 0.466 | 0.970 | 3738 | 0.669 | 0.973 | 2096 | 0.003 | 0.032 | 2034 | 0.322 | 0.844 | 907 | 0.508 | 0.970 | 1020 | 0.364 |
| BRD-K59369769-001-20-3 | tozasertib | 4088 | 0.353 | 0.674 | 7836 | 0.991 | 0.997 | 3284 | 0.271 | 0.590 | 11174 | 0.025 | 0.239 | 5493 | 0.614 | 0.992 | 3958 | 0.945 | 0.997 | 6392 | 0.997 | 0.997 | 4482 | 0.281 | 0.590 | 904 | 0.002 | 0.043 | 2686 | 0.936 | 0.997 | 2331 | 0.208 | 0.590 | 1691 | 0.128 | 0.536 | 830 | 0.883 | 0.997 | 370 | 0.034 |
| BRD-K59369769-001-22-9 | tozasertib | 796 | 0.921 | 0.981 | 1258 | 0.819 | 0.981 | 642 | 0.048 | 0.298 | 2428 | 0.420 | 0.882 | 827 | 0.320 | 0.839 | 700 | 0.981 | 0.981 | 783 | 0.978 | 0.981 | 654 | 0.229 | 0.687 | 231 | 0.017 | 0.298 | 634 | 0.932 | 0.981 | 526 | 0.700 | 0.981 | 112 | 0.057 | 0.298 | 112 | 0.755 | 0.981 | 122 | 0.073 |
| BRD-K59436580-001-01-7 | tedizolid | 1909.5 | 0.862 | 0.955 | 1805 | 0.455 | 0.955 | 1058 | 0.393 | 0.920 | 4526.5 | 0.394 | 0.920 | 2236 | 0.864 | 0.955 | 619 | 0.164 | 0.575 | 1273 | 0.076 | 0.516 | 2503.5 | 0.998 | 0.998 | 618 | 0.163 | 0.575 | 470 | 0.098 | 0.516 | 555 | 0.011 | 0.236 | 602.5 | 0.987 | 0.998 | 358 | 0.546 | 0.819 | 76 | 0.278 |
| BRD-K59753975-001-02-6 | vindesine | 6034.5 | 0.995 | 1.000 | 6605 | 0.727 | 0.935 | 3552 | 0.120 | 0.362 | 15147 | 0.390 | 0.745 | 5287 | 0.488 | 0.789 | 3864 | 0.688 | 0.935 | 7626.5 | 1.000 | 1.000 | 5933 | 0.441 | 0.772 | 1135 | 0.003 | 0.033 | 5142 | 1.000 | 1.000 | 2293 | 0.054 | 0.225 | 1594 | 0.240 | 0.603 | 675 | 0.259 | 0.603 | 894 | 0.298 |
| BRD-K59831625-001-01-6 | LDN-212854 | 3918 | 0.236 | 0.617 | 7164 | 0.908 | 0.908 | 3605 | 0.261 | 0.617 | 15997 | 0.879 | 0.908 | 4688 | 0.382 | 0.617 | 3747 | 0.644 | 0.901 | 6076 | 0.692 | 0.901 | 4614 | 0.098 | 0.617 | 2498 | 0.900 | 0.908 | 2495 | 0.244 | 0.617 | 2151 | 0.262 | 0.617 | 1295 | 0.167 | 0.617 | 717 | 0.339 | 0.617 | 837 | 0.249 |
| BRD-K59866480-001-07-3 | cyclovalone | 3881 | 0.243 | 0.567 | 7528 | 0.929 | 0.990 | 3078 | 0.030 | 0.249 | 16157 | 0.990 | 0.990 | 5049 | 0.548 | 0.767 | 3683 | 0.483 | 0.767 | 4515 | 0.310 | 0.650 | 7078 | 0.973 | 0.990 | 1637 | 0.540 | 0.767 | 3247 | 0.588 | 0.772 | 2093 | 0.237 | 0.567 | 1418 | 0.059 | 0.249 | 937 | 0.700 | 0.864 | 541 | 0.037 |
| BRD-K59983611-001-21-9 | sulfabenzamide | 672.5 | 0.937 | 0.945 | 1262 | 0.565 | 1.000 | 968.5 | 0.243 | 0.850 | 2586.5 | 0.858 | 0.945 | 723 | 0.008 | 0.166 | 370.5 | 0.656 | 1.000 | 341 | 0.102 | 0.713 | 1041 | 0.715 | 1.000 | 116 | 0.228 | 0.850 | 513 | 0.572 | 1.000 | 687.5 | 0.822 | 0.945 | 329 | 0.490 | 1.000 | 357.5 | 0.898 | 0.945 | 143 | 0.362 |
| BRD-K60184833-001-05-6 | tyrphostin-AG-99 | 3075 | 0.905 | 0.977 | 4214.5 | 0.299 | 0.742 | 2417 | 0.495 | 0.742 | 9879.5 | 0.282 | 0.742 | 3297 | 0.154 | 0.742 | 2296 | 0.703 | 0.922 | 4732 | 0.999 | 0.999 | 4284 | 0.786 | 0.970 | 1059 | 0.349 | 0.742 | 2198 | 0.895 | 0.977 | 2041.5 | 0.419 | 0.742 | 834 | 0.248 | 0.742 | 403 | 0.278 | 0.742 | 382 | 0.238 |
| BRD-K60230970-001-17-5 | MG-132 | 5627 | 0.691 | 0.876 | 8234 | 0.783 | 0.876 | 5405 | 0.792 | 0.876 | 16967 | 0.716 | 0.876 | 6599 | 0.768 | 0.876 | 4791 | 0.689 | 0.876 | 7337 | 0.737 | 0.876 | 5974 | 0.104 | 0.436 | 2586 | 0.463 | 0.876 | 3233 | 0.198 | 0.669 | 2406 | 0.042 | 0.436 | 2906 | 0.876 | 0.920 | 830 | 0.409 | 0.860 | 897 | 0.234 |
| BRD-K60341624-001-02-2 | GSK1292263 | 980 | 0.566 | 0.849 | 1877 | 0.923 | 0.990 | 1705 | 0.947 | 0.990 | 3811 | 0.370 | 0.851 | 1453 | 0.362 | 0.851 | 1158 | 0.338 | 0.851 | 1012 | 0.067 | 0.523 | 1601 | 0.950 | 0.990 | 554 | 0.125 | 0.523 | 677 | 0.840 | 0.990 | 1121 | 0.446 | 0.851 | 363 | 0.636 | 0.890 | 388 | 0.717 | 0.942 | | |
| BRD-K60348325-065-09-9 | brilliant-green | 4792 | 0.371 | 0.947 | 7349 | 0.443 | 0.947 | 4692 | 0.565 | 0.947 | 15853 | 0.538 | 0.947 | 5228 | 0.135 | 0.471 | 5718 | 0.997 | 0.999 | 6595 | 0.746 | 0.999 | 7171 | 0.889 | 0.999 | 1841 | 0.047 | 0.248 | 5165 | 0.999 | 0.999 | 2280 | 0.030 | 0.213 | 2283 | 0.632 | 0.947 | 973 | 0.922 | 0.999 | 1077 | 0.486 |
| BRD-K60443845-001-08-6 | chlormidazole | 2211 | 0.690 | 0.960 | 3293 | 0.828 | 0.960 | 2412 | 0.618 | 0.960 | 5785 | 0.040 | 0.170 | 2910.5 | 0.472 | 0.960 | 2226 | 0.839 | 0.960 | 2587 | 0.525 | 0.960 | 3045 | 0.914 | 0.960 | 749 | 0.019 | 0.101 | 1243 | 0.696 | 0.960 | 1512 | 0.476 | 0.960 | 567 | 0.082 | 0.288 | 407 | 0.441 | 0.960 | 453 | 0.568 |
| BRD-K60446698-001-01-2 | ONX-0914 | 6007 | 0.851 | 0.933 | 5824 | 0.015 | 0.165 | 4727 | 0.603 | 0.916 | 16269 | 0.715 | 0.933 | 5665 | 0.573 | 0.916 | 3859 | 0.281 | 0.916 | 7449 | 0.864 | 0.933 | 6038 | 0.452 | 0.916 | 2478 | 0.429 | 0.916 | 4754 | 0.997 | 0.997 | 2842 | 0.142 | 0.743 | 1456 | 0.016 | 0.165 | 496 | 0.247 | 0.916 | 1197 | 0.654 |
| BRD-K60997853-001-02-3 | PHA-848125 | 2478 | 0.937 | 1.000 | 4366 | 0.613 | 0.991 | 2802 | 0.105 | 0.483 | 9267.5 | 0.187 | 0.490 | 2178 | 0.018 | 0.191 | 2925 | 0.963 | 1.000 | 4725.5 | 0.909 | 1.000 | 5463 | 1.000 | 1.000 | 645 | 0.002 | 0.039 | 2169 | 0.236 | 0.550 | 1083 | 0.187 | 0.490 | 1595 | 0.835 | 1.000 | 427 | 0.741 | 1.000 | 575 | 0.115 |
| BRD-K61097567-001-08-5 | SB-218795 | 347.5 | 0.704 | 0.900 | 372 | 0.067 | 0.703 | 418.5 | 0.739 | 0.900 | 1175.5 | 0.450 | 1.000 | 604 | 0.570 | 1.000 | 713 | 0.877 | 0.969 | 477.5 | 0.731 | 0.900 | 866.5 | 0.580 | 1.000 | 276 | 0.664 | 1.000 | 271.5 | 0.340 | 1.000 | 346 | 0.919 | 0.919 | 232 | 0.771 | 0.900 | | | | 10 | 0.075 |
| BRD-K61195623-001-01-4 | combretastatin-A-4 | 4241.5 | 0.407 | 0.810 | 5888.5 | 0.578 | 0.810 | 3924 | 0.574 | 0.810 | 13520.5 | 0.182 | 0.546 | 3483 | 0.014 | 0.291 | 4398 | 0.973 | 0.999 | 6217 | 0.999 | 0.999 | 6039.5 | 0.766 | 0.956 | 1420 | 0.154 | 0.538 | 2540 | 0.943 | 0.999 | 2046 | 0.119 | 0.498 | 994 | 0.036 | 0.308 | 822 | 0.542 | 0.810 | 402 | 0.044 |
| BRD-K61228301-001-01-2 | SGI-1027 | 2607 | 0.232 | 0.609 | 4146 | 0.393 | 0.821 | 2109 | 0.430 | 0.821 | 9407 | 0.906 | 0.994 | 4256 | 0.876 | 0.994 | 1850 | 0.022 | 0.152 | 3929 | 0.860 | 0.994 | 4538 | 0.962 | 0.994 | 1017 | 0.074 | 0.391 | 1371 | 0.967 | 0.994 | 744 | 0.001 | 0.014 | 537 | 0.298 | 0.694 | 571 | 0.702 | 0.994 | 359 | 0.609 |
| BRD-K61314889-001-04-4 | endo-IWR-1 | 2035.5 | 0.808 | 0.999 | 3445 | 0.772 | 0.999 | 1793 | 0.349 | 0.826 | 6915 | 0.222 | 0.826 | 2624 | 0.449 | 0.826 | 1753 | 0.463 | 0.826 | 3053.5 | 0.927 | 1.000 | 1940 | 0.010 | 0.104 | 707 | 0.250 | 0.826 | 1487.5 | 0.456 | 0.826 | 1366.5 | 0.472 | 0.826 | 809 | 0.930 | 1.000 | 82 | 0.008 | 0.104 | 503 | 0.706 |
| BRD-K61323504-001-04-0 | SB-225002 | 3304 | 0.959 | 0.959 | 4036 | 0.793 | 0.979 | 2438 | 0.538 | 0.979 | 7113 | 0.199 | 0.710 | 2698 | 0.179 | 0.710 | 1637 | 0.567 | 0.979 | 3358 | 0.989 | 1.000 | 3431 | 0.731 | 0.979 | 824 | 0.674 | 0.979 | 1404 | 0.994 | 1.000 | 2079 | 0.758 | 0.979 | 579 | 0.002 | 0.046 | 352 | 0.672 | 0.979 | 112 | 0.013 |
| BRD-K61397605-001-03-4 | romidepsin | 6557 | 0.885 | 0.999 | 6575 | 0.043 | 0.353 | 4075 | 0.130 | 0.353 | 16238 | 0.314 | 0.659 | 6650 | 0.846 | 0.999 | 4627 | 0.555 | 0.981 | 8858 | 0.996 | 0.999 | 7900 | 0.906 | 0.999 | 2721 | 0.562 | 0.981 | 4801 | 0.999 | 0.999 | 2984 | 0.084 | 0.353 | 3362 | 0.943 | 0.999 | 365 | 0.090 | 0.353 | 1204 | 0.607 |
| BRD-K61443650-236-13-1 | thiomersal | 682 | 0.168 | 0.560 | 3430 | 0.741 | 0.915 | 2112 | 0.980 | 0.995 | 5549 | 0.638 | 0.915 | 2058 | 0.267 | 0.560 | 2399 | 0.953 | 0.995 | 2792 | 0.995 | 0.995 | 2097 | 0.732 | 0.915 | 314 | 0.006 | 0.032 | 463 | 0.000 | 0.003 | 447 | 0.046 | 0.966 | 495 | 0.216 | 0.560 | 217 | 0.367 | 0.616 | 306 | 0.293 |
| BRD-K61496577-001-20-2 | flumethasone | 436 | 0.638 | 1.000 | 1351 | 0.814 | 1.000 | 626 | 0.495 | 1.000 | 1890 | 0.160 | 0.701 | 957 | 0.426 | 1.000 | 117 | 0.060 | 0.628 | 717 | 0.022 | 0.457 | 1471 | 0.993 | 1.000 | 166 | 0.521 | 1.000 | 158 | 0.147 | 0.701 | 545 | 0.728 | 0.959 | 676 | 0.961 | 1.000 | 175 | 0.575 | 0.880 | 177 | 0.587 |
| BRD-K61536336-001-01-7 | SB-366791 | 627 | 0.492 | 0.778 | 1072.5 | 0.798 | 0.985 | 630 | 0.502 | 0.778 | 2215.5 | 0.881 | 1.000 | 709 | 0.120 | 0.490 | 176 | 0.283 | 0.678 | 622 | 0.476 | 0.778 | 742.5 | 0.291 | 0.678 | 211 | 0.460 | 0.778 | 249 | 0.660 | 0.884 | 212 | 0.181 | 1.000 | | | | 81 | 0.140 | 0.490 | 173 | 0.269 |
| BRD-K61734568-003-02-1 | 2,3-DCPE | 714 | 0.562 | 0.787 | 1132 | 0.551 | 0.787 | 1027.5 | 0.966 | 0.966 | 1951.5 | 0.608 | 0.797 | 1060.5 | 0.542 | 0.787 | 341 | 0.303 | 0.787 | 918 | 0.532 | 0.787 | 848 | 0.533 | 0.787 | 243 | 0.517 | 0.787 | 486.5 | 0.556 | 0.787 | 355.5 | 0.354 | 0.787 | 166 | 0.014 | 0.146 | 127 | 0.839 | 0.966 | 162 | 0.509 |
| BRD-K61936403-001-01-7 | AS-1949490 | 503 | 0.915 | 1.000 | 332 | 0.059 | 0.624 | 602 | 0.845 | 1.000 | 1107 | 0.007 | 0.146 | 722 | 0.681 | 1.000 | 380 | 0.511 | 1.000 | 815 | 0.991 | 0.991 | 438 | 0.292 | 0.961 | 217 | 0.643 | 1.000 | 309 | 0.761 | 1.000 | 465 | 0.961 | 0.991 | 54 | 0.386 | 1.000 | 94 | 0.254 | 0.961 | 70 | 0.134 |
| BRD-K62008436-001-22-1 | paclitaxel | 7068 | 0.962 | 1.000 | 9620 | 0.957 | 1.000 | 3813 | 0.024 | 0.130 | 16067 | 0.120 | 0.313 | 4540 | 0.024 | 0.130 | 5587 | 0.889 | 1.000 | 7662 | 0.859 | 1.000 | 7878 | 0.860 | 1.000 | 2050 | 0.077 | 0.271 | 6299.5 | 1.000 | 1.000 | 2987 | 0.134 | 0.313 | 2668 | 0.472 | 0.902 | 1176 | 0.836 | 1.000 | 1228 | 0.614 |
| BRD-K62008436-001-23-9 | paclitaxel | 1126 | 0.785 | 0.970 | 1705 | 0.417 | 0.800 | 1076 | 0.002 | 0.046 | 5045 | 0.208 | 0.734 | 1543 | 0.210 | 0.734 | 1324 | 0.962 | 1.000 | 2119 | 0.906 | 1.000 | 1889 | 0.525 | 0.800 | 1021 | 0.605 | 0.847 | 2344 | 1.000 | 1.000 | 1448 | 0.533 | 0.800 | 664 | 0.350 | 0.800 | 455 | 0.753 | 0.970 | 213 | 0.367 |
| BRD-K62196610-001-01-6 | AVN-944 | 815.5 | 0.452 | 0.730 | 1890.5 | 0.969 | 0.996 | 544 | 0.351 | 0.637 | 2501 | 0.524 | 0.787 | 733 | 0.013 | 0.045 | 1154 | 0.996 | 0.996 | 1097 | 0.989 | 0.996 | 617 | 0.953 | 0.996 | 212 | 0.007 | 0.030 | 666 | 0.274 | 0.575 | 590.5 | 0.273 | 0.575 | 674.5 | 0.907 | 0.996 | 384 | 0.921 | 0.996 | 106 | 0.039 |
| BRD-K62304154-001-04-2 | deforolimus | 5339 | 0.998 | 0.998 | 7061 | 0.830 | 0.998 | 3522 | 0.103 | 0.319 | 15757 | 0.821 | 0.998 | 4286 | 0.044 | 0.308 | 4543 | 0.837 | 0.998 | 7408 | 0.997 | 0.998 | 5483 | 0.127 | 0.333 | 1407 | 0.106 | 0.319 | 3327 | 0.723 | 0.998 | 2429 | 0.090 | 0.319 | 2038 | 0.156 | 0.363 | 635 | 0.204 | 0.429 | 1206 | 0.937 |
| BRD-K62310379-001-10-5 | fluticasone-propionate | 1289 | 0.835 | 1.000 | 1163 | 0.909 | 1.000 | 652 | 0.128 | 0.794 | 2953 | 0.367 | 0.855 | 1414 | 0.316 | 0.855 | 673 | 0.532 | 0.932 | 1274 | 0.666 | 1.000 | 1129 | 0.958 | 1.000 | 271 | 0.158 | 0.794 | 824 | 0.145 | 0.794 | 796 | 0.393 | 0.855 | 457 | 0.744 | 1.000 | 183 | 0.448 | 0.855 | 302 | 0.913 |
| BRD-K62374002-001-01-5 | PI3K-IN-2 | 5323 | 0.978 | 1.000 | 8211 | 0.846 | 0.988 | 3824 | 0.092 | 0.275 | 16174 | 0.423 | 0.888 | 4835 | 0.045 | 0.254 | 4890 | 0.693 | 0.970 | 6510 | 0.599 | 0.898 | 8431 | 1.000 | 1.000 | 1358 | 0.062 | 0.260 | 3735 | 0.754 | 0.974 | 2194 | 0.020 | 0.254 | 2183 | 0.534 | 0.898 | 667 | 0.217 | 0.570 | 1136 | 0.567 |
| BRD-K62627508-001-01-5 | idasanutlin | 1582.5 | 0.812 | 1.000 | 1579.5 | 0.000 | 0.003 | 1235.5 | 0.743 | 1.000 | 3567 | 0.951 | 1.000 | 1303 | 0.117 | 0.693 | 820.5 | 0.502 | 1.000 | 1640 | 1.000 | 1.000 | 1689 | 0.393 | 1.000 | 656.5 | 0.161 | 0.693 | 772 | 0.006 | 0.058 | 769 | 0.197 | 0.693 | 324.5 | 0.215 | 0.693 | 332.5 | 0.570 | 1.000 | 149 | 0.231 |
| BRD-K62696033-001-01-4 | G-1 | 412.5 | 0.723 | 0.981 | 710 | 0.376 | 1.000 | 776.5 | 0.748 | 0.981 | 1873 | 0.612 | 1.000 | 933.5 | 0.490 | 1.000 | 853.5 | 0.967 | 1.000 | 750.5 | 0.948 | 1.000 | 810 | 0.485 | 1.000 | 87 | 0.038 | 0.289 | 688.5 | 0.962 | 1.000 | 270.5 | 0.768 | 0.768 | 90 | 0.041 | 0.289 | | | | 230 | 0.247 |
| BRD-K62779383-001-05-5 | cholecalciferol | 675 | 0.772 | 0.954 | 2104 | 0.851 | 0.992 | 606 | 0.172 | 0.769 | 3782 | 0.621 | 0.850 | 983 | 0.036 | 0.759 | 603.5 | 0.589 | 0.850 | 923.5 | 0.295 | 0.769 | 1112 | 0.337 | 0.769 | 229 | 0.083 | 0.769 | 938 | 0.996 | 1.000 | 874.5 | 0.366 | 0.769 | 322 | 0.626 | 0.850 | 119 | 0.647 | 0.850 | 144 | 0.264 |
| BRD-K62814476-001-03-3 | BMS-265246 | 2209 | 0.061 | 1.000 | 6792 | 1.000 | 1.000 | 2735 | 0.400 | 0.895 | 9276 | 0.008 | 0.044 | 2701 | 0.002 | 0.017 | 3384 | 0.962 | 1.000 | 3677 | 0.798 | 0.994 | 4867 | 0.990 | 1.000 | 851 | 0.476 | 0.909 | 2024 | 0.315 | 0.828 | 1685 | 0.289 | 0.828 | 1505 | 0.681 | 0.953 | 489 | 0.426 | 0.895 | 558 | 0.583 |
| BRD-K62965247-001-04-9 | tipifarnib | 4355.5 | 0.970 | 0.971 | 7360 | 0.958 | 0.971 | 2826 | 0.155 | 0.565 | 10841.5 | 0.068 | 0.565 | 4269 | 0.161 | 0.565 | 3364 | 0.971 | 0.971 | 5239 | 0.919 | 0.971 | 4500 | 0.389 | 0.877 | 1128 | 0.016 | 0.339 | 2127.5 | 0.772 | 0.971 | 2185 | 0.084 | 0.565 | 1472 | 0.460 | 0.877 | 715 | 0.420 | 0.877 | 618.5 | 0.598 |
| BRD-K63068307-001-08-9 | ZSTK-474 | 2211 | 0.987 | 0.987 | 2640.5 | 0.501 | 0.874 | 950.5 | 0.201 | 0.642 | 5420.5 | 0.533 | 0.874 | 1949 | 0.074 | 0.519 | 1567.5 | 0.605 | 0.907 | 3449.5 | 0.978 | 0.987 | 1593 | 0.182 | 0.642 | 345 | 0.011 | 0.200 | 1320.5 | 0.242 | 0.642 | 968 | 0.019 | 0.200 | 567.5 | 0.328 | 0.690 | 157 | 0.168 | 0.642 | 270.5 | 0.541 |
| BRD-K63126190-003-01-6 | epinephrine | 822 | 0.949 | 0.995 | 966 | 0.354 | 0.995 | 871 | 0.442 | 0.995 | 1596 | 0.055 | 0.311 | 364 | 0.074 | 0.311 | 941 | 0.995 | 0.995 | 803 | 0.633 | 0.995 | 904 | 0.852 | 0.995 | 401 | 0.545 | 0.995 | 416 | 0.602 | 0.995 | 597 | 0.686 | 0.995 | 125 | 0.836 | 0.995 | 212 | 0.793 | 0.995 | 129 | 0.857 |
| BRD-K63504947-001-15-4 | semaxanib | 1721 | 0.964 | 0.964 | 1741 | 0.013 | 0.281 | 1784 | 0.509 | 1.000 | 6859 | 0.363 | 1.000 | 1959 | 0.059 | 0.302 | 985 | 0.045 | 0.302 | 2627 | 0.945 | 1.000 | 2349 | 0.623 | 1.000 | 1126 | 0.854 | 1.000 | 1919 | 0.978 | 1.000 | 1250 | 0.645 | 1.000 | 578 | 0.134 | 0.471 | 475 | 0.720 | 1.000 | 4 | 0.048 |
| BRD-K63736853-001-04-8 | acipimox | 583 | 0.978 | 0.978 | 648 | 0.016 | 0.264 | 386 | 0.473 | 1.000 | 1732 | 0.911 | 1.000 | 219 | 0.275 | 0.826 | 513 | 0.717 | 1.000 | 627 | 0.673 | 1.000 | 681 | 0.650 | 1.000 | 364 | 0.897 | 1.000 | 273 | 0.258 | 0.826 | 233 | 0.133 | 0.699 | 222 | 0.625 | 1.000 | | | | | |
| BRD-K63750851-001-20-7 | mycophenolic-acid | 1787 | 0.656 | 0.911 | 3511 | 0.879 | 0.930 | 1829 | 0.244 | 0.644 | 6598 | 0.640 | 0.911 | 2379 | 0.482 | 0.844 | 1806 | 0.930 | 0.930 | 2603 | 0.738 | 0.911 | 2204 | 0.922 | 0.930 | 464 | 0.041 | 0.261 | 1137 | 0.246 | 0.644 | 1222 | 0.050 | 0.261 | 378 | 0.166 | 0.581 | 246 | 0.420 | 0.811 | 44 | 0.166 |
| BRD-K63784565-001-06-2 | 10-hydroxycamptothecin | 5075 | 0.205 | 0.478 | 10539 | 0.999 | 0.999 | 4833 | 0.392 | 0.747 | 16762 | 0.427 | 0.747 | 4917 | 0.052 | 0.240 | 5529 | 0.944 | 0.999 | 8028 | 0.917 | 0.999 | 7306 | 0.656 | 0.960 | 1560 | 0.007 | 0.144 | 4712 | 0.964 | 0.999 | 3220 | 0.420 | 0.747 | 2019 | 0.073 | 0.257 | 918 | 0.515 | 0.832 | 520 | 0.017 |
| BRD-K63923597-001-10-0 | barasertib-HQPA | 1916 | 0.256 | 0.699 | 5110.5 | 0.997 | 0.997 | 1615 | 0.027 | 0.283 | 6747 | 0.240 | 1.000 | 2841.5 | 0.300 | 0.699 | 1294 | 0.706 | 0.976 | 3915.5 | 0.987 | 0.997 | 3027.5 | 0.481 | 0.841 | 93 | 0.002 | 0.007 | 1385 | 0.946 | 0.997 | 1180 | 0.013 | 0.273 | 1236 | 0.394 | 0.790 | 416 | 0.837 | 0.976 | 552 | 0.414 |
| BRD-K64052750-001-17-5 | gefitinib | 874 | 0.383 | 0.894 | 1303 | 0.989 | 0.989 | 505 | 0.006 | 0.063 | 2877 | 0.758 | 0.989 | 1312 | 0.903 | 0.989 | 863 | 0.358 | 0.894 | 1272 | 0.588 | 0.989 | 629 | 0.956 | 0.989 | 139 | 0.836 | 0.989 | 630 | 0.584 | 0.989 | 629 | 0.186 | 0.649 | 261 | 0.877 | 0.989 | 231 | 0.357 | 0.894 | 116 | 0.703 |
| BRD-K64052750-001-22-5 | gefitinib | 446 | 0.933 | 0.994 | 500 | 0.984 | 0.994 | 532 | 0.064 | 0.670 | 1433 | 0.843 | 0.994 | 774 | 0.802 | 0.994 | 331 | 0.055 | 0.192 | 471 | 0.837 | 0.994 | 360 | 0.994 | 0.994 | 170 | 0.768 | 0.994 | 229 | 0.353 | 0.824 | 114 | 0.111 | 0.775 | 215 | 0.941 | 0.994 | 92 | 0.756 | 0.994 | | |
| BRD-K64120610-001-01-4 | vinflunine | 5942 | 0.968 | 0.997 | 8113 | 0.932 | 0.997 | 3610 | 0.115 | 0.484 | 13738 | 0.051 | 0.269 | 5318 | 0.328 | 0.574 | 4330 | 0.801 | 0.997 | 6915 | 0.997 | 0.997 | 6944 | 0.734 | 0.997 | 994 | 0.002 | 0.048 | 3889 | 0.997 | 0.997 | 2151 | 0.051 | 0.269 | 2270 | 0.297 | 0.574 | 792 | 0.414 | 0.646 | 595 | 0.047 |
| BRD-K64538373-001-01-4 | Ro-4987655 | 2839 | 0.004 | 0.046 | 3307 | 0.000 | 0.000 | 3316 | 0.138 | 0.413 | 11799 | 0.953 | 1.000 | 4750 | 0.967 | 1.000 | 3623 | 0.998 | 1.000 | 3933 | 0.009 | 0.039 | 5678 | 1.000 | 1.000 | 1867 | 0.568 | 1.000 | 2101 | 0.808 | 1.000 | 2010 | 0.051 | 0.179 | 1087 | 0.950 | 1.000 | 535 | 0.862 | 1.000 | 949 | 0.553 |
| BRD-K64800655-001-07-4 | PHA-793887 | 1277.5 | 0.673 | 0.996 | 3544.5 | 0.975 | 0.996 | 1500 | 0.429 | 0.818 | 5720.5 | 0.089 | 0.311 | 2154 | 0.051 | 0.219 | 2830 | 0.996 | 0.996 | 2000.5 | 0.618 | 0.996 | 2076.5 | 0.842 | 0.996 | 589 | 0.052 | 0.219 | 714.5 | 0.881 | 0.996 | 993 | 0.018 | 0.378 | 468 | 0.131 | 0.393 | 360 | 0.383 | 0.804 | 319 | 0.273 |
| BRD-K64866502-001-03-9 | CYC116 | 963.5 | 0.741 | 0.915 | 1536.5 | 0.996 | 0.996 | 330 | 0.003 | 0.069 | 2496 | 0.680 | 0.908 | 684 | 0.125 | 0.876 | 511 | 0.835 | 0.922 | 798 | 0.198 | 0.520 | 786 | 0.692 | 0.908 | 222.5 | 0.384 | 0.897 | 767.5 | 0.644 | 0.908 | 405 | 0.014 | 0.151 | 370.5 | 0.673 | 0.908 | 27 | 0.124 | 0.371 | 168.5 | 0.518 |
| BRD-K64881305-001-03-7 | ispinesib | 1722 | 0.230 | 0.603 | 3719 | 0.947 | 0.982 | 1221 | 0.095 | 0.333 | 5877 | 0.127 | 0.382 | 2132 | 0.434 | 0.702 | 1627 | 0.969 | 0.982 | 2709 | 0.982 | 0.982 | 1849 | 0.377 | 0.668 | 287 | 0.054 | 0.225 | 1479 | 0.971 | 0.982 | 1346 | 0.348 | 0.668 | 598 | 0.017 | 0.120 | 238 | 0.382 | 0.668 | 254 | 0.034 |
| BRD-K64888243-001-03-7 | AZ960 | 3361 | 0.903 | 0.998 | 4162 | 0.980 | 0.998 | 2133 | 0.018 | 0.217 | 8823 | 0.201 | 0.591 | 3060 | 0.198 | 0.591 | 2879 | 0.998 | 0.998 | 4196 | 0.917 | 0.998 | 3457 | 0.523 | 0.733 | 1277 | 0.461 | 0.691 | 2677 | 0.958 | 0.998 | 2309 | 0.410 | 0.691 | 879 | 0.021 | 0.217 | 475 | 0.450 | 0.691 | 202 | 0.039 |
| BRD-K64890080-001-13-8 | BI-2536 | 5094 | 0.639 | 0.874 | 7620 | 0.919 | 0.984 | 4448.5 | 0.583 | 0.874 | 13254 | 0.080 | 0.829 | 4994 | 0.335 | 0.874 | 3948 | 0.874 | 0.984 | 5190 | 0.698 | 0.874 | 5920 | 0.690 | 0.874 | 2199 | 0.472 | 0.874 | 2597.5 | 0.984 | 0.984 | 2904 | 0.265 | 0.874 | 1409 | 0.118 | 0.829 | 915 | 0.649 | 0.874 | 1173 | 0.708 |
| BRD-K64925568-001-01-8 | AMG-232 | 433 | 0.101 | 1.000 | 1186 | 0.004 | 0.081 | 998 | 0.824 | 0.996 | 2979.5 | 0.980 | 0.996 | 1070 | 0.193 | 0.578 | 488 | 0.394 | 0.828 | 703.5 | 0.990 | 0.996 | 1164 | 0.352 | 0.822 | 132 | 0.004 | 0.020 | 561 | 0.005 | 0.021 | 666.5 | 0.250 | 0.655 | 93 | 0.123 | 0.432 | 329 | 0.764 | 0.996 | 81 | 0.439 |
| BRD-K65814004-003-02-9 | diphenyleneiodonium | 5199 | 0.703 | 0.907 | 7655 | 0.777 | 0.907 | 4315 | 0.508 | 0.907 | 14624 | 0.327 | 0.907 | 5297 | 0.206 | 0.721 | 4071 | 0.753 | 0.907 | 5606 | 0.366 | 0.907 | 7829 | 0.984 | 0.993 | 2206 | 0.400 | 0.907 | 4751 | 0.993 | 0.993 | 2012 | 0.056 | 0.469 | 2615 | 0.773 | 0.907 | 980 | 0.689 | 0.907 | 1037 | 0.462 |
| BRD-K65877271-001-01-8 | halobetasol-propionate | 589 | 0.263 | 0.789 | 1096.5 | 0.572 | 0.858 | 714 | 0.387 | 0.858 | 2474.5 | 0.568 | 0.858 | 949.5 | 0.140 | 0.590 | 857.5 | 0.560 | 0.858 | 1042 | 0.793 | 0.942 | 1521.5 | 0.953 | 0.953 | 300 | 0.679 | 0.892 | 294.5 | 0.301 | 0.789 | 569.5 | 0.040 | 0.843 | 476.5 | 0.667 | 0.892 | | | | 179.5 | 0.529 |
| BRD-K65924316-001-03-1 | serdemetan | 1105 | 0.956 | 0.956 | 2216 | 0.989 | 1.000 | 1258 | 0.435 | 0.937 | 4148 | 0.545 | 0.937 | 889 | 0.138 | 0.579 | 1022 | 0.728 | 0.937 | 1153 | 0.411 | 0.937 | 1546 | 0.618 | 0.937 | 342 | 0.098 | 0.516 | 1064 | 0.613 | 0.937 | 684 | 0.673 | 0.937 | 387 | 0.758 | 0.937 | 257 | 0.096 | 0.516 | 35 | 0.004 |
| BRD-K66094457-001-01-5 | CPP | 1201.5 | 0.859 | 0.953 | 1876.5 | 0.918 | 0.953 | 936 | 0.367 | 0.892 | 3734.5 | 0.781 | 0.953 | 1165.5 | 0.095 | 0.470 | 1409 | 0.728 | 0.953 | 958 | 0.064 | 0.470 | 1634.5 | 0.953 | 0.953 | 617 | 0.515 | 0.892 | 838 | 0.356 | 0.892 | 895 | 0.155 | 0.543 | 504.5 | 0.780 | 0.953 | 29 | 0.108 | 0.470 | 183 | 0.112 |
| BRD-K66097663-001-01-8 | TMC-353121 | 507 | 0.549 | 0.953 | 2778 | 0.268 | 0.849 | 919 | 0.019 | 0.205 | 5455.5 | 0.713 | 0.953 | 1866 | 0.356 | 0.849 | 1648 | 0.505 | 0.953 | 2089.5 | 0.955 | 0.955 | 2062 | 0.762 | 0.953 | 222 | 0.031 | 0.214 | 1179 | 0.680 | 0.953 | 1487 | 0.778 | 0.953 | 338 | 0.816 | 0.953 | 263 | 0.565 | 0.953 | 85 | 0.296 |
| BRD-K66175015-001-09-0 | afatinib | 1537 | 0.322 | 0.753 | 4416 | 1.000 | 1.000 | 1392 | 0.040 | 0.280 | 6682 | 0.939 | 1.000 | 2224 | 0.323 | 0.753 | 1340 | 0.222 | 0.666 | 2142 | 0.050 | 0.211 | 2783 | 1.000 | 1.000 | 672 | 0.529 | 0.926 | 1193 | 0.984 | 1.000 | 1242 | 0.128 | 0.447 | 159 | 0.629 | 0.988 | 250 | 0.440 | 0.924 | 378 | 0.845 |
| BRD-K66175015-001-12-4 | afatinib | 1702 | 0.402 | 0.845 | 3854 | 1.000 | 1.000 | 1623 | 0.004 | 0.031 | 7999 | 0.938 | 1.000 | 2565 | 0.276 | 0.644 | 942 | 0.017 | 0.060 | 2881 | 0.733 | 1.000 | 3956 | 1.000 | 1.000 | 1317 | 0.810 | 1.000 | 1492 | 0.843 | 1.000 | 1021 | 0.081 | 0.244 | 264 | 0.927 | 1.000 | 563 | 0.827 | 1.000 | 773 | 0.895 |
| BRD-K66296774-236-11-3 | fluvastatin | 4167 | 0.982 | 0.995 | 4441 | 0.008 | 0.082 | 3740 | 0.878 | 0.995 | 12768.5 | 0.348 | 0.913 | 5053 | 0.343 | 0.913 | 4377 | 0.995 | 0.995 | 5680.5 | 0.831 | 0.995 | 3501 | 0.000 | 0.005 | 1271 | 0.012 | 0.087 | 2933 | 0.743 | 0.995 | 3144 | 0.877 | 0.995 | 1731 | 0.499 | 0.995 | 919 | 0.731 | 0.995 | 1325 | 0.992 |
| BRD-K66527818-001-02-6 | VU0364770 | 1271 | 0.679 | 0.965 | 1985 | 0.303 | 0.797 | 1498 | 0.204 | 0.797 | 5050.5 | 0.275 | 0.797 | 1860.5 | 0.274 | 0.797 | 1077 | 0.538 | 0.965 | 2592 | 0.991 | 0.991 | 1581 | 0.187 | 0.797 | 587.5 | 0.476 | 0.965 | 1443 | 0.968 | 0.991 | 1253 | 0.827 | 0.965 | 131 | 0.006 | 0.067 | 390 | 0.932 | 0.991 | 51 | 0.005 |
| BRD-K66788707-001-14-3 | fludarabine | 612 | 0.853 | 1.000 | 773 | 0.078 | 0.818 | 579 | 0.163 | 0.854 | 1805 | 0.027 | 0.576 | 1096 | 0.247 | 1.000 | 507 | 0.562 | 1.000 | 1019 | 0.682 | 1.000 | 1312 | 0.903 | 0.952 | 519 | 0.845 | 1.000 | 434 | 0.590 | 1.000 | 374 | 0.677 | 1.000 | 285 | 0.668 | 1.000 | 116 | 0.747 | 1.000 | 95 | 0.149 |
| BRD-K66808046-065-03-7 | oxyquinoline | 2767 | 0.690 | 0.940 | 5032 | 0.582 | 0.940 | 3253 | 0.530 | 0.940 | 12179 | 0.940 | 0.940 | 3585 | 0.289 | 0.607 | 3335 | 0.740 | 0.940 | 4810 | 0.746 | 0.940 | 4970 | 0.901 | 0.940 | 721 | 0.092 | 0.465 | 2021 | 0.631 | 0.940 | 2189 | 0.164 | 0.545 | 977 | 0.027 | 0.275 | 546 | 0.216 | 0.568 | 177 | 0.111 |
| BRD-K67261995-001-11-7 | iodipamide | 912 | 0.366 | 0.716 | 1975 | 0.162 | 1.000 | 983 | 0.101 | 1.000 | 3663 | 0.507 | 0.716 | 1746 | 0.927 | 0.927 | 953 | 0.843 | 0.927 | 1561 | 0.731 | 0.904 | 1226 | 0.459 | 0.716 | 975 | 0.873 | 0.927 | 445 | 0.919 | 0.927 | 792 | 0.511 | 0.716 | 249 | 0.300 | 0.700 | | | | 63 | 0.258 |
| BRD-K67298865-001-09-0 | SB-431542 | 1674 | 0.043 | 0.621 | 4137 | 0.566 | 0.897 | 2419 | 0.807 | 0.897 | 10078 | 0.897 | 0.897 | 2664 | 0.214 | 0.640 | 2312 | 0.546 | 0.897 | 3083 | 0.734 | 0.897 | 3267 | 0.120 | 0.628 | 1040 | 0.894 | 0.897 | 1824 | 0.183 | 0.640 | 1573 | 0.200 | 0.640 | 918 | 0.244 | 0.640 | 262 | 0.346 | 0.805 | 834 | 0.865 |
| BRD-K67511046-001-02-3 | naloxone | 568 | 0.179 | 1.000 | 1595 | 0.551 | 1.000 | 1043 | 0.526 | 1.000 | 2341 | 0.096 | 1.000 | 374 | 0.267 | 1.000 | 1132 | 0.855 | 0.897 | 892 | 0.217 | 1.000 | 1332 | 0.954 | 0.954 | 494 | 0.379 | 1.000 | 598 | 0.692 | 0.897 | 562 | 0.836 | 0.897 | 554 | 0.563 | 0.844 | 134 | 0.263 | 0.502 | 165 | 0.413 |
| BRD-K67536197-003-11-4 | cyclocytidine | 2652 | 0.047 | 0.997 | 6027 | 0.859 | 0.958 | 3038 | 0.607 | 0.950 | 10238 | 0.164 | 1.000 | 3427 | 0.263 | 1.000 | 3280 | 0.958 | 0.958 | 4182 | 0.368 | 1.000 | 4696 | 0.953 | 0.958 | 1512 | 0.228 | 0.479 | 3017 | 0.857 | 0.958 | 2733 | 0.787 | 0.958 | 747 | 0.011 | 0.027 | 670 | 0.771 | 0.958 | 775 | 0.311 |
| BRD-K67566344-001-07-5 | ku-0063794 | 3252.5 | 0.965 | 0.999 | 5524 | 0.740 | 0.864 | 2563 | 0.227 | 0.680 | 10869.5 | 0.632 | 0.864 | 3416 | 0.044 | 0.307 | 2037.5 | 0.094 | 0.494 | 4219.5 | 0.342 | 0.697 | 5425.5 | 0.999 | 0.999 | 1388.5 | 0.147 | 0.514 | 2760 | 0.706 | 0.864 | 1307 | 0.041 | 0.307 | 622 | 0.134 | 0.514 | 572 | 0.263 | 0.690 | 318 | 0.406 |
| BRD-K67772619-001-05-1 | rheochrysidin | 516 | 0.622 | 0.829 | 1407 | 0.040 | 0.420 | 966 | 0.390 | 1.000 | 3728 | 0.909 | 0.919 | 1626 | 0.804 | 0.894 | 576 | 0.282 | 1.000 | 1055 | 0.399 | 1.000 | 1886 | 0.809 | 0.894 | 536 | 0.681 | 0.842 | 234 | 0.286 | 0.547 | 608 | 0.611 | 1.000 | 203 | 0.547 | 0.829 | 213 | 0.597 | 0.829 | 104 | 0.029 |
| BRD-K67844266-003-01-9 | pevonedistat | 2684 | 0.988 | 0.988 | 3455 | 0.905 | 0.988 | 1317 | 0.141 | 0.353 | 6163 | 0.134 | 0.353 | 2052 | 0.029 | 0.153 | 2185 | 0.981 | 0.988 | 3057 | 0.981 | 0.988 | 2295 | 0.711 | 0.988 | 1196 | 0.454 | 0.867 | 1251 | 0.760 | 0.988 | 1164 | 0.112 | 1.000 | 740 | 0.151 | 0.353 | 111 | 0.072 | 0.300 | 153 | 0.566 |
| BRD-K67868012-001-07-6 | PI-103 | 1664 | 0.521 | 0.837 | 4576 | 0.989 | 0.989 | 1709 | 0.253 | 1.000 | 6140 | 0.058 | 1.000 | 2994 | 0.457 | 1.000 | 2111 | 0.746 | 0.922 | 2799 | 0.936 | 0.989 | 2736 | 0.961 | 0.989 | 752 | 0.151 | 0.351 | 1225 | 0.459 | 0.837 | 1170 | 0.099 | 1.000 | 181 | 0.008 | 0.023 | 284 | 0.498 | 0.837 | 445 | 0.240 |
| BRD-K68065987-300-09-1 | MK-2206 | 4307 | 0.803 | 0.992 | 5426.5 | 0.052 | 0.379 | 3610 | 0.200 | 0.466 | 14862 | 0.801 | 0.992 | 3522 | 0.092 | 0.379 | 3716 | 0.382 | 0.730 | 6858 | 0.998 | 0.998 | 6357.5 | 0.800 | 0.992 | 723 | 0.000 | 0.006 | 3902 | 0.984 | 0.998 | 2256 | 0.126 | 0.379 | 2329 | 0.939 | 0.998 | 509 | 0.103 | 0.379 | 838 | 0.555 |
| BRD-K68103045-001-02-9 | CGS-20625 | 506 | 0.841 | 0.981 | 761.5 | 0.624 | 0.883 | 423.5 | 0.324 | 0.618 | 1590 | 0.204 | 0.618 | 615 | 0.114 | 0.598 | 161 | 0.079 | 0.598 | 845 | 0.981 | 0.981 | 802.5 | 0.736 | 0.965 | 224 | 0.263 | 0.618 | 175 | 0.320 | 0.618 | 123 | 0.114 | 0.598 | 109 | 0.299 | 0.618 | 240 | 0.959 | 0.981 | 106 | 0.281 |
| BRD-K68164687-001-01-6 | epothilone-b | 5214 | 0.534 | 0.883 | 9488 | 0.994 | 0.995 | 3077 | 0.011 | 0.226 | 14223 | 0.134 | 0.422 | 4681 | 0.141 | 0.422 | 5621 | 0.995 | 0.995 | 6683 | 0.607 | 0.891 | 7007 | 0.679 | 0.891 | 2026 | 0.387 | 0.850 | 4414 | 0.995 | 0.995 | 2786 | 0.212 | 0.555 | 2451 | 0.405 | 0.850 | 898 | 0.546 | 0.883 | 771 | 0.139 |
| BRD-K68174511-001-06-6 | torin-2 | 3988 | 0.981 | 0.993 | 6845 | 0.993 | 0.993 | 2923 | 0.364 | 0.650 | 12524 | 0.819 | 0.955 | 3537 | 0.082 | 0.389 | 3267 | 0.372 | 0.650 | 6279 | 0.965 | 0.993 | 5056 | 0.487 | 0.736 | 1303 | 0.133 | 0.465 | 2304 | 0.267 | 0.560 | 1809 | 0.032 | 0.670 | 1074 | 0.093 | 0.389 | 371 | 0.162 | 0.486 | 893 | 0.738 |
| BRD-K68202742-001-16-5 | trichostatin-a | 2906 | 0.019 | 0.137 | 5749 | 0.413 | 0.722 | 4288 | 0.725 | 0.896 | 12426 | 0.298 | 0.626 | 4596 | 0.225 | 0.590 | 4396 | 0.977 | 1.000 | 7332 | 1.000 | 1.000 | 6766 | 0.955 | 1.000 | 1908 | 0.454 | 0.734 | 4350 | 0.998 | 1.000 | 1730 | 0.028 | 0.146 | 1280 | 0.081 | 0.341 | 852 | 0.600 | 0.882 | 464 | 0.020 |
| BRD-K68336408-001-10-9 | tyrphostin-AG-1478 | 3579 | 0.634 | 0.951 | 6499.5 | 0.945 | 1.000 | 2048 | 0.004 | 0.043 | 12608 | 0.293 | 0.631 | 4197 | 0.334 | 0.631 | 3643 | 0.808 | 1.000 | 5566.5 | 0.998 | 1.000 | 6422 | 1.000 | 1.000 | 1576 | 0.830 | 1.000 | 3150.5 | 0.931 | 1.000 | 2052 | 0.200 | 0.525 | 689 | 0.391 | 0.631 | 243 | 0.045 | 0.236 | 679 | 0.373 |

| broad_id | name | al_nervous_system_PVAL_ADJ | fibroblast_MWU_STAT | fibroblast_PVAL | fibroblast_PVAL_ADJ | bile_duct_MWU_STAT | bile_duct_PVAL | bile_duct_PVAL_ADJ | thyroid_MWU_STAT | thyroid_PVAL | thyroid_PVAL_ADJ | upper_aerodigestive_MWU_STAT | upper_aerodigestive_PVAL | upper_aerodigestive_PVAL_ADJ | uterus_MWU_STAT | uterus_PVAL | uterus_PVAL_ADJ | liver_MWU_STAT | liver_PVAL | liver_PVAL_ADJ | esophagus_MWU_STAT | esophagus_PVAL | esophagus_PVAL_ADJ |
|---|---|---|---|---|---|---|---|---|---|---|---|---|---|---|---|---|---|---|---|---|---|---|---|
| BRD-K56981171-001-02-8 | brigatinib | 0.822 | | | | 36 | 0.180 | 1.000 | 160 | 0.534 | 0.822 | 1145 | 0.401 | 0.799 | 387 | 0.274 | 0.720 | 345 | 0.670 | 0.828 | 743 | 0.202 | 1.000 |
| BRD-K57080016-001-15-9 | selumetinib | 0.983 | 150.5 | 0.869 | 0.983 | 186 | 0.164 | 0.689 | 397 | 0.627 | 0.983 | 742 | 0.018 | 0.123 | 968.5 | 0.962 | 0.983 | 671 | 0.971 | 0.983 | 271 | 0.487 | 0.983 |
| BRD-K57136142-001-02-4 | CCT129202 | 0.277 | | | | 216 | 0.417 | 0.673 | 523 | 0.344 | 0.656 | 1277 | 0.060 | 0.277 | 1322 | 0.084 | 0.294 | 470 | 0.227 | 0.477 | 1995 | 0.985 | 0.985 |
| BRD-K57169635-001-04-5 | dacomitinib | 0.916 | | | | 178 | 0.193 | 0.507 | 1373 | 0.996 | 1.000 | 992 | 0.000 | 0.000 | 1223 | 0.249 | 0.552 | 1107 | 0.263 | 0.552 | 1280 | 0.000 | 0.004 |
| BRD-K57284066-001-01-6 | Ro-9187 | 0.781 | 37 | 0.164 | 0.687 | 320.5 | 0.766 | 0.902 | 580.5 | 0.919 | 0.919 | 741 | 0.485 | 0.781 | 305 | 0.142 | 0.687 | 240 | 0.155 | 0.687 | 1023.5 | 0.773 | 0.902 |
| BRD-K58049875-001-04-7 | thiostrepton | 0.981 | 19 | 0.072 | 0.488 | 360 | 0.856 | 0.981 | 635 | 0.981 | 0.981 | 1232 | 0.101 | 0.488 | 949 | 0.121 | 0.488 | 497 | 0.028 | 0.488 | 1521 | 0.169 | 0.507 |
| BRD-K58122457-001-02-2 | hesperadin | 0.638 | 113 | 0.295 | 0.688 | 1021 | 0.973 | 0.991 | 1161 | 0.544 | 0.991 | 1760 | 0.046 | 0.242 | 2361 | 0.658 | 0.991 | 2176.5 | 0.909 | 0.991 | 3360 | 0.598 | 0.991 |
| BRD-K58148589-001-03-6 | betamethasone-dipropionate | 0.889 | | | | 153 | 0.529 | 1.000 | 557 | 0.881 | 0.889 | 589 | 0.191 | 0.990 | 608 | 0.825 | 0.889 | 415 | 0.702 | 0.889 | 792 | 0.889 | 0.889 |
| BRD-K58262659-001-09-7 | doxifluridine | 0.945 | | | | 15 | 0.096 | 1.000 | 125 | 0.481 | 0.842 | 299 | 0.437 | 0.835 | 206 | 0.598 | 0.918 | 489.5 | 0.730 | 0.918 | 343.5 | 0.378 | 1.000 |
| BRD-K58435339-001-03-0 | AT13387 | 0.601 | 301 | 0.914 | 0.986 | 1095 | 0.986 | 0.986 | 1609 | 0.672 | 0.986 | 2857 | 0.338 | 0.601 | 2123 | 0.343 | 0.601 | 1672.5 | 0.197 | 0.601 | 2058 | 0.044 | 0.462 |
| BRD-K58466253-003-14-0 | palmatine-chloride | 0.940 | | | | 286 | 0.827 | 0.940 | 77 | 0.089 | 0.469 | 422 | 0.245 | 0.784 | 373 | 0.261 | 0.784 | 216 | 0.792 | 0.940 | 563 | 0.202 | 0.784 |
| BRD-K58529924-001-01-5 | ONC201 | 0.971 | 142.5 | 0.925 | 0.971 | 65 | 0.391 | 0.821 | 330.5 | 0.901 | 0.971 | 467.5 | 0.313 | 0.744 | 529.5 | 0.760 | 0.971 | 421.5 | 0.190 | 0.744 | 935.5 | 0.997 | 0.997 |
| BRD-K58550667-001-08-7 | FK-866 | 0.139 | 29 | 0.073 | 0.263 | 820 | 0.342 | 0.654 | 1742 | 0.634 | 0.951 | 2823 | 0.327 | 0.654 | 2394 | 0.075 | 0.263 | 2123 | 0.043 | 0.223 | 3382 | 0.263 | 0.615 |
| BRD-K58770988-001-01-8 | NMS-E973 | 0.831 | 22 | 0.053 | 0.223 | 1522 | 0.946 | 0.999 | 1272 | 0.015 | 0.107 | 3445 | 0.102 | 0.307 | 2551 | 0.011 | 0.107 | 2953 | 0.512 | 0.827 | 5036 | 0.577 | 0.831 |
| BRD-K58772419-001-07-0 | AZD6482 | 0.518 | 331 | 0.802 | 0.993 | 1563 | 0.875 | 0.993 | 1593 | 0.259 | 0.877 | 3353 | 0.074 | 0.518 | 3700 | 0.608 | 0.993 | 2894 | 0.751 | 0.993 | 5241 | 0.994 | 0.994 |
| BRD-K59146805-001-03-7 | AZ3146 | 0.763 | | | | 721 | 0.983 | 0.983 | 836 | 0.974 | 0.983 | 624 | 0.012 | 0.245 | 1045 | 0.423 | 0.763 | 766 | 0.133 | 0.350 | 1475 | 0.201 | 1.000 |
| BRD-K59317601-001-05-5 | MLN0128 | 0.761 | 181 | 0.532 | 0.734 | 527 | 0.523 | 0.734 | 1427 | 0.950 | 1.000 | 2714 | 0.559 | 0.734 | 2762 | 0.309 | 0.734 | 1715 | 0.020 | 0.211 | 2822 | 0.509 | 0.734 |
| BRD-K59325863-001-02-8 | delanzomib | 0.850 | 128 | 0.284 | 0.844 | 1934 | 0.852 | 0.973 | 1297 | 0.093 | 0.491 | 5469 | 0.880 | 0.973 | 4951 | 0.852 | 0.973 | 4708 | 0.990 | 0.990 | 4454 | 0.304 | 0.844 |
| BRD-K59369769-001-20-3 | tozasertib | 0.239 | 73 | 0.187 | 0.590 | 1326 | 0.912 | 0.997 | 1890 | 0.670 | 0.997 | 3602 | 0.256 | 0.590 | 2135 | 0.048 | 0.254 | 2393 | 0.591 | 0.992 | 4745 | 0.887 | 0.997 |
| BRD-K59369769-001-22-9 | tozasertib | 0.305 | 78 | 0.483 | 0.922 | 208 | 0.771 | 0.981 | 210 | 0.363 | 0.846 | 333 | 0.114 | 0.399 | 47 | 0.042 | 0.298 | 432 | 0.652 | 0.981 | 640 | 0.793 | 0.981 |
| BRD-K59436580-001-01-7 | tedizolid | 0.450 | | | | 311.5 | 0.803 | 0.955 | 363 | 0.026 | 0.270 | 1208 | 0.303 | 0.908 | 1329.5 | 0.849 | 0.955 | 1054 | 0.593 | 1.000 | 1831 | 0.643 | 0.900 |
| BRD-K59753975-001-02-6 | vindesine | 0.626 | 67 | 0.121 | 0.362 | 1518 | 0.826 | 0.963 | 1185 | 0.033 | 0.171 | 2242 | 0.003 | 0.033 | 2356 | 0.006 | 0.043 | 2743 | 0.590 | 0.885 | 4369 | 0.757 | 0.935 |
| BRD-K59831625-001-01-6 | LDN-212854 | 0.617 | 8 | 0.022 | 0.457 | 1641 | 0.751 | 0.901 | 1710 | 0.376 | 0.617 | 4033 | 0.571 | 0.857 | 3431 | 0.264 | 0.617 | 2947 | 0.307 | 0.617 | 4524 | 0.772 | 0.901 |
| BRD-K59866480-001-07-3 | cyclovalone | 0.249 | 35 | 0.088 | 0.308 | 424 | 0.051 | 0.249 | 1577 | 0.469 | 0.767 | 3168 | 0.043 | 0.249 | 3673 | 0.475 | 0.767 | 3112 | 0.788 | 0.919 | 3641 | 0.127 | 0.381 |
| BRD-K59983611-001-21-9 | sulfabenzamide | 1.000 | 162.5 | 0.945 | 0.945 | | | | 305 | 0.393 | 1.000 | 542 | 0.044 | 0.462 | 463 | 0.201 | 0.850 | 702.5 | 0.851 | 0.945 | 915 | 0.778 | 1.000 |
| BRD-K60184833-001-05-6 | tyrphostin-AG-99 | 0.742 | 163 | 0.480 | 0.742 | 886 | 0.598 | 0.838 | 1070 | 0.365 | 0.742 | 3140 | 0.931 | 0.977 | 1701 | 0.011 | 0.118 | 1197 | 0.011 | 0.118 | 2411.5 | 0.489 | 0.742 |
| BRD-K60230970-001-17-5 | MG-132 | 0.669 | 128 | 0.287 | 0.669 | 1578 | 0.777 | 0.876 | 1642 | 0.085 | 0.436 | 5488 | 0.955 | 0.955 | 3726 | 0.091 | 0.436 | 2366 | 0.076 | 0.436 | 4160 | 0.277 | 0.669 |
| BRD-K60341624-001-02-2 | GSK1292263 | | | | | 197 | 0.410 | 0.851 | 161 | 0.016 | 0.336 | 1271 | 0.230 | 0.804 | 1206 | 0.082 | 0.523 | 635 | 0.115 | 0.523 | 1982 | 0.990 | 0.990 |
| BRD-K60348325-065-09-9 | brilliant-green | 0.947 | 342 | 0.780 | 0.999 | 1920 | 0.885 | 0.999 | 1708 | 0.133 | 0.471 | 3359 | 0.017 | 0.174 | 4137 | 0.600 | 0.947 | 3304 | 0.596 | 0.947 | 2971 | 0.006 | 0.126 |
| BRD-K60443845-001-08-6 | chlormidazole | 0.960 | | | | 716.5 | 0.977 | 0.977 | 998 | 0.521 | 0.960 | 2873 | 0.897 | 0.960 | 1843 | 0.739 | 0.960 | 719 | 0.014 | 0.100 | 1722 | 0.404 | 1.000 |
| BRD-K60446698-001-01-2 | ONX-0914 | 0.916 | 246 | 0.564 | 0.916 | 1430 | 0.888 | 0.933 | 1774 | 0.339 | 0.916 | 3159 | 0.064 | 0.449 | 3936 | 0.468 | 0.916 | 3320 | 0.775 | 0.933 | 4940 | 0.612 | 0.916 |
| BRD-K60997853-001-02-3 | PHA-848125 | 0.483 | 169 | 0.500 | 0.930 | 467 | 0.148 | 0.490 | 1255 | 0.856 | 1.000 | 2410 | 0.115 | 0.483 | 2776 | 0.531 | 0.930 | 2171 | 0.729 | 1.000 | 2698 | 0.453 | 0.930 |
| BRD-K61097567-001-08-5 | SB-218795 | 0.112 | | | | | | | 185.5 | 0.500 | 1.000 | 342.5 | 0.206 | 1.000 | 228 | 0.062 | 0.703 | 237.5 | 0.456 | 1.000 | 287.5 | 0.416 | 1.000 |
| BRD-K61195623-001-01-4 | combretastatin-A-4 | 0.308 | 112 | 0.223 | 0.586 | 1856.5 | 0.991 | 0.999 | 1088 | 0.066 | 0.346 | 4002 | 0.493 | 0.810 | 3075 | 0.331 | 0.772 | 2859.5 | 0.455 | 0.810 | 4778 | 0.774 | 0.956 |
| BRD-K61228301-001-01-2 | SGI-1027 | 0.984 | | | | 576 | 0.712 | 0.994 | 1710 | 0.994 | 0.994 | 2085 | 0.150 | 0.450 | 2196 | 0.136 | 0.450 | 1600 | 0.542 | 0.948 | 2322 | 0.132 | 0.450 |
| BRD-K61314889-001-04-4 | endo-IWR-1 | 0.999 | | | | 214 | 0.806 | 0.999 | 637 | 0.357 | 0.826 | 1614 | 0.164 | 0.826 | 1415 | 0.202 | 0.826 | 1449 | 0.599 | 0.967 | 3418 | 1.000 | 1.000 |
| BRD-K61323504-001-04-0 | SB-225002 | 0.091 | | | | 77 | 0.203 | 0.710 | 766 | 0.560 | 0.979 | 2665 | 0.496 | 0.979 | 1121 | 0.008 | 0.079 | 953 | 0.375 | 0.979 | 1945 | 0.279 | 0.836 |
| BRD-K61397605-001-03-4 | romidepsin | 0.981 | 60 | 0.134 | 0.353 | 1568 | 0.754 | 0.999 | 1729 | 0.116 | 0.353 | 2588 | 0.001 | 0.024 | 3818 | 0.182 | 0.424 | 3653 | 0.765 | 0.999 | 4107 | 0.083 | 0.353 |
| BRD-K61443650-236-13-1 | thiomersal | 0.560 | | | | 419 | 0.289 | 0.560 | 1045 | 0.381 | 0.616 | 1601 | 0.709 | 0.915 | 1360 | 0.200 | 0.560 | 516 | 0.099 | 0.416 | 1591 | 0.845 | 0.986 |
| BRD-K61496577-001-20-2 | flumethasone | 0.880 | | | | | | | 420 | 0.308 | 1.000 | 894 | 0.167 | 0.701 | 546 | 0.731 | 0.959 | 551 | 0.495 | 1.000 | 547 | 0.734 | 0.734 |
| BRD-K61536336-001-01-7 | SB-366791 | 0.678 | | | | 270 | 0.979 | 1.000 | 165 | 0.235 | 1.000 | 686 | 0.674 | 0.884 | 157 | 0.016 | 0.344 | 222 | 0.519 | 0.778 | 793.5 | 0.904 | 0.904 |
| BRD-K61734568-003-02-1 | 2,3-DCPE | 0.787 | 22 | 0.107 | 0.748 | 270.5 | 0.951 | 0.966 | 283 | 0.356 | 0.787 | 494 | 0.337 | 0.787 | 306 | 0.002 | 0.052 | 534 | 0.910 | 0.966 | 826 | 0.834 | 0.966 |
| BRD-K61936403-001-01-7 | AS-1949490 | 0.703 | | | | 221 | 0.956 | 0.991 | 347 | 0.888 | 1.000 | 306 | 0.092 | 0.642 | 391 | 0.320 | 0.961 | 155 | 0.683 | 1.000 | 362 | 0.433 | 1.000 |
| BRD-K62008436-001-22-1 | paclitaxel | 1.000 | 36 | 0.073 | 0.271 | 2041 | 0.900 | 1.000 | 2023 | 0.286 | 0.600 | 3639 | 0.025 | 0.130 | 3839 | 0.093 | 0.279 | 4149.5 | 0.866 | 1.000 | 3458 | 0.012 | 0.130 |
| BRD-K62008436-001-23-9 | paclitaxel | 0.800 | 45 | 0.136 | 0.716 | 352 | 0.442 | 0.800 | 977 | 0.519 | 0.800 | 918 | 0.045 | 0.476 | 1600 | 0.422 | 0.800 | 1433 | 0.950 | 1.000 | 1218 | 0.102 | 0.714 |
| BRD-K62196610-001-01-6 | AVN-944 | 0.116 | | | | 393.5 | 0.936 | 0.996 | | | | 413 | 0.208 | 1.000 | 626.5 | 0.364 | 0.637 | 358 | 0.104 | 0.272 | 579 | 0.021 | 0.434 |
| BRD-K62304154-001-04-2 | deforolimus | 0.998 | 304 | 0.780 | 0.998 | 1742 | 0.952 | 0.998 | 2441 | 0.842 | 0.998 | 4047 | 0.382 | 0.730 | 2996 | 0.103 | 0.319 | 2392 | 0.037 | 0.308 | 3123 | 0.023 | 0.308 |
| BRD-K62310379-001-10-5 | fluticasone-propionate | 1.000 | | | | 626 | 0.883 | 1.000 | 100 | 0.006 | 0.122 | 811 | 0.428 | 0.855 | 923 | 0.304 | 0.855 | 613 | 0.189 | 0.794 | 1203 | 0.846 | 0.846 |
| BRD-K62374002-001-01-5 | PI3K-IN-2 | 0.898 | 33 | 0.077 | 0.270 | 1216 | 0.394 | 0.888 | 2557 | 0.949 | 1.000 | 3277 | 0.048 | 0.254 | 4157 | 0.470 | 0.896 | 2596 | 0.026 | 0.254 | 5064 | 0.789 | 0.974 |
| BRD-K62627508-001-01-5 | idasanutlin | 0.693 | | | | 270.5 | 0.763 | 1.000 | 709.5 | 0.926 | 1.000 | 933.5 | 0.985 | 1.000 | 1035 | 0.750 | 1.000 | 774.5 | 0.637 | 1.000 | 730.5 | 0.946 | 0.946 |
| BRD-K62696033-001-01-4 | G-1 | 0.371 | | | | 78 | 0.547 | 1.000 | 173 | 0.087 | 0.457 | 381.5 | 0.603 | 1.000 | 313 | 0.136 | 0.571 | 205.5 | 0.165 | 0.578 | 344.5 | 0.034 | 0.289 |
| BRD-K62779383-001-05-5 | cholecalciferol | 0.769 | | | | 240 | 0.303 | 0.769 | 193 | 0.493 | 0.850 | 1283 | 0.505 | 0.850 | 610.5 | 0.358 | 0.769 | 393 | 0.246 | 0.769 | 1199 | 0.934 | 0.934 |
| BRD-K62814476-001-03-3 | BMS-265246 | 0.941 | | | | 726 | 0.564 | 0.941 | 2162 | 0.997 | 1.000 | 3189 | 0.805 | 0.994 | 2647 | 0.630 | 0.945 | 1617 | 0.053 | 0.224 | 2562 | 0.150 | 0.525 |
| BRD-K62965247-001-04-9 | tipifarnib | 0.966 | 287 | 0.807 | 0.971 | 878 | 0.390 | 0.877 | 1647 | 0.678 | 0.971 | 3302 | 0.152 | 0.565 | 3186 | 0.560 | 0.966 | 2457 | 0.231 | 0.693 | 4645 | 0.905 | 0.971 |
| BRD-K63068307-001-08-9 | ZSTK-474 | 0.874 | | | | 570 | 0.919 | 1.000 | 980.5 | 0.978 | 0.987 | 1879 | 0.251 | 0.642 | 1292.5 | 0.755 | 1.000 | 658 | 0.275 | 0.642 | 2655 | 0.931 | 1.000 |
| BRD-K63126190-003-01-6 | epinephrine | 0.995 | 136 | 0.888 | 0.995 | 156 | 0.950 | 0.995 | 405 | 0.298 | 0.995 | 889 | 0.935 | 0.995 | 351 | 0.060 | 0.311 | 193 | 0.028 | 0.311 | 654 | 0.062 | 0.311 |
| BRD-K63504947-001-15-4 | semaxanib | 0.302 | | | | 572 | 0.905 | 1.000 | 556 | 0.883 | 1.000 | 1265 | 0.459 | 1.000 | 1354 | 0.407 | 1.000 | 938 | 0.072 | 0.302 | 1953 | 0.944 | 1.000 |
| BRD-K63736853-001-04-8 | acipimox | | | | | 200 | 0.881 | 1.000 | 319 | 0.219 | 0.826 | 351 | 0.025 | 0.264 | 304 | 0.384 | 1.000 | 2 | 0.048 | 0.337 | 893 | 0.864 | 1.000 |
| BRD-K63750851-001-20-7 | mycophenolic-acid | 0.581 | | | | 88 | 0.328 | 0.766 | 617 | 0.698 | 0.911 | 1503 | 0.425 | 0.811 | 1698 | 0.854 | 0.930 | 1495 | 0.602 | 0.911 | 1680 | 0.032 | 0.665 |
| BRD-K63784565-001-06-2 | 10-hydroxycamptothecin | 0.182 | 82 | 0.181 | 0.476 | 2116 | 0.938 | 0.999 | 1581 | 0.056 | 0.240 | 3892 | 0.120 | 0.360 | 3652 | 0.057 | 0.240 | 3875 | 0.743 | 0.975 | 5088 | 0.685 | 0.960 |
| BRD-K63923597-001-10-0 | barasertib-HQPA | 0.790 | | | | 730 | 0.793 | 0.976 | 850 | 0.727 | 0.976 | 1977 | 0.197 | 1.000 | 2080 | 0.293 | 0.699 | 1491 | 0.569 | 0.919 | 2418 | 0.812 | 0.976 |
| BRD-K64052750-001-17-5 | gefitinib | 0.989 | | | | 304 | 0.675 | 0.989 | 346 | 0.823 | 0.989 | 289 | 0.001 | 0.018 | 1107 | 0.865 | 0.989 | 205 | 0.079 | 0.333 | 704 | 0.094 | 0.656 |
| BRD-K64052750-001-22-5 | gefitinib | | | | | 192 | 0.485 | 0.994 | 241 | 0.980 | 0.994 | 552 | 0.001 | 0.015 | 190 | 0.186 | 0.489 | 510 | 0.759 | 0.994 | 700 | 0.165 | 0.489 |
| BRD-K64120610-001-01-4 | vinflunine | 0.269 | 131 | 0.308 | 0.574 | 2311 | 0.996 | 0.997 | 1880 | 0.294 | 0.574 | 3447 | 0.203 | 0.574 | 3788 | 0.295 | 0.574 | 2329 | 0.201 | 0.574 | 4176 | 0.430 | 0.646 |
| BRD-K64538373-001-01-4 | Ro-4987655 | 1.000 | | | | 1302 | 0.783 | 1.000 | 1251 | 0.463 | 1.000 | 2256 | 0.008 | 0.057 | 2693 | 0.828 | 1.000 | 3587 | 0.961 | 1.000 | 4663 | 0.952 | 1.000 |
| BRD-K64800655-001-07-4 | PHA-793887 | 0.638 | | | | 516 | 0.806 | 0.996 | 1448.5 | 0.967 | 0.996 | 2195.5 | 0.706 | 0.996 | 1290 | 0.165 | 1.000 | 1260 | 0.836 | 0.996 | 1791 | 0.214 | 0.561 |
| BRD-K64866502-001-03-9 | CYC116 | 0.906 | | | | 221 | 0.792 | 0.922 | 376.5 | 0.388 | 1.000 | 377 | 0.390 | 1.000 | 727 | 0.904 | 0.949 | 160 | 0.468 | 0.906 | 930.5 | 0.488 | 0.906 |
| BRD-K64881305-001-03-7 | ispinesib | 0.180 | | | | 836 | 0.971 | 0.982 | 841 | 0.326 | 0.668 | 2175 | 0.642 | 0.899 | 1954 | 0.516 | 0.774 | 1608 | 0.880 | 0.982 | 1705 | 0.213 | 1.000 |
| BRD-K64888243-001-03-7 | AZ960 | 0.271 | 105 | 0.263 | 0.615 | 1250 | 0.977 | 0.998 | 1085 | 0.408 | 0.691 | 2401 | 0.225 | 0.591 | 1787 | 0.191 | 0.591 | 1488 | 0.180 | 0.591 | 2991 | 0.361 | 0.691 |
| BRD-K64890080-001-13-8 | BI-2536 | 0.874 | 412.5 | 0.956 | 0.984 | 874 | 0.584 | 0.874 | 1484 | 0.331 | 0.874 | 3860 | 0.296 | 0.874 | 2954.5 | 0.028 | 0.582 | 3040 | 0.529 | 0.874 | 4650 | 0.592 | 0.874 |
| BRD-K64925568-001-01-8 | AMG-232 | 0.837 | | | | 160.5 | 0.917 | 0.996 | 623.5 | 0.949 | 0.996 | 609 | 0.934 | 0.996 | 805 | 0.594 | 0.996 | 553.5 | 0.846 | 0.996 | 1174 | 0.996 | 0.996 |
| BRD-K65814004-003-02-9 | diphenyleneiodonium | 0.907 | 287 | 0.668 | 0.907 | 1727 | 0.768 | 0.907 | 2582 | 0.890 | 0.984 | 2859 | 0.011 | 0.226 | 3118 | 0.067 | 0.469 | 2655 | 0.164 | 0.687 | 3753 | 0.161 | 0.687 |
| BRD-K65877271-001-01-8 | halobetasol-propionate | 0.858 | | | | 139.5 | 0.855 | 0.942 | 553 | 0.872 | 0.942 | 998 | 0.211 | 0.737 | 496 | 0.445 | 0.858 | 756.5 | 0.897 | 0.942 | 352.5 | 0.089 | 0.467 |
| BRD-K65924316-001-03-1 | serdemetan | 0.088 | | | | 625 | 0.958 | 1.000 | 359 | 0.321 | 0.937 | 1037 | 0.216 | 0.754 | 870 | 0.626 | 0.937 | 427 | 0.011 | 0.118 | 911 | 0.496 | 0.937 |
| BRD-K66094457-001-01-5 | CPP | 0.470 | 111 | 0.546 | 0.892 | 456.5 | 0.920 | 0.953 | 409 | 0.492 | 0.892 | 914.5 | 0.019 | 0.392 | 829.5 | 0.552 | 0.892 | 793.5 | 0.701 | 0.953 | 788 | 0.454 | 0.892 |
| BRD-K66097663-001-01-8 | TMC-353121 | 0.849 | 31 | 0.099 | 0.520 | 513 | 0.878 | 0.955 | 781 | 0.364 | 0.849 | 1123.5 | 0.010 | 0.205 | 1527.5 | 0.327 | 0.849 | 1102.5 | 0.756 | 0.953 | 2207.5 | 0.953 | 0.955 |
| BRD-K66175015-001-09-0 | afatinib | 1.000 | | | | 659 | 0.799 | 1.000 | 1100 | 0.951 | 1.000 | 803 | 0.000 | 0.000 | 1296 | 0.502 | 0.926 | 1649 | 0.658 | 0.988 | 1117 | 0.000 | 0.001 |
| BRD-K66175015-001-12-4 | afatinib | 1.000 | | | | 821 | 0.736 | 1.000 | 526 | 0.753 | 1.000 | 1309 | 0.001 | 0.012 | 1420 | 0.055 | 0.286 | 809 | 0.097 | 0.254 | 1317 | 0.002 | 0.018 |
| BRD-K66296774-236-11-3 | fluvastatin | 0.995 | 8 | 0.049 | 0.258 | 674 | 0.321 | 0.913 | 1548 | 0.148 | 0.622 | 4465 | 0.860 | 0.995 | 2987 | 0.631 | 0.995 | 2748 | 0.584 | 0.995 | 3323 | 0.749 | 0.995 |
| BRD-K66527818-001-02-6 | VU0364770 | 0.067 | 166 | 0.739 | 0.965 | 207 | 0.364 | 0.850 | 628 | 0.579 | 0.965 | 1219 | 0.127 | 0.797 | 1342.5 | 0.786 | 0.965 | 1172.5 | 0.711 | 0.965 | 1461.5 | 0.785 | 0.965 |
| BRD-K66788707-001-14-3 | fludarabine | 0.854 | | | | 264 | 0.922 | 0.952 | 480 | 0.744 | 1.000 | 935 | 0.322 | 1.000 | 519 | 0.602 | 1.000 | 450 | 0.646 | 1.000 | 868 | 0.952 | 0.952 |
| BRD-K66808046-065-03-7 | oxyquinoline | 0.465 | 84 | 0.182 | 0.545 | 973 | 0.901 | 0.940 | 1179 | 0.902 | 0.940 | 2307 | 0.039 | 0.275 | 2511 | 0.769 | 0.940 | 2338 | 0.275 | 0.607 | 2412 | 0.033 | 0.275 |
| BRD-K67261995-001-11-7 | iodipamide | 0.773 | | | | | | | 254 | 0.105 | 1.000 | 1518 | 0.511 | 0.716 | 319 | 0.564 | 0.741 | 464 | 0.393 | 0.716 | 801 | 0.322 | 1.000 |
| BRD-K67298865-001-09-0 | SB-431542 | 0.897 | | | | 340 | 0.059 | 0.621 | 1015 | 0.383 | 0.805 | 2614 | 0.710 | 0.897 | 1917 | 0.778 | 0.897 | 1313 | 0.611 | 0.897 | 2374 | 0.615 | 0.897 |
| BRD-K67511046-001-02-3 | naloxone | 0.722 | | | | 201 | 0.603 | 0.844 | 571 | 0.854 | 0.897 | 793 | 0.515 | 1.000 | 440 | 0.098 | 1.000 | 636 | 0.788 | 0.897 | 1077 | 0.601 | 0.844 |
| BRD-K67536197-003-11-4 | cyclocytidine | 0.593 | | | | 1156 | 0.875 | 0.958 | 1195 | 0.437 | 1.000 | 2680 | 0.281 | 1.000 | 2752 | 0.342 | 1.000 | 1511 | 0.633 | 0.950 | 3155 | 0.563 | 0.950 |
| BRD-K67566344-001-07-5 | ku-0063794 | 0.710 | 189 | 0.549 | 0.864 | 631 | 0.365 | 0.697 | 1979 | 0.998 | 0.999 | 3417.5 | 0.698 | 0.864 | 2403.5 | 0.354 | 0.697 | 1345 | 0.021 | 0.307 | 3207 | 0.660 | 0.864 |
| BRD-K67772619-001-05-1 | rheochrysidin | 0.061 | | | | 116 | 0.631 | 0.829 | 480 | 0.510 | 1.000 | 934 | 0.510 | 1.000 | 549 | 0.015 | 0.309 | 351 | 0.386 | 0.676 | 1272 | 0.919 | 0.919 |
| BRD-K67844266-003-01-9 | pevonedistat | 0.974 | | | | 312 | 0.615 | 0.974 | 1040 | 0.650 | 0.974 | 1460 | 0.288 | 0.606 | 1485 | 0.098 | 1.000 | 1967 | 0.901 | 0.988 | 1867 | 0.123 | 0.353 |
| BRD-K67868012-001-07-6 | PI-103 | 0.503 | | | | 562 | 0.492 | 1.000 | 646 | 0.681 | 0.894 | 2480 | 0.920 | 0.989 | 1853 | 0.598 | 0.837 | 892 | 0.010 | 0.027 | 2239 | 0.588 | 0.837 |
| BRD-K68065987-300-09-1 | MK-2206 | 0.972 | 100 | 0.191 | 0.466 | 1152 | 0.714 | 0.992 | 2249.5 | 0.765 | 0.992 | 3586 | 0.113 | 0.379 | 2510 | 0.289 | 0.607 | 2106.5 | 0.068 | 0.379 | 4674 | 0.882 | 0.998 |
| BRD-K68103045-001-02-9 | CGS-20625 | 0.618 | 41 | 0.240 | 0.618 | 65 | 0.451 | 0.790 | 290 | 0.571 | 0.883 | 606 | 0.897 | 0.981 | 168 | 0.093 | 0.598 | 302.5 | 0.630 | 0.883 | 975 | 0.834 | 0.981 |
| BRD-K68164687-001-01-6 | epothilone-b | 0.422 | 41 | 0.096 | 0.422 | 1413 | 0.645 | 0.891 | 2418 | 0.759 | 0.938 | 3508 | 0.062 | 0.422 | 3708 | 0.446 | 0.851 | 3829 | 0.819 | 0.956 | 3755 | 0.078 | 0.422 |
| BRD-K68174511-001-06-6 | torin-2 | 0.911 | | | | 1034 | 0.654 | 0.911 | 1869 | 0.736 | 0.911 | 3238 | 0.491 | 0.736 | 3090 | 0.239 | 0.559 | 1441 | 0.008 | 0.055 | 2902 | 0.222 | 0.559 |
| BRD-K68202742-001-16-5 | trichostatin-a | 0.137 | 66 | 0.168 | 0.586 | 1198 | 0.284 | 0.626 | 1873 | 0.630 | 0.882 | 3061 | 0.216 | 0.590 | 1951 | 0.003 | 0.054 | 2163 | 0.338 | 0.646 | 4045 | 0.687 | 0.896 |
| BRD-K68336408-001-10-9 | tyrphostin-AG-1478 | 0.631 | 86 | 0.174 | 0.525 | 1448.5 | 0.983 | 1.000 | 1520 | 0.328 | 0.631 | 2223 | 0.002 | 0.043 | 2516 | 0.183 | 0.525 | 1020 | 0.008 | 0.056 | 3103 | 0.090 | 0.376 |

| broad_id | name | colorectal_MWU_STAT | colorectal_PVAL | colorectal_PVAL_ADJ | skin_MWU_STAT | skin_PVAL | skin_PVAL_ADJ | urinary_tract_MWU_STAT | urinary_tract_PVAL | urinary_tract_PVAL_ADJ | lung_MWU_STAT | lung_PVAL | lung_PVAL_ADJ | ovary_MWU_STAT | ovary_PVAL | ovary_PVAL_ADJ | breast_MWU_STAT | breast_PVAL | breast_PVAL_ADJ | pancreas_MWU_STAT | pancreas_PVAL | pancreas_PVAL_ADJ | nervous_system_MWU_STAT | central_nervous_system_PVAL | nervous_system_PVAL_ADJ | soft_tissue_MWU_STAT | soft_tissue_PVAL | soft_tissue_PVAL_ADJ | kidney_MWU_STAT | kidney_PVAL | kidney_PVAL_ADJ | gastric_MWU_STAT | gastric_PVAL | gastric_PVAL_ADJ | bone_MWU_STAT | bone_PVAL | bone_PVAL_ADJ | prostate_MWU_STAT | prostate_PVAL | prostate_PVAL_ADJ | peripheral_nervous_system_MWU_STAT | peripheral_nervous_system_PVAL |
|---|---|---|---|---|---|---|---|---|---|---|---|---|---|---|---|---|---|---|---|---|---|---|---|---|---|---|---|---|---|---|---|---|---|---|---|---|---|---|---|---|---|---|
| BRD-K68346641-001-01-4 | amsacrine | 2549 | 0.135 | 0.594 | 5414 | 0.782 | 0.970 | 3264 | 0.840 | 0.970 | 8507 | 0.004 | 0.078 | 2438 | 0.045 | 0.470 | 2908 | 0.924 | 0.970 | 3529 | 0.538 | 0.970 | 5058 | 0.996 | 0.996 | 1546 | 0.498 | 0.970 | 2268 | 0.878 | 0.970 | 2179 | 0.303 | 0.908 | 1223 | 0.141 | 0.594 | 679 | 0.465 | 0.970 | 523 | 0.202 |
| BRD-K68392338-003-02-0 | ZK-93426 | 980 | 0.201 | 0.674 | 1455 | 0.859 | 0.979 | 805 | 0.093 | 0.639 | 3841 | 0.462 | 0.840 | 1241 | 0.350 | 0.749 | 651 | 0.665 | 0.979 | 1024 | 0.152 | 0.639 | 1322 | 0.920 | 0.979 | 802 | 0.932 | 0.979 | 625 | 0.975 | 0.979 | 468 | 0.028 | 0.293 | 244 | 0.291 | 0.749 | 176 | 0.901 | 0.979 | 113 | 0.145 |
| BRD-K68395654-001-03-1 | EVP4593 | 5728 | 0.993 | 0.993 | 8048 | 0.972 | 0.993 | 4422 | 0.532 | 0.870 | 15591 | 0.728 | 0.983 | 4541 | 0.105 | 0.368 | 3404 | 0.217 | 0.569 | 5203 | 0.156 | 0.469 | 7209 | 0.993 | 0.993 | 1139 | 0.024 | 0.256 | 2120 | 0.774 | 0.983 | 2103 | 0.104 | 0.368 | 2065 | 0.516 | 0.870 | 697 | 0.295 | 0.620 | 1314 | 0.843 |
| BRD-K68408467-065-01-8 | proflavine-hemisulfate | 626 | 0.021 | 0.108 | 2627 | 0.999 | 0.999 | 1094 | 0.373 | 0.784 | 2693 | 0.373 | 0.784 | 1737 | 0.791 | 0.999 | 362 | 0.460 | 0.807 | 1219 | 0.461 | 0.807 | 812 | 0.159 | 0.558 | 483 | 0.558 | 0.838 | 675 | 0.958 | 0.999 | 381 | 0.003 | 0.072 | 269 | 0.833 | 0.999 | 189 | 0.995 | 0.999 | 104 | 0.550 |
| BRD-K68408772-001-05-4 | carboxyamidotriazole | 249 | 0.270 | 1.000 | 733 | 0.708 | 0.987 | 422 | 0.198 | 1.000 | 1767 | 0.009 | 0.183 | 712 | 0.450 | 0.787 | 434 | 0.228 | 0.798 | 953 | 0.561 | 0.907 | 938 | 0.982 | 0.999 | 264 | 0.128 | 0.335 | 469 | 0.968 | 0.999 | 259 | 0.309 | 0.600 | 108 | 0.755 | 0.987 | 308 | 0.846 | 0.987 | | |
| BRD-K68488863-001-04-9 | ENMD-2076 | 1044 | 0.012 | 0.258 | 3010 | 0.944 | 0.977 | 1472.5 | 0.249 | 0.690 | 6693 | 0.762 | 0.975 | 2190 | 0.296 | 0.690 | 1152 | 0.284 | 0.690 | 2473.5 | 0.865 | 0.977 | 3015 | 0.977 | 0.977 | 807 | 0.546 | 0.881 | 1863 | 0.894 | 0.977 | 1095 | 0.103 | 0.434 | 506 | 0.022 | 0.116 | 117 | 0.418 | 0.749 | 497 | 0.421 |
| BRD-K68502831-001-01-2 | dianhydrogalactitol | 1490 | 0.995 | 0.995 | 2566 | 0.521 | 0.912 | 902 | 0.174 | 0.456 | 4058 | 0.059 | 0.436 | 1612 | 0.673 | 1.000 | 1431 | 0.950 | 1.000 | 1625 | 0.926 | 1.000 | 1154 | 0.418 | 0.873 | 784 | 0.294 | 0.687 | 989 | 0.717 | 1.000 | 817 | 0.088 | 0.436 | 758 | 0.457 | 0.873 | 128 | 0.145 | 0.436 | 9 | 0.057 |
| BRD-K68532323-003-02-8 | SB-743921 | 252 | 0.001 | 0.015 | 348 | 0.897 | 0.991 | 61 | 0.035 | 0.372 | 631 | 0.684 | 0.903 | 311 | 0.513 | 0.903 | 242 | 0.693 | 0.903 | 508 | 0.140 | 0.368 | 523 | 0.791 | 0.923 | 90 | 0.108 | 0.324 | 654 | 0.953 | 0.998 | 52 | 0.486 | 0.903 | 114 | 0.567 | 0.903 | | | | 90 | 0.359 |
| BRD-K68552125-001-05-3 | 12-O-tetradecanoylphorbol-13-acetate | 346 | 0.676 | 0.978 | 426 | 0.791 | 0.978 | 306 | 0.053 | 0.553 | 980 | 0.354 | 0.917 | 356 | 0.480 | 0.917 | 336 | 0.977 | 0.978 | 222 | 0.016 | 0.338 | 568 | 0.894 | 0.978 | 154 | 0.852 | 0.978 | 318 | 0.303 | 0.909 | 95 | 0.116 | 0.593 | 197 | 0.978 | 0.978 | 48 | 0.091 | 0.593 | 99 | 0.923 |
| BRD-K68567222-001-01-2 | BVT-948 | 477 | 0.936 | 1.000 | 1090 | 0.003 | 0.073 | 147 | 0.113 | 0.487 | 3049 | 0.973 | 1.000 | 1278 | 0.615 | 0.966 | 639 | 0.899 | 1.000 | 532 | 0.644 | 0.966 | 906 | 0.594 | 0.966 | 313 | 0.185 | 0.649 | 367 | 0.347 | 0.839 | 380 | 0.392 | 0.839 | 96 | 0.008 | 0.083 | 148 | 0.907 | 1.000 | 149 | 0.400 |
| BRD-K68633617-003-02-9 | ACDPP | 164 | 0.296 | 0.647 | 747 | 0.070 | 0.732 | 412 | 0.339 | 0.647 | 1660 | 0.614 | 0.859 | 545 | 0.395 | 0.692 | 204 | 0.075 | 0.262 | 673 | 0.984 | 0.998 | 910 | 0.998 | 0.998 | 260 | 0.472 | 0.762 | 277 | 0.558 | 0.838 | 183 | 0.046 | 0.732 | 126 | 0.137 | 0.410 | 110 | 0.859 | 0.968 | | |
| BRD-K68747584-001-02-0 | PF-03814735 | 1365 | 0.016 | 0.323 | 3593 | 0.987 | 0.987 | 1605 | 0.042 | 0.323 | 6778 | 0.442 | 0.920 | 2297 | 0.155 | 0.751 | 1752.5 | 0.927 | 0.987 | 2817.5 | 0.671 | 0.987 | 2071.5 | 0.392 | 0.920 | 393.5 | 0.046 | 0.323 | 1724.5 | 0.910 | 0.987 | 1556 | 0.227 | 0.751 | 1331 | 0.250 | 0.751 | 471.5 | 0.947 | 0.987 | 298 | 0.188 |
| BRD-K68904758-001-03-5 | cephalomannine | 6015 | 0.898 | 1.000 | 9466 | 0.981 | 1.000 | 3641 | 0.023 | 0.164 | 14858 | 0.080 | 0.270 | 5786 | 0.273 | 0.705 | 5239 | 0.921 | 1.000 | 6865 | 0.839 | 1.000 | 7587 | 0.877 | 1.000 | 1533 | 0.057 | 0.270 | 3678 | 1.000 | 1.000 | 2957 | 0.302 | 0.705 | 2012 | 0.352 | 0.738 | 964 | 0.628 | 1.000 | 1194 | 0.633 |
| BRD-K68938568-001-01-7 | GDC-0152 | 959 | 0.713 | 0.995 | 975.5 | 0.000 | 0.008 | 896 | 0.372 | 1.000 | 3199.5 | 0.228 | 1.000 | 1797 | 0.923 | 0.995 | 852.5 | 0.284 | 1.000 | 1391 | 0.907 | 0.995 | 1145.5 | 0.008 | 0.020 | 201.5 | 0.515 | 1.000 | 326 | 0.616 | 1.000 | 890 | 0.205 | 1.000 | 695 | 0.945 | 0.995 | 186.5 | 0.933 | 0.995 | 162 | 0.860 |
| BRD-K69001009-001-02-8 | golvatinib | 252 | 0.431 | 0.692 | 913 | 0.357 | 1.000 | 605 | 0.601 | 0.789 | 1267 | 0.119 | 1.000 | 883.5 | 0.980 | 0.982 | 674 | 0.927 | 0.982 | 494 | 0.422 | 0.814 | 805.5 | 0.922 | 0.982 | 81 | 0.039 | 0.118 | 251 | 0.426 | 0.814 | 223 | 0.293 | 1.000 | 343 | 0.982 | 0.982 | | | | 113 | 0.350 |
| BRD-K69116396-001-04-1 | sertindole | 2122 | 0.944 | 0.944 | 3039 | 0.340 | 0.929 | 1288 | 0.132 | 0.725 | 5227 | 0.269 | 0.929 | 1878 | 0.442 | 0.929 | 1372 | 0.541 | 0.929 | 2146 | 0.506 | 0.929 | 1796 | 0.648 | 0.929 | 829 | 0.098 | 0.725 | 1088 | 0.866 | 1.000 | 1235 | 0.715 | 0.929 | 236 | 0.138 | 0.725 | 395 | 0.547 | 0.929 | 220 | 0.377 |
| BRD-K69280563-001-01-8 | vinorelbine | 6520 | 0.987 | 1.000 | 7903 | 0.753 | 1.000 | 4273 | 0.307 | 0.644 | 14351 | 0.041 | 0.390 | 5403 | 0.418 | 0.798 | 5153 | 0.846 | 1.000 | 6881 | 0.993 | 1.000 | 6834 | 0.697 | 1.000 | 2021 | 0.110 | 0.390 | 3191 | 1.000 | 1.000 | 2982 | 0.218 | 0.573 | 1864 | 0.124 | 0.390 | 857 | 0.487 | 0.851 | 717 | 0.277 |
| BRD-K69608737-001-10-2 | tacrolimus | 75 | 0.718 | 0.962 | 710 | 0.917 | 0.962 | 343 | 0.357 | 1.000 | 1303 | 0.543 | 1.000 | 404 | 0.414 | 1.000 | 79 | 0.758 | 0.962 | 524.5 | 0.962 | 0.962 | 609 | 0.925 | 0.962 | 111 | 0.013 | 0.030 | 14 | 0.101 | 0.176 | 102 | 0.129 | 1.000 | 31 | 0.041 | 0.078 | 83 | 0.794 | 0.962 | 100 | 0.402 |
| BRD-K69650333-003-14-0 | idarubicin | 4165 | 0.711 | 0.920 | 5712 | 0.377 | 0.753 | 3801 | 0.711 | 0.920 | 11812 | 0.080 | 0.691 | 3889 | 0.371 | 0.753 | 3565 | 0.802 | 0.920 | 5178 | 0.782 | 0.920 | 6262 | 0.999 | 0.999 | 1775 | 0.197 | 0.691 | 2507 | 0.871 | 0.920 | 2236 | 0.166 | 0.691 | 2611 | 0.809 | 0.920 | 711 | 0.394 | 0.753 | 630 | 0.269 |
| BRD-K69694239-001-02-2 | bardoxolone-methyl | 3787 | 0.341 | 0.675 | 6718 | 0.578 | 0.867 | 2886 | 0.016 | 0.166 | 14535 | 0.868 | 1.000 | 3438 | 0.045 | 0.234 | 4192 | 0.975 | 1.000 | 7550 | 1.000 | 1.000 | 6406 | 0.976 | 1.000 | 1622 | 0.336 | 0.675 | 1847 | 0.213 | 0.638 | 2265 | 0.075 | 0.313 | 1320 | 0.097 | 0.339 | 690 | 0.668 | 0.877 | 518 | 0.033 |
| BRD-K69726342-238-02-4 | atorvastatin | 310 | 0.617 | 0.989 | 489 | 0.000 | 0.001 | 672 | 0.955 | 0.989 | 2457 | 0.989 | 0.989 | 717 | 0.920 | 0.989 | 213 | 0.486 | 1.000 | 474 | 0.696 | 0.989 | 631 | 0.010 | 0.025 | 327 | 0.170 | 0.358 | 375 | 0.861 | 0.989 | 246 | 0.316 | 1.000 | 204 | 0.161 | 0.358 | 78 | 0.553 | 0.967 | | |
| BRD-K69776681-001-03-8 | volasertib | 1327.5 | 0.289 | 0.606 | 2548.5 | 0.895 | 0.979 | 1533 | 0.616 | 0.924 | 4835 | 0.238 | 0.606 | 1554.5 | 0.118 | 0.476 | 954.5 | 0.793 | 0.979 | 2028.5 | 0.967 | 0.979 | 2452 | 0.954 | 0.979 | 800 | 0.266 | 0.606 | 869 | 0.851 | 0.979 | 1023 | 0.486 | 0.785 | 554 | 0.198 | 0.593 | 409.5 | 0.964 | 0.979 | 160 | 0.012 |
| BRD-K69932463-001-10-6 | AZD8055 | 5958 | 0.978 | 0.982 | 8655 | 0.953 | 0.982 | 4333 | 0.229 | 0.534 | 15790 | 0.501 | 0.876 | 3975 | 0.014 | 0.147 | 4575 | 0.636 | 0.982 | 7240 | 0.860 | 0.982 | 7561 | 0.967 | 0.982 | 2358 | 0.710 | 0.982 | 3170 | 0.340 | 0.648 | 2674 | 0.151 | 0.473 | 1788 | 0.088 | 0.464 | 739 | 0.311 | 0.648 | 1478 | 0.918 |
| BRD-K70163715-300-05-7 | vatalanib | 1193.5 | 0.785 | 0.990 | 1648 | 0.704 | 0.990 | 1299.5 | 0.441 | 0.862 | 3817.5 | 0.165 | 0.576 | 1597 | 0.328 | 0.772 | 1053.5 | 0.529 | 0.898 | 1645 | 0.922 | 0.990 | 1264 | 0.137 | 0.576 | 471 | 0.331 | 0.772 | 866.5 | 0.992 | 0.992 | 359 | 0.107 | 0.576 | 151 | 0.057 | 0.576 | 74 | 0.294 | 0.772 | 36 | 0.022 |
| BRD-K70301465-001-05-9 | ibrutinib | 472 | 0.711 | 1.000 | 825 | 0.999 | 1.000 | 518 | 0.021 | 0.153 | 1411 | 0.519 | 0.990 | 645 | 0.135 | 0.334 | 328 | 0.027 | 0.087 | 819 | 0.957 | 1.000 | 1089 | 1.000 | 1.000 | 224 | 0.919 | 1.000 | 319 | 0.676 | 1.000 | 150 | 0.015 | 0.153 | 249 | 0.966 | 1.000 | | | | 125 | 0.902 |
| BRD-K70401845-001-15-7 | erlotinib | 985 | 0.999 | 1.000 | 846 | 0.993 | 1.000 | 270 | 0.013 | 0.136 | 1736 | 0.678 | 1.000 | 500.5 | 0.071 | 0.300 | 338 | 0.841 | 1.000 | 419 | 0.980 | 1.000 | 873.5 | 1.000 | 1.000 | 61 | 0.444 | 0.776 | 245 | 0.176 | 0.369 | 92 | 0.056 | 0.395 | 118.5 | 0.910 | 1.000 | | | | | |
| BRD-K70401845-003-09-6 | erlotinib | 1295.5 | 0.833 | 0.992 | 2592 | 0.982 | 0.992 | 1185 | 0.003 | 0.029 | 5902.5 | 0.781 | 0.992 | 2639.5 | 0.855 | 0.992 | 1291.5 | 0.453 | 0.951 | 2084.5 | 0.947 | 0.992 | 2083 | 0.982 | 0.992 | 691 | 0.683 | 0.992 | 1461 | 0.992 | 0.992 | 937.5 | 0.118 | 0.353 | 106 | 0.082 | 0.320 | 158 | 0.188 | 0.494 | 161 | 0.693 |
| BRD-K70463136-001-01-5 | BAY-87-2243 | 4952 | 0.845 | 0.941 | 7163 | 0.805 | 0.941 | 4410.5 | 0.516 | 0.917 | 14137 | 0.349 | 0.917 | 5011 | 0.419 | 0.917 | 2555 | 0.076 | 0.526 | 6359 | 0.475 | 0.917 | 6807 | 0.929 | 0.975 | 1557 | 0.040 | 0.419 | 3304.5 | 0.980 | 0.980 | 2452 | 0.100 | 0.526 | 2380 | 0.804 | 0.941 | 750 | 0.365 | 0.917 | 1263 | 0.796 |
| BRD-K70511574-001-06-9 | HMN-214 | 3859 | 0.616 | 0.863 | 5288 | 0.921 | 0.996 | 2560.5 | 0.135 | 1.000 | 11557 | 0.389 | 0.628 | 4192 | 0.364 | 0.628 | 3246 | 0.679 | 0.883 | 5756.5 | 0.948 | 0.996 | 4342 | 0.358 | 0.628 | 1448 | 0.268 | 0.628 | 3209 | 1.000 | 1.000 | 1630 | 0.069 | 1.000 | 1192 | 0.031 | 0.108 | 481 | 0.341 | 0.628 | 769 | 0.259 |
| BRD-K70914287-300-02-8 | BIBX-1382 | 265 | 0.806 | 0.996 | 980 | 0.996 | 0.996 | 252 | 0.021 | 0.218 | 1781 | 0.678 | 0.949 | 563 | 0.242 | 1.000 | 200 | 0.471 | 1.000 | 780 | 0.948 | 0.996 | 693 | 0.985 | 0.996 | 394 | 0.937 | 0.996 | 171 | 0.312 | 0.546 | 224 | 0.102 | 0.714 | 160 | 0.258 | 0.493 | | | | 71 | 0.520 |
| BRD-K71106091-001-06-1 | fludarabine-phosphate | 697.5 | 0.691 | 0.893 | 1140.5 | 0.252 | 1.000 | 790 | 0.344 | 1.000 | 2423.5 | 0.699 | 0.893 | 443 | 0.011 | 0.235 | 462 | 0.928 | 0.994 | 704 | 0.168 | 1.000 | 1081 | 0.700 | 0.893 | 147 | 0.120 | 0.252 | 582.5 | 0.814 | 0.949 | 328 | 0.242 | 1.000 | 166 | 0.515 | 0.893 | 165 | 0.509 | 0.893 | 192 | 0.663 |
| BRD-K71164191-001-01-0 | KD-023 | 1731 | 0.459 | 0.926 | 3529 | 0.344 | 0.926 | 1743 | 0.656 | 0.926 | 5771 | 0.010 | 0.216 | 2580 | 0.572 | 0.926 | 1354 | 0.685 | 0.926 | 2466 | 0.844 | 0.958 | 2516 | 0.499 | 0.926 | 1044 | 0.855 | 0.958 | 939 | 0.702 | 0.926 | 1089 | 0.136 | 0.926 | 1498 | 0.993 | 0.993 | 502 | 0.362 | 0.926 | 353 | 0.315 |
| BRD-K71221037-001-01-6 | birinapant | 503 | 0.239 | 0.718 | 1888 | 0.078 | 0.819 | 811 | 0.317 | 0.831 | 3891 | 0.496 | 0.963 | 1120 | 0.576 | 0.963 | 894 | 0.164 | 0.619 | 1209 | 0.742 | 0.963 | 1564 | 0.177 | 0.619 | 279 | 0.043 | 0.224 | 742 | 0.998 | 0.998 | 288 | 0.014 | 0.297 | 530 | 0.569 | 0.963 | 159 | 0.825 | 0.963 | 187 | 0.920 |
| BRD-K71281111-001-04-3 | SNX-2112 | 6249 | 0.698 | 0.952 | 8072 | 0.394 | 0.753 | 4992 | 0.450 | 0.787 | 18085 | 0.725 | 0.952 | 4736 | 0.023 | 0.166 | 5490 | 0.919 | 0.998 | 9020 | 0.995 | 0.998 | 7664 | 0.772 | 0.953 | 1724 | 0.016 | 0.166 | 5395 | 0.998 | 0.998 | 2723 | 0.024 | 0.166 | 2945 | 0.695 | 0.952 | 801 | 0.334 | 0.702 | 796 | 0.124 |
| BRD-K71467466-001-02-3 | KPT-185 | 6547 | 0.964 | 0.991 | 9164 | 0.988 | 0.991 | 3624 | 0.143 | 0.429 | 15744 | 0.465 | 0.976 | 4186 | 0.014 | 0.097 | 3772 | 0.128 | 0.429 | 6518 | 0.585 | 0.991 | 4869 | 0.008 | 0.079 | 2552 | 0.668 | 0.991 | 4595 | 0.991 | 0.991 | 4808 | 0.990 | 0.991 | 2604 | 0.712 | 0.991 | 703 | 0.256 | 0.672 | 1190 | 0.631 |
| BRD-K71480163-001-01-4 | GDC-0068 | 1356 | 0.314 | 0.911 | 1630 | 0.603 | 0.957 | 1544 | 0.625 | 0.957 | 4468 | 0.983 | 0.983 | 1289 | 0.067 | 0.350 | 1590 | 0.158 | 0.555 | 1366 | 0.931 | 0.977 | 2000 | 0.911 | 0.977 | 555 | 0.113 | 0.475 | 973 | 0.513 | 0.957 | 592 | 0.024 | 0.256 | 272 | 0.710 | 0.977 | 189 | 0.361 | 0.911 | 132 | 0.627 |
| BRD-K71512533-066-02-8 | SNS-314 | 1922 | 0.055 | 0.553 | 3959.5 | 0.941 | 0.951 | 2308 | 0.187 | 0.752 | 7964 | 0.374 | 0.880 | 2920 | 0.377 | 0.880 | 1364.5 | 0.729 | 0.951 | 2950.5 | 0.679 | 0.951 | 2730.5 | 0.215 | 0.752 | 831 | 0.161 | 0.752 | 2298 | 0.951 | 0.951 | 2169 | 0.293 | 0.880 | 679.5 | 0.051 | 0.553 | 413 | 0.795 | 0.951 | 483 | 0.079 |
| BRD-K71823332-001-03-7 | epothilone-a | 5085 | 0.360 | 0.824 | 10111 | 0.999 | 0.999 | 3576 | 0.014 | 0.148 | 16425 | 0.392 | 0.824 | 6549 | 0.832 | 0.999 | 6162 | 0.993 | 0.999 | 7113 | 0.812 | 0.999 | 6899 | 0.511 | 0.845 | 2365 | 0.454 | 0.845 | 4053 | 0.982 | 0.999 | 2575 | 0.283 | 0.786 | 2190 | 0.300 | 0.786 | 907 | 0.523 | 0.845 | 670 | 0.065 |
| BRD-K72093121-001-16-3 | vidarabine | 190 | 0.948 | 0.948 | 515.5 | 0.445 | 1.000 | 423 | 0.878 | 1.000 | 1203 | 0.238 | 1.000 | 436 | 0.738 | 1.000 | 213 | 0.026 | 0.540 | 547 | 0.752 | 1.000 | 374 | 0.710 | 1.000 | 186 | 0.616 | 1.000 | 58 | 0.512 | 1.000 | 257 | 0.937 | 1.000 | | | | 129 | 0.629 | 1.000 | 97 | 0.363 |
| BRD-K72167406-001-02-1 | cefditoren-pivoxil | 1139 | 0.986 | 0.986 | 1263 | 0.823 | 0.999 | 827 | 0.370 | 0.864 | 2443 | 0.743 | 0.999 | 1148 | 0.453 | 0.885 | 925 | 0.277 | 0.767 | 1084 | 0.631 | 0.999 | 315 | 0.007 | 0.131 | 149 | 0.114 | 0.767 | 247 | 0.188 | 0.767 | 280 | 0.292 | 0.767 | 307 | 0.977 | 1.000 | 80 | 0.464 | 0.885 | 128 | 0.280 |
| BRD-K72414522-001-06-7 | AZD5438 | 3122 | 0.127 | 1.000 | 5372 | 0.476 | 0.908 | 2724 | 0.261 | 0.790 | 11169 | 0.306 | 0.790 | 3600 | 0.049 | 0.305 | 2322 | 0.684 | 0.982 | 3501 | 0.058 | 0.305 | 5570 | 0.982 | 0.982 | 1499 | 0.567 | 0.982 | 2717 | 0.392 | 0.824 | 3124 | 0.873 | 0.982 | 764 | 0.272 | 0.790 | 756 | 0.861 | 0.982 | 756 | 0.861 |
| BRD-K72420232-001-12-3 | WZ-4002 | 833 | 0.960 | 0.987 | 559 | 0.968 | 0.987 | 753 | 0.707 | 0.987 | 1505 | 0.486 | 0.784 | 338 | 0.017 | 0.073 | 220 | 0.281 | 0.537 | 592 | 0.987 | 0.987 | 628 | 0.964 | 0.987 | 73 | 0.129 | 0.420 | 291 | 0.140 | 0.420 | 149 | 0.222 | 0.537 | 321 | 0.969 | 0.987 | 45 | 0.281 | 0.537 | 119 | 0.393 |
| BRD-K72533376-001-01-2 | etoposide-phosphate | 216 | 0.386 | 1.000 | 866 | 0.148 | 1.000 | 669 | 0.432 | 1.000 | 2720 | 0.702 | 0.991 | 844 | 0.064 | 1.000 | 397 | 0.974 | 0.991 | 354 | 0.652 | 0.991 | 894 | 0.815 | 0.991 | 340 | 0.046 | 0.121 | 546 | 0.747 | 0.991 | 548 | 0.508 | 0.889 | 132 | 0.092 | 0.215 | 253 | 0.918 | 0.991 | 156 | 0.151 |
| BRD-K72703948-001-10-1 | ZM-447439 | 1913 | 0.485 | 0.755 | 4240 | 0.991 | 0.997 | 1517 | 0.091 | 0.953 | 5934 | 0.338 | 1.000 | 3321 | 0.846 | 0.997 | 1759 | 0.294 | 0.685 | 3214 | 0.861 | 0.997 | 2404 | 0.133 | 0.350 | 828 | 0.040 | 0.121 | 1769 | 0.997 | 0.997 | 1031 | 0.013 | 0.274 | 229 | 0.005 | 0.017 | 695 | 0.968 | 0.997 | 110 | 0.344 |
| BRD-K72723676-003-17-8 | benzethonium | 2789 | 0.173 | 0.686 | 2741 | 0.001 | 0.027 | 2513 | 0.056 | 0.391 | 10710 | 0.888 | 0.981 | 3125 | 0.079 | 0.413 | 2764 | 0.528 | 0.913 | 4000 | 0.683 | 0.956 | 3830 | 0.990 | 1.000 | 631 | 0.018 | 0.193 | 3673 | 1.000 | 1.000 | 1822 | 0.196 | 0.686 | 1503 | 0.740 | 0.972 | 530 | 0.559 | 0.913 | 305 | 0.411 |
| BRD-K72726508-001-02-1 | arcyriaflavin-a | 186.5 | 0.073 | 0.255 | 899 | 0.324 | 0.619 | 439.5 | 0.105 | 0.314 | 1644 | 0.934 | 0.981 | 513 | 0.681 | 0.978 | 40 | 0.000 | 0.002 | 526 | 0.894 | 0.981 | 764 | 0.991 | 0.991 | 148.5 | 0.287 | 0.604 | 281 | 0.714 | 0.978 | 162.5 | 0.011 | 0.231 | | | | 33 | 0.210 | 0.548 | 36 | 0.235 |
| BRD-K72815923-034-01-2 | xanomeline | 1202 | 0.874 | 1.000 | 2356 | 0.847 | 1.000 | 774 | 0.010 | 0.070 | 3208 | 0.002 | 0.049 | 2145 | 0.732 | 1.000 | 1283 | 0.673 | 1.000 | 2628 | 0.997 | 1.000 | 1673 | 0.633 | 1.000 | 538 | 0.761 | 1.000 | 1685 | 1.000 | 1.000 | 486 | 0.005 | 0.055 | 455 | 0.096 | 0.334 | 203 | 0.408 | 0.952 | 204 | 0.919 |
| BRD-K72827473-001-01-0 | CEP-37440 | 1814 | 0.561 | 0.907 | 1973 | 0.009 | 0.098 | 1551 | 0.867 | 0.999 | 6028 | 0.475 | 0.831 | 2001 | 0.079 | 0.554 | 1730 | 0.969 | 0.999 | 2334 | 0.474 | 0.831 | 2837 | 0.970 | 0.999 | 412 | 0.237 | 0.711 | 995 | 0.868 | 0.999 | 930 | 0.156 | 0.711 | 375 | 0.442 | 0.831 | 807 | 0.968 | 0.999 | 556 | 0.288 |
| BRD-K72951360-001-01-4 | valrubicin | 5447 | 0.694 | 0.971 | 7649 | 0.547 | 0.870 | 3992 | 0.165 | 0.451 | 13025 | 0.045 | 0.451 | 4673 | 0.091 | 0.451 | 5105 | 0.957 | 0.995 | 7807 | 0.988 | 0.995 | 8438 | 0.995 | 0.995 | 1672 | 0.021 | 0.451 | 4054 | 0.969 | 0.995 | 2800 | 0.068 | 0.451 | 1818 | 0.215 | 0.501 | 399 | 0.132 | 0.451 | 1121 | 0.561 |
| BRD-K73073971-001-01-7 | ZLN005 | 509 | 0.643 | 0.965 | 799 | 0.476 | 0.965 | 371 | 0.516 | 0.965 | 1216 | 0.232 | 1.000 | 511 | 0.902 | 0.970 | 439 | 0.576 | 0.965 | 648 | 0.930 | 0.970 | 973 | 0.970 | 0.970 | 225 | 0.800 | 0.970 | 96 | 0.017 | 0.044 | 230 | 0.348 | 1.000 | | | | 76 | 0.690 | 0.965 | 96 | 0.196 |
| BRD-K73197500-001-02-2 | SNX-5422 | 4574 | 0.765 | 0.984 | 5621 | 0.451 | 0.677 | 3578 | 0.291 | 0.556 | 13221 | 0.843 | 0.984 | 4146 | 0.235 | 0.495 | 3806 | 0.991 | 0.991 | 6457 | 0.984 | 0.991 | 4421 | 0.211 | 0.495 | 1281 | 0.227 | 0.495 | 3714 | 0.984 | 0.991 | 2535 | 0.421 | 0.677 | 1456 | 0.227 | 0.495 | 645 | 0.630 | 0.882 | 500 | 0.353 |
| BRD-K73293050-001-03-1 | WZ-3146 | 2906 | 0.826 | 0.999 | 5436 | 0.999 | 0.999 | 1955 | 0.012 | 0.260 | 9347 | 0.487 | 0.771 | 2889 | 0.171 | 0.607 | 2277 | 0.234 | 0.607 | 3565 | 0.358 | 0.626 | 4891 | 0.985 | 0.999 | 635 | 0.195 | 0.607 | 1840 | 0.230 | 0.607 | 1587 | 0.269 | 0.607 | 995 | 0.514 | 0.771 | 697 | 0.877 | 0.999 | 466 | 0.162 |
| BRD-K73309154-003-02-8 | OSI-420 | 1186.5 | 0.749 | 0.994 | 2369 | 0.994 | 0.994 | 625 | 0.001 | 0.013 | 5747.5 | 0.638 | 0.994 | 2144 | 0.535 | 0.994 | 645 | 0.356 | 0.974 | 1398.5 | 0.714 | 0.994 | 2053 | 0.989 | 0.994 | 857.5 | 0.957 | 0.994 | 1103 | 0.397 | 0.974 | 1012 | 0.128 | 0.670 | 286 | 0.275 | 0.974 | 205 | 0.888 | 0.994 | 484 | 0.852 |
| BRD-K73753850-003-01-6 | hexaminolevulinate | 3548 | 0.992 | 0.999 | 4882 | 0.807 | 0.892 | 2292 | 0.183 | 0.723 | 8410 | 0.241 | 0.723 | 3155 | 0.218 | 0.723 | 1890 | 0.326 | 0.746 | 2928 | 0.647 | 0.876 | 3468 | 0.475 | 0.746 | 758 | 0.024 | 0.512 | 2276 | 0.430 | 0.746 | 1871 | 0.668 | 0.876 | 950 | 0.479 | 0.746 | 405 | 0.716 | 0.885 | 640 | 0.222 |
| BRD-K73838513-003-05-5 | cinacalcet | 2054 | 0.935 | 0.935 | 2809 | 0.200 | 0.886 | 1713.5 | 0.731 | 0.886 | 4425 | 0.463 | 0.886 | 1498 | 0.739 | 0.886 | 1381 | 0.734 | 0.886 | 1105 | 0.072 | 0.506 | 1476.5 | 0.535 | 0.886 | 1000 | 0.863 | 0.906 | 833 | 0.802 | 0.886 | 1049 | 0.343 | 0.886 | 219 | 0.436 | 0.886 | 173 | 0.791 | 0.886 | 439 | 0.773 |
| BRD-K73982490-001-02-9 | BI-78D3 | 3526 | 0.797 | 1.000 | 5861 | 0.511 | 0.825 | 2889 | 0.271 | 0.610 | 12271 | 0.951 | 1.000 | 2940 | 0.035 | 0.275 | 3059 | 0.411 | 0.719 | 4669 | 0.853 | 1.000 | 4305 | 0.194 | 0.610 | 2470 | 0.925 | 1.000 | 2073 | 0.262 | 0.610 | 1649 | 0.039 | 0.275 | 1394 | 0.114 | 0.600 | 580 | 0.248 | 0.610 | 425 | 0.019 |
| BRD-K74057757-001-01-9 | indibulin | 6021 | 0.881 | 0.974 | 8763 | 0.994 | 0.994 | 3997 | 0.180 | 0.630 | 14933 | 0.318 | 0.739 | 4298 | 0.016 | 0.132 | 4249 | 0.839 | 0.974 | 7538 | 0.988 | 0.994 | 6809 | 0.718 | 0.974 | 1870 | 0.136 | 0.630 | 2998 | 0.866 | 0.974 | 3080 | 0.454 | 0.795 | 2419 | 0.595 | 0.961 | 786 | 0.387 | 0.739 | 799 | 0.165 |
| BRD-K74141488-003-11-2 | naftifine | 252 | 0.630 | 1.000 | 653 | 0.250 | 0.764 | 123 | 0.196 | 0.764 | 1075 | 0.195 | 0.764 | 364.5 | 0.201 | 0.764 | 139.5 | 0.282 | 0.764 | 437 | 0.281 | 0.764 | 927.5 | 0.999 | 0.999 | 22 | 0.144 | 0.764 | 243 | 0.291 | 0.764 | 262.5 | 0.687 | 0.850 | 79 | 0.219 | 0.383 | 82 | 0.753 | 0.850 | 22 | 0.025 |
| BRD-K74193284-001-04-8 | phenylmercuric-acetate | 2698 | 0.986 | 0.988 | 4047 | 0.013 | 0.137 | 3768 | 0.934 | 0.988 | 13134 | 0.988 | 0.988 | 3385 | 0.082 | 0.390 | 2834 | 0.549 | 0.883 | 4611 | 0.985 | 0.988 | 4099 | 0.284 | 0.821 | 1495 | 0.097 | 0.390 | 2424 | 0.474 | 0.883 | 2261 | 0.313 | 0.821 | 837 | 0.007 | 0.137 | 635 | 0.688 | 0.904 | 589 | 0.598 |
| BRD-K74305673-001-06-2 | IKK-2-inhibitor-V | 2703 | 0.508 | 1.000 | 5676.5 | 0.762 | 0.915 | 2454.5 | 0.273 | 1.000 | 10864 | 0.521 | 1.000 | 3073 | 0.018 | 0.387 | 3035 | 0.519 | 1.000 | 4887 | 0.990 | 0.990 | 3807 | 0.640 | 0.840 | 1395 | 0.073 | 0.152 | 1391 | 0.302 | 0.488 | 2624 | 0.592 | 0.828 | 265 | 0.070 | 0.152 | 325 | 0.128 | 0.225 | 39 | 0.090 |
| BRD-K74501079-001-18-1 | azithromycin | 1419 | 0.394 | 0.690 | 3097 | 0.384 | 1.000 | 2188 | 0.937 | 0.978 | 7362 | 0.792 | 0.909 | 2499 | 0.059 | 1.000 | 1547 | 0.585 | 0.898 | 2816 | 0.823 | 0.909 | 2785 | 0.681 | 0.909 | 1086 | 0.278 | 0.583 | 1352 | 0.697 | 0.909 | 1037 | 0.214 | 1.000 | 517 | 0.059 | 0.155 | 218 | 0.305 | 0.583 | 517 | 0.750 |
| BRD-K74514084-001-03-9 | pazopanib | 517 | 0.927 | 0.943 | 506 | 0.752 | 0.943 | 360 | 0.606 | 0.926 | 1056 | 0.371 | 0.848 | 660 | 0.872 | 0.943 | 257 | 0.662 | 0.926 | 319 | 0.404 | 0.848 | 550 | 0.522 | 0.926 | 107 | 0.123 | 0.647 | 186 | 0.255 | 0.848 | 186 | 0.090 | 0.629 | 97 | 0.339 | 0.848 | 96 | 0.869 | 0.943 | 191 | 0.943 |
| BRD-K74671368-001-01-8 | idronoxil | 504 | 0.630 | 0.897 | 3024 | 0.901 | 0.998 | 1825 | 0.472 | 0.826 | 5080 | 0.641 | 0.897 | 1487 | 0.092 | 0.387 | 1554 | 0.691 | 0.906 | 1738 | 0.998 | 0.998 | 1079 | 0.134 | 0.403 | 631 | 0.374 | 0.792 | 1499 | 0.605 | 0.897 | 321 | 0.415 | 0.792 | 372 | 0.029 | 0.299 | 319 | 0.408 | 0.792 | 190 | 0.090 |
| BRD-K75009076-001-02-1 | SCH-900776 | 3346 | 0.607 | 0.901 | 5001 | 0.623 | 0.901 | 2611 | 0.657 | 0.901 | 10606 | 0.872 | 0.916 | 2297 | 0.015 | 0.306 | 2780 | 0.664 | 0.901 | 3567 | 0.532 | 0.901 | 4044 | 0.437 | 0.901 | 1205 | 0.317 | 0.739 | 1635 | 0.073 | 0.512 | 1601 | 0.127 | 0.646 | 789 | 0.183 | 0.646 | 475 | 0.816 | 0.901 | 725 | 0.299 |
| BRD-K75295174-001-05-0 | alisertib | 2595 | 0.708 | 0.954 | 3984 | 0.997 | 0.997 | 1909 | 0.141 | 0.594 | 7102 | 0.106 | 0.556 | 2086 | 0.028 | 0.369 | 1828 | 0.920 | 0.997 | 3014 | 0.960 | 0.997 | 3377 | 0.331 | 0.773 | 1202 | 0.557 | 0.954 | 1231 | 0.818 | 0.954 | 1452 | 0.175 | 0.611 | 541 | 0.053 | 0.369 | 406 | 0.813 | 0.954 | 193 | 0.046 |
| BRD-K75352575-001-01-3 | KPT-276 | 5242 | 0.956 | 0.956 | 6982 | 0.854 | 0.956 | 3863 | 0.442 | 0.956 | 14377 | 0.513 | 0.956 | 4264 | 0.017 | 0.160 | 2572 | 0.023 | 0.160 | 5485 | 0.668 | 0.956 | 5142 | 0.068 | 0.309 | 2219 | 0.703 | 0.956 | 3527 | 0.951 | 0.956 | 3382 | 0.907 | 0.956 | 2304 | 0.589 | 0.956 | 884 | 0.607 | 0.956 | 780 | 0.786 |
| BRD-K75649340-001-13-1 | tioguanine | 1465.5 | 0.188 | 0.438 | 1573.5 | 0.307 | 0.587 | 1424 | 0.077 | 0.438 | 6006.5 | 0.180 | 0.438 | 3053 | 0.755 | 0.991 | 2458.5 | 0.999 | 0.999 | 2814 | 0.929 | 0.999 | 2545.5 | 0.744 | 0.991 | 636 | 0.008 | 0.168 | 606 | 0.144 | 0.438 | 1367.5 | 0.109 | 0.438 | 689 | 0.262 | 0.551 | 332 | 0.695 | 0.991 | 60 | 0.029 |
| BRD-K75911534-001-01-6 | SU014813 | 1446.5 | 0.957 | 0.969 | 2781 | 0.770 | 0.969 | 1306 | 0.337 | 0.708 | 4522 | 0.484 | 0.924 | 2118.5 | 0.886 | 0.969 | 701 | 0.580 | 0.969 | 1686.5 | 0.889 | 0.969 | 964 | 0.029 | 0.199 | 498 | 0.053 | 0.199 | 496 | 0.141 | 0.369 | 553 | 0.036 | 0.199 | 201 | 0.007 | 0.151 | 374 | 0.622 | 0.969 | 191 | 0.879 |
| BRD-K75958547-238-01-0 | pitavastatin | 775 | 0.704 | 0.928 | 1075 | 0.000 | 0.009 | 1150 | 0.831 | 0.928 | 4233 | 0.928 | 0.928 | 1652 | 0.321 | 1.000 | 791 | 0.738 | 0.928 | 1747 | 0.928 | 0.928 | 732 | 0.002 | 0.007 | 460 | 0.170 | 0.397 | 905 | 0.763 | 0.928 | 1004 | 0.906 | 0.928 | 438 | 0.319 | 0.671 | 196 | 0.469 | 0.895 | 163 | 0.851 |
| BRD-K76064317-001-07-3 | tyrphostin-AG-1296 | 2373 | 0.904 | 0.995 | 3507 | 0.833 | 0.979 | 1601 | 0.061 | 0.425 | 6760 | 0.181 | 0.670 | 2835 | 0.724 | 0.950 | 2007 | 0.552 | 0.828 | 3241 | 0.984 | 0.995 | 2688 | 0.711 | 0.950 | 636 | 0.136 | 0.670 | 2109 | 0.995 | 0.995 | 1445 | 0.191 | 0.670 | 513 | 0.012 | 0.255 | 200 | 0.055 | 0.425 | 450 | 0.532 |
| BRD-K76180386-001-01-6 | JIB04 | 2746 | 0.549 | 0.915 | 3914 | 0.058 | 0.421 | 2340 | 0.865 | 0.915 | 7644 | 0.566 | 0.915 | 3022 | 0.192 | 0.743 | 1814 | 0.022 | 0.421 | 3714.5 | 0.865 | 0.915 | 2897 | 0.558 | 0.915 | 589 | 0.060 | 0.421 | 2093 | 0.795 | 0.915 | 1748 | 0.390 | 0.915 | 889 | 0.204 | 0.743 | 611 | 0.464 | 0.915 | 503 | 0.248 |
| BRD-K76239644-001-01-8 | BMS-690514 | 2097 | 0.117 | 0.306 | 3264 | 1.000 | 1.000 | 2199 | 0.103 | 0.306 | 9904 | 0.976 | 1.000 | 3702 | 0.570 | 0.921 | 1422 | 0.101 | 0.306 | 3425 | 0.438 | 0.767 | 4405 | 0.999 | 1.000 | 2003 | 0.990 | 1.000 | 1461 | 0.239 | 0.502 | 1237 | 0.005 | 0.032 | 734 | 0.726 | 1.000 | 493 | 0.909 | 1.000 | 782 | 0.979 |
| BRD-K76239644-001-02-6 | BMS-690514 | 478.5 | 0.312 | 0.595 | 2163 | 1.000 | 1.000 | 781.5 | 0.032 | 0.222 | 2846.5 | 0.742 | 1.000 | 1179 | 0.031 | 0.131 | 928 | 0.410 | 0.718 | 857.5 | 0.264 | 0.553 | 799.5 | 0.976 | 1.000 | 417 | 0.936 | 1.000 | 1004.5 | 0.901 | 1.000 | 469.5 | 0.127 | 0.334 | 283.5 | 0.904 | 1.000 | | | | | |
| BRD-K76406695-001-02-9 | CCT128930 | 319.5 | 0.363 | 0.758 | 610.5 | 0.970 | 0.993 | 551 | 0.725 | 0.993 | 2385 | 0.993 | 0.993 | 767 | 0.397 | 0.758 | 228 | 0.029 | 0.102 | 822 | 0.990 | 0.993 | 785.5 | 0.294 | 0.687 | 204 | 0.055 | 0.161 | 280.5 | 0.821 | 0.993 | 546.5 | 0.285 | 1.000 | | | | 150 | 0.544 | 0.878 | 201.5 | 0.836 |
| BRD-K76634210-001-14-7 | idoxuridine | 882 | 0.370 | 0.828 | 577 | 0.005 | 0.103 | 328 | 0.030 | 0.311 | 2624 | 0.723 | 0.949 | 441 | 0.067 | 0.283 | 688 | 0.960 | 0.989 | 1554 | 0.989 | 0.989 | 567 | 0.513 | 0.828 | 146 | 0.111 | 0.387 | 198 | 0.276 | 0.828 | 497 | 0.060 | 0.422 | 435 | 0.604 | 0.879 | 274 | 0.628 | 0.879 | 148 | 0.398 |
| BRD-K76674262-001-03-3 | homoharringtonine | 2779 | 0.443 | 0.903 | 5250 | 0.211 | 0.570 | 3821 | 0.748 | 0.998 | 12141 | 0.494 | 0.903 | 3490 | 0.092 | 0.486 | 2476 | 0.653 | 0.998 | 4587 | 0.938 | 0.998 | 5077 | 0.964 | 0.998 | 515 | 0.001 | 0.019 | 2836 | 0.969 | 0.998 | 2356 | 0.359 | 0.837 | 2339 | 0.945 | 0.998 | 543 | 0.186 | 0.570 | 924 | 0.516 |
| BRD-K76703230-004-04-1 | YM-155 | 6848 | 0.997 | 0.997 | 8049 | 0.498 | 0.997 | 5112 | 0.739 | 0.997 | 14802 | 0.010 | 0.070 | 4088 | 0.006 | 0.064 | 5065 | 0.822 | 0.997 | 7284 | 0.786 | 0.997 | 7582 | 0.887 | 0.997 | 1482 | 0.037 | 0.197 | 4598 | 0.996 | 0.997 | 3225 | 0.304 | 0.799 | 1847 | 0.088 | 0.371 | 965 | 0.602 | 0.997 | 930 | 0.551 |
| BRD-K76840893-003-02-5 | RS-17053 | 2099 | 0.162 | 0.870 | 3815 | 0.920 | 0.932 | 1824 | 0.255 | 0.870 | 8646 | 0.932 | 0.932 | 1827 | 0.008 | 0.177 | 2179 | 0.350 | 0.913 | 2478 | 0.056 | 0.584 | 3752 | 0.725 | 0.913 | 1265 | 0.621 | 0.913 | 1427 | 0.643 | 0.913 | 1116 | 0.204 | 0.870 | 650 | 0.605 | 0.913 | 592 | 0.812 | 0.913 | 907 | 0.784 |
| BRD-K76973696-001-12-6 | tryptanthrin | 3582 | 0.540 | 0.914 | 7875 | 0.979 | 0.997 | 2783 | 0.014 | 0.302 | 12032 | 0.146 | 0.472 | 4724 | 0.267 | 0.702 | 3280 | 0.588 | 0.914 | 5071 | 0.609 | 0.914 | 5532 | 0.921 | 0.997 | 1714 | 0.495 | 0.914 | 2750 | 0.436 | 0.914 | 1655 | 0.054 | 0.472 | 1773 | 0.774 | 0.973 | 379 | 0.157 | 0.472 | 1353 | 0.997 |
| BRD-K77008974-001-03-2 | WYE-354 | 2014 | 0.804 | 0.981 | 4084 | 0.981 | 0.981 | 1186 | 0.146 | 0.615 | 7328.5 | 0.104 | 0.615 | 3801 | 0.762 | 0.981 | 3176 | 0.949 | 0.981 | 2674 | 0.013 | 0.281 | 2366 | 0.717 | 0.981 | 1161 | 0.457 | 0.960 | 1396 | 0.802 | 0.981 | 1567 | 0.262 | 0.689 | 258 | 0.100 | 0.615 | 345 | 0.240 | 0.689 | 341 | 0.622 |
| BRD-K77625799-001-07-7 | vandetanib | 176 | 0.593 | 1.000 | 715 | 0.982 | 1.000 | 391 | 0.220 | 0.923 | 1180 | 0.667 | 0.934 | 346 | 0.215 | 0.923 | 303 | 0.916 | 1.000 | 106 | 0.469 | 1.000 | 289 | 0.989 | 1.000 | 182 | 0.634 | 0.934 | 134 | 0.300 | 0.573 | 178 | 0.607 | 0.934 | | | | | | | 177 | 0.932 |
| BRD-K77634909-001-03-5 | AEG3482 | 749 | 0.930 | 0.976 | 1362 | 0.729 | 0.944 | 504.5 | 0.289 | 0.505 | 1800.5 | 0.474 | 0.766 | 817 | 0.633 | 0.886 | 429.5 | 0.119 | 0.416 | 968.5 | 0.976 | 0.976 | 727.5 | 0.903 | 0.976 | 111 | 0.271 | 0.505 | 448.5 | 0.809 | 0.944 | 274 | 0.173 | 0.455 | | | | 240 | 0.597 | 0.886 | 103 | 0.229 |
| BRD-K77638923-001-02-0 | CUDC-907 | 5057 | 0.169 | 0.507 | 7848 | 0.292 | 0.681 | 4928 | 0.410 | 0.690 | 17164 | 0.335 | 0.690 | 6389 | 0.433 | 0.690 | 6137 | 0.981 | 0.999 | 9017 | 0.999 | 0.999 | 7668 | 0.773 | 0.955 | 2070 | 0.169 | 0.507 | 4843 | 0.992 | 0.999 | 2832 | 0.077 | 0.434 | 1936 | 0.103 | 0.434 | 500 | 0.059 | 0.434 | 647 | 0.047 |
| BRD-K77677632-003-02-6 | SB-200646 | 388 | 0.220 | 1.000 | 953 | 0.588 | 0.824 | 1124 | 0.903 | 0.979 | 1790 | 0.039 | 0.823 | 1128 | 0.508 | 0.762 | 676 | 0.427 | 0.762 | 767 | 0.480 | 0.762 | 863 | 0.877 | 0.979 | 214 | 0.364 | 0.762 | 288 | 0.144 | 0.379 | 508 | 0.359 | 0.762 | 211 | 0.121 | 0.362 | 78 | 0.471 | 0.762 | 17 | 0.088 |
| BRD-K77685957-300-09-1 | dequalinium | 1477 | 0.653 | 0.980 | 3555 | 0.976 | 0.995 | 834 | 0.035 | 0.394 | 6131 | 0.765 | 0.995 | 2406 | 0.246 | 1.000 | 2378 | 0.919 | 0.995 | 1234 | 0.034 | 0.103 | 2450 | 0.537 | 0.868 | 604 | 0.073 | 0.191 | 1234 | 0.850 | 0.995 | 441 | 0.038 | 0.394 | 721 | 0.995 | 0.995 | 620 | 0.962 | 0.995 | 71 | 0.218 |
| BRD-K77791657-003-01-7 | 3-deazaneplanocin-A | 3880 | 0.533 | 0.850 | 7397.5 | 1.000 | 1.000 | 3182 | 0.621 | 0.869 | 8942 | 0.020 | 0.105 | 3032 | 0.046 | 0.192 | 3533 | 0.871 | 0.920 | 4558 | 0.794 | 0.920 | 3600.5 | 0.308 | 0.719 | 1169 | 0.216 | 0.649 | 2644 | 0.868 | 0.920 | 2509.5 | 0.294 | 0.719 | 1125 | 0.356 | 0.748 | 485 | 0.141 | 0.495 | 514 | 0.463 |
| BRD-K77841042-001-14-1 | brefeldin-a | 4295 | 0.181 | 0.550 | 5338 | 0.001 | 0.018 | 4238 | 0.362 | 0.760 | 17399 | 0.771 | 0.991 | 6101 | 0.767 | 0.991 | 5797 | 0.991 | 0.991 | 8003 | 0.990 | 0.991 | 6618 | 0.515 | 0.983 | 1622 | 0.035 | 0.372 | 3287 | 0.255 | 0.595 | 2829 | 0.119 | 0.510 | 1631 | 0.090 | 0.510 | 1425 | 0.986 | 0.991 | 985 | 0.655 |
| BRD-K77908580-001-09-6 | entinostat | 2820 | 0.313 | 0.606 | 4053 | 0.057 | 0.302 | 2074 | 0.178 | 0.534 | 9286 | 0.318 | 0.606 | 3928 | 0.921 | 0.999 | 3663 | 0.976 | 0.999 | 4352 | 0.992 | 0.999 | 3465 | 0.859 | 0.999 | 1249 | 0.216 | 0.567 | 2409 | 0.999 | 0.999 | 1620 | 0.160 | 0.534 | 488 | 0.015 | 0.159 | 409 | 0.097 | 0.406 | 255 | 0.015 |
| BRD-K77943374-001-02-8 | NVP-BVU972 | 250 | 0.842 | 0.909 | 490 | 0.264 | 0.923 | 290 | 0.128 | 0.670 | 957 | 0.689 | 1.000 | 427 | 0.909 | 0.909 | 316 | 0.078 | 0.547 | 378 | 0.674 | 1.000 | 310 | 0.869 | 0.909 | | | | | | | 142 | 0.061 | 0.547 | 189 | 0.807 | 0.909 | 64 | 0.695 | 1.000 | 139 | 0.313 |
| BRD-K77987382-001-13-2 | mebendazole | 2201 | 0.715 | 1.000 | 2419 | 0.033 | 0.303 | 2190 | 0.265 | 0.928 | 6303 | 0.093 | 0.431 | 2716 | 0.317 | 0.952 | 2242 | 0.949 | 1.000 | 3673 | 0.998 | 1.000 | 2485 | 0.931 | 1.000 | 651 | 0.398 | 1.000 | 2044 | 1.000 | 1.000 | 1720 | 0.787 | 1.000 | 130 | 0.002 | 0.004 | 281 | 0.495 | 0.799 | 282 | 0.993 |
| BRD-K78055238-001-01-8 | LY2090314 | 2187 | 0.681 | 0.954 | 2413 | 0.491 | 0.837 | 1782 | 0.116 | 0.406 | 8520 | 0.987 | 0.987 | 3137 | 0.605 | 0.907 | 1409 | 0.102 | 0.406 | 2627 | 0.453 | 0.837 | 2764 | 0.157 | 0.472 | 890 | 0.843 | 0.987 | 1235 | 0.013 | 0.145 | 1909 | 0.498 | 0.837 | 551 | 0.465 | 0.837 | 704 | 0.800 | 0.987 | 432 | 0.518 |
| BRD-K78096648-001-05-7 | oxiperomide | 1410 | 0.180 | 0.757 | 1635 | 0.011 | 0.222 | 1425 | 0.482 | 0.922 | 4628 | 0.527 | 0.922 | 2139 | 0.761 | 0.993 | 912 | 0.518 | 0.922 | 1809 | 0.448 | 0.922 | 1208 | 0.168 | 0.757 | 947 | 0.946 | 0.993 | 923 | 0.927 | 0.993 | 494 | 0.045 | 0.469 | 207 | 0.401 | 0.922 | 408 | 0.693 | 0.971 | 184 | 0.317 |
| BRD-K78126613-001-28-5 | menadione | 448 | 0.062 | 0.437 | 2634 | 0.094 | 0.491 | 1650 | 0.867 | 0.929 | 4412 | 0.391 | 0.929 | 2332 | 0.858 | 0.929 | 1451 | 0.430 | 0.929 | 1786 | 0.764 | 0.929 | 2672 | 0.929 | 0.929 | 471 | 0.219 | 0.790 | 1221 | 0.602 | 0.929 | 1145 | 0.850 | 0.929 | 193 | 0.019 | 0.399 | 369 | 0.529 | 0.929 | 529 | 0.915 |
| BRD-K78177893-001-02-4 | A-674563 | 2515 | 0.030 | 0.209 | 6035 | 0.984 | 0.987 | 3490 | 0.593 | 0.987 | 9981 | 0.007 | 0.149 | 4556 | 0.636 | 0.987 | 2874 | 0.502 | 0.987 | 4815 | 0.586 | 0.987 | 5785 | 0.977 | 0.987 | 1234 | 0.100 | 0.523 | 2642 | 0.896 | 0.987 | 1308 | 0.025 | 0.209 | 1270 | 0.476 | 0.987 | 1035 | 0.910 | 0.987 | 724 | 0.208 |
| BRD-K78318619-001-02-7 | PF-4981517 | 203.5 | 0.604 | 0.845 | 289 | 0.078 | 0.817 | 558.5 | 0.588 | 1.000 | 1618.5 | 0.678 | 0.890 | 598 | 0.530 | 1.000 | 349 | 0.920 | 0.964 | 369 | 0.516 | 1.000 | 412.5 | 0.252 | 1.000 | 386.5 | 0.839 | 0.927 | 163 | 0.767 | 0.927 | 143 | 0.074 | 0.817 | 110.5 | 0.385 | 0.635 | 69 | 0.136 | 0.260 | 111.5 | 0.393 |
| BRD-K78431006-001-10-2 | crizotinib | 2601 | 0.145 | 0.815 | 4992 | 0.945 | 0.945 | 3179 | 0.357 | 0.815 | 9378 | 0.427 | 0.815 | 4036 | 0.593 | 0.831 | 2033 | 0.585 | 0.831 | 4506 | 0.891 | 0.936 | 3862 | 0.571 | 0.831 | 1048 | 0.160 | 0.815 | 1661 | 0.729 | 0.919 | 1722 | 0.236 | 0.815 | 1244 | 0.204 | 0.815 | 448 | 0.775 | 0.919 | 793 | 0.427 |

| broad_id | name | al_nervous_system_PVAL_ADJ | fibroblast_MWU_STAT | fibroblast_PVAL | fibroblast_PVAL_ADJ | bile_duct_MWU_STAT | bile_duct_PVAL | bile_duct_PVAL_ADJ | thyroid_MWU_STAT | thyroid_PVAL | thyroid_PVAL_ADJ | upper_aerodigestive_MWU_STAT | upper_aerodigestive_PVAL | upper_aerodigestive_PVAL_ADJ | uterus_MWU_STAT | uterus_PVAL | uterus_PVAL_ADJ | liver_MWU_STAT | liver_PVAL | liver_PVAL_ADJ | esophagus_MWU_STAT | esophagus_PVAL | esophagus_PVAL_ADJ |
|---|---|---|---|---|---|---|---|---|---|---|---|---|---|---|---|---|---|---|---|---|---|---|---|
| BRD-K68346641-001-01-4 | amsacrine | 0.708 | 309 | 0.878 | 0.970 | 1390 | 0.919 | 0.970 | 1793 | 0.792 | 0.970 | 3634 | 0.857 | 0.970 | 2779 | 0.583 | 0.970 | 1573 | 0.087 | 0.594 | 3351 | 0.660 | 0.970 |
| BRD-K68392338-003-02-0 | ZK-93426 | 0.639 | 79 | 0.357 | 0.749 | 161 | 0.849 | 0.979 | 400 | 0.840 | 0.979 | 1312 | 0.480 | 0.840 | 371 | 0.020 | 0.293 | 570 | 0.225 | 0.674 | 1856 | 0.979 | 0.979 |
| BRD-K68395654-001-03-1 | EVP4593 | 0.983 | 287 | 0.686 | 0.983 | 1271 | 0.539 | 0.870 | 1839 | 0.289 | 0.620 | 3318 | 0.054 | 0.340 | 3196 | 0.065 | 0.340 | 2052 | 0.016 | 0.256 | 4921 | 0.837 | 0.983 |
| BRD-K68408467-065-01-8 | proflavine-hemisulfate | 0.838 | | | | 162 | 0.371 | 0.784 | 29 | 0.013 | 0.139 | 540 | 0.240 | 0.719 | 1036 | 0.910 | 0.999 | 362 | 0.076 | 0.319 | 1237 | 0.662 | 0.927 |
| BRD-K68408772-001-05-4 | carboxyamidotriazole | | | | | 118 | 0.819 | 0.987 | 174 | 0.239 | 1.000 | 1060 | 0.999 | 0.999 | 397 | 0.314 | 0.600 | 702 | 0.255 | 0.596 | 790 | 0.837 | 0.987 |
| BRD-K68488863-001-04-9 | ENMD-2076 | 0.749 | | | | 717 | 0.637 | 0.908 | 119 | 0.428 | 0.749 | 1585 | 0.142 | 0.498 | 1162 | 0.077 | 0.812 | 1213 | 0.789 | 0.975 | 1885 | 0.648 | 0.908 |
| BRD-K68502831-001-01-2 | dianhydrogalactitol | 0.436 | | | | 40 | 0.133 | 0.436 | 291 | 0.114 | 0.436 | 1960 | 0.842 | 1.000 | 672 | 0.779 | 1.000 | 970 | 0.143 | 0.436 | 1412 | 0.840 | 1.000 |
| BRD-K68532323-003-02-8 | SB-743921 | 1.000 | | | | 238 | 0.671 | 0.903 | 67 | 0.202 | 1.000 | 323 | 0.574 | 0.903 | 377 | 0.998 | 0.998 | 356 | 0.731 | 0.903 | 266 | 0.536 | 0.903 |
| BRD-K68552125-001-05-3 | 12-O-tetradecanoylphorbol-13-acetate | 0.978 | 30 | 0.224 | 0.784 | 69 | 0.682 | 0.978 | 121 | 0.619 | 0.978 | 20 | 0.141 | 0.593 | 157 | 0.478 | 0.917 | 408 | 0.899 | 0.978 | 244 | 0.398 | 0.917 |
| BRD-K68567222-001-01-2 | BVT-948 | 0.839 | | | | 174 | 0.543 | 0.966 | 494 | 0.783 | 1.000 | 482 | 0.116 | 0.487 | 504 | 0.311 | 0.839 | 264 | 0.089 | 0.487 | 1326 | 0.991 | 0.991 |
| BRD-K68633617-003-02-9 | ACDPP | | | | | 105 | 0.298 | 0.647 | 261 | 0.813 | 0.968 | 288 | 0.317 | 0.647 | 521 | 0.742 | 0.968 | 220 | 0.035 | 0.146 | 838 | 0.875 | 0.968 |
| BRD-K68747584-001-02-0 | PF-03814735 | 0.751 | 127 | 0.428 | 0.920 | 569 | 0.846 | 0.987 | 556 | 0.482 | 0.920 | 2193 | 0.542 | 0.925 | 1841 | 0.889 | 0.987 | 1426.5 | 0.573 | 0.925 | 2604.5 | 0.715 | 0.987 |
| BRD-K68904758-001-03-5 | cephalomannine | 1.000 | 5 | 0.014 | 0.142 | 1755 | 0.921 | 1.000 | 2126 | 0.462 | 0.881 | 3846 | 0.090 | 0.270 | 2180 | 0.001 | 0.021 | 3908 | 0.973 | 1.000 | 3971 | 0.076 | 0.270 |
| BRD-K68938568-001-01-7 | GDC-0152 | 0.995 | | | | | | | 454 | 0.391 | 1.000 | 1211 | 0.979 | 0.995 | 574 | 0.745 | 0.995 | 712.5 | 0.589 | 1.000 | 1640.5 | 0.995 | 0.995 |
| BRD-K69001009-001-02-8 | golvatinib | 0.814 | | | | | | | 343.5 | 0.841 | 0.982 | 532 | 0.560 | 0.784 | 152 | 0.006 | 0.119 | 185 | 0.152 | 0.398 | 444.5 | 0.461 | 0.692 |
| BRD-K69116396-001-04-1 | sertindole | 0.929 | | | | 426 | 0.636 | 0.929 | 432 | 0.653 | 0.929 | 1743 | 0.950 | 1.000 | 1386 | 0.752 | 0.929 | 1335 | 0.677 | 0.929 | 1609 | 0.076 | 0.725 |
| BRD-K69280563-001-01-8 | vinorelbine | 0.644 | 62 | 0.109 | 0.390 | 1892 | 0.974 | 1.000 | 2188 | 0.554 | 0.895 | 3117 | 0.006 | 0.128 | 3548 | 0.130 | 0.390 | 3683 | 0.940 | 1.000 | 3669 | 0.060 | 0.390 |
| BRD-K69608737-001-10-2 | tacrolimus | 0.603 | | | | | | | 164.5 | 0.489 | 1.000 | 314 | 0.452 | 1.000 | 112 | 0.169 | 1.000 | 220 | 0.035 | 0.072 | 444 | 0.937 | 0.962 |
| BRD-K69650333-003-14-0 | idarubicin | 0.750 | 194 | 0.499 | 0.873 | 1691 | 0.877 | 0.920 | 1228 | 0.172 | 0.691 | 3817 | 0.722 | 0.920 | 2889 | 0.159 | 0.691 | 1665 | 0.024 | 0.506 | 3431 | 0.286 | 0.750 |
| BRD-K69694239-001-02-2 | bardoxolone-methyl | 0.228 | 215 | 0.537 | 0.867 | 1463 | 0.831 | 1.000 | 1116 | 0.403 | 0.704 | 2655 | 0.005 | 0.099 | 3652 | 0.251 | 0.659 | 2555 | 0.354 | 0.675 | 4389 | 0.654 | 0.877 |
| BRD-K69726342-238-02-4 | atorvastatin | | 11 | 0.070 | 0.733 | 306 | 0.897 | 0.989 | 318 | 0.346 | 1.000 | 429 | 0.791 | 0.989 | 134 | 0.429 | 1.000 | 570 | 0.550 | 0.967 | 731 | 0.937 | 0.989 |
| BRD-K69776681-001-03-8 | volasertib | 0.064 | | | | 433.5 | 0.979 | 0.979 | 531.5 | 0.370 | 0.648 | 855.5 | 0.369 | 0.648 | 1223.5 | 0.285 | 1.000 | 611 | 0.136 | 0.476 | 884 | 0.115 | 1.000 |
| BRD-K69932463-001-10-6 | AZD8055 | 0.982 | 78 | 0.180 | 0.473 | 1597 | 0.832 | 0.982 | 2973 | 0.982 | 0.982 | 3604 | 0.149 | 0.473 | 3139 | 0.060 | 0.417 | 2113 | 0.014 | 0.147 | 4044 | 0.179 | 0.473 |
| BRD-K70163715-300-05-7 | vatalanib | 0.453 | 207.5 | 0.943 | 0.990 | 228 | 0.556 | 0.898 | 649.5 | 0.800 | 0.990 | 1359.5 | 0.146 | 0.576 | 988 | 0.802 | 0.990 | 515 | 0.451 | 0.862 | 2074 | 0.929 | 0.990 |
| BRD-K70301465-001-05-9 | ibrutinib | 1.000 | | | | 67 | 0.457 | 0.959 | 372 | 0.987 | 1.000 | 715 | 0.022 | 0.153 | 194 | 0.143 | 0.334 | 386 | 0.655 | 1.000 | 734 | 0.029 | 0.087 |
| BRD-K70401845-001-15-7 | erlotinib | | | | | 278 | 0.585 | 0.946 | 174.5 | 0.779 | 1.000 | 372 | 0.000 | 0.006 | 222 | 0.303 | 0.578 | 126 | 0.038 | 0.101 | 510 | 0.082 | 0.192 |
| BRD-K70401845-003-09-6 | erlotinib | 0.992 | | | | 568.5 | 0.943 | 0.992 | 318 | 0.752 | 0.992 | 1074 | 0.000 | 0.006 | 1281.5 | 0.268 | 0.625 | 1006 | 0.091 | 0.320 | 1687 | 0.045 | 0.313 |
| BRD-K70463136-001-01-5 | BAY-87-2243 | 0.941 | 314 | 0.805 | 0.941 | 1609 | 0.692 | 0.941 | 2451 | 0.852 | 0.941 | 3584 | 0.325 | 0.917 | 3146 | 0.171 | 0.717 | 2372 | 0.035 | 0.419 | 4611 | 0.524 | 0.917 |
| BRD-K70511574-001-06-9 | HMN-214 | 0.628 | | | | 608 | 0.606 | 0.863 | 1644 | 0.720 | 0.883 | 3369.5 | 0.149 | 1.000 | 2971.5 | 0.296 | 1.000 | 2620 | 0.757 | 0.883 | 3494 | 0.322 | 0.628 |
| BRD-K70914287-300-02-8 | BIBX-1382 | 1.000 | | | | 194 | 0.840 | 0.996 | 285 | 0.567 | 0.851 | 225 | 0.000 | 0.001 | 406 | 0.518 | 1.000 | 269 | 0.222 | 0.467 | 953 | 0.543 | 0.851 |
| BRD-K71106091-001-06-1 | fludarabine-phosphate | 0.893 | | | | 446 | 0.993 | 0.994 | 464.5 | 0.723 | 0.893 | 774 | 0.306 | 1.000 | 332 | 0.254 | 1.000 | 342 | 0.117 | 0.252 | 871.5 | 0.994 | 0.994 |
| BRD-K71164191-001-01-0 | KD-023 | 0.926 | 232 | 0.866 | 0.958 | 417 | 0.489 | 0.926 | 533 | 0.184 | 0.926 | 2576 | 0.706 | 0.926 | 1498 | 0.182 | 0.926 | 1292 | 0.590 | 0.926 | 2874 | 0.967 | 0.993 |
| BRD-K71221037-001-01-6 | birinapant | 0.998 | 128 | 0.662 | 0.963 | | | | 215 | 0.545 | 0.963 | 1450 | 0.809 | 0.963 | 1079 | 0.686 | 0.963 | 697 | 0.736 | 0.963 | 1773 | 0.992 | 0.998 |
| BRD-K71281111-001-04-3 | SNX-2112 | 0.474 | 63 | 0.138 | 0.474 | 1928 | 0.821 | 0.958 | 1849 | 0.158 | 0.474 | 3631 | 0.045 | 0.235 | 4167 | 0.321 | 0.702 | 3654 | 0.553 | 0.894 | 4790 | 0.222 | 0.583 |
| BRD-K71467466-001-02-3 | KPT-185 | 0.991 | 38 | 0.088 | 0.371 | 1155 | 0.585 | 0.991 | 1876 | 0.425 | 0.976 | 3613 | 0.042 | 0.223 | 4480 | 0.895 | 0.991 | 4335 | 0.968 | 0.991 | 3195 | 0.003 | 0.068 |
| BRD-K71480163-001-01-4 | GDC-0068 | 0.957 | | | | 492 | 0.926 | 0.977 | 93 | 0.390 | 0.911 | 927 | 0.638 | 0.957 | 992 | 0.000 | 0.004 | 997 | 0.921 | 0.977 | 1618 | 0.862 | 0.977 |
| BRD-K71512533-066-02-8 | SNS-314 | 0.553 | 179 | 0.608 | 0.951 | 622 | 0.850 | 0.951 | 1257 | 0.572 | 0.951 | 2686.5 | 0.554 | 0.951 | 1835.5 | 0.882 | 0.951 | 1612.5 | 0.650 | 0.951 | 2982 | 0.825 | 0.951 |
| BRD-K71823332-001-03-7 | epothilone-a | 0.340 | 407 | 0.907 | 0.999 | 1877 | 0.827 | 0.999 | 1980 | 0.295 | 0.786 | 2783 | 0.001 | 0.013 | 3538 | 0.048 | 0.339 | 3129 | 0.564 | 0.846 | 4256 | 0.141 | 0.592 |
| BRD-K72093121-001-16-3 | vidarabine | 1.000 | | | | | | | 283.5 | 0.818 | 1.000 | 269 | 0.097 | 1.000 | 255 | 0.177 | 1.000 | 103 | 0.915 | 1.000 | 426 | 0.285 | 1.000 |
| BRD-K72167406-001-02-1 | cefditoren-pivoxil | 0.767 | | | | 178 | 0.554 | 0.970 | 339 | 0.849 | 0.999 | 520 | 0.168 | 0.767 | 286 | 0.012 | 0.131 | 589 | 0.786 | 0.999 | 1129 | 0.856 | 0.999 |
| BRD-K72414522-001-06-7 | AZD5438 | 0.982 | | | | 819 | 0.654 | 0.982 | 2164 | 0.956 | 0.982 | 3897 | 0.950 | 0.982 | 3443 | 0.733 | 0.982 | 1917 | 0.144 | 0.603 | 3145 | 0.339 | 0.790 |
| BRD-K72420232-001-12-3 | WZ-4002 | 0.688 | | | | 454 | 0.936 | 0.987 | 108 | 0.860 | 0.987 | 520 | 0.084 | 1.000 | 266 | 0.233 | 0.537 | 90 | 0.002 | 0.012 | 582 | 0.193 | 1.000 |
| BRD-K72533376-001-01-2 | etoposide-phosphate | 0.318 | 46 | 0.232 | 0.442 | | | | 505 | 0.978 | 0.991 | 1109 | 0.991 | 0.991 | 466 | 0.119 | 1.000 | 113 | 0.014 | 0.042 | 667 | 0.840 | 0.991 |
| BRD-K72703948-001-10-1 | ZM-447439 | 0.722 | | | | 949 | 0.902 | 0.997 | 711 | 0.503 | 0.755 | 2425 | 0.771 | 0.997 | 1736 | 0.268 | 1.000 | 1489 | 0.442 | 0.755 | 2227 | 0.400 | 0.755 |
| BRD-K72723676-003-17-8 | benzethonium | 0.910 | 284 | 0.841 | 0.981 | 936 | 0.675 | 0.956 | 995 | 0.262 | 0.787 | 2005 | 0.372 | 0.910 | 2801 | 0.565 | 0.913 | 2080 | 0.806 | 0.981 | 2367 | 0.433 | 0.910 |
| BRD-K72726508-001-02-1 | arcyriaflavin-a | 0.548 | | | | | | | 125 | 0.520 | 0.840 | 274 | 0.934 | 0.981 | 102.5 | 0.877 | 0.981 | 472.5 | 0.745 | 0.978 | 330 | 0.377 | 0.660 |
| BRD-K72815923-034-01-2 | xanomeline | 1.000 | | | | 219 | 0.948 | 1.000 | 568 | 0.276 | 0.724 | 774 | 0.063 | 0.331 | 1336 | 0.892 | 1.000 | 589 | 0.865 | 1.000 | 1102 | 0.199 | 0.596 |
| BRD-K72827473-001-01-0 | CEP-37440 | 0.717 | 81 | 0.307 | 0.717 | 750 | 0.999 | 0.999 | 638 | 0.225 | 0.711 | 1072 | 0.002 | 0.051 | 2445 | 0.992 | 0.999 | 832 | 0.175 | 0.711 | 2295 | 0.710 | 0.999 |
| BRD-K72951360-001-01-4 | valrubicin | 0.870 | 252 | 0.580 | 0.870 | 1979 | 0.924 | 0.995 | 2146 | 0.517 | 0.870 | 3827 | 0.402 | 0.843 | 3671 | 0.112 | 0.451 | 2325 | 0.172 | 0.451 | 5233 | 0.795 | 0.995 |
| BRD-K73073971-001-01-7 | ZLN005 | 0.685 | | | | 151 | 0.754 | 0.970 | 76 | 0.690 | 0.965 | 231 | 0.000 | 0.003 | 168 | 0.318 | 1.000 | 298 | 0.834 | 0.970 | 461 | 0.373 | 0.871 |
| BRD-K73197500-001-02-2 | SNX-5422 | 0.618 | 56 | 0.147 | 0.495 | 1190 | 0.820 | 0.984 | 1184 | 0.146 | 0.495 | 2389 | 0.004 | 0.087 | 2912 | 0.184 | 0.495 | 1930 | 0.197 | 0.495 | 3278 | 0.206 | 0.495 |
| BRD-K73293050-001-03-1 | WZ-3146 | 0.607 | 333 | 0.997 | 0.999 | 740 | 0.351 | 0.626 | 1949 | 0.992 | 0.999 | 2005 | 0.037 | 0.383 | 2273 | 0.231 | 0.607 | 1567 | 0.637 | 0.892 | 2916 | 0.289 | 0.607 |
| BRD-K73309154-003-02-8 | OSI-420 | 0.994 | 180 | 0.805 | 0.994 | 611 | 0.982 | 0.994 | 641 | 0.883 | 0.994 | 1148 | 0.001 | 0.013 | 866 | 0.361 | 0.974 | 895 | 0.418 | 0.974 | 1763 | 0.064 | 0.448 |
| BRD-K73753850-003-01-6 | hexaminolevulinate | 0.723 | 250 | 0.768 | 0.892 | 803 | 0.497 | 0.746 | 898 | 0.392 | 0.746 | 3700 | 0.999 | 0.999 | 1782 | 0.219 | 0.723 | 1401 | 0.064 | 0.675 | 2565 | 0.427 | 0.746 |
| BRD-K73838513-003-05-5 | cinacalcet | 0.886 | 161 | 0.738 | 0.886 | 563 | 0.763 | 0.886 | 353 | 0.508 | 0.886 | 1054 | 0.047 | 0.498 | 884 | 0.672 | 0.886 | 494 | 0.014 | 0.284 | 1936 | 0.596 | 0.886 |
| BRD-K73982490-001-02-9 | BI-78D3 | 0.275 | | | | 713 | 0.805 | 1.000 | 1001 | 0.371 | 0.708 | 3335 | 0.788 | 1.000 | 2431 | 0.290 | 0.610 | 2180 | 0.218 | 0.610 | 4653 | 0.982 | 0.982 |
| BRD-K74057757-001-01-9 | indibulin | 0.630 | 131 | 0.250 | 0.739 | 1558 | 0.819 | 0.974 | 2230 | 0.807 | 0.974 | 3163 | 0.019 | 0.132 | 3944 | 0.367 | 0.739 | 2585 | 0.373 | 0.739 | 3637 | 0.016 | 0.132 |
| BRD-K74141488-003-11-2 | naftifine | 0.038 | | | | | | | 240 | 0.562 | 1.000 | 437.5 | 0.699 | 0.850 | 350.5 | 0.809 | 0.850 | 283.5 | 0.787 | 0.850 | 404.5 | 0.789 | 0.850 |
| BRD-K74193284-001-04-8 | phenylmercuric-acetate | 0.883 | 127 | 0.352 | 0.821 | 792 | 0.631 | 0.883 | 1217 | 0.111 | 0.390 | 3559 | 0.736 | 0.909 | 2851 | 0.409 | 0.858 | 1454 | 0.033 | 0.233 | 3629 | 0.524 | 0.883 |
| BRD-K74305673-001-06-2 | IKK-2-inhibitor-V | 0.172 | | | | 1451 | 0.951 | 0.990 | 868 | 0.100 | 1.000 | 3387 | 0.902 | 0.990 | 2697 | 0.350 | 1.000 | 1783 | 0.385 | 0.577 | 3842 | 0.784 | 0.915 |
| BRD-K74501079-001-18-1 | azithromycin | 0.909 | | | | 878 | 0.598 | 0.898 | 700 | 0.264 | 1.000 | 1889 | 0.226 | 1.000 | 1189 | 0.250 | 1.000 | 547 | 0.080 | 0.186 | 2472 | 0.978 | 0.978 |
| BRD-K74514084-001-03-9 | pazopanib | 0.943 | 11 | 0.078 | 0.629 | 62 | 0.542 | 0.926 | 177 | 0.213 | 0.848 | 255 | 0.922 | 0.943 | 299 | 0.057 | 0.629 | 256 | 0.656 | 0.926 | 360 | 0.357 | 0.848 |
| BRD-K74671368-001-01-8 | idronoxil | 0.387 | 19 | 0.076 | 0.387 | 584 | 0.979 | 0.998 | 632 | 0.377 | 0.792 | 1058 | 0.933 | 0.998 | 1053 | 0.111 | 0.390 | 493 | 0.005 | 0.096 | 1253 | 0.887 | 0.998 |
| BRD-K75009076-001-02-1 | SCH-900776 | 0.739 | 9 | 0.029 | 0.306 | 1219 | 0.802 | 0.901 | 1341 | 0.734 | 0.901 | 3432 | 0.982 | 0.982 | 2730 | 0.764 | 0.901 | 1387 | 0.185 | 0.646 | 2977 | 0.279 | 0.739 |
| BRD-K75295174-001-05-0 | alisertib | 0.369 | 104 | 0.302 | 0.773 | 402 | 0.804 | 0.954 | 902 | 0.539 | 0.954 | 2423 | 0.251 | 0.752 | 2268 | 0.800 | 0.954 | 1248 | 0.630 | 0.954 | 2567 | 0.680 | 0.954 |
| BRD-K75352575-001-01-3 | KPT-276 | 0.956 | 102 | 0.192 | 0.672 | 999 | 0.466 | 0.956 | 2489 | 0.895 | 0.956 | 3183 | 0.074 | 0.309 | 4087 | 0.930 | 0.956 | 3237 | 0.931 | 0.956 | 3185 | 0.009 | 0.160 |
| BRD-K75649340-001-13-1 | tioguanine | 0.301 | 268.5 | 0.949 | 0.999 | 551 | 0.514 | 0.878 | 842 | 0.148 | 0.438 | 2574 | 0.978 | 0.999 | 2414 | 0.975 | 0.999 | 994 | 0.185 | 0.438 | 2366.5 | 0.544 | 0.878 |
| BRD-K75911534-001-01-6 | SU014813 | 0.969 | 9 | 0.057 | 0.199 | 277 | 0.295 | 0.688 | 586 | 0.053 | 0.199 | 1779 | 0.947 | 0.969 | 563 | 0.108 | 0.324 | 912 | 0.936 | 0.969 | 1836.5 | 0.969 | 0.969 |
| BRD-K75958547-238-01-0 | pitavastatin | 0.928 | | | | 508 | 0.818 | 0.928 | 221 | 0.588 | 1.000 | 992 | 0.893 | 0.928 | 749 | 0.643 | 1.000 | 438 | 0.134 | 0.352 | 755 | 0.422 | 1.000 |
| BRD-K76064317-001-07-3 | tyrphostin-AG-1296 | 0.828 | 87 | 0.298 | 0.718 | 588 | 0.839 | 0.979 | 784 | 0.349 | 0.718 | 2598 | 0.340 | 0.718 | 1874 | 0.242 | 0.718 | 1207 | 0.376 | 0.718 | 2270 | 0.546 | 0.828 |
| BRD-K76180386-001-01-6 | JIB04 | 0.743 | 240 | 0.814 | 0.915 | 562 | 0.715 | 0.915 | 1531 | 0.872 | 0.915 | 2669 | 0.760 | 0.915 | 1994 | 0.694 | 0.915 | 1865 | 0.218 | 0.743 | 3206.5 | 0.986 | 0.986 |
| BRD-K76239644-001-01-8 | BMS-690514 | 1.000 | | | | 837 | 0.346 | 0.661 | 898 | 0.935 | 1.000 | 1871 | 0.007 | 0.038 | 1400 | 0.089 | 0.306 | 967 | 0.156 | 0.364 | 1989 | 0.001 | 0.012 |
| BRD-K76239644-001-02-6 | BMS-690514 | | | | | 274 | 0.494 | 0.740 | 521.5 | 0.928 | 1.000 | 982 | 0.031 | 0.222 | 600 | 0.220 | 0.513 | 445.5 | 0.470 | 0.740 | 950 | 0.022 | 0.222 |
| BRD-K76406695-001-02-9 | CCT128930 | 0.993 | | | | 76 | 0.538 | 0.878 | 292.5 | 0.861 | 0.993 | 313 | 0.002 | 0.023 | 176 | 0.002 | 0.023 | 52 | 0.061 | 0.161 | 876.5 | 0.699 | 0.993 |
| BRD-K76634210-001-14-7 | idoxuridine | 0.828 | 144 | 0.898 | 0.989 | | | | 248 | 0.505 | 0.828 | 832 | 0.420 | 0.828 | 644 | 0.915 | 0.989 | 328 | 0.502 | 0.828 | 1235 | 0.969 | 0.989 |
| BRD-K76674262-001-03-3 | homoharringtonine | 0.903 | 324 | 0.904 | 0.998 | 1021 | 0.911 | 0.998 | 2327 | 0.998 | 0.998 | 2399 | 0.040 | 0.419 | 3026 | 0.150 | 0.570 | 1217 | 0.217 | 0.570 | 3024 | 0.086 | 0.486 |
| BRD-K76703230-004-04-1 | YM-155 | 0.997 | 73 | 0.164 | 0.573 | 1691 | 0.863 | 0.997 | 1917 | 0.237 | 0.711 | 6044 | 0.986 | 0.997 | 2094 | 0.000 | 0.001 | 2884 | 0.353 | 0.824 | 6532 | 0.996 | 0.997 |
| BRD-K76840893-003-02-5 | RS-17053 | 0.913 | 228 | 0.751 | 0.913 | 578 | 0.449 | 0.913 | 1247 | 0.594 | 0.913 | 2534 | 0.741 | 0.913 | 1858 | 0.290 | 0.870 | 1576 | 0.826 | 0.913 | 1939 | 0.240 | 0.870 |
| BRD-K76973696-001-12-6 | tryptanthrin | 0.997 | 59 | 0.153 | 0.472 | 1541 | 0.991 | 0.997 | 997 | 0.124 | 0.472 | 3332 | 0.479 | 0.914 | 3162 | 0.788 | 0.973 | 2037 | 0.074 | 0.472 | 3993 | 0.718 | 0.973 |
| BRD-K77008974-001-03-2 | WYE-354 | 0.981 | 259 | 0.889 | 0.981 | 681 | 0.363 | 0.847 | 724 | 0.963 | 0.981 | 2486.5 | 0.544 | 0.981 | 1830 | 0.261 | 0.689 | 1412 | 0.124 | 0.615 | 3071 | 0.830 | 0.981 |
| BRD-K77625799-001-07-7 | vandetanib | 1.000 | | | | | | | 162 | 0.200 | 0.923 | 259 | 0.002 | 0.039 | 147 | 0.045 | 0.472 | 163 | 0.204 | 0.429 | 708 | 0.847 | 0.847 |
| BRD-K77634909-001-03-5 | AEG3482 | 0.503 | | | | | | | 147 | 0.160 | 1.000 | 593 | 0.049 | 0.861 | 519.5 | 0.791 | 0.944 | 105 | 0.239 | 0.503 | 461.5 | 0.082 | 0.861 |
| BRD-K77638923-001-02-0 | CUDC-907 | 0.434 | 410 | 0.886 | 0.999 | 1821 | 0.732 | 0.955 | 2262 | 0.493 | 0.690 | 4780 | 0.447 | 0.690 | 3829 | 0.085 | 0.434 | 3573 | 0.492 | 0.690 | 4898 | 0.275 | 0.681 |
| BRD-K77677632-003-02-6 | SB-200646 | 0.307 | | | | 154 | 0.936 | 0.979 | 805 | 0.979 | 0.979 | 767 | 0.302 | 1.000 | 358 | 0.346 | 1.000 | 261 | 0.929 | 0.979 | 1068 | 0.926 | 0.979 |
| BRD-K77685957-300-09-1 | dequalinium | 0.508 | | | | 251 | 0.945 | 0.995 | 884 | 0.267 | 1.000 | 1640 | 0.228 | 1.000 | 1466 | 0.448 | 0.784 | 767 | 0.264 | 0.554 | 2036 | 0.304 | 0.579 |
| BRD-K77791657-003-01-7 | 3-deazaneplanocin-A | 0.850 | | | | 845.5 | 0.752 | 0.920 | 1240 | 0.523 | 0.850 | 2343 | 0.094 | 1.000 | 2657 | 0.876 | 0.920 | 1617 | 0.567 | 0.850 | 2932 | 0.262 | 1.000 |
| BRD-K77841042-001-14-1 | brefeldin-a | 0.991 | 291 | 0.658 | 0.991 | 1917 | 0.974 | 0.991 | 1701 | 0.121 | 0.510 | 4112 | 0.183 | 0.550 | 4264 | 0.790 | 0.991 | 2837 | 0.218 | 0.573 | 5355 | 0.888 | 0.991 |
| BRD-K77908580-001-09-6 | entinostat | 0.159 | 128 | 0.383 | 0.670 | 781 | 0.718 | 0.984 | 1308 | 0.489 | 0.790 | 2663 | 0.750 | 0.984 | 2041 | 0.042 | 0.291 | 1762 | 0.286 | 0.606 | 3059 | 0.680 | 0.984 |
| BRD-K77943374-001-02-8 | NVP-BVU972 | 0.939 | 64 | 0.695 | 1.000 | 64 | 0.695 | 1.000 | 64 | 0.695 | 1.000 | 140 | 0.056 | 0.547 | 189 | 0.807 | 0.909 | 255 | 0.163 | 0.686 | 587 | 0.786 | 1.000 |
| BRD-K77987382-001-13-2 | mebendazole | 1.000 | | | | 657 | 0.713 | 1.000 | 671 | 0.440 | 1.000 | 1155 | 0.042 | 0.303 | 1270 | 0.043 | 0.303 | 1907 | 0.932 | 1.000 | 1860 | 0.103 | 0.431 |
| BRD-K78055238-001-01-8 | LY2090314 | 0.837 | 28 | 0.083 | 0.406 | 796 | 0.919 | 0.987 | 667 | 0.419 | 0.837 | 1737 | 0.976 | 0.987 | 859 | 0.021 | 0.145 | 626 | 0.017 | 0.145 | 2115 | 0.951 | 0.987 |
| BRD-K78096648-001-05-7 | oxiperomide | 0.922 | | | | 613 | 0.993 | 0.993 | 760 | 0.892 | 0.993 | 1829 | 0.628 | 0.943 | 1309 | 0.817 | 0.993 | 779 | 0.467 | 0.922 | 1298 | 0.629 | 0.943 |
| BRD-K78126613-001-28-5 | menadione | 0.929 | 156 | 0.646 | 0.929 | 169 | 0.700 | 0.929 | 531 | 0.644 | 0.929 | 1840 | 0.690 | 0.929 | 813 | 0.053 | 0.437 | 685 | 0.226 | 0.790 | 1499 | 0.343 | 0.929 |
| BRD-K78177893-001-02-4 | A-674563 | 0.874 | 369 | 0.987 | 0.987 | 834 | 0.661 | 0.987 | 1902 | 0.923 | 0.987 | 3532 | 0.485 | 0.987 | 3154 | 0.311 | 0.987 | 2485 | 0.458 | 0.987 | 3524 | 0.865 | 0.987 |
| BRD-K78318619-001-02-7 | PF-4981517 | 0.635 | | | | 218 | 0.964 | 0.964 | 112.5 | 0.400 | 1.000 | 476 | 0.289 | 1.000 | 345 | 0.204 | 1.000 | 198 | 0.570 | 0.845 | 629 | 0.812 | 0.927 |
| BRD-K78431006-001-10-2 | crizotinib | 0.815 | 133 | 0.395 | 0.815 | 644 | 0.788 | 0.919 | 524 | 0.079 | 0.815 | 2257 | 0.197 | 0.815 | 2605 | 0.522 | 0.831 | 1767 | 0.835 | 0.923 | 3043 | 0.365 | 0.815 |

| broad_id | name | colorectal_MWU_STAT | colorectal_PVAL | colorectal_PVAL_ADJ | skin_MWU_STAT | skin_PVAL | skin_PVAL_ADJ | urinary_tract_MWU_STAT | urinary_tract_PVAL | urinary_tract_PVAL_ADJ | lung_MWU_STAT | lung_PVAL | lung_PVAL_ADJ | ovary_MWU_STAT | ovary_PVAL | ovary_PVAL_ADJ | breast_MWU_STAT | breast_PVAL | breast_PVAL_ADJ | pancreas_MWU_STAT | pancreas_PVAL | pancreas_PVAL_ADJ | nervous_system_MWU_STAT | central_nervous_system_PVAL | nervous_system_PVAL_ADJ | soft_tissue_MWU_STAT | soft_tissue_PVAL | soft_tissue_PVAL_ADJ | kidney_MWU_STAT | kidney_PVAL | kidney_PVAL_ADJ | gastric_MWU_STAT | gastric_PVAL | gastric_PVAL_ADJ | bone_MWU_STAT | bone_PVAL | bone_PVAL_ADJ | prostate_MWU_STAT | prostate_PVAL | prostate_PVAL_ADJ | al_nervous_system_MWU_STAT | peripheral_nervous_system_PVAL |
|---|---|---|---|---|---|---|---|---|---|---|---|---|---|---|---|---|---|---|---|---|---|---|---|---|---|---|---|---|---|---|---|---|---|---|---|---|---|---|---|---|---|---|
| BRD-K78431006-001-15-1 | crizotinib | 263 | 0.508 | 0.932 | 1139 | 0.899 | 0.940 | 336 | 0.021 | 0.440 | 1413 | 0.769 | 0.940 | 703 | 0.886 | 0.940 | 410 | 0.841 | 0.940 | 351 | 0.621 | 0.932 | 449 | 0.293 | 0.615 | 229 | 0.015 | 0.045 | 274 | 0.278 | 0.615 | 273 | 0.559 | 0.932 | 283 | 0.610 | 0.932 | 127 | 0.940 | 0.940 | 1 | 0.046 |
| BRD-K78567475-001-01-4 | dolastatin-10 | 5145 | 0.613 | 0.871 | 7995 | 0.959 | 0.999 | 3043 | 0.064 | 0.511 | 13949 | 0.090 | 0.511 | 5132 | 0.364 | 0.637 | 4293 | 0.809 | 0.999 | 6795 | 0.991 | 0.999 | 5582 | 0.419 | 0.676 | 1942 | 0.209 | 0.611 | 4753 | 0.999 | 0.999 | 1930 | 0.019 | 0.401 | 2226 | 0.283 | 0.611 | 746 | 0.358 | 0.637 | 574 | 0.146 |
| BRD-K78659596-001-03-9 | ixazomib | 3617 | 0.326 | 0.651 | 5861 | 0.120 | 0.504 | 2788 | 0.341 | 0.651 | 13482 | 0.861 | 0.952 | 3702 | 0.109 | 0.504 | 4507 | 0.957 | 0.996 | 6360 | 0.996 | 0.996 | 3353 | 0.004 | 0.076 | 1420 | 0.619 | 0.952 | 2437 | 0.336 | 0.651 | 2380 | 0.287 | 0.651 | 1044 | 0.067 | 0.504 | 884 | 0.696 | 0.952 | 630 | 0.099 |
| BRD-K78815826-001-14-6 | suxibuzone | 438 | 0.858 | 1.000 | 693 | 0.261 | 0.784 | 327 | 0.084 | 0.509 | 1508 | 0.254 | 0.784 | 730 | 0.357 | 0.891 | 512.5 | 0.858 | 0.990 | 831.5 | 0.804 | 1.000 | 365.5 | 0.156 | 0.656 | 37 | 0.037 | 0.388 | 315 | 0.382 | 0.891 | 573 | 0.631 | 1.000 | 405.5 | 0.947 | 0.990 | 65 | 0.097 | 0.509 | 101 | 0.777 |
| BRD-K78960041-001-05-7 | oxaliplatin | 1702 | 0.127 | 0.532 | 3026 | 0.954 | 0.998 | 1118 | 0.081 | 0.425 | 5819 | 0.757 | 0.998 | 1869 | 0.177 | 0.532 | 1517 | 0.575 | 0.909 | 2720 | 0.969 | 0.998 | 2183 | 0.402 | 0.767 | 368 | 0.016 | 0.114 | 856 | 0.250 | 0.584 | 748 | 0.002 | 0.043 | 289 | 0.237 | 0.584 | 415 | 0.606 | 0.909 | 390 | 0.902 |
| BRD-K79131256-001-17-9 | albendazole | 1469 | 0.133 | 0.534 | 2855 | 0.929 | 0.974 | 1566 | 0.220 | 0.534 | 5742 | 0.160 | 0.534 | 2241 | 0.229 | 0.534 | 1909 | 0.974 | 0.974 | 2992 | 0.971 | 0.974 | 2699 | 0.964 | 0.974 | 865 | 0.400 | 0.708 | 967 | 0.963 | 0.974 | 1090 | 0.198 | 0.534 | 1010 | 0.439 | 0.708 | 131 | 0.487 | 0.731 | 519 | 0.207 |
| BRD-K79239947-001-06-8 | PF-573228 | 747.5 | 0.776 | 0.940 | 1102 | 0.241 | 1.000 | 719 | 0.703 | 0.925 | 2409 | 0.927 | 0.940 | 622.5 | 0.232 | 1.000 | 458.5 | 0.557 | 0.836 | 280 | 0.410 | 0.713 | 860.5 | 0.705 | 0.925 | 230.5 | 0.210 | 0.550 | 132.5 | 0.377 | 0.713 | 599.5 | 0.182 | 1.000 | 33 | 0.005 | 0.014 | 246 | 0.940 | 0.940 | 142.5 | 0.441 |
| BRD-K79254416-001-21-8 | decitabine | 633 | 0.580 | 0.870 | 753 | 0.327 | 0.772 | 658 | 0.257 | 0.772 | 2122 | 0.467 | 0.811 | 482 | 0.070 | 0.492 | 660 | 0.444 | 0.811 | 854 | 0.905 | 0.974 | 1368 | 0.983 | 0.983 | 96 | 0.170 | 0.699 | 208 | 0.127 | 0.665 | 435 | 0.200 | 0.699 | 312 | 0.502 | 0.811 | 59 | 0.331 | 0.772 | 305 | 0.813 |
| BRD-K79254416-001-22-6 | decitabine | 2192 | 0.379 | 0.795 | 2516 | 0.748 | 0.866 | 1833 | 0.150 | 0.794 | 7355 | 0.594 | 0.866 | 2405 | 0.344 | 0.795 | 1665 | 0.209 | 0.794 | 2628 | 0.454 | 0.866 | 2852.5 | 0.983 | 0.983 | 596 | 0.110 | 0.794 | 1443 | 0.784 | 0.866 | 1378 | 0.303 | 0.794 | 1110 | 0.726 | 0.866 | 188 | 0.205 | 0.794 | 476 | 0.294 |
| BRD-K79259477-001-04-9 | NVP-BHG712 | 468 | 0.315 | 0.609 | 957.5 | 0.857 | 0.983 | 637 | 0.128 | 0.622 | 2724.5 | 0.998 | 0.998 | 369 | 0.001 | 0.030 | 444 | 0.469 | 0.758 | 1010 | 0.682 | 0.983 | 853 | 0.922 | 0.983 | 442.5 | 0.937 | 0.983 | 411 | 0.167 | 0.411 | 191 | 0.030 | 0.208 | 20 | 0.102 | 0.307 | 146 | 0.935 | 0.983 | 136 | 0.386 |
| BRD-K79584249-001-01-3 | CGM097 | 139 | 0.170 | 1.000 | 300 | 0.039 | 0.811 | 130 | 0.370 | 0.578 | 1024 | 0.967 | 0.994 | 462 | 0.361 | 0.578 | 132 | 0.385 | 0.578 | 139 | 0.840 | 0.980 | 221 | 0.074 | 0.194 | 190 | 0.476 | 0.666 | 254 | 0.158 | 0.369 | 285 | 0.279 | 0.578 | 243 | 0.522 | 0.685 | | | | 21 | 0.163 |
| BRD-K79612754-001-19-1 | inosine | 1984 | 0.577 | 0.895 | 4008.5 | 0.927 | 0.973 | 1707 | 0.071 | 0.320 | 7097 | 0.805 | 0.973 | 2495 | 0.071 | 0.320 | 2521 | 0.851 | 0.973 | 1284 | 0.091 | 0.320 | 3339.5 | 0.858 | 0.973 | 1040 | 0.597 | 0.895 | 1571 | 0.562 | 0.895 | 1545 | 0.199 | 0.465 | 489 | 0.314 | 0.659 | 153 | 0.129 | 0.337 | 347 | 0.090 |
| BRD-K79633074-001-01-8 | xilobam | 1400.5 | 0.681 | 0.953 | 2145 | 0.333 | 0.778 | 2068.5 | 0.981 | 1.000 | 3058 | 0.281 | 0.778 | 946.5 | 0.205 | 0.778 | 1052 | 0.578 | 0.953 | 1378.5 | 0.643 | 0.953 | 1137.5 | 0.126 | 0.778 | 330.5 | 0.238 | 0.778 | 929 | 0.326 | 0.778 | 1078.5 | 0.436 | 0.916 | 119 | 0.032 | 0.682 | 184 | 0.899 | 1.000 | 344 | 0.610 |
| BRD-K79821389-001-03-5 | rubitecan | 5973 | 0.784 | 0.969 | 10378 | 0.998 | 0.998 | 3773 | 0.040 | 0.213 | 17104 | 0.200 | 0.420 | 4973 | 0.030 | 0.213 | 6135 | 0.983 | 0.998 | 8313 | 0.975 | 0.998 | 7204 | 0.767 | 0.969 | 1646 | 0.028 | 0.213 | 4037 | 0.682 | 0.969 | 2966 | 0.065 | 0.275 | 2156 | 0.117 | 0.372 | 623 | 0.142 | 0.372 | 797 | 0.127 |
| BRD-K79930101-300-03-7 | GW-583340 | 639 | 0.070 | 0.246 | 1194 | 0.903 | 0.995 | 306 | 0.000 | 0.005 | 3176 | 0.833 | 0.995 | 1228 | 0.698 | 0.995 | 590 | 0.872 | 0.995 | 1345 | 0.609 | 0.995 | 1309 | 0.999 | 0.999 | 526 | 0.933 | 0.995 | 568 | 0.578 | 0.995 | 601 | 0.044 | 0.310 | 535 | 0.943 | 0.995 | 218 | 0.670 | 0.995 | 180 | 0.948 |
| BRD-K80060353-303-03-9 | IEM1754 | 531 | 0.685 | 0.890 | 998 | 0.847 | 0.890 | 761 | 0.764 | 0.890 | 1609.5 | 0.387 | 1.000 | 661 | 0.173 | 1.000 | 444 | 0.620 | 1.000 | 364 | 0.574 | 1.000 | 780.5 | 0.924 | 0.924 | 199 | 0.433 | 1.000 | 70 | 0.490 | 1.000 | 443 | 0.369 | 1.000 | 98 | 0.742 | 0.890 | 178 | 0.733 | 0.890 | 87 | 0.176 |
| BRD-K80343549-001-02-6 | TAK-285 | 1150.5 | 0.891 | 0.993 | 1430 | 0.846 | 0.993 | 776 | 0.097 | 0.408 | 3444.5 | 0.301 | 0.616 | 1589.5 | 0.352 | 0.616 | 610 | 0.233 | 0.569 | 2092.5 | 0.898 | 0.993 | 2742.5 | 1.000 | 1.000 | 757.5 | 0.810 | 0.993 | 1138 | 0.965 | 1.000 | 619.5 | 0.017 | 0.179 | 331.5 | 0.552 | 0.892 | 266.5 | 0.732 | 0.993 | 152 | 0.244 |
| BRD-K80431395-001-07-3 | triciribine | 3348 | 0.888 | 1.000 | 5430 | 0.988 | 1.000 | 2366 | 0.108 | 0.455 | 10335 | 0.808 | 0.998 | 3801 | 0.550 | 0.722 | 2432 | 0.237 | 0.623 | 3243 | 0.138 | 0.481 | 5409 | 1.000 | 1.000 | 634 | 0.005 | 0.061 | 2353 | 0.963 | 1.000 | 1338 | 0.033 | 0.234 | 746 | 0.358 | 0.537 | 378 | 0.245 | 0.428 | 633 | 0.439 |
| BRD-K80480517-001-02-4 | repsox | 1079 | 0.858 | 0.968 | 1617.5 | 0.956 | 0.968 | 647 | 0.005 | 0.099 | 3995 | 0.793 | 0.968 | 986 | 0.094 | 0.397 | 1375.5 | 0.709 | 0.968 | 1394 | 0.580 | 0.968 | 1653.5 | 0.919 | 0.968 | 505.5 | 0.801 | 0.968 | 684 | 0.251 | 0.878 | 655 | 0.388 | 0.968 | 319.5 | 0.554 | 0.968 | 70 | 0.293 | 0.879 | 124 | 0.641 |
| BRD-K80608265-001-01-6 | erteberel | 254 | 0.205 | 0.465 | 1121 | 0.234 | 0.465 | 454 | 0.309 | 0.499 | 1526 | 0.869 | 0.960 | 793 | 0.350 | 0.524 | 328 | 0.266 | 0.465 | 325 | 0.255 | 0.465 | 749 | 0.853 | 0.960 | 85 | 0.683 | 0.896 | 76 | 0.596 | 0.835 | 7 | 0.063 | 1.000 | 81 | 0.174 | 0.456 | | | | 5 | 0.057 |
| BRD-K80700417-001-04-2 | SGI-1776 | 842 | 0.282 | 0.845 | 1236 | 0.116 | 0.809 | 957 | 0.383 | 0.934 | 2010 | 0.251 | 0.845 | 546 | 0.446 | 0.934 | 501 | 0.965 | 1.000 | 595 | 0.095 | 0.809 | 766 | 0.641 | 0.934 | 93 | 0.582 | 0.934 | 283 | 0.667 | 0.934 | 603 | 0.843 | 1.000 | 301 | 0.977 | 1.000 | 257 | 0.548 | 0.934 | 359 | 0.629 |
| BRD-K80725821-300-02-9 | RS-16566 | 813 | 0.032 | 0.539 | 2401 | 0.959 | 0.959 | 1443 | 0.707 | 0.869 | 4098 | 0.330 | 0.770 | 1839 | 0.518 | 0.869 | 1031 | 0.766 | 0.869 | 1980 | 0.212 | 0.555 | 1757 | 0.697 | 0.869 | 171 | 0.068 | 0.358 | 607 | 0.152 | 0.476 | 1050 | 0.594 | 0.869 | 119 | 0.121 | 0.476 | 135 | 0.159 | 0.476 | 432 | 0.780 |
| BRD-K81197548-003-01-4 | NSC-632839 | 809 | 0.942 | 0.942 | 1370 | 0.300 | 1.000 | 955.5 | 0.491 | 1.000 | 3515 | 0.359 | 1.000 | 1326 | 0.045 | 0.542 | 508 | 0.549 | 1.000 | 1519 | 0.923 | 0.942 | 1268 | 0.744 | 0.927 | 456 | 0.942 | 0.942 | 885 | 0.750 | 0.927 | 772 | 0.491 | 1.000 | 150 | 0.070 | 0.123 | 194 | 0.473 | 0.765 | 296 | 0.880 |
| BRD-K81330143-001-04-8 | tideglusib | 676 | 0.426 | 0.880 | 1205 | 0.248 | 1.000 | 812 | 0.769 | 0.983 | 2745 | 0.461 | 0.880 | 1432 | 0.959 | 0.983 | 826 | 0.797 | 0.983 | 1011 | 0.637 | 0.983 | 1118 | 0.225 | 0.592 | 47 | 0.007 | 0.030 | 710 | 0.165 | 0.495 | 519 | 0.105 | 1.000 | 391 | 0.133 | 0.467 | 281 | 0.908 | 0.983 | 139 | 0.285 |
| BRD-K81405859-300-01-4 | JTE-607 | 1055 | 0.760 | 0.972 | 2456 | 0.480 | 0.876 | 1432 | 0.294 | 0.706 | 4616 | 0.046 | 0.602 | 1925 | 0.972 | 0.972 | 732 | 0.336 | 0.706 | 1992 | 0.892 | 0.972 | 1525 | 0.098 | 0.602 | 502 | 0.123 | 0.602 | 1264 | 0.876 | 0.972 | 1303 | 0.911 | 0.972 | 659 | 0.203 | 0.706 | 150 | 0.679 | 0.972 | 168 | 0.243 |
| BRD-K81418486-001-44-2 | vorinostat | 723.5 | 0.365 | 0.698 | 2367 | 0.869 | 0.989 | 789.5 | 0.167 | 0.876 | 3171 | 0.216 | 0.907 | 1126 | 0.098 | 0.876 | 1154 | 0.989 | 0.989 | 821.5 | 0.814 | 0.989 | 848 | 0.857 | 0.989 | 529 | 0.605 | 0.907 | 1391 | 0.962 | 0.989 | 886.5 | 0.532 | 0.907 | 861 | 0.972 | 0.989 | 50 | 0.034 | 0.090 | 196 | 0.153 |
| BRD-K81418486-001-47-5 | vorinostat | 4070 | 0.690 | 0.852 | 6005 | 0.350 | 0.667 | 3634 | 0.497 | 0.742 | 13245 | 0.454 | 0.733 | 4780 | 0.314 | 0.667 | 3492 | 0.530 | 0.742 | 7070 | 0.999 | 0.999 | 5750 | 0.934 | 0.981 | 1981 | 0.328 | 0.667 | 2894 | 0.813 | 0.948 | 1809 | 0.041 | 0.581 | 1580 | 0.148 | 0.581 | 565 | 0.166 | 0.581 | 523 | 0.125 |
| BRD-K81458380-001-04-9 | CX-4945 | 419 | 0.109 | 1.000 | 479 | 0.034 | 0.709 | 491 | 0.697 | 0.967 | 842 | 0.307 | 1.000 | 631 | 0.802 | 0.967 | 479 | 0.844 | 0.967 | 365 | 0.197 | 0.590 | 428 | 0.664 | 0.967 | 288 | 0.977 | 0.977 | 337 | 0.493 | 0.871 | 236 | 0.539 | 0.871 | 97 | 0.875 | 0.967 | | | | | |
| BRD-K81473043-001-14-6 | tanespimycin | 5896 | 0.627 | 1.000 | 7477.5 | 0.162 | 0.677 | 3798 | 0.022 | 0.153 | 14144 | 0.001 | 0.022 | 5535 | 0.139 | 0.677 | 5305 | 0.771 | 1.000 | 6904 | 0.614 | 1.000 | 8164 | 0.926 | 1.000 | 2639 | 0.637 | 1.000 | 5823 | 1.000 | 1.000 | 2351 | 0.010 | 0.104 | 4045 | 0.998 | 1.000 | 912 | 0.492 | 1.000 | 1732 | 0.976 |
| BRD-K81473043-001-19-5 | tanespimycin | 1693 | 0.855 | 0.998 | 2134 | 0.189 | 0.661 | 1329 | 0.011 | 0.117 | 3843 | 0.001 | 0.012 | 2295 | 0.749 | 0.998 | 1909 | 0.923 | 0.998 | 2501 | 0.836 | 0.998 | 2025 | 0.970 | 0.998 | 527 | 0.118 | 0.622 | 1850 | 0.998 | 0.998 | 570 | 0.067 | 0.468 | 432 | 0.686 | 0.998 | 149 | 0.165 | 0.661 | 547 | 0.920 |
| BRD-K81548480-001-01-2 | AC-264613 | 967 | 0.267 | 0.702 | 2318 | 0.998 | 0.998 | 1338 | 0.244 | 0.702 | 4321 | 0.478 | 0.669 | 1720 | 0.254 | 0.702 | 684 | 0.058 | 0.702 | 1260 | 0.268 | 0.702 | 2242 | 0.982 | 0.998 | 476 | 0.115 | 0.702 | 1211 | 0.962 | 0.998 | 757 | 0.455 | 0.669 | 223 | 0.042 | 0.081 | 268 | 0.672 | 0.882 | 436 | 0.459 |
| BRD-K81613478-001-01-8 | gastrodin | 657 | 0.348 | 0.664 | 1684.5 | 0.871 | 0.988 | 951.5 | 0.294 | 1.000 | 2420.5 | 0.504 | 0.757 | 978 | 0.116 | 0.810 | 993 | 0.888 | 0.988 | 995 | 0.382 | 0.668 | 1175 | 0.960 | 0.988 | 460.5 | 0.264 | 0.617 | 1244 | 0.985 | 0.988 | 567 | 0.328 | 0.664 | 534 | 0.474 | 0.757 | 484 | 0.988 | 0.988 | 93 | 0.114 |
| BRD-K81645907-001-01-1 | captamine | 403.5 | 0.055 | 0.903 | 504.5 | 0.098 | 0.903 | 907.5 | 0.725 | 0.950 | 1955 | 0.129 | 0.903 | 982 | 0.950 | 0.950 | 720.5 | 0.939 | 0.950 | 686 | 0.731 | 0.950 | 1028 | 0.819 | 0.950 | 232.5 | 0.487 | 0.950 | 303 | 0.205 | 0.537 | 603.5 | 0.487 | 0.950 | 300 | 0.794 | 0.950 | 246.5 | 0.914 | 0.950 | 37 | 0.181 |
| BRD-K81694556-003-01-9 | VTP-27999 | 527 | 0.333 | 0.761 | 1342 | 0.990 | 0.990 | 452 | 0.543 | 0.761 | 2220 | 0.689 | 0.761 | 627 | 0.259 | 0.761 | 338 | 0.168 | 0.761 | 596 | 0.554 | 0.761 | 614 | 0.229 | 0.761 | 71 | 0.459 | 0.761 | 377 | 0.275 | 0.761 | 433 | 0.249 | 1.000 | 443 | 0.509 | 0.761 | 93 | 0.654 | 0.761 | 85 | 0.585 |
| BRD-K81728688-001-04-4 | WZ8040 | 1669 | 0.039 | 0.203 | 5651 | 1.000 | 1.000 | 1660 | 0.257 | 0.601 | 6948.5 | 0.473 | 0.828 | 3400 | 0.598 | 0.893 | 1739 | 0.209 | 0.601 | 3515 | 0.941 | 1.000 | 3341 | 0.997 | 1.000 | 670 | 0.680 | 0.893 | 1653 | 0.408 | 0.778 | 1876.5 | 0.524 | 0.846 | 513 | 0.680 | 0.893 | 239 | 0.319 | 0.669 | 827 | 0.917 |
| BRD-K81801188-001-02-8 | PP-121 | 6143 | 0.881 | 0.925 | 8681 | 0.829 | 0.923 | 4409 | 0.177 | 0.866 | 17151 | 0.368 | 0.923 | 4899 | 0.053 | 0.370 | 4279 | 0.206 | 0.866 | 7069 | 0.835 | 0.923 | 7563 | 0.788 | 0.923 | 2264 | 0.336 | 0.923 | 3778 | 0.691 | 0.923 | 2119 | 0.052 | 0.370 | 2594 | 0.633 | 0.923 | 1059 | 0.716 | 0.923 | 1138 | 0.515 |
| BRD-K81916719-001-13-9 | triclabendazole | 2983 | 0.925 | 0.925 | 4535 | 0.509 | 0.822 | 2561 | 0.314 | 0.822 | 9963 | 0.494 | 0.822 | 3183 | 0.150 | 0.787 | 2641 | 0.388 | 0.822 | 3891 | 0.888 | 0.992 | 4298.5 | 0.664 | 0.992 | 981 | 0.461 | 0.822 | 2163 | 0.861 | 0.992 | 1431 | 0.117 | 0.787 | 834.5 | 0.239 | 0.822 | 440 | 0.348 | 0.822 | 475 | 0.426 |
| BRD-K82091397-001-03-0 | SB-239063 | 1546 | 0.292 | 0.681 | 3115 | 0.949 | 0.990 | 1372 | 0.213 | 1.000 | 4780 | 0.062 | 1.000 | 2306 | 0.570 | 0.798 | 1375 | 0.546 | 0.798 | 2078 | 0.417 | 0.796 | 2050 | 0.990 | 0.990 | 854 | 0.914 | 0.990 | 396 | 0.082 | 0.286 | 1046 | 0.105 | 1.000 | 381 | 0.210 | 0.550 | 434 | 0.658 | 0.864 | 121 | 0.116 |
| BRD-K82135108-001-04-3 | elesclomol | 2158 | 0.969 | 0.969 | 3157 | 0.533 | 0.933 | 2589 | 0.783 | 0.865 | 8810 | 0.588 | 0.950 | 2290.5 | 0.122 | 0.852 | 1552 | 0.241 | 0.852 | 2674 | 0.314 | 0.852 | 3156.5 | 0.406 | 0.852 | 1112 | 0.196 | 0.852 | 1946 | 0.728 | 0.865 | 1783 | 0.526 | 0.933 | 777 | 0.123 | 0.852 | 561 | 0.404 | 0.852 | 279 | 0.925 |
| BRD-K82244583-001-01-3 | MEK162 | 2449 | 0.001 | 0.005 | 2814 | 0.000 | 0.000 | 4167 | 0.684 | 1.000 | 12796.5 | 0.987 | 1.000 | 3122 | 0.788 | 1.000 | 3750 | 0.998 | 1.000 | 3150 | 0.004 | 0.013 | 5354 | 1.000 | 1.000 | 1777.5 | 0.713 | 1.000 | 1622 | 0.909 | 1.000 | 2389 | 0.165 | 0.432 | 1361.5 | 0.976 | 1.000 | 845 | 0.952 | 1.000 | 485 | 0.130 |
| BRD-K82746043-001-15-1 | navitoclax | 1928 | 0.532 | 0.859 | 3505 | 0.940 | 0.974 | 1134 | 0.130 | 0.546 | 5250.5 | 0.109 | 0.546 | 2660 | 0.695 | 0.927 | 1100 | 0.104 | 0.546 | 3098.5 | 0.926 | 0.974 | 2233 | 0.468 | 0.828 | 519 | 0.473 | 0.828 | 1572 | 0.369 | 0.775 | 1115.5 | 0.229 | 0.802 | 368 | 0.045 | 0.119 | 451 | 0.652 | 0.927 | 489 | 0.750 |
| BRD-K82746043-001-19-3 | navitoclax | 1289.5 | 0.938 | 0.954 | 789 | 0.954 | 0.954 | 1050 | 0.257 | 1.000 | 1822.5 | 0.011 | 0.223 | 1032 | 0.513 | 1.000 | 851.5 | 0.430 | 1.000 | 1005 | 0.786 | 1.000 | 760 | 0.586 | 1.000 | 131 | 0.827 | 1.000 | 758.5 | 0.582 | 1.000 | 446 | 0.342 | 1.000 | 214 | 0.329 | 1.000 | | | | | |
| BRD-K82818427-001-02-2 | batimastat | 5168 | 0.830 | 1.000 | 6267 | 0.425 | 0.767 | 3432 | 0.288 | 0.756 | 14014 | 0.438 | 0.767 | 4657 | 0.284 | 0.756 | 3430 | 0.287 | 0.756 | 6722.5 | 0.988 | 1.000 | 6248 | 0.932 | 1.000 | 1577 | 0.056 | 0.390 | 2966.5 | 1.000 | 1.000 | 1927 | 0.011 | 0.116 | 1180 | 0.230 | 0.756 | 903 | 0.651 | 0.911 | 1297 | 0.860 |
| BRD-K82823804-001-04-1 | PD-407824 | 1832 | 0.647 | 0.900 | 3466 | 0.925 | 0.995 | 1223 | 0.033 | 0.645 | 6212 | 0.690 | 0.900 | 1668 | 0.267 | 0.645 | 1121 | 0.072 | 0.645 | 1610 | 0.333 | 0.645 | 1896 | 0.729 | 0.900 | 596 | 0.183 | 0.645 | 889 | 0.283 | 0.645 | 1269 | 0.098 | 0.645 | 605 | 0.722 | 0.900 | 253 | 0.479 | 0.774 | 386 | 0.205 |
| BRD-K82850175-001-01-1 | navarixin | 1124 | 0.557 | 0.756 | 2576 | 0.534 | 1.000 | 1468 | 0.291 | 1.000 | 4421.5 | 0.576 | 0.756 | 1330 | 0.551 | 1.000 | 1697 | 0.962 | 0.976 | 1791 | 0.231 | 0.970 | 1732 | 0.544 | 1.000 | 306 | 0.137 | 0.961 | 1086 | 0.964 | 0.976 | 825 | 0.041 | 0.428 | 348 | 0.220 | 0.420 | 351 | 0.909 | 0.976 | 50 | 0.168 |
| BRD-K82861010-001-02-5 | daclatasvir | 281 | 0.154 | 1.000 | 1204 | 0.875 | 0.919 | 524 | 0.266 | 1.000 | 2743 | 0.753 | 0.890 | 866 | 0.806 | 0.890 | 521 | 0.126 | 0.442 | 585 | 0.252 | 0.589 | 795 | 0.934 | 0.934 | 293 | 0.183 | 0.481 | 387 | 0.803 | 0.890 | 176 | 0.618 | 0.890 | 264 | 0.650 | 0.890 | | | | 31 | 0.150 |
| BRD-K82928847-001-04-7 | ACY-1215 | 878 | 0.422 | 0.943 | 2209 | 0.686 | 0.960 | 1138 | 0.319 | 0.837 | 5424 | 0.894 | 0.986 | 2214 | 0.560 | 0.960 | 1170 | 0.763 | 0.967 | 2468 | 0.986 | 0.986 | 1855 | 0.629 | 0.960 | 760 | 0.658 | 0.960 | 430 | 0.979 | 0.986 | 994 | 0.062 | 0.436 | 169 | 0.062 | 0.260 | 341 | 0.468 | 0.943 | 285 | 0.289 |
| BRD-K82967180-001-01-6 | LY3023414 | 8616 | 1.000 | 1.000 | 9384 | 0.906 | 1.000 | 4104 | 0.058 | 0.243 | 19560 | 0.855 | 1.000 | 4248 | 0.007 | 0.071 | 4441 | 0.351 | 0.686 | 8943 | 0.992 | 1.000 | 6268 | 0.120 | 0.420 | 1720 | 0.201 | 0.491 | 3667 | 0.392 | 0.686 | 3834 | 0.513 | 0.770 | 2679 | 0.462 | 0.746 | 656 | 0.166 | 0.491 | 1580 | 0.921 |
| BRD-K83029223-001-01-3 | litronesib | 1388 | 0.561 | 0.906 | 3445 | 0.906 | 0.992 | 1733 | 0.202 | 0.607 | 6930 | 0.275 | 0.641 | 1985 | 0.131 | 0.607 | 2187 | 0.944 | 0.992 | 1613 | 0.695 | 0.973 | 2445 | 0.856 | 0.992 | 1010 | 0.366 | 0.715 | 1468 | 0.992 | 0.992 | 1138 | 0.200 | 0.607 | 1042 | 0.420 | 0.735 | 322 | 0.652 | 0.973 | 417 | 0.201 |
| BRD-K83699324-001-01-1 | NMS-1286937 | 2754 | 0.226 | 0.678 | 6524 | 0.976 | 1.000 | 2321 | 0.396 | 0.756 | 11037 | 0.519 | 0.856 | 3287 | 0.181 | 0.635 | 3370 | 0.617 | 0.910 | 3979 | 0.818 | 0.964 | 4600 | 0.693 | 0.910 | 934 | 0.138 | 0.635 | 3824 | 1.000 | 1.000 | 1808 | 0.112 | 0.635 | 1555 | 0.274 | 0.720 | 205 | 0.166 | 0.635 | 634 | 0.359 |
| BRD-K83834509-001-01-1 | PTC-209 | 731 | 0.958 | 0.958 | 718 | 0.077 | 0.807 | 885 | 0.702 | 0.921 | 1877 | 0.513 | 0.828 | 634 | 0.055 | 0.807 | 495 | 0.871 | 0.958 | 1013 | 0.674 | 0.921 | 950 | 0.689 | 0.921 | 20 | 0.017 | 0.045 | 698 | 0.926 | 0.958 | 293 | 0.187 | 0.898 | 524 | 0.922 | 0.958 | 79 | 0.509 | 0.828 | 39 | 0.195 |
| BRD-K83837640-001-04-8 | JNJ-26481585 | 3046 | 0.521 | 0.730 | 4164 | 0.037 | 0.223 | 2111 | 0.133 | 0.350 | 11458 | 0.920 | 0.999 | 2851 | 0.043 | 0.223 | 3705 | 0.886 | 0.999 | 4397 | 0.991 | 0.999 | 5382 | 0.987 | 0.999 | 1153 | 0.412 | 0.665 | 3764 | 0.999 | 0.999 | 2180 | 0.297 | 0.665 | 1778 | 0.573 | 0.752 | 97 | 0.039 | 0.223 | 479 | 0.401 |
| BRD-K83963101-001-06-9 | MLN-8054 | 1986 | 0.525 | 0.803 | 4221 | 0.959 | 0.959 | 1859 | 0.535 | 0.803 | 6717 | 0.106 | 0.803 | 2970 | 0.725 | 0.881 | 2592 | 0.749 | 0.881 | 2838 | 0.918 | 0.959 | 2375 | 0.369 | 0.803 | 691 | 0.203 | 0.803 | 1753 | 0.755 | 0.881 | 1539 | 0.468 | 0.803 | 914 | 0.166 | 0.803 | 233 | 0.318 | 0.803 | 170 | 0.034 |
| BRD-K83988098-001-02-0 | alvespimycin | 1542 | 0.904 | 0.997 | 1843 | 0.511 | 0.997 | 939 | 0.002 | 0.044 | 3343 | 0.008 | 0.088 | 988 | 0.204 | 0.718 | 1115 | 0.927 | 0.997 | 1797 | 0.727 | 0.997 | 2187 | 0.985 | 0.997 | 349 | 0.027 | 0.190 | 1073 | 0.997 | 0.997 | 1216 | 0.914 | 0.997 | 494 | 0.934 | 0.997 | 353 | 0.590 | 0.997 | 75 | 0.292 |
| BRD-K83988098-003-03-4 | alvespimycin | 5137 | 0.966 | 1.000 | 5213 | 0.405 | 0.773 | 2040 | 0.003 | 0.020 | 10050 | 0.022 | 0.113 | 3976 | 0.097 | 0.347 | 3486 | 0.492 | 0.815 | 4449 | 0.156 | 0.468 | 5879 | 0.933 | 1.000 | 1648 | 0.249 | 0.654 | 4773 | 1.000 | 1.000 | 2037 | 0.289 | 0.674 | 2904 | 0.957 | 1.000 | 321 | 0.099 | 0.347 | 1499 | 0.987 |
| BRD-K84163249-001-01-9 | ercalcitriol | 893.5 | 0.269 | 0.712 | 1540 | 0.876 | 0.936 | 1303.5 | 0.693 | 0.936 | 3380.5 | 0.240 | 1.000 | 1033.5 | 0.056 | 0.390 | 1253.5 | 0.774 | 0.936 | 1465 | 0.640 | 0.936 | 1529.5 | 0.866 | 0.936 | 690.5 | 0.690 | 0.936 | 967.5 | 0.815 | 0.936 | 774.5 | 0.019 | 0.279 | 288.5 | 0.821 | 0.936 | 249 | 0.271 | 0.712 | 197 | 0.442 |
| BRD-K84564571-001-02-4 | telatinib | 150 | 0.867 | 1.000 | 304.5 | 0.035 | 0.729 | 385 | 0.404 | 0.942 | 865 | 0.538 | 0.964 | 333.5 | 0.374 | 0.942 | 367.5 | 0.962 | 1.000 | 532.5 | 0.955 | 1.000 | 498 | 0.180 | 0.801 | 246.5 | 0.175 | 0.801 | 111.5 | 0.551 | 0.964 | 258.5 | 0.815 | 1.000 | 77.5 | 0.229 | 0.801 | | | | 107.5 | 0.510 |
| BRD-K84683831-001-01-0 | radezolid | 1456 | 0.990 | 0.990 | 1899 | 0.718 | 0.990 | 1380 | 0.478 | 1.000 | 4002 | 0.669 | 0.990 | 1532 | 0.113 | 0.790 | 679 | 0.206 | 1.000 | 1827 | 0.932 | 0.990 | 1969 | 0.982 | 0.990 | 411 | 0.079 | 0.185 | 376 | 0.146 | 0.306 | 352 | 0.109 | 0.790 | 304 | 0.860 | 0.990 | 186 | 0.905 | 0.990 | 215 | 0.521 |
| BRD-K84868168-001-01-0 | erdafitinib | 93 | 0.551 | 0.921 | 1613 | 0.961 | 0.976 | 1025 | 0.776 | 0.959 | 2858.5 | 0.771 | 0.959 | 1349 | 0.378 | 1.000 | 487 | 0.712 | 0.959 | 669 | 0.909 | 0.976 | 640 | 0.054 | 0.152 | 345 | 0.093 | 0.195 | 721 | 0.065 | 0.152 | 352.5 | 0.260 | 1.000 | 63 | 0.062 | 0.152 | | | | 18 | 0.088 |
| BRD-K84937637-001-09-9 | sirolimus | 4611 | 0.999 | 0.999 | 5572 | 0.720 | 0.999 | 2287 | 0.483 | 0.845 | 9991 | 0.420 | 0.801 | 2684 | 0.015 | 0.159 | 2793 | 0.813 | 0.999 | 4597 | 0.991 | 0.999 | 2608 | 0.040 | 0.208 | 1045 | 0.556 | 0.897 | 2319 | 0.936 | 0.999 | 1081 | 0.029 | 0.206 | 1099 | 0.387 | 0.801 | 256 | 0.073 | 0.306 | 1076 | 0.849 |
| BRD-K85014146-001-02-5 | AZD2461 | 202.5 | 0.671 | 1.000 | 606 | 0.207 | 1.000 | 398.5 | 0.926 | 1.000 | 1515 | 0.822 | 1.000 | 375.5 | 0.647 | 1.000 | 133.5 | 0.232 | 1.000 | 454.5 | 0.737 | 1.000 | 109 | 0.008 | 0.021 | 170.5 | 0.175 | 0.368 | 412 | 0.795 | 1.000 | 217.5 | 0.061 | 1.000 | 46 | 0.013 | 0.031 | | | | 85 | 0.243 |
| BRD-K85140930-001-12-7 | aphidicolin | 4127 | 0.369 | 0.920 | 5868 | 0.866 | 0.972 | 2870 | 0.049 | 0.403 | 9740 | 0.076 | 0.403 | 4280 | 0.485 | 0.940 | 3428 | 0.972 | 0.972 | 5373 | 0.959 | 0.972 | 3836 | 0.184 | 0.773 | 1019 | 0.394 | 0.920 | 2521 | 0.709 | 0.972 | 2345 | 0.361 | 0.920 | 858 | 0.025 | 0.403 | 579 | 0.566 | 0.972 | 286 | 0.077 |
| BRD-K85146014-314-05-4 | foscarnet | 652.5 | 0.971 | 0.971 | 979.5 | 0.376 | 0.865 | 332.5 | 0.502 | 0.865 | 1137.5 | 0.295 | 0.865 | 467 | 0.164 | 0.863 | 300.5 | 0.160 | 0.863 | 627.5 | 0.838 | 0.978 | 880.5 | 0.925 | 1.000 | 51 | 0.066 | 0.863 | 220 | 0.273 | 0.865 | 414 | 0.334 | 0.865 | 336 | 0.808 | 0.978 | 170 | 0.731 | 0.978 | 216.5 | 0.584 |
| BRD-K85307935-236-09-5 | novobiocin | 116 | 0.016 | 0.328 | 1111 | 0.099 | 0.521 | 931 | 0.749 | 0.962 | 2188 | 0.950 | 0.984 | 1296.5 | 0.870 | 0.984 | 484 | 0.316 | 0.903 | 587 | 0.115 | 0.402 | 958.5 | 0.917 | 0.984 | 411.5 | 0.984 | 0.984 | 192 | 0.689 | 0.962 | 496 | 0.352 | 0.740 | 305 | 0.455 | 0.807 | | | | 133 | 0.344 |
| BRD-K85402309-001-06-6 | dovitinib | 1248 | 0.219 | 0.602 | 2865 | 0.757 | 0.935 | 1300 | 0.161 | 0.602 | 5225 | 0.459 | 0.839 | 1717 | 0.096 | 0.602 | 1082 | 0.480 | 0.839 | 2755 | 0.985 | 0.985 | 1839 | 0.590 | 0.870 | 576 | 0.180 | 0.602 | 1364 | 0.753 | 0.935 | 756 | 0.022 | 0.469 | 335 | 0.134 | 0.602 | 86 | 0.296 | 0.691 | 280 | 0.229 |
| BRD-K85402309-043-01-9 | dovitinib | 121.5 | 0.299 | 1.000 | 522.5 | 0.430 | 1.000 | 236 | 0.758 | 0.937 | 934.5 | 0.758 | 0.937 | 462.5 | 0.343 | 1.000 | 312 | 0.861 | 0.950 | 277.5 | 0.995 | 0.995 | 208 | 0.129 | 0.270 | 155 | 0.088 | 0.206 | 306 | 0.365 | 0.639 | 110 | 0.600 | 0.900 | 9 | 0.014 | 0.038 | | | | 214 | 0.905 |
| BRD-K85554912-001-06-3 | ivermectin | 1240.5 | 0.854 | 0.898 | 1289 | 0.800 | 0.898 | 934.5 | 0.425 | 1.000 | 3367.5 | 0.653 | 0.898 | 897 | 0.343 | 1.000 | 1278 | 0.963 | 0.963 | 988 | 0.733 | 0.898 | 789 | 0.015 | 0.036 | 350 | 0.442 | 0.794 | 200 | 0.004 | 0.011 | 353 | 0.454 | 0.794 | 236 | 0.336 | 0.705 | 260 | 0.839 | 0.898 | 265 | 0.855 |
| BRD-K85606544-001-09-1 | neratinib | 323 | 0.336 | 0.706 | 730.5 | 0.996 | 1.000 | 405.5 | 0.124 | 0.871 | 1209.5 | 0.844 | 0.997 | 458.5 | 0.135 | 0.473 | 138.5 | 0.070 | 0.294 | 669.5 | 0.794 | 0.997 | 972 | 1.000 | 1.000 | 130 | 0.551 | 0.842 | 265 | 0.619 | 0.866 | 309 | 0.280 | 0.652 | 108.5 | 0.902 | 0.997 | | | | | |
| BRD-K85751432-001-03-3 | refametinib | 1568 | 0.003 | 0.028 | 2714 | 0.000 | 0.002 | 2100 | 0.264 | 0.562 | 9970 | 0.913 | 0.999 | 3618 | 0.963 | 0.999 | 2546 | 0.859 | 0.999 | 2597 | 0.077 | 0.268 | 4777 | 0.999 | 0.999 | 427 | 0.037 | 0.154 | 2215 | 0.855 | 0.999 | 1617 | 0.207 | 0.544 | 1538 | 0.846 | 0.999 | 153 | 0.114 | 0.342 | 757 | 0.725 |
| BRD-K85849709-001-01-5 | AZD2858 | 1433 | 0.104 | 0.439 | 1723 | 0.586 | 0.904 | 974 | 0.069 | 0.361 | 5563 | 0.620 | 0.904 | 2737.5 | 0.623 | 0.904 | 1233 | 0.541 | 0.904 | 2491 | 0.999 | 0.999 | 2352.5 | 0.410 | 0.862 | 680 | 0.899 | 0.999 | 1039.5 | 0.402 | 0.862 | 1015 | 0.018 | 0.361 | 927 | 0.406 | 0.862 | 421.5 | 0.953 | 0.999 | 241 | 0.146 |
| BRD-K85872723-001-15-9 | kynurenic-acid | 412 | 0.248 | 0.836 | 463.5 | 0.582 | 0.836 | 139 | 0.150 | 0.836 | 1265.5 | 0.627 | 0.836 | 338.5 | 0.453 | 0.836 | 74.5 | 0.069 | 0.728 | 407 | 0.057 | 0.728 | 571.5 | 0.604 | 0.836 | 272 | 0.353 | 0.836 | 205.5 | 0.251 | 0.836 | 396 | 0.846 | 0.846 | 68.5 | 0.665 | 0.836 | 202.5 | 0.763 | 0.836 | 68.5 | 0.665 |
| BRD-K85920262-001-02-1 | WAY-600 | 3749 | 0.658 | 0.987 | 7057 | 0.934 | 0.998 | 3060 | 0.252 | 0.662 | 14307 | 0.905 | 0.998 | 3467 | 0.002 | 0.018 | 4324 | 0.998 | 0.998 | 5612 | 0.881 | 0.998 | 5422 | 0.878 | 0.998 | 1726 | 0.498 | 0.918 | 1768 | 0.003 | 0.018 | 1099 | 0.000 | 0.007 | 1200 | 0.146 | 0.437 | 612 | 0.553 | 0.918 | 951 | 0.472 |
| BRD-K86003836-001-10-5 | flubendazole | 4314 | 0.149 | 0.414 | 7122 | 0.870 | 1.000 | 3530 | 0.115 | 0.414 | 14793 | 0.404 | 0.743 | 4885 | 0.111 | 0.414 | 3917 | 0.730 | 1.000 | 6701 | 0.996 | 1.000 | 5550 | 0.664 | 0.997 | 2808 | 0.923 | 1.000 | 4339 | 1.000 | 1.000 | 2133 | 0.120 | 0.414 | 1877 | 0.178 | 0.414 | 1084 | 0.848 | 1.000 | 353 | 0.023 |
| BRD-K86525559-001-07-8 | AZD7762 | 5921 | 0.545 | 0.875 | 8430 | 0.610 | 0.875 | 4661 | 0.399 | 0.837 | 17549 | 0.573 | 0.875 | 5000 | 0.062 | 0.326 | 5908 | 0.985 | 0.995 | 7448 | 0.708 | 0.875 | 7422 | 0.786 | 0.917 | 2407 | 0.269 | 0.755 | 4032 | 0.690 | 0.875 | 2278 | 0.003 | 0.060 | 2098 | 0.097 | 0.409 | 1124 | 0.973 | 0.995 | 546 | 0.022 |
| BRD-K86687746-001-01-3 | R306465 | 3953 | 0.237 | 0.621 | 4874 | 0.100 | 0.492 | 4358 | 0.765 | 0.892 | 11990 | 0.508 | 0.769 | 4255 | 0.325 | 0.758 | 2829 | 0.715 | 0.884 | 7247 | 1.000 | 1.000 | 5698 | 0.892 | 0.986 | 1623 | 0.642 | 0.855 | 3455 | 0.982 | 1.000 | 1946 | 0.116 | 0.492 | 1702 | 0.513 | 0.769 | 345 | 0.407 | 0.769 | 66 | 0.001 |
| BRD-K86727142-001-12-4 | embelin | 348 | 0.966 | 0.991 | 1142 | 0.991 | 0.991 | 654 | 0.801 | 1.000 | 1745 | 0.209 | 0.668 | 526.5 | 0.223 | 0.668 | 399 | 0.005 | 0.105 | 878.5 | 0.834 | 1.000 | 655 | 0.136 | 0.572 | 173 | 0.089 | 0.572 | 487 | 0.491 | 0.859 | 211 | 0.475 | 0.859 | 117 | 0.322 | 0.846 | 102 | 0.757 | 1.000 | 24 | 0.126 |
| BRD-K86797399-001-05-1 | SB-939 | 5538 | 0.453 | 0.717 | 8168 | 0.499 | 0.717 | 4397 | 0.512 | 0.717 | 18367 | 0.734 | 0.964 | 4733 | 0.052 | 0.218 | 5405 | 0.916 | 1.000 | 9041 | 0.997 | 1.000 | 8500 | 0.990 | 1.000 | 1662 | 0.033 | 0.174 | 4985 | 1.000 | 1.000 | 2102 | 0.008 | 0.112 | 2541 | 0.383 | 0.670 | 573 | 0.107 | 0.324 | 762 | 0.108 |
| BRD-K86816506-001-01-8 | PF-04457845 | 534 | 0.184 | 0.966 | 1991 | 0.978 | 0.978 | 845 | 0.161 | 0.966 | 3443 | 0.501 | 1.000 | 1430 | 0.873 | 0.917 | 561 | 0.043 | 0.896 | 1636 | 0.794 | 0.898 | 1118 | 0.687 | 0.898 | 434 | 0.153 | 0.321 | 414 | 0.295 | 0.517 | 675 | 0.764 | 0.898 | 324 | 0.106 | 0.247 | 255 | 0.354 | 0.571 | 7 | 0.056 |
| BRD-K86887724-001-10-6 | dofetilide | 656 | 0.759 | 0.885 | 897 | 0.043 | 0.448 | 579 | 0.748 | 0.885 | 1998 | 0.445 | 0.885 | 952 | 0.904 | 0.922 | 484 | 0.431 | 0.885 | 968 | 0.922 | 0.922 | 934 | 0.598 | 0.885 | 399 | 0.640 | 0.885 | 71 | 0.104 | 0.570 | 139 | 0.136 | 0.570 | 73 | 0.110 | 0.570 | 81 | 0.559 | 0.885 | 158 | 0.561 |
| BRD-K86930074-001-05-0 | cediranib | 4941 | 0.996 | 0.996 | 6029 | 0.995 | 0.996 | 3366 | 0.520 | 0.909 | 9257.5 | 0.003 | 0.053 | 4528 | 0.653 | 0.996 | 3439.5 | 0.971 | 0.996 | 5073 | 0.963 | 0.996 | 4429 | 0.699 | 0.996 | 1482 | 0.322 | 0.616 | 1909 | 0.031 | 0.323 | 1962 | 0.169 | 0.443 | 1191 | 0.193 | 0.449 | 464 | 0.103 | 0.378 | 592 | 0.254 |
| BRD-K87112191-001-04-5 | PP242 | 6215 | 0.991 | 0.991 | 6684 | 0.348 | 0.804 | 4446 | 0.732 | 0.885 | 12644 | 0.038 | 0.725 | 4544 | 0.083 | 0.725 | 3836 | 0.715 | 0.885 | 5204 | 0.496 | 0.804 | 5649 | 0.982 | 0.991 | 1685 | 0.188 | 0.725 | 3154 | 0.482 | 0.804 | 2434 | 0.207 | 0.725 | 1825 | 0.306 | 0.804 | 990 | 0.759 | 0.885 | 368 | 0.122 |
| BRD-K87124298-001-03-4 | OSI-027 | 6424 | 0.998 | 1.000 | 5945 | 0.024 | 0.167 | 3600 | 0.246 | 0.645 | 16685 | 0.614 | 0.904 | 5280 | 0.108 | 0.379 | 4514 | 0.465 | 0.904 | 6795 | 0.850 | 0.974 | 8190 | 1.000 | 1.000 | 2037 | 0.230 | 0.645 | 3521 | 0.462 | 0.904 | 1977 | 0.007 | 0.142 | 1403 | 0.032 | 0.168 | 963 | 0.646 | 0.904 | 696 | 0.578 |
| BRD-K87349682-347-03-8 | norepinephrine | 1578 | 0.529 | 0.968 | 2411 | 0.346 | 1.000 | 1591 | 0.241 | 1.000 | 4370 | 0.009 | 0.187 | 2340 | 0.649 | 0.968 | 1827 | 0.982 | 0.982 | 1527 | 0.450 | 1.000 | 3110 | 0.982 | 0.982 | 597 | 0.445 | 0.968 | 963 | 0.704 | 0.968 | 758 | 0.138 | 0.969 | 747 | 0.957 | 0.982 | 399 | 0.577 | 0.968 | 525 | 0.575 |
| BRD-K87510569-001-02-8 | RS-504393 | 680 | 0.901 | 0.986 | 893 | 0.864 | 0.986 | 588 | 0.464 | 0.649 | 2354 | 0.555 | 0.705 | 634 | 0.394 | 0.644 | 468 | 0.306 | 0.644 | 959 | 0.985 | 0.986 | 654 | 0.146 | 0.644 | 82 | 0.126 | 0.644 | 572 | 0.414 | 0.644 | 364 | 0.429 | 0.644 | 171 | 0.219 | 0.644 | 30 | 0.146 | 0.644 | 17 | 0.091 |
| BRD-K87512222-019-01-7 | taltobulin | 4729 | 0.629 | 0.887 | 5556 | 0.399 | 0.645 | 3884 | 0.383 | 0.645 | 10688 | 0.067 | 0.367 | 4011 | 0.161 | 0.483 | 2950 | 0.633 | 0.887 | 7338 | 1.000 | 1.000 | 5672 | 0.860 | 1.000 | 1112 | 0.097 | 0.409 | 3420 | 0.973 | 1.000 | 2953 | 0.787 | 0.972 | 1343 | 0.136 | 0.475 | 287 | 0.070 | 0.367 | 257 | 0.050 |
| BRD-K87696786-003-04-4 | LY456236 | 685.5 | 0.550 | 0.888 | 2603 | 0.472 | 1.000 | 910.5 | 0.042 | 0.876 | 4013 | 0.276 | 1.000 | 1595.5 | 0.498 | 1.000 | 1417 | 0.853 | 0.941 | 1652 | 0.741 | 0.928 | 1662 | 0.754 | 0.928 | 591 | 0.592 | 0.888 | 951 | 0.963 | 0.963 | 1030 | 0.385 | 1.000 | 380.5 | 0.304 | 0.639 | 304 | 0.386 | 0.736 | 252 | 0.227 |
| BRD-K87700323-003-05-1 | cetylpyridinium | 1745 | 0.393 | 0.635 | 6011 | 1.000 | 1.000 | 1489 | 0.030 | 0.229 | 8501 | 0.521 | 0.782 | 3083 | 0.241 | 0.723 | 2700 | 0.958 | 1.000 | 3880 | 0.801 | 1.000 | 3647 | 0.908 | 1.000 | 1152 | 0.090 | 0.189 | 1494 | 0.068 | 0.160 | 1359 | 0.028 | 0.229 | 842 | 0.877 | 1.000 | 393 | 0.321 | 0.562 | 64 | 0.205 |
| BRD-K87737963-001-06-0 | cyt387 | 475 | 0.525 | 1.000 | 749 | 0.906 | 0.970 | 405 | 0.056 | 0.817 | 1618 | 0.213 | 1.000 | 478 | 0.078 | 0.817 | 531 | 0.902 | 0.970 | 303 | 0.643 | 1.000 | 758 | 0.797 | 0.970 | 221 | 0.030 | 0.063 | 266 | 0.027 | 0.063 | 259 | 0.774 | 0.970 | 260 | 0.433 | 0.700 | 229 | 0.943 | 0.970 | 31 | 0.172 |
| BRD-K87782578-001-01-4 | AVL-292 | 379 | 0.581 | 1.000 | 677 | 0.947 | 0.976 | 884 | 0.819 | 0.976 | 1548 | 0.067 | 0.707 | 507 | 0.014 | 0.296 | 564 | 0.728 | 0.976 | 226 | 0.540 | 1.000 | 954 | 0.976 | 0.976 | 159 | 0.063 | 0.146 | 590 | 0.376 | 0.719 | 418 | 0.934 | 0.976 | 358 | 0.790 | 0.976 | 128 | 0.898 | 0.976 | 101 | 0.739 |
| BRD-K87827419-001-16-8 | melphalan | 929 | 0.692 | 0.994 | 2002 | 0.832 | 0.994 | 995 | 0.815 | 0.994 | 2770 | 0.214 | 0.748 | 1262 | 0.117 | 0.490 | 409 | 0.042 | 0.221 | 1317 | 0.861 | 0.994 | 1792 | 0.978 | 0.994 | 446 | 0.025 | 0.065 | 523 | 0.635 | 0.994 | 162 | 0.001 | 0.030 | 462 | 0.086 | 0.180 | 218 | 0.615 | 0.994 | 187 | 0.039 |
| BRD-K87909389-003-03-4 | alvocidib | 3447 | 0.111 | 0.716 | 6869 | 0.667 | 0.925 | 3489 | 0.127 | 0.716 | 13093 | 0.184 | 0.773 | 5253 | 0.429 | 0.925 | 4005 | 0.829 | 0.925 | 4895 | 0.672 | 0.925 | 6266 | 0.881 | 0.925 | 1270 | 0.063 | 0.716 | 3653 | 0.855 | 0.925 | 2524 | 0.441 | 0.925 | 2336 | 0.933 | 0.933 | 899 | 0.636 | 0.925 | 1094 | 0.615 |
| BRD-K88429204-001-35-1 | pyrimethamine | 1396 | 0.202 | 0.607 | 2202 | 0.032 | 0.340 | 1305 | 0.380 | 0.798 | 6408 | 0.655 | 0.975 | 2206 | 0.513 | 0.874 | 1967 | 0.991 | 0.991 | 2126 | 0.832 | 0.975 | 2453 | 0.541 | 0.874 | 535 | 0.036 | 0.187 | 1266 | 0.882 | 0.975 | 768 | 0.018 | 0.340 | 375 | 0.175 | 0.607 | 606 | 0.953 | 0.991 | 764 | 0.766 |
| BRD-K88506063-001-01-8 | KD025 | 2523 | 1.000 | 1.000 | 2974 | 0.993 | 1.000 | 1732 | 0.485 | 0.862 | 6165 | 0.613 | 0.919 | 1902 | 0.003 | 0.069 | 916 | 0.156 | 0.547 | 1970 | 0.493 | 0.862 | 2356 | 0.548 | 0.885 | 681 | 0.140 | 0.547 | 1472 | 0.984 | 1.000 | 659 | 0.044 | 0.233 | 282 | 0.016 | 0.165 | 478 | 0.750 | 1.000 | 511 | 0.823 |
| BRD-K88510285-001-11-1 | bortezomib | 5874 | 0.525 | 0.989 | 6680 | 0.028 | 0.162 | 6426 | 0.989 | 0.989 | 17451 | 0.509 | 0.989 | 6572 | 0.786 | 0.989 | 5785 | 0.977 | 0.989 | 7649 | 0.871 | 0.989 | 5657 | 0.031 | 0.162 | 2489 | 0.336 | 0.989 | 2268 | 0.007 | 0.076 | 2085 | 0.007 | 0.076 | 2884 | 0.672 | 0.989 | 1085 | 0.740 | 0.989 | 751 | 0.100 |
| BRD-K88510285-001-17-8 | bortezomib | 4729 | 0.893 | 0.954 | 5317 | 0.006 | 0.132 | 4013 | 0.220 | 0.788 | 14411 | 0.785 | 0.954 | 4610 | 0.622 | 0.933 | 3886 | 0.877 | 0.954 | 6469 | 0.941 | 0.954 | 5068 | 0.106 | 0.740 | 1314 | 0.083 | 0.740 | 2817 | 0.241 | 0.788 | 3217 | 0.563 | 0.909 | 2658 | 0.888 | 0.954 | 527 | 0.353 | 0.788 | 732 | 0.375 |
| BRD-K88625236-001-04-2 | nonoxynol-9 | 438 | 0.681 | 0.961 | 1210 | 0.136 | 0.678 | 790 | 0.369 | 0.892 | 3265 | 0.653 | 0.961 | 950 | 0.106 | 0.678 | 796 | 0.382 | 0.892 | 1603 | 0.764 | 0.961 | 1830 | 0.961 | 0.961 | 567 | 0.253 | 0.759 | 563 | 0.472 | 0.907 | 644 | 0.238 | 0.759 | 522 | 0.889 | 0.961 | 94 | 0.475 | 0.907 | 60 | 0.045 |

| broad_id | name | al_nervous_system_PVAL_ADJ | fibroblast_MWU_STAT | fibroblast_PVAL | fibroblast_PVAL_ADJ | bile_duct_MWU_STAT | bile_duct_PVAL | bile_duct_PVAL_ADJ | thyroid_MWU_STAT | thyroid_PVAL | thyroid_PVAL_ADJ | upper_aerodigestive_MWU_STAT | upper_aerodigestive_PVAL | upper_aerodigestive_PVAL_ADJ | uterus_MWU_STAT | uterus_PVAL | uterus_PVAL_ADJ | liver_MWU_STAT | liver_PVAL | liver_PVAL_ADJ | esophagus_MWU_STAT | esophagus_PVAL | esophagus_PVAL_ADJ |
|---|---|---|---|---|---|---|---|---|---|---|---|---|---|---|---|---|---|---|---|---|---|---|---|
| BRD-K78431006-001-15-1 | crizotinib | 0.122 | | | | | | | 379 | 0.936 | 0.940 | 367 | 0.095 | 0.998 | 307 | 0.198 | 1.000 | 368 | 0.917 | 0.940 | 710 | 0.416 | 1.000 |
| BRD-K78567475-001-01-4 | dolastatin-10 | 0.511 | 122 | 0.291 | 0.611 | 1502 | 0.954 | 0.999 | 1480 | 0.146 | 0.511 | 3447 | 0.138 | 0.511 | 3802 | 0.622 | 0.871 | 3629 | 0.782 | 0.999 | 3519 | 0.270 | 0.611 |
| BRD-K78659596-001-03-9 | ixazomib | 0.504 | 131 | 0.341 | 0.651 | 1191 | 0.824 | 0.952 | 1153 | 0.278 | 0.651 | 3586 | 0.432 | 0.756 | 3006 | 0.832 | 0.952 | 2454 | 0.855 | 0.952 | 4338 | 0.737 | 0.952 |
| BRD-K78815826-001-14-6 | suxibuzone | 1.000 | | | | 370.5 | 0.990 | 0.990 | 264 | 0.443 | 0.931 | 610 | 0.747 | 1.000 | 152 | 0.017 | 0.359 | 240 | 0.676 | 1.000 | 688.5 | 0.914 | 0.990 |
| BRD-K78960041-001-05-7 | oxaliplatin | 0.998 | | | | 194 | 0.815 | 0.998 | 947 | 0.860 | 0.998 | 2469 | 0.998 | 0.998 | 1002 | 0.157 | 0.532 | 1036 | 0.578 | 0.909 | 1469 | 0.328 | 0.688 |
| BRD-K79131256-001-17-9 | albendazole | 0.534 | 112 | 0.391 | 0.708 | 398 | 0.885 | 0.974 | 747 | 0.694 | 0.972 | 1563 | 0.122 | 0.534 | 1727 | 0.420 | 0.708 | 690 | 0.132 | 0.534 | 1632 | 0.176 | 0.534 |
| BRD-K79239947-001-06-8 | PF-573228 | 0.713 | | | | 246 | 0.940 | 0.940 | 247.5 | 0.271 | 1.000 | 551.5 | 0.394 | 0.713 | 216.5 | 0.166 | 1.000 | 246 | 0.940 | 0.940 | 614.5 | 0.383 | 0.713 |
| BRD-K79254416-001-21-8 | decitabine | 0.949 | | | | 350 | 0.928 | 0.974 | 210 | 0.792 | 0.949 | 535 | 0.500 | 0.811 | 741 | 0.679 | 0.949 | 529 | 0.730 | 0.949 | 381 | 0.037 | 0.770 |
| BRD-K79254416-001-22-6 | decitabine | 0.794 | 285.5 | 0.958 | 0.983 | 607 | 0.599 | 0.866 | 661 | 0.719 | 0.866 | 1648 | 0.193 | 0.794 | 1833 | 0.255 | 0.794 | 1499 | 0.672 | 0.866 | 2319 | 0.681 | 0.866 |
| BRD-K79259477-001-04-9 | NVP-BHG712 | 0.676 | | | | 149 | 0.148 | 0.622 | 320 | 0.874 | 0.983 | 669 | 0.319 | 0.609 | 209 | 0.013 | 0.135 | 158 | 0.176 | 0.411 | 926.5 | 0.803 | 0.983 |
| BRD-K79584249-001-01-3 | CGM097 | 0.570 | | | | | | | | | | 79 | 0.316 | 0.578 | 381 | 0.918 | 0.994 | 115 | 0.657 | 0.812 | 515 | 0.994 | 0.994 |
| BRD-K79612754-001-19-1 | inosine | 0.320 | 13 | 0.059 | 0.320 | 510 | 0.714 | 0.973 | 795 | 0.913 | 0.973 | 1714 | 0.074 | 0.320 | 1462 | 0.403 | 0.770 | 956 | 0.109 | 0.328 | 3243 | 0.990 | 0.990 |
| BRD-K79633074-001-01-8 | xilobam | 0.953 | | | | 346.5 | 0.942 | 1.000 | 363 | 0.328 | 0.778 | 906 | 0.152 | 0.778 | 604 | 0.733 | 0.962 | 813.5 | 0.486 | 0.928 | 1674.5 | 0.959 | 0.959 |
| BRD-K79821389-001-03-5 | rubitecan | 0.372 | 75 | 0.165 | 0.384 | 2080 | 0.916 | 0.998 | 1256 | 0.008 | 0.171 | 4703 | 0.549 | 0.960 | 4085 | 0.283 | 0.541 | 3916 | 0.747 | 0.969 | 5290 | 0.769 | 0.969 |
| BRD-K79930101-300-03-7 | GW-583340 | 0.995 | | | | 171 | 0.430 | 0.995 | 319 | 0.680 | 0.995 | 744 | 0.020 | 0.210 | 767 | 0.635 | 0.995 | 256 | 0.014 | 0.059 | 1286 | 0.347 | 0.995 |
| BRD-K80060353-303-03-9 | IEM1754 | 0.284 | | | | 88 | 0.659 | 1.000 | 197 | 0.015 | 0.319 | 737 | 0.213 | 1.000 | 450.5 | 0.209 | 1.000 | 252 | 0.717 | 0.890 | 505.5 | 0.821 | 0.890 |
| BRD-K80343549-001-02-6 | TAK-285 | 0.569 | | | | 18 | 0.076 | 0.408 | 568 | 0.625 | 0.937 | 966 | 0.004 | 0.091 | 690.5 | 0.097 | 0.408 | 1122 | 0.323 | 0.616 | 725 | 0.134 | 0.401 |
| BRD-K80431395-001-07-3 | triciribine | 0.615 | | | | 882 | 0.328 | 0.766 | 543 | 0.097 | 0.455 | 1727 | 0.006 | 0.061 | 1963 | 0.207 | 0.621 | 1281 | 0.125 | 0.238 | 2554 | 0.344 | 0.537 |
| BRD-K80480517-001-02-4 | repsox | 0.968 | | | | 342.5 | 0.949 | 0.968 | 400.5 | 0.483 | 0.968 | 1041 | 0.037 | 0.260 | 871 | 0.968 | 0.968 | 464.5 | 0.691 | 0.968 | 1199 | 0.019 | 0.204 |
| BRD-K80608265-001-01-6 | erteberel | 0.200 | | | | 33 | 0.195 | 1.000 | 169 | 0.747 | 0.923 | 473 | 0.204 | 1.000 | 364 | 0.203 | 1.000 | 429 | 0.985 | 0.985 | 942 | 0.982 | 0.985 |
| BRD-K80700417-001-04-2 | SGI-1776 | 0.934 | | | | 65 | 0.070 | 0.809 | 378 | 0.178 | 0.845 | 702 | 0.278 | 0.845 | 600 | 0.616 | 0.934 | 570 | 0.937 | 1.000 | 1240 | 0.970 | 0.970 |
| BRD-K80725821-300-02-9 | RS-16566 | 0.869 | | | | 416 | 0.736 | 0.869 | 806 | 0.786 | 0.869 | 1725 | 0.901 | 0.946 | 950 | 0.051 | 0.539 | 928 | 0.569 | 0.869 | 1932 | 0.785 | 0.869 |
| BRD-K81197548-003-01-4 | NSC-632839 | 0.942 | | | | 5 | 0.052 | 0.542 | 387 | 0.471 | 1.000 | 1333.5 | 0.151 | 1.000 | 816 | 0.599 | 1.000 | 546 | 0.662 | 0.927 | 1282 | 0.604 | 0.907 |
| BRD-K81330143-001-04-8 | tideglusib | 0.665 | | | | 176 | 0.945 | 0.983 | 240 | 0.039 | 0.818 | 1110 | 0.670 | 0.983 | 805 | 0.983 | 0.983 | 290 | 0.582 | 0.983 | 935 | 0.829 | 0.983 |
| BRD-K81405859-300-01-4 | JTE-607 | 0.706 | 10 | 0.058 | 0.602 | 482 | 0.542 | 0.876 | 322 | 0.143 | 0.602 | 1371 | 0.954 | 0.972 | 1465 | 0.505 | 0.876 | 621 | 0.327 | 0.706 | 1325 | 0.927 | 0.972 |
| BRD-K81418486-001-44-2 | vorinostat | 0.356 | | | | 266.5 | 0.788 | 0.989 | 503 | 0.268 | 0.939 | 911.5 | 0.590 | 0.907 | 871.5 | 0.166 | 0.876 | 710.5 | 0.173 | 0.364 | 623 | 0.074 | 0.876 |
| BRD-K81418486-001-47-5 | vorinostat | 0.581 | 129 | 0.323 | 0.667 | 955 | 0.445 | 0.733 | 1720 | 0.669 | 0.852 | 4829 | 0.890 | 0.981 | 2848 | 0.104 | 0.581 | 2328 | 0.097 | 0.581 | 3753 | 0.210 | 0.631 |
| BRD-K81458380-001-04-9 | CX-4945 | | | | | 28 | 0.182 | 1.000 | 232 | 0.845 | 0.967 | 291 | 0.535 | 0.871 | 261 | 0.936 | 0.977 | 210 | 0.384 | 0.807 | 414 | 0.374 | 0.807 |
| BRD-K81473043-001-14-6 | tanespimycin | 1.000 | 238 | 0.522 | 1.000 | 2035 | 0.895 | 1.000 | 2412 | 0.824 | 1.000 | 4122 | 0.193 | 0.677 | 5020 | 0.843 | 1.000 | 4502.5 | 0.961 | 1.000 | 5047 | 0.365 | 1.000 |
| BRD-K81473043-001-19-5 | tanespimycin | 0.998 | 151 | 0.643 | 0.998 | 656 | 0.871 | 0.998 | 382 | 0.534 | 0.998 | 1766 | 0.658 | 0.998 | 1196 | 0.485 | 0.998 | 821 | 0.901 | 0.998 | 1874 | 0.649 | 0.998 |
| BRD-K81548480-001-01-2 | AC-264613 | 0.669 | | | | 224 | 0.953 | 0.998 | 483 | 0.301 | 0.702 | 981 | 0.155 | 0.702 | 1462 | 0.283 | 0.702 | 720 | 0.372 | 0.650 | 1871 | 0.744 | 0.919 |
| BRD-K81613478-001-01-8 | gastrodin | 0.342 | | | | 480.5 | 0.859 | 0.988 | 419 | 0.939 | 0.988 | 462 | 0.045 | 0.471 | 409 | 0.007 | 0.139 | 294 | 0.252 | 0.617 | 911.5 | 0.223 | 1.000 |
| BRD-K81645907-001-01-1 | captamine | 0.537 | | | | 272.5 | 0.682 | 0.950 | 96 | 0.644 | 0.950 | 896.5 | 0.225 | 1.000 | 671.5 | 0.692 | 0.950 | 232 | 0.192 | 0.537 | 539 | 0.513 | 0.950 |
| BRD-K81694556-003-01-9 | VTP-27999 | 0.761 | | | | 89 | 0.620 | 0.761 | 213 | 0.842 | 0.884 | 692 | 0.434 | 0.761 | 203 | 0.044 | 0.919 | 411 | 0.665 | 0.761 | 743 | 0.583 | 0.761 |
| BRD-K81728688-001-04-4 | WZ8040 | 1.000 | | | | 325 | 0.213 | 0.601 | 1250 | 0.844 | 1.000 | 1345 | 0.000 | 0.009 | 1139 | 0.128 | 0.537 | 1130 | 0.247 | 0.601 | 2047 | 0.002 | 0.025 |
| BRD-K81801188-001-02-8 | PP-121 | 0.923 | 211 | 0.465 | 0.923 | 1693 | 0.635 | 0.923 | 2252 | 0.522 | 0.923 | 2925 | 0.003 | 0.056 | 5980 | 0.992 | 0.992 | 3680 | 0.622 | 0.923 | 5089 | 0.571 | 0.923 |
| BRD-K81916719-001-13-9 | triclabendazole | 0.822 | | | | 1083 | 0.867 | 0.992 | 814 | 0.454 | 0.822 | 2227 | 0.092 | 0.787 | 2869 | 0.758 | 0.992 | 1783 | 0.038 | 0.787 | 3751 | 0.897 | 0.992 |
| BRD-K82091397-001-03-0 | SB-239063 | 0.348 | | | | 789 | 0.975 | 0.990 | 459 | 0.385 | 0.796 | 1732 | 0.557 | 0.798 | 1124 | 0.177 | 1.000 | 1243 | 0.523 | 0.798 | 2650 | 0.901 | 0.990 |
| BRD-K82135108-001-04-3 | elesclomol | 0.969 | | | | 540 | 0.359 | 0.852 | 1014 | 0.703 | 0.865 | 2289 | 0.776 | 0.865 | 958 | 0.352 | 0.852 | 1238 | 0.190 | 0.852 | 3316 | 0.685 | 0.865 |
| BRD-K82244583-001-01-3 | MEK162 | 0.390 | | | | 1276 | 0.534 | 1.000 | 864.5 | 0.080 | 0.421 | 2655 | 0.042 | 0.292 | 3135 | 0.988 | 1.000 | 2496 | 0.712 | 1.000 | 4258 | 0.961 | 1.000 |
| BRD-K82746043-001-15-1 | navitoclax | 0.927 | | | | 634 | 0.751 | 0.927 | 685 | 0.127 | 0.546 | 2330 | 0.973 | 0.974 | 936 | 0.069 | 0.546 | 607.5 | 0.168 | 0.393 | 2473 | 0.974 | 0.974 |
| BRD-K82746043-001-19-3 | navitoclax | | 158.5 | 0.933 | 0.954 | 241 | 0.852 | 1.000 | 450.5 | 0.629 | 1.000 | 624 | 0.653 | 1.000 | 775 | 0.423 | 1.000 | 552 | 0.111 | 1.000 | 357 | 0.893 | 1.000 |
| BRD-K82818427-001-02-2 | batimastat | 1.000 | 223 | 0.564 | 0.846 | 957 | 0.417 | 0.767 | 1681 | 0.363 | 0.767 | 2874 | 0.011 | 0.116 | 3301 | 0.206 | 0.756 | 3451 | 0.861 | 1.000 | 4463 | 0.509 | 0.823 |
| BRD-K82823804-001-04-1 | PD-407824 | 0.645 | 79 | 0.307 | 0.645 | 793 | 0.970 | 0.995 | 329 | 0.338 | 0.645 | 2155 | 0.995 | 0.995 | 1391 | 0.203 | 0.645 | 1101 | 0.442 | 0.773 | 2082 | 0.799 | 0.932 |
| BRD-K82850175-001-01-1 | navarixin | 0.253 | | | | | | | 244 | 0.204 | 0.970 | 733 | 0.005 | 0.115 | 858.5 | 0.882 | 0.976 | 772 | 0.490 | 0.857 | 2093 | 0.976 | 0.976 |
| BRD-K82861010-001-02-5 | daclatasvir | 0.451 | | | | 55 | 0.304 | 1.000 | 513 | 0.689 | 0.890 | 487 | 0.034 | 0.711 | 355 | 0.692 | 0.890 | 390 | 0.522 | 0.890 | 686 | 0.339 | 0.647 |
| BRD-K82928847-001-04-7 | ACY-1215 | 0.837 | | | | 233 | 0.494 | 0.943 | 740 | 0.855 | 0.986 | 1250 | 0.025 | 0.266 | 714 | 0.009 | 0.182 | 944 | 0.782 | 0.967 | 1247 | 0.104 | 0.364 |
| BRD-K82967180-001-01-6 | LY3023414 | 1.000 | 97 | 0.210 | 0.491 | 1959 | 0.837 | 1.000 | 1573 | 0.047 | 0.243 | 5185 | 0.688 | 0.963 | 3422 | 0.019 | 0.134 | 2347 | 0.004 | 0.071 | 4915 | 0.388 | 0.686 |
| BRD-K83029223-001-01-3 | litronesib | 0.607 | | | | 668 | 0.965 | 0.992 | 657 | 0.079 | 0.832 | 2011 | 0.375 | 0.715 | 1920 | 0.271 | 0.641 | 1905 | 0.947 | 0.992 | 1095 | 0.012 | 0.258 |
| BRD-K83699324-001-01-1 | NMS-1286937 | 0.754 | 248 | 0.692 | 0.910 | 1427 | 0.924 | 1.000 | 1081 | 0.530 | 0.856 | 2370 | 0.025 | 0.257 | 2761 | 0.357 | 0.754 | 2420 | 0.826 | 0.964 | 1971 | 0.005 | 0.106 |
| BRD-K83834509-001-01-1 | PTC-209 | 0.454 | | | | 28 | 0.135 | 0.898 | 181 | 0.257 | 0.898 | 450 | 0.254 | 0.898 | 508 | 0.432 | 0.826 | 763 | 0.904 | 0.958 | 586 | 0.263 | 0.552 |
| BRD-K83837640-001-04-8 | JNJ-26481585 | 0.665 | 20 | 0.059 | 0.249 | 957 | 0.367 | 0.665 | 1225 | 0.516 | 0.730 | 2329 | 0.025 | 0.223 | 2162 | 0.088 | 0.264 | 1634 | 0.394 | 0.665 | 2426 | 0.079 | 0.264 |
| BRD-K83963101-001-06-9 | MLN-8054 | 0.719 | 175 | 0.597 | 0.835 | 901 | 0.819 | 0.905 | 841 | 0.452 | 0.803 | 2251 | 0.372 | 0.803 | 1427 | 0.314 | 0.803 | 981 | 0.454 | 0.803 | 2212 | 0.330 | 0.803 |
| BRD-K83988098-001-02-0 | alvespimycin | 0.877 | 50 | 0.175 | 0.718 | 435 | 0.834 | 0.997 | 237 | 0.205 | 0.718 | 1392 | 0.392 | 0.997 | 1137 | 0.829 | 0.997 | 452 | 0.552 | 0.997 | 1338 | 0.465 | 0.997 |
| BRD-K83988098-003-03-4 | alvespimycin | 1.000 | | | | 1307 | 0.920 | 1.000 | 1519 | 0.504 | 0.815 | 3014 | 0.386 | 0.773 | 3396 | 0.836 | 1.000 | 3625 | 0.975 | 1.000 | 2579 | 0.027 | 0.566 |
| BRD-K84163249-001-01-9 | ercalcitriol | 0.936 | | | | 565 | 0.891 | 0.936 | 387.5 | 0.761 | 0.936 | 1035.5 | 0.027 | 0.279 | 700.5 | 0.249 | 1.000 | 617 | 0.770 | 0.936 | 1730.5 | 0.941 | 0.941 |
| BRD-K84564571-001-02-4 | telatinib | 0.964 | | | | 94.5 | 0.379 | 0.942 | 164 | 0.203 | 0.801 | 128.5 | 0.069 | 0.729 | 150 | 0.867 | 1.000 | 229.5 | 0.643 | 1.000 | 456 | 0.918 | 0.918 |
| BRD-K84683831-001-01-0 | radezolid | 0.911 | | | | 395 | 0.776 | 0.990 | 205 | 0.475 | 1.000 | 686 | 0.006 | 0.128 | 1017 | 0.709 | 0.990 | 596 | 0.014 | 0.036 | 1115 | 0.509 | 0.911 |
| BRD-K84868168-001-01-0 | erdafitinib | 0.613 | 7 | 0.058 | 0.607 | 314 | 0.742 | 0.959 | 528 | 0.570 | 0.921 | 1222 | 0.976 | 0.976 | 358 | 0.014 | 0.284 | 631 | 0.397 | 0.758 | 551 | 0.871 | 0.976 |
| BRD-K84937637-001-09-9 | sirolimus | 0.999 | 101 | 0.297 | 0.781 | 919 | 0.355 | 0.801 | 1876 | 0.974 | 0.999 | 3435 | 0.803 | 0.999 | 1675 | 0.179 | 0.536 | 802 | 0.001 | 0.016 | 2680 | 0.172 | 0.536 |
| BRD-K85014146-001-02-5 | AZD2461 | 0.464 | | | | 99.5 | 0.879 | 1.000 | 197.5 | 0.640 | 1.000 | 414.5 | 0.803 | 1.000 | 215.5 | 0.162 | 1.000 | 131.5 | 0.606 | 1.000 | 777 | 0.944 | 0.944 |
| BRD-K85140930-001-12-7 | aphidicolin | 0.403 | 304 | 0.871 | 0.972 | 1104 | 0.961 | 0.972 | 1231 | 0.238 | 0.834 | 3316 | 0.493 | 0.940 | 3465 | 0.865 | 0.972 | 2197 | 0.744 | 0.972 | 3639 | 0.626 | 0.972 |
| BRD-K85146014-314-05-4 | foscarnet | 0.876 | | | | 70 | 0.520 | 0.865 | 309 | 0.396 | 0.865 | 431 | 0.099 | 0.863 | 404 | 0.535 | 0.865 | 195 | 0.459 | 0.865 | 618 | 0.816 | 0.978 |
| BRD-K85307935-236-09-5 | novobiocin | 0.903 | | | | 192.5 | 0.692 | 0.962 | 180.5 | 0.625 | 0.962 | 701 | 0.050 | 0.349 | 269 | 0.039 | 0.349 | 382 | 0.461 | 0.807 | 1092.5 | 0.779 | 0.962 |
| BRD-K85402309-001-06-6 | dovitinib | 0.602 | | | | 240 | 0.943 | 0.985 | 798 | 0.869 | 0.962 | 1975 | 0.622 | 0.870 | 1412 | 0.456 | 0.839 | 1149 | 0.603 | 0.870 | 1965 | 0.871 | 0.962 |
| BRD-K85402309-043-01-9 | dovitinib | 0.950 | | | | | | | 100 | 0.505 | 1.000 | 18 | 0.138 | 1.000 | 194.5 | 0.230 | 1.000 | 156 | 0.248 | 0.473 | 378 | 0.901 | 0.950 |
| BRD-K85554912-001-06-3 | ivermectin | 0.898 | | | | 65 | 0.058 | 0.612 | 65 | 0.058 | 0.612 | 1017 | 0.429 | 1.000 | 854 | 0.822 | 0.898 | 550 | 0.788 | 0.898 | 738.5 | 0.348 | 1.000 |
| BRD-K85606544-001-09-1 | neratinib | | 108.5 | 0.902 | 0.997 | 54 | 0.412 | 0.721 | | | | 361 | 0.001 | 0.026 | 441.5 | 0.381 | 0.721 | 254.5 | 0.561 | 0.842 | 356 | 0.052 | 0.545 |
| BRD-K85751432-001-03-3 | refametinib | 0.999 | | | | 904 | 0.411 | 0.785 | 812 | 0.268 | 0.562 | 2053 | 0.016 | 0.111 | 1951 | 0.591 | 0.999 | 2273 | 0.963 | 0.999 | 3509 | 0.991 | 0.999 |
| BRD-K85849709-001-01-5 | AZD2858 | 0.513 | 9 | 0.055 | 0.361 | 327 | 0.775 | 0.999 | 473 | 0.186 | 0.558 | 1183 | 0.847 | 0.999 | 1531.5 | 0.646 | 0.904 | 913 | 0.041 | 0.361 | 1656.5 | 0.920 | 0.999 |
| BRD-K85872723-001-15-9 | kynurenic-acid | 0.836 | 68.5 | 0.665 | 0.836 | 136 | 0.722 | 0.836 | 68.5 | 0.665 | 0.836 | 268 | 0.796 | 0.836 | 268 | 0.796 | 0.836 | 452.5 | 0.507 | 0.836 | 522 | 0.619 | 0.836 |
| BRD-K85920262-001-02-1 | WAY-600 | 0.918 | 14 | 0.038 | 0.160 | 1167 | 0.509 | 0.918 | 2091 | 0.860 | 0.998 | 5006 | 0.995 | 0.998 | 1830 | 0.004 | 0.020 | 2205 | 0.142 | 0.437 | 3815 | 0.569 | 0.918 |
| BRD-K86003836-001-10-5 | flubendazole | 0.414 | 177 | 0.425 | 0.743 | 1454 | 0.763 | 1.000 | 1670 | 0.161 | 0.414 | 3211 | 0.127 | 0.414 | 2924 | 0.087 | 0.414 | 3203 | 0.647 | 0.997 | 4056 | 0.287 | 0.603 |
| BRD-K86525559-001-07-8 | AZD7762 | 0.226 | 26 | 0.059 | 0.326 | 1778 | 0.706 | 0.875 | 2050 | 0.316 | 0.755 | 5899 | 0.995 | 0.995 | 4106 | 0.199 | 0.696 | 3296 | 0.639 | 0.875 | 4941 | 0.323 | 0.755 |
| BRD-K86687746-001-01-3 | R306465 | 0.021 | 173 | 0.452 | 0.769 | 851 | 0.652 | 0.855 | 974 | 0.043 | 0.456 | 3073 | 0.117 | 0.492 | 2906 | 0.195 | 0.585 | 1829 | 0.141 | 0.493 | 3702 | 0.405 | 0.769 |
| BRD-K86727142-001-12-4 | embelin | 0.572 | | | | 131.5 | 0.919 | 0.991 | 70 | 0.476 | 0.859 | 110 | 0.073 | 0.572 | 396.5 | 0.399 | 0.859 | 595 | 0.955 | 0.991 | 966 | 0.867 | 1.000 |
| BRD-K86797399-001-05-1 | SB-939 | 0.324 | 14 | 0.033 | 0.174 | 1885 | 0.808 | 0.998 | 1750 | 0.246 | 0.517 | 4570 | 0.355 | 0.670 | 3224 | 0.011 | 0.112 | 3060 | 0.180 | 0.421 | 4541 | 0.146 | 0.382 |
| BRD-K86816506-001-01-8 | PF-04457845 | 0.146 | | | | 25 | 0.101 | 0.966 | 379.5 | 0.813 | 0.898 | 583 | 0.284 | 1.000 | 1253.5 | 0.763 | 0.898 | 536 | 0.188 | 0.358 | 1326 | 0.733 | 0.898 |
| BRD-K86887724-001-10-6 | dofetilide | 0.885 | 135 | 0.917 | 0.922 | 65 | 0.414 | 0.885 | 112 | 0.276 | 0.885 | 383 | 0.576 | 0.885 | 690 | 0.457 | 0.885 | 36 | 0.034 | 0.448 | 426 | 0.739 | 0.885 |
| BRD-K86930074-001-05-0 | cediranib | 0.532 | 59 | 0.121 | 0.378 | 1009 | 0.902 | 0.996 | 1141 | 0.063 | 0.366 | 2969 | 0.070 | 0.366 | 2668 | 0.126 | 0.378 | 2404 | 0.887 | 0.996 | 3983 | 0.920 | 0.996 |
| BRD-K87112191-001-04-5 | PP242 | 0.725 | 176 | 0.429 | 0.804 | 1018 | 0.498 | 0.804 | 1875 | 0.564 | 0.847 | 3141 | 0.321 | 0.804 | 5016 | 0.982 | 0.991 | 2420 | 0.198 | 0.725 | 4523 | 0.659 | 0.885 |
| BRD-K87124298-001-03-4 | OSI-027 | 0.904 | 228 | 0.524 | 0.904 | 1311 | 0.523 | 0.904 | 2599 | 0.880 | 0.974 | 3172 | 0.017 | 0.167 | 3913 | 0.864 | 0.974 | 2534 | 0.095 | 0.379 | 5246 | 0.881 | 0.974 |
| BRD-K87349682-347-03-8 | norepinephrine | 0.968 | | | | 597 | 0.753 | 0.968 | 538 | 0.307 | 1.000 | 1047 | 0.023 | 0.242 | 1327 | 0.856 | 0.982 | 834 | 0.701 | 0.968 | 2000 | 0.783 | 0.968 |
| BRD-K87510569-001-02-8 | RS-504393 | 0.644 | 61 | 0.355 | 0.644 | 297 | 0.986 | 0.986 | 326 | 0.289 | 0.644 | 784 | 0.274 | 0.644 | 400 | 0.571 | 0.705 | 277 | 0.368 | 0.644 | 638 | 0.405 | 0.644 |
| BRD-K87512222-019-01-7 | taltobulin | 0.367 | 97 | 0.252 | 0.538 | 1625 | 0.998 | 1.000 | 1284 | 0.227 | 0.538 | 2833 | 0.040 | 0.367 | 3201 | 0.256 | 0.538 | 2126 | 0.362 | 0.645 | 4051 | 0.770 | 0.972 |
| BRD-K87696786-003-04-4 | LY456236 | 0.529 | | | | 518 | 0.705 | 0.928 | 797 | 0.795 | 0.928 | 1225 | 0.127 | 1.000 | 884 | 0.157 | 1.000 | 682.5 | 0.542 | 0.888 | 1575.5 | 0.896 | 0.941 |
| BRD-K87700323-003-05-1 | cetylpyridinium | 0.391 | | | | 72 | 0.230 | 0.723 | 922 | 0.044 | 0.229 | 2047 | 0.044 | 0.229 | 1583 | 0.054 | 0.229 | 1726 | 0.895 | 1.000 | 3063 | 0.726 | 1.000 |
| BRD-K87737963-001-06-0 | cyt387 | 0.300 | | | | | | | 130 | 0.932 | 0.970 | 473 | 0.301 | 1.000 | 300 | 0.629 | 1.000 | 743 | 0.970 | 0.970 | 765 | 0.926 | 0.970 |
| BRD-K87782578-001-01-4 | AVL-292 | 0.976 | | | | 43 | 0.239 | 1.000 | 60 | 0.377 | 1.000 | 996 | 0.776 | 0.976 | 548 | 0.133 | 0.932 | 322 | 0.152 | 0.319 | 627 | 0.704 | 0.976 |
| BRD-K87827419-001-16-8 | melphalan | 0.091 | | | | | | | 928 | 0.962 | 0.994 | 784 | 0.938 | 0.994 | 672 | 0.149 | 1.000 | 791 | 0.589 | 0.994 | 1155 | 0.994 | 0.994 |
| BRD-K87909389-003-03-4 | alvocidib | 0.925 | 317 | 0.774 | 0.925 | 1202 | 0.756 | 0.925 | 2104 | 0.799 | 0.925 | 3341 | 0.136 | 0.716 | 3772 | 0.272 | 0.925 | 3127 | 0.636 | 0.925 | 4348 | 0.551 | 0.925 |
| BRD-K88429204-001-35-1 | pyrimethamine | 0.975 | 39 | 0.114 | 0.481 | | | | 688 | 0.332 | 0.798 | 1658 | 0.871 | 0.975 | 1459 | 0.802 | 0.975 | 934 | 0.350 | 0.798 | 1872 | 0.528 | 0.874 |
| BRD-K88506063-001-01-8 | KD025 | 1.000 | 114 | 0.414 | 0.862 | 188 | 0.776 | 1.000 | 1008 | 0.901 | 1.000 | 1407 | 0.351 | 0.862 | 1375 | 0.483 | 0.862 | 1051 | 0.039 | 0.233 | 1667 | 0.393 | 0.862 |
| BRD-K88510285-001-11-1 | bortezomib | 0.420 | 180 | 0.394 | 0.989 | 2301 | 0.982 | 0.989 | 2364 | 0.806 | 0.989 | 4575 | 0.625 | 0.989 | 4690 | 0.698 | 0.989 | 3597 | 0.702 | 0.989 | 5040 | 0.388 | 0.989 |
| BRD-K88510285-001-17-8 | bortezomib | 0.788 | 112 | 0.276 | 0.788 | 1381 | 0.721 | 0.954 | 1627 | 0.528 | 0.909 | 3705 | 0.360 | 0.788 | 4964 | 0.954 | 0.954 | 1773 | 0.470 | 0.897 | 3204 | 0.160 | 0.788 |
| BRD-K88625236-001-04-2 | nonoxynol-9 | 0.475 | 170 | 0.901 | 0.961 | 418 | 0.906 | 0.961 | 460 | 0.747 | 0.961 | 423 | 0.005 | 0.112 | 879 | 0.785 | 0.961 | 682 | 0.162 | 0.678 | 1400 | 0.937 | 0.961 |

Suppl. Table 9

| broad_id | name | colorectal_MWU_STAT | colorectal_PVAL | colorectal_PVAL_ADJ | skin_MWU_STAT | skin_PVAL | skin_PVAL_ADJ | urinary_tract_MWU_STAT | urinary_tract_PVAL | urinary_tract_PVAL_ADJ | lung_MWU_STAT | lung_PVAL | lung_PVAL_ADJ | ovary_MWU_STAT | ovary_PVAL | ovary_PVAL_ADJ | breast_MWU_STAT | breast_PVAL | breast_PVAL_ADJ | pancreas_MWU_STAT | pancreas_PVAL | pancreas_PVAL_ADJ | nervous_system_MWU_STAT | central_nervous_system_PVAL | nervous_system_PVAL_ADJ | soft_tissue_MWU_STAT | soft_tissue_PVAL | soft_tissue_PVAL_ADJ | kidney_MWU_STAT | kidney_PVAL | kidney_PVAL_ADJ | gastric_MWU_STAT | gastric_PVAL | gastric_PVAL_ADJ | bone_MWU_STAT | bone_PVAL | bone_PVAL_ADJ | prostate_MWU_STAT | prostate_PVAL | prostate_PVAL_ADJ | peripheral_nervous_system_MWU_STAT | peripheral_nervous_system_PVAL |
|---|---|---|---|---|---|---|---|---|---|---|---|---|---|---|---|---|---|---|---|---|---|---|---|---|---|---|---|---|---|---|---|---|---|---|---|---|---|---|---|---|---|---|
| BRD-K88677950-001-03-9 | PD-198306 | 1488 | 0.000 | 0.004 | 2817 | 0.000 | 0.004 | 1994 | 0.397 | 0.758 | 8329 | 0.970 | 0.996 | 3692 | 0.971 | 0.996 | 1830 | 0.850 | 0.996 | 2753 | 0.285 | 0.665 | 2098 | 0.831 | 0.996 | 696 | 0.215 | 0.565 | 657 | 0.374 | 0.758 | 1093 | 0.114 | 0.399 | 832.5 | 0.731 | 0.996 | 483 | 0.948 | 0.996 | 435 | 0.505 |
| BRD-K88807631-001-01-3 | elacridar | 986 | 0.471 | 0.856 | 2366 | 0.465 | 1.000 | 716 | 0.011 | 0.221 | 2960 | 0.303 | 1.000 | 1524 | 0.608 | 0.912 | 1358.5 | 0.799 | 0.956 | 924 | 0.912 | 0.956 | 1934 | 0.553 | 0.893 | 195 | 0.136 | 0.356 | 1484 | 0.832 | 0.956 | 802 | 0.487 | 0.856 | 147 | 0.240 | 0.559 | 207.5 | 0.956 | 0.956 | | |
| BRD-K89014967-001-04-3 | AS-703026 | 1294 | 0.006 | 0.061 | 977 | 0.000 | 0.000 | 699 | 0.045 | 0.262 | 6499 | 1.000 | 1.000 | 625 | 0.592 | 1.000 | 1896 | 0.987 | 1.000 | 2013 | 0.164 | 0.492 | 1615 | 0.984 | 1.000 | 550 | 0.398 | 0.853 | 1595 | 0.980 | 1.000 | 882 | 0.214 | 0.562 | 992 | 0.996 | 1.000 | | | | 441 | 0.406 |
| BRD-K89015388-001-01-7 | GSK923295 | 1754 | 0.630 | 0.903 | 2967 | 0.412 | 1.000 | 2303 | 0.556 | 1.000 | 5882 | 0.160 | 0.840 | 1104 | 0.008 | 0.172 | 2062 | 0.976 | 1.000 | 2322 | 0.720 | 0.903 | 2378 | 0.504 | 1.000 | 543 | 0.033 | 0.346 | 1946 | 1.000 | 1.000 | 1063 | 0.080 | 0.558 | 720 | 0.358 | 0.578 | 339 | 0.328 | 0.575 | 137 | 0.119 |
| BRD-K89053832-001-01-0 | VLX600 | 2119.5 | 0.318 | 0.668 | 3856 | 0.950 | 0.961 | 1852.5 | 0.187 | 1.000 | 7813 | 0.484 | 0.782 | 2872 | 0.209 | 0.549 | 2191.5 | 0.851 | 0.961 | 3354.5 | 0.926 | 0.961 | 2623.5 | 0.440 | 0.771 | 450 | 0.067 | 0.234 | 1400.5 | 0.247 | 0.576 | 1104 | 0.098 | 1.000 | 703.5 | 0.195 | 0.549 | 649.5 | 0.627 | 0.878 | 105 | 0.057 |
| BRD-K89121095-001-01-7 | tedizolid-phosphate | 2436 | 0.987 | 0.987 | 4216 | 0.847 | 0.987 | 2200 | 0.141 | 0.611 | 7613 | 0.132 | 0.611 | 3220 | 0.827 | 0.987 | 2352 | 0.385 | 0.897 | 2129 | 0.437 | 0.918 | 3337 | 0.811 | 0.987 | 1014 | 0.236 | 0.708 | 390 | 0.337 | 0.886 | 1189 | 0.146 | 0.611 | 1103 | 0.963 | 0.987 | 907 | 0.958 | 0.987 | 251 | 0.092 |
| BRD-K89299012-001-06-0 | DCEBIO | 2948 | 0.701 | 0.982 | 3762 | 0.097 | 0.554 | 2427 | 0.205 | 0.554 | 8417 | 0.157 | 0.554 | 2122 | 0.180 | 0.554 | 1768 | 0.063 | 0.554 | 4651 | 0.997 | 0.997 | 3672 | 0.576 | 0.982 | 1417 | 0.896 | 0.984 | 2075 | 0.881 | 0.984 | 2182 | 0.694 | 0.982 | 724 | 0.163 | 0.554 | 778 | 0.768 | 0.984 | 393 | 0.098 |
| BRD-K89561498-001-01-7 | SN-38 | 863 | 0.448 | 0.867 | 2624 | 0.578 | 0.867 | 1129 | 0.557 | 0.867 | 4799 | 0.712 | 0.880 | 1184 | 0.291 | 1.000 | 1759 | 0.978 | 0.978 | 1954 | 0.707 | 0.880 | 1793 | 0.878 | 0.971 | 317 | 0.012 | 0.033 | 1281 | 0.811 | 0.946 | 1238 | 0.563 | 0.867 | 608 | 0.619 | 0.867 | 216 | 0.147 | 0.342 | | |
| BRD-K89704198-001-12-3 | vincamine | 685 | 0.831 | 0.989 | 501 | 0.133 | 1.000 | 469 | 0.791 | 0.989 | 1553 | 0.321 | 0.750 | 382 | 0.007 | 0.028 | 468 | 0.946 | 0.989 | 312 | 0.420 | 0.801 | 926 | 0.912 | 0.989 | 131 | 0.041 | 0.124 | 207 | 0.538 | 0.942 | 253 | 0.188 | 0.493 | 354 | 0.989 | 0.989 | 237 | 0.968 | 0.989 | | |
| BRD-K89714990-001-01-5 | tepoxalin | 1319 | 0.409 | 1.000 | 2312 | 0.623 | 0.956 | 1523 | 0.749 | 0.956 | 4347 | 0.622 | 0.956 | 1459 | 0.121 | 0.364 | 976 | 0.901 | 0.956 | 1224 | 0.911 | 0.956 | 1750 | 0.233 | 0.544 | 660 | 0.785 | 0.956 | 628 | 0.005 | 0.025 | 732 | 0.436 | 0.917 | 214 | 0.040 | 0.166 | 305 | 0.820 | 0.956 | 152 | 0.223 |
| BRD-K89839824-001-05-3 | raltitrexed | 4161 | 0.021 | 0.324 | 8872 | 0.959 | 0.978 | 3511 | 0.092 | 0.386 | 15240 | 0.144 | 0.436 | 6017 | 0.483 | 0.845 | 5618 | 0.978 | 0.978 | 6779 | 0.786 | 0.978 | 6553 | 0.460 | 0.845 | 2084 | 0.406 | 0.845 | 3798 | 0.761 | 0.978 | 2494 | 0.031 | 0.324 | 1826 | 0.357 | 0.834 | 1008 | 0.686 | 0.978 | 884 | 0.223 |
| BRD-K90027355-001-13-3 | spironolactone | 3352 | 0.511 | 0.826 | 4351 | 0.046 | 0.344 | 2111 | 0.069 | 0.344 | 10197 | 0.470 | 0.822 | 4340 | 0.678 | 0.902 | 2342 | 0.303 | 0.795 | 4720 | 0.973 | 0.990 | 4925 | 0.757 | 0.935 | 1336 | 0.672 | 0.902 | 1712 | 0.082 | 0.344 | 2135 | 0.412 | 0.822 | 1004 | 0.438 | 0.822 | 372 | 0.200 | 0.656 | 181 | 0.024 |
| BRD-K90374350-001-09-2 | chlorquinaldol | 416 | 0.106 | 0.318 | 994 | 0.025 | 0.527 | 1076 | 0.959 | 0.998 | 2291 | 0.221 | 0.516 | 1353 | 0.830 | 0.970 | 545 | 0.402 | 0.844 | 1061 | 0.865 | 0.970 | 1421 | 0.820 | 0.970 | 132 | 0.078 | 0.318 | 739 | 0.998 | 0.998 | 142 | 0.097 | 1.000 | 141 | 0.095 | 0.318 | 251 | 0.878 | 0.970 | 69 | 0.076 |
| BRD-K90382497-001-10-7 | GW-843682X | 2520 | 0.015 | 0.321 | 6113 | 0.999 | 0.999 | 2864 | 0.138 | 0.322 | 9652 | 0.094 | 0.281 | 4561 | 0.888 | 0.981 | 3740 | 0.952 | 0.999 | 4366 | 0.792 | 0.981 | 5013.5 | 0.874 | 0.981 | 1595 | 0.765 | 0.981 | 1343 | 0.680 | 0.981 | 1841 | 0.440 | 0.840 | 1349.5 | 0.888 | 0.981 | 668 | 0.786 | 0.981 | 230 | 0.010 |
| BRD-K90497590-001-04-2 | AR-12 | 2169 | 0.742 | 0.947 | 4390 | 0.966 | 0.992 | 2659 | 0.683 | 0.947 | 7455 | 0.841 | 0.982 | 2092 | 0.214 | 0.644 | 2193 | 0.766 | 0.947 | 2491 | 0.358 | 0.684 | 3606 | 0.489 | 0.734 | 395 | 0.010 | 0.069 | 1706 | 0.992 | 0.992 | 1726 | 0.220 | 0.644 | 763 | 0.313 | 0.684 | 260 | 0.396 | 0.693 | 283 | 0.154 |
| BRD-K90948141-001-01-4 | indiplon | 372 | 0.978 | 0.978 | 551 | 0.028 | 0.582 | 728 | 0.867 | 1.000 | 1626 | 0.977 | 1.000 | 522.5 | 0.230 | 1.000 | 163 | 0.411 | 1.000 | 267 | 0.934 | 1.000 | 711.5 | 0.693 | 0.909 | 147 | 0.311 | 0.503 | 64 | 0.134 | 0.234 | 191 | 0.595 | 0.834 | 0 | 0.044 | 0.093 | 125 | 0.553 | 0.829 | 50 | 0.082 |
| BRD-K91370081-001-25-1 | anisomycin | 4367 | 0.067 | 0.354 | 7994 | 0.680 | 0.952 | 4557 | 0.536 | 0.913 | 16205 | 0.243 | 0.913 | 3725 | 0.001 | 0.026 | 5624 | 0.990 | 0.990 | 7130 | 0.819 | 0.990 | 7893 | 0.955 | 0.990 | 2325 | 0.417 | 0.913 | 3879 | 0.787 | 0.990 | 2277 | 0.053 | 0.354 | 3223 | 0.971 | 0.990 | 844 | 0.430 | 0.913 | 1028 | 0.389 |
| BRD-K91601245-001-12-0 | mercaptopurine | 681 | 0.728 | 0.949 | 988 | 0.833 | 0.949 | 753 | 0.892 | 0.949 | 1502.5 | 0.008 | 0.043 | 756.5 | 0.377 | 1.000 | 279 | 0.838 | 0.949 | 577 | 0.607 | 0.949 | 841.5 | 0.799 | 0.949 | 362.5 | 0.560 | 0.949 | 67 | 0.086 | 0.258 | 426.5 | 0.291 | 1.000 | 108.5 | 0.827 | 0.949 | 55 | 0.337 | 0.884 | | |
| BRD-K91623615-001-05-0 | ABT-751 | 5260 | 0.370 | 0.706 | 8329 | 0.895 | 0.996 | 4462 | 0.247 | 0.673 | 15787 | 0.235 | 0.673 | 4977 | 0.090 | 0.542 | 5517 | 0.996 | 0.996 | 7884 | 0.981 | 0.996 | 6850 | 0.618 | 0.865 | 2396 | 0.314 | 0.673 | 4581 | 0.987 | 0.996 | 3213 | 0.479 | 0.773 | 2218 | 0.188 | 0.673 | 795 | 0.710 | 0.931 | 592 | 0.037 |
| BRD-K91701654-001-03-1 | U-0126 | 314 | 0.165 | 0.577 | 1275 | 0.011 | 0.229 | 1169 | 0.760 | 0.887 | 2726 | 0.760 | 0.887 | 1131 | 0.372 | 0.853 | 659 | 0.718 | 0.887 | 795 | 0.437 | 0.853 | 1087 | 0.754 | 0.887 | 415 | 0.475 | 0.853 | 364 | 0.597 | 0.887 | 350 | 0.036 | 0.381 | 369 | 0.316 | 0.830 | 88 | 0.516 | 0.853 | | |
| BRD-K91825936-001-01-2 | ZK811752 | 1572 | 0.613 | 0.920 | 4314 | 0.983 | 0.983 | 1660 | 0.372 | 0.867 | 8011 | 0.971 | 0.983 | 2452 | 0.307 | 0.867 | 1162 | 0.020 | 0.139 | 2278 | 0.253 | 0.867 | 2496 | 0.764 | 0.963 | 672 | 0.085 | 0.446 | 1287.5 | 0.795 | 0.963 | 1571 | 0.612 | 0.920 | 829.5 | 0.772 | 0.963 | 739 | 0.868 | 0.963 | 459 | 0.610 |
| BRD-K92000912-001-12-6 | AM-251 | 714 | 0.755 | 0.925 | 888.5 | 0.793 | 0.925 | 647 | 0.787 | 0.925 | 2123 | 0.519 | 1.000 | 1396.5 | 0.994 | 0.994 | 457 | 0.102 | 0.723 | 546 | 0.138 | 0.723 | 645.5 | 0.567 | 1.000 | 617.5 | 0.896 | 0.941 | 280 | 0.014 | 0.030 | 333 | 0.270 | 1.000 | 332.5 | 0.555 | 0.897 | 327 | 0.854 | 0.941 | 200 | 0.307 |
| BRD-K92093830-003-30-8 | doxorubicin | 6389 | 0.816 | 0.989 | 7561 | 0.234 | 0.679 | 3989 | 0.097 | 0.634 | 16997 | 0.403 | 0.693 | 5227 | 0.129 | 0.634 | 5537 | 0.950 | 0.989 | 8442 | 0.989 | 0.989 | 7878 | 0.936 | 0.989 | 1276 | 0.001 | 0.022 | 4643 | 0.985 | 0.989 | 2789 | 0.151 | 0.634 | 2600 | 0.444 | 0.693 | 1034 | 0.684 | 0.958 | 1098 | 0.462 |
| BRD-K92150287-066-01-2 | exatecan-mesylate | 703 | 0.217 | 0.608 | 1881 | 0.940 | 0.988 | 638 | 0.232 | 0.608 | 2067 | 0.421 | 0.681 | 535 | 0.348 | 0.620 | 1213 | 0.918 | 0.988 | 1358 | 0.988 | 0.988 | 842 | 0.545 | 0.817 | 440 | 0.297 | 0.620 | 734 | 0.865 | 0.988 | 590 | 0.293 | 0.620 | 428 | 0.977 | 0.988 | 61 | 0.354 | 0.620 | | |
| BRD-K92241597-001-06-0 | tosedostat | 6528 | 0.981 | 0.996 | 4963 | 0.000 | 0.001 | 4224 | 0.238 | 0.757 | 15120.5 | 0.094 | 0.496 | 6211 | 0.816 | 0.996 | 4630 | 0.629 | 0.996 | 8587.5 | 0.995 | 0.996 | 5668 | 0.144 | 0.604 | 1994 | 0.086 | 0.496 | 3126 | 0.962 | 0.996 | 3588 | 0.469 | 0.996 | 2601 | 0.859 | 0.996 | 1219 | 0.908 | 0.996 | 692 | 0.556 |
| BRD-K92428153-001-04-4 | mycophenolate-mofetil | 385 | 0.336 | 0.874 | 464 | 0.426 | 0.874 | 138 | 0.017 | 0.088 | 1755 | 0.866 | 0.874 | 561 | 0.579 | 0.874 | 405 | 0.874 | 0.874 | 501 | 0.361 | 0.874 | 443 | 0.793 | 0.874 | 59 | 0.003 | 0.019 | 405 | 0.874 | 0.874 | 170 | 0.045 | 0.936 | 117 | 0.429 | 0.874 | 164 | 0.749 | 0.874 | 143 | 0.248 |
| BRD-K92428232-001-10-6 | GSK461364 | 3312 | 0.201 | 0.587 | 5543.5 | 0.922 | 0.984 | 2806.5 | 0.168 | 0.587 | 11368.5 | 0.737 | 0.984 | 3871.5 | 0.272 | 0.587 | 3613.5 | 0.984 | 0.984 | 3811 | 0.336 | 0.587 | 4767.5 | 0.970 | 0.984 | 1111 | 0.329 | 0.587 | 1465 | 0.805 | 0.984 | 2194 | 0.281 | 0.587 | 824 | 0.023 | 0.241 | 493 | 0.147 | 0.587 | 315 | 0.007 |
| BRD-K92657060-001-05-7 | zaleplon | 349 | 0.172 | 0.672 | 1530 | 0.479 | 0.887 | 395 | 0.135 | 0.672 | 2031 | 0.475 | 0.887 | 1164 | 0.724 | 0.887 | 400 | 0.854 | 0.944 | 651 | 0.456 | 0.887 | 487 | 0.624 | 0.887 | 100 | 0.047 | 0.495 | 369 | 0.760 | 0.887 | 851 | 0.919 | 0.919 | 149 | 0.466 | 0.887 | 106 | 0.224 | 0.672 | 41 | 0.210 |
| BRD-K92723993-001-06-7 | imatinib | 1581 | 0.821 | 1.000 | 2615 | 0.114 | 0.547 | 1993 | 0.672 | 0.927 | 6172 | 0.221 | 0.663 | 2558 | 0.453 | 0.927 | 2100 | 0.644 | 0.927 | 2772 | 0.689 | 0.927 | 2981 | 0.653 | 0.927 | 471 | 0.118 | 0.547 | 1951 | 0.999 | 0.999 | 1322 | 0.456 | 0.927 | 841 | 0.140 | 0.547 | 283 | 0.519 | 0.927 | 261 | 0.142 |
| BRD-K92991072-001-04-9 | PAC-1 | 405.5 | 0.468 | 0.867 | 561 | 0.270 | 1.000 | 491 | 0.818 | 0.903 | 1300.5 | 0.536 | 0.867 | 341.5 | 0.086 | 0.361 | 241 | 0.497 | 0.867 | 473.5 | 0.759 | 0.903 | 582 | 0.733 | 0.903 | 176.5 | 0.459 | 0.867 | 260 | 0.116 | 0.270 | 309.5 | 0.280 | 1.000 | | | | 45 | 0.312 | 0.935 | 66 | 0.116 |
| BRD-K93034159-001-25-8 | cladribine | 3652 | 0.945 | 0.945 | 3886 | 0.161 | 0.847 | 3438 | 0.927 | 1.000 | 8917 | 0.577 | 0.960 | 2716 | 0.026 | 0.370 | 2419 | 0.963 | 1.000 | 3305 | 0.516 | 0.960 | 2685 | 0.210 | 0.881 | 977 | 0.312 | 0.936 | 1767 | 0.392 | 0.960 | 2460 | 0.681 | 0.960 | 321 | 0.045 | 0.370 | 468 | 0.480 | 0.960 | 229 | 0.717 |
| BRD-K93123848-001-04-1 | RAF265 | 1488 | 0.739 | 0.997 | 2203 | 0.000 | 0.000 | 2582 | 0.960 | 0.997 | 8251.5 | 0.868 | 0.997 | 3005 | 0.298 | 1.000 | 2918.5 | 0.997 | 0.997 | 2412 | 0.336 | 1.000 | 3586.5 | 0.992 | 0.997 | 905 | 0.020 | 0.047 | 1379 | 0.377 | 0.659 | 1485 | 0.530 | 1.000 | 580 | 0.072 | 0.151 | 452 | 0.507 | 0.818 | 631 | 0.576 |
| BRD-K93779381-001-01-9 | ingenol-mebutate | 578 | 0.908 | 1.000 | 755 | 0.809 | 1.000 | 558 | 0.130 | 0.727 | 1943 | 0.172 | 0.727 | 766 | 0.663 | 1.000 | 566 | 0.698 | 1.000 | 861 | 0.391 | 0.821 | 548 | 0.229 | 0.802 | 194 | 0.350 | 0.821 | 338 | 0.173 | 0.727 | 602 | 0.374 | 0.821 | | | | 0 | 0.044 | 0.727 | 264 | 0.968 |
| BRD-K93869735-001-01-1 | taltirelin | 149 | 0.708 | 0.991 | 940 | 0.963 | 1.000 | 466 | 0.311 | 0.987 | 1012 | 0.236 | 0.987 | 599 | 0.423 | 0.987 | 128 | 0.551 | 1.000 | 970 | 0.979 | 1.000 | 385 | 0.397 | 0.987 | 170 | 0.833 | 1.000 | 48 | 0.358 | 0.987 | 148 | 0.295 | 0.987 | 252 | 0.570 | 1.000 | 202 | 0.945 | 1.000 | 104 | 0.364 |
| BRD-K94176593-001-09-4 | TWS-119 | 925.5 | 0.909 | 0.997 | 1471 | 0.997 | 0.997 | 851 | 0.234 | 0.684 | 2610.5 | 0.151 | 0.652 | 1118 | 0.102 | 0.652 | 416 | 0.359 | 0.754 | 1465 | 0.997 | 0.997 | 1390 | 0.963 | 0.997 | 248 | 0.131 | 0.652 | 666 | 0.185 | 0.652 | 379 | 0.035 | 0.652 | 166 | 0.395 | 0.754 | 124 | 0.712 | 0.928 | 114 | 0.173 |
| BRD-K94441233-001-13-0 | mevastatin | 2428 | 0.673 | 0.993 | 3499 | 0.001 | 0.021 | 3012 | 0.945 | 0.993 | 9751 | 0.742 | 0.993 | 4232 | 0.528 | 0.993 | 2301 | 0.943 | 0.993 | 3961 | 0.928 | 0.993 | 3670 | 0.018 | 0.126 | 925 | 0.076 | 0.397 | 2068 | 0.632 | 0.993 | 2729 | 0.963 | 0.993 | 1025 | 0.143 | 0.502 | 540 | 0.584 | 0.993 | 633 | 0.993 |
| BRD-K94455792-001-01-8 | ICG-001 | 1682 | 0.939 | 0.997 | 1928 | 0.293 | 1.000 | 1166.5 | 0.545 | 0.733 | 4681 | 0.987 | 0.997 | 1037 | 0.001 | 0.006 | 1039 | 0.305 | 0.733 | 1537.5 | 0.815 | 0.951 | 1373 | 0.394 | 0.733 | 99 | 0.020 | 0.069 | 920 | 0.997 | 0.997 | 348 | 0.029 | 0.606 | 30 | 0.107 | 0.321 | 393 | 0.738 | 0.912 | 318 | 0.480 |
| BRD-K94485812-001-02-9 | LDN-57444 | 1465 | 0.564 | 0.741 | 3850 | 1.000 | 1.000 | 1319 | 0.339 | 1.000 | 5682 | 0.099 | 1.000 | 2223 | 0.205 | 1.000 | 2393 | 0.949 | 0.997 | 2465 | 0.844 | 0.997 | 2655 | 0.547 | 0.741 | 671 | 0.042 | 0.088 | 1204 | 0.019 | 0.045 | 1136 | 0.434 | 1.000 | 574 | 0.897 | 0.997 | 305 | 0.250 | 0.437 | 461 | 0.330 |
| BRD-K95142244-001-01-5 | talazoparib | 1715 | 0.672 | 0.961 | 2519 | 0.959 | 0.961 | 1048 | 0.294 | 1.000 | 5031 | 0.380 | 0.726 | 1809 | 0.044 | 0.131 | 1924 | 0.961 | 0.961 | 2185 | 0.830 | 0.961 | 1470 | 0.300 | 0.700 | 441 | 0.017 | 0.060 | 934 | 0.908 | 0.961 | 733 | 0.143 | 1.000 | 451 | 0.006 | 0.024 | 210 | 0.901 | 0.961 | 299 | 0.098 |
| BRD-K95292280-001-01-3 | SB-657510 | 364.5 | 0.765 | 0.765 | 762.5 | 0.898 | 0.992 | 249.5 | 0.081 | 0.850 | 875.5 | 0.537 | 1.000 | 525 | 0.788 | 0.957 | 347.5 | 0.688 | 0.957 | 562.5 | 0.576 | 1.000 | 432.5 | 0.821 | 0.957 | 63 | 0.615 | 0.957 | 108 | 0.044 | 0.102 | 167.5 | 0.081 | 0.850 | 12 | 0.086 | 0.180 | | | | 20 | 0.134 |
| BRD-K95435023-001-08-5 | PHA-665752 | 841 | 0.086 | 0.903 | 1685 | 0.007 | 0.147 | 1328 | 0.473 | 0.950 | 5058 | 0.572 | 0.950 | 1550 | 0.213 | 0.747 | 1311 | 0.805 | 0.950 | 1982 | 0.672 | 0.950 | 1992 | 0.905 | 0.950 | 866 | 0.857 | 0.950 | 884 | 0.055 | 0.290 | 607 | 0.138 | 0.579 | 800 | 0.994 | 0.994 | 189 | 0.855 | 0.950 | 204 | 0.899 |
| BRD-K95573441-001-01-2 | PFK-015 | 773 | 0.584 | 0.901 | 2589 | 0.939 | 0.946 | 955 | 0.332 | 0.901 | 3577 | 0.160 | 0.838 | 1780 | 0.613 | 0.901 | 997 | 0.414 | 0.901 | 1442 | 0.525 | 0.901 | 1801 | 0.882 | 0.946 | 446 | 0.559 | 0.901 | 895 | 0.404 | 0.901 | 945 | 0.314 | 0.901 | 558 | 0.852 | 0.946 | 96 | 0.424 | 0.901 | 132 | 0.649 |
| BRD-K95581532-001-01-5 | RN-1734 | 3350 | 0.311 | 0.725 | 5394 | 0.476 | 0.909 | 2677 | 0.051 | 0.477 | 9705 | 0.215 | 0.901 | 4987 | 0.886 | 0.976 | 3227 | 0.758 | 0.976 | 4510 | 0.896 | 0.976 | 5291 | 0.929 | 0.976 | 1507 | 0.220 | 0.576 | 2325 | 0.737 | 0.976 | 1603 | 0.082 | 0.477 | 1152 | 0.984 | 0.984 | 505 | 0.432 | 0.907 | 51 | 0.011 |
| BRD-K95773607-004-03-8 | darifenacin | 790 | 0.996 | 0.996 | 1019.5 | 0.712 | 0.947 | 343 | 0.114 | 0.716 | 2728.5 | 0.857 | 0.947 | 829 | 0.152 | 0.716 | 448.5 | 0.384 | 1.000 | 1072 | 0.813 | 0.947 | 739.5 | 0.380 | 0.726 | 110 | 0.210 | 0.441 | 141 | 0.006 | 0.017 | 286 | 0.132 | 0.716 | 677 | 0.995 | 0.996 | 262.5 | 0.582 | 0.947 | 102 | 0.658 |
| BRD-K96006502-001-02-6 | deflazacort | 713 | 0.241 | 0.643 | 1769 | 0.861 | 0.952 | 1334 | 0.515 | 0.643 | 3963 | 0.785 | 0.916 | 1697.5 | 0.242 | 0.643 | 561.5 | 0.145 | 0.643 | 1377.5 | 0.425 | 0.643 | 2057 | 0.969 | 0.969 | 616 | 0.457 | 0.643 | 974.5 | 0.378 | 0.643 | 1021 | 0.472 | 0.643 | 469.5 | 0.141 | 0.643 | 171 | 0.311 | 0.643 | 174 | 0.323 |
| BRD-K96104201-001-01-5 | azalomycin-b | 3925 | 0.805 | 0.978 | 5290 | 0.173 | 0.518 | 3163 | 0.231 | 0.539 | 12640 | 0.336 | 0.642 | 3813 | 0.088 | 0.518 | 4477 | 0.978 | 0.978 | 3658 | 0.230 | 0.539 | 6236 | 0.960 | 0.978 | 1775 | 0.578 | 0.933 | 3756 | 0.960 | 0.978 | 1777 | 0.103 | 0.518 | 1179 | 0.543 | 0.933 | 426 | 0.064 | 0.518 | 1243 | 0.867 |
| BRD-K96123349-236-02-8 | brequinar | 1195 | 0.512 | 0.903 | 1593 | 0.895 | 0.972 | 572 | 0.079 | 0.277 | 2753 | 0.833 | 0.972 | 876 | 0.516 | 0.903 | 976 | 0.996 | 0.996 | 1373 | 0.926 | 0.972 | 1194 | 0.677 | 0.964 | 377 | 0.012 | 0.082 | 492 | 0.901 | 0.972 | 414 | 0.170 | 0.511 | 485 | 0.630 | 0.964 | 242 | 0.796 | 0.972 | 107 | 0.033 |
| BRD-K96159438-001-06-3 | adefovir-dipivoxil | 2145 | 0.594 | 0.832 | 2615 | 0.576 | 0.832 | 2284 | 0.754 | 0.880 | 6058 | 0.331 | 0.832 | 2370 | 0.427 | 0.832 | 1715 | 0.217 | 0.832 | 2466 | 0.404 | 0.832 | 1733 | 0.543 | 0.832 | 738 | 0.361 | 0.832 | 1320 | 0.170 | 0.832 | 1457 | 0.508 | 0.832 | 347 | 0.032 | 0.662 | 374 | 0.403 | 0.832 | 490 | 0.372 |
| BRD-K96233303-001-01-9 | TMS | 270 | 0.066 | 1.000 | 849 | 0.597 | 0.836 | 512 | 0.299 | 1.000 | 1559 | 0.960 | 0.982 | 599 | 0.270 | 1.000 | 157 | 0.348 | 1.000 | 239 | 0.497 | 0.746 | 634 | 0.870 | 0.982 | 106 | 0.111 | 0.212 | 146 | 0.093 | 0.195 | 343 | 0.451 | 0.729 | 18 | 0.004 | 0.011 | 108 | 0.905 | 0.982 | 11 | 0.081 |
| BRD-K96253961-046-03-1 | metaraminol | 3943 | 0.395 | 0.642 | 6918 | 0.693 | 0.857 | 2886 | 0.386 | 1.000 | 11872 | 0.140 | 1.000 | 5293 | 0.624 | 0.819 | 3972 | 0.807 | 0.924 | 6354 | 0.998 | 0.998 | 5318 | 0.836 | 0.924 | 1333 | 0.262 | 0.610 | 3742 | 0.982 | 0.998 | 1918 | 0.085 | 1.000 | 1218 | 0.331 | 0.633 | 575 | 0.192 | 0.505 | 847 | 0.319 |
| BRD-K96344439-002-02-7 | tofogliflozin | 475 | 0.498 | 1.000 | 1023.5 | 0.896 | 0.990 | 922.5 | 0.841 | 0.990 | 1591.5 | 0.186 | 0.499 | 685 | 0.232 | 0.499 | 220 | 0.086 | 0.300 | 128 | 0.034 | 0.141 | 797.5 | 0.238 | 0.499 | 122 | 0.028 | 0.141 | 396 | 0.228 | 0.499 | 241.5 | 0.674 | 0.990 | 207 | 0.874 | 0.990 | 82 | 0.610 | 0.990 | 68 | 0.476 |
| BRD-K96358241-001-01-9 | harringtonine | 3791 | 0.302 | 0.839 | 6390 | 0.854 | 0.998 | 4341 | 0.708 | 0.998 | 10685 | 0.008 | 0.169 | 4660 | 0.440 | 0.839 | 4269 | 0.992 | 0.998 | 5163 | 0.774 | 0.998 | 5689 | 0.854 | 0.998 | 1183 | 0.020 | 0.071 | 2362 | 0.903 | 0.998 | 2489 | 0.359 | 0.839 | 1803 | 0.939 | 0.998 | 731 | 0.428 | 0.839 | 721 | 0.411 |
| BRD-K96631475-001-02-4 | 10-deacetylbaccatin | 1783 | 0.960 | 1.000 | 4772 | 1.000 | 1.000 | 867 | 0.004 | 0.076 | 6428 | 0.253 | 0.663 | 2565 | 0.472 | 0.991 | 1542 | 0.144 | 0.660 | 2011 | 0.252 | 0.663 | 3047 | 0.829 | 1.000 | 799 | 0.235 | 0.663 | 1097 | 0.989 | 1.000 | 1755 | 0.537 | 0.863 | 299 | 0.578 | 0.863 | 597 | 0.617 | 0.863 | 403 | 0.470 |
| BRD-K97010173-001-04-1 | deferasirox | 278 | 0.931 | 0.996 | 788 | 0.996 | 0.996 | 106 | 0.063 | 0.263 | 623 | 0.021 | 0.110 | 185 | 0.003 | 0.021 | 78 | 0.307 | 0.587 | 373 | 0.900 | 0.996 | 532.5 | 0.814 | 0.996 | 55 | 0.143 | 0.333 | 130 | 0.133 | 0.333 | 140 | 0.174 | 0.366 | 262.5 | 0.990 | 0.996 | 63 | 0.679 | 0.996 | 228 | 0.950 |
| BRD-K97365803-001-02-1 | PI-828 | 228 | 0.244 | 1.000 | 403 | 0.808 | 0.893 | 298 | 0.160 | 1.000 | 1532 | 0.989 | 0.989 | 561 | 0.602 | 0.843 | 267 | 0.733 | 0.855 | 529 | 0.102 | 0.268 | 483.5 | 0.872 | 0.916 | 263.5 | 0.415 | 0.792 | 110 | 0.148 | 0.346 | 350 | 0.346 | 0.726 | 40 | 0.059 | 0.176 | 122 | 0.563 | 0.843 | | |
| BRD-K97746869-001-27-0 | chlorpropamide | 207 | 0.416 | 0.873 | 1118 | 0.136 | 1.000 | 592 | 0.554 | 0.995 | 1683 | 0.028 | 0.589 | 629 | 0.149 | 1.000 | 192 | 0.340 | 1.000 | 887 | 0.788 | 0.995 | 1012 | 0.764 | 0.995 | 277 | 0.981 | 0.995 | 139 | 0.928 | 0.995 | 238 | 0.254 | 1.000 | 17 | 0.093 | 0.225 | 221 | 0.877 | 0.995 | 18 | 0.096 |
| BRD-K97764662-001-17-3 | PD-173074 | 271 | 0.809 | 0.914 | 260.5 | 0.463 | 1.000 | 85 | 0.020 | 0.418 | 1244 | 0.746 | 0.914 | 547 | 0.817 | 0.914 | 224 | 0.553 | 0.914 | 302 | 0.914 | 0.914 | 362 | 0.476 | 0.914 | 132 | 0.028 | 0.098 | 251 | 0.081 | 0.212 | 235.5 | 0.328 | 1.000 | 133.5 | 0.297 | 0.692 | 76 | 0.743 | 0.914 | 169.5 | 0.907 |
| BRD-K97799481-001-16-0 | theophylline | 318 | 0.057 | 0.301 | 1363 | 0.169 | 0.508 | 313 | 0.053 | 0.301 | 2752 | 0.586 | 0.904 | 1041 | 0.445 | 0.904 | 595 | 0.502 | 0.904 | 1703 | 0.995 | 1.000 | 950 | 0.590 | 0.904 | 629 | 0.602 | 0.904 | 623 | 0.585 | 0.904 | 532 | 0.818 | 1.000 | 389 | 0.928 | 1.000 | 51 | 0.237 | 0.623 | 221 | 0.741 |
| BRD-K97810537-001-13-6 | beclomethasone-dipropionate | 983 | 0.777 | 0.959 | 1764 | 0.590 | 0.959 | 1278 | 0.636 | 0.959 | 3346 | 0.696 | 0.959 | 963 | 0.206 | 0.541 | 624 | 0.191 | 0.541 | 765 | 0.032 | 0.401 | 1656 | 0.959 | 0.959 | 454 | 0.714 | 0.959 | 281 | 0.055 | 0.401 | 293 | 0.198 | 0.541 | 493 | 0.818 | 0.959 | 327 | 0.633 | 0.959 | 281 | 0.450 |
| BRD-K98203492-003-04-1 | GSK-J4 | 2170 | 0.816 | 0.927 | 4171.5 | 0.839 | 0.927 | 2048 | 0.228 | 1.000 | 7120 | 0.152 | 1.000 | 2668 | 0.326 | 1.000 | 1629 | 0.525 | 1.000 | 2865 | 0.887 | 0.927 | 3001 | 0.531 | 1.000 | 1132 | 0.657 | 0.892 | 2103.5 | 0.753 | 0.927 | 1575 | 0.650 | 0.892 | 757 | 0.041 | 0.072 | 419 | 0.416 | 0.673 | 681 | 0.680 |
| BRD-K98384930-003-01-5 | finafloxacin | 402 | 0.534 | 0.960 | 476 | 0.004 | 0.085 | 284 | 0.091 | 0.479 | 778 | 0.217 | 0.910 | 526 | 0.812 | 0.960 | 264 | 0.946 | 0.960 | 715 | 0.871 | 0.960 | 480 | 0.846 | 0.960 | 226 | 0.466 | 0.960 | 145 | 0.727 | 0.960 | 368 | 0.373 | 1.000 | 89 | 0.824 | 0.960 | 143 | 0.713 | 0.960 | 50 | 0.390 |
| BRD-K98548675-001-05-9 | parthenolide | 1566 | 0.325 | 1.000 | 3007 | 0.074 | 0.520 | 2033 | 0.167 | 0.770 | 8030 | 0.694 | 0.911 | 3349 | 0.895 | 0.940 | 1955 | 0.658 | 0.911 | 3316 | 0.792 | 0.924 | 2447 | 0.598 | 0.911 | 405 | 0.046 | 0.120 | 1785 | 0.997 | 0.997 | 2018 | 0.866 | 0.940 | 787 | 0.626 | 0.911 | 274 | 0.439 | 0.911 | 269 | 0.133 |
| BRD-K98572433-001-02-9 | AZD8931 | 952.5 | 0.844 | 1.000 | 2204 | 1.000 | 1.000 | 485.5 | 0.008 | 0.054 | 2400.5 | 0.327 | 0.687 | 1279.5 | 0.495 | 0.945 | 529.5 | 0.107 | 0.449 | 1186 | 0.632 | 1.000 | 1294 | 0.999 | 1.000 | 155.5 | 0.886 | 1.000 | 682 | 0.868 | 1.000 | 506.5 | 0.192 | 0.673 | 697 | 0.982 | 1.000 | 66 | 0.311 | 0.687 | 155.5 | 0.886 |
| BRD-K98795921-001-01-7 | SB-2343 | 6756 | 0.955 | 0.957 | 8459 | 0.713 | 0.957 | 4846 | 0.387 | 0.804 | 17614 | 0.372 | 0.804 | 5686 | 0.208 | 0.637 | 5342 | 0.806 | 0.957 | 8090 | 0.922 | 0.957 | 6876 | 0.728 | 0.957 | 2565 | 0.587 | 0.957 | 2448 | 0.017 | 0.182 | 2335 | 0.010 | 0.182 | 2082 | 0.091 | 0.384 | 857 | 0.421 | 0.804 | 785 | 0.660 |
| BRD-K99023089-001-03-7 | AZD5363 | 1059 | 0.045 | 0.238 | 2072 | 0.860 | 0.951 | 1087 | 0.394 | 0.873 | 5601 | 0.986 | 0.986 | 2359 | 0.906 | 0.951 | 1297 | 0.034 | 0.238 | 1564 | 0.483 | 0.923 | 1886 | 0.241 | 0.842 | 596 | 0.044 | 0.238 | 474 | 0.845 | 0.951 | 1041 | 0.314 | 0.873 | 906 | 0.893 | 0.951 | 143 | 0.168 | 0.706 | 175 | 0.794 |
| BRD-K99113996-001-02-0 | AZD2014 | 1759.5 | 0.999 | 0.999 | 2402 | 0.089 | 0.930 | 1343 | 0.323 | 1.000 | 5132.5 | 0.323 | 1.000 | 1406.5 | 0.003 | 0.073 | 1481.5 | 0.363 | 1.000 | 1826.5 | 0.539 | 1.000 | 2485.5 | 0.943 | 0.990 | 781 | 0.157 | 1.000 | 1452 | 0.840 | 0.990 | 1382 | 0.755 | 0.990 | 817 | 0.881 | 0.990 | 543 | 0.904 | 0.990 | 182 | 0.249 |
| BRD-K99170882-001-04-3 | NSC-319726 | 7127 | 0.998 | 0.998 | 7599 | 0.727 | 0.998 | 4119 | 0.150 | 0.393 | 16266.5 | 0.807 | 0.998 | 5573 | 0.432 | 0.647 | 3330 | 0.214 | 0.449 | 7449 | 0.982 | 0.998 | 6885 | 0.829 | 0.998 | 1201 | 0.003 | 0.065 | 2680.5 | 0.089 | 0.393 | 3099 | 0.310 | 0.542 | 1317 | 0.020 | 0.213 | 405 | 0.133 | 0.393 | 711 | 0.098 |
| BRD-K99475619-001-01-2 | genz-644282 | 4718 | 0.073 | 0.366 | 10574 | 0.998 | 0.998 | 5785 | 0.877 | 0.998 | 15296 | 0.030 | 0.317 | 5055 | 0.066 | 0.366 | 5501 | 0.925 | 0.998 | 7743 | 0.809 | 0.998 | 7455 | 0.778 | 0.998 | 1375 | 0.002 | 0.038 | 5050 | 0.990 | 0.998 | 3629 | 0.383 | 0.893 | 2308 | 0.192 | 0.577 | 916 | 0.498 | 0.950 | 765 | 0.105 |
| BRD-K99498722-001-02-6 | plinabulin | 6256 | 0.905 | 0.994 | 9931 | 0.994 | 0.994 | 4772 | 0.355 | 0.678 | 15964 | 0.091 | 0.383 | 5122 | 0.032 | 0.224 | 5730 | 0.939 | 0.994 | 8071 | 0.991 | 0.994 | 7074 | 0.534 | 0.862 | 2247 | 0.313 | 0.678 | 4103 | 0.939 | 0.994 | 3483 | 0.625 | 0.938 | 1525 | 0.016 | 0.165 | 726 | 0.253 | 0.678 | 314 | 0.001 |
| BRD-K99545815-001-06-3 | PF-562271 | 3325 | 0.181 | 0.455 | 5716 | 0.943 | 0.996 | 3046 | 0.195 | 0.455 | 9741 | 0.168 | 0.455 | 4113 | 0.527 | 0.996 | 3868 | 0.996 | 0.996 | 4838 | 0.709 | 0.996 | 4948 | 0.919 | 0.996 | 846 | 0.024 | 0.109 | 3255 | 0.996 | 0.996 | 1766 | 0.007 | 0.141 | 1309 | 0.584 | 0.996 | 645 | 0.976 | 0.996 | 1146 | 0.852 |
| BRD-K99604664-001-01-1 | tanaproget | 373 | 0.230 | 0.728 | 1611 | 0.010 | 0.215 | 1766 | 0.749 | 0.992 | 5216 | 0.828 | 0.992 | 2354 | 0.944 | 0.992 | 1107 | 0.404 | 0.728 | 1222 | 0.416 | 0.728 | 1939 | 0.270 | 0.728 | 887 | 0.401 | 0.728 | 938 | 0.296 | 0.728 | 818 | 0.132 | 1.000 | 193 | 0.021 | 0.087 | 363 | 0.895 | 0.992 | 437 | 0.744 |
| BRD-K99792991-001-34-9 | hexachlorophene | 2129 | 0.757 | 0.935 | 3416 | 0.738 | 0.935 | 1661 | 0.056 | 0.302 | 7555 | 0.958 | 0.958 | 2644 | 0.600 | 0.900 | 1773 | 0.481 | 0.860 | 2180 | 0.143 | 0.499 | 2611 | 0.822 | 0.948 | 943 | 0.224 | 0.587 | 1461 | 0.409 | 0.860 | 1505 | 0.474 | 0.860 | 424 | 0.492 | 0.860 | 312 | 0.211 | 0.587 | 306 | 0.057 |
| BRD-K99946902-001-07-5 | hexylresorcinol | 126 | 0.125 | 0.711 | 620.5 | 0.590 | 0.954 | 338 | 0.248 | 0.711 | 1884 | 0.941 | 1.000 | 691 | 0.271 | 0.711 | 396.5 | 0.467 | 0.892 | 795 | 0.955 | 1.000 | 481 | 0.171 | 0.711 | 212 | 0.222 | 0.711 | 328 | 0.217 | 0.711 | 333 | 0.770 | 0.770 | 418.5 | 0.811 | 1.000 | 24 | 0.132 | 0.711 | 153.5 | 0.603 |
| BRD-K99964838-001-11-9 | bosutinib | 626 | 0.981 | 0.981 | 763 | 0.980 | 0.981 | 374 | 0.006 | 0.068 | 1402 | 0.408 | 0.857 | 518 | 0.216 | 0.559 | 468 | 0.860 | 0.981 | 814 | 0.911 | 0.981 | 614 | 0.974 | 0.981 | 101 | 0.844 | 0.981 | 149 | 0.071 | 0.299 | 166 | 0.110 | 0.384 | 300 | 0.960 | 0.981 | 39 | 0.240 | 0.559 | 65 | 0.506 |
| BRD-M92675308-003-07-1 | efonidipine-monoethanolate | 468 | 0.960 | 0.960 | 898 | 0.543 | 1.000 | 530 | 0.140 | 0.845 | 1826 | 0.246 | 0.845 | 653 | 0.436 | 0.966 | 405 | 0.144 | 0.845 | 953 | 0.680 | 1.000 | 1478 | 0.873 | 1.000 | 159 | 0.172 | 0.845 | 326 | 0.282 | 0.845 | 497 | 0.636 | 1.000 | 147 | 0.933 | 1.000 | 150 | 0.460 | 0.966 | | |
| BRD-M97302542-001-03-6 | dichloroacetate | 772 | 0.272 | 0.635 | 1940 | 0.530 | 0.839 | 1190 | 0.751 | 0.839 | 3518 | 0.907 | 0.907 | 1509 | 0.662 | 0.839 | 696 | 0.759 | 0.839 | 808 | 0.345 | 0.725 | 1635 | 0.839 | 0.881 | 374 | 0.178 | 0.534 | 695 | 0.757 | 0.839 | 482 | 0.091 | 0.963 | 318 | 0.250 | 0.635 | 112 | 0.157 | 0.534 | 292 | 0.467 |

| broad_id | name | al_nervous_system_PVAL_ADJ | fibroblast_MWU_STAT | fibroblast_PVAL | fibroblast_PVAL_ADJ | bile_duct_MWU_STAT | bile_duct_PVAL | bile_duct_PVAL_ADJ | thyroid_MWU_STAT | thyroid_PVAL | thyroid_PVAL_ADJ | upper_aerodigestive_MWU_STAT | upper_aerodigestive_PVAL | upper_aerodigestive_PVAL_ADJ | uterus_MWU_STAT | uterus_PVAL | uterus_PVAL_ADJ | liver_MWU_STAT | liver_PVAL | liver_PVAL_ADJ | esophagus_MWU_STAT | esophagus_PVAL | esophagus_PVAL_ADJ |
|---|---|---|---|---|---|---|---|---|---|---|---|---|---|---|---|---|---|---|---|---|---|---|---|
| BRD-K88677950-001-03-9 | PD-198306 | 0.885 | | | | 124 | 0.017 | 0.118 | 504 | 0.042 | 0.221 | 1794 | 0.187 | 0.561 | 1894 | 0.968 | 0.996 | 2073.5 | 0.973 | 0.996 | 3068 | 0.996 | 0.996 |
| BRD-K88807631-001-01-3 | elacridar | | 24 | 0.093 | 0.813 | 601 | 0.942 | 0.956 | 835 | 0.785 | 0.956 | 1028 | 0.905 | 0.956 | 809 | 0.155 | 0.813 | 803 | 0.489 | 0.856 | 534 | 0.133 | 0.813 |
| BRD-K89014967-001-04-3 | AS-703026 | 0.853 | 211 | 0.907 | 1.000 | 232 | 0.148 | 0.621 | 695 | 0.755 | 1.000 | 1451 | 0.050 | 0.262 | 1342 | 0.929 | 1.000 | 1267 | 0.700 | 1.000 | 1055 | 0.739 | 1.000 |
| BRD-K89015388-001-01-7 | GSK923295 | 0.227 | | | | 334 | 0.731 | 0.903 | 690 | 0.851 | 0.940 | 1859 | 0.438 | 1.000 | 975 | 0.613 | 1.000 | 1101 | 0.818 | 0.940 | 2268 | 0.658 | 0.903 |
| BRD-K89053832-001-01-0 | VLX600 | 0.234 | | | | 710.5 | 0.961 | 0.961 | 1553 | 0.942 | 0.961 | 2111 | 0.193 | 1.000 | 1955.5 | 0.434 | 0.771 | 1335 | 0.533 | 0.800 | 2463 | 0.706 | 0.927 |
| BRD-K89121095-001-01-7 | tedizolid-phosphate | 0.611 | 273 | 0.915 | 0.987 | 785 | 0.849 | 0.987 | 1331 | 0.891 | 0.987 | 2398 | 0.190 | 0.665 | 2186 | 0.505 | 0.964 | 1102 | 0.827 | 0.987 | 1934 | 0.008 | 0.163 |
| BRD-K89299012-001-06-0 | DCEBIO | 0.554 | 100 | 0.313 | 0.657 | 487 | 0.883 | 0.984 | 624 | 0.211 | 0.554 | 2565 | 0.618 | 0.982 | 1274 | 0.308 | 0.657 | 1278 | 0.528 | 0.982 | 3650 | 0.937 | 0.984 |
| BRD-K89561498-001-01-7 | SN-38 | | | | | 662 | 0.477 | 0.867 | 209 | 0.423 | 1.000 | 482 | 0.013 | 0.138 | 850 | 0.231 | 1.000 | 1155 | 0.925 | 0.971 | 891 | 0.013 | 0.138 |
| BRD-K89704198-001-12-3 | vincamine | | | | | 156 | 0.651 | 0.989 | 58 | 0.404 | 0.801 | 635 | 0.866 | 0.989 | 333 | 0.116 | 1.000 | 314 | 0.954 | 0.989 | 519 | 0.168 | 1.000 |
| BRD-K89714990-001-01-5 | tepoxalin | 0.544 | | | | 521 | 0.956 | 0.956 | 772 | 0.942 | 0.956 | 858 | 0.146 | 1.000 | 1027 | 0.818 | 0.956 | 672 | 0.061 | 0.213 | 1385 | 0.942 | 0.956 |
| BRD-K89839824-001-05-3 | raltitrexed | 0.587 | 27 | 0.065 | 0.386 | 2177 | 0.971 | 0.978 | 1619 | 0.082 | 0.386 | 3853 | 0.145 | 0.436 | 5121 | 0.937 | 0.978 | 4194 | 0.932 | 0.978 | 5217 | 0.721 | 0.978 |
| BRD-K90027355-001-13-3 | spironolactone | 0.344 | 28 | 0.082 | 0.344 | 986 | 0.687 | 0.902 | 1736 | 0.975 | 0.990 | 2845 | 0.219 | 0.656 | 2572 | 0.371 | 0.822 | 2400 | 0.841 | 0.982 | 3628 | 0.990 | 0.990 |
| BRD-K90374350-001-09-2 | chlorquinaldol | 0.318 | | | | 120 | 0.760 | 0.970 | 326 | 0.462 | 0.881 | 664 | 0.525 | 0.918 | 540 | 0.200 | 0.516 | 183 | 0.576 | 0.931 | 927 | 0.779 | 0.970 |
| BRD-K90382497-001-10-7 | GW-843682X | 0.055 | | | | 327 | 0.129 | 0.322 | 789 | 0.055 | 0.582 | 3793 | 0.840 | 0.981 | 2668 | 0.310 | 0.651 | 1212 | 0.021 | 0.089 | 2734 | 0.084 | 0.281 |
| BRD-K90497590-001-04-2 | AR-12 | 0.644 | | | | 676 | 0.946 | 0.992 | 705 | 0.460 | 0.734 | 1976 | 0.351 | 0.684 | 1498 | 0.245 | 0.644 | 642 | 0.016 | 0.084 | 2147 | 0.094 | 1.000 |
| BRD-K90948141-001-01-4 | indiplon | 0.156 | | | | | | | 123 | 0.536 | 1.000 | 199 | 0.308 | 1.000 | 324 | 0.403 | 1.000 | 142 | 0.027 | 0.064 | 236 | 0.514 | 1.000 |
| BRD-K91370081-001-25-1 | anisomycin | 0.913 | 273 | 0.609 | 0.913 | 1724 | 0.890 | 0.990 | 2250 | 0.549 | 0.913 | 4797 | 0.560 | 0.913 | 4662 | 0.607 | 0.913 | 3611 | 0.608 | 0.913 | 4165 | 0.063 | 0.354 |
| BRD-K91601245-001-12-0 | mercaptopurine | | | | | 216 | 0.908 | 0.949 | 296 | 0.574 | 0.949 | 322.5 | 0.949 | 0.949 | 273 | 0.188 | 1.000 | 265 | 0.416 | 0.949 | 607.5 | 0.486 | 0.949 |
| BRD-K91623615-001-05-0 | ABT-751 | 0.388 | 131 | 0.295 | 0.673 | 1767 | 0.919 | 0.996 | 1996 | 0.320 | 0.673 | 3174 | 0.012 | 0.259 | 3556 | 0.103 | 0.542 | 2940 | 0.425 | 0.743 | 4993 | 0.577 | 0.865 |
| BRD-K91701654-001-03-1 | U-0126 | | | | | 579 | 0.992 | 0.992 | 312 | 0.160 | 1.000 | 512 | 0.528 | 0.853 | 496 | 0.942 | 0.992 | 131 | 0.283 | 0.830 | 899 | 0.955 | 0.992 |
| BRD-K91825936-001-01-2 | ZK811752 | 0.920 | 233 | 0.871 | 0.963 | 526 | 0.423 | 0.889 | 763 | 0.367 | 0.867 | 1213 | 0.144 | 0.606 | 1023 | 0.015 | 0.139 | 694 | 0.004 | 0.088 | 2473 | 0.604 | 0.920 |
| BRD-K92000912-001-12-6 | AM-251 | 0.538 | | | | 307.5 | 0.793 | 0.925 | 219 | 0.394 | 1.000 | 650 | 0.105 | 0.723 | 392 | 0.101 | 0.723 | 408 | 0.283 | 0.538 | 912 | 0.680 | 0.925 |
| BRD-K92093830-003-30-8 | doxorubicin | 0.693 | 88 | 0.195 | 0.679 | 1082 | 0.441 | 0.693 | 2057 | 0.338 | 0.693 | 4054 | 0.298 | 0.693 | 3761 | 0.150 | 0.634 | 2982 | 0.259 | 0.679 | 5604 | 0.748 | 0.982 |
| BRD-K92150287-066-01-2 | exatecan-mesylate | | | | | 138 | 0.794 | 0.988 | 251 | 0.190 | 0.999 | 486 | 0.039 | 0.412 | 569 | 0.020 | 0.412 | 735 | 0.682 | 0.955 | 461 | 0.071 | 0.498 |
| BRD-K92241597-001-06-0 | tosedostat | 0.996 | 232 | 0.534 | 0.996 | 2138 | 0.996 | 0.996 | 1902 | 0.252 | 0.757 | 3726 | 0.059 | 0.496 | 3881 | 0.686 | 0.996 | 2927 | 0.431 | 0.996 | 5992 | 0.940 | 0.996 |
| BRD-K92428153-001-04-4 | mycophenolate-mofetil | 0.869 | | | | 225 | 0.706 | 0.874 | 77 | 0.605 | 0.874 | 494 | 0.755 | 0.874 | 277 | 0.641 | 0.874 | 283 | 0.174 | 0.730 | 447 | 0.805 | 0.874 |
| BRD-K92428232-001-10-6 | GSK461364 | 0.141 | 315 | 0.906 | 0.984 | 1296.5 | 0.963 | 0.984 | 707 | 0.082 | 0.571 | 3080 | 0.511 | 0.825 | 2820.5 | 0.575 | 0.863 | 1987 | 0.238 | 0.587 | 3133 | 0.285 | 0.587 |
| BRD-K92657060-001-05-7 | zaleplon | 0.672 | | | | 88 | 0.592 | 0.887 | 147 | 0.142 | 0.672 | 733 | 0.695 | 0.887 | 586 | 0.700 | 0.887 | 40 | 0.037 | 0.495 | 887 | 0.719 | 0.887 |
| BRD-K92723993-001-06-7 | imatinib | 0.547 | | | | 406 | 0.854 | 0.944 | 620 | 0.156 | 0.547 | 1622 | 0.707 | 0.927 | 1874 | 0.356 | 0.927 | 948 | 0.055 | 0.547 | 2356 | 0.953 | 0.999 |
| BRD-K92991072-001-04-9 | PAC-1 | 0.406 | | | | 92.5 | 0.822 | 0.903 | 137.5 | 0.624 | 0.903 | 349.5 | 0.234 | 1.000 | 327.5 | 0.903 | 0.903 | 458.5 | 0.897 | 0.903 | 513.5 | 0.691 | 0.903 |
| BRD-K93034159-001-25-8 | cladribine | 0.960 | | | | 793 | 0.507 | 0.960 | 455 | 0.053 | 0.370 | 1944 | 0.786 | 0.971 | 2092 | 0.609 | 0.960 | 1659 | 0.269 | 0.936 | 2663 | 0.732 | 0.960 |
| BRD-K93123848-001-04-1 | RAF265 | 0.864 | | | | 389 | 0.763 | 0.997 | 1275 | 0.652 | 1.000 | 2197.5 | 0.546 | 1.000 | 2207.5 | 0.398 | 1.000 | 1181 | 0.292 | 0.558 | 2727.5 | 0.957 | 0.997 |
| BRD-K93779381-001-01-9 | ingenol-mebutate | 1.000 | | | | 85 | 0.590 | 1.000 | 99 | 0.708 | 1.000 | 449 | 0.961 | 1.000 | 446 | 0.296 | 0.821 | 233 | 0.083 | 0.727 | 1065 | 0.980 | 0.980 |
| BRD-K93869735-001-01-1 | taltirelin | 0.546 | | | | | | | 74 | 0.642 | 1.000 | 308 | 0.000 | 0.008 | 246 | 0.258 | 0.987 | 91 | 0.272 | 0.987 | 687 | 0.860 | 0.860 |
| BRD-K94176593-001-09-4 | TWS-119 | 0.260 | | | | 182 | 0.477 | 0.834 | 240 | 0.341 | 0.754 | 1174 | 0.260 | 0.684 | 898 | 0.186 | 0.652 | 541.5 | 0.752 | 0.928 | 1315.5 | 0.678 | 0.928 |
| BRD-K94441233-001-13-0 | mevastatin | 0.993 | 36 | 0.109 | 0.460 | 458 | 0.797 | 0.993 | 737 | 0.005 | 0.054 | 2427 | 0.672 | 0.993 | 2811 | 0.731 | 0.993 | 1685 | 0.559 | 0.993 | 3191 | 0.780 | 0.993 |
| BRD-K94455792-001-01-8 | ICG-001 | 0.733 | | | | 210 | 0.471 | 0.733 | 661 | 0.558 | 0.733 | 1112 | 0.272 | 1.000 | 1067 | 0.546 | 0.733 | 755 | 0.533 | 0.733 | 1546 | 0.375 | 0.733 |
| BRD-K94485812-001-02-9 | LDN-57444 | 0.534 | | | | 386 | 0.461 | 1.000 | 544 | 0.253 | 1.000 | 1686 | 0.945 | 0.997 | 1121 | 0.409 | 1.000 | 946 | 0.161 | 0.308 | 2021 | 0.486 | 0.729 |
| BRD-K95142244-001-01-5 | talazoparib | 0.257 | | | | 363 | 0.894 | 0.961 | 495 | 0.871 | 0.961 | 1385 | 0.948 | 0.961 | 1054 | 0.304 | 1.000 | 991 | 0.819 | 0.961 | 1733 | 0.380 | 0.726 |
| BRD-K95292280-001-01-3 | SB-657510 | 0.256 | | | | 141.5 | 0.784 | 0.957 | 37 | 0.290 | 1.000 | 235 | 0.344 | 1.000 | 149.5 | 0.153 | 1.000 | 134.5 | 0.734 | 0.957 | 346.5 | 0.683 | 0.957 |
| BRD-K95435023-001-08-5 | PHA-665752 | 0.950 | | | | 291 | 0.311 | 0.932 | 752 | 0.648 | 0.950 | 1207 | 0.643 | 0.950 | 1066 | 0.792 | 0.950 | 846 | 0.600 | 0.950 | 1716 | 0.744 | 0.950 |
| BRD-K95573441-001-01-2 | PFK-015 | 0.901 | 27 | 0.099 | 0.838 | 209 | 0.946 | 0.946 | 673 | 0.844 | 0.946 | 928 | 0.033 | 0.703 | 1030 | 0.686 | 0.901 | 792 | 0.629 | 0.901 | 995 | 0.129 | 0.838 |
| BRD-K95581532-001-01-5 | RN-1734 | 0.032 | | | | 402 | 0.068 | 0.477 | 1538 | 0.659 | 0.976 | 2812 | 0.091 | 0.477 | 2821 | 0.552 | 0.966 | 2417 | 0.813 | 0.976 | 3782 | 0.668 | 0.976 |
| BRD-K95773607-004-03-8 | darifenacin | 0.947 | | | | 229.5 | 0.832 | 0.947 | 492.5 | 0.797 | 0.947 | 621 | 0.134 | 0.716 | 644 | 0.170 | 0.716 | 379 | 0.074 | 0.172 | 751.5 | 0.615 | 0.947 |
| BRD-K96006502-001-02-6 | deflazacort | 0.643 | | | | 65 | 0.249 | 0.643 | 392.5 | 0.395 | 0.643 | 1399.5 | 0.312 | 0.643 | 1323 | 0.932 | 0.969 | 326 | 0.520 | 0.643 | 1184.5 | 0.149 | 1.000 |
| BRD-K96104201-001-01-5 | azalomycin-b | 0.978 | 60 | 0.157 | 0.518 | 1144 | 0.764 | 0.978 | 1002 | 0.131 | 0.518 | 3308 | 0.333 | 0.642 | 2621 | 0.062 | 0.518 | 3132 | 0.890 | 0.978 | 4299 | 0.808 | 0.978 |
| BRD-K96123349-236-02-8 | brequinar | 0.175 | | | | 195 | 0.198 | 0.519 | 315 | 0.689 | 0.964 | 411 | 0.377 | 0.792 | 791 | 0.312 | 0.728 | 554 | 0.063 | 0.265 | 709 | 0.074 | 1.000 |
| BRD-K96159438-001-06-3 | adefovir-dipivoxil | 0.832 | 180 | 0.708 | 0.875 | 461 | 0.305 | 0.832 | 902 | 0.902 | 0.924 | 1577 | 0.686 | 0.875 | 2201.5 | 0.800 | 0.884 | 1073.5 | 0.511 | 0.832 | 2104 | 0.924 | 0.924 |
| BRD-K96233303-001-01-9 | TMS | 0.189 | | | | | | | 89 | 0.783 | 0.982 | 333 | 0.907 | 0.982 | 166 | 0.149 | 1.000 | 211 | 0.343 | 0.601 | 603 | 0.982 | 0.982 |
| BRD-K96253961-046-03-1 | metaraminol | 0.633 | | | | 891 | 0.381 | 1.000 | 1506 | 0.459 | 0.642 | 3315 | 0.450 | 0.642 | 2958 | 0.302 | 1.000 | 1703 | 0.011 | 0.034 | 3950 | 0.400 | 0.642 |
| BRD-K96344439-002-02-7 | tofogliflozin | 0.908 | | | | 327.5 | 0.730 | 0.990 | 669 | 0.995 | 0.995 | 328 | 0.952 | 0.995 | 255 | 0.163 | 1.000 | 204 | 0.863 | 0.990 | 495 | 0.803 | 0.990 |
| BRD-K96358241-001-01-9 | harringtonine | 0.839 | | | | 1239 | 0.858 | 0.998 | 2429 | 0.998 | 0.998 | 2781 | 0.059 | 0.411 | 2668 | 0.017 | 0.181 | 2630 | 0.928 | 0.998 | 3714 | 0.252 | 1.000 |
| BRD-K96631475-001-02-4 | 10-deacetylbaccatin | 0.822 | | | | 574 | 0.876 | 1.000 | 784 | 0.430 | 0.991 | 1522 | 0.032 | 0.333 | 1343 | 0.087 | 0.612 | 1163 | 0.961 | 1.000 | 1902 | 0.157 | 0.660 |
| BRD-K97010173-001-04-1 | deferasirox | 0.996 | | | | | | | | | | 170.5 | 0.686 | 0.996 | 232 | 0.957 | 0.996 | 102 | 0.534 | 0.935 | 250.5 | 0.147 | 1.000 |
| BRD-K97365803-001-02-1 | PI-828 | | | | | 103 | 0.033 | 0.694 | 73 | 0.677 | 0.843 | 198 | 0.682 | 0.843 | 156 | 0.149 | 1.000 | 230 | 0.527 | 0.843 | 255 | 0.671 | 0.843 |
| BRD-K97746869-001-27-0 | chlorpropamide | 0.225 | | | | 139 | 0.928 | 0.995 | 347 | 0.724 | 0.995 | 621 | 0.647 | 0.995 | 431 | 0.252 | 1.000 | 459 | 0.830 | 0.995 | 1053 | 0.995 | 0.995 |
| BRD-K97764662-001-17-3 | PD-173074 | 0.914 | 10 | 0.081 | 0.212 | | | | 188.5 | 0.678 | 0.914 | 124 | 0.626 | 0.914 | 346 | 0.400 | 1.000 | 279 | 0.841 | 0.914 | 429.5 | 0.563 | 0.914 |
| BRD-K97799481-001-16-0 | theophylline | 1.000 | | | | 36 | 0.157 | 0.508 | 174 | 0.163 | 0.508 | 1114 | 0.766 | 1.000 | 211 | 0.027 | 0.301 | 449 | 0.055 | 0.301 | 714 | 0.949 | 0.949 |
| BRD-K97810537-001-13-6 | beclomethasone-dipropionate | 0.959 | 185 | 0.934 | 0.959 | 153 | 0.078 | 0.408 | 376 | 0.196 | 0.541 | 1522 | 0.864 | 0.959 | 573 | 0.484 | 0.959 | 1083 | 0.912 | 0.959 | 741 | 0.057 | 0.401 |
| BRD-K98203492-003-04-1 | GSK-J4 | 0.892 | | | | | | | 1207 | 0.538 | 1.000 | 2138 | 0.114 | 1.000 | 1311 | 0.147 | 1.000 | 2342 | 0.927 | 0.927 | 2832 | 0.632 | 1.000 |
| BRD-K98384930-003-01-5 | finafloxacin | 0.960 | | | | 14 | 0.082 | 0.479 | 201 | 0.680 | 0.960 | 373 | 0.089 | 0.479 | 264 | 0.946 | 0.960 | 309 | 0.622 | 0.960 | 541 | 0.960 | 0.960 |
| BRD-K98548675-001-05-9 | parthenolide | 0.310 | | | | 300 | 0.183 | 0.770 | 863 | 0.762 | 0.924 | 1855 | 0.069 | 0.520 | 1148 | 0.068 | 0.520 | 1498 | 0.609 | 0.911 | 2471 | 0.625 | 0.911 |
| BRD-K98572433-001-02-9 | AZD8931 | 1.000 | | | | 673.5 | 0.853 | 1.000 | 463.5 | 0.981 | 1.000 | 429 | 0.000 | 0.000 | 445 | 0.229 | 0.687 | 558 | 0.310 | 0.687 | 756 | 0.004 | 0.039 |
| BRD-K98795921-001-01-7 | SB-2343 | 0.957 | 180 | 0.393 | 0.804 | 784 | 0.037 | 0.194 | 2860 | 0.929 | 0.957 | 4332 | 0.212 | 0.637 | 5216 | 0.920 | 0.957 | 2561 | 0.029 | 0.194 | 5863 | 0.957 | 0.957 |
| BRD-K99023089-001-03-7 | AZD5363 | 0.951 | 156 | 0.707 | 0.951 | 91 | 0.345 | 0.873 | 712 | 0.798 | 0.951 | 1446 | 0.805 | 0.951 | 794 | 0.001 | 0.019 | 663 | 0.416 | 0.873 | 1982 | 0.772 | 0.951 |
| BRD-K99113996-001-02-0 | AZD2014 | 0.435 | | | | 381 | 0.892 | 0.990 | 524.5 | 0.877 | 0.990 | 981.5 | 0.282 | 1.000 | 2173.5 | 0.821 | 0.990 | 1362 | 0.208 | 0.397 | 1273.5 | 0.388 | 1.000 |
| BRD-K99170882-001-04-3 | NSC-319726 | 0.393 | 167 | 0.348 | 0.562 | 1864 | 0.971 | 0.998 | 1905 | 0.292 | 0.542 | 3490 | 0.118 | 0.393 | 2838 | 0.041 | 0.288 | 2510 | 0.176 | 0.411 | 5770 | 0.963 | 0.998 |
| BRD-K99475619-001-01-2 | genz-644282 | 0.366 | 228 | 0.495 | 0.950 | 2053 | 0.901 | 0.998 | 2351 | 0.582 | 0.998 | 4243 | 0.254 | 0.665 | 3849 | 0.093 | 0.366 | 3686 | 0.739 | 0.998 | 5690 | 0.746 | 0.998 |
| BRD-K99498722-001-02-6 | plinabulin | 0.031 | 143 | 0.314 | 0.678 | 1551 | 0.724 | 0.994 | 2064 | 0.336 | 0.678 | 3825 | 0.055 | 0.286 | 4140 | 0.225 | 0.678 | 3974 | 0.800 | 0.994 | 4887 | 0.426 | 0.746 |
| BRD-K99545815-001-06-3 | PF-562271 | 0.996 | | | | 894 | 0.790 | 0.996 | 1561 | 0.868 | 0.996 | 3036 | 0.295 | 0.619 | 2619 | 0.128 | 1.000 | 1131 | 0.026 | 0.109 | 2989 | 0.162 | 0.455 |
| BRD-K99604664-001-01-1 | tanaproget | 0.992 | | | | 807 | 0.992 | 0.992 | 932 | 0.908 | 0.992 | 1436 | 0.416 | 0.728 | 1758 | 0.865 | 0.992 | 495 | 0.001 | 0.006 | 2133 | 0.950 | 0.992 |
| BRD-K99792991-001-34-9 | hexachlorophene | 0.302 | 51 | 0.134 | 0.499 | 782 | 0.903 | 0.948 | 653 | 0.697 | 0.935 | 2344 | 0.549 | 0.887 | 1552 | 0.057 | 0.302 | 944 | 0.043 | 0.302 | 2842 | 0.886 | 0.948 |
| BRD-K99946902-001-07-5 | hexylresorcinol | 0.954 | | | | 134.5 | 0.945 | 1.000 | 242.5 | 0.350 | 0.817 | 438.5 | 0.636 | 0.954 | 620 | 0.391 | 0.821 | 348.5 | 0.019 | 0.407 | 539 | 0.748 | 1.000 |
| BRD-K99964838-001-11-9 | bosutinib | 0.965 | | | | 201 | 0.923 | 0.981 | 84 | 0.190 | 0.559 | 291 | 0.002 | 0.046 | 396 | 0.848 | 0.981 | 226 | 0.711 | 0.981 | 202 | 0.023 | 0.162 |
| BRD-M92675308-003-07-1 | efonidipine-monoethanolate | | | | | 64 | 0.376 | 0.966 | 351 | 0.932 | 1.000 | 630 | 0.106 | 0.845 | 662 | 0.279 | 0.845 | 339 | 0.627 | 1.000 | 983 | 0.875 | 1.000 |
| BRD-M97302542-001-03-6 | dichloroacetate | 0.839 | | | | 238 | 0.663 | 0.839 | 463 | 0.184 | 1.000 | 967 | 0.092 | 0.963 | 1209 | 0.614 | 0.839 | 583 | 0.113 | 0.473 | 1355 | 0.710 | 0.839 |

| Property (why it matters) | Resource<br>L1000 | Perturb-seq | GDSC | PRISM (secondary screen) |
|---|---|---|---|---|
| **How the data are generated (scale)** | Bulk transcriptional profiles in ~20–30 cell lines (mostly immortalised, non-ovarian); ~1.3 M compound signatures across ~20,000 perturbations; 978 measured landmark genes (rest imputed). | Single-cell transcriptomes after CRISPRi/CRISPRa/KOs; typically thousands to tens of thousands of cells per perturbation with hundreds of perturbations assayed per experiment in one or a few host cell lines. | Dose–response viability (IC50/AUC) to ~500 oncology-focused compounds across ~1,000 individually plated cancer cell lines spanning many tissue types (incl. ~50 ovarian/urogenital). | Each compound tested individually across 1–8 doses in each of ~500 barcoded cell lines (incl. ~30 ovarian/urogenital) with viability readout. |
| **Readout type (mechanistic vs outcome)** | Mechanistic (bulk transcriptome) but mixes on/off-target effects for small molecules. | Mechanistic & causal (gene→program change) at single-cell level; closest to pathway mechanism. | Outcome (viability) only; no direct mechanism readout. | Outcome (viability) only; no direct mechanism readout. |
| **Perturbation type (genetic vs chemical)** | Both genetic & chemical; chemical polypharmacology common. | Genetic only → clean target→program links. | Chemical only. | Chemical only. |
| **Dose/timepoint design (detectability of selective effects)** | Often fixed dose, short (6–24 h) exposure; may miss late programs. | Variable timepoints capturing early rewiring. | Full dose–response; robust effect size estimates. | Multiple doses but typically fewer than GDSC. |
| **Tissue/context alignment (ovarian relevance)** | Limited cell-line diversity; many profiles from a few canonical non-ovarian lines; context mismatch for HGSC. | Depends on host line; typically non-ovarian but mechanism generalizes because it is genetic. | Broad panel with some ovarian/urogenital lines; can subset/weight by lineage. | Broad panel includes ovarian/urogenital lines; can subset/weight by lineage. |
| **Polypharmacology confounding (false positives from off-target effects) hurting mechanistic insight** | High for small-molecule profiles; off-target signatures common. | Low (genetic perturbation isolates target/pathway). | Moderate–high: pan-killers look "good" across many lines, can be filtered by contrasting different cell lines. | High: pan-killers look "good" across many lines, filtering more difficult with noisier data. |
| **Pooling/deconvolution artefacts (context specificity)** | N/A (bulk per line). | N/A (single cell). | N/A (per-line assays). | Pooling cell lines adds variability/noise. |
| **Direct evidence for adjuvant synergy or selective activity** | None, synergy inference possible (e.g., overlap between vulnerability and reversal signatures) at bulk level. | None, synergy inference possible (e.g., overlap between vulnerability and reversal signatures) at single cell level. | None, selective activity inference possible (e.g., selective potency in relevant lines). | None, selective activity inference possible (e.g., selective potency in relevant lines). |
| **Compatibility with scRNA-seq signatures (ease of mapping)** | Good for signature reversal; bulk may obscure subclones. | Excellent: single-cell to single-cell/program mapping. | Indirect: map signature-high lines → differential viability. | Indirect and noisier due to limited dose range and barcoding legacy. |
| **Support for signature mapping to targets/MoA** | Strong (MoA labels for many compounds; also genetic KO/knockdown). | Strong (gene identities, interactomes). | Strong for targets/MoA; clinical stage info often available. | Moderate; MoA present but assay design weakens mechanistic interpretation. |
| **Drug selection and assay bias toward broadly cytotoxic agents** | Moderate (compound set skew; short timepoints). | Low (genetic not cytotoxic by default). | High (pan-killers rise to top in viability). | High (even after secondary screen, broadly cytotoxic drugs still appear active). |
| **Relevant data about carboplatin context** | None. | None. | Carboplatin IC50/AUC across cell lines allows stratification. | Carboplatin IC50/AUC across cell lines allows stratification, but noisier due to assay design. |
| **Bottom line – general solid-tumor adjuvant discovery** | Broad hypothesis generator for signature reversal; bulk/lineage mismatch and polypharmacology need to be controlled. | Mechanism-proximal map program→target; ideal to nominate classes that mimic predicted vulnerabilities. Requires manual selection of compounds. | Translational triage for selective sensitivity in relevant lineages; good gatekeeper before wet-lab testing. | Scale and chemical breadth, but noisier and more biased toward broadly cytotoxic agents than GDSC; better for coarse screens than precise adjuvant picking. |
| **Bottom line – carboplatin adjuvants in HGSC** | Useful for mechanism hypotheses but weak context match to HGSC or carboplatin response. | Strong for mapping resistance programs to actionable targets but requires chemical follow-up. | Most informative for carboplatin adjuvant nomination because of per-line, multi-dose viability including ovarian lines. | Valuable for breadth but limited precision for HGSC carboplatin adjuvants; secondary screen improves but does not eliminate noise. |

**Supplementary Table 10. Comparative utility of four drug-response resources for predicting adjuvants from single-cell resistance programs.**
**Abbreviations**. MoA, mechanism of action; scRNA-seq, single-cell RNA sequencing. "Tissue/context alignment" indicates how well the training lines match ovarian/urogenital biology. The "Bottom line"
